# Inferring binding specificities of human transcription factors with the wisdom of crowds

**DOI:** 10.1101/2025.11.16.688692

**Authors:** Nikita Gryzunov, Dmitry Penzar, Vasilii Kamenets, Valery Vyaltsev, Ivan Kozin, Irina A. Eliseeva, Vladimir Nozdrin, Ilya E. Vorontsov, Sergey Bushuev, Vadim Strekalovskikh, Arsenii Zinkevich, Gregory Andrews, Matwej Bedarew, Ido Blass, Dmitry Frolov, Iuliia Lariushina, Jill Moore, Yaron Orenstein, German Roev, Danil Salimov, Noam Shimshoviz, Ido Tziony, Zhiping Weng, IBIS Consortium, GRECO-BIT/Codebook Consortium, Philipp Bucher, Bart Deplancke, Oriol Fornes, Jan Grau, Ivo Grosse, Arttu Jolma, Fedor A. Kolpakov, Vsevolod J. Makeev, Timothy R. Hughes, Ivan V. Kulakovskiy

## Abstract

DNA motif discovery and, particularly, computational modeling of transcription factor binding motifs, has been a mecca of algorithmic bioinformatics for several decades. Here, we report the results of the largest open community challenge in Inferring <u>B</u>Inding <u>S</u>pecificities (IBIS), where participants all over the world were invited to construct binding specificity models from multi-assay experimental data for poorly studied human transcription factors. The submissions were rigorously tested against a rich held-out dataset. Benchmarking demonstrated a consistent advantage of properly designed deep learning models over traditional positional weight matrices and other machine learning methods. Yet, the positional weight matrices displayed a surprisingly strong performance out of the box, being only slightly behind the best deep learning models. A post-challenge assessment of a selection of other deep learning methods further solidified this finding. IBIS highlights the power of benchmarking in finding adequate DNA motif representations, emphasizes the pros and cons of various machine learning methods applied to DNA motif modeling, and establishes a rich dataset, benchmarking protocols, and computational framework for a fair cross-platform evaluation of future models of transcription factor binding motifs in DNA sequences.

**Graphical Abstract:** 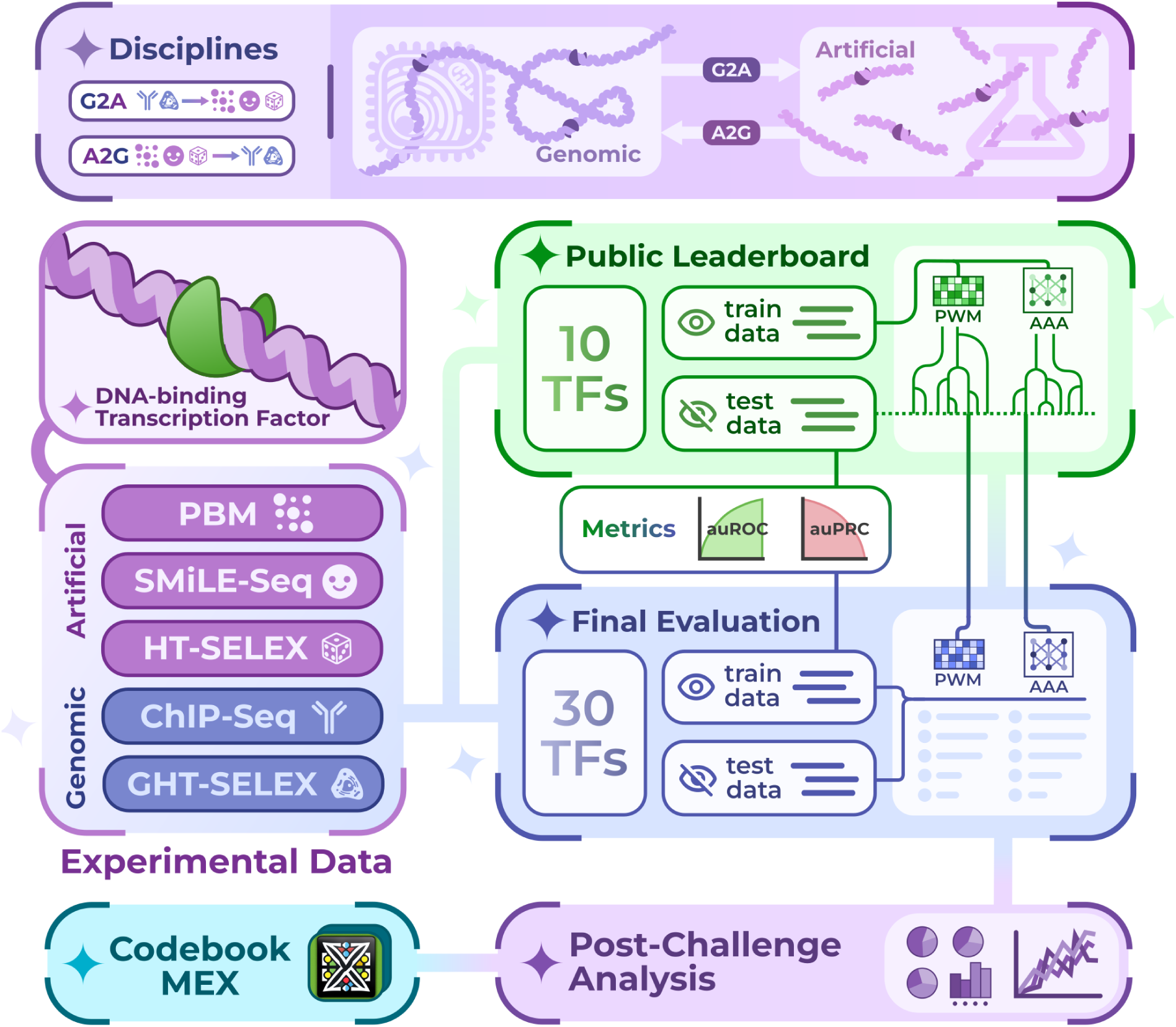

## Introduction

Transcription factors, or TFs, are among the nuts and bolts of gene regulatory mechanisms. The regulation of transcription by TFs is driven by their ability to recognize short nucleotide patterns, the sequence motifs, whose occurrences in gene regulatory regions define TF binding at their cognate binding sites (TFBS), the transcriptional response of the target genes, and, consequently, the global gene regulatory network^1^. There is a plethora of experimental methods for mapping DNA-protein interactions to identify transcription factor binding sites, both in genomic regions and artificial sequences^2^. Yet, sequence-based prediction of TFBS nonetheless requires computational modelling of DNA binding specificities, accounting for biases and peculiarities of particular experimental techniques. Efficient and reliable computational models of transcription factor binding sites have remained in the focus of computational genomics for many decades^3,4^. The effort in improving TFBS modeling was stimulated by the demand from the areas of regulatory biology and genetics, fuelled by progress in experimental techniques, the rapidly growing amount of available experimental data, the development of efficient algorithms for handling sequences and recognizing recurrent text patterns, and the availability of computational power^5^.

The classic model of the TF binding specificity is the position weight matrix, PWM, which assumes independent nucleotide contributions to the binding specificity^4^. Many previous algorithms were developed to build the optimal PWMs from various types of experimental data; PWMs are widely represented in literature and databases and come with a rich software toolbox. Yet, the assumption of independence has usually been considered a fundamental limitation of PWMs, expected to cap their predictive power. Various more advanced models have been introduced to account for interdependent contributions of neighboring and distant nucleotides within binding sites^5^. Surprisingly, despite many individual examples of advanced models outperforming PWMs, it is still unclear if the advanced models provide a considerable gain across the whole spectrum of transcription factors and practical tasks. Furthermore, even the optimal selection of the best algorithm for the construction of PWMs is still unclear, as benchmarking studies remained sporadic and usually were focused on synthetic data^6^, data from one experimental platform^7,8^, or evaluated existing non-optimized PWMs^9,10^.

All in all, the diversity of the TFBS models, the lack of an established codebase for practical applications of advanced models, and the absence of commonly accepted benchmarking data and protocols altogether limit our understanding of the quantitative underperformance of standard PWM models and the potential of advanced models built upon different principles.

Recently, we have performed a large-scale benchmarking of PWMs and DNA motif discovery tools for PWM derivation^11^. The benchmarking was empowered by rich experimental data for several hundred human transcription factors for which the DNA binding specificity was assessed in five distinct experimental assays. However, this study included only a limited selection of tools, and several top-performing tools were run by their authors, which apparently put them into a privileged position. Furthermore, the entire study was focused solely on PWMs, without any systematic analysis of complex models. To fill these gaps, we organized IBIS, the open challenge in Inferring Binding Specificities of transcription factors, which was inspired by the success of the DREAM and CAGI challenges^12,13^, which continuously bring the wisdom of crowds to the fair benchmarking table.

In IBIS, we utilized previously unpublished experimental data for 40 transcription factors, including poorly studied proteins and a few well-studied positive controls. The data were obtained using five experimental platforms assaying protein binding to genomic and artificial DNA sequences. We rigorously tested PWMs and more complex models constructed using <u>a</u>rbitrary <u>a</u>dvanced <u>a</u>pproaches (AAA, Triple-A) with a diverse set of benchmarking protocols and performance metrics, which were openly described before the challenge started. Here, we report IBIS results, discuss the best-performing motif discovery tools, explore the performance gain of triple-A models over PWMs, highlight the general power of benchmarking in selecting reliably the best models, and provide a software platform and standardized data for further development of DNA motif discovery and TFBS modeling tools in the years to come.

## Results

### Design and implementation of the IBIS Challenge

We conceptualised the IBIS challenge to comply with several key principles. First, both classes of models, PWM and triple-A, developed by the participants must be comparatively assessed on the same data, with clear separation of the train and test data slices to prevent any information leakage; the test data labels must be open to participants upon completion of the challenge. Second, for fair benchmarking and interpretable results, the final performance metrics should be announced before the challenge and applied to all models and across all data types, whenever possible. Third, the models of DNA binding specificity must be inferred from the DNA sequence, without referring to extra layers and other types of genomic information such as chromatin accessibility or evolutionary conservation. Fourth, the comparison must be cross-platform: we would like to evaluate the models’ capacity to describe the genuine binding sequence specificity of a transcription factor, not the surrounding genomic context or specific bias of a single experimental platform.

To avoid information leakage^14^ from previously published experiments, we focused on poorly studied human TFs, the experimental data on which were recently obtained by the Codebook consortium^15^, but were not disclosed until the challenge ended. The nature of Codebook data allowed us to perform a cross-platform evaluation, that is, the inferred models were primarily benchmarked on experimental data obtained from platforms not included in training. IBIS relied on the data from five platforms: two assays with genomic DNA (ChIP-Seq^16^ and HT-SELEX with genomic DNA fragments^17^, abbreviated below as CHS and GHTS, respectively), and three assays with synthetic (artificial) oligonucleotides (simple HT-SELEX, HTS^18^, SMiLE-Seq, SMS^19^, and protein-binding microarray, PBM^20^).

Based on these data, we introduced two primary IBIS disciplines: transferring knowledge from artificial to genomic sequences (A2G, with binding specificity models inferred at synthetic data and tested at ChIP-Seq and GHT-SELEX) and vice versa (G2A, with models inferred at ChIP-Seq and GHT-SELEX and tested at the synthetic data). The A2G setup was designed to reflect a practical use case of predicting binding regions in a genome using a model of protein-binding specificity measured *in vitro* against artificial oligonucleotides. The G2A setup to predict TF binding to artificial oligos using models trained on genomic binding seems less practical, but it yields important information on whether the models can capture genuine binding specificity in a complex genomic context. Additionally, as a basic check of models’ predictive capabilities (secondary IBIS disciplines), we performed an optional within-experiment-type assessment (WET) with the model tested at independent experimental replicates obtained with the same platform but excluded from the training data. In total, the IBIS data included results of 372 experiments for 40 human TFs (see **Data Availability** and **Supplementary Data SD1**).

To establish fair and straightforward benchmarking, we preprocessed the data for all TFs uniformly for each platform, arranged train/test splits, and then merged and anonymized the test data across TFs for blind evaluation (see **Methods** and **Supplementary Figure SF1**). Primarily, we considered the binary classification between bound (positive class) and unbound (negative class) sequences, employing several alternative strategies to assemble the negative class datasets, controlling for basic confounders (such as GC composition or local genomic context), see Methods and the extensive IBIS online documentation for more details (https://ibis.autosome.org/docs/). For HT-SELEX, we additionally estimated rank correlation between predictions and HT-SELEX cycles, assuming stronger binding sites to preferably belong to later cycles.

The IBIS challenge ran from March to September 2024 in two stages. First, we opened the public Leaderboard where everyone could test their models on a small number of TF, five in A2G and five in G2A, see **Supplementary Table ST1**. 4 of 10 TFs were the well-studied positive controls (GABPA, NFKB1, LEF1, and RORB) to help the participants judge whether their pipelines converged properly to the binding motifs, similar to known previously. 28 teams participated in the Leaderboard stage, and the total number of scored Leaderboard submissions at the end of the challenge reached 1905 (1448 Triple-A, 457 PWMs, excluding the precomputed baseline solutions). The Leaderboard was envisioned to allow participants to test and adapt multiple approaches; thus, we did not impose a hard limit on the number of submissions from a single team. In the end, the absolute performance of the top-scoring solutions could be inflated due to multiple tests. Thus, from here on, we focus on the results of the Final stage. It is worth noting that the Final winner and runner-up teams matched closely with the Leaderboard ratings.

The Final stage started in midsummer 2024 and ended in September 2024. Only one Final submission per team was allowed, and as a handicap for the PWMs against the triple-As, a single submission could contain up to four alternative PWMs for each TF. The PWM and triple-A solutions were ranked independently in each discipline to highlight the best method in each category, and the PWMs versus triple-As comparison (described below in a separate section) did not affect the team rankings in the challenge.

The Final data covered 30 TFs without any overlaps with the Leaderboard, and only 2 of the 30 Final TFs were previously studied ’positive controls’. Thirty Final TFs (**Supplementary Table ST1**) belonged to 13 TFClass^21^ structural families, including both the well-studied bHLH-ZIP TFs (USF3) and less explored CG-binding SAND domain (SP140L) and CG-1 TFs (CAMTA1), which bind extra-short motifs (just the CG pair in a short context), and also a plethora of TFs with diverse zinc fingers, many of which bind prolonged and complex sequence motifs (**Table 1**). GCM1 (GCM-family TF, A2G) and MYF6 (MyoD-ASC-related TF, G2A) were included as well-studied positive controls. Our cross-platform train-test setup explicitly penalized the models for fitting the artifacts or biases characteristic of the particular assays of the training data. The primary motif subtypes were highly concordant in the training and test data (**Table 1**), but the range of tested binding affinities and varying diversity of the binding sites between platforms provided an extra challenge, especially for the triple-As, which, compared to PWMs, could be more prone to overfitting.

**Table 1.** Overview of the transcription factors of the IBIS challenge Final stage. The motif logos represent the top-ranking representative motifs for the particular type of assay obtained by motif discovery and benchmarking independently from IBIS and kept hidden during the challenge.

| TF (A2G) | DBD | ChIP-Seq (Test) | GHT-SELEX (Test) | HT-SELEX (Train/Test) | SMiLE-Seq (Train/Test) | PBM (Train/Test) |
| --- | --- | --- | --- | --- | --- | --- |
| CREB3L3 | bZIP |  |  |  |  | - |
| FIZ1 | C2H2 ZF |  |  |  |  | - |
| GCM1 | GCM |  |  |  |  |  |
| MKX | Homeod. |  |  |  |  |  |
| MSANTD1 | MADF |  |  |  |  |  |
| MYPOP | Myb/SANT |  |  |  | - |  |
| SP140L | SAND |  |  |  |  |  |
| TPRX1 | Homeod. |  |  |  |  |  |
| ZBTB47 | C2H2 ZF |  |  |  | - | - |
| ZFTA | BED ZF |  |  |  |  |  |
| ZNF286B | C2H2 ZF |  |  |  | - | - |
| ZNF500 | C2H2 ZF |  |  |  |  | - |
| ZNF721 | C2H2 ZF |  |  |  | - | - |
| ZNF780B | C2H2 ZF |  |  |  |  | - |
| ZNF831 | C2H2 ZF |  |  |  |  | - |
| TF (G2A) | DBD | ChIP-Seq (Train/Test) | GHT-SELEX (Train/Test) | HT-SELEX (Test) | SMiLE-Seq (Test) | PBM (Test) |
| CAMTA1 | CG-1 |  |  |  |  | - |
| LEUTX | Homeod. |  |  |  | - |  |
| MYF6 | bHLH |  |  |  | - |  |
| PRDM13 | C2H2 ZF |  |  |  |  | - |
| SALL3 | C2H2 ZF |  |  |  | - | - |
| USF3 | bHLH |  |  |  |  |  |
| ZBED2 | BED ZF |  |  |  |  |  |
| ZBED5 | BED ZF |  |  |  |  |  |
| ZNF20 | C2H2 ZF |  |  |  | - | - |
| ZNF251 | C2H2 ZF |  |  |  |  | - |
| ZNF367 | C2H2 ZF |  |  |  |  | - |
| ZNF395 | C2H2 ZF |  |  |  |  | - |
| ZNF493 | C2H2 ZF |  |  |  |  | - |
| ZNF518B | C2H2 ZF |  |  |  | - | - |
| ZNF648 | C2H2 ZF |  |  |  |  | - |

### IBIS Solutions and Winners

19 teams submitted their solutions for the Final stage to compete in four primary disciplines (A2G-AAA, G2A-AAA, A2G-PWM, G2A-PWM) and the secondary WET disciplines (to explore performance in the context of a single assay); see **Supplementary Table ST2,** and the Final teams list at the IBIS website (https://ibis.autosome.org/challenge_teams/final). To identify the best solutions and rank teams, we reused the hierarchical ranking strategy (see **Methods**) from the Codebook Motif Explorer^11^ based on individual ranks obtained from multiple classification performance metrics and several types of negative (unbound) sequence sets. For benchmarking, we conducted PWM scanning of the test sequences at the organisers’ side, whereas for triple-As, we asked the participants to submit the predictions for the test data.

In total, 19 independent teams submitted solutions across four primary disciplines, drawing on a diverse array of approaches. For PWMs, IBIS participants employed multiple classic tools (MEME^22^, STREME^23^, RSAT^24^, HOMER^25^, SeSiMCMC^26^, ChIPMunk^27^), which were competing with more advanced tools based on motif representations extracted from convolution kernels of neural networks, as well as unorthodox approaches involving protein structure-based predictions or Kolmogorov-Arnold Networks (**Table 2**). In turn, triple-As included traditional machine learning approaches, such as gradient boosting^28^, as well as several types of deep learning solutions circling around Convolutional Neural Networks (CNNs), including CNN-Recurrent Neural Network (CNN-RNN) hybrids^29^ and CNNs with squeeze-excitation blocks^30^. Several teams employed ensembles of models to boost the performance of proposed solutions further (**Table 3**). Notably, while transformer-based and DNA language models were explored by participants during the Leaderboard stage, none of these solutions were brought to the Final stage.

**Table 2.** IBIS Final PWM solutions.

| Team | Training data platforms | Overall strategy | Special feature(s) | Motif discovery software | Probabilistic/PWM based | K-mer/word enumeration strategy | Internal benchmarking | Differential motif discovery | References |
| --- | --- | --- | --- | --- | --- | --- | --- | --- | --- |
| Bench Pressers | HTS, CHS, GHTS | A2G: ZMotif (CNN)<br>G2A: ZMotif (CNN) → STREME | 2-component Gaussian mixture model for HTS | ZMotif, STREME | ✗ | k-mers & suffix trees | ✓ | ✓ | 23,31 |
| Biology Impostor | All | A2G: RAP (PBM), HTS-IBIS (HTS),<br>k-mer frequencies (SMS)<br>G2A: k-mer frequencies | PWM length extension using information content | RAP, HTS-IBIS | ✗ | k-mers | ✗ | ✗ | 64,65 |
| callitmagic | All | MEME → AME | Multiple runs of MEME over an extensive grid of parameter settings | MEME, AME | ✓ | ✗ | ✓ | ✗ | 22 |
| ChatGPTusers | HTS, GHTS | A2G: Autoseed<br>G2A: HOMER | Multinomial method for generating PWMs | Autoseed, HOMER | ✗ | gapped subsequences, k-mers | ✗ | ✗ | 25,66 |
| Post Bioinformatic Disorder | CHS, GHTS | ChIPMunk applied to CHS, GHTS, or both | Peak summit as an informative prior for motif location | ChIPMunk | ✓ | ✗ | ✗ | ✗ | 27 |
| RSAT | All | peak-motifs → matrix-clustering → optimize-matrices-GA | PWMs optimization using a genetic algorithm | RSAT | ✗ | k-mers | ✓ | ✓ | 24,67 |
| sbi two | HTS, SMS, CHS, GHTS | MEME ↔ ModCRE ↔ AlphaFold3 | Protein structure-based approach | ModCRE, MEME | ✗ | ✗ | ✓ | ✗ | 68 |
| StochasticChaos | All | SeSiMCMC | MCMC-powered Gibbs sampler | SeSiMCMC | ✓ | ✗ | ✗ | ✗ | 26 |
| The Motifvators | GHTS | HOMER → STIMULUS | STIMULUS augmented Kolmogorov-Arnold Networks | HOMER, STIMULUS + KAN | ✗ | k-mers | ✗ | ✓ | 25,69 |
| Trituration | CHS | STREME | Motif length optimization based on performance estimates | STREME | ✓ | k-mers & suffix trees | ✓ | ✓ | 23 |

**Table 3.** IBIS Final Triple-A solutions.

| Team | Training data | Prim. disc. | Overall design | Special feature(s) | AAA / NN type | Optimizer, LR scheduler | Regularization | Framework / Software | Negatives, augmentations | Refs. | Loss function |
| --- | --- | --- | --- | --- | --- | --- | --- | --- | --- | --- | --- |
| Bench Pressers | CHS, GHTS, HTS | A2G, G2A | PWMs derived from CNNs and refined with STREME | Initialization with enriched gapped k-mers, distillation, Gaussian mixture model for filtering train sequences | CNN | Adam, SGD with warm restarts | Dropout, L1, L2 | TensorFlow, Keras, STREME | Dinucleotide shuffled negatives, shift, Gaussian blurring | 31,70 | Binary cross-entropy |
| Biology Impostor | HTS | A2G | Ensemble of CNNs with variable-length convolution filters | Averaging predictions for windows of different lengths, sliding windows | CNN | Adam, constant | Dropout, L1, L2, Early stopping | TensorFlow, Keras | Negatives: shuffled & alien; both strands | 71 | MSE |
| callitmagic | All | A2G, G2A | Motif finding (FIMO) with PWMs | Max score for top 12 PWMs | PWM-based | ✗ | ✗ | MEME Suite | All types of negatives | 22 | ✗ |
| ChatGPTusers | GTHS, HTS | A2G, G2A | Motif discovery (Autoseed) followed by PWM scanning (MOODS) | Multinomial PWM generation | PWM-based | ✗ | ✗ | Autoseed, Homer, MOODS | ✗ | 72 | ✗ |
| LovelsBind | CHS, GHTS | G2A | Gradient boosting on k-mer occurrences | TF name as feature, family-level models | CatBoost | ✗ | ✗ | scikit-learn, CatBoost | Negatives: random & alien (other prot. families) | 28 | ✗ |
| Medici | CHS, GHTS | G2A | CNN with SE blocks | Parallel convolutions, EfficientNet-like blocks | CNN | AdamW, LinearLR | Weight decay | PyTorch | Reverse complement, cyclic shift | 73 | Cross-entropy |
| mj | All | A2G, G2A | Two strand-scoring TCN | Combines forward/revcomp scores, accounting for confidence | TCN (CNN) | Adam, constant | Dropout | TensorFlow, Keras | Negatives: random & alien; reverse complement | 74 | Binary cross-entropy |
| Natural Killer | HTS | A2G | ResNet-like CNN | Pseudo-labelling, maximum from sliding windows | CNN | Adam | ✗ | PyTorch | Alien negatives | 75 | Binary cross-entropy |
| Pap | CHS, GHTS | G2A | Two-layer CNN | Several candidate models, voting from the ensemble | CNN | Adam | Dropout, L1, L2, Early stopping | TensorFlow | Shuffled negatives | 76,77 | Cross-entropy |
| pwmsandme | CHS, GHTS | G2A | Gradient boosting following joined logistic regression & k-nearest neighbors | TF-IDF embeddings on n-grams of equivariant k-mer tokens | CatBoost | ✗ | ✗ | scikit-learn, CatBoost | Each token represents both strands; alien negatives | 28 | Cross-entropy |
| Salimov and Frolov Laboratory | HTS, PBM | A2G | Hybrid CNN-RNN with skip connections | Multitask setting, sliding windows, parallel convolutions, Gaussian mixture model for reweighting and denoising PBM signal | CNN + biGRU | AdamW, cosine annealing | Weight decay, Early stopping | TensorFlow, Keras | Reverse complement, alien negatives for HTS | 78,79 | MSE for PBM, Cross Entropy for HTS |
| sbi two | CHS, GHTS, HTS, SMS | A2G, G2A | Structure-based PWM generation with ModCRE, combining multiple PWM hit counts using NN | Combination of multiple PWMs, built from structural predictions of TF interactions, with every possible DNA sequence | PWM-based + fully connected NN | Adam | ✗ | TensorFlow, Keras, FIMO, ModCRE | Alien negatives | 68 | Binary cross-entropy |
| The Motifvators | GHTS | G2A | Initialization with HOMER PWMs | Learnable spline-based activations on edges (KAN) | CNN + KAN | Adam / SGD | ✗ | PyTorch, STIMULUS | Alien negatives, reverse complement, random masking | 25,69 | Binary cross-entropy |
| Transcriptome | CHS, GHTS | G2A | Two-layer CNN with LSTM | Hybrid CNN-RNN | CNN + LSTM | Adam | Dropout | TensorFlow | Negatives: random, shades | 80 | Binary cross-entropy |
| Trituration | CHS | ✗ | Bayesian Markov Model | Search for optimal motif length | Markov Model | ✗ | ✗ | BaMM, STREME | Random negatives (dinucl. matched) | 34 | ✗ |
| chiCkeN pox gaNg | HTS | ✗ | EfficientNet-like CNN | EfficientNet-like architecture, grouped convolutions | CNN | AdamW, OneCycleLR | Dropout, weight decay | PyTorch, PyTorch Lightning | GC-matched aliens, Reverse complement | 35 | Binary cross-entropy |

**Table 4.** The IBIS Challenge timeline.

| Phase | Start | End | Description |
| --- | --- | --- | --- |
| User registration | March 8, 2024 | July 15, 2024 | Participants gained access to the challenge data by registering online via a GitHub account. |
| Online leaderboard | March 13, 2024 | Sep 1, 2024 | Participants performed *de novo* motif discovery and submitted PWMs and advanced model predictions for online evaluation. |
| Final submissions | Jul 15, 2024 | Sep 1, 2024 | Participants submitted their solutions for final evaluation: PWMs and/or predictions from advanced models. |
| Offline evaluation | Sep 1, 2024 | Nov 27, 2024 | The organizers benchmarked and verified solutions and determined the winning and runner-up teams. |
| IBIS Final Conference | Nov 27, 2024 | Nov 27, 2024 | An online conference announcing the winners and allowing them to present their solutions in scientific talks. |
| Post-challenge analysis | Fall 2024 | Fall 2025 | The members of the top-performing teams were analyzing the solutions in-depth and preparing the manuscript. |

To ensure the stability of the Final ranking (**Supplementary Table ST2**), we estimated the resulting ranks for each team across 1000 random subsamples obtained by dropping out any <TF, platform> combination with a probability of 0.25 and thus retaining on average 75% of the total pool (**Figure 1**). In the end, the ranking of the Winner and Runner-up teams remained unchanged, with one exception of the A2G-PWM discipline, where the bronze-ranked solution was treading on the heels of the runner-up team. Overall, the PWM winner and runner-up teams, *Bench Pressers* and *callitmagic*, took the lead in A2G, as well as G2A. Both teams relied on internal benchmarking schemes on the available training data to prioritize more reliable motif models, but the methods of initial PWM generation were completely different. The runner-up team, *callitmagic*, built their motif analysis pipeline fully on top of the classic MEME suite^22^. However, the winner, *Bench Pressers*, performed motif discovery using a recently developed CNN-based approach (ZMotif^31^), followed by refinement with STREME^23^.

**Figure 1.**
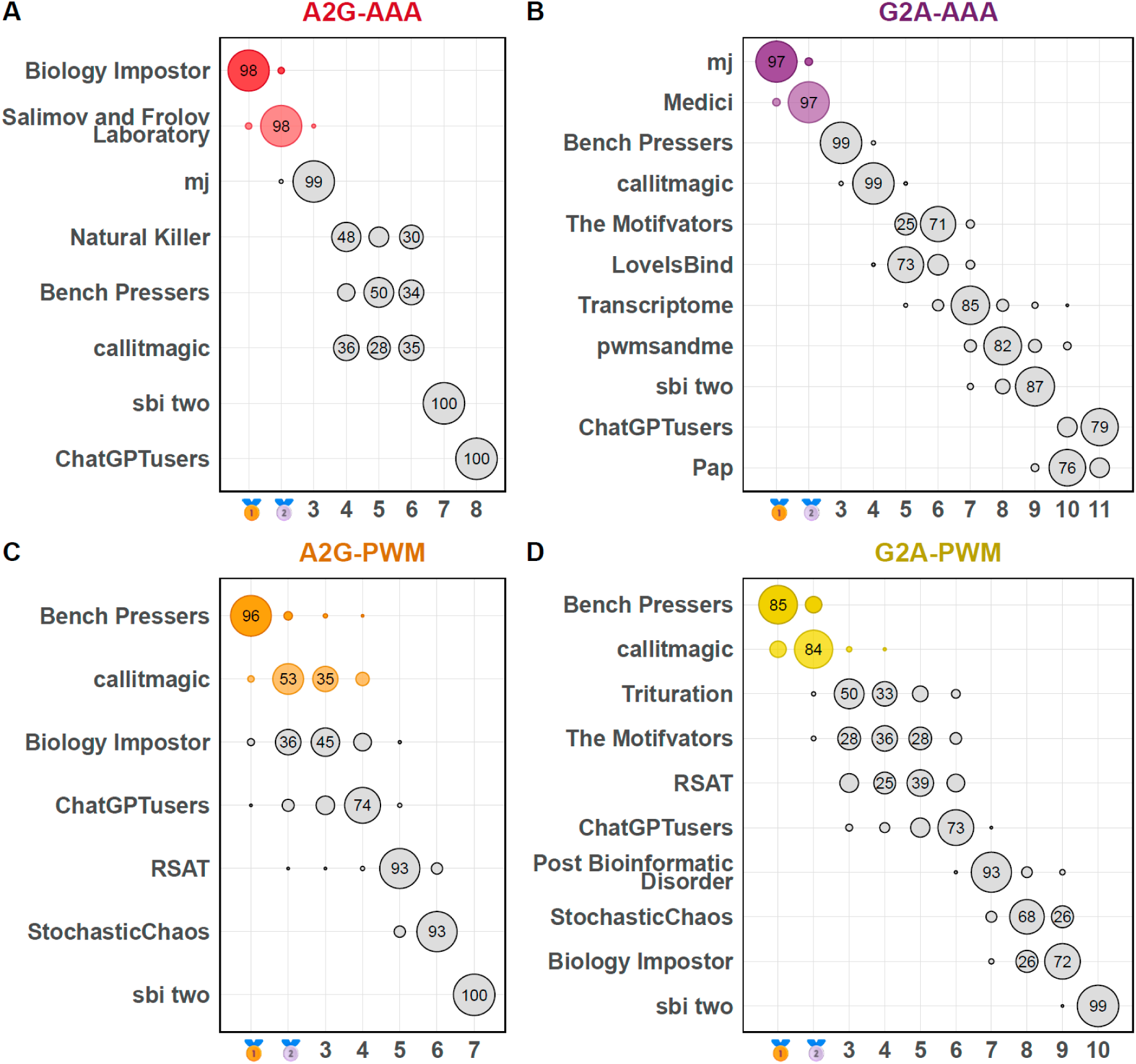
IBIS Final participants and winners, and the stability of the final rankings. The bubble size is proportional to the percentage of cases (the number in the bubble) where the team received a particular rank across 1000 random subsamples, where each combination of a TF and a data type had a 25% chance to be excluded from ranking. X-axis: ranks. **A**, **B**, **C**, **D**: primary IBIS disciplines, A2G-AAA/PWM and G2A-AAA/PWM.

Compared to PWMs, Triple-A approaches are easier to overfit and end up with a platform-specific model. Thus, they required a more focused effort in each discipline, and the winners and runner-up teams were different in the A2G and G2A disciplines, although all of them used variations of CNNs. A2G was dominated by *Biology Impostor* (winner) and *Salimov and Frolov Laboratory* (runner-up) with a CNN and a hybrid CNN-RNN, respectively. In turn, teams *mj* and *Medici* won G2A with a TCN (temporal CNN^32^, which *mj* trained as a CNN with dilation) and a CNN with a Mobile Inverted Bottleneck Convolution^33^.

Evaluation within the experiment type (WET) brought about several noteworthy cases. For PWMs, *RSAT* showed its versatility and took the bronze in WET-CHS, -GHTS, and -PBM; *Trituration* (with STREME) was the runner-up team in WET-GHTS; *StochasticChaos* (with SeSiMCMC^26^) was breathing down the leader’s neck in WET-PBM; and *ChatGPTusers*, using an interpretable deterministic algorithm based on local maxima of k-mers, placed second in WET-HTS (**Supplementary Figure SF2**). For TripleAs, *Trituration* (with a Bayesian Markov Model^34^) got the bronze in WET-CHTS and WET-GHTS, while *chICken pox gaNg* (with LegNet^35^) was the runner-up team in WET-GHTS. However, overall, the A2G and G2A winner and runner-up teams dominated in the secondary WET disciplines, solidifying the evidence of their superior performance.

### Triple-A models outperform PWMs, but by a limited margin and only when thoroughly designed

The ranking was convenient to identify the top-performing teams, but it did not show differences in the quantitative performance of solutions. For that, metrics should be quantitatively comparable across different datasets. We achieved this by normalizing each performance metric against a strong baseline. For the baseline, we assembled the *MEX1* collection of PWMs reusing the strategy of the Codebook Motif Explorer^11^ with IBIS training data only. For each TF, the *MEX1* collection contains a single best PWM selected from PWMs obtained from IBIS training data with diverse motif discovery tools and tested in a series of benchmarks used in Codebook, again limited to IBIS training data^11^. Next, *MEX1* motifs were benchmarked along with the IBIS submissions.

For the normalized performance score of a motif, we used the log_2_-odds value computed against the same metric of the respective *MEX1* motif for the same TF. Positive values correspond to improved performance. Such normalization makes it possible to compare different performance metrics as well as metrics for different TFs and to visualize the average performance difference, be it gain or loss, across TFs (A2G and G2A in **Figure 2**, WET in **Supplementary Figure SF3**). For PWMs, we averaged the obtained performance scores across matrices of each team’s solution.

**Figure 2.**
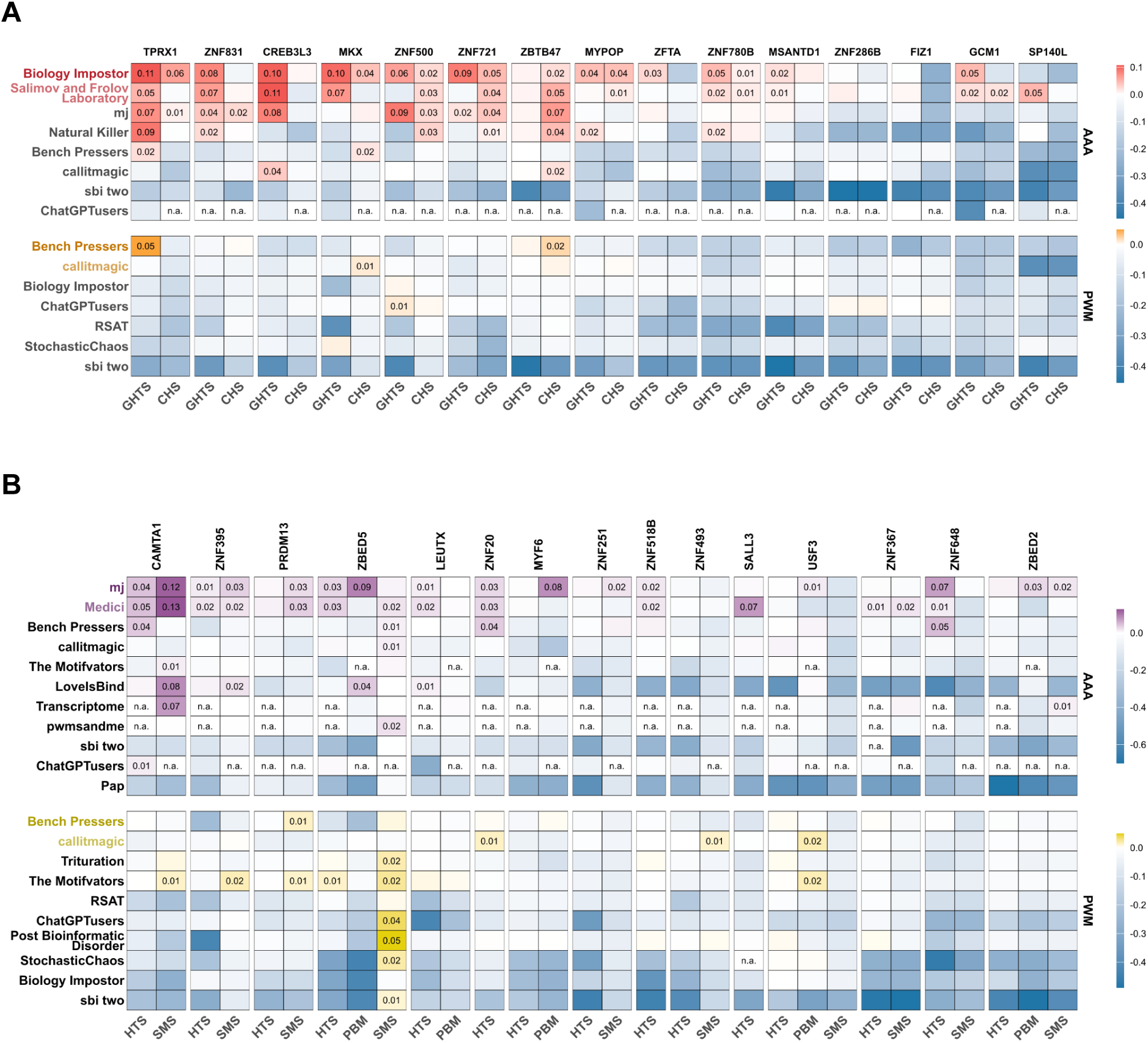
Normalized performance of different teams across TFs. **A**: A2G, **B**: G2A. Color scale: mean value across test datasets and different performance metrics. The values are log_2_-ratios versus the top-ranked PWMs obtained from IBIS data by the organizers; n.a.: solution for the combination of a TF and a data type not submitted for evaluation. Teams are ordered according to their rankings in the challenge. Values exceeding zero (improvement over the strong PWM baseline) are explicitly labeled.

On the one hand, PWM produced by IBIS participants performed on par with or even slightly worse than *MEX1*, thus failing to show any major improvement across the board. On the other hand, teams with the best PWM solutions relied on internal benchmarking to select a set of representative (submitted) PWMs for each TF from a wider set of candidates, and such benchmarking-backed approaches ultimately reached the *MEX1* level.

Considering triple-As, the best models outperformed PWMs not only in WET but in both A2G and G2A primary disciplines. Raw non-normalized values of individual performance metrics also support this observation (**Supplementary Figure SF4**). However, while the top-scoring triple-A models clearly outperformed PWM-based solutions from the competition and MEX, numerous deep learning–based approaches failed substantially in cross-experiment validation, in some cases performing far worse than the best PWMs. These results demonstrate that the application of deep learning to regulatory genomics remains highly challenging and is particularly susceptible to overfitting, noise, and experiment-specific artifacts.

It is also worth noting that the models’ performance varied heavily not only between TFs (which could have been expected) but also between different experimental platforms, such as CHS versus GHTS or HTS versus PBM. We attribute this variability to real differences in binding preferences measured in different experiments (see the representative motifs in **Tables 1-2**). The binding sites detected with different platforms vary in terms of the sequence diversity, distribution of their affinities, or even the presence of different motif subtypes (**Table 1**). Thus, we find it encouraging for further advances in the TFBS prediction accuracy that, despite platform-specific peculiarities, the top-performing Triple-A models were successful not only in WET but also in the primary IBIS cross-platform disciplines (A2G and G2A).

### Better TFBS prediction does not necessarily improve recognition of regulatory variant effects

An important application of DNA motif models is the prediction of variant effects in the regulatory regions of a genome^8^. The better performance of triple-A models suggests that such models сould outperform PWMs in this context as well. To compare the performance of Triple-A models with that of PWMs in this setting, we asked the participants to predict allele-specific TF binding in genomic regions containing regulatory single-nucleotide polymorphisms (rSNPs) detected directly in CHS and GHTS data^15^.

For benchmarking, we focused on the concordance of predicted and observed allelic preferences, i.e., whether the difference in predicted binding between alternating alleles agreed with the direction of the allelic imbalance (preferred binding to the reference allele, Ref, or the alternative allele, Alt) observed in ChIP-Seq or genomic HT-SELEX experiment. This approach allows for measuring the quantitative performance of a model without relying on a reference negative set of neutral SNPs.

Technically, for each model, we compared predictions for Ref and Alt alleles, and estimated the area under the ’concordance curve’, which reflects the fraction of the concordant cases, i.e., how often the true allelic preference towards Ref or Alt was correctly recognized by the model depending on the predicted binding specificity for the region encompassing the rSNP (the maximal predicted score of Ref and Alt), see **Methods** and examples in **Supplementary Figure 5**. For Triple-As, the scoring was performed by the authors of each solution, whereas for PWMs, we scored the variants with PERFECTOS-APE^36^. For each submission, we calculated the area under the concordance curve, where the X-axis reflects the number of SNPs with a score passing a given threshold (number of hits) and the Y-axis reflects the fraction of concordant hits. The results are shown in **Figure 3**; for PWM submissions, the values were averaged over 4 PWMs. Note that here the MEX set used for reference comprised the top 4 PWMs per TF (instead of a single PWM used for normalization in **Figure 2**) for a fair comparison to the IBIS submissions of 4 PWMs. As the number of rSNPs varies dramatically between TFs, we used the weighted mean area under the curve to obtain the global estimate for each team.

**Figure 3.**
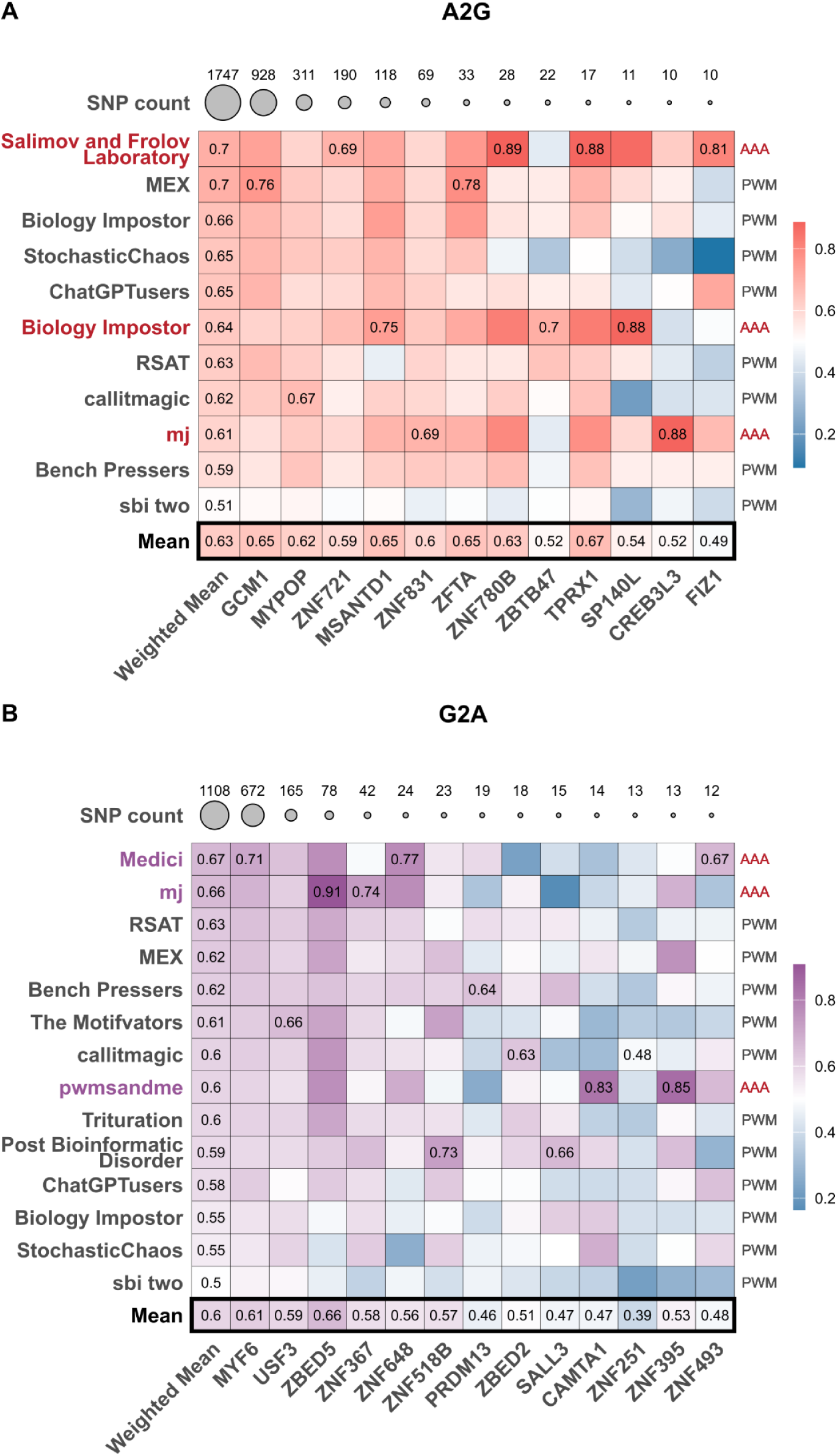
IBIS models performance in recognizing the allelic preferences of individual transcription factors. **A**: A2G models, **B**: G2A models. Color scale shows the area under the concordance curve (0.5 is the expected level for random predictions; 1 is perfect performance). Maximum values reached for each TF (column) are labeled explicitly. The number of tested SNPs for each TF is labeled on top and illustrated by the circle size.

The rSNP evaluation results partly agreed with those of the main benchmark, especially in G2A, where top-performing triple-A solutions took the medals, followed by MEX (**Figure 3**) and RSAT, which became the best of the PWM solutions. The layout in A2G changed dramatically: PWMs performed surprisingly well, with *MEX* being the second best and losing only to the Triple-A solution of the original runner-up team.

Special attention should be paid to the lack in this layout of any differences in the AUC scale between A2G and G2A: surprisingly, models trained on the artificial data were not less successful in recognizing regulatory variant effects in genomic regions than the CHS/GHTS-based models. Not only does this underline the generalization and predictive power of A2G models, but it also highlights the raw value of large-scale *in vitro* data.

### Insights on the benchmarking layout for TFBS motif models

The main goal of this study is to compare TFBS models based on different principles using data obtained at different experimental platforms in a fair manner. Thus, the challenge benchmarking layout was not only announced in advance, and included alternative scoring modes (for PWMs), multiple performance metrics, and multiple variants of experiment-specific negative datasets (both for PWMs and Triple-A models). This approach made it meaningless for participants to narrowly focus on a set performance estimate, and also allowed us to explore the tested alternatives in the post-challenge assessment.

First, we explored the relation between quantitative (regression) and qualitative (classification) performance metrics with HT-SELEX data. For SELEX, the affinity of protein binding sites rapidly increases with the selection cycles^37^. In IBIS, we computed the Kendall rank correlation (**τ_b_**) of the model score predicted for each read and the number of the HT-SELEX cycle in which the read was obtained. Across all TFs, **τ_b_** were highly correlated with AUPRC and AUROC of binary classifications into bound and unbound reads (Pearson ρ of about 0.9, see **Supplementary Figure SF6**). This was observed both for PWMs and Triple-A models. Such correlation additionally justifies using AUPRC and AUROC as a performance measure for HT-SELEX, and strongly supports using these quantities for other experimental platforms.

Next, we focused on PWM scanning methods. In IBIS, we evaluated PWMs using two scanning methods: the total occupancy score (the *sum-occupancy* of a sequence estimated with the position frequency matrix)^38^, and the highest scoring words (the *best-hit* in the sequence estimated with the log-odds weight matrix), see Methods. The two scanning methods displayed a comparable performance, but on average, the *sum-occupancy* scanning achieved a higher benchmarking performance (**Supplementary Figure SF7**, the strongest outliers are Homeodomain TFs MKX and TPRX1), especially exhibited for GHT-SELEX and, to a lesser extent, for ChIP-Seq data. This agrees well with the recent study reporting the contribution of multiple overlapping binding sites to TF occupancy^39^. Yet, for some combinations of TFs and datasets the best hit score still had an edge on the sum occupancy (e.g., ZFTA in ChIP-Seq). Also, in some practical scenarios the exact coordinates of the strongest binding sites of the *best-hit* might give additional information, so we did not discard the best hit score from our collection of metrics.

When preparing the IBIS competition, we designed multiple negative sets to gather the evidence on which type of negative set is harder to distinguish from positives (i.e., lowering the dataset-induced systematic classification error). For artificial sequences, the negative sets consisted of the input library (*input*) and the samples from read sets of non-relevant proteins (*aliens*). The performance estimates against different negative sets displayed similar efficiency (with slightly different distribution for AUPRC for SMiLE-Seq, see **Supplementary Figure SF8**), which concludes that any of the two approaches can be used, and in practice the selection should be made depending on the sequencing depth of the input library, and availability of experimental data for non-relevant TFs obtained in the same experimental batch with a comparable number of SELEX cycles. In turn, for genomic platforms, the regions in the vicinity of the peaks (the *shades* negative set, see Methods) comprised the least reliable control data, with many cases of inflated performance values, and the increased variability of performance estimates across TFs and models. The explanation could be that the selection of neighboring regions was not balanced by GC composition by design, and consequently was strongly biased by genomic fluctuations of nucleotide content. Using peaks of non-relevant proteins (*aliens*, see Methods) does not differ so much from randomly sampled genomic regions; it was still noisier, as it is non-realistic to ideally balance the nucleotide composition as thoroughly as for random genomic regions.

### Deeper exploration of the Triple-A models with the IBIS benchmarking suite

Setting up proper benchmarking is a complicated task in general and in regulatory genomics in particular. For IBIS, we developed *bibis*, a Python package with a set of utilities enabling flexible, information leakage-resistant, and efficient preparation of TF-binding datasets from diverse experimental data, and running a unified benchmarking for simple models such as PWMs and advanced arbitrary approaches. In addition, *bibis* also offers useful primitives such as an efficient genome-scale GC balancing strategy, allowing for custom schemes. Together with preprocessed Codebook data and documented benchmarking protocols for TFBS models, *bibis* sets the ground for further developments in the field. To further explore deep learning solutions on top of those prepared by IBIS participants and showcase future applications of *bibis*, we tested multiple existing architectures of neural networks, which are popular in regulatory genomics.

First, we included several approaches, which performed well in the Random Promoter DREAM Challenge 2022^40^, namely, LegNet, UnlockDNA, BHI, and DREAM-RNN. Second, we included Malinois, which is often applied specifically for modeling short nucleotide sequences^41,42^. Third, we evaluated three DNA language models with accessible fine-tuning protocols: DNABERT2^43^, GENA-LM^44^, and Nucleotide Transformer^45^.

To explore alternative architectural components of a single model, we started with LegNet and tested its several derivatives: LegNet-LSTM with the biLSTM layer, LegNet-DiffConvs with convolutions of different sizes (3, 6, 9, 12); LegNet-Max (based on LegNet-DiffConvs), in which the global average pooling was replaced with global max pooling; LegNet-Max-LSTM (based on LegNet-Max) with the addition of the biLSTM layer; LegNet-WE (based on LegNet-DiffConvs), with the SE block replaced by a WindowedSE block and global average pooling replaced with a windowed average pooling. Further, it has been shown recently that performance and transferability^11,46^ of machine learning in genomics can be enhanced by incorporating position weight matrices (PWMs). Therefore, we tested two modifications of the LegNet-Max architecture that included the top-4 MEX PWMs for the corresponding transcription factors in the first convolutional layers. However, this did not yield a performance improvement compared with other LegNet variants (**Supplementary Figure SF9**). Overall (**Figure 4**), LegNet-Max and LegNet-WE were the best among all tested models in both A2G and G2A settings, although the overall difference between approaches was limited (**Supplementary Figure SF9**), and most of them were competitive with the best IBIS submissions. It is noteworthy that LegNet-Max has only a few minor modifications compared to the original LegNet (LegNet-Pure, **Supplementary Figure SF9**) such as a variable kernel size in the first convolutional layer and global average pooling replaced with global max pooling in the SE block. Yet, it largely outperformed the original model for many TFs and platforms. This underscores that in neural networks, even minor architectural changes can substantially affect the generalizability of the learned signal representations.

**Figure 4.**
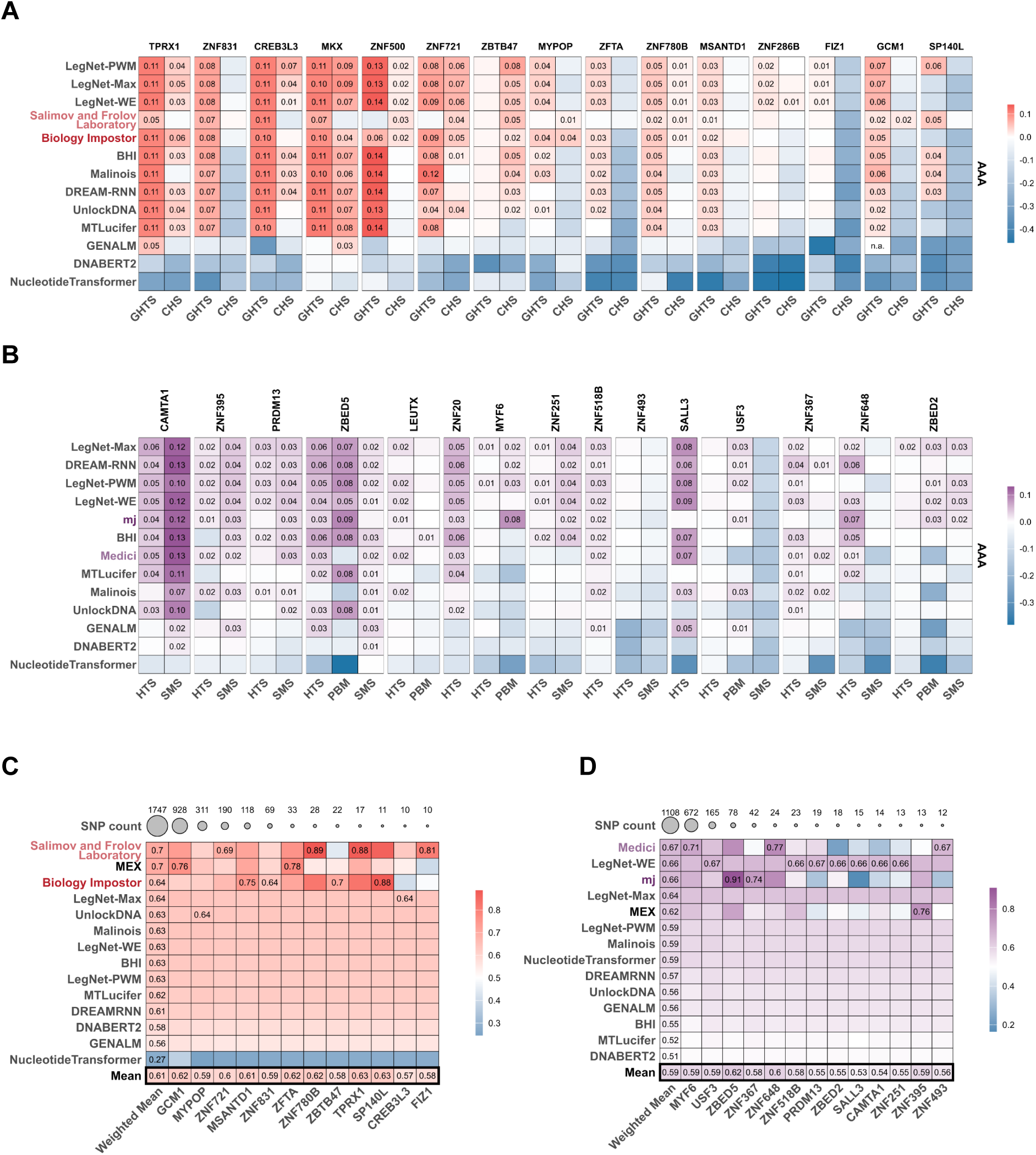
IBIS post-challenge benchmarking of diverse deep learning models. **A, B**: IBIS winning teams are included for comparison (A - A2G, B - G2A). Color scale: mean value across test datasets and different performance metrics. The values are log_2_-ratios versus the top-ranked PWMs obtained from IBIS data. Values exceeding zero (improvement over the strong PWM baseline) are explicitly labeled. **C, D:** IBIS models performance in recognizing the allelic preferences of individual transcription factors (C - A2G, D - G2A). Colors and plot structure are the same as in Figure 3.

Consistent with studies highlighting the limitations of applying DNA language models to regulatory tasks^47,48^, all three fine-tuned models performed dramatically worse than fully supervised approaches, generally achieving performance comparable to or below that of the top-1 MEX PWMs.

Despite global success in the primary IBIS disciplines, the post-challenge models did not truly succeed in predicting rSNP effects: once again, better TFBS prediction did not translate into better quantitative estimates of the variant effects. Specifically, all post-challenge models, despite better performance in A2G and G2A, were lagging behind the best IBIS solutions in the rSNP analysis.

## Discussion

The original concept of IBIS stems from GREEKC (https://www.greekc.org/), the COST Action that was paving the road to interoperable Knowledge Commons in the area of Gene Regulation information^1^. Among many facets of gene regulation, modeling TF binding specificity has been evolving for several decades, yet, till now, there is no consensus on the best motif discovery tools or the practical applicability of advanced models. The final arguments here can be found only by a technically fair evaluation of benchmarking results on previously unseen data. As the human TF binding motif dictionary is approaching its completion with uncovering binding specificities of the full inventory of TFs, the data from the large-scale Codebook project^15^ might have been the last chance to perform a large-scale fair evaluation of motif models for human TFs. Being able to use the Codebook data for running IBIS before the data has been published in open access, we profited from a rich, reliable, and diverse dataset to call for the wisdom of crowds in a joint attempt to identify the best-performing tools and decide if the advanced models truly learn the binding specificity and not the specifics of a particular experimental platform.

The design of the IBIS primary disciplines is highly demanding for participants, as there is a clear difference in preferred signals even between two genomic platforms, GHT-SELEX and ChIP-Seq (**Table 1**), while the gap between artificial and genomic data is even larger. Despite huge platform-specific effects and the increased risk of overfitting, the best Triple-A models successfully learned a generalized representation of binding specificity and outperformed PWMs not just in the original challenge setup, but also in annotation or regulatory SNPs. However, it proved to be much more difficult to build a working Triple-As, as many Triple-As failed to reach the PWM performance for particular TFs or even across the board.

The performance in binary classification (bound/unbound) in IBIS was highly correlated with the performance in the regression task of HT-SELEX cycles. Thus, one could expect that a better classification accuracy would translate directly into a better performance in quantitative estimation of changes in binding specificity upon single-nucleotide substitutions in TF binding sites. Yet, recognizing rSNPs affecting the binding of a particular TF remains the most challenging task, and the ranking of methods was reflecting the main IBIS benchmarks only to a limited degree. In the case of Triple-A models, the culprit could be the internal scoring scheme or pooling layers, which prohibit proper focusing on the single affected genomic binding site due to the presence of stronger binding sites in the close vicinity. Other explanations for the poor performance of triple-A models at genomic rSNPs could be overfitting that remains undetectable without validation datasets incorporating single-nucleotide mutation effects, or architectural choices in neural networks limiting the models’ capacity to capture subtle nucleotide-level sequence changes. This observation is consistent with previous findings in chromatin accessibility prediction, where a model well-performing on sequence-based accessibility prediction did not necessarily provide an accurate inference of SNP-induced effects on chromatin accessibility^49^.

Considering the diversity of approaches, convolutional neural networks were the top choice for modeling short regulatory sequences such as TFBS of individual TFs. This finding is consistent with numerous studies demonstrating the superiority of CNNs in modeling regulatory genomic grammar^40,47,50^. Most of the solutions profited from the reverse-complement data augmentation and model ensembling to improve the performance, although there was no single best method to preprocess the data or prepare the negative dataset. Unfortunately, far-out strategies such as the structure-based approach of *sbi two* or KAN of *The Motifvators* failed to perform competitively, although they overcame the simple consensus baseline during the Leaderboard stage. In IBIS, we limited the participants to use only the human genome and the training data from TF binding specificity assays, except for protein sequence information and DNA shape features^51^. Unfortunately, the former was not helpful, and the latter was not explored by any of the solutions. Although numerous studies have explored language models for DNA, their overall utility remains uncertain^48^, including their applicability to regulatory genomics^49,52^ and their potential to outperform well-designed supervised models^47^. While IBIS solutions did not include such approaches, the additional post-challenge assessment was unable to provide any arguments to support employing them for TFBS modeling at the time being.

## Perspectives and Conclusions

Modeling binding sites and binding specificities of individual transcription factors is indispensable in a protein-centric research universe, where the function of a transcription factor directly relates to its gene targets and TFBS locations in the genome regulatory regions. This "bottom-up" approach is deciphering the structure of regulatory sequences from individual binding sites, their heterotypic combinations, composite elements, and homotypic clusters. In this setting, TFBS modeling and prediction for a particular transcription factor have been essential and highly invested in. Nowadays, this approach is challenged with the new paradigm brought about by progress in machine learning, coupled with the genome-scale and even larger synthetic data from functional omics experiments profiling chromatin accessibility or transcription initiation, and massively parallel reporter assays. In this "top-down" approach, all regulatory regions are modelled as a whole, whereas the functional sequence motifs and involved TFs are identified *a posteriori* by model interpretation techniques. This paradigm shift diminishes the practical importance of genome-scale TFBS prediction for an individual TF, but does not discard the importance of TF-centric motif models, which remain highly valuable for diverse tasks, from focused annotation of regulatory variant effects to interpreting the top-down black-box models.

Running the IBIS challenge allowed us to revisit classic motif discovery methods and put them against Triple-A models in a fair benchmarking setup. In the end, we realized that the dispute of universal necessity and wider applicability of Triple-As as a substitute for PWMs is not over. In the complex setup of knowledge transfer between genomic and artificial sequences, on the one hand, cleverly designed CNNs backed by a proper data processing strategy were able to outperform PWMs across the board. On the other hand, PWM motif discovery was also quite successful, but the top-performing teams relied heavily on internal benchmarking procedures, which suggests that the motif discovery tools are to be amended with some motif selection strategy, and this stage is no less important than the motif discovery per se. All in all, the dictionary of non-studied motifs is almost exhausted in humans; thus, the next generation of TFBS modeling challenges will likely rely on the data from other species or use completely different setups.

To execute IBIS, we developed and fully tested a comprehensive framework for systematic evaluation of transcription factor (TF) binding prediction methods. IBIS provides extensively preprocessed, ready-to-use datasets derived from diverse experimental sources, featuring well-defined train/test splits across a wide range of TFs and a fair automated benchmarking system. The IBIS online benchmarking platform welcomes new submissions for Leaderboard TFs (https://ibis.autosome.org) and maintains the hierarchical ranking scheme enabling direct comparison against baseline and IBIS participants’ models. In turn, the wider set of data and benchmarks for Final TFs remains suitable for offline usage with *bibis*. All in all, built upon a solid and transparent set of evaluation metrics and accompanied by detailed documentation, the IBIS framework establishes a robust foundation for testing and advancing future sequence-level models of DNA binding specificity.

## Data Availability

The complete IBIS Challenge data, including train and test datasets, solutions submitted for final evaluation, and benchmarking results, are available at Zenodo^53,54^.

## Code Availability

The IBIS benchmarking package, *bibis*, is available at Zenodo^54^ along with the test data labels. The package source code is available on GitHub (https://github.com/autosome-ru/ibis-challenge). Code repositories prepared by the challenge participants are listed in **Supplementary Table ST2**. Code for evaluating IBIS models against the allele-specific binding sites is available on GitHub (https://github.com/autosome-ru/IBIS-rSNP). The implementation of the post-challenge deep learning models is available on GitHub (https://github.com/autosome-ru/postibis).

## Author contributions

**Writing - Original Draft, Writing - Review & Editing:** N.G., D.P., V.K., V.V., I.K., I.A.E., V.N., I.E.V., G.A., M.B., I.B., D.F., I.L., J.M., Y.O., G.R., D.S., N.S., I.T., Z.W., P.B., B.D., O.F., J.G., I.G., A.J., F.A.K., V.J.M., T.R.H., I.V.K.

**Data Curation, Investigation, Methodology:** N.G., D.P., V.K., V.V., I.K., I.A.E., V.N., I.E.V., P.B., B.D., O.F., J.G., I.G., A.J., F.A.K., V.J.M., T.R.H., I.V.K.

**Formal Analysis, Software, Investigation, Methodology:** N.G., D.P., V.K., V.V., I.K., I.A.E., V.N., I.E.V., S.B., G.A., M.B., I.B., D.F., I.L., J.M., Y.O., G.R., D.S., N.S., I.T., Z.W., V.S., P.B., B.D., O.F., J.G., I.G., A.J., F.A.K., V.J.M., T.R.H., I.V.K.

**Project Administration, Funding Acquisition, Resources:** P.B., B.D., O.F., J.G., I.G., A.J., F.A.K., V.J.M., T.R.H., I.V.K.

**Supervision:** D.P, P.B., B.D., O.F., J.G., I.G., A.J., F.A.K., V.J.M., T.R.H., I.V.K.

## Acknowledgments

We wholeheartedly thank the IT Group of the Institute of Computer Science at Martin Luther University Halle-Wittenberg for computational resources and, personally, Maximilian Biermann, for valuable technical support. Ivan V. Kulakovskiy personally thanks Vivien Marx for her kind encouragement and valuable advice throughout the project. We thank members of the GRECO consortium and, personally, Martin Kuiper, for supporting this project at its early stages within the GREEKC COST action.

## Methods

### Challenge overview

The IBIS Challenge took place in 2024. It was fully online and proceeded from March 8 to November 27. The primary interest of IBIS was the cross-platform modeling of transcription factor binding specificities, i.e., transferring knowledge from artificial to genomic sequences (A2G) or from genomic to artificial sequences (G2A). The secondary within-experiment-type (WET) disciplines were focused on models trained and tested with the data from the same platform. The challenge was built on top of unpublished data for 40 human transcription factors with 3-5 different types of experimental data (platforms) for each, with ChIP-Seq (CHS), HT-SELEX (HTS), and genomic HT-SELEX (GHTS) data available for all IBIS TFs. Experimental data were preprocessed and split between training and test slices (see below). The training data were made available upon user registration at the start of the challenge. The test data was made publicly available post-challenge. Several performance measures were used to evaluate the models for each type of experimental data, aimed at reducing the risks of overfitting models to a particular benchmark. Description and implementation of benchmarking protocols and performance measures for each type of experimental data were made public from the start of the challenge. During the Leaderboard stage, oversimplified baseline solutions were obtained with consensus binding sequence from the Codebook Motif Explorer (MEX)^11^. As the ’consensus’ baseline was easily beaten by all participants during the Leaderboard stage, we skipped it in the Final stage and compared PWMs between each other and, for assessing the overall improvement versus state-of-the-art PWMs, against the best of MEX PWMs built and tested solely on IBIS data. An overview of the IBIS challenge design is given in the **Graphical Abstract** and **Supplementary Figure SF1**. An overview of Codebook datasets borrowed by IBIS is given in **Supplementary Data SD1**. An overview of the solutions submitted by the participants of the Final stage is given in **Supplementary Methods**.

### General rules of IBIS

As IBIS was focused on binding sequence specificity, we expected the submitted models to rely on the nucleotide sequence only and on the training data supplied for this challenge only. Simultaneous usage of training data for multiple TFs and/or multiple types of experiments was allowed. External annotations were not allowed. Using arbitrary genomic regions (e.g., random genomic samples, or regions in the vicinity of the training regions) was allowed. Mixing training data for multiple TFs was allowed. Ensemble models were allowed. Mixing training data from multiple types of experiments was allowed and encouraged for the primary (knowledge transfer) disciplines (G2A and A2G). The IBIS used the data mapped to the hg38 genome assembly. Using an arbitrary genome or random sequences to pre-train an artificial neural network or to extract features was allowed if performed from scratch using the challenge data only, including any complex embeddings (if obtained solely from the hg38 human genome assembly). The exception was the usage of the precomputed biophysical features derived from the DNA sequences, such as the DNA shape features^55–57^, which were explicitly allowed. It was also allowed to use the RepeatMasker track for the hg38 genome assembly hosted at UCSC. Finally, it was allowed to use the protein-level metadata on transcription factors (including but not limited to protein sequence and domain information) available directly in UniProt (any human proteins, not only those used in IBIS). The features relying on external data (e.g., epigenetics tracks, predictions from pre-trained neural networks, motifs derived from third-party data, predefined sets of genomic regions, etc) were not allowed in all disciplines.

### Ensuring fair play and fair evaluation

To ensure fair evaluation of IBIS solutions, we followed several principles.

1. The test data was hidden until the announcement of the winners.
2. There was no overlap between Leaderboard and Final TFs.
3. Tests on genomic intervals relied on the whole-chromosome holdouts.
4. The organizers reserved the right to assess their solutions at the post-challenge stage, but these solutions were not included in the model ranking and did not affect the selection of the winners of the challenge.
5. All finalists must have provided a method write-up accompanying the Final submission.
6. The winning teams of the Final stage must have provided a reproducible pipeline to derive the submitted models from the training data.

### Ranking Strategy

In each discipline, for each PWM or AAA submission, we ran several benchmarks. AAA and PWM models were considered and ranked independently.

First, the ranks of individual submissions were obtained:

1. On the individual <experiment type, benchmark, TF> level;
2. On top of that, by rank aggregation across individual benchmarks (performance measures), yielding <experiment type, TF>-level ranks.

Then, for each discipline:

1. The teams’ best submissions were identified and re-ranked for each <experiment type, TF>;
2. For each team, the resulting ranks were aggregated across TFs and experiment types, forming the overall rank of the team.

For each team, any missing (i.e., non-submitted) models for particular TFs were considered as if they had the lowest possible ranks.

Note that PWMs of a single submission were scored independently, and each submitted PWM for a particular TF was scored in all benchmarks available for the TF and participated in all TF-relevant disciplines.

The rank aggregation was performed with the procedure borrowed from the DREAM-ENCODE challenge (https://www.synapse.org/#!Synapse:syn6131484/wiki/402026) and also used in Codebook MEX^11^: the ranking score on a particular test data set was computed as the sum of normalized ranking measures of *−log(r/(N+1))* across different benchmarking protocols (performance measures), where *r* is the rank of a particular solution for a specific performance measure, and *N* is the total number of solutions. The log-normalized rank-sum was hierarchically averaged following the iterative procedure as specified above, by re-ranking at each level and re-applying the log-transformation.

### Experimental data preprocessing

#### Protein-Binding Microarrays

There were data from two designs of protein-binding microarrays (PBMs) available, ME and HK: ME provided for model training, and HK was exclusive for the test data. These arrays have different numbers and layouts of the probes and were normalized independently.

The PBM data were preprocessed to account for systematic biases, e.g., arising from probe layout on the microarray. We used two types of preprocessing strategies: SD, spatial detrending with a window size 11×11, as tested in Weirauch *et al*.^7^, and QNZS, quantile normalization followed by probe-level Z-score estimation with the mean and std.dev. assessed for each probe across all available PBMs. Note that QNZS normalization was performed across all ME and HK Codebook PBMs^11^, not limited to the IBIS TFs.

The probe intensity values were log_10_-transformed before quantile normalization; the resulting SD-processed files contain the log_10_-transformed values in the ‘mean_signal_intensity’ column; the QNZS-normalized files contain Z-scores in the same column. The rest of the original content of the PBM files and file format were kept intact.

**Resulting file format of the train and test data:** original PBM file format with the normalized values in the ‘mean_signal_intensity’ column.

ChIP-Seq and genomic HT-SELEX

The peak calling and related analysis were performed with the unified GTRD ChIP-Seq pipeline^58^ as in Codebook MEX^11^.

**Reads preprocessing and alignment**

Both for ChIP-Seq and genomic HT-SELEX, the read alignment was performed with bowtie2 (default parameters and fixed --seed 0). For paired-end reads, we additionally specified --no-mixed --no-discordant --maxins 1000. Reported alignments were filtered by MAPQ score with samtools -q 10. For paired-end data, we additionally marked and removed PCR duplicates with Picard MarkDuplicates. Specifically for the genomic HT-SELEX data, before read mapping, we performed adapter trimming with cutadapt 1.15 (default parameters, AGATCGGAAGAGC as the adapter sequence: -a AGATCGGAAGAGC -A AGATCGGAAGAGC -o out.R1.fastq.gz -p out.R2.fastq.gz in.R1.fastq.gz in.R2.fastq.gz).

To have a balanced sequencing depth between experiments and controls and reduce computational load, the peak calling was performed against randomly sampled control data (10% of the total pooled set of control reads from the matching batch, sampling performed after the alignment step). For ChIP-Seq, the ’input DNA’ samples were used as the control. For genomic HT-SELEX, the ’zero-cycle unselected’ reads were used as the control. Paired-end control data were prioritized for paired-end ChIP-Seq when available in the same batch.

For peak calling, four peak calling tools (*macs2*, *pics*, *gem*, *sissrs*) were executed with default settings, except for *macs2*. For the latter, for single-end reads, we externally estimated the expected fragment length *$frag_len* using a strand cross-correlation approach with *run_spp.R* script from the ENCODE pipeline (dated Aug 29, 2016). Next, *macs2* was executed with --no-model --extsize *$frag_len* for single-end read alignments. For paired-end reads, we ran *macs2* in the paired-end mode (-f BAMPE --nomodel). Single-end peak callers (*pics*, *gem*, *sissrs*) were executed on paired-end data using alignments of the first reads in pairs (*samtools* -F 128 paired.bam).

### Identifying technically reproducible peak calls

The reference peak calls for each data set were obtained with *macs2*. Next, the technically reproducible peaks were selected by checking for overlap between the peaks of *macs2* and the peaks of all other peak callers (*pics*, *sissrs*, *gem*). The resulting *macs2* peaks supported by other peak callers were saved in the *macs2* peak format with an additional label specifying a peak-confirming peak caller.

For genomic HT-SELEX, the peak calling was performed separately for reads originating from each cycle. For the test data, the cycle with the highest number of reproducible peaks was selected as the most informative representative peak set; the rest were discarded. Peak sets from all cycles were provided in the train data.

**Resulting file format of the training and test data**: *macs2* peak calls with an additional column regarding supporting evidence from our peak callers.

### HT-SELEX and SMiLE-Seq

Complete unfiltered HT-SELEX data were provided for model training. The reads from each cycle of each experimental replicate were provided as raw, unprocessed FASTQ files. For HT-SELEX, binding sites may overhang the constant parts of the oligonucleotides that were physically present during the binding experiments, i.e., the binding sites may include parts of the primers and/or barcodes, which vary from experiment to experiment. Thus, we provided the sequences of the constant parts and barcodes.

The sequence design of the HT-SELEX reads was the following:

5’ ACACTCTTTCCCTACACGACGCTCTTCCGATCT

[BAR1] (40N) [BAR2]

AGATCGGAAGAGCACACGTCTGAACTCCAGTCAC 3’

where 40N is a random insert with a length of 40 bp. For convenience, we explicitly provide the sequences adjacent to the random insert on the 5’ and 3’ flanks, as this information may be useful for model training and when making predictions. Note that the experiment-specific barcode substrings are masked in the case of the test data.

The preparation of the HT-SELEX test data followed a different protocol. First, for each collection of HT-SELEX reads for a particular TF and cycle, we discarded duplicate reads. Next, across all such deduplicated collections, we kept only the reads that can be unambiguously attributed to a particular TF, replicate, and cycle. Next, the resulting reads were pooled across replicates.

The procedure for SMiLE-Seq was the same as for HT-SELEX, except there was only a single dataset (no ’multiple cycles’) for each replicate and TF.

The sequence design of SMiLE-Seq reads was the following:

5’ CGTCGGCAGCGTCAGATGTGTATAAGAGACAG

[BAR1] (40N)

CTGTCTCTTATACACATCTCCGAGCCCA 3’

where 40N is a random insert with a length of 40 bp. For a few proteins, the training data were taken from the previously published datasets (identifiable with SRA IDs, SRR*). The sequence design of previously published SMiLE-Seq data was the following:

5’ ACACTCTTTCCCTACACGACGCTCTTCCGATCT

[BC-half1] (30N) [BC-half2]

GATCGGAAGAGCTCGTATGCCGTCTTCTGCTTG 3’

with a 30 bp random insert.

For convenience, we explicitly provided the sequences adjacent to the random insert on the 5’ and 3’ flanks, as this information may be useful for model training and when making predictions. Note that the experiment-specific barcode substrings were masked in the test data.

**Resulting file format of the training and test data:** FASTQ.

### IBIS benchmarking protocols

For all protocols, the primary performance measures (AUROC, AUPRC) were computed with PRROC^59^. Kendall rank correlation for HT-SELEX was computed using the implementation provided in SciPy.

### General notes on position matrices

Only position frequency matrices (PFMs) were accepted. Before benchmarking, a pseudocount of 0.00001 was added to each value of a PFM. Internally, PFMs were additionally converted to log-odds position weight matrices (PWMs): log2([PFM + 0.00001] / 0.25). In each benchmarking protocol, PWM scanning was performed with PWMEval^10,60^ in two modes: PWM best-hit and PFM sum-occupancy. For PWMs, PWMEval accepts only integer values, so each PWM value was truncated to 5 digits after the decimal point and multiplied by 10^5^. The sum-occupancy^38^ estimation was computed as in^10^. The accepted PWM width (i.e., the minimal length of the scorable nucleotide ’substring’) was 5 to 30.

### Benchmarking on protein binding microarray probe intensities

This benchmark assessed the performance of models in solving the binary classification problem of discriminating high-intensity (positive) PBM probes from the rest. The benchmarking was conducted twice for SD- and QNZS-normalized PBMs, see the respective section in the data preprocessing description above.

**Positives:** for SD-preprocessed PBMs, probes passing ’mean + 4 std.dev.’ intensity threshold were considered positives as in the PBM DREAM challenge of Weirauch *et al*.^7^ (see ’Online Methods, AUROC of probe intensity predictions’). For QNZS-normalized PBMs, probes with a Z-score above 4 were considered as ’positives’. In case these estimates provided fewer than 50 positives, a minimum of 50 top probes was used instead.

**Negatives:** a random sample of the rest of the probes to obtain a matched GC-content distribution and a 1:10 (positives-to-negatives) class balance.

**Preventing overfitting and ensuring fair play:** results of independent PBM experiments of two different designs were used in the assessment, one for training and the other for testing. Predictions for probes lacking TF-specific positive or negative class labels were ignored in the assessment.

**Handling replicates:** when available, independent replicates were provided in the training data. At the test stage, replicates were scored independently, and the scores were averaged before the global rank-aggregation (see above).

**Performance metrics**: mean area under the precision-recall curve (mean AUPRC), mean area under the receiver operating characteristic (mean AUROC).

### Benchmarking on genomic regions using ChIP-Seq and genomic HT-SELEX peaks

This benchmark assessed the performance in solving the binary classification problem of discriminating ChIP-Seq/GHT-SELEX peaks (positives) from non-relevant negative sequences.

**Positives:** 301-bp long regions centered on the peak summits of the technically reproducible ChIP-Seq peaks, which were the *macs2* peak calls supported by (i.e., overlapping with) peak calls of any other peak callers (*sissrs*, *cpics*, *gem*).

**Negatives:** (1) ’shades’: regions located in the vicinity of the ChIP-Seq peaks. To generate shades, full-length peaks shorter than 300bp were extended in both directions to cover 300bp regions. For each resulting region, we created one 300bp shade region located at a random distance of 300-600bp from the region borders; the exact location and upstream/downstream placement were chosen randomly; (2) *’aliens’*: peaks of non-related proteins not overlapping any reproducible peaks of the target transcription factor; (3) *’random’*: random genomic regions with matched GC% composition.

Aliens were selected to reflect a distribution of %GC content of the positive set. Performance in reference to the shades, aliens, or random genomic regions was evaluated independently. For shades, positives and negatives were balanced 1:1. For aliens and random regions, positives and negatives were (im)balanced 1:2. When generating control regions, we ensured that there was no overlap with the original full-length peaks, and additionally required at least 300bp spacer between any positive and negative regions.

**Preventing overfitting and ensuring fair play:** train and test splits were based on chromosome holdouts: peaks at odd-numbered autosomes were provided for training, and peaks at even-numbered autosomes were used for testing. Positives and negatives of different proteins were mixed in a single set. For each particular TF, the predictions for sequences lacking TF-specific positive or negative class labels were ignored in the assessment.

**Handling replicates**: the data from experimental replicates were provided independently for training, when available. As test data, we used the single replicate with the largest number of peak calls. For GHT-SELEX, we used the results from a single cycle of a single replicate that yielded the largest number of peaks.

**Performance metrics:** area under the precision-recall curve (AUPRC), area under the receiver operating characteristic (AUROC).

### Benchmarking on SMiLE-Seq reads

This benchmark assessed the performance in solving the binary classification problem of discriminating positive (sequence in a SMiLE-Seq experiment) from negative (non-relevant) reads. For each dataset, duplicate reads were discarded before processing. Across TFs, no reads from the train data were included in the test data.

**Positives**: a complete subset of SMiLE-Seq reads. **Negatives**: *’aliens’*: reads of non-related proteins excluding the reads identical to those found in the positive set; *’input’*: reads originating from sequencing the SMiLE-Seq control libraries. Positives and negatives were (im)balanced 1:5. GC composition of the aliens and input negative data were matched to that of the positive set.

**Preventing overfitting and ensuring fair play**: positives and negatives of all proteins were mixed together in a single test set, where each entry should have been scored. When evaluating a particular model for a particular TF, predictions for sequences without class labels for the TF were ignored in the assessment.

For a particular protein, the reads present in the training data were explicitly excluded from the test data. Identical reads found in datasets of different proteins were also excluded from the test data to avoid uncertainty or information leakage.

**Handling replicates**: data from experimental replicates were provided independently for training, when available. For testing, the data from multiple replicates were pooled.

**Performance metrics**: area under the precision-recall curve (AUPRC), area under the receiver operating characteristic (AUROC).

SMiLE-Seq experiments yield high- and low-affinity binding sites in a variable proportion. To improve the benchmarking reliability across all TFs and datasets, in addition to the basic auROC and auPRC, we followed the ideas of Ambrosini *et al*.^10^ and computed these quantities taking the top 25% and top 50% predictions: the true positives and true negatives were ordered by the predicted score, the top predictions were selected from each list independently, and the auROC and auPRC scores were computed on the selected subset. The reported mean AUROC and mean AUPRC benchmarking scores are averages of (*auROC, auROC-25%, auROC-50%*) and (*auPRC, auPRC-25%, auPRC-50%*), respectively.

**A special note on PWMs**: when benchmarking PWMs, 20bp from constant flanking nucleotides were concatenated to the 5’ and 3’ ends of each read at the scoring stage to allow predicting the binding sites overhanging the constant part of the library, see the data preparation for details.

### Benchmarking on HT-SELEX reads

This benchmark assessed the performance in solving (1) the regression problem of predicting the SELEX cycle that yielded a particular read and (2) the binary classification problem of discriminating between positives and negatives.

Duplicate reads within each cycle were discarded before processing. Identical reads present in multiple SELEX cycles were discarded to avoid ambiguity. Across TFs, reads from the train data were explicitly excluded from the test data.

**Positives**: a random subset of 100,000 HT-SELEX reads of each cycle. An equal number of reads is sampled from each cycle, when possible. **Negatives**: *’aliens’*: reads of non-related proteins; *’input’*: reads originating from sequencing the HT-SELEX zero-cycle libraries. Positives and negatives were (im)balanced 1:2. GC composition of the aliens and zeros negative data were matched to that of the respective positive set.

**Preventing overfitting and ensuring fair play**: reads of all proteins were mixed together in a single test set, where each entry must have been scored. When evaluating a particular model for a particular TF, predictions for sequences w/o class labels for the TF were ignored in the assessment.

**Handling replicates**: when available, the complete data from experimental replicates were provided independently for training. For testing, the data from multiple replicates were pooled.

**Performance measures**: Kendall rank correlation (Kendall’s **τ_b_**, allows for ties in the ranks) for the regression problem; area under the precision-recall curve (AUPRC) and area under the receiver operating characteristic (AUROC) for the binary classification problem.

**Note:** similarly to the SMiLE-Seq benchmarking, the reported mean AUROC (MAUROC) and mean AUPRC (MAUPRC) benchmarking scores are averages of (*auROC, auROC-25%, auROC-50%*) and (*auPRC, auPRC-25%, auPRC-50%*), respectively. Further, different HT-SELEX replicates may have different numbers of cycles and different selection efficiency, hence the reported Kendall rank correlation value is the average of values computed separately for read sets selected from each replicate.

**A special note on PWMs:** similar to HT-SELEX, when benchmarking PWMs, 20bp from constant flanking nucleotides were appended to the 5’ and 3’ ends of each read at the scoring stage (see the data preparation for details).

### Post-challenge benchmarking against the allele-specific binding sites

The allele-specific binding sites of the TFs of interest were provided to participants in the context of 301bp genomic regions centered on the particular single-nucleotide polymorphism. We took into account that (1) the allele-specific binding sites may contain passenger variants that do not alter binding sites directly and (2) variant effects in stronger binding sites should be both easier to capture in allele-specific analysis and easier to predict for the models. With this in mind, for each model, we sorted the SNPs by the maximal reached score (considering both Ref and Alt alleles), and computed the fraction of concordant cases among SNPs passing the score threshold. For a ’random guess’ prediction the area under such curve is 0.5, and the curve to some extent resembles the precision-recall curve (although the sensitivity and specificity are assessed for different quantities): the fraction of concordant variants is, to some extent, analogous to ’precision’ (fraction of correct predictions), and the number of sites scoring above the threshold and tested for concordance can be thought of as ’recall’ (fraction of events captured). For PWMs, we scored the same set of SNPs using PERFECTOS-APE (with motif P-values estimated against the uniform nucleotide distribution). The lowest motif P-value (between the alternating alleles) was used to select the subset of ’predicted binding sites’, and the log-ratio of P-values was compared against the true allelic preferences. Since each team provided up to 4 matrices per TF, we used all of them independently and averaged the AUC values for visualization (Figure 3). The Triple-A model predictions were provided by the participating teams.

### Triple-A SNP scoring strategies

**Team *Biology Impostor*.** The predictions were generated by scoring three windows and averaging. For the GHTS experiments, we used the central (with respect to the read) 301, 101, and 51 bp, while for the CHS experiments, we used the central 301, 151, and 51 bp.

**Team *mj***. The predictions were generated for 41bp windows surrounding the variant positions via TCN with global max pooling.

**Team *Medici***. The sequences were cropped to a central 40 bp window (±20 bp around the midpoint) to mimic the sequence length used for HTS data evaluation in G2A. The resulting cropped sequences were then analyzed using the CNN models developed for the HTS data.

**Team *pwmsandme***. The predictions were generated by scoring 41bp windows centered on the variant positions.

**Team *Salimov and Frolov Laboratory*.** The predictions were based on the model output for the central windows of 59 bp (±29 bp around the SNPs).

### Post-challenge deep learning models

The implementation of the post-challenge deep learning models is available on GitHub (https://github.com/autosome-ru/postibis). To train each model, we used the same scheme based on the best practices of the top-performing and runner-up challenge solutions.

For training the A2G models, the positive sets were used ’as is’ and only HT-SELEX data were utilized. The negative set consisted of (1) sequences bound by other proteins and (2) sequences generated by single-nucleotide shuffling of the sequences from the positive set. The dataset was balanced in a 1:1:1 ratio for training:validation:test. The models were tasked to predict the highest cycle number in which a particular sequence was observed. All negative examples were assigned a label of 0.

For the G2A, the positive sets were used ’as is’. The negative set consisted of sequences bound by other proteins and sequences generated by single-nucleotide shuffling of the positive sequences, maintaining a 1:1:1 ratio, as for A2G. Both ChIP-Seq and GHT-SELEX datasets were used as training data, and the models were tasked to perform the binary classification.

For supervised models, for each protein we performed the cross-validation-style training of five models, each built with 80% of the available data, and their predictions were averaged. For G2A, the non-overlapping splits were arranged on the level of chromosomes; for A2G, the splits were random. For A2G, all HT-SELEX cycles were included in the training set.

All models were optimized with AdamW and scheduled with OneCycleLR, and the checkpoint with the best validation metric was saved for submission (Lightning’s ModelCheckpoint callback). For each model, hyperparameters were selected to maximize performance under cross-validation and kept consistent across transcription factors to reduce the risk of overfitting.

Following the ideas of the top-ranked A2G solution, for each A2G model, predictions were generated using 51-, 101-, and 301-nucleotide windows, and the final outputs were obtained by averaging the results across these window sizes as well as forward and reverse-complement strands. For G2A, we averaged predictions for forward and reverse-complementary strands. For rSNP effect prediction, each model was tasked with predicting TF binding within 41-bp windows centered on the variant, separately for the reference and the alternative alleles.

### DREAM Challenge models

These models represent the top performers and post-challenge models of the DREAM Promoter Challenge 2022, where the participants were tasked to predict promoter activity measured in massively parallel reporter assays using relatively short regulatory sequences^40^.

**UnlockDNA.** Hybrid CNN + Attention. To improve transferability between sequences of different lengths, the sinusoidal positional encoding was replaced with attention using linear biases^61^.

**BHI.** Hybrid CNN + LSTM. The *flatten* layer at the head of the neural network was replaced with global average pooling to enable operation on sequences of varying lengths.

**DREAM-RNN.** Hybrid CNN + LSTM architecture. The *flatten* layer at the head of the neural network was replaced with global average pooling to enable operation on sequences of varying lengths.

### LegNet variants

**LegNet-Pure.** The LegNet model used in Agarwal *et al*.^62^

**LegNet-DiffConvs.** A variant of LegNet in which the first convolutional block uses convolutions of different kernel sizes (3, 6, 9, 12) instead of a single uniform size for all convolutions, a modification inspired by the A2G solution of *Biology Impostor*.

**LegNet-LSTM.** The original architecture with a biLSTM layer inserted between the convolutional component and the final MLP layers.

**LegNet-Max.** A variant based on LegNet-DiffConvs, with the global average pooling in the SE block replaced by global max pooling to mitigate the effects of differing sequence lengths in the training and test datasets (see the explanation in **Supplementary Figure SF10**).

**LegNet-Max-LSTM.** A variant based on LegNet-Max, with the addition of a biLSTM layer. **LegNet-WE.** A variant based on LegNet-DiffConvs, with the SE block replaced by a WindowedSE block, substituting global average pooling with windowed average pooling to improve generalization across sequences of varying lengths.

**LegNet-PWM.** Convolutional layers were pre-initialized using the top 4 MEX position weight matrices (PWMs), and their weights were frozen during training. The outputs of these PWM-initialized layers were concatenated with the outputs of the mapper block.

**LegNet-PWM-Stem.** The outputs of the PWM-initialized layers were concatenated with the outputs of the stem block instead.

### Other fully supervised models

**Malinois.** CNN architecture from Gosai *et al*.^41^

**MTLucifer.** Hybrid CNN + Attention architecture from Reddy *et al*.^63^, optimized for short sequences.

### DNA Language Models

In all cases, while optimizing performance for each model, we adhered as closely as possible to the fine-tuning procedures recommended by the original models’ authors.

**DNABERT-2.** A pretrained 117M parameter transformer model from Zhou *et al*.^43^ During training, default hyperparameter values were retained, as they yielded the best average validation performance. The batch size was set to 512 for training and 64 for validation. Training was performed for five epochs to reduce the risk of overfitting.

**Nucleotide Transformer.** Pretrained transformer model from Dalla-Torre *et al*.^45^ We used the 500M-parameter version of this model pretrained on the human reference genome. Hyperparameters for NT were selected with consideration of its large size (500M parameters). The training and validation batch sizes were 128 and 64, respectively. The number of training epochs was limited to two, as additional epochs led to a measurable decline in predictive performance.

**GENA-LM.** Pretrained transformer model. We used the *gena-lm-bert-base-t2t* version of this model, pretrained on the T2T human genome assembly. To enhance model performance, the batch size was increased to 1024. Training was carried out for four epochs, with validation metrics used for model selection. The learning rate was set to 5 × 10^−4^ with a warmup ratio of 0.75. In addition, LayerNorm–Linear–SiLU blocks were incorporated as the final classifier.

## The IBIS Consortium

### IBIS Challenge Final Stage participants in alphabetical order

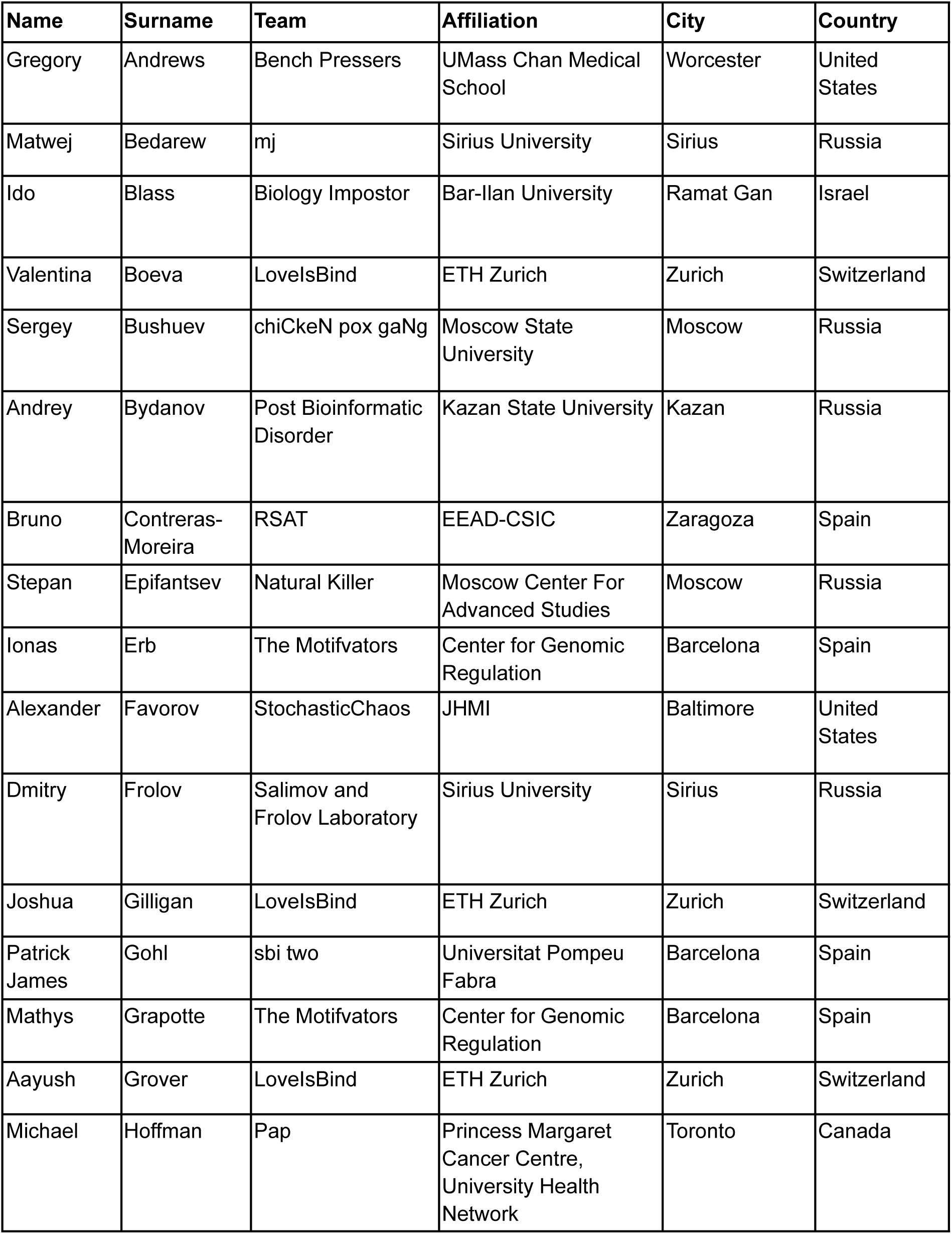
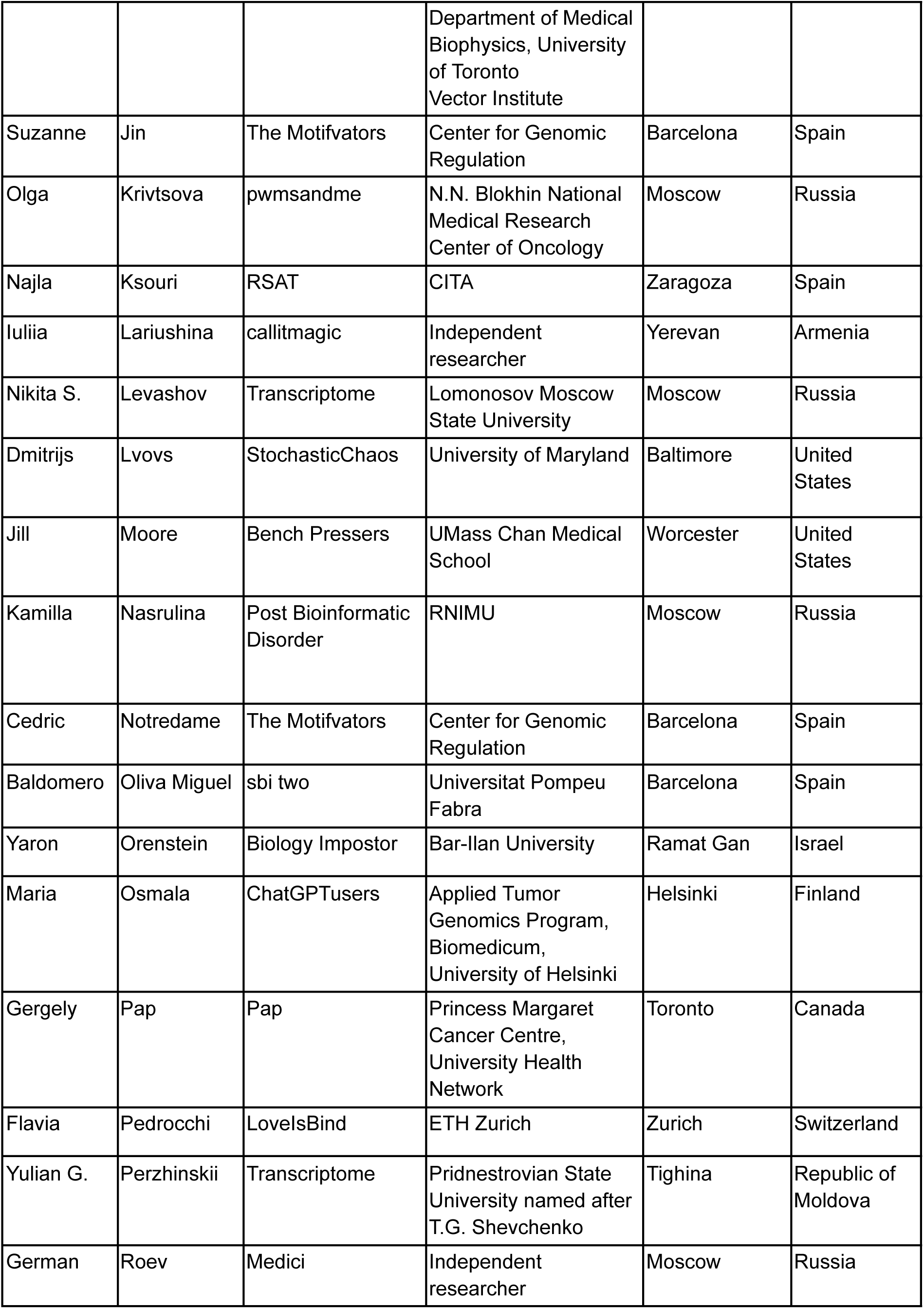
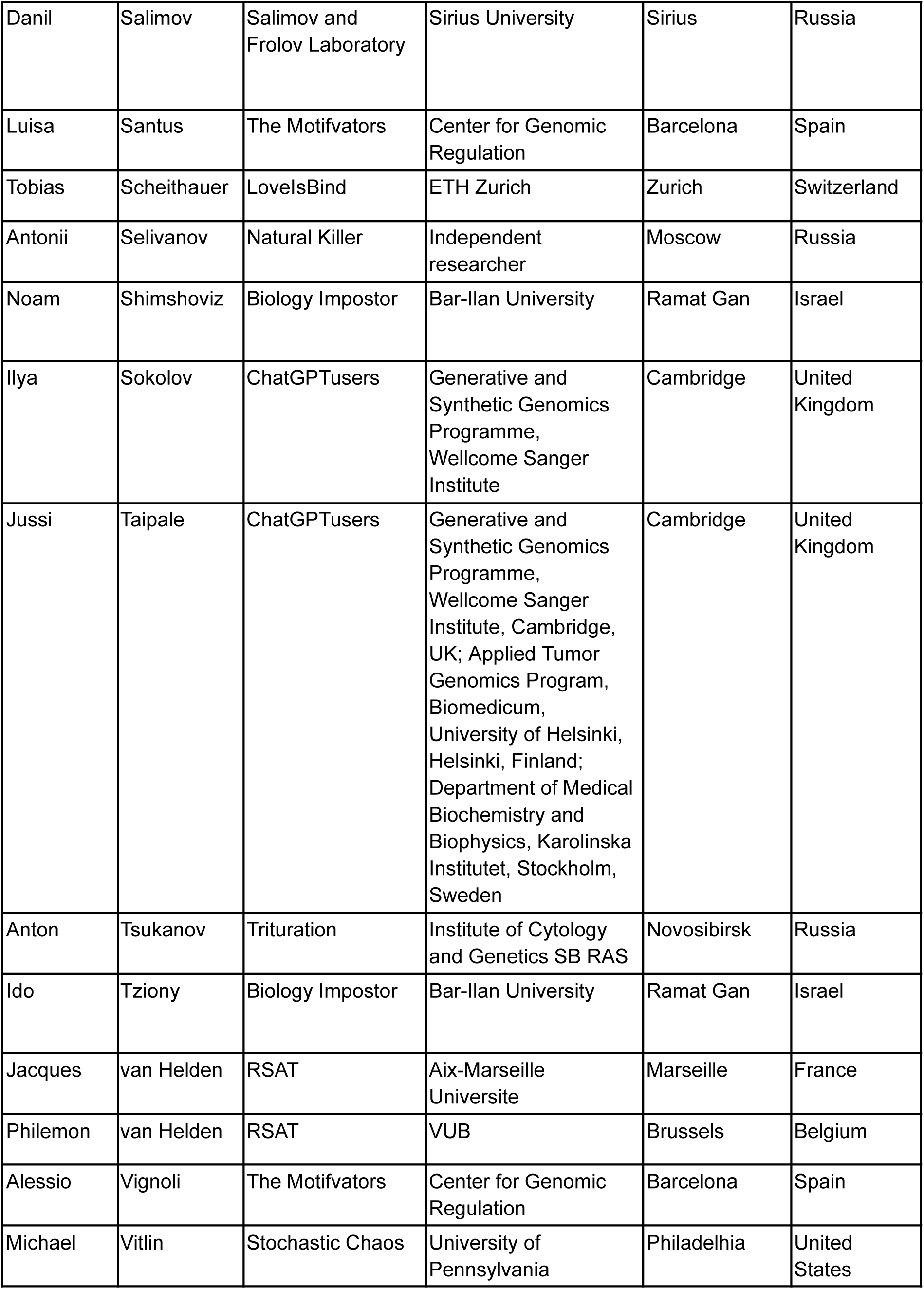
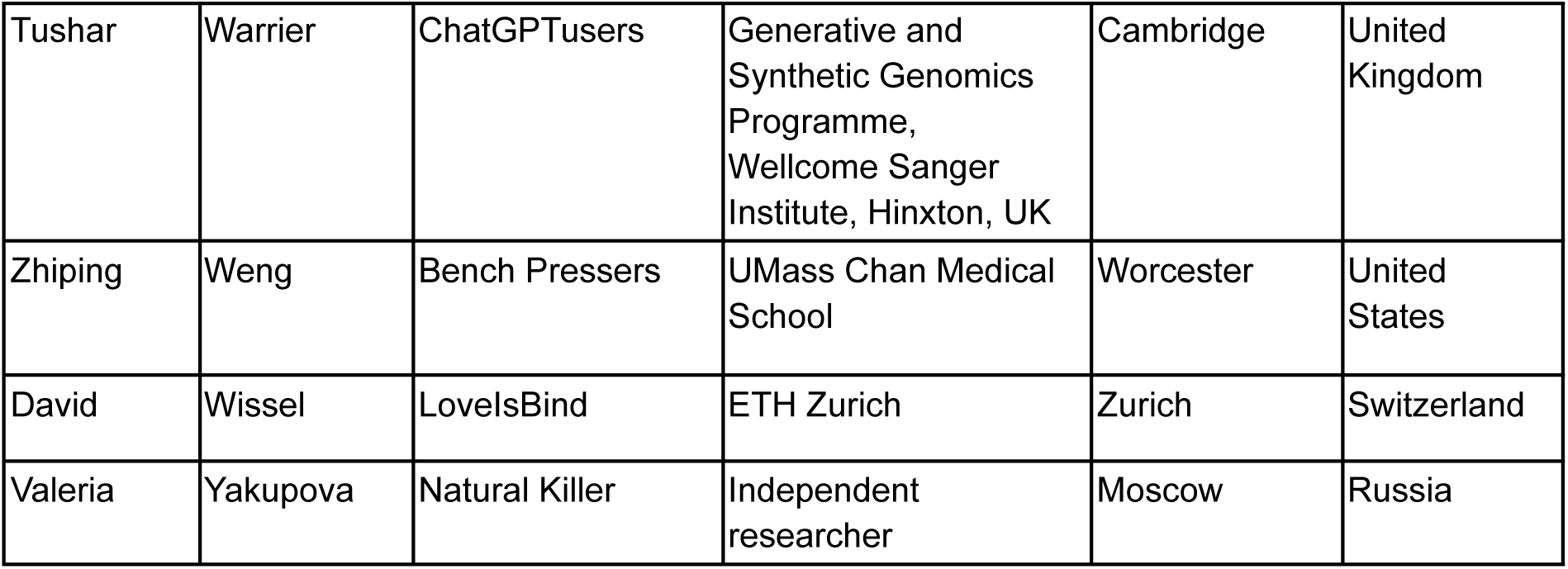

| Name | Surname | Team | Affiliation | City | Country |
| --- | --- | --- | --- | --- | --- |
| Gregory | Andrews | Bench Pressers | UMass Chan Medical School | Worcester | United States |
| Matwej | Bedarew | mj | Sirius University | Sirius | Russia |
| Ido | Blass | Biology Impostor | Bar-Ilan University | Ramat Gan | Israel |
| Valentina | Boeva | LovelsBind | ETH Zurich | Zurich | Switzerland |
| Sergey | Bushuev | chiCkeN pox gaNg | Moscow State University | Moscow | Russia |
| Andrey | Bydanov | Post Bioinformatic Disorder | Kazan State University | Kazan | Russia |
| Bruno | Contreras-Moreira | RSAT | EEAD-CSIC | Zaragoza | Spain |
| Stepan | Epifantsev | Natural Killer | Moscow Center For Advanced Studies | Moscow | Russia |
| Ionas | Erb | The Motifvators | Center for Genomic Regulation | Barcelona | Spain |
| Alexander | Favorov | StochasticChaos | JHMI | Baltimore | United States |
| Dmitry | Frolov | Salimov and Frolov Laboratory | Sirius University | Sirius | Russia |
| Joshua | Gilligan | LovelsBind | ETH Zurich | Zurich | Switzerland |
| Patrick James | Gohl | sbi two | Universitat Pompeu Fabra | Barcelona | Spain |
| Mathys | Grapotte | The Motifvators | Center for Genomic Regulation | Barcelona | Spain |
| Aayush | Grover | LovelsBind | ETH Zurich | Zurich | Switzerland |
| Michael | Hoffman | Pap | Princess Margaret Cancer Centre, University Health Network | Toronto | Canada |
|  |  |  | Department of Medical Biophysics, University of Toronto<br>Vector Institute |  |  |
| Suzanne | Jin | The Motifvators | Center for Genomic Regulation | Barcelona | Spain |
| Olga | Krivtsova | pwmsandme | N.N. Blokhin National Medical Research Center of Oncology | Moscow | Russia |
| Najla | Ksouri | RSAT | CITA | Zaragoza | Spain |
| Iuliia | Lariushina | callitmagic | Independent researcher | Yerevan | Armenia |
| Nikita S. | Levashov | Transcriptome | Lomonosov Moscow State University | Moscow | Russia |
| Dmitrijs | Lvovs | StochasticChaos | University of Maryland | Baltimore | United States |
| Jill | Moore | Bench Pressers | UMass Chan Medical School | Worcester | United States |
| Kamilla | Nasrulina | Post Bioinformatic Disorder | RNIMU | Moscow | Russia |
| Cedric | Notredame | The Motifvators | Center for Genomic Regulation | Barcelona | Spain |
| Baldomero | Oliva Miguel | sbi two | Universitat Pompeu Fabra | Barcelona | Spain |
| Yaron | Orenstein | Biology Impostor | Bar-Ilan University | Ramat Gan | Israel |
| Maria | Osmala | ChatGPTusers | Applied Tumor Genomics Program, Biomedicum, University of Helsinki | Helsinki | Finland |
| Gergely | Pap | Pap | Princess Margaret Cancer Centre, University Health Network | Toronto | Canada |
| Flavia | Pedrocchi | LovelsBind | ETH Zurich | Zurich | Switzerland |
| Yulian G. | Perzhinskii | Transcriptome | Pridnestrovian State University named after T.G. Shevchenko | Tighina | Republic of Moldova |
| German | Roev | Medici | Independent researcher | Moscow | Russia |
| Danil | Salimov | Salimov and Frolov Laboratory | Sirius University | Sirius | Russia |
| Luisa | Santus | The Motifvators | Center for Genomic Regulation | Barcelona | Spain |
| Tobias | Scheithauer | LoveIsBind | ETH Zurich | Zurich | Switzerland |
| Antonii | Selivanov | Natural Killer | Independent researcher | Moscow | Russia |
| Noam | Shimshoviz | Biology Impostor | Bar-Ilan University | Ramat Gan | Israel |
| Ilya | Sokolov | ChatGPTusers | Generative and Synthetic Genomics Programme, Wellcome Sanger Institute | Cambridge | United Kingdom |
| Jussi | Taipale | ChatGPTusers | Generative and Synthetic Genomics Programme, Wellcome Sanger Institute, Cambridge, UK; Applied Tumor Genomics Program, Biomedicum, University of Helsinki, Helsinki, Finland; Department of Medical Biochemistry and Biophysics, Karolinska Institutet, Stockholm, Sweden | Cambridge | United Kingdom |
| Anton | Tsukanov | Trituration | Institute of Cytology and Genetics SB RAS | Novosibirsk | Russia |
| Ido | Tziony | Biology Impostor | Bar-Ilan University | Ramat Gan | Israel |
| Jacques | van Helden | RSAT | Aix-Marseille Universite | Marseille | France |
| Philemon | van Helden | RSAT | VUB | Brussels | Belgium |
| Alessio | Vignoli | The Motifvators | Center for Genomic Regulation | Barcelona | Spain |
| Michael | Vitlin | Stochastic Chaos | University of Pennsylvania | Philadelphia | United States |
| Tushar | Warrier | ChatGPTusers | Generative and Synthetic Genomics Programme, Wellcome Sanger Institute, Hinxton, UK | Cambridge | United Kingdom |
| Zhiping | Weng | Bench Pressers | UMass Chan Medical School | Worcester | United States |
| David | Wissel | LovelsBind | ETH Zurich | Zurich | Switzerland |
| Valeria | Yakupova | Natural Killer | Independent researcher | Moscow | Russia |

### The Codebook / GRECO-BIT Consortium

#### Principal investigators (Steering committee)

Philipp Bucher, Bart Deplancke, Oriol Fornes, Jan Grau, Ivo Grosse, Timothy R. Hughes, Arttu Jolma, Fedor A. Kolpakov, Ivan V. Kulakovskiy, Vsevolod J. Makeev

### Analysis Centers

**University of Toronto (Data production and analysis):** Mihai Albu, Marjan Barazandeh, Alexander Brechalov, Zhenfeng Deng, Ali Fathi, Arttu Jolma, Chun Hu, Timothy R. Hughes, Samuel A. Lambert, Kaitlin U. Laverty, Zain M. Patel, Sara E. Pour, Rozita Razavi, Mikhail Salnikov, Ally W.H. Yang, Isaac Yellan, Hong Zheng **Institute of Protein Research (Data analysis):** Ivan V. Kulakovskiy, Georgy Meshcheryakov

**EPFL, École polytechnique fédérale de Lausanne (Data production and analysis):**

Giovanna Ambrosini, Bart Deplancke, Antoni J. Gralak, Sachi Inukai, Judith F. Kribelbauer-Swietek

**Martin Luther University Halle-Wittenberg (Data analysis):** Jan Grau, Ivo Grosse, Marie-Luise Plescher

**Sirius University of Science and Technology (Data analysis):** Semyon Kolmykov, Fedor Kolpakov

**Biosoft.Ru (Data analysis):** Ivan Yevshin

**Faculty of Bioengineering and Bioinformatics, Lomonosov Moscow State University (Data analysis):** Nikita Gryzunov, Ivan Kozin, Mikhail Nikonov, Vladimir Nozdrin, Arsenii Zinkevich

**Institute of Organic Chemistry and Biochemistry (Data analysis):** Katerina Faltejskova

**Max Planck Institute of Biochemistry (Data analysis):** Pavel Kravchenko **Swiss Institute for Bioinformatics (Data analysis):** Philipp Bucher **University of British Columbia (Data analysis):** Oriol Fornes

**Vavilov Institute of General Genetics (Data analysis):** Sergey Abramov, Alexandr Boytsov, Vasilii Kamenets, Vsevolod J. Makeev, Dmitry Penzar, Anton Vlasov, Ilya E. Vorontsov

**McGill University (Data analysis):** Aldo Hernandez-Corchado, Hamed S. Najafabadi

**Memorial Sloan Kettering (Data production and analysis):** Kaitlin U. Laverty, Quaid Morris

**Cincinnati Children’s Hospital (Data analysis):** Xiaoting Chen, Matthew T. Weirauch

