## Supplementary Data SD1 for "Inferring binding specificities of human transcription factors with the wisdom of crowds": ibis_motifs_Final_A2G.html

 

| TF | Platform | Slice | Replicate ID | logo-direct | logo-revcomp | Construct type | tfclass:id | tfclass:superclass | tfclass:class | tfclass:family | tfclass:subfamily | Motif length | Information content | GC% | Motif |
| --- | --- | --- | --- | --- | --- | --- | --- | --- | --- | --- | --- | --- | --- | --- | --- |
| CREB3L3 | CHS | test | THC\_0565 |  |  | FL | 1.1.7.2.4 | Basic domains | Basic leucine zipper factors (bZIP) | CREB-related | CREB3-like | 12 | 3.96 | 53.57 | CREB3L3.FL@AFS.Lys@beady-lemon-olm@Halle.Dimont@Motif\_1\_imw20\_astrained.ppm |
| CREB3L3 | GHTS.Lys | test | YWN\_B\_AffSeq\_F2\_CREB3L3 |  |  | FL | 1.1.7.2.4 | Basic domains | Basic leucine zipper factors (bZIP) | CREB-related | CREB3-like | 20 | 7.29 | 58.39 | CREB3L3.FL@AFS.GFPIVT@sloppy-puce-numbat@Halle.Dimont@Motif\_1\_w20\_astrained.ppm |
| CREB3L3 | HTS.IVT | train | YWC\_B\_GA40NGGTCAT |  |  | DBD | 1.1.7.2.4 | Basic domains | Basic leucine zipper factors (bZIP) | CREB-related | CREB3-like | 8 | 6.88 | 51.86 | CREB3L3.DBD@PBM.HK@randy-myrtle-hyena@Halle.Dimont@Motif\_1\_astrained.ppm |
| CREB3L3 | HTS.GFPIVT | test | YWM\_A\_AG40NCTTTTC |  |  | FL | 1.1.7.2.4 | Basic domains | Basic leucine zipper factors (bZIP) | CREB-related | CREB3-like | 12 | 3.96 | 53.57 | CREB3L3.FL@AFS.Lys@beady-lemon-olm@Halle.Dimont@Motif\_1\_imw20\_astrained.ppm |
| CREB3L3 | HTS.Lys | test | YWO\_A\_GG40NGGTTTA |  |  | FL | 1.1.7.2.4 | Basic domains | Basic leucine zipper factors (bZIP) | CREB-related | CREB3-like | 12 | 3.71 | 53.21 | CREB3L3.FL@AFS.Lys@breezy-taupe-rhinoceros@Halle.Dimont@Motif\_1\_imw20\_astrained.ppm |
| CREB3L3 | HTS.Lys | train | YWN\_A\_GG40NGGTTTA |  |  | FL | 1.1.7.2.4 | Basic domains | Basic leucine zipper factors (bZIP) | CREB-related | CREB3-like | 12 | 1.12 | 51.83 | CREB3L3.FL@HTS.Lys@scummy-ecru-coyote+nippy-lemon-corgi+stinky-persimmon-seahorse+snippy-cyan-dragon@faltejsk.ProBound@motif\_without\_ns.ppm |
| CREB3L3 | SMS | train | UT380-042 |  |  | FL | 1.1.7.2.4 | Basic domains | Basic leucine zipper factors (bZIP) | CREB-related | CREB3-like | 12 | 15.71 | 62.51 | CREB3L3.FL@AFS.GFPIVT@sickly-chestnut-wolf+stuffy-bistre-ragdoll@HughesLab.MEME@topk\_cycle=C3+C4\_k=5\_top=10000\_Motif1\_min3max30.ppm |
| ZNF831 | CHS | test | THC\_0507 |  |  | DBD | 2.3.2.4.257 | Zinc-coordinating DNA-binding domains | C2H2 zinc finger factors | Other with up to three adjacent zinc fingers | Other | 12 | 5.91 | 57.83 | ZNF831.DBD@AFS.GFPIVT@sunny-russet-vole@Halle.Dimont@Motif\_1\_imw20\_astrained.ppm |
| ZNF831 | GHTS.GFPIVT | test | YWS\_B\_AffSeq\_G10\_ZNF831-DBD |  |  | DBD | 2.3.2.4.257 | Zinc-coordinating DNA-binding domains | C2H2 zinc finger factors | Other with up to three adjacent zinc fingers | Other | 20 | 7.27 | 58.96 | ZNF831.DBD@AFS.GFPIVT@sunny-russet-vole+tacky-wisteria-crane+clammy-alizarin-mandrill@Halle.Dimont@Motif\_1\_w20\_astrained.ppm |
| ZNF831 | HTS.IVT | train | YWI\_A\_TC40NGCGATT |  |  | DBD | 2.3.2.4.257 | Zinc-coordinating DNA-binding domains | C2H2 zinc finger factors | Other with up to three adjacent zinc fingers | Other | 12 | 2.38 | 52.42 | ZNF831.DBD@HTS.IVT@tacky-cobalt-rattlesnake+trippy-linen-raccoon+nerdy-seashell-albatross+beady-charcoal-paradise@faltejsk.ProBound@motif\_without\_ns.ppm |
| ZNF831 | HTS.GFPIVT | test | YWS\_A\_TC40NAATCTC |  |  | NA | 2.3.2.4.257 | Zinc-coordinating DNA-binding domains | C2H2 zinc finger factors | Other with up to three adjacent zinc fingers | Other | 12 | 1.79 | 52.48 | ZNF831.NA@SMS@wheezy-corn-otter@faltejsk.ProBound@motif\_without\_ns.ppm |
| ZNF831 | SMS | train | UT380-477 |  |  | DBD | 2.3.2.4.257 | Zinc-coordinating DNA-binding domains | C2H2 zinc finger factors | Other with up to three adjacent zinc fingers | Other | 12 | 4.19 | 54.4 | ZNF831.DBD@HTS.GFPIVT@hazy-rust-falcon+dorky-denim-bison+sunny-tangerine-snake@faltejsk.ProBound@motif\_without\_ns.ppm |
| ZNF500 | CHS | test | THC\_0210 |  |  | FL | 2.3.3.19.1 | Zinc-coordinating DNA-binding domains | C2H2 zinc finger factors | More than 3 adjacent zinc fingers | ZNF500-like | 20 | 8.0 | 57.0 | ZNF500.FL@AFS.GFPIVT@pokey-copper-wrasse+hazy-cobalt-oriole+leaky-emerald-setter@Halle.Dimont@Motif\_1\_w20\_astrained.ppm |
| ZNF500 | GHTS.GFPIVT | test | YWR\_B\_AffSeq\_B8\_ZNF500-FL |  |  | FL | 2.3.3.19.1 | Zinc-coordinating DNA-binding domains | C2H2 zinc finger factors | More than 3 adjacent zinc fingers | ZNF500-like | 20 | 8.0 | 57.0 | ZNF500.FL@AFS.GFPIVT@pokey-copper-wrasse+hazy-cobalt-oriole+leaky-emerald-setter@Halle.Dimont@Motif\_1\_w20\_astrained.ppm |
| ZNF500 | HTS.GFPIVT | train | YWR\_A\_AT40NTCCCGA |  |  | FL | 2.3.3.19.1 | Zinc-coordinating DNA-binding domains | C2H2 zinc finger factors | More than 3 adjacent zinc fingers | ZNF500-like | 14 | 3.62 | 52.1 | ZNF500.FL@AFS.GFPIVT@hazy-cobalt-oriole@Halle.Dimont@Motif\_1\_imw20\_astrained.ppm |
| ZNF500 | SMS | train | UT380-357 |  |  | FL | 2.3.3.19.1 | Zinc-coordinating DNA-binding domains | C2H2 zinc finger factors | More than 3 adjacent zinc fingers | ZNF500-like | 17 | 16.88 | 55.66 | ZNF500.FL@HTS.GFPIVT@wimpy-xanthic-bonobo@AJolma.Autoseed@Multinom2\_Onehit\_Seed\_GGNHRGTGTAGACGSKBackgroundCyc1.ppm |
| ZBTB47 | CHS | test | THC\_0114 |  |  | FL | 2.3.3.29.2 | Zinc-coordinating DNA-binding domains | C2H2 zinc finger factors | More than 3 adjacent zinc fingers | ZNF652-like | 20 | 7.78 | 50.63 | ZBTB47.FL@AFS.Lys@cheeky-jade-quokka@Halle.Dimont@Motif\_1\_w20\_astrained.ppm |
| ZBTB47 | GHTS.GFPIVT | test | YWQ\_B\_AffSeq\_E6\_ZBTB47-DBD |  |  | FL | 2.3.3.29.2 | Zinc-coordinating DNA-binding domains | C2H2 zinc finger factors | More than 3 adjacent zinc fingers | ZNF652-like | 20 | 7.78 | 50.63 | ZBTB47.FL@AFS.Lys@cheeky-jade-quokka@Halle.Dimont@Motif\_1\_w20\_astrained.ppm |
| ZBTB47 | HTS.GFPIVT | test | YWP\_A\_GT40NTGTGGA |  |  | FL | 2.3.3.29.2 | Zinc-coordinating DNA-binding domains | C2H2 zinc finger factors | More than 3 adjacent zinc fingers | ZNF652-like | 12 | 1.96 | 49.06 | ZBTB47.FL@HTS.Lys@muggy-beige-collie+scummy-malachite-fox+hilly-turquoise-coral+homely-myrtle-koala@faltejsk.ProBound@motif\_without\_ns.ppm |
| ZBTB47 | HTS.GFPIVT | train | YWQ\_A\_GC40NTATAGG |  |  | FL | 2.3.3.29.2 | Zinc-coordinating DNA-binding domains | C2H2 zinc finger factors | More than 3 adjacent zinc fingers | ZNF652-like | 12 | 1.96 | 49.06 | ZBTB47.FL@HTS.Lys@muggy-beige-collie+scummy-malachite-fox+hilly-turquoise-coral+homely-myrtle-koala@faltejsk.ProBound@motif\_without\_ns.ppm |
| ZBTB47 | HTS.Lys | test | YWN\_A\_AC40NTCCTTG |  |  | FL | 2.3.3.29.2 | Zinc-coordinating DNA-binding domains | C2H2 zinc finger factors | More than 3 adjacent zinc fingers | ZNF652-like | 12 | 1.96 | 49.06 | ZBTB47.FL@HTS.Lys@muggy-beige-collie+scummy-malachite-fox+hilly-turquoise-coral+homely-myrtle-koala@faltejsk.ProBound@motif\_without\_ns.ppm |
| ZBTB47 | HTS.Lys | train | YWO\_A\_AC40NTCCTTG |  |  | FL | 2.3.3.29.2 | Zinc-coordinating DNA-binding domains | C2H2 zinc finger factors | More than 3 adjacent zinc fingers | ZNF652-like | 12 | 1.96 | 49.06 | ZBTB47.FL@HTS.Lys@muggy-beige-collie+scummy-malachite-fox+hilly-turquoise-coral+homely-myrtle-koala@faltejsk.ProBound@motif\_without\_ns.ppm |
| ZNF286B | CHS | test | THC\_0898 |  |  | DBD | 2.3.3.49.2 | Zinc-coordinating DNA-binding domains | C2H2 zinc finger factors | More than 3 adjacent zinc fingers | ZNF286 | 20 | 7.63 | 52.36 | ZNF286B.DBD@AFS.GFPIVT@leaky-khaki-slug@Halle.Dimont@Motif\_1\_w20\_astrained.ppm |
| ZNF286B | GHTS.GFPIVT | test | YWR\_B\_AffSeq\_G3\_ZNF286B-DBD |  |  | DBD | 2.3.3.49.2 | Zinc-coordinating DNA-binding domains | C2H2 zinc finger factors | More than 3 adjacent zinc fingers | ZNF286 | 20 | 7.63 | 52.36 | ZNF286B.DBD@AFS.GFPIVT@leaky-khaki-slug@Halle.Dimont@Motif\_1\_w20\_astrained.ppm |
| ZNF286B | HTS.GFPIVT | train | YWR\_A\_TG40NTGATCT |  |  | FL | 2.3.3.49.2 | Zinc-coordinating DNA-binding domains | C2H2 zinc finger factors | More than 3 adjacent zinc fingers | ZNF286 | 17 | 14.8 | 48.17 | ZNF286B.FL@HTS.GFPIVT@nippy-ruby-worm+seedy-goldenrod-rattlesnake+sleepy-magnolia-ray@autosome-ru.ChIPMunk@topk\_cycle=C1+C2+C3\_k=5\_top=10000.pcm |
| ZNF286B | HTS.GFPIVT | test | YWR\_A\_TA40NAGGCTT |  |  | FL | 2.3.3.49.2 | Zinc-coordinating DNA-binding domains | C2H2 zinc finger factors | More than 3 adjacent zinc fingers | ZNF286 | 16 | 19.45 | 49.93 | ZNF286B.FL@HTS.GFPIVT@nippy-ruby-worm+seedy-goldenrod-rattlesnake+sleepy-magnolia-ray@HughesLab.MEME@topk\_cycle=C1+C2+C3\_k=5\_top=10000\_Motif1\_min3max30.ppm |
| ZNF780B | CHS | test | THC\_0502 |  |  | FL | 2.3.3.67.12 | Zinc-coordinating DNA-binding domains | C2H2 zinc finger factors | More than 3 adjacent zinc fingers | ZNF283-like | 20 | 7.08 | 55.08 | ZNF780B.FL@AFS.GFPIVT@woozy-peach-kiwi+crabby-azure-drever+stealthy-tomato-swan@Halle.Dimont@Motif\_1\_w20\_astrained.ppm |
| ZNF780B | GHTS.GFPIVT | test | YWS\_B\_AffSeq\_H8\_ZNF780B-DBD1 |  |  | DBD | 2.3.3.67.12 | Zinc-coordinating DNA-binding domains | C2H2 zinc finger factors | More than 3 adjacent zinc fingers | ZNF283-like | 20 | 6.62 | 54.85 | ZNF780B.DBD@AFS.GFPIVT@snippy-russet-tamarin+clammy-carmine-warthog+cloudy-violet-rabbit@Halle.Dimont@Motif\_1\_w20\_astrained.ppm |
| ZNF780B | HTS.GFPIVT | test | YWS\_A\_TG40NCTGTGG |  |  | DBD | 2.3.3.67.12 | Zinc-coordinating DNA-binding domains | C2H2 zinc finger factors | More than 3 adjacent zinc fingers | ZNF283-like | 12 | 2.88 | 46.59 | ZNF780B.DBD@HTS.GFPIVT@cloudy-coral-javanese+squirrely-ochre-goose+hilly-tangerine-elephant@faltejsk.ProBound@motif\_without\_ns.ppm |
| ZNF780B | HTS.GFPIVT | train | YWS\_A\_AT40NAGCCTC |  |  | NA | 2.3.3.67.12 | Zinc-coordinating DNA-binding domains | C2H2 zinc finger factors | More than 3 adjacent zinc fingers | ZNF283-like | 12 | 11.22 | 50.77 | ZNF780B.NA@SMS@snappy-bistre-baboon@autosome-ru.ChIPMunk@topk\_k=5\_top=1000.pcm |
| ZNF780B | SMS | train | UT380-462 |  |  | NA | 2.3.3.67.12 | Zinc-coordinating DNA-binding domains | C2H2 zinc finger factors | More than 3 adjacent zinc fingers | ZNF283-like | 13 | 13.68 | 50.18 | ZNF780B.NA@SMS@snappy-bistre-baboon@autosome-ru.ChIPMunk@topk\_k=5\_top=500.pcm |
| ZNF721 | CHS | test | THC\_0688 |  |  | FL | 2.3.3.77.2 | Zinc-coordinating DNA-binding domains | C2H2 zinc finger factors | More than 3 adjacent zinc fingers | ZNF595-like | 17 | 14.99 | 42.12 | ZNF721.FL@CHS@foggy-rust-fly@autosome-ru.ChIPMunk@406seq\_7to21\_m1.pcm |
| ZNF721 | GHTS.GFPIVT | test | YWS\_B\_AffSeq\_A6\_ZNF721-DBD1 |  |  | DBD | 2.3.3.77.2 | Zinc-coordinating DNA-binding domains | C2H2 zinc finger factors | More than 3 adjacent zinc fingers | ZNF595-like | 20 | 7.76 | 48.81 | ZNF721.DBD@AFS.GFPIVT@stealthy-tangerine-birman@Halle.Dimont@Motif\_1\_w20\_astrained.ppm |
| ZNF721 | HTS.GFPIVT | test | YWS\_A\_AA40NGTGGTG |  |  | DBD | 2.3.3.77.2 | Zinc-coordinating DNA-binding domains | C2H2 zinc finger factors | More than 3 adjacent zinc fingers | ZNF595-like | 20 | 7.76 | 48.81 | ZNF721.DBD@AFS.GFPIVT@stealthy-tangerine-birman@Halle.Dimont@Motif\_1\_w20\_astrained.ppm |
| ZNF721 | HTS.GFPIVT | train | YWS\_A\_AG40NCTTTTC |  |  | DBD | 2.3.3.77.2 | Zinc-coordinating DNA-binding domains | C2H2 zinc finger factors | More than 3 adjacent zinc fingers | ZNF595-like | 17 | 12.9 | 46.77 | ZNF721.DBD@HTS.GFPIVT@crappy-wisteria-spoonbill@AJolma.Autoseed@Multinom2\_Onehit\_Seed\_VNAAARCAGBRTHNCCBackgroundCyc1.ppm |
| ZNF721 | HTS.GFPIVT | test | YWS\_A\_CA40NGCCTTT |  |  | FL | 2.3.3.77.2 | Zinc-coordinating DNA-binding domains | C2H2 zinc finger factors | More than 3 adjacent zinc fingers | ZNF595-like | 16 | 7.42 | 45.31 | ZNF721.FL@HTS.GFPIVT@gamy-malachite-orangutan+craggy-orange-yak+hasty-burgundy-turtle@Halle.Dimont@Motif\_1\_sampled\_e4\_astrained.ppm |
| FIZ1 | CHS | test | THC\_0758 |  |  | FL | 2.3.4.0.38 | Zinc-coordinating DNA-binding domains | C2H2 zinc finger factors | Multiple dispersed zinc fingers | Unclassified | 20 | 6.82 | 55.14 | FIZ1.FL@CHS@stinky-purple-kingfisher@Halle.Dimont@Motif\_1\_w20\_astrained.ppm |
| FIZ1 | CHS | test | THC\_0758 |  |  | FL | 2.3.4.0.38 | Zinc-coordinating DNA-binding domains | C2H2 zinc finger factors | Multiple dispersed zinc fingers | Unclassified | 20 | 6.7 | 56.82 | FIZ1.FL@CHS@thirsty-maroon-moorhen@Halle.Dimont@Motif\_1\_w20\_astrained.ppm |
| FIZ1 | GHTS.Lys | test | YWK\_B\_AffSeq\_G8\_FIZ1 |  |  | FL | 2.3.4.0.38 | Zinc-coordinating DNA-binding domains | C2H2 zinc finger factors | Multiple dispersed zinc fingers | Unclassified | 19 | 18.86 | 49.78 | FIZ1.FL@HTS.Lys@dorky-xanthic-flounder+muggy-malachite-blue@HughesLab.Streme@topk\_cycle=C3+C4\_k=5\_top=10000\_Motif1\_min3max30.ppm |
| FIZ1 | HTS.Lys | test | YWK\_A\_TC40NGCGATT |  |  | NA | 2.3.4.0.38 | Zinc-coordinating DNA-binding domains | C2H2 zinc finger factors | Multiple dispersed zinc fingers | Unclassified | 15 | 15.38 | 48.45 | FIZ1.NA@SMS@boozy-sangria-caiman@ |
| FIZ1 | HTS.Lys | train | YWK\_C\_TC40NGCGATT |  |  | NA | 2.3.4.0.38 | Zinc-coordinating DNA-binding domains | C2H2 zinc finger factors | Multiple dispersed zinc fingers | Unclassified | 15 | 15.38 | 48.45 | FIZ1.NA@SMS@boozy-sangria-caiman@ |
| FIZ1 | SMS | train | UT380-070-2 |  |  | NA | 2.3.4.0.38 | Zinc-coordinating DNA-binding domains | C2H2 zinc finger factors | Multiple dispersed zinc fingers | Unclassified | 17 | 12.11 | 47.82 | FIZ1.NA@SMS@boozy-sangria-caiman@autosome-ru.ChIPMunk@topk\_k=5\_top=10000.pcm |
| ZFTA | CHS | test | THC\_0197 |  |  | FL | 2.3.5.0.257 | Zinc-coordinating DNA-binding domains | C2H2 zinc finger factors | BED zinc finger |  | 11 | 14.34 | 76.4 | ZFTA.FL@CHS@foggy-red-dalmatian@autosome-ru.ChIPMunk@500seq\_15to7\_m0.pcm |
| ZFTA | GHTS.GFPIVT | test | YWM\_B\_AffSeq\_A4\_C11orf95-FL |  |  | FL | 2.3.5.0.257 | Zinc-coordinating DNA-binding domains | C2H2 zinc finger factors | BED zinc finger |  | 20 | 8.33 | 64.03 | ZFTA.FL@AFS.GFPIVT@squeaky-bronze-ostrich@Halle.Dimont@Motif\_1\_w20\_astrained.ppm |
| ZFTA | PBM | train | PBM13825 |  |  | FL | 2.3.5.0.257 | Zinc-coordinating DNA-binding domains | C2H2 zinc finger factors | BED zinc finger |  | 11 | 13.53 | 74.41 | ZFTA.FL@AFS.Lys@cloudy-salmon-macaque@ |
| ZFTA | PBM | test | PBM13841 |  |  | FL | 2.3.5.0.257 | Zinc-coordinating DNA-binding domains | C2H2 zinc finger factors | BED zinc finger |  | 11 | 13.53 | 74.41 | ZFTA.FL@AFS.Lys@cloudy-salmon-macaque@ |
| ZFTA | HTS.GFPIVT | train | YWM\_A\_AA40NGTGGTC |  |  | FL | 2.3.5.0.257 | Zinc-coordinating DNA-binding domains | C2H2 zinc finger factors | BED zinc finger |  | 12 | 1.5 | 53.13 | ZFTA.FL@HTS.GFPIVT@frumpy-myrtle-stingray+skanky-cyan-sparrow+smelly-champagne-bongo+messy-burgundy-guppy@faltejsk.ProBound@motif\_without\_ns.ppm |
| ZFTA | SMS | train | UT380-025 |  |  | FL | 2.3.5.0.257 | Zinc-coordinating DNA-binding domains | C2H2 zinc finger factors | BED zinc finger |  | 12 | 15.31 | 69.7 | ZFTA.FL@HTS.GFPIVT@frumpy-myrtle-stingray+messy-burgundy-guppy+skanky-cyan-sparrow+smelly-champagne-bongo@HughesLab.MEME@topk\_cycle=C1+C2+C3+C4\_k=5\_top=10000\_Motif1\_min3max30.ppm |
| TPRX1 | CHS | test | THC\_0336.Rep-MICHELLE\_0314 |  |  | DBD | 3.1.3.26.1 | Helix-turn-helix domains | Homeo domain factors | Paired-related HD | TPRX | 12 | 3.01 | 42.06 | TPRX1.DBD@HTS.GFPIVT@crabby-celadon-bonobo+skinny-cardinal-oriole+chummy-charcoal-urchin+skinny-purple-chipmunk@faltejsk.ProBound@motif\_without\_ns.ppm |
| TPRX1 | GHTS.Lys | test | YWL\_B\_AffSeq\_D4\_TPRX1 |  |  | FL | 3.1.3.26.1 | Helix-turn-helix domains | Homeo domain factors | Paired-related HD | TPRX | 20 | 7.82 | 49.19 | TPRX1.FL@AFS.Lys@flimsy-magnolia-eel+homey-cyan-ladybird+shaky-violet-capuchin@Halle.Dimont@Motif\_1\_w20\_astrained.ppm |
| TPRX1 | GHTS.Lys | test | YWL\_B\_AffSeq\_D4\_TPRX1 |  |  | FL | 3.1.3.26.1 | Helix-turn-helix domains | Homeo domain factors | Paired-related HD | TPRX | 20 | 8.08 | 50.19 | TPRX1.FL@AFS.Lys@shaky-violet-capuchin@Halle.Dimont@Motif\_1\_w20\_astrained.ppm |
| TPRX1 | PBM | test | PBM13675 |  |  | DBD | 3.1.3.26.1 | Helix-turn-helix domains | Homeo domain factors | Paired-related HD | TPRX | 11 | 3.36 | 40.17 | TPRX1.DBD@PBM.HK@smelly-cardinal-toucan@Halle.Dimont@Motif\_1\_w0\_01\_bg4\_imw15\_astrained.ppm |
| TPRX1 | PBM | test | PBM13675 |  |  | DBD | 3.1.3.26.1 | Helix-turn-helix domains | Homeo domain factors | Paired-related HD | TPRX | 6 | 5.34 | 30.6 | TPRX1.DBD@PBM.HK@smelly-cardinal-toucan@Halle.Dimont@Motif\_1\_astrained.ppm |
| TPRX1 | PBM | train | PBM13659 |  |  | DBD | 3.1.3.26.1 | Helix-turn-helix domains | Homeo domain factors | Paired-related HD | TPRX | 11 | 3.36 | 40.17 | TPRX1.DBD@PBM.HK@smelly-cardinal-toucan@Halle.Dimont@Motif\_1\_w0\_01\_bg4\_imw15\_astrained.ppm |
| TPRX1 | PBM | train | PBM13659 |  |  | DBD | 3.1.3.26.1 | Helix-turn-helix domains | Homeo domain factors | Paired-related HD | TPRX | 15 | 12.77 | 36.62 | TPRX1.DBD@HTS.IVT@flaky-orchid-paradise+snippy-periwinkle-grouse@autosome-ru.ChIPMunk@topk\_cycle=C2+C3\_k=5\_top=10000.pcm |
| TPRX1 | PBM | test | PBM13944 |  |  | FL | 3.1.3.26.1 | Helix-turn-helix domains | Homeo domain factors | Paired-related HD | TPRX | 9 | 12.55 | 46.76 | TPRX1.FL@HTS.Lys@pokey-linen-spider+tacky-crimson-caiman@HughesLab.Streme@topk\_cycle=C3+C4\_k=5\_top=500\_Motif1.ppm |
| TPRX1 | HTS.IVT | test | YWC\_B\_TG40NGACCTT |  |  | FL | 3.1.3.26.1 | Helix-turn-helix domains | Homeo domain factors | Paired-related HD | TPRX | 12 | 2.92 | 43.33 | TPRX1.FL@HTS.Lys@pokey-linen-spider+hilly-brown-woodlouse+thirsty-harlequin-pinscher+tacky-crimson-caiman@faltejsk.ProBound@motif\_without\_ns.ppm |
| TPRX1 | HTS.GFPIVT | test | YWP\_A\_AG40NTGGCTA |  |  | FL | 3.1.3.26.1 | Helix-turn-helix domains | Homeo domain factors | Paired-related HD | TPRX | 12 | 2.92 | 43.33 | TPRX1.FL@HTS.Lys@pokey-linen-spider+hilly-brown-woodlouse+thirsty-harlequin-pinscher+tacky-crimson-caiman@faltejsk.ProBound@motif\_without\_ns.ppm |
| TPRX1 | HTS.GFPIVT | train | YWM\_A\_TC40NACGGGA |  |  | FL | 3.1.3.26.1 | Helix-turn-helix domains | Homeo domain factors | Paired-related HD | TPRX | 12 | 2.92 | 43.33 | TPRX1.FL@HTS.Lys@pokey-linen-spider+hilly-brown-woodlouse+thirsty-harlequin-pinscher+tacky-crimson-caiman@faltejsk.ProBound@motif\_without\_ns.ppm |
| TPRX1 | HTS.Lys | train | YWL\_A\_CT40NCGGAGA |  |  | FL | 3.1.3.26.1 | Helix-turn-helix domains | Homeo domain factors | Paired-related HD | TPRX | 12 | 2.92 | 43.33 | TPRX1.FL@HTS.Lys@pokey-linen-spider+hilly-brown-woodlouse+thirsty-harlequin-pinscher+tacky-crimson-caiman@faltejsk.ProBound@motif\_without\_ns.ppm |
| TPRX1 | SMS | train | UT380-233-2 |  |  | FL | 3.1.3.26.1 | Helix-turn-helix domains | Homeo domain factors | Paired-related HD | TPRX | 12 | 2.92 | 43.33 | TPRX1.FL@HTS.Lys@pokey-linen-spider+hilly-brown-woodlouse+thirsty-harlequin-pinscher+tacky-crimson-caiman@faltejsk.ProBound@motif\_without\_ns.ppm |
| TPRX1 | SMS | test | UT380-233-1 |  |  | FL | 3.1.3.26.1 | Helix-turn-helix domains | Homeo domain factors | Paired-related HD | TPRX | 12 | 2.92 | 43.33 | TPRX1.FL@HTS.Lys@pokey-linen-spider+hilly-brown-woodlouse+thirsty-harlequin-pinscher+tacky-crimson-caiman@faltejsk.ProBound@motif\_without\_ns.ppm |
| MKX | CHS | test | THC\_0462.Rep-MICHELLE\_0314 |  |  | FL | 3.1.4.3.1 | Helix-turn-helix domains | Homeo domain factors | TALE-type HD | MKX | 10 | 10.11 | 49.36 | MKX.FL@CHS@droopy-tomato-affenpinscher@ |
| MKX | GHTS.Lys | test | YWK\_B\_AffSeq\_C7\_MKX |  |  | FL | 3.1.4.3.1 | Helix-turn-helix domains | Homeo domain factors | TALE-type HD | MKX | 16 | 4.76 | 43.63 | MKX.FL@AFS.Lys@randy-denim-camel@Halle.Dimont@Motif\_1\_imw20\_astrained.ppm |
| MKX | PBM | test | PBM13673 |  |  | FL | 3.1.4.3.1 | Helix-turn-helix domains | Homeo domain factors | TALE-type HD | MKX | 12 | 1.43 | 46.43 | MKX.FL@HTS.Lys@wiggy-flax-gorilla+lanky-white-tarantula+messy-green-mongoose+thirsty-cerulean-bandicoot@faltejsk.ProBound@motif\_without\_ns.ppm |
| MKX | PBM | train | PBM13657 |  |  | DBD | 3.1.4.3.1 | Helix-turn-helix domains | Homeo domain factors | TALE-type HD | MKX | 9 | 3.75 | 33.74 | MKX.DBD@PBM.ME@sickly-denim-capybara@Halle.Dimont@Motif\_1\_astrained.ppm |
| MKX | PBM | train | PBM13657 |  |  | DBD | 3.1.4.3.1 | Helix-turn-helix domains | Homeo domain factors | TALE-type HD | MKX | 10 | 3.17 | 35.59 | MKX.DBD@PBM.ME@cloudy-cinnamon-quoll@Halle.Dimont@Motif\_1\_astrained.ppm |
| MKX | HTS.IVT | train | YWC\_B\_GT40NGGCATT |  |  | FL | 3.1.4.3.1 | Helix-turn-helix domains | Homeo domain factors | TALE-type HD | MKX | 12 | 2.1 | 49.18 | MKX.FL@AFS.Lys@scaly-sangria-buffalo+bumpy-periwinkle-mouse+hasty-crimson-chameleon+slaphappy-turquoise-fowl@faltejsk.ProBound@motif\_without\_ns.ppm |
| MKX | HTS.IVT | test | YWF\_A\_CC40NTGAGTA |  |  | FL | 3.1.4.3.1 | Helix-turn-helix domains | Homeo domain factors | TALE-type HD | MKX | 12 | 2.1 | 49.18 | MKX.FL@AFS.Lys@scaly-sangria-buffalo+bumpy-periwinkle-mouse+hasty-crimson-chameleon+slaphappy-turquoise-fowl@faltejsk.ProBound@motif\_without\_ns.ppm |
| MKX | HTS.GFPIVT | test | YWP\_A\_TA40NTCACTC |  |  | FL | 3.1.4.3.1 | Helix-turn-helix domains | Homeo domain factors | TALE-type HD | MKX | 12 | 3.98 | 39.96 | MKX.FL@HTS.Lys@wiggy-flax-gorilla+lanky-white-tarantula+messy-green-mongoose+thirsty-cerulean-bandicoot@faltejsk.ProBound@motif\_with\_ns.ppm |
| MKX | HTS.Lys | train | YWK\_A\_CC40NTGAGTA |  |  | FL | 3.1.4.3.1 | Helix-turn-helix domains | Homeo domain factors | TALE-type HD | MKX | 10 | 5.09 | 42.42 | MKX.FL@HTS.Lys@wiggy-flax-gorilla+lanky-white-tarantula+messy-green-mongoose+thirsty-cerulean-bandicoot@Halle.Dimont@Motif\_1\_sampled\_e4\_astrained.ppm |
| MKX | HTS.Lys | test | YWK\_C\_CC40NTGAGTA |  |  | FL | 3.1.4.3.1 | Helix-turn-helix domains | Homeo domain factors | TALE-type HD | MKX | 10 | 5.09 | 42.42 | MKX.FL@HTS.Lys@wiggy-flax-gorilla+lanky-white-tarantula+messy-green-mongoose+thirsty-cerulean-bandicoot@Halle.Dimont@Motif\_1\_sampled\_e4\_astrained.ppm |
| MKX | SMS | train | UT380-113 |  |  | FL | 3.1.4.3.1 | Helix-turn-helix domains | Homeo domain factors | TALE-type HD | MKX | 12 | 2.1 | 49.18 | MKX.FL@AFS.Lys@scaly-sangria-buffalo+bumpy-periwinkle-mouse+hasty-crimson-chameleon+slaphappy-turquoise-fowl@faltejsk.ProBound@motif\_without\_ns.ppm |
| MSANTD1 | CHS | test | THC\_0610 |  |  | FL | 3.5.1.0.1 | Helix-turn-helix domains | Tryptophan cluster factors | Myb/SANT domain | Other Myb-like | 20 | 4.8 | 48.52 | MSANTD1.FL@CHS@crappy-tomato-monster@Halle.Dimont@Motif\_1\_w20\_astrained.ppm |
| MSANTD1 | GHTS.Lys | test | YWL\_B\_AffSeq\_H6\_MSANTD1 |  |  | FL | 3.5.1.0.1 | Helix-turn-helix domains | Tryptophan cluster factors | Myb/SANT domain | Other Myb-like | 15 | 15.71 | 36.25 | MSANTD1.FL@AFS.Lys@shabby-brown-dugong@autosome-ru.ChIPMunk@979seq\_21to7\_m0.pcm |
| MSANTD1 | GHTS.Lys | test | YWL\_B\_AffSeq\_H6\_MSANTD1 |  |  | FL | 3.5.1.0.1 | Helix-turn-helix domains | Tryptophan cluster factors | Myb/SANT domain | Other Myb-like | 15 | 16.72 | 38.19 | MSANTD1.FL@AFS.Lys@smelly-champagne-russel@autosome-ru.ChIPMunk@103seq\_7to15\_m0.pcm |
| MSANTD1 | PBM | test | PBM13677 |  |  | FL | 3.5.1.0.1 | Helix-turn-helix domains | Tryptophan cluster factors | Myb/SANT domain | Other Myb-like | 22 | 16.01 | 36.8 | MSANTD1.FL@AFS.Lys@shabby-brown-dugong@HughesLab.Homer@500\_fa\_homer\_minw3\_maxw\_40\_Motif1.ppm |
| MSANTD1 | PBM | test | PBM13677 |  |  | FL | 3.5.1.0.1 | Helix-turn-helix domains | Tryptophan cluster factors | Myb/SANT domain | Other Myb-like | 21 | 15.64 | 35.66 | MSANTD1.FL@AFS.Lys@shabby-brown-dugong@HughesLab.Homer@1000\_fa\_homer\_minw3\_maxw\_40\_Motif1.ppm |
| MSANTD1 | PBM | train | PBM13661 |  |  | FL | 3.5.1.0.1 | Helix-turn-helix domains | Tryptophan cluster factors | Myb/SANT domain | Other Myb-like | 16 | 15.41 | 38.99 | MSANTD1.FL@AFS.Lys@shabby-brown-dugong@HughesLab.Streme@1000\_fa\_streme\_minw3\_maxw30\_Motif1.ppm |
| MSANTD1 | HTS.IVT | train | YWC\_B\_AA40NGTGGTG |  |  | FL | 3.5.1.0.1 | Helix-turn-helix domains | Tryptophan cluster factors | Myb/SANT domain | Other Myb-like | 20 | 8.68 | 45.39 | MSANTD1.FL@AFS.Lys@shabby-brown-dugong@Halle.Dimont@Motif\_1\_w20\_astrained.ppm |
| MSANTD1 | HTS.GFPIVT | test | YWQ\_A\_GT40NGAAGTA |  |  | FL | 3.5.1.0.1 | Helix-turn-helix domains | Tryptophan cluster factors | Myb/SANT domain | Other Myb-like | 14 | 18.29 | 42.0 | MSANTD1.FL@HTS.Lys@hilly-ruby-kingfisher+ugly-ruby-olm@HughesLab.MEME@topk\_cycle=C3+C4\_k=5\_top=1000\_Motif1.ppm |
| MSANTD1 | HTS.Lys | test | YWL\_A\_TG40NTGTGTT |  |  | FL | 3.5.1.0.1 | Helix-turn-helix domains | Tryptophan cluster factors | Myb/SANT domain | Other Myb-like | 12 | 1.77 | 47.48 | MSANTD1.FL@HTS.Lys@hilly-ruby-kingfisher+slimy-purple-opossum+ugly-ruby-olm+cheeky-champagne-alligator@faltejsk.ProBound@motif\_without\_ns.ppm |
| MSANTD1 | SMS | train | UT380-114 |  |  | FL | 3.5.1.0.1 | Helix-turn-helix domains | Tryptophan cluster factors | Myb/SANT domain | Other Myb-like | 20 | 8.68 | 45.39 | MSANTD1.FL@AFS.Lys@shabby-brown-dugong@Halle.Dimont@Motif\_1\_w20\_astrained.ppm |
| MYPOP | CHS | test | THC\_0710 |  |  | DBD | 3.5.1.0.5 | Helix-turn-helix domains | Tryptophan cluster factors | Myb/SANT domain | Other Myb-like | 7 | 3.53 | 60.11 | MYPOP.DBD@PBM.ME@stuffy-myrtle-walrus@Halle.Dimont@Motif\_1\_w0\_01\_bg4\_imw15\_astrained.ppm |
| MYPOP | GHTS.Lys | test | YWO\_B\_AffSeq\_C7\_MYPOP-FL |  |  | FL | 3.5.1.0.5 | Helix-turn-helix domains | Tryptophan cluster factors | Myb/SANT domain | Other Myb-like | 20 | 7.59 | 56.43 | MYPOP.FL@AFS.Lys@scanty-cornflower-gecko+craggy-orchid-tarantula+sleepy-aquamarine-urchin+muggy-emerald-booby@Halle.Dimont@Motif\_1\_w20\_astrained.ppm |
| MYPOP | GHTS.Lys | test | YWO\_B\_AffSeq\_C7\_MYPOP-FL |  |  | FL | 3.5.1.0.5 | Helix-turn-helix domains | Tryptophan cluster factors | Myb/SANT domain | Other Myb-like | 7 | 5.84 | 58.56 | MYPOP.FL@AFS.Lys@bumpy-buff-clam@Halle.Dimont@Motif\_1\_imw20\_astrained.ppm |
| MYPOP | PBM | test | PBM13680 |  |  | FL | 3.5.1.0.5 | Helix-turn-helix domains | Tryptophan cluster factors | Myb/SANT domain | Other Myb-like | 12 | 6.72 | 54.1 | MYPOP.FL@HTS.GFPIVT@wimpy-auburn-frigatebird+sickly-ecru-tamarin+bumpy-red-terrier@faltejsk.ProBound@motif\_with\_ns.ppm |
| MYPOP | PBM | train | PBM13664 |  |  | FL | 3.5.1.0.5 | Helix-turn-helix domains | Tryptophan cluster factors | Myb/SANT domain | Other Myb-like | 12 | 6.72 | 54.1 | MYPOP.FL@HTS.GFPIVT@wimpy-auburn-frigatebird+sickly-ecru-tamarin+bumpy-red-terrier@faltejsk.ProBound@motif\_with\_ns.ppm |
| MYPOP | HTS.IVT | train | YWF\_A\_TC40NTGCATA |  |  | FL | 3.5.1.0.5 | Helix-turn-helix domains | Tryptophan cluster factors | Myb/SANT domain | Other Myb-like | 12 | 2.99 | 53.71 | MYPOP.FL@HTS.GFPIVT@wimpy-auburn-frigatebird+sickly-ecru-tamarin+bumpy-red-terrier@faltejsk.ProBound@motif\_without\_ns.ppm |
| MYPOP | HTS.IVT | test | YWC\_B\_GC40NTATAGG |  |  | FL | 3.5.1.0.5 | Helix-turn-helix domains | Tryptophan cluster factors | Myb/SANT domain | Other Myb-like | 12 | 2.99 | 53.71 | MYPOP.FL@HTS.GFPIVT@wimpy-auburn-frigatebird+sickly-ecru-tamarin+bumpy-red-terrier@faltejsk.ProBound@motif\_without\_ns.ppm |
| MYPOP | HTS.GFPIVT | test | YWP\_A\_GC40NCACTTC |  |  | FL | 3.5.1.0.5 | Helix-turn-helix domains | Tryptophan cluster factors | Myb/SANT domain | Other Myb-like | 12 | 2.12 | 52.03 | MYPOP.FL@HTS.Lys@wimpy-sangria-stingray+ready-eggplant-crane+frumpy-lilac-numbat+sickly-gamboge-affenpinscher@faltejsk.ProBound@motif\_without\_ns.ppm |
| MYPOP | HTS.Lys | train | YWN\_A\_CC40NTGAGTA |  |  | FL | 3.5.1.0.5 | Helix-turn-helix domains | Tryptophan cluster factors | Myb/SANT domain | Other Myb-like | 12 | 2.12 | 52.03 | MYPOP.FL@HTS.Lys@wimpy-sangria-stingray+ready-eggplant-crane+frumpy-lilac-numbat+sickly-gamboge-affenpinscher@faltejsk.ProBound@motif\_without\_ns.ppm |
| MYPOP | HTS.Lys | test | YWO\_A\_CC40NTGAGTA |  |  | FL | 3.5.1.0.5 | Helix-turn-helix domains | Tryptophan cluster factors | Myb/SANT domain | Other Myb-like | 12 | 2.12 | 52.03 | MYPOP.FL@HTS.Lys@wimpy-sangria-stingray+ready-eggplant-crane+frumpy-lilac-numbat+sickly-gamboge-affenpinscher@faltejsk.ProBound@motif\_without\_ns.ppm |
| SP140L | CHS | test | THC\_0072 |  |  | FL | 5.3.5.1.2 | alpha-Helices exposed by beta-structures | SAND domain factors | Sp140-Sp100 | Sp140 | 20 | 5.22 | 49.04 | SP140L.FL@CHS@queasy-pumpkin-gopher@Halle.Dimont@Motif\_1\_w20\_astrained.ppm |
| SP140L | GHTS.GFPIVT | test | YWP\_B\_AffSeq\_F6\_SP140L-DBD |  |  | DBD | 5.3.5.1.2 | alpha-Helices exposed by beta-structures | SAND domain factors | Sp140-Sp100 | Sp140 | 20 | 6.81 | 56.55 | SP140L.DBD@AFS.GFPIVT@sickly-coral-eel@Halle.Dimont@Motif\_1\_w20\_astrained.ppm |
| SP140L | PBM | train | PBM13763 |  |  | NA | 5.3.5.1.2 | alpha-Helices exposed by beta-structures | SAND domain factors | Sp140-Sp100 | Sp140 | 12 | 3.22 | 49.48 | SP140L.NA@SMS@hazy-plum-akbash@faltejsk.ProBound@motif\_with\_ns.ppm |
| SP140L | PBM | test | PBM13779 |  |  | FL | 5.3.5.1.2 | alpha-Helices exposed by beta-structures | SAND domain factors | Sp140-Sp100 | Sp140 | 12 | 2.18 | 47.78 | SP140L.FL@HTS.GFPIVT@hilly-white-wolverine+leaky-brass-bobcat+crappy-gamboge-buffalo@faltejsk.ProBound@motif\_without\_ns.ppm |
| SP140L | HTS.IVT | test | YWC\_B\_GT40NTGTGGA |  |  | DBD | 5.3.5.1.2 | alpha-Helices exposed by beta-structures | SAND domain factors | Sp140-Sp100 | Sp140 | 8 | 4.38 | 53.84 | SP140L.DBD@HTS.IVT@wiggy-blue-lemming+sleazy-wheat-maltese+sickly-fuchsia-guppy@Halle.Dimont@Motif\_1\_sampled\_e1\_5\_astrained.ppm |
| SP140L | HTS.IVT | train | YWJ\_A\_CC40NCCAATA |  |  | DBD | 5.3.5.1.2 | alpha-Helices exposed by beta-structures | SAND domain factors | Sp140-Sp100 | Sp140 | 8 | 4.38 | 53.84 | SP140L.DBD@HTS.IVT@wiggy-blue-lemming+sleazy-wheat-maltese+sickly-fuchsia-guppy@Halle.Dimont@Motif\_1\_sampled\_e1\_5\_astrained.ppm |
| SP140L | HTS.GFPIVT | test | YWP\_A\_TG40NGACCTT |  |  | NA | 5.3.5.1.2 | alpha-Helices exposed by beta-structures | SAND domain factors | Sp140-Sp100 | Sp140 | 12 | 2.77 | 49.53 | SP140L.NA@SMS@hazy-plum-akbash@faltejsk.ProBound@motif\_without\_ns.ppm |
| SP140L | HTS.GFPIVT | train | YWP\_A\_GT40NGTTCTC |  |  | NA | 5.3.5.1.2 | alpha-Helices exposed by beta-structures | SAND domain factors | Sp140-Sp100 | Sp140 | 12 | 2.77 | 49.53 | SP140L.NA@SMS@hazy-plum-akbash@faltejsk.ProBound@motif\_without\_ns.ppm |
| SP140L | SMS | train | UT380-204-2 |  |  | NA | 5.3.5.1.2 | alpha-Helices exposed by beta-structures | SAND domain factors | Sp140-Sp100 | Sp140 | 12 | 2.77 | 49.53 | SP140L.NA@SMS@hazy-plum-akbash@faltejsk.ProBound@motif\_without\_ns.ppm |
| GCM1 | CHS | test | THC\_0621 |  |  | NA | 7.2.1.0.1 | beta-Hairpin exposed by an alpha/beta-scaffold | GCM domain factors | GCM |  | 13 | 5.87 | 62.6 | GCM1.NA@CHS@geeky-zucchini-shrew@Halle.Dimont@Motif\_1\_imw20\_astrained.ppm |
| GCM1 | GHTS.IVT | test | YWF\_B\_AffSeq\_F12\_GCM1 |  |  | NA | 7.2.1.0.1 | beta-Hairpin exposed by an alpha/beta-scaffold | GCM domain factors | GCM |  | 11 | 13.18 | 66.72 | GCM1.NA@AFS.IVT@skanky-ecru-chicken@HughesLab.Streme@1000\_fa\_streme\_Motif1.ppm |
| GCM1 | PBM | test | PBM14359 |  |  | NA | 7.2.1.0.1 | beta-Hairpin exposed by an alpha/beta-scaffold | GCM domain factors | GCM |  | 10 | 8.97 | 60.87 | GCM1.NA@PBM.ME@tacky-zucchini-pinscher@autosome-ru.ChIPMunk@s\_6-16\_flat.pcm |
| GCM1 | PBM | train | PBM14343 |  |  | NA | 7.2.1.0.1 | beta-Hairpin exposed by an alpha/beta-scaffold | GCM domain factors | GCM |  | 10 | 9.16 | 59.67 | GCM1.NA@PBM.HK@woolly-purple-shark@autosome-ru.ChIPMunk@s\_6-16\_flat.pcm |
| GCM1 | HTS.IVT | test | YWF\_A\_TA40NGTTAGC |  |  | NA | 7.2.1.0.1 | beta-Hairpin exposed by an alpha/beta-scaffold | GCM domain factors | GCM |  | 12 | 6.36 | 57.09 | GCM1.NA@HTS.IVT@cranky-salmon-butterfly+grumpy-pear-affenpinscher+zippy-orchid-gharial+baggy-gamboge-whippet@Halle.Dimont@Motif\_1\_sampled\_e4\_astrained.ppm |
| GCM1 | HTS.Lys | train | YWK\_A\_TC40NTAAGTG |  |  | NA | 7.2.1.0.1 | beta-Hairpin exposed by an alpha/beta-scaffold | GCM domain factors | GCM |  | 10 | 9.16 | 59.67 | GCM1.NA@PBM.HK@woolly-purple-shark@autosome-ru.ChIPMunk@s\_6-16\_flat.pcm |
| GCM1 | HTS.Lys | test | YWK\_C\_TC40NTAAGTG |  |  | NA | 7.2.1.0.1 | beta-Hairpin exposed by an alpha/beta-scaffold | GCM domain factors | GCM |  | 10 | 9.16 | 59.67 | GCM1.NA@PBM.HK@woolly-purple-shark@autosome-ru.ChIPMunk@s\_6-16\_flat.pcm |
| GCM1 | SMS | train | SRR3405056 |  |  | NA | 7.2.1.0.1 | beta-Hairpin exposed by an alpha/beta-scaffold | GCM domain factors | GCM |  | 15 | 5.89 | 55.38 | GCM1.NA@HTS.IVT@cranky-salmon-butterfly+grumpy-pear-affenpinscher+zippy-orchid-gharial+baggy-gamboge-whippet@Halle.Dimont@Motif\_1\_sampled\_e1\_5\_astrained.ppm |
