## Supplementary Data SD1 for "Inferring binding specificities of human transcription factors with the wisdom of crowds": ibis_motifs_Final_G2A.html

 

| TF | Platform | Slice | Replicate ID | logo-direct | logo-revcomp | Construct type | tfclass:id | tfclass:superclass | tfclass:class | tfclass:family | tfclass:subfamily | Motif length | Information content | GC% | Motif |
| --- | --- | --- | --- | --- | --- | --- | --- | --- | --- | --- | --- | --- | --- | --- | --- |
| CAMTA1 | CHS | train | THC\_0249 |  |  | FL | 0.6.1.0.1 | Yet undefined DNA-binding domains | CG-1 domain factors | CAMTA |  | 12 | 11.18 | 70.02 | CAMTA1.FL@CHS@grumpy-emerald-tiger@ |
| CAMTA1 | GHTS.GFPIVT | test | YWM\_B\_AffSeq\_B4\_CAMTA1-DBD |  |  | DBD | 0.6.1.0.1 | Yet undefined DNA-binding domains | CG-1 domain factors | CAMTA |  | 20 | 7.38 | 57.34 | CAMTA1.DBD@AFS.GFPIVT@flaky-xanthic-nightingale+zippy-violet-tarantula+sleepy-red-quoll+surly-jade-warthog@Halle.Dimont@Motif\_1\_w20\_astrained.ppm |
| CAMTA1 | GHTS.Lys | train | YWK\_D\_AffSeq\_G3\_CAMTA1 |  |  | FL | 0.6.1.0.1 | Yet undefined DNA-binding domains | CG-1 domain factors | CAMTA |  | 20 | 7.75 | 58.88 | CAMTA1.FL@AFS.Lys@stuffy-sangria-manatee@Halle.Dimont@Motif\_1\_w20\_astrained.ppm |
| CAMTA1 | GHTS.Lys | train | YWK\_B\_AffSeq\_G3\_CAMTA1 |  |  | FL | 0.6.1.0.1 | Yet undefined DNA-binding domains | CG-1 domain factors | CAMTA |  | 20 | 7.75 | 58.88 | CAMTA1.FL@AFS.Lys@stuffy-sangria-manatee@Halle.Dimont@Motif\_1\_w20\_astrained.ppm |
| CAMTA1 | HTS.GFPIVT | test | YWM\_A\_AG40NTGGCTA |  |  | DBD | 0.6.1.0.1 | Yet undefined DNA-binding domains | CG-1 domain factors | CAMTA |  | 15 | 5.26 | 53.81 | CAMTA1.DBD@AFS.GFPIVT@zippy-violet-tarantula@Halle.Dimont@Motif\_1\_imw20\_astrained.ppm |
| CAMTA1 | HTS.Lys | test | YWK\_A\_TA40NAGGCTT |  |  | FL | 0.6.1.0.1 | Yet undefined DNA-binding domains | CG-1 domain factors | CAMTA |  | 12 | 3.65 | 52.93 | CAMTA1.FL@HTS.Lys@thirsty-seashell-drever+gimpy-bistre-dalmatian+crappy-cobalt-molly+cheeky-thistle-woodpecker@faltejsk.ProBound@motif\_with\_ns.ppm |
| CAMTA1 | HTS.Lys | test | YWK\_C\_TA40NAGGCTT |  |  | FL | 0.6.1.0.1 | Yet undefined DNA-binding domains | CG-1 domain factors | CAMTA |  | 11 | 4.93 | 54.02 | CAMTA1.FL@AFS.Lys@foggy-smalt-cat@Halle.Dimont@Motif\_1\_imw20\_astrained.ppm |
| CAMTA1 | SMS | test | UT380-026 |  |  | DBD | 0.6.1.0.1 | Yet undefined DNA-binding domains | CG-1 domain factors | CAMTA |  | 15 | 5.23 | 54.11 | CAMTA1.DBD@AFS.GFPIVT@sleepy-red-quoll@Halle.Dimont@Motif\_1\_imw20\_astrained.ppm |
| MYF6 | CHS | train | THC\_0721 |  |  | NA | 1.2.2.1.4 | Basic domains | Basic helix-loop-helix factors (bHLH) | MyoD-ASC-related | Myogenic TFs | 11 | 11.89 | 65.43 | MYF6.NA@CHS@sloppy-lavender-tarantula@HughesLab.MEME@1000\_fa\_memechip\_Motif1.ppm |
| MYF6 | CHS | test | THC\_0817 |  |  | NA | 1.2.2.1.4 | Basic domains | Basic helix-loop-helix factors (bHLH) | MyoD-ASC-related | Myogenic TFs | 11 | 11.89 | 65.43 | MYF6.NA@CHS@sloppy-lavender-tarantula@HughesLab.MEME@1000\_fa\_memechip\_Motif1.ppm |
| MYF6 | GHTS.Lys | train | YWK\_D\_AffSeq\_B12\_MYF6 |  |  | NA | 1.2.2.1.4 | Basic domains | Basic helix-loop-helix factors (bHLH) | MyoD-ASC-related | Myogenic TFs | 20 | 8.08 | 55.16 | MYF6.NA@AFS.Lys@sleazy-purple-millipede@Halle.Dimont@Motif\_1\_w20\_astrained.ppm |
| MYF6 | GHTS.Lys | test | YWK\_B\_AffSeq\_B12\_MYF6 |  |  | NA | 1.2.2.1.4 | Basic domains | Basic helix-loop-helix factors (bHLH) | MyoD-ASC-related | Myogenic TFs | 20 | 8.51 | 53.25 | MYF6.NA@AFS.Lys@homey-myrtle-wolf@Halle.Dimont@Motif\_1\_w20\_astrained.ppm |
| MYF6 | PBM | test | PBM14377 |  |  | NA | 1.2.2.1.4 | Basic domains | Basic helix-loop-helix factors (bHLH) | MyoD-ASC-related | Myogenic TFs | 11 | 9.04 | 54.82 | MYF6.NA@PBM.ME@cranky-persimmon-wrasse@autosome-ru.ChIPMunk@s\_6-16\_flat.pcm |
| MYF6 | HTS.IVT | test | YWF\_A\_GC40NCACTTC |  |  | NA | 1.2.2.1.4 | Basic domains | Basic helix-loop-helix factors (bHLH) | MyoD-ASC-related | Myogenic TFs | 12 | 7.34 | 51.33 | MYF6.NA@HTS.IVT@snoopy-goldenrod-uguisu+cranky-vermilion-chow+freaky-charcoal-beaver+leaky-rust-echidna@Halle.Dimont@Motif\_1\_sampled\_e1\_5\_astrained.ppm |
| MYF6 | HTS.Lys | test | YWK\_C\_AT40NCCTTGG |  |  | NA | 1.2.2.1.4 | Basic domains | Basic helix-loop-helix factors (bHLH) | MyoD-ASC-related | Myogenic TFs | 12 | 6.53 | 51.86 | MYF6.NA@HTS.Lys@craggy-russet-rattlesnake+homey-amaranth-macaque+stuffy-xanthic-rat+homely-vermilion-deer@faltejsk.ProBound@motif\_with\_ns.ppm |
| MYF6 | HTS.Lys | test | YWK\_A\_AT40NCCTTGG |  |  | NA | 1.2.2.1.4 | Basic domains | Basic helix-loop-helix factors (bHLH) | MyoD-ASC-related | Myogenic TFs | 12 | 6.53 | 51.86 | MYF6.NA@HTS.Lys@craggy-russet-rattlesnake+homey-amaranth-macaque+stuffy-xanthic-rat+homely-vermilion-deer@faltejsk.ProBound@motif\_with\_ns.ppm |
| USF3 | CHS | train | THC\_0757 |  |  | FL | 1.2.6.2.3 | Basic domains | Basic helix-loop-helix factors (bHLH) | bHLH-ZIP | USF | 11 | 14.02 | 71.56 | USF3.FL@CHS@shabby-razzmatazz-uakari@HughesLab.MEME@1000\_fa\_memechip\_Motif1.ppm |
| USF3 | CHS | test | THC\_0661 |  |  | FL | 1.2.6.2.3 | Basic domains | Basic helix-loop-helix factors (bHLH) | bHLH-ZIP | USF | 15 | 14.99 | 72.42 | USF3.FL@CHS@ready-orchid-uguisu@ |
| USF3 | GHTS.GFPIVT | test | YWP\_B\_AffSeq\_F9\_USF3-DBD |  |  | DBD | 1.2.6.2.3 | Basic domains | Basic helix-loop-helix factors (bHLH) | bHLH-ZIP | USF | 35 | 16.26 | 58.69 | USF3.DBD@AFS.GFPIVT@droopy-vermilion-chow@HughesLab.Homer@1000\_fa\_homer\_minw3\_maxw\_40\_Motif1.ppm |
| USF3 | GHTS.Lys | train | YWL\_B\_AffSeq\_G10\_USF3 |  |  | FL | 1.2.6.2.3 | Basic domains | Basic helix-loop-helix factors (bHLH) | bHLH-ZIP | USF | 12 | 10.34 | 61.44 | USF3.FL@HTS.Lys@stuffy-sangria-mongrel+cranky-olivine-akita+bluesy-magnolia-capybara+scanty-pear-barracuda@faltejsk.ProBound@motif\_with\_ns.ppm |
| USF3 | PBM | test | PBM13848 |  |  | DBD | 1.2.6.2.3 | Basic domains | Basic helix-loop-helix factors (bHLH) | bHLH-ZIP | USF | 8 | 14.28 | 71.5 | USF3.DBD@AFS.GFPIVT@stuffy-charcoal-numbat@HughesLab.Homer@topk\_cycle=C3\_k=5\_top=1000\_fasta\_homer\_Motif1.ppm |
| USF3 | HTS.IVT | test | YWC\_B\_CG40NACGGTT |  |  | FL | 1.2.6.2.3 | Basic domains | Basic helix-loop-helix factors (bHLH) | bHLH-ZIP | USF | 12 | 10.34 | 61.44 | USF3.FL@HTS.Lys@stuffy-sangria-mongrel+cranky-olivine-akita+bluesy-magnolia-capybara+scanty-pear-barracuda@faltejsk.ProBound@motif\_with\_ns.ppm |
| USF3 | HTS.GFPIVT | test | YWP\_A\_GT40NTTGGTG |  |  | FL | 1.2.6.2.3 | Basic domains | Basic helix-loop-helix factors (bHLH) | bHLH-ZIP | USF | 12 | 10.34 | 61.44 | USF3.FL@HTS.Lys@stuffy-sangria-mongrel+cranky-olivine-akita+bluesy-magnolia-capybara+scanty-pear-barracuda@faltejsk.ProBound@motif\_with\_ns.ppm |
| USF3 | HTS.Lys | test | YWL\_A\_TC40NAATCTC |  |  | FL | 1.2.6.2.3 | Basic domains | Basic helix-loop-helix factors (bHLH) | bHLH-ZIP | USF | 12 | 10.34 | 61.44 | USF3.FL@HTS.Lys@stuffy-sangria-mongrel+cranky-olivine-akita+bluesy-magnolia-capybara+scanty-pear-barracuda@faltejsk.ProBound@motif\_with\_ns.ppm |
| USF3 | SMS | test | UT380-241 |  |  | FL | 1.2.6.2.3 | Basic domains | Basic helix-loop-helix factors (bHLH) | bHLH-ZIP | USF | 12 | 10.34 | 61.44 | USF3.FL@HTS.Lys@stuffy-sangria-mongrel+cranky-olivine-akita+bluesy-magnolia-capybara+scanty-pear-barracuda@faltejsk.ProBound@motif\_with\_ns.ppm |
| ZNF395 | CHS | train | THC\_0294.Rep-DIANA\_0293 |  |  | DBD | 2.3.2.255.2 | Zinc-coordinating DNA-binding domains | C2H2 zinc finger factors | Other with up to three adjacent zinc fingers | CR1-CR2 | 12 | 5.93 | 65.05 | ZNF395.DBD@HTS.IVT@sleazy-indigo-boar+stinky-periwinkle-hornet+greasy-aquamarine-robin+crappy-cerulean-impala@faltejsk.ProBound@motif\_with\_ns.ppm |
| ZNF395 | CHS | test | THC\_0294.Rep-MICHELLE\_0314 |  |  | DBD | 2.3.2.255.2 | Zinc-coordinating DNA-binding domains | C2H2 zinc finger factors | Other with up to three adjacent zinc fingers | CR1-CR2 | 12 | 5.93 | 65.05 | ZNF395.DBD@HTS.IVT@sleazy-indigo-boar+stinky-periwinkle-hornet+greasy-aquamarine-robin+crappy-cerulean-impala@faltejsk.ProBound@motif\_with\_ns.ppm |
| ZNF395 | GHTS.IVT | train | YWH\_B\_AffSeq\_B11\_ZNF395 |  |  | DBD | 2.3.2.255.2 | Zinc-coordinating DNA-binding domains | C2H2 zinc finger factors | Other with up to three adjacent zinc fingers | CR1-CR2 | 10 | 12.76 | 87.75 | ZNF395.DBD@AFS.IVT@randy-ecru-macaw@autosome-ru.ChIPMunk@995seq\_7to21\_m0.pcm |
| ZNF395 | GHTS.GFPIVT | train | YWR\_B\_AffSeq\_H5\_ZNF395-FL |  |  | DBD | 2.3.2.255.2 | Zinc-coordinating DNA-binding domains | C2H2 zinc finger factors | Other with up to three adjacent zinc fingers | CR1-CR2 | 20 | 7.13 | 61.01 | ZNF395.DBD@AFS.GFPIVT@pasty-ochre-rabbit+foggy-champagne-bison+tasty-ivory-sheepdog@Halle.Dimont@Motif\_1\_w20\_astrained.ppm |
| ZNF395 | GHTS.GFPIVT | test | YWM\_B\_AffSeq\_E12\_ZNF395-DBD |  |  | DBD | 2.3.2.255.2 | Zinc-coordinating DNA-binding domains | C2H2 zinc finger factors | Other with up to three adjacent zinc fingers | CR1-CR2 | 20 | 7.13 | 61.01 | ZNF395.DBD@AFS.GFPIVT@pasty-ochre-rabbit+foggy-champagne-bison+tasty-ivory-sheepdog@Halle.Dimont@Motif\_1\_w20\_astrained.ppm |
| ZNF395 | GHTS.Lys | train | YWK\_D\_AffSeq\_A6\_ZNF395 |  |  | FL | 2.3.2.255.2 | Zinc-coordinating DNA-binding domains | C2H2 zinc finger factors | Other with up to three adjacent zinc fingers | CR1-CR2 | 10 | 15.21 | 92.3 | ZNF395.FL@AFS.GFPIVT@chummy-white-chipmunk@autosome-ru.ChIPMunk@969seq\_7to21\_m0.pcm |
| ZNF395 | GHTS.Lys | train | YWK\_B\_AffSeq\_A6\_ZNF395 |  |  | DBD | 2.3.2.255.2 | Zinc-coordinating DNA-binding domains | C2H2 zinc finger factors | Other with up to three adjacent zinc fingers | CR1-CR2 | 20 | 7.13 | 61.01 | ZNF395.DBD@AFS.GFPIVT@pasty-ochre-rabbit+foggy-champagne-bison+tasty-ivory-sheepdog@Halle.Dimont@Motif\_1\_w20\_astrained.ppm |
| ZNF395 | HTS.IVT | test | YWH\_A\_AT40NCGGGTT |  |  | DBD | 2.3.2.255.2 | Zinc-coordinating DNA-binding domains | C2H2 zinc finger factors | Other with up to three adjacent zinc fingers | CR1-CR2 | 12 | 2.64 | 57.83 | ZNF395.DBD@HTS.IVT@sleazy-indigo-boar+stinky-periwinkle-hornet+greasy-aquamarine-robin+crappy-cerulean-impala@faltejsk.ProBound@motif\_without\_ns.ppm |
| ZNF395 | HTS.GFPIVT | test | YWR\_A\_TG40NGACCTT |  |  | FL | 2.3.2.255.2 | Zinc-coordinating DNA-binding domains | C2H2 zinc finger factors | Other with up to three adjacent zinc fingers | CR1-CR2 | 12 | 1.52 | 49.1 | ZNF395.FL@HTS.Lys@sloppy-carmine-whippet+droopy-scarlet-macaque+gamy-xanthic-cow+paltry-purple-fox@faltejsk.ProBound@motif\_without\_ns.ppm |
| ZNF395 | HTS.GFPIVT | test | YWM\_A\_GG40NGTGAGA |  |  | DBD | 2.3.2.255.2 | Zinc-coordinating DNA-binding domains | C2H2 zinc finger factors | Other with up to three adjacent zinc fingers | CR1-CR2 | 12 | 2.64 | 57.83 | ZNF395.DBD@HTS.IVT@sleazy-indigo-boar+stinky-periwinkle-hornet+greasy-aquamarine-robin+crappy-cerulean-impala@faltejsk.ProBound@motif\_without\_ns.ppm |
| ZNF395 | HTS.Lys | test | YWK\_C\_AA40NGTGGTG |  |  | FL | 2.3.2.255.2 | Zinc-coordinating DNA-binding domains | C2H2 zinc finger factors | Other with up to three adjacent zinc fingers | CR1-CR2 | 12 | 3.61 | 53.36 | ZNF395.FL@HTS.GFPIVT@greasy-denim-opossum+grumpy-cyan-scorpion+scanty-lime-bird@faltejsk.ProBound@motif\_without\_ns.ppm |
| ZNF395 | HTS.Lys | test | YWK\_A\_AA40NGTGGTG |  |  | FL | 2.3.2.255.2 | Zinc-coordinating DNA-binding domains | C2H2 zinc finger factors | Other with up to three adjacent zinc fingers | CR1-CR2 | 12 | 3.61 | 53.36 | ZNF395.FL@HTS.GFPIVT@greasy-denim-opossum+grumpy-cyan-scorpion+scanty-lime-bird@faltejsk.ProBound@motif\_without\_ns.ppm |
| ZNF395 | SMS | test | UT380-338 |  |  | NA | 2.3.2.255.2 | Zinc-coordinating DNA-binding domains | C2H2 zinc finger factors | Other with up to three adjacent zinc fingers | CR1-CR2 | 15 | 15.52 | 69.77 | ZNF395.NA@SMS@lumpy-indigo-hound@autosome-ru.ChIPMunk@topk\_k=5\_top=500.pcm |
| ZNF395 | SMS | test | UT380-337 |  |  | NA | 2.3.2.255.2 | Zinc-coordinating DNA-binding domains | C2H2 zinc finger factors | Other with up to three adjacent zinc fingers | CR1-CR2 | 8 | 8.06 | 68.68 | ZNF395.NA@SMS@lumpy-indigo-hound@autosome-ru.ChIPMunk@topk\_k=5\_top=10000.pcm |
| ZNF367 | CHS | test | THC\_0045 |  |  | FL | 2.3.2.4.256 | Zinc-coordinating DNA-binding domains | C2H2 zinc finger factors | Other with up to three adjacent zinc fingers | Other | 12 | 13.23 | 33.78 | ZNF367.FL@CHS@goopy-razzmatazz-armadillo@autosome-ru.ChIPMunk@500seq\_25to7\_m0.pcm |
| ZNF367 | CHS | train | THC\_0046 |  |  | FL | 2.3.2.4.256 | Zinc-coordinating DNA-binding domains | C2H2 zinc finger factors | Other with up to three adjacent zinc fingers | Other | 12 | 13.23 | 33.78 | ZNF367.FL@CHS@goopy-razzmatazz-armadillo@autosome-ru.ChIPMunk@500seq\_25to7\_m0.pcm |
| ZNF367 | GHTS.GFPIVT | train | YWR\_B\_AffSeq\_E5\_ZNF367-FL |  |  | FL | 2.3.2.4.256 | Zinc-coordinating DNA-binding domains | C2H2 zinc finger factors | Other with up to three adjacent zinc fingers | Other | 20 | 8.59 | 43.42 | ZNF367.FL@AFS.GFPIVT@flimsy-heliotrope-mandrill+whiny-turquoise-bear+whiny-purple-dachsbracke@Halle.Dimont@Motif\_1\_w20\_astrained.ppm |
| ZNF367 | GHTS.Lys | test | YWD\_B\_AffSeq\_H08\_ZNF367 |  |  | FL | 2.3.2.4.256 | Zinc-coordinating DNA-binding domains | C2H2 zinc finger factors | Other with up to three adjacent zinc fingers | Other | 20 | 8.59 | 43.42 | ZNF367.FL@AFS.GFPIVT@flimsy-heliotrope-mandrill+whiny-turquoise-bear+whiny-purple-dachsbracke@Halle.Dimont@Motif\_1\_w20\_astrained.ppm |
| ZNF367 | HTS.GFPIVT | test | YWR\_A\_GA40NACTTTG |  |  | FL | 2.3.2.4.256 | Zinc-coordinating DNA-binding domains | C2H2 zinc finger factors | Other with up to three adjacent zinc fingers | Other | 12 | 4.14 | 37.34 | ZNF367.FL@HTS.Lys@gloppy-russet-goat+flimsy-bronze-dragonfly+bumpy-myrtle-angelfish+bumpy-myrtle-zebu@faltejsk.ProBound@motif\_without\_ns.ppm |
| ZNF367 | HTS.Lys | test | YWD\_A\_TG40NCTGTGG |  |  | NA | 2.3.2.4.256 | Zinc-coordinating DNA-binding domains | C2H2 zinc finger factors | Other with up to three adjacent zinc fingers | Other | 12 | 5.39 | 33.54 | ZNF367.NA@SMS@bluesy-purple-drever@faltejsk.ProBound@motif\_without\_ns.ppm |
| ZNF367 | SMS | test | UT380-333 |  |  | FL | 2.3.2.4.256 | Zinc-coordinating DNA-binding domains | C2H2 zinc finger factors | Other with up to three adjacent zinc fingers | Other | 12 | 2.25 | 47.13 | ZNF367.FL@AFS.Lys@woolly-brass-barnacle+gimpy-scarlet-shrew+geeky-coral-tarsier+nippy-indigo-warthog@faltejsk.ProBound@motif\_without\_ns.ppm |
| ZNF493 | CHS | train | THC\_0389.Rep-MICHELLE\_0314 |  |  | FL | 2.3.3.0.229 | Zinc-coordinating DNA-binding domains | C2H2 zinc finger factors | More than 3 adjacent zinc fingers | Unclassified | 18 | 3.09 | 53.14 | ZNF493.FL@AFS.GFPIVT@chummy-viridian-wasp@Halle.Dimont@Motif\_1\_imw20\_astrained.ppm |
| ZNF493 | CHS | train | THC\_0334.Rep-DIANA\_0293 |  |  | DBD | 2.3.3.0.229 | Zinc-coordinating DNA-binding domains | C2H2 zinc finger factors | More than 3 adjacent zinc fingers | Unclassified | 18 | 3.82 | 53.67 | ZNF493.DBD@AFS.GFPIVT@smelly-chartreuse-gecko@Halle.Dimont@Motif\_1\_imw20\_astrained.ppm |
| ZNF493 | CHS | train | THC\_0334.Rep-MICHELLE\_0314 |  |  | FL | 2.3.3.0.229 | Zinc-coordinating DNA-binding domains | C2H2 zinc finger factors | More than 3 adjacent zinc fingers | Unclassified | 20 | 7.01 | 59.27 | ZNF493.FL@AFS.GFPIVT@shaky-eggplant-okapi@Halle.Dimont@Motif\_1\_w20\_astrained.ppm |
| ZNF493 | CHS | test | THC\_0891 |  |  | FL | 2.3.3.0.229 | Zinc-coordinating DNA-binding domains | C2H2 zinc finger factors | More than 3 adjacent zinc fingers | Unclassified | 20 | 7.44 | 59.14 | ZNF493.FL@AFS.GFPIVT@chummy-viridian-wasp@Halle.Dimont@Motif\_1\_w20\_astrained.ppm |
| ZNF493 | GHTS.GFPIVT | train | YWR\_B\_AffSeq\_H7\_ZNF493-FL |  |  | DBD | 2.3.3.0.229 | Zinc-coordinating DNA-binding domains | C2H2 zinc finger factors | More than 3 adjacent zinc fingers | Unclassified | 20 | 7.05 | 58.14 | ZNF493.DBD@AFS.GFPIVT@smelly-chartreuse-gecko@Halle.Dimont@Motif\_1\_w20\_astrained.ppm |
| ZNF493 | GHTS.GFPIVT | test | YWR\_B\_AffSeq\_G7\_ZNF493-DBD2 |  |  | DBD | 2.3.3.0.229 | Zinc-coordinating DNA-binding domains | C2H2 zinc finger factors | More than 3 adjacent zinc fingers | Unclassified | 20 | 6.92 | 57.97 | ZNF493.DBD@AFS.GFPIVT@sunny-periwinkle-molly@Halle.Dimont@Motif\_1\_w20\_astrained.ppm |
| ZNF493 | HTS.GFPIVT | test | YWR\_A\_TG40NTCTTGC |  |  | FL | 2.3.3.0.229 | Zinc-coordinating DNA-binding domains | C2H2 zinc finger factors | More than 3 adjacent zinc fingers | Unclassified | 15 | 18.01 | 58.42 | ZNF493.FL@HTS.GFPIVT@crabby-smalt-wolfhound+craggy-auburn-catfish+squirrely-carmine-squirt@HughesLab.MEME@topk\_cycle=C1+C2+C3\_k=5\_top=10000\_Motif1.ppm |
| ZNF493 | HTS.GFPIVT | test | YWR\_A\_TC40NTGCATA |  |  | DBD | 2.3.3.0.229 | Zinc-coordinating DNA-binding domains | C2H2 zinc finger factors | More than 3 adjacent zinc fingers | Unclassified | 15 | 21.63 | 65.05 | ZNF493.DBD@HTS.GFPIVT@silly-burgundy-dragon@HughesLab.MEME@topk\_cycle=C3\_k=5\_top=1000\_Motif1.ppm |
| ZNF493 | SMS | test | UT380-354 |  |  | DBD | 2.3.3.0.229 | Zinc-coordinating DNA-binding domains | C2H2 zinc finger factors | More than 3 adjacent zinc fingers | Unclassified | 15 | 18.13 | 64.14 | ZNF493.DBD@HTS.GFPIVT@boozy-beige-peccary+silly-burgundy-dragon+tasty-firebrick-turtle@HughesLab.MEME@topk\_cycle=C1+C2+C3\_k=5\_top=10000\_Motif1.ppm |
| ZNF493 | SMS | test | UT380-354-2 |  |  | DBD | 2.3.3.0.229 | Zinc-coordinating DNA-binding domains | C2H2 zinc finger factors | More than 3 adjacent zinc fingers | Unclassified | 17 | 16.48 | 61.21 | ZNF493.DBD@HTS.GFPIVT@tasty-firebrick-turtle@AJolma.Autoseed@Multinom2\_Onehit\_Seed\_DDGGGCACTGDBHYGGBackgroundCyc1.ppm |
| ZNF648 | CHS | test | THC\_0679 |  |  | FL | 2.3.3.0.94 | Zinc-coordinating DNA-binding domains | C2H2 zinc finger factors | More than 3 adjacent zinc fingers | Unclassified | 20 | 7.59 | 64.1 | ZNF648.FL@AFS.GFPIVT@messy-magenta-bat@Halle.Dimont@Motif\_1\_w20\_astrained.ppm |
| ZNF648 | CHS | train | THC\_0775 |  |  | FL | 2.3.3.0.94 | Zinc-coordinating DNA-binding domains | C2H2 zinc finger factors | More than 3 adjacent zinc fingers | Unclassified | 15 | 13.66 | 70.56 | ZNF648.FL@CHS@wimpy-myrtle-dollar@ |
| ZNF648 | GHTS.GFPIVT | test | YWS\_B\_AffSeq\_A2\_ZNF648-FL |  |  | FL | 2.3.3.0.94 | Zinc-coordinating DNA-binding domains | C2H2 zinc finger factors | More than 3 adjacent zinc fingers | Unclassified | 31 | 20.31 | 68.16 | ZNF648.FL@AFS.GFPIVT@freaky-tan-tamarin+jumpy-cyan-capybara+shabby-khaki-rhinoceros@HughesLab.Homer@topk\_cycle=C1+C2+C3\_k=5\_top=10000\_fasta\_homer\_minw3\_maxw\_40\_Motif2.ppm |
| ZNF648 | GHTS.Lys | train | YWL\_B\_AffSeq\_D10\_ZNF648 |  |  | FL | 2.3.3.0.94 | Zinc-coordinating DNA-binding domains | C2H2 zinc finger factors | More than 3 adjacent zinc fingers | Unclassified | 26 | 16.17 | 61.06 | ZNF648.FL@HTS.GFPIVT@sleepy-persimmon-quetzal@AJolma.Autoseed@Multinom2\_Onehit\_Seed\_DRGGGGRGNNNNRDNVNNNYGCTGRBackgroundCyc1.ppm |
| ZNF648 | HTS.GFPIVT | test | YWS\_A\_AA40NCCGCGT |  |  | FL | 2.3.3.0.94 | Zinc-coordinating DNA-binding domains | C2H2 zinc finger factors | More than 3 adjacent zinc fingers | Unclassified | 26 | 22.04 | 62.91 | ZNF648.FL@HTS.GFPIVT@scummy-lemon-guppy@autosome-ru.ChIPMunk@topk\_cycle=C3\_k=5\_top=1000.pcm |
| ZNF648 | SMS | test | UT380-406 |  |  | FL | 2.3.3.0.94 | Zinc-coordinating DNA-binding domains | C2H2 zinc finger factors | More than 3 adjacent zinc fingers | Unclassified | 26 | 24.48 | 62.23 | ZNF648.FL@HTS.GFPIVT@crappy-persimmon-insect+sleepy-persimmon-quetzal+scummy-lemon-guppy@OFornes.ExplaiNN@filter76\_1.ppm |
| ZNF648 | SMS | test | UT380-405 |  |  | FL | 2.3.3.0.94 | Zinc-coordinating DNA-binding domains | C2H2 zinc finger factors | More than 3 adjacent zinc fingers | Unclassified | 26 | 24.48 | 62.23 | ZNF648.FL@HTS.GFPIVT@crappy-persimmon-insect+sleepy-persimmon-quetzal+scummy-lemon-guppy@OFornes.ExplaiNN@filter76\_1.ppm |
| ZNF20 | CHS | train | THC\_0341.Rep-DIANA\_0293 |  |  | FL | 2.3.3.33.17 | Zinc-coordinating DNA-binding domains | C2H2 zinc finger factors | More than 3 adjacent zinc fingers | ZNF763-like | 17 | 3.76 | 51.97 | ZNF20.FL@AFS.GFPIVT@breezy-sapphire-oriole@Halle.Dimont@Motif\_1\_imw20\_astrained.ppm |
| ZNF20 | CHS | train | THC\_0395.Rep-DIANA\_0293 |  |  | FL | 2.3.3.33.17 | Zinc-coordinating DNA-binding domains | C2H2 zinc finger factors | More than 3 adjacent zinc fingers | ZNF763-like | 20 | 7.4 | 57.74 | ZNF20.FL@CHS@stuffy-khaki-ray@Halle.Dimont@Motif\_1\_w20\_astrained.ppm |
| ZNF20 | CHS | train | THC\_0341.Rep-MICHELLE\_0314 |  |  | FL | 2.3.3.33.17 | Zinc-coordinating DNA-binding domains | C2H2 zinc finger factors | More than 3 adjacent zinc fingers | ZNF763-like | 20 | 7.4 | 57.74 | ZNF20.FL@CHS@stuffy-khaki-ray@Halle.Dimont@Motif\_1\_w20\_astrained.ppm |
| ZNF20 | CHS | test | THC\_0395.Rep-MICHELLE\_0314 |  |  | FL | 2.3.3.33.17 | Zinc-coordinating DNA-binding domains | C2H2 zinc finger factors | More than 3 adjacent zinc fingers | ZNF763-like | 20 | 7.4 | 57.74 | ZNF20.FL@CHS@stuffy-khaki-ray@Halle.Dimont@Motif\_1\_w20\_astrained.ppm |
| ZNF20 | GHTS.GFPIVT | train | YWR\_B\_AffSeq\_B1\_ZNF20-FL |  |  | FL | 2.3.3.33.17 | Zinc-coordinating DNA-binding domains | C2H2 zinc finger factors | More than 3 adjacent zinc fingers | ZNF763-like | 20 | 7.67 | 55.17 | ZNF20.FL@AFS.GFPIVT@lumpy-cobalt-panda@Halle.Dimont@Motif\_1\_w20\_astrained.ppm |
| ZNF20 | HTS.GFPIVT | test | YWR\_A\_AG40NTCGACT |  |  | FL | 2.3.3.33.17 | Zinc-coordinating DNA-binding domains | C2H2 zinc finger factors | More than 3 adjacent zinc fingers | ZNF763-like | 20 | 7.05 | 55.15 | ZNF20.FL@AFS.GFPIVT@seedy-asparagus-markhor+breezy-sapphire-oriole+lumpy-cobalt-panda@Halle.Dimont@Motif\_1\_w20\_astrained.ppm |
| PRDM13 | CHS | train | THC\_0223 |  |  | FL | 2.3.4.0.12 | Zinc-coordinating DNA-binding domains | C2H2 zinc finger factors | Multiple dispersed zinc fingers | Unclassified | 10 | 11.44 | 69.01 | PRDM13.FL@CHS@leaky-buff-wolf@autosome-ru.ChIPMunk@999seq\_15to7\_m0.pcm |
| PRDM13 | CHS | test | THC\_0231 |  |  | FL | 2.3.4.0.12 | Zinc-coordinating DNA-binding domains | C2H2 zinc finger factors | Multiple dispersed zinc fingers | Unclassified | 20 | 7.6 | 59.13 | PRDM13.FL@AFS.Lys@sloppy-celadon-bandicoot@Halle.Dimont@Motif\_1\_w20\_astrained.ppm |
| PRDM13 | GHTS.IVT | train | YWH\_B\_AffSeq\_B05\_PRDM13\_DBD |  |  | DBD | 2.3.4.0.12 | Zinc-coordinating DNA-binding domains | C2H2 zinc finger factors | Multiple dispersed zinc fingers | Unclassified | 20 | 7.73 | 58.65 | PRDM13.DBD@AFS.GFPIVT@wiggy-seashell-ant+jumpy-pumpkin-koala+sleepy-platinum-dormouse@Halle.Dimont@Motif\_1\_w20\_astrained.ppm |
| PRDM13 | GHTS.GFPIVT | train | YWQ\_B\_AffSeq\_G8\_PRDM13-FL |  |  | DBD | 2.3.4.0.12 | Zinc-coordinating DNA-binding domains | C2H2 zinc finger factors | Multiple dispersed zinc fingers | Unclassified | 20 | 8.17 | 62.41 | PRDM13.DBD@AFS.IVT@stealthy-pink-hare@Halle.Dimont@Motif\_1\_w20\_astrained.ppm |
| PRDM13 | GHTS.GFPIVT | train | YWP\_B\_AffSeq\_C12\_PRDM13-DBD |  |  | DBD | 2.3.4.0.12 | Zinc-coordinating DNA-binding domains | C2H2 zinc finger factors | Multiple dispersed zinc fingers | Unclassified | 12 | 10.17 | 61.56 | PRDM13.DBD@HTS.GFPIVT@nippy-burgundy-oriole+queasy-eggplant-mayfly+flaky-brown-horse@faltejsk.ProBound@motif\_with\_ns.ppm |
| PRDM13 | GHTS.Lys | train | YWK\_D\_AffSeq\_F1\_PRDM13 |  |  | FL | 2.3.4.0.12 | Zinc-coordinating DNA-binding domains | C2H2 zinc finger factors | Multiple dispersed zinc fingers | Unclassified | 20 | 7.6 | 59.13 | PRDM13.FL@AFS.Lys@sloppy-celadon-bandicoot@Halle.Dimont@Motif\_1\_w20\_astrained.ppm |
| PRDM13 | GHTS.Lys | test | YWK\_B\_AffSeq\_F1\_PRDM13 |  |  | FL | 2.3.4.0.12 | Zinc-coordinating DNA-binding domains | C2H2 zinc finger factors | Multiple dispersed zinc fingers | Unclassified | 20 | 7.6 | 59.13 | PRDM13.FL@AFS.Lys@sloppy-celadon-bandicoot@Halle.Dimont@Motif\_1\_w20\_astrained.ppm |
| PRDM13 | HTS.IVT | test | YWH\_A\_AG40NGCTCTG |  |  | DBD | 2.3.4.0.12 | Zinc-coordinating DNA-binding domains | C2H2 zinc finger factors | Multiple dispersed zinc fingers | Unclassified | 10 | 8.17 | 59.71 | PRDM13.DBD@SMS@muggy-platinum-orangutan@AJolma.Autoseed@Multinom1\_OneHitAllReads\_Seed\_GNYACCWGYN.ppm |
| PRDM13 | HTS.GFPIVT | test | YWP\_A\_CG40NAATAGC |  |  | FL | 2.3.4.0.12 | Zinc-coordinating DNA-binding domains | C2H2 zinc finger factors | Multiple dispersed zinc fingers | Unclassified | 12 | 2.88 | 54.81 | PRDM13.FL@HTS.GFPIVT@flaky-jade-eel+crappy-emerald-mastiff+chummy-turquoise-newfoundland@faltejsk.ProBound@motif\_without\_ns.ppm |
| PRDM13 | HTS.GFPIVT | test | YWQ\_A\_TC40NGCGATT |  |  | FL | 2.3.4.0.12 | Zinc-coordinating DNA-binding domains | C2H2 zinc finger factors | Multiple dispersed zinc fingers | Unclassified | 12 | 2.88 | 54.81 | PRDM13.FL@HTS.GFPIVT@flaky-jade-eel+crappy-emerald-mastiff+chummy-turquoise-newfoundland@faltejsk.ProBound@motif\_without\_ns.ppm |
| PRDM13 | HTS.Lys | test | YWK\_A\_GG40NCGTAGT |  |  | FL | 2.3.4.0.12 | Zinc-coordinating DNA-binding domains | C2H2 zinc finger factors | Multiple dispersed zinc fingers | Unclassified | 12 | 2.88 | 54.81 | PRDM13.FL@HTS.GFPIVT@flaky-jade-eel+crappy-emerald-mastiff+chummy-turquoise-newfoundland@faltejsk.ProBound@motif\_without\_ns.ppm |
| PRDM13 | HTS.Lys | test | YWK\_C\_GG40NCGTAGT |  |  | FL | 2.3.4.0.12 | Zinc-coordinating DNA-binding domains | C2H2 zinc finger factors | Multiple dispersed zinc fingers | Unclassified | 12 | 2.88 | 54.81 | PRDM13.FL@HTS.GFPIVT@flaky-jade-eel+crappy-emerald-mastiff+chummy-turquoise-newfoundland@faltejsk.ProBound@motif\_without\_ns.ppm |
| PRDM13 | SMS | test | UT380-154 |  |  | DBD | 2.3.4.0.12 | Zinc-coordinating DNA-binding domains | C2H2 zinc finger factors | Multiple dispersed zinc fingers | Unclassified | 9 | 7.81 | 59.56 | PRDM13.DBD@SMS@muggy-platinum-orangutan@Halle.Dimont@Motif\_1\_posneg\_astrained.ppm |
| ZNF251 | CHS | train | THC\_0300.Rep-DIANA\_0293 |  |  | FL | 2.3.4.0.52 | Zinc-coordinating DNA-binding domains | C2H2 zinc finger factors | Multiple dispersed zinc fingers | Unclassified | 20 | 5.83 | 53.87 | ZNF251.FL@CHS@sleazy-thistle-sheepdog@Halle.Dimont@Motif\_1\_w20\_astrained.ppm |
| ZNF251 | CHS | train | THC\_0445.Rep-DIANA\_0293 |  |  | FL | 2.3.4.0.52 | Zinc-coordinating DNA-binding domains | C2H2 zinc finger factors | Multiple dispersed zinc fingers | Unclassified | 20 | 5.83 | 53.87 | ZNF251.FL@CHS@sleazy-thistle-sheepdog@Halle.Dimont@Motif\_1\_w20\_astrained.ppm |
| ZNF251 | CHS | test | THC\_0300.Rep-MICHELLE\_0314 |  |  | FL | 2.3.4.0.52 | Zinc-coordinating DNA-binding domains | C2H2 zinc finger factors | Multiple dispersed zinc fingers | Unclassified | 20 | 5.83 | 53.87 | ZNF251.FL@CHS@sleazy-thistle-sheepdog@Halle.Dimont@Motif\_1\_w20\_astrained.ppm |
| ZNF251 | CHS | train | THC\_0445.Rep-MICHELLE\_0314 |  |  | FL | 2.3.4.0.52 | Zinc-coordinating DNA-binding domains | C2H2 zinc finger factors | Multiple dispersed zinc fingers | Unclassified | 20 | 5.83 | 53.87 | ZNF251.FL@CHS@sleazy-thistle-sheepdog@Halle.Dimont@Motif\_1\_w20\_astrained.ppm |
| ZNF251 | GHTS.GFPIVT | test | YWR\_B\_AffSeq\_H2\_ZNF251-DBD |  |  | DBD | 2.3.4.0.52 | Zinc-coordinating DNA-binding domains | C2H2 zinc finger factors | Multiple dispersed zinc fingers | Unclassified | 20 | 5.92 | 54.52 | ZNF251.DBD@AFS.GFPIVT@sleazy-cardinal-buzzard+snoopy-wisteria-havanese+flaky-beige-kiwi@Halle.Dimont@Motif\_1\_w20\_astrained.ppm |
| ZNF251 | GHTS.GFPIVT | train | YWR\_B\_AffSeq\_A3\_ZNF251-FL |  |  | DBD | 2.3.4.0.52 | Zinc-coordinating DNA-binding domains | C2H2 zinc finger factors | Multiple dispersed zinc fingers | Unclassified | 20 | 5.92 | 54.52 | ZNF251.DBD@AFS.GFPIVT@sleazy-cardinal-buzzard+snoopy-wisteria-havanese+flaky-beige-kiwi@Halle.Dimont@Motif\_1\_w20\_astrained.ppm |
| ZNF251 | HTS.GFPIVT | test | YWR\_A\_AA40NAGTGGT |  |  | DBD | 2.3.4.0.52 | Zinc-coordinating DNA-binding domains | C2H2 zinc finger factors | Multiple dispersed zinc fingers | Unclassified | 20 | 13.0 | 47.83 | ZNF251.DBD@HTS.GFPIVT@squeaky-ruby-caiman@AJolma.Autoseed@Multinom2\_Onehit\_Seed\_RNWANBGCGYTAAGYGHNWBackgroundCyc1.ppm |
| ZNF251 | HTS.GFPIVT | test | YWR\_A\_TC40NGTTTTG |  |  | DBD | 2.3.4.0.52 | Zinc-coordinating DNA-binding domains | C2H2 zinc finger factors | Multiple dispersed zinc fingers | Unclassified | 20 | 13.0 | 47.83 | ZNF251.DBD@HTS.GFPIVT@squeaky-ruby-caiman@AJolma.Autoseed@Multinom2\_Onehit\_Seed\_RNWANBGCGYTAAGYGHNWBackgroundCyc1.ppm |
| ZNF251 | SMS | test | UT380-309 |  |  | DBD | 2.3.4.0.52 | Zinc-coordinating DNA-binding domains | C2H2 zinc finger factors | Multiple dispersed zinc fingers | Unclassified | 15 | 3.65 | 48.47 | ZNF251.DBD@HTS.GFPIVT@pokey-wheat-bison+squeaky-ruby-caiman+nerdy-tangerine-crane@Halle.Dimont@Motif\_1\_sampled\_e4\_astrained.ppm |
| ZNF518B | CHS | train | THC\_0669 |  |  | FL | 2.3.4.18.2 | Zinc-coordinating DNA-binding domains | C2H2 zinc finger factors | Multiple dispersed zinc fingers | ZNF518 | 20 | 5.43 | 52.77 | ZNF518B.FL@CHS@smelly-silver-heron@Halle.Dimont@Motif\_1\_w20\_astrained.ppm |
| ZNF518B | CHS | test | THC\_0765 |  |  | FL | 2.3.4.18.2 | Zinc-coordinating DNA-binding domains | C2H2 zinc finger factors | Multiple dispersed zinc fingers | ZNF518 | 20 | 5.43 | 52.77 | ZNF518B.FL@CHS@smelly-silver-heron@Halle.Dimont@Motif\_1\_w20\_astrained.ppm |
| ZNF518B | GHTS.GFPIVT | train | YWR\_B\_AffSeq\_C9\_ZNF518B-DBD |  |  | FL | 2.3.4.18.2 | Zinc-coordinating DNA-binding domains | C2H2 zinc finger factors | Multiple dispersed zinc fingers | ZNF518 | 17 | 17.76 | 42.85 | ZNF518B.FL@AFS.Lys@seedy-buff-maltese@autosome-ru.ChIPMunk@968seq\_7to21\_m0.pcm |
| ZNF518B | GHTS.Lys | train | YWN\_B\_AffSeq\_H1\_ZNF518B |  |  | FL | 2.3.4.18.2 | Zinc-coordinating DNA-binding domains | C2H2 zinc finger factors | Multiple dispersed zinc fingers | ZNF518 | 20 | 7.24 | 53.94 | ZNF518B.FL@AFS.Lys@frumpy-puce-ostrich+seedy-buff-maltese+gloppy-thistle-cuscus+snazzy-peach-albatross@Halle.Dimont@Motif\_1\_w20\_astrained.ppm |
| ZNF518B | GHTS.Lys | test | YWO\_B\_AffSeq\_H1\_ZNF518B |  |  | FL | 2.3.4.18.2 | Zinc-coordinating DNA-binding domains | C2H2 zinc finger factors | Multiple dispersed zinc fingers | ZNF518 | 20 | 7.99 | 54.18 | ZNF518B.FL@AFS.Lys@gloppy-thistle-cuscus@Halle.Dimont@Motif\_1\_w20\_astrained.ppm |
| ZNF518B | HTS.GFPIVT | test | YWR\_A\_CC40NGACATG |  |  | FL | 2.3.4.18.2 | Zinc-coordinating DNA-binding domains | C2H2 zinc finger factors | Multiple dispersed zinc fingers | ZNF518 | 12 | 9.14 | 45.89 | ZNF518B.FL@HTS.Lys@boozy-harlequin-rattlesnake+shaggy-cornflower-fossa+cheeky-salmon-setter+boozy-cerise-mau@faltejsk.ProBound@motif\_with\_ns.ppm |
| ZNF518B | HTS.Lys | test | YWO\_A\_TC40NGGCTGT |  |  | FL | 2.3.4.18.2 | Zinc-coordinating DNA-binding domains | C2H2 zinc finger factors | Multiple dispersed zinc fingers | ZNF518 | 12 | 9.14 | 45.89 | ZNF518B.FL@HTS.Lys@boozy-harlequin-rattlesnake+shaggy-cornflower-fossa+cheeky-salmon-setter+boozy-cerise-mau@faltejsk.ProBound@motif\_with\_ns.ppm |
| ZNF518B | HTS.Lys | test | YWN\_A\_TC40NGGCTGT |  |  | FL | 2.3.4.18.2 | Zinc-coordinating DNA-binding domains | C2H2 zinc finger factors | Multiple dispersed zinc fingers | ZNF518 | 12 | 1.34 | 50.99 | ZNF518B.FL@HTS.Lys@boozy-harlequin-rattlesnake+shaggy-cornflower-fossa+cheeky-salmon-setter+boozy-cerise-mau@faltejsk.ProBound@motif\_without\_ns.ppm |
| SALL3 | CHS | train | THC\_0314.Rep-DIANA\_0293 |  |  | FL | 2.3.4.3.3 | Zinc-coordinating DNA-binding domains | C2H2 zinc finger factors | Multiple dispersed zinc fingers | SAL-like | 19 | 17.32 | 13.02 | SALL3.FL@AFS.Lys@snappy-denim-hound@autosome-ru.ChIPMunk@998seq\_21to7\_m0.pcm |
| SALL3 | CHS | test | THC\_0314.Rep-MICHELLE\_0314 |  |  | FL | 2.3.4.3.3 | Zinc-coordinating DNA-binding domains | C2H2 zinc finger factors | Multiple dispersed zinc fingers | SAL-like | 19 | 17.32 | 13.02 | SALL3.FL@AFS.Lys@snappy-denim-hound@autosome-ru.ChIPMunk@998seq\_21to7\_m0.pcm |
| SALL3 | GHTS.IVT | train | YWE\_B\_AffSeq\_B02\_SALL3 |  |  | DBD | 2.3.4.3.3 | Zinc-coordinating DNA-binding domains | C2H2 zinc finger factors | Multiple dispersed zinc fingers | SAL-like | 15 | 7.73 | 31.44 | SALL3.DBD@HTS.IVT@trippy-tan-dollar+freaky-aquamarine-gerbil+muzzy-black-avocet+scaly-chestnut-coyote@Halle.Dimont@Motif\_1\_sampled\_e1\_5\_astrained.ppm |
| SALL3 | GHTS.IVT | train | YWF\_B\_AffSeq\_B02\_SALL3 |  |  | DBD | 2.3.4.3.3 | Zinc-coordinating DNA-binding domains | C2H2 zinc finger factors | Multiple dispersed zinc fingers | SAL-like | 13 | 17.82 | 15.47 | SALL3.DBD@AFS.IVT@lovely-chartreuse-cat@autosome-ru.ChIPMunk@343seq\_21to7\_m0.pcm |
| SALL3 | GHTS.GFPIVT | train | YWQ\_B\_AffSeq\_C6\_SALL3-DBD1 |  |  | FL | 2.3.4.3.3 | Zinc-coordinating DNA-binding domains | C2H2 zinc finger factors | Multiple dispersed zinc fingers | SAL-like | 18 | 18.65 | 19.9 | SALL3.FL@HTS.Lys@scaly-sepia-quail+skanky-silver-fly+whiny-eggplant-reindeer+wimpy-khaki-lynx@autosome-ru.ChIPMunk@topk\_cycle=C1+C2+C3+C4\_k=5\_top=2500.pcm |
| SALL3 | GHTS.GFPIVT | train | YWP\_B\_AffSeq\_B10\_SALL3-DBD2 |  |  | DBD | 2.3.4.3.3 | Zinc-coordinating DNA-binding domains | C2H2 zinc finger factors | Multiple dispersed zinc fingers | SAL-like | 11 | 16.96 | 13.29 | SALL3.DBD@AFS.IVT@lovely-chartreuse-cat@HughesLab.MEME@346\_fa\_memechip\_Motif1.ppm |
| SALL3 | GHTS.Lys | test | YWK\_B\_AffSeq\_H6\_SALL3 |  |  | FL | 2.3.4.3.3 | Zinc-coordinating DNA-binding domains | C2H2 zinc finger factors | Multiple dispersed zinc fingers | SAL-like | 19 | 16.66 | 14.89 | SALL3.FL@AFS.Lys@lumpy-gamboge-chamois@autosome-ru.ChIPMunk@988seq\_21to7\_m0.pcm |
| SALL3 | GHTS.Lys | train | YWK\_B\_AffSeq\_A1\_SALL3 |  |  | FL | 2.3.4.3.3 | Zinc-coordinating DNA-binding domains | C2H2 zinc finger factors | Multiple dispersed zinc fingers | SAL-like | 17 | 16.65 | 19.59 | SALL3.FL@HTS.Lys@breezy-olive-caiman+bumpy-wheat-mongoose@autosome-ru.ChIPMunk@topk\_cycle=C3+C4\_k=5\_top=10000.pcm |
| SALL3 | GHTS.Lys | train | YWK\_D\_AffSeq\_H6\_SALL3 |  |  | FL | 2.3.4.3.3 | Zinc-coordinating DNA-binding domains | C2H2 zinc finger factors | Multiple dispersed zinc fingers | SAL-like | 19 | 17.32 | 13.02 | SALL3.FL@AFS.Lys@snappy-denim-hound@autosome-ru.ChIPMunk@998seq\_21to7\_m0.pcm |
| SALL3 | GHTS.Lys | train | YWK\_D\_AffSeq\_A1\_SALL3 |  |  | FL | 2.3.4.3.3 | Zinc-coordinating DNA-binding domains | C2H2 zinc finger factors | Multiple dispersed zinc fingers | SAL-like | 19 | 17.51 | 20.72 | SALL3.FL@HTS.Lys@breezy-olive-caiman+bumpy-wheat-mongoose+sleazy-flax-jaguar+surly-ultramarine-jackal@autosome-ru.ChIPMunk@topk\_cycle=C1+C2+C3+C4\_k=5\_top=2500.pcm |
| SALL3 | HTS.IVT | test | YWF\_A\_AG40NACCATA |  |  | DBD | 2.3.4.3.3 | Zinc-coordinating DNA-binding domains | C2H2 zinc finger factors | Multiple dispersed zinc fingers | SAL-like | 12 | 2.59 | 40.26 | SALL3.DBD@HTS.IVT@ugly-corn-jaguar+hilly-apricot-dragonfly+trippy-amaranth-bonobo+gimpy-bistre-akita@faltejsk.ProBound@motif\_without\_ns.ppm |
| SALL3 | HTS.IVT | test | YWE\_A\_AG40NACCATA |  |  | DBD | 2.3.4.3.3 | Zinc-coordinating DNA-binding domains | C2H2 zinc finger factors | Multiple dispersed zinc fingers | SAL-like | 12 | 2.52 | 46.39 | SALL3.DBD@AFS.GFPIVT@sickly-taupe-kakapo+jumpy-aqua-cockroach+scummy-cyan-octopus@faltejsk.ProBound@motif\_without\_ns.ppm |
| SALL3 | HTS.GFPIVT | test | YWP\_A\_AT40NTCTGTC |  |  | DBD | 2.3.4.3.3 | Zinc-coordinating DNA-binding domains | C2H2 zinc finger factors | Multiple dispersed zinc fingers | SAL-like | 15 | 19.83 | 20.13 | SALL3.DBD@HTS.GFPIVT@paltry-ruby-quoll@HughesLab.MEME@topk\_cycle=C3\_k=5\_top=10000\_Motif1.ppm |
| SALL3 | HTS.GFPIVT | test | YWQ\_A\_CA40NGCCTTT |  |  | DBD | 2.3.4.3.3 | Zinc-coordinating DNA-binding domains | C2H2 zinc finger factors | Multiple dispersed zinc fingers | SAL-like | 15 | 16.6 | 16.15 | SALL3.DBD@HTS.GFPIVT@cozy-chartreuse-cassowary@AJolma.Autoseed@Multinom2\_Onehit\_Seed\_ATATKAWWTMATAWBackgroundCyc1.ppm |
| SALL3 | HTS.Lys | test | YWK\_C\_TG40NTGTGTT |  |  | FL | 2.3.4.3.3 | Zinc-coordinating DNA-binding domains | C2H2 zinc finger factors | Multiple dispersed zinc fingers | SAL-like | 12 | 2.44 | 44.69 | SALL3.FL@HTS.Lys@breezy-olive-caiman+sleazy-flax-jaguar+bumpy-wheat-mongoose+surly-ultramarine-jackal@faltejsk.ProBound@motif\_without\_ns.ppm |
| SALL3 | HTS.Lys | test | YWK\_A\_TG40NTGTGTT |  |  | FL | 2.3.4.3.3 | Zinc-coordinating DNA-binding domains | C2H2 zinc finger factors | Multiple dispersed zinc fingers | SAL-like | 12 | 2.44 | 44.69 | SALL3.FL@HTS.Lys@breezy-olive-caiman+sleazy-flax-jaguar+bumpy-wheat-mongoose+surly-ultramarine-jackal@faltejsk.ProBound@motif\_without\_ns.ppm |
| SALL3 | HTS.Lys | test | YWK\_A\_AA40NTATCGC |  |  | FL | 2.3.4.3.3 | Zinc-coordinating DNA-binding domains | C2H2 zinc finger factors | Multiple dispersed zinc fingers | SAL-like | 12 | 2.44 | 44.69 | SALL3.FL@HTS.Lys@breezy-olive-caiman+sleazy-flax-jaguar+bumpy-wheat-mongoose+surly-ultramarine-jackal@faltejsk.ProBound@motif\_without\_ns.ppm |
| SALL3 | HTS.Lys | test | YWK\_C\_AA40NTATCGC |  |  | FL | 2.3.4.3.3 | Zinc-coordinating DNA-binding domains | C2H2 zinc finger factors | Multiple dispersed zinc fingers | SAL-like | 12 | 2.44 | 44.69 | SALL3.FL@HTS.Lys@breezy-olive-caiman+sleazy-flax-jaguar+bumpy-wheat-mongoose+surly-ultramarine-jackal@faltejsk.ProBound@motif\_without\_ns.ppm |
| ZBED2 | CHS | train | THC\_0902 |  |  | FL | 2.3.5.0.2 | Zinc-coordinating DNA-binding domains | C2H2 zinc finger factors | BED zinc finger |  | 12 | 7.15 | 51.26 | ZBED2.FL@HTS.Lys@whiny-emerald-wallaby+paltry-amethyst-zebu+tacky-violet-quoll+muggy-russet-neanderthal@faltejsk.ProBound@motif\_with\_ns.ppm |
| ZBED2 | CHS | test | THC\_0101 |  |  | FL | 2.3.5.0.2 | Zinc-coordinating DNA-binding domains | C2H2 zinc finger factors | BED zinc finger |  | 9 | 10.43 | 52.28 | ZBED2.FL@SMS@muzzy-teal-armadillo@HughesLab.MEME@topk\_k=5\_top=2500\_Motif1\_min3max30.ppm |
| ZBED2 | GHTS.GFPIVT | train | YWM\_B\_AffSeq\_H11\_ZBED2-FL |  |  | FL | 2.3.5.0.2 | Zinc-coordinating DNA-binding domains | C2H2 zinc finger factors | BED zinc finger |  | 20 | 8.32 | 53.43 | ZBED2.FL@AFS.Lys@homey-copper-ferret+sleazy-coral-bombay+sleazy-salmon-maltese@Halle.Dimont@Motif\_1\_w20\_astrained.ppm |
| ZBED2 | GHTS.Lys | train | YWN\_B\_AffSeq\_G11\_ZBED2-FL |  |  | FL | 2.3.5.0.2 | Zinc-coordinating DNA-binding domains | C2H2 zinc finger factors | BED zinc finger |  | 20 | 8.32 | 53.43 | ZBED2.FL@AFS.Lys@homey-copper-ferret+sleazy-coral-bombay+sleazy-salmon-maltese@Halle.Dimont@Motif\_1\_w20\_astrained.ppm |
| ZBED2 | GHTS.Lys | test | YWO\_B\_AffSeq\_G11\_ZBED2-FL |  |  | FL | 2.3.5.0.2 | Zinc-coordinating DNA-binding domains | C2H2 zinc finger factors | BED zinc finger |  | 19 | 7.22 | 51.28 | ZBED2.FL@HTS.Lys@greasy-green-rabbit+stuffy-chartreuse-wolfhound+greasy-ultramarine-capybara+tasty-salmon-owl@Halle.Dimont@Motif\_1\_dtrue\_htsversion.ppm |
| ZBED2 | PBM | test | PBM13709 |  |  | FL | 2.3.5.0.2 | Zinc-coordinating DNA-binding domains | C2H2 zinc finger factors | BED zinc finger |  | 14 | 7.86 | 52.98 | ZBED2.FL@HTS.GFPIVT@squeaky-tangerine-crocodile+surly-heliotrope-lemming+goopy-champagne-liger+smelly-periwinkle-wolverine@Halle.Dimont@Motif\_1\_dtrue\_htsversion.ppm |
| ZBED2 | HTS.GFPIVT | test | YWM\_A\_TT40NTAGCAG |  |  | FL | 2.3.5.0.2 | Zinc-coordinating DNA-binding domains | C2H2 zinc finger factors | BED zinc finger |  | 12 | 4.84 | 49.44 | ZBED2.FL@SMS@silly-mauve-iguana@faltejsk.ProBound@motif\_without\_ns.ppm |
| ZBED2 | HTS.Lys | test | YWN\_A\_TC40NACGGGA |  |  | FL | 2.3.5.0.2 | Zinc-coordinating DNA-binding domains | C2H2 zinc finger factors | BED zinc finger |  | 12 | 4.84 | 49.44 | ZBED2.FL@SMS@silly-mauve-iguana@faltejsk.ProBound@motif\_without\_ns.ppm |
| ZBED2 | HTS.Lys | test | YWO\_A\_TC40NACGGGA |  |  | FL | 2.3.5.0.2 | Zinc-coordinating DNA-binding domains | C2H2 zinc finger factors | BED zinc finger |  | 12 | 4.84 | 49.44 | ZBED2.FL@SMS@silly-mauve-iguana@faltejsk.ProBound@motif\_without\_ns.ppm |
| ZBED2 | SMS | test | UT380-242-2 |  |  | FL | 2.3.5.0.2 | Zinc-coordinating DNA-binding domains | C2H2 zinc finger factors | BED zinc finger |  | 7 | 5.22 | 53.13 | ZBED2.FL@CHS@jumpy-viridian-crocodile@Halle.Dimont@Motif\_1\_imw20\_astrained.ppm |
| ZBED2 | SMS | test | UT380-242 |  |  | FL | 2.3.5.0.2 | Zinc-coordinating DNA-binding domains | C2H2 zinc finger factors | BED zinc finger |  | 9 | 8.8 | 49.67 | ZBED2.FL@SMS@silly-mauve-iguana@AJolma.Autoseed@Multinom1\_OneHitAllReads\_Seed\_NCGAAACYN.ppm |
| ZBED5 | CHS | test | THC\_0081 |  |  | DBD | 2.3.5.0.5 | Zinc-coordinating DNA-binding domains | C2H2 zinc finger factors | BED zinc finger |  | 16 | 14.3 | 71.5 | ZBED5.DBD@AFS.GFPIVT@wiggy-cyan-kudu@HughesLab.Streme@500\_fa\_streme\_minw3\_maxw30\_Motif1.ppm |
| ZBED5 | CHS | train | THC\_0278 |  |  | DBD | 2.3.5.0.5 | Zinc-coordinating DNA-binding domains | C2H2 zinc finger factors | BED zinc finger |  | 16 | 14.3 | 71.5 | ZBED5.DBD@AFS.GFPIVT@wiggy-cyan-kudu@HughesLab.Streme@500\_fa\_streme\_minw3\_maxw30\_Motif1.ppm |
| ZBED5 | GHTS.GFPIVT | test | YWM\_B\_AffSeq\_C12\_ZBED5-DBD |  |  | DBD | 2.3.5.0.5 | Zinc-coordinating DNA-binding domains | C2H2 zinc finger factors | BED zinc finger |  | 20 | 8.37 | 63.36 | ZBED5.DBD@AFS.GFPIVT@wiggy-cyan-kudu@Halle.Dimont@Motif\_1\_w20\_astrained.ppm |
| ZBED5 | GHTS.Lys | train | YWO\_B\_AffSeq\_A1\_ZBED5-FL |  |  | FL | 2.3.5.0.5 | Zinc-coordinating DNA-binding domains | C2H2 zinc finger factors | BED zinc finger |  | 20 | 7.86 | 62.81 | ZBED5.FL@AFS.Lys@gamy-ochre-millipede@Halle.Dimont@Motif\_1\_w20\_astrained.ppm |
| ZBED5 | PBM | test | PBM13711 |  |  | DBD | 2.3.5.0.5 | Zinc-coordinating DNA-binding domains | C2H2 zinc finger factors | BED zinc finger |  | 10 | 9.42 | 64.21 | ZBED5.DBD@PBM.ME@scaly-russet-donkey@autosome-ru.ChIPMunk@s\_6-16\_flat.pcm |
| ZBED5 | PBM | test | PBM13711 |  |  | DBD | 2.3.5.0.5 | Zinc-coordinating DNA-binding domains | C2H2 zinc finger factors | BED zinc finger |  | 12 | 15.52 | 71.22 | ZBED5.DBD@HTS.IVT@lovely-grey-worm@AJolma.Autoseed@Cyc2\_MultiNom1\_SeedNNRCGGAACCCC.ppm |
| ZBED5 | HTS.IVT | test | YWC\_A\_GA40NTGTATC |  |  | DBD | 2.3.5.0.5 | Zinc-coordinating DNA-binding domains | C2H2 zinc finger factors | BED zinc finger |  | 12 | 1.85 | 52.48 | ZBED5.DBD@HTS.IVT@slimy-asparagus-newt+ugly-scarlet-gorilla+lovely-grey-worm@faltejsk.ProBound@motif\_without\_ns.ppm |
| ZBED5 | HTS.GFPIVT | test | YWQ\_A\_CG40NGTTATT |  |  | FL | 2.3.5.0.5 | Zinc-coordinating DNA-binding domains | C2H2 zinc finger factors | BED zinc finger |  | 11 | 9.68 | 66.19 | ZBED5.FL@HTS.GFPIVT@cloudy-rust-numbat+grumpy-platinum-newt+smelly-red-squirt@HughesLab.Homer@topk\_cycle=C1+C2+C3\_k=5\_top=2500\_fasta\_homer\_minw3\_maxw\_40\_Motif1.ppm |
| ZBED5 | HTS.GFPIVT | test | YWM\_A\_CG40NAATAGC |  |  | DBD | 2.3.5.0.5 | Zinc-coordinating DNA-binding domains | C2H2 zinc finger factors | BED zinc finger |  | 12 | 1.85 | 52.48 | ZBED5.DBD@HTS.IVT@slimy-asparagus-newt+ugly-scarlet-gorilla+lovely-grey-worm@faltejsk.ProBound@motif\_without\_ns.ppm |
| ZBED5 | HTS.Lys | test | YWO\_A\_AA40NTATCGC |  |  | FL | 2.3.5.0.5 | Zinc-coordinating DNA-binding domains | C2H2 zinc finger factors | BED zinc finger |  | 16 | 7.19 | 58.88 | ZBED5.FL@HTS.Lys@hazy-seashell-guppy+homely-scarlet-insect+cheeky-magnolia-stoat+gimpy-thistle-tapir@Halle.Dimont@Motif\_1\_sampled\_e4\_astrained.ppm |
| ZBED5 | HTS.Lys | test | YWN\_A\_AA40NTATCGC |  |  | FL | 2.3.5.0.5 | Zinc-coordinating DNA-binding domains | C2H2 zinc finger factors | BED zinc finger |  | 11 | 10.22 | 68.45 | ZBED5.FL@HTS.Lys@cozy-olive-ragdoll+frumpy-beige-dachshund+nerdy-olivine-forest+snazzy-mustard-dog@HughesLab.Homer@topk\_cycle=C1+C2+C3+C4\_k=5\_top=10000\_fasta\_homer\_minw3\_maxw\_40\_Motif1.ppm |
| ZBED5 | SMS | test | UT380-245-2 |  |  | DBD | 2.3.5.0.5 | Zinc-coordinating DNA-binding domains | C2H2 zinc finger factors | BED zinc finger |  | 20 | 8.37 | 63.36 | ZBED5.DBD@AFS.GFPIVT@wiggy-cyan-kudu@Halle.Dimont@Motif\_1\_w20\_astrained.ppm |
| ZBED5 | SMS | test | UT380-245-3 |  |  | DBD | 2.3.5.0.5 | Zinc-coordinating DNA-binding domains | C2H2 zinc finger factors | BED zinc finger |  | 11 | 11.1 | 72.25 | ZBED5.DBD@AFS.GFPIVT@wiggy-cyan-kudu@HughesLab.Homer@500\_fa\_homer\_minw3\_maxw\_40\_Motif1.ppm |
| LEUTX | CHS | train | THC\_0455.Rep-DIANA\_0293 |  |  | FL | 3.1.3.13.1 | Helix-turn-helix domains | Homeo domain factors | Paired-related HD | LEUTX | 8 | 8.73 | 37.26 | LEUTX.FL@PBM.ME@sickly-pink-quoll@autosome-ru.ChIPMunk@s\_6-16\_flat.pcm |
| LEUTX | CHS | train | THC\_0312.Rep-DIANA\_0293 |  |  | FL | 3.1.3.13.1 | Helix-turn-helix domains | Homeo domain factors | Paired-related HD | LEUTX | 10 | 7.56 | 39.06 | LEUTX.FL@HTS.GFPIVT@scaly-carmine-setter+geeky-plum-weasel+ugly-maroon-okapi@Halle.Dimont@Motif\_1\_sampled\_e4\_astrained.ppm |
| LEUTX | CHS | train | THC\_0312.Rep-MICHELLE\_0314 |  |  | FL | 3.1.3.13.1 | Helix-turn-helix domains | Homeo domain factors | Paired-related HD | LEUTX | 10 | 7.56 | 39.06 | LEUTX.FL@HTS.GFPIVT@scaly-carmine-setter+geeky-plum-weasel+ugly-maroon-okapi@Halle.Dimont@Motif\_1\_sampled\_e4\_astrained.ppm |
| LEUTX | CHS | test | THC\_0455.Rep-MICHELLE\_0314 |  |  | FL | 3.1.3.13.1 | Helix-turn-helix domains | Homeo domain factors | Paired-related HD | LEUTX | 8 | 8.22 | 35.37 | LEUTX.FL@PBM.ME@lanky-aquamarine-echidna@autosome-ru.ChIPMunk@s\_6-16\_flat.pcm |
| LEUTX | GHTS.GFPIVT | train | YWP\_B\_AffSeq\_H3\_LEUTX-FL |  |  | FL | 3.1.3.13.1 | Helix-turn-helix domains | Homeo domain factors | Paired-related HD | LEUTX | 20 | 9.31 | 49.46 | LEUTX.FL@AFS.GFPIVT@tacky-champagne-lemur@Halle.Dimont@Motif\_1\_w20\_astrained.ppm |
| LEUTX | GHTS.Lys | train | YWK\_D\_AffSeq\_B7\_LEUTX |  |  | FL | 3.1.3.13.1 | Helix-turn-helix domains | Homeo domain factors | Paired-related HD | LEUTX | 10 | 5.83 | 38.88 | LEUTX.FL@HTS.Lys@ready-cerulean-collie+beady-plum-cichlid+messy-flax-raccoon+scanty-orchid-scorpion@Halle.Dimont@Motif\_1\_dtrue\_htsversion.ppm |
| LEUTX | GHTS.Lys | test | YWK\_B\_AffSeq\_B7\_LEUTX |  |  | FL | 3.1.3.13.1 | Helix-turn-helix domains | Homeo domain factors | Paired-related HD | LEUTX | 20 | 9.31 | 49.46 | LEUTX.FL@AFS.GFPIVT@tacky-champagne-lemur@Halle.Dimont@Motif\_1\_w20\_astrained.ppm |
| LEUTX | PBM | test | PBM13720 |  |  | FL | 3.1.3.13.1 | Helix-turn-helix domains | Homeo domain factors | Paired-related HD | LEUTX | 12 | 4.69 | 40.18 | LEUTX.FL@AFS.Lys@craggy-rust-sponge+muzzy-chartreuse-dane+dorky-asparagus-fowl+blurry-cerulean-wolfhound@faltejsk.ProBound@motif\_with\_ns.ppm |
| LEUTX | PBM | test | PBM14323 |  |  | FL | 3.1.3.13.1 | Helix-turn-helix domains | Homeo domain factors | Paired-related HD | LEUTX | 10 | 5.13 | 36.36 | LEUTX.FL@HTS.Lys@cloudy-wisteria-catfish+wiggy-gamboge-warthog+lumpy-rust-blue+skimpy-russet-dollar@Halle.Dimont@Motif\_1\_dtrue\_htsversion.ppm |
| LEUTX | HTS.GFPIVT | test | YWP\_A\_TG40NTGATCT |  |  | FL | 3.1.3.13.1 | Helix-turn-helix domains | Homeo domain factors | Paired-related HD | LEUTX | 12 | 4.33 | 39.5 | LEUTX.FL@HTS.Lys@messy-flax-raccoon+beady-plum-cichlid+ready-cerulean-collie+scanty-orchid-scorpion@faltejsk.ProBound@motif\_with\_ns.ppm |
| LEUTX | HTS.Lys | test | YWK\_A\_AT40NGAGAGG |  |  | FL | 3.1.3.13.1 | Helix-turn-helix domains | Homeo domain factors | Paired-related HD | LEUTX | 10 | 7.56 | 39.06 | LEUTX.FL@HTS.GFPIVT@scaly-carmine-setter+geeky-plum-weasel+ugly-maroon-okapi@Halle.Dimont@Motif\_1\_sampled\_e4\_astrained.ppm |
| LEUTX | HTS.Lys | test | YWK\_C\_AT40NGAGAGG |  |  | FL | 3.1.3.13.1 | Helix-turn-helix domains | Homeo domain factors | Paired-related HD | LEUTX | 12 | 4.33 | 39.5 | LEUTX.FL@HTS.Lys@messy-flax-raccoon+beady-plum-cichlid+ready-cerulean-collie+scanty-orchid-scorpion@faltejsk.ProBound@motif\_with\_ns.ppm |
