## Supplementary Data SD1 for "Inferring binding specificities of human transcription factors with the wisdom of crowds": ibis_motifs_Leaderboard_A2G.html

 

| TF | Platform | Slice | Replicate ID | logo-direct | logo-revcomp | Construct type | tfclass:id | tfclass:superclass | tfclass:class | tfclass:family | tfclass:subfamily | Motif length | Information content | GC% | Motif |
| --- | --- | --- | --- | --- | --- | --- | --- | --- | --- | --- | --- | --- | --- | --- | --- |
| RORB | CHS | test | THC\_0813 |  |  | NA | 2.1.2.6.2 | Zinc-coordinating DNA-binding domains | Nuclear receptors with C4 zinc fingers | Thyroid hormone receptor-related | ROR (NR1F) | 20 | 8.5 | 54.42 | RORB.NA@AFS.Lys@surly-brown-serval@Halle.Dimont@Motif\_1\_w20\_astrained.ppm |
| RORB | GHTS.IVT | test | YWF\_B\_AffSeq\_C12\_RORB |  |  | NA | 2.1.2.6.2 | Zinc-coordinating DNA-binding domains | Nuclear receptors with C4 zinc fingers | Thyroid hormone receptor-related | ROR (NR1F) | 11 | 14.36 | 40.19 | RORB.NA@HTS.Lys@droopy-flax-rat+sleazy-champagne-octopus+squeaky-alizarin-rattlesnake+tacky-charcoal-donkey@HughesLab.MEME@topk\_cycle=C1+C2+C3+C4\_k=5\_top=10000\_Motif1.ppm |
| RORB | PBM | test | PBM14353 |  |  | NA | 2.1.2.6.2 | Zinc-coordinating DNA-binding domains | Nuclear receptors with C4 zinc fingers | Thyroid hormone receptor-related | ROR (NR1F) | 12 | 14.95 | 40.13 | RORB.NA@HTS.Lys@droopy-flax-rat+sleazy-champagne-octopus+squeaky-alizarin-rattlesnake+tacky-charcoal-donkey@HughesLab.MEME@topk\_cycle=C1+C2+C3+C4\_k=5\_top=10000\_Motif1\_min3max30.ppm |
| RORB | PBM | test | PBM14353 |  |  | NA | 2.1.2.6.2 | Zinc-coordinating DNA-binding domains | Nuclear receptors with C4 zinc fingers | Thyroid hormone receptor-related | ROR (NR1F) | 10 | 13.48 | 41.47 | RORB.NA@HTS.Lys@droopy-flax-rat+sleazy-champagne-octopus+squeaky-alizarin-rattlesnake+tacky-charcoal-donkey@HughesLab.Homer@topk\_cycle=C1+C2+C3+C4\_k=5\_top=10000\_fasta\_homer\_minw3\_maxw\_40\_Motif1.ppm |
| RORB | PBM | train | PBM14337 |  |  | NA | 2.1.2.6.2 | Zinc-coordinating DNA-binding domains | Nuclear receptors with C4 zinc fingers | Thyroid hormone receptor-related | ROR (NR1F) | 12 | 14.95 | 40.13 | RORB.NA@HTS.Lys@droopy-flax-rat+sleazy-champagne-octopus+squeaky-alizarin-rattlesnake+tacky-charcoal-donkey@HughesLab.MEME@topk\_cycle=C1+C2+C3+C4\_k=5\_top=10000\_Motif1\_min3max30.ppm |
| RORB | HTS.IVT | test | YWF\_A\_CG40NAATAGC |  |  | NA | 2.1.2.6.2 | Zinc-coordinating DNA-binding domains | Nuclear receptors with C4 zinc fingers | Thyroid hormone receptor-related | ROR (NR1F) | 11 | 14.36 | 40.19 | RORB.NA@HTS.Lys@droopy-flax-rat+sleazy-champagne-octopus+squeaky-alizarin-rattlesnake+tacky-charcoal-donkey@HughesLab.MEME@topk\_cycle=C1+C2+C3+C4\_k=5\_top=10000\_Motif1.ppm |
| RORB | HTS.Lys | train | YWL\_A\_CA40NGCCTTT |  |  | NA | 2.1.2.6.2 | Zinc-coordinating DNA-binding domains | Nuclear receptors with C4 zinc fingers | Thyroid hormone receptor-related | ROR (NR1F) | 12 | 2.87 | 53.85 | RORB.NA@HTS.Lys@sleazy-champagne-octopus+droopy-flax-rat+tacky-charcoal-donkey+squeaky-alizarin-rattlesnake@faltejsk.ProBound@motif\_with\_ns.ppm |
| RORB | SMS | train | SRR3405080 |  |  | NA | 2.1.2.6.2 | Zinc-coordinating DNA-binding domains | Nuclear receptors with C4 zinc fingers | Thyroid hormone receptor-related | ROR (NR1F) | 12 | 4.86 | 49.82 | RORB.NA@PBM.HK@flimsy-scarlet-frigatebird@Halle.Dimont@Motif\_1\_w0\_01\_bg4\_imw15\_astrained.ppm |
| TIGD3 | CHS | test | THC\_0146 |  |  | FL | 3.0.256.0.2 | Helix-turn-helix domains | Other | CENPB |  | 15 | 11.32 | 37.05 | TIGD3.FL@CHS@sunny-asparagus-boar@autosome-ru.ChIPMunk@999seq\_21to7\_m0.pcm |
| TIGD3 | GHTS.Lys | test | YWO\_B\_AffSeq\_E11\_TIGD3-FL |  |  | FL | 3.0.256.0.2 | Helix-turn-helix domains | Other | CENPB |  | 14 | 19.75 | 44.91 | TIGD3.FL@HTS.GFPIVT@bumpy-lilac-mandrill+sleazy-smalt-dachsbracke+whiny-xanthic-bear@HughesLab.MEME@topk\_cycle=C1+C2+C3\_k=5\_top=10000\_Motif1.ppm |
| TIGD3 | PBM | test | PBM13715 |  |  | DBD | 3.0.256.0.2 | Helix-turn-helix domains | Other | CENPB |  | 7 | 5.95 | 32.85 | TIGD3.DBD@PBM.ME@scummy-dandelion-retriever@Halle.Dimont@Motif\_1\_astrained.ppm |
| TIGD3 | PBM | test | PBM13933 |  |  | DBD | 3.0.256.0.2 | Helix-turn-helix domains | Other | CENPB |  | 14 | 3.61 | 43.49 | TIGD3.DBD@PBM.HK@stuffy-gold-jackal@Halle.Dimont@Motif\_1\_w0\_01\_bg4\_imw15\_astrained.ppm |
| TIGD3 | PBM | train | PBM13699 |  |  | DBD | 3.0.256.0.2 | Helix-turn-helix domains | Other | CENPB |  | 7 | 5.95 | 32.85 | TIGD3.DBD@PBM.ME@scummy-dandelion-retriever@Halle.Dimont@Motif\_1\_astrained.ppm |
| TIGD3 | HTS.GFPIVT | test | YWQ\_A\_AA40NAGTGGT |  |  | FL | 3.0.256.0.2 | Helix-turn-helix domains | Other | CENPB |  | 20 | 8.61 | 49.57 | TIGD3.FL@AFS.Lys@randy-tan-octopus@Halle.Dimont@Motif\_1\_w20\_astrained.ppm |
| TIGD3 | HTS.Lys | train | YWN\_A\_GC40NCACTTC |  |  | FL | 3.0.256.0.2 | Helix-turn-helix domains | Other | CENPB |  | 12 | 2.82 | 49.04 | TIGD3.FL@HTS.Lys@slimy-thistle-llama+snippy-lemon-bee+leaky-cardinal-shrew+skanky-puce-serval@faltejsk.ProBound@motif\_without\_ns.ppm |
| TIGD3 | HTS.Lys | test | YWO\_A\_GC40NCACTTC |  |  | FL | 3.0.256.0.2 | Helix-turn-helix domains | Other | CENPB |  | 12 | 2.82 | 49.04 | TIGD3.FL@HTS.Lys@slimy-thistle-llama+snippy-lemon-bee+leaky-cardinal-shrew+skanky-puce-serval@faltejsk.ProBound@motif\_without\_ns.ppm |
| TIGD3 | SMS | train | UT380-225 |  |  | FL | 3.0.256.0.2 | Helix-turn-helix domains | Other | CENPB |  | 14 | 19.75 | 44.91 | TIGD3.FL@HTS.GFPIVT@bumpy-lilac-mandrill+sleazy-smalt-dachsbracke+whiny-xanthic-bear@HughesLab.MEME@topk\_cycle=C1+C2+C3\_k=5\_top=10000\_Motif1.ppm |
| NACC2 | CHS | test | THC\_0171 |  |  | DBD | 3.0.257.0.1 | Helix-turn-helix domains | Other | BEN |  | 20 | 6.8 | 54.27 | NACC2.DBD@AFS.GFPIVT@frumpy-heliotrope-walrus@Halle.Dimont@Motif\_1\_w20\_astrained.ppm |
| NACC2 | GHTS.GFPIVT | test | YWM\_B\_AffSeq\_A11\_NACC2-DBD |  |  | DBD | 3.0.257.0.1 | Helix-turn-helix domains | Other | BEN |  | 20 | 7.21 | 52.51 | NACC2.DBD@AFS.GFPIVT@frumpy-heliotrope-walrus+wimpy-copper-capybara+paltry-saffron-mongrel+ready-peach-jaguar@Halle.Dimont@Motif\_1\_w20\_astrained.ppm |
| NACC2 | PBM | test | PBM13741 |  |  | DBD | 3.0.257.0.1 | Helix-turn-helix domains | Other | BEN |  | 10 | 8.36 | 40.53 | NACC2.DBD@PBM.HK@foggy-cornflower-wolfhound@autosome-ru.ChIPMunk@s\_6-16\_flat.pcm |
| NACC2 | PBM | train | PBM13725 |  |  | DBD | 3.0.257.0.1 | Helix-turn-helix domains | Other | BEN |  | 26 | 12.45 | 44.87 | NACC2.DBD@PBM.ME@beady-sangria-ant@OFornes.ExplaiNN@filter35\_1.ppm |
| NACC2 | HTS.IVT | train | YWC\_B\_TG40NTGTGTT |  |  | DBD | 3.0.257.0.1 | Helix-turn-helix domains | Other | BEN |  | 12 | 2.43 | 46.36 | NACC2.DBD@SMS@breezy-cinnabar-shrimp@faltejsk.ProBound@motif\_without\_ns.ppm |
| NACC2 | HTS.IVT | test | YWF\_A\_GC40NCTTGTA |  |  | FL | 3.0.257.0.1 | Helix-turn-helix domains | Other | BEN |  | 7 | 3.32 | 44.65 | NACC2.FL@HTS.IVT@zippy-tan-lionfish+cloudy-mustard-okapi+beady-periwinkle-akbash+dorky-zucchini-dollar@Halle.Dimont@Motif\_2\_sampled\_e1\_5\_astrained.ppm |
| NACC2 | HTS.GFPIVT | test | YWM\_A\_AC40NTGTTAC |  |  | DBD | 3.0.257.0.1 | Helix-turn-helix domains | Other | BEN |  | 15 | 20.92 | 32.09 | NACC2.DBD@HTS.GFPIVT@boozy-puce-binturong+flimsy-amber-mouse@HughesLab.Streme@topk\_cycle=C3+C4\_k=5\_top=2500\_Motif1.ppm |
| NACC2 | HTS.Lys | train | YWD\_A\_GA40NGGTCAT |  |  | DBD | 3.0.257.0.1 | Helix-turn-helix domains | Other | BEN |  | 12 | 2.43 | 46.36 | NACC2.DBD@SMS@breezy-cinnabar-shrimp@faltejsk.ProBound@motif\_without\_ns.ppm |
| NACC2 | SMS | train | UT380-127-3 |  |  | DBD | 3.0.257.0.1 | Helix-turn-helix domains | Other | BEN |  | 22 | 23.16 | 38.96 | NACC2.DBD@HTS.GFPIVT@boozy-puce-binturong+cozy-dandelion-cichlid+flimsy-amber-mouse+gloppy-azure-chamois@autosome-ru.ChIPMunk@topk\_cycle=C1+C2+C3+C4\_k=5\_top=10000.pcm |
| NACC2 | SMS | test | UT380-127-2 |  |  | DBD | 3.0.257.0.1 | Helix-turn-helix domains | Other | BEN |  | 9 | 6.25 | 43.99 | NACC2.DBD@PBM.HK@squeaky-champagne-vole@Halle.Dimont@Motif\_1\_astrained.ppm |
| LEF1 | CHS | test | THC\_0722 |  |  | NA | 4.1.3.0.4 | Other all-alpha-helical DNA-binding domains | High-mobility group (HMG) domain factors | TCF7-related |  | 20 | 8.29 | 49.77 | LEF1.NA@CHS@beady-olivine-goat@Halle.Dimont@Motif\_1\_w20\_astrained.ppm |
| LEF1 | GHTS.Lys | test | YWL\_B\_AffSeq\_B9\_LEF1 |  |  | NA | 4.1.3.0.4 | Other all-alpha-helical DNA-binding domains | High-mobility group (HMG) domain factors | TCF7-related |  | 20 | 9.33 | 47.23 | LEF1.NA@AFS.Lys@flabby-sepia-audemer@Halle.Dimont@Motif\_1\_w20\_astrained.ppm |
| LEF1 | PBM | test | PBM14349 |  |  | NA | 4.1.3.0.4 | Other all-alpha-helical DNA-binding domains | High-mobility group (HMG) domain factors | TCF7-related |  | 15 | 18.43 | 39.22 | LEF1.NA@AFS.Lys@flabby-sepia-audemer@HughesLab.Streme@1000\_fa\_streme\_Motif1.ppm |
| LEF1 | PBM | train | PBM14333 |  |  | NA | 4.1.3.0.4 | Other all-alpha-helical DNA-binding domains | High-mobility group (HMG) domain factors | TCF7-related |  | 15 | 18.43 | 39.22 | LEF1.NA@AFS.Lys@flabby-sepia-audemer@HughesLab.Streme@1000\_fa\_streme\_Motif1.ppm |
| LEF1 | HTS.IVT | test | YWF\_A\_AC40NTCCTTG |  |  | NA | 4.1.3.0.4 | Other all-alpha-helical DNA-binding domains | High-mobility group (HMG) domain factors | TCF7-related |  | 13 | 4.03 | 40.73 | LEF1.NA@HTS.IVT@gummy-yellow-urchin+cloudy-harlequin-serval+dorky-rose-termite+gummy-aquamarine-crocodile@Halle.Dimont@Motif\_1\_dtrue\_htsversion.ppm |
| LEF1 | HTS.Lys | train | YWL\_A\_AT40NAGCCTC |  |  | NA | 4.1.3.0.4 | Other all-alpha-helical DNA-binding domains | High-mobility group (HMG) domain factors | TCF7-related |  | 20 | 9.25 | 45.74 | LEF1.NA@AFS.Lys@flabby-sepia-audemer+crabby-emerald-whale+stuffy-ultramarine-elephant@Halle.Dimont@Motif\_1\_w20\_astrained.ppm |
| LEF1 | SMS | train | SRR3405063 |  |  | NA | 4.1.3.0.4 | Other all-alpha-helical DNA-binding domains | High-mobility group (HMG) domain factors | TCF7-related |  | 12 | 5.28 | 42.55 | LEF1.NA@HTS.IVT@gummy-yellow-urchin+cloudy-harlequin-serval+dorky-rose-termite+gummy-aquamarine-crocodile@Halle.Dimont@Motif\_1\_sampled\_e4\_astrained.ppm |
| NFKB1 | CHS | test | THC\_0624 |  |  | NA | 6.1.1.1.1 | Immunoglobulin fold | Rel homology region (RHR) factors | NFkappaB-related | NFkappaB p50-like | 13 | 17.46 | 59.44 | NFKB1.NA@CHS@slimy-sepia-ferret@autosome-ru.ChIPMunk@999seq\_7to15\_m0.pcm |
| NFKB1 | GHTS.Lys | test | YWL\_B\_AffSeq\_A9\_NFKB1 |  |  | NA | 6.1.1.1.1 | Immunoglobulin fold | Rel homology region (RHR) factors | NFkappaB-related | NFkappaB p50-like | 12 | 13.09 | 66.88 | NFKB1.NA@HTS.Lys@cheeky-champagne-chimpanzee+sleazy-maroon-newt+scummy-turquoise-barb+sleazy-heliotrope-gibbon@faltejsk.ProBound@motif\_with\_ns.ppm |
| NFKB1 | HTS.IVT | test | YWF\_A\_TT40NCTCGTC |  |  | NA | 6.1.1.1.1 | Immunoglobulin fold | Rel homology region (RHR) factors | NFkappaB-related | NFkappaB p50-like | 12 | 3.72 | 55.25 | NFKB1.NA@HTS.Lys@cheeky-champagne-chimpanzee+sleazy-maroon-newt+scummy-turquoise-barb+sleazy-heliotrope-gibbon@faltejsk.ProBound@motif\_without\_ns.ppm |
| NFKB1 | HTS.Lys | train | YWL\_A\_AC40NGCTGCT |  |  | NA | 6.1.1.1.1 | Immunoglobulin fold | Rel homology region (RHR) factors | NFkappaB-related | NFkappaB p50-like | 12 | 3.72 | 55.25 | NFKB1.NA@HTS.Lys@cheeky-champagne-chimpanzee+sleazy-maroon-newt+scummy-turquoise-barb+sleazy-heliotrope-gibbon@faltejsk.ProBound@motif\_without\_ns.ppm |
| NFKB1 | SMS | train | SRR3405069 |  |  | NA | 6.1.1.1.1 | Immunoglobulin fold | Rel homology region (RHR) factors | NFkappaB-related | NFkappaB p50-like | 12 | 13.09 | 66.88 | NFKB1.NA@HTS.Lys@cheeky-champagne-chimpanzee+sleazy-maroon-newt+scummy-turquoise-barb+sleazy-heliotrope-gibbon@faltejsk.ProBound@motif\_with\_ns.ppm |
