## Supplementary Data SD1 for "Inferring binding specificities of human transcription factors with the wisdom of crowds": ibis_motifs_Leaderboard_G2A.html

 

| TF | Platform | Slice | Replicate ID | logo-direct | logo-revcomp | Construct type | tfclass:id | tfclass:superclass | tfclass:class | tfclass:family | tfclass:subfamily | Motif length | Information content | GC% | Motif |
| --- | --- | --- | --- | --- | --- | --- | --- | --- | --- | --- | --- | --- | --- | --- | --- |
| PRDM5 | CHS | train | THC\_0307.Rep-MICHELLE\_0314 |  |  | DBD | 2.3.3.0.195 | Zinc-coordinating DNA-binding domains | C2H2 zinc finger factors | More than 3 adjacent zinc fingers | Unclassified | 14 | 9.19 | 59.62 | PRDM5.DBD@SMS@shabby-aqua-eel@Halle.Dimont@Motif\_1\_posneg\_bg4\_w15\_astrained.ppm |
| PRDM5 | CHS | test | THC\_0904 |  |  | DBD | 2.3.3.0.195 | Zinc-coordinating DNA-binding domains | C2H2 zinc finger factors | More than 3 adjacent zinc fingers | Unclassified | 14 | 9.19 | 59.62 | PRDM5.DBD@SMS@shabby-aqua-eel@Halle.Dimont@Motif\_1\_posneg\_bg4\_w15\_astrained.ppm |
| PRDM5 | CHS | train | THC\_0307.Rep-DIANA\_0293 |  |  | FL | 2.3.3.0.195 | Zinc-coordinating DNA-binding domains | C2H2 zinc finger factors | More than 3 adjacent zinc fingers | Unclassified | 15 | 11.38 | 65.87 | PRDM5.FL@CHS@sunny-chestnut-slug@HughesLab.MEME@500\_fa\_memechip\_Motif1.ppm |
| PRDM5 | GHTS.GFPIVT | test | YWQ\_B\_AffSeq\_B12\_PRDM5-DBD |  |  | FL | 2.3.3.0.195 | Zinc-coordinating DNA-binding domains | C2H2 zinc finger factors | More than 3 adjacent zinc fingers | Unclassified | 29 | 12.97 | 59.3 | PRDM5.FL@HTS.GFPIVT@lumpy-maroon-lemur@HughesLab.Homer@topk\_cycle=C3\_k=5\_top=10000\_fasta\_homer\_minw3\_maxw\_40\_Motif1.ppm |
| PRDM5 | GHTS.GFPIVT | train | YWQ\_B\_AffSeq\_F7\_PRDM5-FL |  |  | FL | 2.3.3.0.195 | Zinc-coordinating DNA-binding domains | C2H2 zinc finger factors | More than 3 adjacent zinc fingers | Unclassified | 29 | 12.97 | 59.3 | PRDM5.FL@HTS.GFPIVT@lumpy-maroon-lemur@HughesLab.Homer@topk\_cycle=C3\_k=5\_top=10000\_fasta\_homer\_minw3\_maxw\_40\_Motif1.ppm |
| PRDM5 | GHTS.Lys | train | YWK\_D\_AffSeq\_B1\_PRDM5 |  |  | FL | 2.3.3.0.195 | Zinc-coordinating DNA-binding domains | C2H2 zinc finger factors | More than 3 adjacent zinc fingers | Unclassified | 32 | 21.77 | 63.61 | PRDM5.FL@HTS.Lys@snazzy-tangerine-eel@AJolma.Autoseed@Multinom3\_Onehit\_Seed\_YDGGNKBBMARGGNNCGNVHGGWGVDYVGGKBackgroundCyc3.ppm |
| PRDM5 | GHTS.Lys | train | YWK\_B\_AffSeq\_B1\_PRDM5 |  |  | FL | 2.3.3.0.195 | Zinc-coordinating DNA-binding domains | C2H2 zinc finger factors | More than 3 adjacent zinc fingers | Unclassified | 29 | 12.97 | 59.3 | PRDM5.FL@HTS.GFPIVT@lumpy-maroon-lemur@HughesLab.Homer@topk\_cycle=C3\_k=5\_top=10000\_fasta\_homer\_minw3\_maxw\_40\_Motif1.ppm |
| PRDM5 | HTS.Lys | test | YWK\_C\_AG40NTCGACT |  |  | FL | 2.3.3.0.195 | Zinc-coordinating DNA-binding domains | C2H2 zinc finger factors | More than 3 adjacent zinc fingers | Unclassified | 26 | 16.43 | 58.24 | PRDM5.FL@HTS.Lys@jumpy-cardinal-jackal+silly-black-cat@autosome-ru.ChIPMunk@topk\_cycle=C3+C4\_k=5\_top=2500.pcm |
| PRDM5 | HTS.Lys | test | YWK\_A\_AG40NTCGACT |  |  | FL | 2.3.3.0.195 | Zinc-coordinating DNA-binding domains | C2H2 zinc finger factors | More than 3 adjacent zinc fingers | Unclassified | 26 | 16.43 | 58.24 | PRDM5.FL@HTS.Lys@jumpy-cardinal-jackal+silly-black-cat@autosome-ru.ChIPMunk@topk\_cycle=C3+C4\_k=5\_top=2500.pcm |
| PRDM5 | SMS | test | UT380-158 |  |  | FL | 2.3.3.0.195 | Zinc-coordinating DNA-binding domains | C2H2 zinc finger factors | More than 3 adjacent zinc fingers | Unclassified | 29 | 12.97 | 59.3 | PRDM5.FL@HTS.GFPIVT@lumpy-maroon-lemur@HughesLab.Homer@topk\_cycle=C3\_k=5\_top=10000\_fasta\_homer\_minw3\_maxw\_40\_Motif1.ppm |
| ZNF362 | CHS | train | THC\_0364.Rep-MICHELLE\_0314 |  |  | FL | 2.3.3.37.1 | Zinc-coordinating DNA-binding domains | C2H2 zinc finger factors | More than 3 adjacent zinc fingers | ZNF362-like | 12 | 6.67 | 28.82 | ZNF362.FL@HTS.GFPIVT@ready-carmine-husky+stinky-crimson-squirrel+lumpy-tan-bonobo@faltejsk.ProBound@motif\_with\_ns.ppm |
| ZNF362 | CHS | train | THC\_0364.Rep-DIANA\_0293 |  |  | FL | 2.3.3.37.1 | Zinc-coordinating DNA-binding domains | C2H2 zinc finger factors | More than 3 adjacent zinc fingers | ZNF362-like | 12 | 5.8 | 30.55 | ZNF362.FL@HTS.Lys@nippy-puce-ant+snappy-azure-ferret+skinny-chestnut-alligator+pokey-bronze-squid@faltejsk.ProBound@motif\_with\_ns.ppm |
| ZNF362 | CHS | test | THC\_0411.Rep-MICHELLE\_0314 |  |  | FL | 2.3.3.37.1 | Zinc-coordinating DNA-binding domains | C2H2 zinc finger factors | More than 3 adjacent zinc fingers | ZNF362-like | 12 | 6.67 | 28.82 | ZNF362.FL@HTS.GFPIVT@ready-carmine-husky+stinky-crimson-squirrel+lumpy-tan-bonobo@faltejsk.ProBound@motif\_with\_ns.ppm |
| ZNF362 | CHS | train | THC\_0411.Rep-DIANA\_0293 |  |  | FL | 2.3.3.37.1 | Zinc-coordinating DNA-binding domains | C2H2 zinc finger factors | More than 3 adjacent zinc fingers | ZNF362-like | 7 | 11.03 | 3.14 | ZNF362.FL@SMS@smelly-firebrick-dalmatian@HughesLab.Streme@topk\_k=5\_top=500\_Motif1\_min3max30.ppm |
| ZNF362 | GHTS.IVT | train | YWH\_B\_AffSeq\_F10\_ZNF362 |  |  | FL | 2.3.3.37.1 | Zinc-coordinating DNA-binding domains | C2H2 zinc finger factors | More than 3 adjacent zinc fingers | ZNF362-like | 20 | 2.21 | 49.72 | ZNF362.FL@AFS.GFPIVT@foggy-khaki-quail+wheezy-brass-chamois+cheeky-celadon-spider@Halle.Dimont@Motif\_2\_imw20\_astrained.ppm |
| ZNF362 | GHTS.GFPIVT | test | YWR\_B\_AffSeq\_C5\_ZNF362-FL |  |  | FL | 2.3.3.37.1 | Zinc-coordinating DNA-binding domains | C2H2 zinc finger factors | More than 3 adjacent zinc fingers | ZNF362-like | 20 | 6.25 | 46.53 | ZNF362.FL@AFS.GFPIVT@foggy-khaki-quail+wheezy-brass-chamois+cheeky-celadon-spider@Halle.Dimont@Motif\_1\_w20\_astrained.ppm |
| ZNF362 | GHTS.Lys | train | YWK\_D\_AffSeq\_E5\_ZNF362 |  |  | FL | 2.3.3.37.1 | Zinc-coordinating DNA-binding domains | C2H2 zinc finger factors | More than 3 adjacent zinc fingers | ZNF362-like | 33 | 21.15 | 30.04 | ZNF362.FL@HTS.Lys@hazy-myrtle-scorpion+skinny-alizarin-fly@HughesLab.Homer@topk\_cycle=C3+C4\_k=5\_top=500\_fasta\_homer\_minw3\_maxw\_40\_Motif4.ppm |
| ZNF362 | GHTS.Lys | train | YWK\_B\_AffSeq\_E5\_ZNF362 |  |  | FL | 2.3.3.37.1 | Zinc-coordinating DNA-binding domains | C2H2 zinc finger factors | More than 3 adjacent zinc fingers | ZNF362-like | 20 | 3.23 | 40.28 | ZNF362.FL@CHS@geeky-tangerine-hornet@Halle.Dimont@Motif\_1\_w20\_astrained.ppm |
| ZNF362 | HTS.IVT | test | YWH\_A\_GT40NCATTCT |  |  | FL | 2.3.3.37.1 | Zinc-coordinating DNA-binding domains | C2H2 zinc finger factors | More than 3 adjacent zinc fingers | ZNF362-like | 8 | 12.78 | 9.36 | ZNF362.FL@SMS@smelly-firebrick-dalmatian@HughesLab.MEME@topk\_k=5\_top=2500\_Motif1.ppm |
| ZNF362 | HTS.GFPIVT | test | YWR\_A\_CA40NCTTTGA |  |  | FL | 2.3.3.37.1 | Zinc-coordinating DNA-binding domains | C2H2 zinc finger factors | More than 3 adjacent zinc fingers | ZNF362-like | 9 | 6.84 | 26.21 | ZNF362.FL@HTS.GFPIVT@lumpy-tan-bonobo@AJolma.Autoseed@Multinom1\_Onehit\_Seed\_AAAAAAACBackgroundCyc0.ppm |
| ZNF362 | HTS.Lys | test | YWK\_A\_GA40NACTTTG |  |  | FL | 2.3.3.37.1 | Zinc-coordinating DNA-binding domains | C2H2 zinc finger factors | More than 3 adjacent zinc fingers | ZNF362-like | 9 | 6.84 | 26.21 | ZNF362.FL@HTS.GFPIVT@lumpy-tan-bonobo@AJolma.Autoseed@Multinom1\_Onehit\_Seed\_AAAAAAACBackgroundCyc0.ppm |
| ZNF362 | HTS.Lys | test | YWK\_C\_GA40NACTTTG |  |  | FL | 2.3.3.37.1 | Zinc-coordinating DNA-binding domains | C2H2 zinc finger factors | More than 3 adjacent zinc fingers | ZNF362-like | 9 | 6.84 | 26.21 | ZNF362.FL@HTS.GFPIVT@lumpy-tan-bonobo@AJolma.Autoseed@Multinom1\_Onehit\_Seed\_AAAAAAACBackgroundCyc0.ppm |
| ZNF362 | SMS | test | UT380-331-2 |  |  | FL | 2.3.3.37.1 | Zinc-coordinating DNA-binding domains | C2H2 zinc finger factors | More than 3 adjacent zinc fingers | ZNF362-like | 8 | 12.78 | 9.36 | ZNF362.FL@SMS@smelly-firebrick-dalmatian@HughesLab.MEME@topk\_k=5\_top=2500\_Motif1.ppm |
| ZNF407 | CHS | train | THC\_0668 |  |  | DBD | 2.3.4.0.73 | Zinc-coordinating DNA-binding domains | C2H2 zinc finger factors | Multiple dispersed zinc fingers | Unclassified | 22 | 16.18 | 61.93 | ZNF407.DBD@AFS.GFPIVT@muzzy-scarlet-beaver@autosome-ru.ChIPMunk@topk\_cycle=C3\_k=5\_top=10000.pcm |
| ZNF407 | GHTS.GFPIVT | test | YWR\_B\_AffSeq\_A6\_ZNF407-DBD |  |  | DBD | 2.3.4.0.73 | Zinc-coordinating DNA-binding domains | C2H2 zinc finger factors | Multiple dispersed zinc fingers | Unclassified | 22 | 16.18 | 61.93 | ZNF407.DBD@AFS.GFPIVT@muzzy-scarlet-beaver@autosome-ru.ChIPMunk@topk\_cycle=C3\_k=5\_top=10000.pcm |
| ZNF407 | GHTS.Lys | train | YWL\_B\_AffSeq\_A11\_ZNF407 |  |  | DBD | 2.3.4.0.73 | Zinc-coordinating DNA-binding domains | C2H2 zinc finger factors | Multiple dispersed zinc fingers | Unclassified | 22 | 16.18 | 61.93 | ZNF407.DBD@AFS.GFPIVT@muzzy-scarlet-beaver@autosome-ru.ChIPMunk@topk\_cycle=C3\_k=5\_top=10000.pcm |
| ZNF407 | HTS.IVT | test | YWF\_A\_AG40NTGGCTA |  |  | DBD | 2.3.4.0.73 | Zinc-coordinating DNA-binding domains | C2H2 zinc finger factors | Multiple dispersed zinc fingers | Unclassified | 20 | 7.53 | 58.86 | ZNF407.DBD@AFS.GFPIVT@flaky-platinum-mouse+shabby-mauve-corgi@Halle.Dimont@Motif\_1\_w20\_astrained.ppm |
| ZNF407 | HTS.GFPIVT | test | YWR\_A\_AA40NGTGGTG |  |  | DBD | 2.3.4.0.73 | Zinc-coordinating DNA-binding domains | C2H2 zinc finger factors | Multiple dispersed zinc fingers | Unclassified | 26 | 23.76 | 57.13 | ZNF407.DBD@HTS.IVT@breezy-zucchini-scorpion+skimpy-carmine-armadillo+nerdy-jade-frise+snazzy-mustard-oyster@OFornes.ExplaiNN@filter69\_1.ppm |
| ZNF407 | HTS.Lys | test | YWL\_A\_AC40NTGTTAC |  |  | DBD | 2.3.4.0.73 | Zinc-coordinating DNA-binding domains | C2H2 zinc finger factors | Multiple dispersed zinc fingers | Unclassified | 20 | 4.92 | 53.07 | ZNF407.DBD@AFS.GFPIVT@flaky-platinum-mouse+shabby-mauve-corgi@Halle.Dimont@Motif\_1\_imw20\_astrained.ppm |
| ZNF407 | SMS | test | UT380-340 |  |  | DBD | 2.3.4.0.73 | Zinc-coordinating DNA-binding domains | C2H2 zinc finger factors | Multiple dispersed zinc fingers | Unclassified | 20 | 7.53 | 58.86 | ZNF407.DBD@AFS.GFPIVT@flaky-platinum-mouse+shabby-mauve-corgi@Halle.Dimont@Motif\_1\_w20\_astrained.ppm |
| GABPA | CHS | train | THC\_0866 |  |  | NA | 3.5.2.1.5 | Helix-turn-helix domains | Tryptophan cluster factors | Ets-related | ETS-like | 12 | 6.74 | 57.96 | GABPA.NA@HTS.IVT@scaly-wisteria-centipede+muggy-violet-foxhound+wiggy-ochre-llama+sleazy-pear-dane@Halle.Dimont@Motif\_1\_sampled\_e1\_5\_astrained.ppm |
| GABPA | CHS | test | THC\_0864 |  |  | NA | 3.5.2.1.5 | Helix-turn-helix domains | Tryptophan cluster factors | Ets-related | ETS-like | 12 | 4.35 | 52.01 | GABPA.NA@HTS.IVT@scaly-wisteria-centipede+muggy-violet-foxhound+wiggy-ochre-llama+sleazy-pear-dane@Halle.Dimont@Motif\_1\_sampled\_e1\_5\_bg4\_astrained.ppm |
| GABPA | GHTS.IVT | train | YWE\_B\_AffSeq\_C12\_GABPA |  |  | NA | 3.5.2.1.5 | Helix-turn-helix domains | Tryptophan cluster factors | Ets-related | ETS-like | 12 | 4.35 | 52.01 | GABPA.NA@HTS.IVT@scaly-wisteria-centipede+muggy-violet-foxhound+wiggy-ochre-llama+sleazy-pear-dane@Halle.Dimont@Motif\_1\_sampled\_e1\_5\_bg4\_astrained.ppm |
| GABPA | GHTS.Lys | test | YWL\_B\_AffSeq\_G8\_GABPA |  |  | NA | 3.5.2.1.5 | Helix-turn-helix domains | Tryptophan cluster factors | Ets-related | ETS-like | 15 | 5.88 | 53.82 | GABPA.NA@HTS.Lys@lumpy-magenta-burmese+queasy-yellow-blue+silly-asparagus-crane+shaggy-wheat-walrus@Halle.Dimont@Motif\_2\_dtrue\_htsversion.ppm |
| GABPA | PBM | test | PBM14352 |  |  | NA | 3.5.2.1.5 | Helix-turn-helix domains | Tryptophan cluster factors | Ets-related | ETS-like | 15 | 5.88 | 53.82 | GABPA.NA@HTS.Lys@lumpy-magenta-burmese+queasy-yellow-blue+silly-asparagus-crane+shaggy-wheat-walrus@Halle.Dimont@Motif\_2\_dtrue\_htsversion.ppm |
| GABPA | HTS.IVT | test | YWE\_A\_CG40NAATAGC |  |  | NA | 3.5.2.1.5 | Helix-turn-helix domains | Tryptophan cluster factors | Ets-related | ETS-like | 15 | 5.88 | 53.82 | GABPA.NA@HTS.Lys@lumpy-magenta-burmese+queasy-yellow-blue+silly-asparagus-crane+shaggy-wheat-walrus@Halle.Dimont@Motif\_2\_dtrue\_htsversion.ppm |
| GABPA | HTS.Lys | test | YWL\_A\_TC40NGCGATT |  |  | NA | 3.5.2.1.5 | Helix-turn-helix domains | Tryptophan cluster factors | Ets-related | ETS-like | 10 | 6.43 | 56.0 | GABPA.NA@AFS.Lys@blurry-seashell-gorilla+flimsy-carmine-butterfly+sickly-fuchsia-urchin+sunny-charcoal-beaver@HughesLab.Homer@topk\_cycle=C1+C2+C3+C4\_k=5\_top=500\_fasta\_homer\_minw3\_maxw\_40\_Motif1.ppm |
| SP140 | CHS | train | THC\_0193 |  |  | DBD | 5.3.5.1.1 | alpha-Helices exposed by beta-structures | SAND domain factors | Sp140-Sp100 | Sp140 | 11 | 4.42 | 52.13 | SP140.DBD@HTS.GFPIVT@trippy-lime-rattlesnake+sunny-auburn-turtle+fuzzy-purple-jaguar@Halle.Dimont@Motif\_1\_sampled\_e4\_astrained.ppm |
| SP140 | GHTS.GFPIVT | train | YWQ\_B\_AffSeq\_G2\_SP140-DBD |  |  | DBD | 5.3.5.1.1 | alpha-Helices exposed by beta-structures | SAND domain factors | Sp140-Sp100 | Sp140 | 12 | 4.76 | 51.42 | SP140.DBD@AFS.GFPIVT@sunny-persimmon-gar+baggy-mustard-terrier@Halle.Dimont@Motif\_1\_imw20\_astrained.ppm |
| SP140 | PBM | test | PBM13973 |  |  | DBD | 5.3.5.1.1 | alpha-Helices exposed by beta-structures | SAND domain factors | Sp140-Sp100 | Sp140 | 10 | 4.96 | 49.07 | SP140.DBD@PBM.HK@stinky-salmon-pinscher@Halle.Dimont@Motif\_1\_astrained.ppm |
| SP140 | HTS.GFPIVT | test | YWQ\_A\_TC40NTAAGTG |  |  | DBD | 5.3.5.1.1 | alpha-Helices exposed by beta-structures | SAND domain factors | Sp140-Sp100 | Sp140 | 12 | 3.58 | 54.08 | SP140.DBD@HTS.GFPIVT@fuzzy-purple-jaguar+sunny-auburn-turtle+trippy-lime-rattlesnake@faltejsk.ProBound@motif\_without\_ns.ppm |
| SP140 | HTS.GFPIVT | test | YWQ\_A\_TA40NTCACTC |  |  | DBD | 5.3.5.1.1 | alpha-Helices exposed by beta-structures | SAND domain factors | Sp140-Sp100 | Sp140 | 12 | 3.58 | 54.08 | SP140.DBD@HTS.GFPIVT@fuzzy-purple-jaguar+sunny-auburn-turtle+trippy-lime-rattlesnake@faltejsk.ProBound@motif\_without\_ns.ppm |
| SP140 | SMS | test | UT380-202-2 |  |  | NA | 5.3.5.1.1 | alpha-Helices exposed by beta-structures | SAND domain factors | Sp140-Sp100 | Sp140 | 12 | 1.8 | 50.69 | SP140.NA@SMS@flimsy-white-havanese@faltejsk.ProBound@motif\_without\_ns.ppm |
