## Supplementary Data SD1 for "Inferring binding specificities of human transcription factors with the wisdom of crowds": jquery.dragtable.mod.min.js

```
(function(factory){if (typeof define === 'function' && define.amd){define(['jquery'], factory);} else if (typeof module === 'object' && typeof module.exports === 'object'){module.exports = factory(require('jquery'));} else {factory(jQuery);}}(function(jQuery){
/*! Dragtable Mod for TableSorter - updated 10/31/2015 (v2.24.0) */
!function(T){"use strict";var u=T.tablesorter,t=(u.dragtable={create:function(t){var a,e=t.originalTable.el,i=t.options.dragHandle.replace(".","");e.children("thead").children().children("th,td").each(function(){var e=T(this);e.find(t.options.dragHandle+",."+i+"-disabled").length||(a=!t.options.dragaccept||e.hasClass(t.options.dragaccept.replace(".","")),e.wrapInner('<div class="'+t.options.sortClass.replace(".","")+'"/>').prepend('<div class="'+i+(a?"":"-disabled")+'"></div>'))})},start:function(e){(e=T(e)[0])&&e.config&&(e.config.widgetOptions.dragtableLast={search:T(e).data("lastSearch"),order:u.dragtable.getOrder(e)})},update:function(e){var a,i,t=e.originalTable,r=t.el[0],n=T(r),o=r.config,l=o&&o.widgetOptions,s=t.startIndex-1,t=t.endIndex-1,d=u.dragtable.getOrder(r)||[],c=u.hasWidget(n,"filter")||!1,h=l&&l.dragtableLast||{},b=[];(h.order||[]).join("")!==d.join("")&&(o.sortList.length&&(a=T.extend(!0,[],o.sortList),T.each(d,function(e,t){i=u.isValueInArray(parseInt(t,10),a),t!==h.order[e]&&0<=i&&(o.sortList[i][0]=e)})),c&&T.each(h.search||[],function(e){b[e]=h.search[d[e]]}),(r=!!u.hasWidget(o.$table,"editable")&&l.editable_columnsArray)&&(o.widgetOptions.editable_columnsArray=u.dragtable.reindexArrayItem(r,s,t)),(r=!!u.hasWidget(o.$table,"math")&&l.math_ignore)&&(o.widgetOptions.math_ignore=u.dragtable.reindexArrayItem(r,s,t)),(r=!!u.hasWidget(o.$table,"resizable")&&l.resizable_widths)&&(l.resizable_widths=u.dragtable.moveArrayItem(r,s,t)),u.updateAll(o,!1,function(){c&&setTimeout(function(){o.lastCombinedFilter=null,o.$table.data("lastSearch",b),u.setFilters(n,b),T.isFunction(e.options.tablesorterComplete)&&e.options.tablesorterComplete(o.table)},10)}))},getOrder:function(e){return T(e).children("thead").children("."+u.css.headerRow).children().map(function(){return T(this).attr("data-column")}).get()||[]},startColumnMove:function(e){var t,a=e.el[0].config,i=e.startIndex-1,r=e.endIndex-1,e=a.columns-1,n=r!=e&&r<=i,e=a.$table.children().children("tr");a.debug&&console.log("Inserting column "+i+(n?" before":" after")+" column "+r),e.each(function(){(t=T(this).children()).eq(i)[n?"insertBefore":"insertAfter"](t.eq(r))}),(t=a.$table.children("colgroup").children()).eq(i)[n?"insertBefore":"insertAfter"](t.eq(r))},swapNodes:function(e,t){for(var a,i,r=e.length,n=0;n<r;n++)a=e[n].parentNode,i=e[n].nextSibling===t[n]?e[n]:e[n].nextSibling,t[n].parentNode.insertBefore(e[n],t[n]),a.insertBefore(t[n],i)},moveArrayItem:function(e,t,a){var i,r=e.length;if(r<=a)for(i=a-r;1+i--;)e.push(void 0);return e.splice(a,0,e.splice(t,1)[0]),e},reindexArrayItem:function(e,a,i){var t=T.inArray(i,e),r=T.inArray(a,e),n=(Math.max.apply(Math,e),[]);return 0<=t&&0<=r?e:(T.each(e,function(e,t){i<a?n.push(i<=t?t+(t<a?1:0):t):a<i&&(t===a?n.push(i):t<i&&a<=t?n.push(t-1):t<=i?n.push(t):a<t&&n.push(t+(t<i?0:1)))}),n.sort())}},
/*! dragtable v2.0.14 Mod */
T.widget("akottr.dragtable",{options:{revert:!1,dragHandle:".table-handle",maxMovingRows:40,excludeFooter:!1,onlyHeaderThreshold:100,dragaccept:null,persistState:null,restoreState:null,exact:!0,clickDelay:10,containment:null,cursor:"move",cursorAt:!1,distance:0,tolerance:"pointer",axis:"x",beforeStart:T.noop,beforeMoving:T.noop,beforeReorganize:T.noop,beforeStop:T.noop,tablesorterComplete:null,sortClass:".sorter"},originalTable:{el:null,selectedHandle:null,sortOrder:null,startIndex:0,endIndex:0},sortableTable:{el:T(),selectedHandle:T(),movingRow:T()},persistState:function(){var t=this;this.originalTable.el.find("th").each(function(e){""!==this.id&&(t.originalTable.sortOrder[this.id]=e)}),T.ajax({url:this.options.persistState,data:this.originalTable.sortOrder})},_restoreState:function(e){for(var t in e)t in e&&(this.originalTable.startIndex=T("#"+t).closest("th").prevAll().length+1,this.originalTable.endIndex=parseInt(e[t],10)+1,this._bubbleCols())},_bubbleCols:function(){u.dragtable.startColumnMove(this.originalTable)},_rearrangeTableBackroundProcessing:function(){var e=this;return function(){e._bubbleCols(),e.options.beforeStop(e.originalTable),e.sortableTable.el.remove(),T("#__dragtable_disable_text_selection__").remove(),t?T(document.body).attr("onselectstart",t):T(document.body).removeAttr("onselectstart"),a?T(document.body).attr("unselectable",a):T(document.body).removeAttr("unselectable"),u.dragtable.update(e),T.isFunction(e.options.persistState)?e.options.persistState(e.originalTable):e.persistState()}},_rearrangeTable:function(){var e=this;return function(){e.originalTable.selectedHandle.removeClass("dragtable-handle-selected"),e.sortableTable.el.sortable("disable"),e.sortableTable.el.addClass("dragtable-disabled"),e.options.beforeReorganize(e.originalTable,e.sortableTable),e.originalTable.endIndex=e.sortableTable.movingRow.prevAll().length+1,setTimeout(e._rearrangeTableBackroundProcessing(),50)}},_generateSortable:function(e){e.cancelBubble?e.cancelBubble=!0:e.stopPropagation();for(var t=this,a=this.originalTable.el[0].attributes,i="",r=0;r<a.length;r++)(a[r].value||a[r].nodeValue)&&"id"!=a[r].nodeName&&"width"!=a[r].nodeName&&(i+=a[r].nodeName+'="'+(a[r].value||a[r].nodeValue)+'" ');for(var n,o=[],l=[],s=(t.originalTable.el.children("thead, tbody").children("tr:visible").slice(0,t.options.maxMovingRow).each(function(){for(var e=this.attributes,t="",a=0;a<e.length;a++)(e[a].value||e[a].nodeValue)&&"id"!=e[a].nodeName&&(t+=" "+e[a].nodeName+'="'+(e[a].value||e[a].nodeValue)+'"');o.push(t),l.push(T(this).height())}),[]),d=0,c=t.originalTable.el.children(),h=c.filter("thead").children("tr:visible"),b=c.filter("tbody").children("tr:visible"),u=(h.eq(0).children("th, td").filter(":visible").each(function(){var e=T(this).outerWidth();s.push(e),d+=e}),t.options.exact&&(g=d-t.originalTable.el.outerWidth(),s[0]-=g),d+=2,0),g=(c.filter("caption").each(function(){u+=T(this).outerHeight()}),'<ul class="dragtable-sortable" style="position:absolute; width:'+d+'px;">'),p=[],f=h.eq(0).children("th, td").length,r=0;r<f;r++){var m=h.children(":nth-child("+(r+1)+")");m.is(":visible")&&(n=0,p[r]='<li style="width:'+m.outerWidth()+'px;"><table '+i+">"+(u?'<caption style="height:'+u+'px;"></caption>':"")+"<thead>",h.each(function(e){p[r]+="<tr "+o[n++]+(l[e]?' style="height:'+l[e]+'px;"':"")+">"+m[e].outerHTML+"</tr>"}),p[r]+="</thead><tbody>",m=b.children(":nth-child("+(r+1)+")"),(m=1<t.options.maxMovingRows?m.add(b.children(":nth-child("+(r+1)+")").slice(0,t.options.maxMovingRows-1)):m).each(function(e){p[r]+="<tr "+o[n++]+(l[e]?' style="height:'+l[e]+'px;"':"")+">"+this.outerHTML+"</tr>"}),p[r]+="</tbody>",t.options.excludeFooter||(p[r]+="<tfoot><tr "+o[n++]+">"+c.filter("tfoot").children("tr:visible").children()[r].outerHTML+"</tr></tfoot>"),p[r]+="</table></li>")}g+=p.join("")+"</ul>",this.sortableTable.el=this.originalTable.el.before(g).prev(),this.sortableTable.el.find("> li > table").each(function(e){T(this).css("width",s[e]+"px")}),this.sortableTable.selectedHandle=this.sortableTable.el.find("th .dragtable-handle-selected");var g=this.options.dragaccept?"li:has("+this.options.dragaccept+")":"li",e=(this.sortableTable.el.sortable({items:g,stop:this._rearrangeTable(),revert:this.options.revert,tolerance:this.options.tolerance,containment:this.options.containment,cursor:this.options.cursor,cursorAt:this.options.cursorAt,distance:this.options.distance,axis:this.options.axis}),this.originalTable.startIndex=T(e.target).closest("th,td").prevAll().length+1,this.options.beforeMoving(this.originalTable,this.sortableTable),this.sortableTable.movingRow=this.sortableTable.el.children("li:nth-child("+this.originalTable.startIndex+")"),g=T('<style id="__dragtable_disable_text_selection__" type="text/css">body { -ms-user-select:none;-moz-user-select:-moz-none;-khtml-user-select:none;-webkit-user-select:none;user-select:none; }</style>'),T(document.head).append(g),T(document.body).attr("onselectstart","return false;").attr("unselectable","on"),window.getSelection?window.getSelection().removeAllRanges():document.selection.empty(),this.sortableTable.movingRow.trigger(T.extend(T.Event(e.type),{which:1,clientX:e.clientX,clientY:e.clientY,pageX:e.pageX,pageY:e.pageY,screenX:e.screenX,screenY:e.screenY})),this.sortableTable.el.find(".ui-sortable-placeholder"));0<e.height()&&e.css("height",this.sortableTable.el.find(".ui-sortable-helper").height()),e.html('<div class="outer" style="height:100%;"><div class="inner" style="height:100%;"></div></div>')},bindTo:{},_create:function(){var t=this;t.originalTable={el:t.element,selectedHandle:T(),sortOrder:{},startIndex:0,endIndex:0},u.dragtable.create(t),t.bindTo="> thead > tr > "+(t.options.dragaccept||"th, td"),t.element.find(t.bindTo).find(t.options.dragHandle).length&&(t.bindTo+=" "+t.options.dragHandle),T.isFunction(t.options.restoreState)?t.options.restoreState(t.originalTable):t._restoreState(t.options.restoreState),t.originalTable.el.on("mousedown.dragtable",t.bindTo,function(e){1===e.which&&(u.dragtable.start(t.originalTable.el),!1!==t.options.beforeStart(t.originalTable))&&(clearTimeout(t.downTimer),t.downTimer=setTimeout(function(){t.originalTable.selectedHandle=T(t),t.originalTable.selectedHandle.addClass("dragtable-handle-selected"),t._generateSortable(e)},t.options.clickDelay))}).on("mouseup.dragtable",t.options.dragHandle,function(){clearTimeout(t.downTimer)})},redraw:function(){this.destroy(),this._create()},destroy:function(){this.originalTable.el.off("mousedown.dragtable mouseup.dragtable",this.bindTo),T.Widget.prototype.destroy.apply(this,arguments)}}),T(document.body).attr("onselectstart")),a=T(document.body).attr("unselectable")}(jQuery);return jQuery;}));
```
