## Supplementary Data SD1 for "Inferring binding specificities of human transcription factors with the wisdom of crowds": jquery.metadata.min.js

```
(function(factory){if (typeof define === 'function' && define.amd){define(['jquery'], factory);} else if (typeof module === 'object' && typeof module.exports === 'object'){module.exports = factory(require('jquery'));} else {factory(jQuery);}}(function(jQuery){
!function($){$.extend({metadata:{defaults:{type:"class",name:"metadata",cre:/(\{.*\})/,single:"metadata"},setType:function(t,e){this.defaults.type=t,this.defaults.name=e},get:function(elem,opts){var data,m,e,attr,settings=$.extend({},this.defaults,opts);if(settings.single.length||(settings.single="metadata"),data=$.data(elem,settings.single),!data){if(data="{}","class"===settings.type)m=settings.cre.exec(elem.className),m&&(data=m[1]);else if("elem"===settings.type){if(!elem.getElementsByTagName)return;e=elem.getElementsByTagName(settings.name),e.length&&(data=$.trim(e[0].innerHTML))}else void 0!==elem.getAttribute&&(attr=elem.getAttribute(settings.name),attr)&&(data=attr);data.indexOf("{")<0&&(data="{"+data+"}"),data=eval("("+data+")"),$.data(elem,settings.single,data)}return data}}}),$.fn.metadata=function(t){return $.metadata.get(this[0],t)}}(jQuery);return jQuery;}));
```
