## Supplementary Data SD1 for "Inferring binding specificities of human transcription factors with the wisdom of crowds": jquery.tablesorter.pager.min.js

```
(function(factory){if (typeof define === 'function' && define.amd){define(['jquery'], factory);} else if (typeof module === 'object' && typeof module.exports === 'object'){module.exports = factory(require('jquery'));} else {factory(jQuery);}}(function(jQuery){
/*!
* tablesorter (FORK) pager plugin
* updated 2020-03-03 (v2.31.3)
*/
!function(E){"use strict";var L=E.tablesorter;E.extend({tablesorterPager:new function(){this.defaults={container:null,ajaxUrl:null,customAjaxUrl:function(e,t){return t},ajaxError:null,ajaxObject:{dataType:"json"},processAjaxOnInit:!0,ajaxProcessing:function(e){return e},output:"{startRow} to {endRow} of {totalRows} rows",updateArrows:!0,page:0,pageReset:0,size:10,maxOptionSize:20,savePages:!0,storageKey:"tablesorter-pager",fixedHeight:!1,countChildRows:!1,removeRows:!1,cssFirst:".first",cssPrev:".prev",cssNext:".next",cssLast:".last",cssGoto:".gotoPage",cssPageDisplay:".pagedisplay",cssPageSize:".pagesize",cssErrorRow:"tablesorter-errorRow",cssDisabled:"disabled",totalRows:0,totalPages:0,filteredRows:0,filteredPages:0,ajaxCounter:0,currentFilters:[],startRow:0,endRow:0,$size:null,last:{}};function l(e,t,a,i,s,r){if("function"==typeof a.ajaxProcessing){t.config.$tbodies.eq(0).empty();var o,n,l,g,c,d,p,f,u,h,w,b,R=t.config,z=R.$table,x="",e=a.ajaxProcessing(e,t,i)||[0,[]];if(L.showError(t),r)L.debug(R,"pager")&&console.error("Pager >> Ajax Error",i,s,r),L.showError(t,i,s,r),R.$tbodies.eq(0).children("tr").detach(),a.totalRows=0;else{if(E.isArray(e)?(s=e[(i=isNaN(e[0])&&!isNaN(e[1]))?1:0],a.totalRows=isNaN(s)?a.totalRows||0:s,R.totalRows=R.filteredRows=a.filteredRows=a.totalRows,h=0!==a.totalRows&&e[i?0:1]||[],u=e[2]):(a.ajaxData=e,R.totalRows=a.totalRows=e.total,R.filteredRows=a.filteredRows=void 0!==e.filteredRows?e.filteredRows:e.total,u=e.headers,h=e.rows||[]),w=h&&h.length,h instanceof E)a.processAjaxOnInit&&(R.$tbodies.eq(0).empty(),R.$tbodies.eq(0).append(h));else if(w){for(o=0;o<w;o++){for(x+="<tr>",n=0;n<h[o].length;n++)x+=/^\s*<td/.test(h[o][n])?E.trim(h[o][n]):"<td>"+h[o][n]+"</td>";x+="</tr>"}a.processAjaxOnInit&&R.$tbodies.eq(0).html(x)}if(a.processAjaxOnInit=!0,u)for(c=(l=z.hasClass("hasStickyHeaders"))?R.widgetOptions.$sticky.children("thead:first").children("tr:not(."+R.cssIgnoreRow+")").children():"",g=z.find("tfoot tr:first").children(),b=(d=R.$headers.filter("th ")).length,n=0;n<b;n++)(p=d.eq(n)).find("."+L.css.icon).length?(f=p.find("."+L.css.icon).clone(!0),p.find("."+L.css.headerIn).html(u[n]).append(f),l&&c.length&&(f=c.eq(n).find("."+L.css.icon).clone(!0),c.eq(n).find("."+L.css.headerIn).html(u[n]).append(f))):(p.find("."+L.css.headerIn).html(u[n]),l&&c.length&&(a.$container=a.$container.add(R.widgetOptions.$sticky),c.eq(n).find("."+L.css.headerIn).html(u[n]))),g.eq(n).html(u[n])}R.showProcessing&&L.isProcessing(t),r=N(a,a.size,"get"),a.totalPages="all"===r?1:Math.ceil(a.totalRows/r),a.last.totalRows=a.totalRows,a.last.currentFilters=a.currentFilters,a.last.sortList=(R.sortList||[]).join(","),j(t,a,!1),L.updateCache(R,function(){a.initialized&&setTimeout(function(){L.debug(R,"pager")&&console.log("Pager >> Triggering pagerChange"),z.triggerHandler("pagerChange",a),L.applyWidget(t),j(t,a,!0)},0)})}a.initialized||F(t,a)}function d(e,t){t.page=0,S(e,t)}function u(e,t){t.page=$(e,t)-1,S(e,t)}function p(e,t){t.page++;var a=$(e,t)-1;t.page>=a&&(t.page=a),S(e,t)}function f(e,t){t.page--,t.page<=0&&(t.page=0),S(e,t)}function h(e,t,a){var i,s,r=e.config;t.$container.find(t.cssGoto+","+t.cssPageSize+",.ts-startRow, .ts-page").removeClass(t.cssDisabled).removeAttr("disabled").each(function(){this.ariaDisabled=!1}),t.isDisabled=!1,t.page=E.data(e,"pagerLastPage")||t.page||0,i=(s=t.$container.find(t.cssPageSize)).find("option[selected]").val(),t.size=E.data(e,"pagerLastSize")||N(t,i,"get"),t.totalPages="all"===t.size?1:Math.ceil($(e,t)/t.size),A(e,t.size,t),e.id&&!r.$table.attr("aria-describedby")&&((i=(s=t.$container.find(t.cssPageDisplay)).attr("id"))||(i=e.id+"_pager_info",s.attr("id",i)),r.$table.attr("aria-describedby",i)),P(e,t),a&&(L.update(r),A(e,t.size,t),S(e,t),m(e,t),L.debug(r,"pager"))&&console.log("Pager >> Enabled")}var w=this,b=function(e,t,a){var i="addClass",s="removeClass",r=t.cssDisabled,a=!!a,o=a||0===t.page,e=$(e,t),n=a||t.page===e-1||0===e;t.updateArrows&&((a=t.$container.find(t.cssFirst+","+t.cssPrev))[o?i:s](r),a.each(function(){this.ariaDisabled=o}),(a=t.$container.find(t.cssNext+","+t.cssLast))[n?i:s](r),a.each(function(){this.ariaDisabled=n}))},R=function(e,t){var a,i,s,r=e.config,e=r.$table.hasClass("hasFilters");if(e&&!t.ajax)if(L.isEmptyObject(r.cache))t.filteredRows=t.totalRows=r.$tbodies.eq(0).children("tr").not(t.countChildRows?"":"."+r.cssChildRow).length;else for(t.filteredRows=0,s=(a=r.cache[0].normalized).length,i=0;i<s;i++)t.filteredRows+=t.regexRows.test(a[i][r.columns].$row[0].className)?0:1;else e||(t.filteredRows=t.totalRows)},j=function(e,o,t){if(!o.initializing){var a,i,s,r,n,l,g,c=e.config,d=c.namespace+"pager",p=N(o,o.size,"get");if("all"===p&&(p=o.totalRows),o.countChildRows&&(a[a.length]=c.cssChildRow),o.totalPages=Math.ceil(o.totalRows/p),c.totalRows=o.totalRows,I(e,o),R(e,o),c.filteredRows=o.filteredRows,o.filteredPages=Math.ceil(o.filteredRows/p)||0,0<=$(e,o)){if(a=p*o.page>o.filteredRows&&t,o.page=a?o.pageReset||0:o.page,o.startRow=!a&&0===o.filteredRows?0:p*o.page+1,o.endRow=Math.min(o.filteredRows,o.totalRows,p*(o.page+1)),i=o.$container.find(o.cssPageDisplay),g="function"==typeof o.output?o.output(e,o):(g=i.attr("data-pager-output"+(o.filteredRows<o.totalRows?"-filtered":""))||o.output,(o.ajaxData&&o.ajaxData.output&&o.ajaxData.output||g).replace(/\{page([\-+]\d+)?\}/gi,function(e,t){return o.totalPages?o.page+(t?parseInt(t,10):1):0}).replace(/\{\w+(\s*:\s*\w+)?\}/gi,function(e){var t,a,e=e.replace(/[{}\s]/g,""),i=e.split(":"),s=o.ajaxData,r=/(rows?|pages?)$/i.test(e)?0:"";return/(startRow|page)/.test(i[0])&&"input"===i[1]?(t=(""+("page"===i[0]?o.totalPages:o.totalRows)).length,a="page"===i[0]?o.page+1:o.startRow,'<input type="text" class="ts-'+i[0]+'" style="max-width:'+t+'em" value="'+a+'"/>'):1<i.length&&s&&s[i[0]]?s[i[0]][i[1]]:o[e]||(s?s[e]:r)||r})),(s=o.$container.find(o.cssGoto)).length){for(a="",n=(l=z(e,o)).length,r=0;r<n;r++)a+='<option value="'+l[r]+'">'+l[r]+"</option>";s.html(a).val(o.page+1)}i.length&&(i["INPUT"===i[0].nodeName?"val":"html"](g),i.find(".ts-startRow, .ts-page").unbind("change"+d).bind("change"+d,function(){var e=E(this).val(),e=E(this).hasClass("ts-startRow")?Math.floor(e/p)+1:e;c.$table.triggerHandler("pageSet"+d,[e])}))}b(e,o),x(e,o),o.initialized&&!1!==t&&(L.debug(c,"pager")&&console.log("Pager >> Triggering pagerComplete"),c.$table.triggerHandler("pagerComplete",o),o.savePages)&&L.storage&&L.storage(e,o.storageKey,{page:o.page,size:p===o.totalRows?"all":p})}},z=function(e,t){for(var a,i,s=$(e,t)||1,r=5*Math.ceil(s/t.maxOptionSize/5),o=s>t.maxOptionSize,e=t.page+1,n=r,l=s-r,g=[1],c=o?r:1;c<=s;)g[g.length]=c,c+=o?r:1;if(g[g.length]=s,o){for(a=[],s<(l=e+(i=Math.max(Math.floor(t.maxOptionSize/r)-1,5)))&&(l=s),c=n=(n=e-i)<1?1:n;c<=l;c++)a[a.length]=c;r/2<(e=(g=E.grep(g,function(e,t){return E.inArray(e,g)===t})).length)-(i=a.length)&&e+i>t.maxOptionSize&&(n=Math.floor(e/2)-Math.floor(i/2),Array.prototype.splice.apply(g,[n,i])),g=g.concat(a)}return g=E.grep(g,function(e,t){return E.inArray(e,g)===t}).sort(function(e,t){return e-t})},x=function(e,t){var a,i,s=e.config,r=s.$tbodies.eq(0);r.find("tr.pagerSavedHeightSpacer").remove(),t.fixedHeight&&!t.isDisabled&&(a=E.data(e,"pagerSavedHeight"))&&(i=0,1<E(e).css("border-spacing").split(" ").length&&(i=E(e).css("border-spacing").split(" ")[1].replace(/[^-\d\.]/g,"")),5<(a=a-r.height()+i*t.size-i))&&E.data(e,"pagerLastSize")===t.size&&r.children("tr:visible").length<("all"===t.size?t.totalRows:t.size)&&r.append('<tr class="pagerSavedHeightSpacer '+s.selectorRemove.slice(1)+'" style="height:'+a+'px;"></tr>')},P=function(e,t){var a=e.config,i=a.$tbodies.eq(0);i.find("tr.pagerSavedHeightSpacer").remove(),i.children("tr:visible").length||i.append('<tr class="pagerSavedHeightSpacer '+a.selectorRemove.slice(1)+'"><td>&nbsp</td></tr>'),a=i.children("tr").eq(0).height()*("all"===t.size?t.totalRows:t.size),E.data(e,"pagerSavedHeight",a),x(e,t),E.data(e,"pagerLastSize",t.size)},v=function(e,t){if(!t.ajaxUrl){var a,i=0,s=e.config,r=s.$tbodies.eq(0).children("tr"),o=r.length,e="all"===t.size?t.totalRows:t.size,n=t.page*e,l=n+e,g=-1,c=0;for(t.cacheIndex=[],a=0;a<o;a++)t.regexFiltered.test(r[a].className)||(c===n&&r[a].className.match(s.cssChildRow)?r[a].style.display="none":(r[a].style.display=n<=c&&c<l?"":"none",g!==c&&n<=c&&c<l&&(t.cacheIndex[t.cacheIndex.length]=a,g=c),(c+=r[a].className.match(s.cssChildRow+"|"+s.selectorRemove.slice(1))&&!t.countChildRows?0:1)===l&&"none"!==r[a].style.display&&r[a].className.match(L.css.cssHasChild)&&(i=a)));if(0<i&&r[i].className.match(L.css.cssHasChild))for(;++i<o&&r[i].className.match(s.cssChildRow);)r[i].style.display=""}},m=function(e,t){t.size=N(t,t.$container.find(t.cssPageSize).val(),"get"),A(e,t.size,t),b(e,t),t.removeRows||(v(e,t),E(e).bind("sortEnd filterEnd ".split(" ").join(e.config.namespace+"pager "),function(){v(e,t)}))},n=function(s,r){var i,e=function(e,a){var t,i,s=e.config,r=a.ajaxUrl?a.ajaxUrl.replace(/\{page([\-+]\d+)?\}/,function(e,t){return a.page+(t?parseInt(t,10):0)}).replace(/\{size\}/g,a.size):"",o=s.sortList,n=a.currentFilters||E(e).data("lastSearch")||[],l=r.match(/\{\s*sort(?:List)?\s*:\s*(\w*)\s*\}/),g=r.match(/\{\s*filter(?:List)?\s*:\s*(\w*)\s*\}/),c=[];if(l){l=l[1];i=o.length;for(t=0;t<i;t++)c[c.length]=l+"["+o[t][0]+"]="+o[t][1];r=r.replace(/\{\s*sort(?:List)?\s*:\s*(\w*)\s*\}/g,c.length?c.join("&"):l);c=[]}if(g){g=g[1];i=n.length;for(t=0;t<i;t++)if(n[t])c[c.length]=g+"["+t+"]="+encodeURIComponent(n[t]);r=r.replace(/\{\s*filter(?:List)?\s*:\s*(\w*)\s*\}/g,c.length?c.join("&"):g);a.currentFilters=n}if(typeof a.customAjaxUrl==="function")r=a.customAjaxUrl(e,r);if(L.debug(s,"pager"))console.log("Pager >> Ajax url = "+r);return r}(s,r),o=E(document),t=s.config,n=t.namespace+"pager";""!==e&&(t.showProcessing&&L.isProcessing(s,!0),o.bind("ajaxError"+n,function(e,t,a,i){l(null,s,r,t,a,i),o.unbind("ajaxError"+n)}),i=++r.ajaxCounter,r.last.ajaxUrl=e,r.ajaxObject.url=e,r.ajaxObject.success=function(e,t,a){i<r.ajaxCounter||(l(e,s,r,a),o.unbind("ajaxError"+n),"function"==typeof r.oldAjaxSuccess&&r.oldAjaxSuccess(e))},L.debug(t,"pager")&&console.log("Pager >> Ajax initialized",r.ajaxObject),E.ajax(r.ajaxObject))},g=function(e,t,a){var i,s,r,o,n=E(e),l=e.config,g=L.debug(l,"pager"),c=l.$table.hasClass("hasFilters"),d=t&&t.length||0,p="all"===a.size?a.totalRows:a.size,f=a.page*p;if(d<1)g&&console.warn("Pager >> No rows for pager to render");else{if(a.page>=a.totalPages&&u(e,a),a.cacheIndex=[],a.isDisabled=!1,a.initialized&&(g&&console.log("Pager >> Triggering pagerChange"),n.triggerHandler("pagerChange",a)),a.removeRows){for(L.clearTableBody(e),i=L.processTbody(e,l.$tbodies.eq(0),!0),r=s=c?0:f,o=0;o<p&&s<t.length;)c&&a.regexFiltered.test(t[s][0].className)||f<++r&&o<=p&&(o++,a.cacheIndex[a.cacheIndex.length]=s,i.append(t[s])),s++;L.processTbody(e,i,!1)}else v(e,a);j(e,a),e.isUpdating&&(g&&console.log("Pager >> Triggering updateComplete"),n.triggerHandler("updateComplete",[e,!0]))}},y=function(e,t){var a,i,s;for(t.ajax?b(e,t,!0):(E.data(e,"pagerLastPage",t.page),E.data(e,"pagerLastSize",t.size),t.page=0,t.size=t.totalRows,t.totalPages=1,E(e).addClass("pagerDisabled").removeAttr("aria-describedby").find("tr.pagerSavedHeightSpacer").remove(),g(e,e.config.rowsCopy,t),t.isDisabled=!0,L.applyWidget(e),L.debug(e.config,"pager")&&console.log("Pager >> Disabled")),s=(i=t.$container.find(t.cssGoto+","+t.cssPageSize+", .ts-startRow, .ts-page")).length,a=0;a<s;a++)i.eq(a).addClass(t.cssDisabled)[0].disabled=!0,i[a].ariaDisabled=!0},C=function(i){var s=i.config,r=s.pager;L.updateCache(s,function(){var e,t=[],a=i.config.cache[0].normalized;for(r.totalRows=a.length,e=0;e<r.totalRows;e++)t[t.length]=a[e][s.columns].$row;s.rowsCopy=t,S(i,r,!0)})},S=function(e,t,a){var i,s,r,o;if(!t.isDisabled)return i=e.config,s=L.debug(i,"pager"),r=E(e),o=t.last,!1!==a&&t.initialized&&L.isEmptyObject(i.cache)?C(e):void(t.ajax&&L.hasWidget(e,"filter")&&!i.widgetOptions.filter_initialized||(I(e,t),R(e,t),o.currentFilters=""===(o.currentFilters||[]).join("")?[]:o.currentFilters,t.currentFilters=""===(t.currentFilters||[]).join("")?[]:t.currentFilters,o.page===t.page&&o.size===t.size&&o.totalRows===t.totalRows&&(o.currentFilters||[]).join(",")===(t.currentFilters||[]).join(",")&&(o.ajaxUrl||"")===(t.ajaxObject.url||"")&&(o.optAjaxUrl||"")===(t.ajaxUrl||"")&&o.sortList===(i.sortList||[]).join(","))||(s&&console.log("Pager >> Changing to page "+t.page),t.last={page:t.page,size:t.size,sortList:(i.sortList||[]).join(","),totalRows:t.totalRows,currentFilters:t.currentFilters||[],ajaxUrl:t.ajaxObject.url||"",optAjaxUrl:t.ajaxUrl||""},t.ajax?(t.processAjaxOnInit||L.isEmptyObject(t.initialRows)?n:(t.processAjaxOnInit=!0,o=t.initialRows,t.totalRows=void 0!==o.total?o.total:s&&console.error("Pager >> No initial total page set!")||0,t.filteredRows=void 0!==o.filtered?o.filtered:s&&console.error("Pager >> No initial filtered page set!")||0,F))(e,t):t.ajax||g(e,i.rowsCopy,t),E.data(e,"pagerLastPage",t.page),t.initialized&&!1!==a&&(s&&console.log("Pager >> Triggering pageMoved"),r.triggerHandler("pageMoved",t),L.applyWidget(e),e.isUpdating)&&(s&&console.log("Pager >> Triggering updateComplete"),r.triggerHandler("updateComplete",[e,!0]))))},$=function(e,t){return L.hasWidget(e,"filter")?Math.min(t.totalPages,t.filteredPages):t.totalPages},I=function(e,t){e=$(e,t)-1;return t.page=parseInt(t.page,10),(t.page<0||isNaN(t.page))&&(t.page=0),t.page>e&&0<=e&&(t.page=e),t.page},N=function(e,t,a){var i=parseInt(t,10)||e.size||e.settings.size||10;return e.initialized&&(/all/i.test(i+" "+t)||i===e.totalRows)?e.$container.find(e.cssPageSize+' option[value="all"]').length?"all":e.totalRows:"get"===a?i:e.size},A=function(e,t,a){a.size=N(a,t,"get"),a.$container.find(a.cssPageSize).val(a.size),E.data(e,"pagerLastPage",I(e,a)),E.data(e,"pagerLastSize",a.size),a.totalPages="all"===a.size?1:Math.ceil(a.totalRows/a.size),a.filteredPages="all"===a.size?1:Math.ceil(a.filteredRows/a.size)},F=function(e,t){t.initialized=!0,t.initializing=!1,L.debug(e.config,"pager")&&console.log("Pager >> Triggering pagerInitialized"),E(e).triggerHandler("pagerInitialized",t),L.applyWidget(e),j(e,t)},O=function(r,e){var s,o,i=r.config,t=i.widgetOptions,a=L.debug(i,"pager"),n=i.pager=E.extend(!0,{},E.tablesorterPager.defaults,e),l=i.$table,g=i.namespace+"pager",c=n.$container=E(n.container).addClass("tablesorter-pager").show();n.settings=E.extend(!0,{},E.tablesorterPager.defaults,e),a&&console.log("Pager >> Initializing"),n.oldAjaxSuccess=n.oldAjaxSuccess||n.ajaxObject.success,i.appender=w.appender,n.initializing=!0,n.savePages&&L.storage&&(e=L.storage(r,n.storageKey)||{},n.page=(isNaN(e.page)?n:e).page,n.size="all"===e.size?e.size:(isNaN(e.size)?n:e).size||n.setSize||10,A(r,n.size,n)),n.regexRows=new RegExp("("+(t.filter_filteredRow||"filtered")+"|"+i.selectorRemove.slice(1)+"|"+i.cssChildRow+")"),n.regexFiltered=new RegExp(t.filter_filteredRow||"filtered"),l.unbind("filterInit filterStart filterEnd sortEnd disablePager enablePager destroyPager updateComplete pageSize pageSet pageAndSize pagerUpdate refreshComplete ".split(" ").join(g+" ").replace(/\s+/g," ")).bind("filterInit filterStart ".split(" ").join(g+" "),function(e,t){if(n.currentFilters=E.isArray(t)?t:i.$table.data("lastSearch"),n.ajax&&"filterInit"===e.type)return S(r,n,!1);t=L.filter.equalFilters?L.filter.equalFilters(i,i.lastSearch,n.currentFilters):(i.lastSearch||[]).join("")!==(n.currentFilters||[]).join(""),"filterStart"!==e.type||!1===n.pageReset||t||(n.page=n.pageReset)}).bind("filterEnd sortEnd ".split(" ").join(g+" "),function(){n.currentFilters=i.$table.data("lastSearch"),(n.initialized||n.initializing)&&(i.delayInit&&i.rowsCopy&&0===i.rowsCopy.length&&C(r),j(r,n,!1),S(r,n,!1),L.applyWidget(r))}).bind("disablePager"+g,function(e){e.stopPropagation(),y(r,n)}).bind("enablePager"+g,function(e){e.stopPropagation(),h(r,n,!0)}).bind("destroyPager"+g,function(e){var t,a,i,s;e.stopPropagation(),e=n,a=(t=r).config,i=a.namespace+"pager",s=[e.cssFirst,e.cssPrev,e.cssNext,e.cssLast,e.cssGoto,e.cssPageSize].join(","),y(t,e),e.$container.hide().find(s).unbind(i),a.appender=null,a.$table.unbind(i),L.storage&&L.storage(t,e.storageKey,""),delete a.pager,delete a.rowsCopy}).bind("resetToLoadState"+g,function(e){var t;e.stopPropagation(),e=n,(t=r).config.pager=E.extend(!0,{},E.tablesorterPager.defaults,e.settings),O(t,e.settings)}).bind("updateComplete"+g,function(e,t,a){e.stopPropagation(),!t||a||n.ajax||(e=i.$tbodies.eq(0).children("tr").not(i.selectorRemove),n.totalRows=e.length-(n.countChildRows?0:e.filter("."+i.cssChildRow).length),n.totalPages="all"===n.size?1:Math.ceil(n.totalRows/n.size),e.length&&i.rowsCopy&&0===i.rowsCopy.length&&C(t),n.page>=n.totalPages&&u(t,n),v(t,n),P(t,n),j(t,n,!0))}).bind("pageSize refreshComplete ".split(" ").join(g+" "),function(e,t){e.stopPropagation(),A(r,N(n,t,"get"),n),S(r,n),v(r,n),j(r,n,!1)}).bind("pageSet pagerUpdate ".split(" ").join(g+" "),function(e,t){e.stopPropagation(),"pagerUpdate"===e.type&&(t=void 0===t?n.page+1:t,n.last.page=!0),n.page=(parseInt(t,10)||1)-1,S(r,n,!0),j(r,n,!1)}).bind("pageAndSize"+g,function(e,t,a){e.stopPropagation(),n.page=(parseInt(t,10)||1)-1,A(r,N(n,a,"get"),n),S(r,n,!0),v(r,n),j(r,n,!1)}),s=[n.cssFirst,n.cssPrev,n.cssNext,n.cssLast],o=[d,f,p,u],a&&!c.length&&console.warn('Pager >> "container" not found'),c.find(s.join(",")).attr("tabindex",0).unbind("click"+g).bind("click"+g,function(e){e.stopPropagation();var t,a=E(this),i=s.length;if(!a.hasClass(n.cssDisabled))for(t=0;t<i;t++)if(a.is(s[t])){o[t](r,n);break}}),(e=c.find(n.cssGoto)).length?e.unbind("change"+g).bind("change"+g,function(){n.page=E(this).val()-1,S(r,n,!0),j(r,n,!1)}):a&&console.warn('Pager >> "goto" selector not found'),(e=c.find(n.cssPageSize)).length?(e.find("option").removeAttr("selected"),e.unbind("change"+g).bind("change"+g,function(){var e;return E(this).hasClass(n.cssDisabled)||(e=E(this).val(),A(r,e,n),S(r,n),P(r,n)),!1})):a&&console.warn('Pager >> "size" selector not found'),n.initialized=!1,l.triggerHandler("pagerBeforeInitialized",n),h(r,n,!1),("string"==typeof n.ajaxUrl?(n.ajax=!0,i.widgetOptions.filter_serversideFiltering=!0,i.serverSideSorting=!0,S):(n.ajax=!1,L.appendCache(i,!0),m))(r,n),n.ajax||n.initialized||(n.initializing=!1,n.initialized=!0,A(r,n.size,n),S(r,n),a&&console.log("Pager >> Triggering pagerInitialized"),i.$table.triggerHandler("pagerInitialized",n),i.widgetOptions.filter_initialized&&L.hasWidget(r,"filter"))||j(r,n,!1),i.widgetInit.pager=!0};w.appender=function(e,t){var a=e.config,i=a.pager;i.ajax||(a.rowsCopy=t,i.totalRows=(i.countChildRows?a.$tbodies.eq(0).children("tr"):t).length,i.size=E.data(e,"pagerLastSize")||i.size||i.settings.size||10,i.totalPages="all"===i.size?1:Math.ceil(i.totalRows/i.size),g(e,t,i),j(e,i,!1))},w.construct=function(e){return this.each(function(){this.config&&this.hasInitialized&&O(this,e)})}}}),L.showError=function(e,t,a,i){function s(){r.$table.find("thead").find(r.selectorRemove).remove()}var e=E(e),r=e[0].config,o=r&&r.widgetOptions,n=r.pager&&r.pager.cssErrorRow||o&&o.pager_css&&o.pager_css.errorRow||"tablesorter-errorRow",l=typeof t,g=!0,c="";if(e.length){if("function"==typeof r.pager.ajaxError){if(!1===(g=r.pager.ajaxError(r,t,a,i)))return s();c=g}else if("function"==typeof o.pager_ajaxError){if(!1===(g=o.pager_ajaxError(r,t,a,i)))return s();c=g}if(""===c)if("object"==l)c=0===t.status?"Not connected, verify Network":404===t.status?"Requested page not found [404]":500===t.status?"Internal Server Error [500]":"parsererror"===i?"Requested JSON parse failed":"timeout"===i?"Time out error":"abort"===i?"Ajax Request aborted":"Uncaught error: "+t.statusText+" ["+t.status+"]";else{if("string"!=l)return s();c=t}E(/tr\>/.test(c)?c:'<tr><td colspan="'+r.columns+'">'+c+"</td></tr>").click(function(){E(this).remove()}).appendTo(r.$table.find("thead:first")).addClass(n+" "+r.selectorRemove.slice(1)).attr({role:"alert","aria-live":"assertive"})}else console.error("tablesorter showError: no table parameter passed")},E.fn.extend({tablesorterPager:E.tablesorterPager.construct})}(jQuery);return jQuery;}));
```
