## Supplementary Data SD1 for "Inferring binding specificities of human transcription factors with the wisdom of crowds": semver-mod.min.js

```
/*! Modified semver.js for node.js (v4.3.3, 3/27/2015) */
!function(){var n={exports:{}}.exports=b,c="object"==typeof process&&process.env&&process.env.NODE_DEBUG&&/\bsemver\b/i.test(process.env.NODE_DEBUG)?function(){var e=Array.prototype.slice.call(arguments,0);e.unshift("SEMVER"),console.log.apply(console,e)}:function(){},t=(n.SEMVER_SPEC_VERSION="2.0.0",256),i=Number.MAX_SAFE_INTEGER||9007199254740991,s=n.re=[],e=n.src=[],r=0,o=(r++,e[0]="0|[1-9]\\d*",r++,e[1]="[0-9]+",r++,e[2]="\\d*[a-zA-Z-][a-zA-Z0-9-]*",r++,e[3]="("+e[0]+")\\.("+e[0]+")\\.("+e[0]+")",r++,e[4]="("+e[1]+")\\.("+e[1]+")\\.("+e[1]+")",r++,e[5]="(?:"+e[0]+"|"+e[2]+")",r++,e[6]="(?:"+e[1]+"|"+e[2]+")",r++,e[7]="(?:-("+e[5]+"(?:\\."+e[5]+")*))",r++,e[8]="(?:-?("+e[6]+"(?:\\."+e[6]+")*))",r++,e[9]="[0-9A-Za-z-]+",r++,e[10]="(?:\\+("+e[9]+"(?:\\."+e[9]+")*))",r++),a="v?"+e[3]+e[7]+"?"+e[10]+"?",p=(e[o]="^"+a+"$","[v=\\s]*"+e[4]+e[8]+"?"+e[10]+"?"),u=r++,h=(e[u]="^"+p+"$",r++),f=(e[h]="((?:<|>)?=?)",r++,e[14]=e[1]+"|x|X|\\*",r++,e[15]=e[0]+"|x|X|\\*",r++),l=(e[f]="[v=\\s]*("+e[15]+")(?:\\.("+e[15]+")(?:\\.("+e[15]+")(?:"+e[7]+")?"+e[10]+"?)?)?",r++),m=(e[l]="[v=\\s]*("+e[14]+")(?:\\.("+e[14]+")(?:\\.("+e[14]+")(?:"+e[8]+")?"+e[10]+"?)?)?",r++),v=(e[m]="^"+e[h]+"\\s*"+e[f]+"$",r++),g=(e[v]="^"+e[h]+"\\s*"+e[l]+"$",r++,e[20]="(?:~>?)",r++,e[21]="(\\s*)"+e[20]+"\\s+",s[21]=new RegExp(e[21],"g"),r++),w=(e[g]="^"+e[20]+e[f]+"$",r++),y=(e[w]="^"+e[20]+e[l]+"$",r++,e[24]="(?:\\^)",r++,e[25]="(\\s*)"+e[24]+"\\s+",s[25]=new RegExp(e[25],"g"),r++),d=(e[y]="^"+e[24]+e[f]+"$",r++),D=(e[d]="^"+e[24]+e[l]+"$",r++,e[28]="^"+e[h]+"\\s*("+p+")$|^$",r++,e[29]="^"+e[h]+"\\s*("+a+")$|^$",r++,e[30]="(\\s*)"+e[h]+"\\s*("+p+"|"+e[f]+")",s[30]=new RegExp(e[30],"g"),r++,e[31]="^\\s*("+e[f]+")\\s+-\\s+("+e[f]+")\\s*$",r++,e[32]="^\\s*("+e[l]+")\\s+-\\s+("+e[l]+")\\s*$",+r);e[D]="(<|>)?=?\\s*\\*";for(var j=0;j<34;j++)c(j,e[j]),s[j]||(s[j]=new RegExp(e[j]));function E(e,r){if(e instanceof b)return e;if("string"!=typeof e)return null;if(e.length>t)return null;if(!(r?s[u]:s[o]).test(e))return null;try{return new b(e,r)}catch(e){return null}}function b(e,r){if(e instanceof b){if(e.loose===r)return e;e=e.version}else if("string"!=typeof e)throw new TypeError("Invalid Version: "+e);if(e.length>t)throw new TypeError("version is longer than "+t+" characters");if(!(this instanceof b))return new b(e,r);c("SemVer",e,r),this.loose=r;r=e.trim().match(r?s[u]:s[o]);if(!r)throw new TypeError("Invalid Version: "+e);if(this.raw=e,this.major=+r[1],this.minor=+r[2],this.patch=+r[3],this.major>i||this.major<0)throw new TypeError("Invalid major version");if(this.minor>i||this.minor<0)throw new TypeError("Invalid minor version");if(this.patch>i||this.patch<0)throw new TypeError("Invalid patch version");r[4]?this.prerelease=r[4].split(".").map(function(e){if(/^[0-9]+$/.test(e)){var r=+e;if(0<=r&&r<i)return r}return e}):this.prerelease=[],this.build=r[5]?r[5].split("."):[],this.format()}n.parse=E,n.valid=function(e,r){e=E(e,r);return e?e.version:null},n.clean=function(e,r){e=E(e.trim().replace(/^[=v]+/,""),r);return e?e.version:null},(window.semver=n.SemVer=b).prototype.format=function(){return this.version=this.major+"."+this.minor+"."+this.patch,this.prerelease.length&&(this.version+="-"+this.prerelease.join(".")),this.version},b.prototype.inspect=function(){return'<SemVer "'+this+'">'},b.prototype.toString=function(){return this.version},b.prototype.compare=function(e){return c("SemVer.compare",this.version,this.loose,e),e instanceof b||(e=new b(e,this.loose)),this.compareMain(e)||this.comparePre(e)},b.prototype.compareMain=function(e){return e instanceof b||(e=new b(e,this.loose)),$(this.major,e.major)||$(this.minor,e.minor)||$(this.patch,e.patch)},b.prototype.comparePre=function(e){if(e instanceof b||(e=new b(e,this.loose)),this.prerelease.length&&!e.prerelease.length)return-1;if(!this.prerelease.length&&e.prerelease.length)return 1;if(!this.prerelease.length&&!e.prerelease.length)return 0;var r=0;do{var t=this.prerelease[r],n=e.prerelease[r];if(c("prerelease compare",r,t,n),void 0===t&&void 0===n)return 0;if(void 0===n)return 1;if(void 0===t)return-1;if(t!==n)return $(t,n)}while(++r)},b.prototype.inc=function(e,r){switch(e){case"premajor":this.prerelease.length=0,this.patch=0,this.minor=0,this.major++,this.inc("pre",r);break;case"preminor":this.prerelease.length=0,this.patch=0,this.minor++,this.inc("pre",r);break;case"prepatch":this.prerelease.length=0,this.inc("patch",r),this.inc("pre",r);break;case"prerelease":0===this.prerelease.length&&this.inc("patch",r),this.inc("pre",r);break;case"major":0===this.minor&&0===this.patch&&0!==this.prerelease.length||this.major++,this.minor=0,this.patch=0,this.prerelease=[];break;case"minor":0===this.patch&&0!==this.prerelease.length||this.minor++,this.patch=0,this.prerelease=[];break;case"patch":0===this.prerelease.length&&this.patch++,this.prerelease=[];break;case"pre":if(0===this.prerelease.length)this.prerelease=[0];else{for(var t=this.prerelease.length;0<=--t;)"number"==typeof this.prerelease[t]&&(this.prerelease[t]++,t=-2);-1===t&&this.prerelease.push(0)}r&&(this.prerelease[0]!==r||isNaN(this.prerelease[1]))&&(this.prerelease=[r,0]);break;default:throw new Error("invalid increment argument: "+e)}return this.format(),this},n.inc=function(e,r,t,n){"string"==typeof t&&(n=t,t=void 0);try{return new b(e,t).inc(r,n).version}catch(e){return null}},n.diff=function(e,r){{if(x(e,r))return null;var t=E(e),n=E(r);if(t.prerelease.length||n.prerelease.length){for(var i in t)if(("major"===i||"minor"===i||"patch"===i)&&t[i]!==n[i])return"pre"+i;return"prerelease"}for(i in t)if(("major"===i||"minor"===i||"patch"===i)&&t[i]!==n[i])return i}},n.compareIdentifiers=$;var X=/^[0-9]+$/;function $(e,r){var t=X.test(e),n=X.test(r);return t&&n&&(e=+e,r=+r),t&&!n?-1:n&&!t?1:e<r?-1:r<e?1:0}function k(e,r,t){return new b(e,t).compare(r)}function z(e,r,t){return k(r,e,t)}function R(e,r,t){return 0<k(e,r,t)}function S(e,r,t){return k(e,r,t)<0}function x(e,r,t){return 0===k(e,r,t)}function G(e,r,t){return 0!==k(e,r,t)}function I(e,r,t){return 0<=k(e,r,t)}function T(e,r,t){return k(e,r,t)<=0}function O(e,r,t,n){var i;switch(r){case"===":i=(e="object"==typeof e?e.version:e)===(t="object"==typeof t?t.version:t);break;case"!==":i=(e="object"==typeof e?e.version:e)!==(t="object"==typeof t?t.version:t);break;case"":case"=":case"==":i=x(e,t,n);break;case"!=":i=G(e,t,n);break;case">":i=R(e,t,n);break;case">=":i=I(e,t,n);break;case"<":i=S(e,t,n);break;case"<=":i=T(e,t,n);break;default:throw new TypeError("Invalid operator: "+r)}return i}function V(e,r){if(e instanceof V){if(e.loose===r)return e;e=e.value}if(!(this instanceof V))return new V(e,r);c("comparator",e,r),this.loose=r,this.parse(e),this.semver===A?this.value="":this.value=this.operator+this.semver.version,c("comp",this)}n.rcompareIdentifiers=function(e,r){return $(r,e)},n.major=function(e,r){return new b(e,r).major},n.minor=function(e,r){return new b(e,r).minor},n.patch=function(e,r){return new b(e,r).patch},n.compare=k,n.compareLoose=function(e,r){return k(e,r,!0)},n.rcompare=z,n.sort=function(e,t){return e.sort(function(e,r){return n.compare(e,r,t)})},n.rsort=function(e,t){return e.sort(function(e,r){return n.rcompare(e,r,t)})},n.gt=R,n.lt=S,n.eq=x,n.neq=G,n.gte=I,n.lte=T,n.cmp=O,n.Comparator=V;var A={};function C(e,r){if(e instanceof C&&e.loose===r)return e;if(!(this instanceof C))return new C(e,r);if(this.loose=r,this.raw=e,this.set=e.split(/\s*\|\|\s*/).map(function(e){return this.parseRange(e.trim())},this).filter(function(e){return e.length}),!this.set.length)throw new TypeError("Invalid SemVer Range: "+e);this.format()}function N(e){return!e||"x"===e.toLowerCase()||"*"===e}function P(e,r,t,n,i,o,s,a,p,c,u,h,f){return((r=N(t)?"":N(n)?">="+t+".0.0":N(i)?">="+t+"."+n+".0":">="+r)+" "+(a=N(p)?"":N(c)?"<"+(+p+1)+".0.0":N(u)?"<"+p+"."+(+c+1)+".0":h?"<="+p+"."+c+"."+u+"-"+h:"<="+a)).trim()}function M(e,r,t){try{r=new C(r,t)}catch(e){return!1}return r.test(e)}function _(e,r,t,n){var i,o,s,a,p;switch(e=new b(e,n),r=new C(r,n),t){case">":i=R,o=T,s=S,a=">",p=">=";break;case"<":i=S,o=I,s=R,a="<",p="<=";break;default:throw new TypeError('Must provide a hilo val of "<" or ">"')}if(M(e,r,n))return!1;for(var c=0;c<r.set.length;++c){var u=r.set[c],h=null,f=null;if(u.forEach(function(e){h=h||e,f=f||e,i(e.semver,h.semver,n)?h=e:s(e.semver,f.semver,n)&&(f=e)}),h.operator===a||h.operator===p)return!1;if((!f.operator||f.operator===a)&&o(e,f.semver))return!1;if(f.operator===p&&s(e,f.semver))return!1}return!0}V.prototype.parse=function(e){var r=this.loose?s[28]:s[29],r=e.match(r);if(!r)throw new TypeError("Invalid comparator: "+e);this.operator=r[1],"="===this.operator&&(this.operator=""),r[2]?this.semver=new b(r[2],this.loose):this.semver=A},V.prototype.inspect=function(){return'<SemVer Comparator "'+this+'">'},V.prototype.toString=function(){return this.value},V.prototype.test=function(e){return c("Comparator.test",e,this.loose),this.semver===A||O(e="string"==typeof e?new b(e,this.loose):e,this.operator,this.semver,this.loose)},(n.Range=C).prototype.inspect=function(){return'<SemVer Range "'+this.range+'">'},C.prototype.format=function(){return this.range=this.set.map(function(e){return e.join(" ").trim()}).join("||").trim(),this.range},C.prototype.toString=function(){return this.range},C.prototype.parseRange=function(e){var t=this.loose,r=(e=e.trim(),c("range",e,t),t?s[32]:s[31]),n=(e=e.replace(r,P),c("hyphen replace",e),e=e.replace(s[30],"$1$2$3"),c("comparator trim",e,s[30]),e=(e=(e=e.replace(s[21],"$1~")).replace(s[25],"$1^")).split(/\s+/).join(" "),t?s[28]:s[29]),r=e.split(" ").map(function(e){return r=t,c("comp",e=e),e=function(e,r){return e.trim().split(/\s+/).map(function(e){var o=e,e=r;return c("caret",o,e),e=e?s[d]:s[y],o.replace(e,function(e,r,t,n,i){return c("caret",o,e,r,t,n,i),e=N(r)?"":N(t)?">="+r+".0.0 <"+(+r+1)+".0.0":N(n)?"0"===r?">="+r+"."+t+".0 <"+r+"."+(+t+1)+".0":">="+r+"."+t+".0 <"+(+r+1)+".0.0":i?(c("replaceCaret pr",i),"-"!==i.charAt(0)&&(i="-"+i),"0"===r?"0"===t?">="+r+"."+t+"."+n+i+" <"+r+"."+t+"."+(+n+1):">="+r+"."+t+"."+n+i+" <"+r+"."+(+t+1)+".0":">="+r+"."+t+"."+n+i+" <"+(+r+1)+".0.0"):(c("no pr"),"0"===r?"0"===t?">="+r+"."+t+"."+n+" <"+r+"."+t+"."+(+n+1):">="+r+"."+t+"."+n+" <"+r+"."+(+t+1)+".0":">="+r+"."+t+"."+n+" <"+(+r+1)+".0.0"),c("caret return",e),e})}).join(" ")}(e,r),c("caret",e),e=function(e,r){return e.trim().split(/\s+/).map(function(e){var o=e,e=r;return e=r?s[w]:s[g],o.replace(e,function(e,r,t,n,i){return c("tilde",o,e,r,t,n,i),e=N(r)?"":N(t)?">="+r+".0.0 <"+(+r+1)+".0.0":N(n)?">="+r+"."+t+".0 <"+r+"."+(+t+1)+".0":i?(c("replaceTilde pr",i),">="+r+"."+t+"."+n+(i="-"!==i.charAt(0)?"-"+i:i)+" <"+r+"."+(+t+1)+".0"):">="+r+"."+t+"."+n+" <"+r+"."+(+t+1)+".0",c("tilde return",e),e})}).join(" ")}(e,r),c("tildes",e),e=function(e,r){return c("replaceXRanges",e,r),e.split(/\s+/).map(function(e){var p=e,e=r;return p=p.trim(),e=e?s[v]:s[m],p.replace(e,function(e,r,t,n,i,o){c("xRange",p,e,r,t,n,i,o);var o=N(t),s=o||N(n),a=s||N(i);return"="===r&&a&&(r=""),o?e=">"===r||"<"===r?"<0.0.0":"*":r&&a?(s&&(n=0),a&&(i=0),">"===r?(r=">=",s?(t=+t+1,i=n=0):a&&(n=+n+1,i=0)):"<="===r&&(r="<",s?t=+t+1:n=+n+1),e=r+t+"."+n+"."+i):s?e=">="+t+".0.0 <"+(+t+1)+".0.0":a&&(e=">="+t+"."+n+".0 <"+t+"."+(+n+1)+".0"),c("xRange return",e),e})}).join(" ")}(e,r),c("xrange",e),e=function(e,r){return c("replaceStars",e,r),e.trim().replace(s[D],"")}(e,r),c("stars",e),e;var r}).join(" ").split(/\s+/);return r=(r=this.loose?r.filter(function(e){return!!e.match(n)}):r).map(function(e){return new V(e,t)})},n.toComparators=function(e,r){return new C(e,r).set.map(function(e){return e.map(function(e){return e.value}).join(" ").trim().split(" ")})},C.prototype.test=function(e){if(e){"string"==typeof e&&(e=new b(e,this.loose));for(var r=0;r<this.set.length;r++)if(function(e,r){for(var t=0;t<e.length;t++)if(!e[t].test(r))return;if(r.prerelease.length){for(t=0;t<e.length;t++){if(c(e[t].semver),e[t].semver===A)return 1;if(0<e[t].semver.prerelease.length){var n=e[t].semver;if(n.major===r.major&&n.minor===r.minor&&n.patch===r.patch)return 1}}return}return 1}(this.set[r],e))return!0}return!1},n.satisfies=M,n.maxSatisfying=function(e,r,t){return e.filter(function(e){return M(e,r,t)}).sort(function(e,r){return z(e,r,t)})[0]||null},n.validRange=function(e,r){try{return new C(e,r).range||"*"}catch(e){return null}},n.ltr=function(e,r,t){return _(e,r,"<",t)},n.gtr=function(e,r,t){return _(e,r,">",t)},n.outside=_,"function"==typeof define&&define.amd&&define(n)}();
```
