## Supplementary Data SD1 for "Inferring binding specificities of human transcription factors with the wisdom of crowds": jquery.tablesorter.combined.js

```
/*! tablesorter (FORK) - updated 2024-08-13 (v2.32.0)*/
/* Includes widgets ( storage,uitheme,columns,filter,stickyHeaders,resizable,saveSort ) */
(function(factory){if (typeof define === 'function' && define.amd){define(['jquery'], factory);} else if (typeof module === 'object' && typeof module.exports === 'object'){module.exports = factory(require('jquery'));} else {factory(jQuery);}}(function(jQuery) {
/*! TableSorter (FORK) v2.32.0 *//*
* Client-side table sorting with ease!
* @requires jQuery v1.2.6+
*
* Copyright (c) 2007 Christian Bach
* fork maintained by Rob Garrison
*
* Examples and original docs at: http://tablesorter.com
* Dual licensed under the MIT and GPL licenses:
* http://www.opensource.org/licenses/mit-license.php
* http://www.gnu.org/licenses/gpl.html
*
* @type jQuery
* @name tablesorter (FORK)
* @cat Plugins/Tablesorter
* @author Christian Bach -
* @contributor Rob Garrison - https://github.com/Mottie/tablesorter
* @docs (fork) - https://mottie.github.io/tablesorter/docs/
*/
/*jshint browser:true, jquery:true, unused:false, expr: true */
;( function( $ ) {
	'use strict';
	var ts = $.tablesorter = {

		version : '2.32.0',

		parsers : [],
		widgets : [],
		defaults : {

			// *** appearance
			theme            : 'default',  // adds tablesorter-{theme} to the table for styling
			widthFixed       : false,      // adds colgroup to fix widths of columns
			showProcessing   : false,      // show an indeterminate timer icon in the header when the table is sorted or filtered.

			headerTemplate   : '{content}',// header layout template (HTML ok); {content} = innerHTML, {icon} = <i></i> // class from cssIcon
			onRenderTemplate : null,       // function( index, template ) { return template; }, // template is a string
			onRenderHeader   : null,       // function( index ) {}, // nothing to return

			// *** functionality
			cancelSelection  : true,       // prevent text selection in the header
			tabIndex         : true,       // add tabindex to header for keyboard accessibility
			dateFormat       : 'mmddyyyy', // other options: 'ddmmyyy' or 'yyyymmdd'
			sortMultiSortKey : 'shiftKey', // key used to select additional columns
			sortResetKey     : 'ctrlKey',  // key used to remove sorting on a column
			usNumberFormat   : true,       // false for German '1.234.567,89' or French '1 234 567,89'
			delayInit        : false,      // if false, the parsed table contents will not update until the first sort
			serverSideSorting: false,      // if true, server-side sorting should be performed because client-side sorting will be disabled, but the ui and events will still be used.
			resort           : true,       // default setting to trigger a resort after an 'update', 'addRows', 'updateCell', etc has completed

			// *** sort options
			headers          : null,       // set sorter, string, empty, locked order, sortInitialOrder, filter, etc.
			ignoreCase       : true,       // ignore case while sorting
			sortForce        : null,       // column(s) first sorted; always applied
			sortList         : [],         // Initial sort order; applied initially; updated when manually sorted
			sortAppend       : null,       // column(s) sorted last; always applied
			sortStable       : false,      // when sorting two rows with exactly the same content, the original sort order is maintained

			sortInitialOrder : 'asc',      // sort direction on first click
			sortLocaleCompare: false,      // replace equivalent character (accented characters)
			sortReset        : false,      // third click on the header will reset column to default - unsorted
			sortRestart      : false,      // restart sort to 'sortInitialOrder' when clicking on previously unsorted columns

			emptyTo          : 'bottom',   // sort empty cell to bottom, top, none, zero, emptyMax, emptyMin
			stringTo         : 'max',      // sort strings in numerical column as max, min, top, bottom, zero
			duplicateSpan    : true,       // colspan cells in the tbody will have duplicated content in the cache for each spanned column
			textExtraction   : 'basic',    // text extraction method/function - function( node, table, cellIndex ) {}
			textAttribute    : 'data-text',// data-attribute that contains alternate cell text (used in default textExtraction function)
			textSorter       : null,       // choose overall or specific column sorter function( a, b, direction, table, columnIndex ) [alt: ts.sortText]
			numberSorter     : null,       // choose overall numeric sorter function( a, b, direction, maxColumnValue )

			// *** widget options
			initWidgets      : true,       // apply widgets on tablesorter initialization
			widgetClass      : 'widget-{name}', // table class name template to match to include a widget
			widgets          : [],         // method to add widgets, e.g. widgets: ['zebra']
			widgetOptions    : {
				zebra : [ 'even', 'odd' ]  // zebra widget alternating row class names
			},

			// *** callbacks
			initialized      : null,       // function( table ) {},

			// *** extra css class names
			tableClass       : '',
			cssAsc           : '',
			cssDesc          : '',
			cssNone          : '',
			cssHeader        : '',
			cssHeaderRow     : '',
			cssProcessing    : '', // processing icon applied to header during sort/filter

			cssChildRow      : 'tablesorter-childRow', // class name indiciating that a row is to be attached to its parent
			cssInfoBlock     : 'tablesorter-infoOnly', // don't sort tbody with this class name (only one class name allowed here!)
			cssNoSort        : 'tablesorter-noSort',   // class name added to element inside header; clicking on it won't cause a sort
			cssIgnoreRow     : 'tablesorter-ignoreRow',// header row to ignore; cells within this row will not be added to c.$headers

			cssIcon          : 'tablesorter-icon', // if this class does not exist, the {icon} will not be added from the headerTemplate
			cssIconNone      : '', // class name added to the icon when there is no column sort
			cssIconAsc       : '', // class name added to the icon when the column has an ascending sort
			cssIconDesc      : '', // class name added to the icon when the column has a descending sort
			cssIconDisabled  : '', // class name added to the icon when the column has a disabled sort

			// *** events
			pointerClick     : 'click',
			pointerDown      : 'mousedown',
			pointerUp        : 'mouseup',

			// *** selectors
			selectorHeaders  : '> thead th, > thead td',
			selectorSort     : 'th, td', // jQuery selector of content within selectorHeaders that is clickable to trigger a sort
			selectorRemove   : '.remove-me',

			// *** advanced
			debug            : false,

			// *** Internal variables
			headerList: [],
			empties: {},
			strings: {},
			parsers: [],

			// *** parser options for validator; values must be falsy!
			globalize: 0,
			imgAttr: 0

			// removed: widgetZebra: { css: ['even', 'odd'] }

		},

		// internal css classes - these will ALWAYS be added to
		// the table and MUST only contain one class name - fixes #381
		css : {
			table      : 'tablesorter',
			cssHasChild: 'tablesorter-hasChildRow',
			childRow   : 'tablesorter-childRow',
			colgroup   : 'tablesorter-colgroup',
			header     : 'tablesorter-header',
			headerRow  : 'tablesorter-headerRow',
			headerIn   : 'tablesorter-header-inner',
			icon       : 'tablesorter-icon',
			processing : 'tablesorter-processing',
			sortAsc    : 'tablesorter-headerAsc',
			sortDesc   : 'tablesorter-headerDesc',
			sortNone   : 'tablesorter-headerUnSorted'
		},

		// labels applied to sortable headers for accessibility (aria) support
		language : {
			sortAsc      : 'Ascending sort applied, ',
			sortDesc     : 'Descending sort applied, ',
			sortNone     : 'No sort applied, ',
			sortDisabled : 'sorting is disabled',
			nextAsc      : 'activate to apply an ascending sort',
			nextDesc     : 'activate to apply a descending sort',
			nextNone     : 'activate to remove the sort'
		},

		regex : {
			templateContent : /\{content\}/g,
			templateIcon    : /\{icon\}/g,
			templateName    : /\{name\}/i,
			spaces          : /\s+/g,
			nonWord         : /\W/g,
			formElements    : /(input|select|button|textarea)/i,

			// *** sort functions ***
			// regex used in natural sort
			// chunk/tokenize numbers & letters
			chunk  : /(^([+\-]?(?:\d*)(?:\.\d*)?(?:[eE][+\-]?\d+)?)?$|^0x[0-9a-f]+$|\d+)/gi,
			// replace chunks @ ends
			chunks : /(^\\0|\\0$)/,
			hex    : /^0x[0-9a-f]+$/i,

			// *** formatFloat ***
			comma                : /,/g,
			digitNonUS           : /[\s|\.]/g,
			digitNegativeTest    : /^\s*\([.\d]+\)/,
			digitNegativeReplace : /^\s*\(([.\d]+)\)/,

			// *** isDigit ***
			digitTest    : /^[\-+(]?\d+[)]?$/,
			digitReplace : /[,.'"\s]/g

		},

		// digit sort, text location
		string : {
			max      : 1,
			min      : -1,
			emptymin : 1,
			emptymax : -1,
			zero     : 0,
			none     : 0,
			'null'   : 0,
			top      : true,
			bottom   : false
		},

		keyCodes : {
			enter : 13
		},

		// placeholder date parser data (globalize)
		dates : {},

		// These methods can be applied on table.config instance
		instanceMethods : {},

		/*
		▄█████ ██████ ██████ ██  ██ █████▄
		▀█▄    ██▄▄     ██   ██  ██ ██▄▄██
		   ▀█▄ ██▀▀     ██   ██  ██ ██▀▀▀
		█████▀ ██████   ██   ▀████▀ ██
		*/

		setup : function( table, c ) {
			// if no thead or tbody, or tablesorter is already present, quit
			if ( !table || !table.tHead || table.tBodies.length === 0 || table.hasInitialized === true ) {
				if ( ts.debug(c, 'core') ) {
					if ( table.hasInitialized ) {
						console.warn( 'Stopping initialization. Tablesorter has already been initialized' );
					} else {
						console.error( 'Stopping initialization! No table, thead or tbody', table );
					}
				}
				return;
			}

			var tmp = '',
				$table = $( table ),
				meta = $.metadata;
			// initialization flag
			table.hasInitialized = false;
			// table is being processed flag
			table.isProcessing = true;
			// make sure to store the config object
			table.config = c;
			// save the settings where they read
			$.data( table, 'tablesorter', c );
			if ( ts.debug(c, 'core') ) {
				console[ console.group ? 'group' : 'log' ]( 'Initializing tablesorter v' + ts.version );
				$.data( table, 'startoveralltimer', new Date() );
			}

			// removing this in version 3 (only supports jQuery 1.7+)
			c.supportsDataObject = ( function( version ) {
				version[ 0 ] = parseInt( version[ 0 ], 10 );
				return ( version[ 0 ] > 1 ) || ( version[ 0 ] === 1 && parseInt( version[ 1 ], 10 ) >= 4 );
			})( $.fn.jquery.split( '.' ) );
			// ensure case insensitivity
			c.emptyTo = c.emptyTo.toLowerCase();
			c.stringTo = c.stringTo.toLowerCase();
			c.last = { sortList : [], clickedIndex : -1 };
			// add table theme class only if there isn't already one there
			if ( !/tablesorter\-/.test( $table.attr( 'class' ) ) ) {
				tmp = ( c.theme !== '' ? ' tablesorter-' + c.theme : '' );
			}

			// give the table a unique id, which will be used in namespace binding
			if ( !c.namespace ) {
				c.namespace = '.tablesorter' + Math.random().toString( 16 ).slice( 2 );
			} else {
				// make sure namespace starts with a period & doesn't have weird characters
				c.namespace = '.' + c.namespace.replace( ts.regex.nonWord, '' );
			}

			c.table = table;
			c.$table = $table
				// add namespace to table to allow bindings on extra elements to target
				// the parent table (e.g. parser-input-select)
				.addClass( ts.css.table + ' ' + c.tableClass + tmp + ' ' + c.namespace.slice(1) )
				.attr( 'role', 'grid' );
			c.$headers = $table.find( c.selectorHeaders );

			c.$table.children().children( 'tr' ).attr( 'role', 'row' );
			c.$tbodies = $table.children( 'tbody:not(.' + c.cssInfoBlock + ')' ).attr({
				'aria-live' : 'polite',
				'aria-relevant' : 'all'
			});
			if ( c.$table.children( 'caption' ).length ) {
				tmp = c.$table.children( 'caption' )[ 0 ];
				if ( !tmp.id ) { tmp.id = c.namespace.slice( 1 ) + 'caption'; }
				c.$table.attr( 'aria-labelledby', tmp.id );
			}
			c.widgetInit = {}; // keep a list of initialized widgets
			// change textExtraction via data-attribute
			c.textExtraction = c.$table.attr( 'data-text-extraction' ) || c.textExtraction || 'basic';
			// build headers
			ts.buildHeaders( c );
			// fixate columns if the users supplies the fixedWidth option
			// do this after theme has been applied
			ts.fixColumnWidth( table );
			// add widgets from class name
			ts.addWidgetFromClass( table );
			// add widget options before parsing (e.g. grouping widget has parser settings)
			ts.applyWidgetOptions( table );
			// try to auto detect column type, and store in tables config
			ts.setupParsers( c );
			// start total row count at zero
			c.totalRows = 0;
			// only validate options while debugging. See #1528
			if (c.debug) {
				ts.validateOptions( c );
			}
			// build the cache for the tbody cells
			// delayInit will delay building the cache until the user starts a sort
			if ( !c.delayInit ) { ts.buildCache( c ); }
			// bind all header events and methods
			ts.bindEvents( table, c.$headers, true );
			ts.bindMethods( c );
			// get sort list from jQuery data or metadata
			// in jQuery < 1.4, an error occurs when calling $table.data()
			if ( c.supportsDataObject && typeof $table.data().sortlist !== 'undefined' ) {
				c.sortList = $table.data().sortlist;
			} else if ( meta && ( $table.metadata() && $table.metadata().sortlist ) ) {
				c.sortList = $table.metadata().sortlist;
			}
			// apply widget init code
			ts.applyWidget( table, true );
			// if user has supplied a sort list to constructor
			if ( c.sortList.length > 0 ) {
				// save sortList before any sortAppend is added
				c.last.sortList = c.sortList;
				ts.sortOn( c, c.sortList, {}, !c.initWidgets );
			} else {
				ts.setHeadersCss( c );
				if ( c.initWidgets ) {
					// apply widget format
					ts.applyWidget( table, false );
				}
			}

			// show processesing icon
			if ( c.showProcessing ) {
				$table
				.unbind( 'sortBegin' + c.namespace + ' sortEnd' + c.namespace )
				.bind( 'sortBegin' + c.namespace + ' sortEnd' + c.namespace, function( e ) {
					clearTimeout( c.timerProcessing );
					ts.isProcessing( table );
					if ( e.type === 'sortBegin' ) {
						c.timerProcessing = setTimeout( function() {
							ts.isProcessing( table, true );
						}, 500 );
					}
				});
			}

			// initialized
			table.hasInitialized = true;
			table.isProcessing = false;
			if ( ts.debug(c, 'core') ) {
				console.log( 'Overall initialization time:' + ts.benchmark( $.data( table, 'startoveralltimer' ) ) );
				if ( ts.debug(c, 'core') && console.groupEnd ) { console.groupEnd(); }
			}
			$table.triggerHandler( 'tablesorter-initialized', table );
			if ( typeof c.initialized === 'function' ) {
				c.initialized( table );
			}
		},

		bindMethods : function( c ) {
			var $table = c.$table,
				namespace = c.namespace,
				events = ( 'sortReset update updateRows updateAll updateHeaders addRows updateCell updateComplete ' +
					'sorton appendCache updateCache applyWidgetId applyWidgets refreshWidgets destroy mouseup ' +
					'mouseleave ' ).split( ' ' )
					.join( namespace + ' ' );
			// apply easy methods that trigger bound events
			$table
			.unbind( events.replace( ts.regex.spaces, ' ' ) )
			.bind( 'sortReset' + namespace, function( e, callback ) {
				e.stopPropagation();
				// using this.config to ensure functions are getting a non-cached version of the config
				ts.sortReset( this.config, function( table ) {
					if (table.isApplyingWidgets) {
						// multiple triggers in a row... filterReset, then sortReset - see #1361
						// wait to update widgets
						setTimeout( function() {
							ts.applyWidget( table, '', callback );
						}, 100 );
					} else {
						ts.applyWidget( table, '', callback );
					}
				});
			})
			.bind( 'updateAll' + namespace, function( e, resort, callback ) {
				e.stopPropagation();
				ts.updateAll( this.config, resort, callback );
			})
			.bind( 'update' + namespace + ' updateRows' + namespace, function( e, resort, callback ) {
				e.stopPropagation();
				ts.update( this.config, resort, callback );
			})
			.bind( 'updateHeaders' + namespace, function( e, callback ) {
				e.stopPropagation();
				ts.updateHeaders( this.config, callback );
			})
			.bind( 'updateCell' + namespace, function( e, cell, resort, callback ) {
				e.stopPropagation();
				ts.updateCell( this.config, cell, resort, callback );
			})
			.bind( 'addRows' + namespace, function( e, $row, resort, callback ) {
				e.stopPropagation();
				ts.addRows( this.config, $row, resort, callback );
			})
			.bind( 'updateComplete' + namespace, function() {
				this.isUpdating = false;
			})
			.bind( 'sorton' + namespace, function( e, list, callback, init ) {
				e.stopPropagation();
				ts.sortOn( this.config, list, callback, init );
			})
			.bind( 'appendCache' + namespace, function( e, callback, init ) {
				e.stopPropagation();
				ts.appendCache( this.config, init );
				if ( $.isFunction( callback ) ) {
					callback( this );
				}
			})
			// $tbodies variable is used by the tbody sorting widget
			.bind( 'updateCache' + namespace, function( e, callback, $tbodies ) {
				e.stopPropagation();
				ts.updateCache( this.config, callback, $tbodies );
			})
			.bind( 'applyWidgetId' + namespace, function( e, id ) {
				e.stopPropagation();
				ts.applyWidgetId( this, id );
			})
			.bind( 'applyWidgets' + namespace, function( e, callback ) {
				e.stopPropagation();
				// apply widgets (false = not initializing)
				ts.applyWidget( this, false, callback );
			})
			.bind( 'refreshWidgets' + namespace, function( e, all, dontapply ) {
				e.stopPropagation();
				ts.refreshWidgets( this, all, dontapply );
			})
			.bind( 'removeWidget' + namespace, function( e, name, refreshing ) {
				e.stopPropagation();
				ts.removeWidget( this, name, refreshing );
			})
			.bind( 'destroy' + namespace, function( e, removeClasses, callback ) {
				e.stopPropagation();
				ts.destroy( this, removeClasses, callback );
			})
			.bind( 'resetToLoadState' + namespace, function( e ) {
				e.stopPropagation();
				// remove all widgets
				ts.removeWidget( this, true, false );
				var tmp = $.extend( true, {}, c.originalSettings );
				// restore original settings; this clears out current settings, but does not clear
				// values saved to storage.
				c = $.extend( true, {}, ts.defaults, tmp );
				c.originalSettings = tmp;
				this.hasInitialized = false;
				// setup the entire table again
				ts.setup( this, c );
			});
		},

		bindEvents : function( table, $headers, core ) {
			table = $( table )[ 0 ];
			var tmp,
				c = table.config,
				namespace = c.namespace,
				downTarget = null;
			if ( core !== true ) {
				$headers.addClass( namespace.slice( 1 ) + '_extra_headers' );
				tmp = ts.getClosest( $headers, 'table' );
				if ( tmp.length && tmp[ 0 ].nodeName === 'TABLE' && tmp[ 0 ] !== table ) {
					$( tmp[ 0 ] ).addClass( namespace.slice( 1 ) + '_extra_table' );
				}
			}
			tmp = ( c.pointerDown + ' ' + c.pointerUp + ' ' + c.pointerClick + ' sort keyup ' )
				.replace( ts.regex.spaces, ' ' )
				.split( ' ' )
				.join( namespace + ' ' );
			// apply event handling to headers and/or additional headers (stickyheaders, scroller, etc)
			$headers
			// http://stackoverflow.com/questions/5312849/jquery-find-self;
			.find( c.selectorSort )
			.add( $headers.filter( c.selectorSort ) )
			.unbind( tmp )
			.bind( tmp, function( e, external ) {
				var $cell, cell, temp,
					$target = $( e.target ),
					// wrap event type in spaces, so the match doesn't trigger on inner words
					type = ' ' + e.type + ' ';
				// only recognize left clicks
				if ( ( ( e.which || e.button ) !== 1 && !type.match( ' ' + c.pointerClick + ' | sort | keyup ' ) ) ||
					// allow pressing enter
					( type === ' keyup ' && e.which !== ts.keyCodes.enter ) ||
					// allow triggering a click event (e.which is undefined) & ignore physical clicks
					( type.match( ' ' + c.pointerClick + ' ' ) && typeof e.which !== 'undefined' ) ) {
					return;
				}
				// ignore mouseup if mousedown wasn't on the same target
				if ( type.match( ' ' + c.pointerUp + ' ' ) && downTarget !== e.target && external !== true ) {
					return;
				}
				// set target on mousedown
				if ( type.match( ' ' + c.pointerDown + ' ' ) ) {
					downTarget = e.target;
					// preventDefault needed or jQuery v1.3.2 and older throws an
					// "Uncaught TypeError: handler.apply is not a function" error
					temp = $target.jquery.split( '.' );
					if ( temp[ 0 ] === '1' && temp[ 1 ] < 4 ) { e.preventDefault(); }
					return;
				}
				downTarget = null;
				$cell = ts.getClosest( $( this ), '.' + ts.css.header );
				// prevent sort being triggered on form elements
				if ( ts.regex.formElements.test( e.target.nodeName ) ||
					// nosort class name, or elements within a nosort container
					$target.hasClass( c.cssNoSort ) || $target.parents( '.' + c.cssNoSort ).length > 0 ||
					// disabled cell directly clicked
					$cell.hasClass( 'sorter-false' ) ||
					// elements within a button
					$target.parents( 'button' ).length > 0 ) {
					return !c.cancelSelection;
				}
				if ( c.delayInit && ts.isEmptyObject( c.cache ) ) {
					ts.buildCache( c );
				}
				// use column index from data-attribute or index of current row; fixes #1116
				c.last.clickedIndex = $cell.attr( 'data-column' ) || $cell.index();
				cell = c.$headerIndexed[ c.last.clickedIndex ][0];
				if ( cell && !cell.sortDisabled ) {
					ts.initSort( c, cell, e );
				}
			});
			if ( c.cancelSelection ) {
				// cancel selection
				$headers
					.attr( 'unselectable', 'on' )
					.bind( 'selectstart', false )
					.css({
						'user-select' : 'none',
						'MozUserSelect' : 'none' // not needed for jQuery 1.8+
					});
			}
		},

		buildHeaders : function( c ) {
			var $temp, icon, timer, indx;
			c.headerList = [];
			c.headerContent = [];
			c.sortVars = [];
			if ( ts.debug(c, 'core') ) {
				timer = new Date();
			}
			// children tr in tfoot - see issue #196 & #547
			// don't pass table.config to computeColumnIndex here - widgets (math) pass it to "quickly" index tbody cells
			c.columns = ts.computeColumnIndex( c.$table.children( 'thead, tfoot' ).children( 'tr' ) );
			// add icon if cssIcon option exists
			icon = c.cssIcon ?
				'<i class="' + ( c.cssIcon === ts.css.icon ? ts.css.icon : c.cssIcon + ' ' + ts.css.icon ) + '"></i>' :
				'';
			// redefine c.$headers here in case of an updateAll that replaces or adds an entire header cell - see #683
			c.$headers = $( $.map( c.$table.find( c.selectorHeaders ), function( elem, index ) {
				var configHeaders, header, column, template, tmp,
					$elem = $( elem );
				// ignore cell (don't add it to c.$headers) if row has ignoreRow class
				if ( ts.getClosest( $elem, 'tr' ).hasClass( c.cssIgnoreRow ) ) { return; }
				// transfer data-column to element if not th/td - #1459
				if ( !/(th|td)/i.test( elem.nodeName ) ) {
					tmp = ts.getClosest( $elem, 'th, td' );
					$elem.attr( 'data-column', tmp.attr( 'data-column' ) );
				}
				// make sure to get header cell & not column indexed cell
				configHeaders = ts.getColumnData( c.table, c.headers, index, true );
				// save original header content
				c.headerContent[ index ] = $elem.html();
				// if headerTemplate is empty, don't reformat the header cell
				if ( c.headerTemplate !== '' && !$elem.find( '.' + ts.css.headerIn ).length ) {
					// set up header template
					template = c.headerTemplate
						.replace( ts.regex.templateContent, $elem.html() )
						.replace( ts.regex.templateIcon, $elem.find( '.' + ts.css.icon ).length ? '' : icon );
					if ( c.onRenderTemplate ) {
						header = c.onRenderTemplate.apply( $elem, [ index, template ] );
						// only change t if something is returned
						if ( header && typeof header === 'string' ) {
							template = header;
						}
					}
					$elem.html( '<div class="' + ts.css.headerIn + '">' + template + '</div>' ); // faster than wrapInner
				}
				if ( c.onRenderHeader ) {
					c.onRenderHeader.apply( $elem, [ index, c, c.$table ] );
				}
				column = parseInt( $elem.attr( 'data-column' ), 10 );
				elem.column = column;
				tmp = ts.getOrder( ts.getData( $elem, configHeaders, 'sortInitialOrder' ) || c.sortInitialOrder );
				// this may get updated numerous times if there are multiple rows
				c.sortVars[ column ] = {
					count : -1, // set to -1 because clicking on the header automatically adds one
					order : tmp ?
						( c.sortReset ? [ 1, 0, 2 ] : [ 1, 0 ] ) : // desc, asc, unsorted
						( c.sortReset ? [ 0, 1, 2 ] : [ 0, 1 ] ),  // asc, desc, unsorted
					lockedOrder : false,
					sortedBy : ''
				};
				tmp = ts.getData( $elem, configHeaders, 'lockedOrder' ) || false;
				if ( typeof tmp !== 'undefined' && tmp !== false ) {
					c.sortVars[ column ].lockedOrder = true;
					c.sortVars[ column ].order = ts.getOrder( tmp ) ? [ 1, 1 ] : [ 0, 0 ];
				}
				// add cell to headerList
				c.headerList[ index ] = elem;
				$elem.addClass( ts.css.header + ' ' + c.cssHeader );
				// add to parent in case there are multiple rows
				ts.getClosest( $elem, 'tr' )
					.addClass( ts.css.headerRow + ' ' + c.cssHeaderRow )
					.attr( 'role', 'row' );
				// allow keyboard cursor to focus on element
				if ( c.tabIndex ) {
					$elem.attr( 'tabindex', 0 );
				}
				return elem;
			}) );
			// cache headers per column
			c.$headerIndexed = [];
			for ( indx = 0; indx < c.columns; indx++ ) {
				// colspan in header making a column undefined
				if ( ts.isEmptyObject( c.sortVars[ indx ] ) ) {
					c.sortVars[ indx ] = {};
				}
				// Use c.$headers.parent() in case selectorHeaders doesn't point to the th/td
				$temp = c.$headers.filter( '[data-column="' + indx + '"]' );
				// target sortable column cells, unless there are none, then use non-sortable cells
				// .last() added in jQuery 1.4; use .filter(':last') to maintain compatibility with jQuery v1.2.6
				c.$headerIndexed[ indx ] = $temp.length ?
					$temp.not( '.sorter-false' ).length ?
						$temp.not( '.sorter-false' ).filter( ':last' ) :
						$temp.filter( ':last' ) :
					$();
			}
			c.$table.find( c.selectorHeaders ).attr({
				scope: 'col',
				role : 'columnheader'
			});
			// enable/disable sorting
			ts.updateHeader( c );
			if ( ts.debug(c, 'core') ) {
				console.log( 'Built headers:' + ts.benchmark( timer ) );
				console.log( c.$headers );
			}
		},

		// Use it to add a set of methods to table.config which will be available for all tables.
		// This should be done before table initialization
		addInstanceMethods : function( methods ) {
			$.extend( ts.instanceMethods, methods );
		},

		/*
		█████▄ ▄████▄ █████▄ ▄█████ ██████ █████▄ ▄█████
		██▄▄██ ██▄▄██ ██▄▄██ ▀█▄    ██▄▄   ██▄▄██ ▀█▄
		██▀▀▀  ██▀▀██ ██▀██     ▀█▄ ██▀▀   ██▀██     ▀█▄
		██     ██  ██ ██  ██ █████▀ ██████ ██  ██ █████▀
		*/
		setupParsers : function( c, $tbodies ) {
			var rows, list, span, max, colIndex, indx, header, configHeaders,
				noParser, parser, extractor, time, tbody, len,
				table = c.table,
				tbodyIndex = 0,
				debug = ts.debug(c, 'core'),
				debugOutput = {};
			// update table bodies in case we start with an empty table
			c.$tbodies = c.$table.children( 'tbody:not(.' + c.cssInfoBlock + ')' );
			tbody = typeof $tbodies === 'undefined' ? c.$tbodies : $tbodies;
			len = tbody.length;
			if ( len === 0 ) {
				return debug ? console.warn( 'Warning: *Empty table!* Not building a parser cache' ) : '';
			} else if ( debug ) {
				time = new Date();
				console[ console.group ? 'group' : 'log' ]( 'Detecting parsers for each column' );
			}
			list = {
				extractors: [],
				parsers: []
			};
			while ( tbodyIndex < len ) {
				rows = tbody[ tbodyIndex ].rows;
				if ( rows.length ) {
					colIndex = 0;
					max = c.columns;
					for ( indx = 0; indx < max; indx++ ) {
						header = c.$headerIndexed[ colIndex ];
						if ( header && header.length ) {
							// get column indexed table cell; adding true parameter fixes #1362 but
							// it would break backwards compatibility...
							configHeaders = ts.getColumnData( table, c.headers, colIndex ); // , true );
							// get column parser/extractor
							extractor = ts.getParserById( ts.getData( header, configHeaders, 'extractor' ) );
							parser = ts.getParserById( ts.getData( header, configHeaders, 'sorter' ) );
							noParser = ts.getData( header, configHeaders, 'parser' ) === 'false';
							// empty cells behaviour - keeping emptyToBottom for backwards compatibility
							c.empties[colIndex] = (
								ts.getData( header, configHeaders, 'empty' ) ||
								c.emptyTo || ( c.emptyToBottom ? 'bottom' : 'top' ) ).toLowerCase();
							// text strings behaviour in numerical sorts
							c.strings[colIndex] = (
								ts.getData( header, configHeaders, 'string' ) ||
								c.stringTo ||
								'max' ).toLowerCase();
							if ( noParser ) {
								parser = ts.getParserById( 'no-parser' );
							}
							if ( !extractor ) {
								// For now, maybe detect someday
								extractor = false;
							}
							if ( !parser ) {
								parser = ts.detectParserForColumn( c, rows, -1, colIndex );
							}
							if ( debug ) {
								debugOutput[ '(' + colIndex + ') ' + header.text() ] = {
									parser : parser.id,
									extractor : extractor ? extractor.id : 'none',
									string : c.strings[ colIndex ],
									empty  : c.empties[ colIndex ]
								};
							}
							list.parsers[ colIndex ] = parser;
							list.extractors[ colIndex ] = extractor;
							span = header[ 0 ].colSpan - 1;
							if ( span > 0 ) {
								colIndex += span;
								max += span;
								while ( span + 1 > 0 ) {
									// set colspan columns to use the same parsers & extractors
									list.parsers[ colIndex - span ] = parser;
									list.extractors[ colIndex - span ] = extractor;
									span--;
								}
							}
						}
						colIndex++;
					}
				}
				tbodyIndex += ( list.parsers.length ) ? len : 1;
			}
			if ( debug ) {
				if ( !ts.isEmptyObject( debugOutput ) ) {
					console[ console.table ? 'table' : 'log' ]( debugOutput );
				} else {
					console.warn( '  No parsers detected!' );
				}
				console.log( 'Completed detecting parsers' + ts.benchmark( time ) );
				if ( console.groupEnd ) { console.groupEnd(); }
			}
			c.parsers = list.parsers;
			c.extractors = list.extractors;
		},

		addParser : function( parser ) {
			var indx,
				len = ts.parsers.length,
				add = true;
			for ( indx = 0; indx < len; indx++ ) {
				if ( ts.parsers[ indx ].id.toLowerCase() === parser.id.toLowerCase() ) {
					add = false;
				}
			}
			if ( add ) {
				ts.parsers[ ts.parsers.length ] = parser;
			}
		},

		getParserById : function( name ) {
			/*jshint eqeqeq:false */ // eslint-disable-next-line eqeqeq
			if ( name == 'false' ) { return false; }
			var indx,
				len = ts.parsers.length;
			for ( indx = 0; indx < len; indx++ ) {
				if ( ts.parsers[ indx ].id.toLowerCase() === ( name.toString() ).toLowerCase() ) {
					return ts.parsers[ indx ];
				}
			}
			return false;
		},

		detectParserForColumn : function( c, rows, rowIndex, cellIndex ) {
			var cur, $node, row,
				indx = ts.parsers.length,
				node = false,
				nodeValue = '',
				debug = ts.debug(c, 'core'),
				keepLooking = true;
			while ( nodeValue === '' && keepLooking ) {
				rowIndex++;
				row = rows[ rowIndex ];
				// stop looking after 50 empty rows
				if ( row && rowIndex < 50 ) {
					if ( row.className.indexOf( ts.cssIgnoreRow ) < 0 ) {
						node = rows[ rowIndex ].cells[ cellIndex ];
						nodeValue = ts.getElementText( c, node, cellIndex );
						$node = $( node );
						if ( debug ) {
							console.log( 'Checking if value was empty on row ' + rowIndex + ', column: ' +
								cellIndex + ': "' + nodeValue + '"' );
						}
					}
				} else {
					keepLooking = false;
				}
			}
			while ( --indx >= 0 ) {
				cur = ts.parsers[ indx ];
				// ignore the default text parser because it will always be true
				if ( cur && cur.id !== 'text' && cur.is && cur.is( nodeValue, c.table, node, $node ) ) {
					return cur;
				}
			}
			// nothing found, return the generic parser (text)
			return ts.getParserById( 'text' );
		},

		getElementText : function( c, node, cellIndex ) {
			if ( !node ) { return ''; }
			var tmp,
				extract = c.textExtraction || '',
				// node could be a jquery object
				// http://jsperf.com/jquery-vs-instanceof-jquery/2
				$node = node.jquery ? node : $( node );
			if ( typeof extract === 'string' ) {
				// check data-attribute first when set to 'basic'; don't use node.innerText - it's really slow!
				// http://www.kellegous.com/j/2013/02/27/innertext-vs-textcontent/
				if ( extract === 'basic' && typeof ( tmp = $node.attr( c.textAttribute ) ) !== 'undefined' ) {
					return $.trim( tmp );
				}
				return $.trim( node.textContent || $node.text() );
			} else {
				if ( typeof extract === 'function' ) {
					return $.trim( extract( $node[ 0 ], c.table, cellIndex ) );
				} else if ( typeof ( tmp = ts.getColumnData( c.table, extract, cellIndex ) ) === 'function' ) {
					return $.trim( tmp( $node[ 0 ], c.table, cellIndex ) );
				}
			}
			// fallback
			return $.trim( $node[ 0 ].textContent || $node.text() );
		},

		// centralized function to extract/parse cell contents
		getParsedText : function( c, cell, colIndex, txt ) {
			if ( typeof txt === 'undefined' ) {
				txt = ts.getElementText( c, cell, colIndex );
			}
			// if no parser, make sure to return the txt
			var val = '' + txt,
				parser = c.parsers[ colIndex ],
				extractor = c.extractors[ colIndex ];
			if ( parser ) {
				// do extract before parsing, if there is one
				if ( extractor && typeof extractor.format === 'function' ) {
					txt = extractor.format( txt, c.table, cell, colIndex );
				}
				// allow parsing if the string is empty, previously parsing would change it to zero,
				// in case the parser needs to extract data from the table cell attributes
				val = parser.id === 'no-parser' ? '' :
					// make sure txt is a string (extractor may have converted it)
					parser.format( '' + txt, c.table, cell, colIndex );
				if ( c.ignoreCase && typeof val === 'string' ) {
					val = val.toLowerCase();
				}
			}
			return val;
		},

		/*
		▄████▄ ▄████▄ ▄████▄ ██  ██ ██████
		██  ▀▀ ██▄▄██ ██  ▀▀ ██▄▄██ ██▄▄
		██  ▄▄ ██▀▀██ ██  ▄▄ ██▀▀██ ██▀▀
		▀████▀ ██  ██ ▀████▀ ██  ██ ██████
		*/
		buildCache : function( c, callback, $tbodies ) {
			var cache, val, txt, rowIndex, colIndex, tbodyIndex, $tbody, $row,
				cols, $cells, cell, cacheTime, totalRows, rowData, prevRowData,
				colMax, span, cacheIndex, hasParser, max, len, index,
				table = c.table,
				parsers = c.parsers,
				debug = ts.debug(c, 'core');
			// update tbody variable
			c.$tbodies = c.$table.children( 'tbody:not(.' + c.cssInfoBlock + ')' );
			$tbody = typeof $tbodies === 'undefined' ? c.$tbodies : $tbodies,
			c.cache = {};
			c.totalRows = 0;
			// if no parsers found, return - it's an empty table.
			if ( !parsers ) {
				return debug ? console.warn( 'Warning: *Empty table!* Not building a cache' ) : '';
			}
			if ( debug ) {
				cacheTime = new Date();
			}
			// processing icon
			if ( c.showProcessing ) {
				ts.isProcessing( table, true );
			}
			for ( tbodyIndex = 0; tbodyIndex < $tbody.length; tbodyIndex++ ) {
				colMax = []; // column max value per tbody
				cache = c.cache[ tbodyIndex ] = {
					normalized: [] // array of normalized row data; last entry contains 'rowData' above
					// colMax: #   // added at the end
				};

				totalRows = ( $tbody[ tbodyIndex ] && $tbody[ tbodyIndex ].rows.length ) || 0;
				for ( rowIndex = 0; rowIndex < totalRows; ++rowIndex ) {
					rowData = {
						// order: original row order #
						// $row : jQuery Object[]
						child: [], // child row text (filter widget)
						raw: []    // original row text
					};
					/** Add the table data to main data array */
					$row = $( $tbody[ tbodyIndex ].rows[ rowIndex ] );
					cols = [];
					// ignore "remove-me" rows
					if ( $row.hasClass( c.selectorRemove.slice(1) ) ) {
						continue;
					}
					// if this is a child row, add it to the last row's children and continue to the next row
					// ignore child row class, if it is the first row
					if ( $row.hasClass( c.cssChildRow ) && rowIndex !== 0 ) {
						len = cache.normalized.length - 1;
						prevRowData = cache.normalized[ len ][ c.columns ];
						prevRowData.$row = prevRowData.$row.add( $row );
						// add 'hasChild' class name to parent row
						if ( !$row.prev().hasClass( c.cssChildRow ) ) {
							$row.prev().addClass( ts.css.cssHasChild );
						}
						// save child row content (un-parsed!)
						$cells = $row.children( 'th, td' );
						len = prevRowData.child.length;
						prevRowData.child[ len ] = [];
						// child row content does not account for colspans/rowspans; so indexing may be off
						cacheIndex = 0;
						max = c.columns;
						for ( colIndex = 0; colIndex < max; colIndex++ ) {
							cell = $cells[ colIndex ];
							if ( cell ) {
								prevRowData.child[ len ][ colIndex ] = ts.getParsedText( c, cell, colIndex );
								span = $cells[ colIndex ].colSpan - 1;
								if ( span > 0 ) {
									cacheIndex += span;
									max += span;
								}
							}
							cacheIndex++;
						}
						// go to the next for loop
						continue;
					}
					rowData.$row = $row;
					rowData.order = rowIndex; // add original row position to rowCache
					cacheIndex = 0;
					max = c.columns;
					for ( colIndex = 0; colIndex < max; ++colIndex ) {
						cell = $row[ 0 ].cells[ colIndex ];
						if ( cell && cacheIndex < c.columns ) {
							hasParser = typeof parsers[ cacheIndex ] !== 'undefined';
							if ( !hasParser && debug ) {
								console.warn( 'No parser found for row: ' + rowIndex + ', column: ' + colIndex +
									'; cell containing: "' + $(cell).text() + '"; does it have a header?' );
							}
							val = ts.getElementText( c, cell, cacheIndex );
							rowData.raw[ cacheIndex ] = val; // save original row text
							// save raw column text even if there is no parser set
							txt = ts.getParsedText( c, cell, cacheIndex, val );
							cols[ cacheIndex ] = txt;
							if ( hasParser && ( parsers[ cacheIndex ].type || '' ).toLowerCase() === 'numeric' ) {
								// determine column max value (ignore sign)
								colMax[ cacheIndex ] = Math.max( Math.abs( txt ) || 0, colMax[ cacheIndex ] || 0 );
							}
							// allow colSpan in tbody
							span = cell.colSpan - 1;
							if ( span > 0 ) {
								index = 0;
								while ( index <= span ) {
									// duplicate text (or not) to spanned columns
									// instead of setting duplicate span to empty string, use textExtraction to try to get a value
									// see http://stackoverflow.com/q/36449711/145346
									txt = c.duplicateSpan || index === 0 ?
										txt :
										typeof c.textExtraction !== 'string' ?
											ts.getElementText( c, cell, cacheIndex + index ) || '' :
											'';
									rowData.raw[ cacheIndex + index ] = txt;
									cols[ cacheIndex + index ] = txt;
									index++;
								}
								cacheIndex += span;
								max += span;
							}
						}
						cacheIndex++;
					}
					// ensure rowData is always in the same location (after the last column)
					cols[ c.columns ] = rowData;
					cache.normalized[ cache.normalized.length ] = cols;
				}
				cache.colMax = colMax;
				// total up rows, not including child rows
				c.totalRows += cache.normalized.length;

			}
			if ( c.showProcessing ) {
				ts.isProcessing( table ); // remove processing icon
			}
			if ( debug ) {
				len = Math.min( 5, c.cache[ 0 ].normalized.length );
				console[ console.group ? 'group' : 'log' ]( 'Building cache for ' + c.totalRows +
					' rows (showing ' + len + ' rows in log) and ' + c.columns + ' columns' +
					ts.benchmark( cacheTime ) );
				val = {};
				for ( colIndex = 0; colIndex < c.columns; colIndex++ ) {
					for ( cacheIndex = 0; cacheIndex < len; cacheIndex++ ) {
						if ( !val[ 'row: ' + cacheIndex ] ) {
							val[ 'row: ' + cacheIndex ] = {};
						}
						val[ 'row: ' + cacheIndex ][ c.$headerIndexed[ colIndex ].text() ] =
							c.cache[ 0 ].normalized[ cacheIndex ][ colIndex ];
					}
				}
				console[ console.table ? 'table' : 'log' ]( val );
				if ( console.groupEnd ) { console.groupEnd(); }
			}
			if ( $.isFunction( callback ) ) {
				callback( table );
			}
		},

		getColumnText : function( table, column, callback, rowFilter ) {
			table = $( table )[0];
			var tbodyIndex, rowIndex, cache, row, tbodyLen, rowLen, raw, parsed, $cell, result,
				hasCallback = typeof callback === 'function',
				allColumns = column === 'all',
				data = { raw : [], parsed: [], $cell: [] },
				c = table.config;
			if ( ts.isEmptyObject( c ) ) {
				if ( ts.debug(c, 'core') ) {
					console.warn( 'No cache found - aborting getColumnText function!' );
				}
			} else {
				tbodyLen = c.$tbodies.length;
				for ( tbodyIndex = 0; tbodyIndex < tbodyLen; tbodyIndex++ ) {
					cache = c.cache[ tbodyIndex ].normalized;
					rowLen = cache.length;
					for ( rowIndex = 0; rowIndex < rowLen; rowIndex++ ) {
						row = cache[ rowIndex ];
						if ( rowFilter && !row[ c.columns ].$row.is( rowFilter ) ) {
							continue;
						}
						result = true;
						parsed = ( allColumns ) ? row.slice( 0, c.columns ) : row[ column ];
						row = row[ c.columns ];
						raw = ( allColumns ) ? row.raw : row.raw[ column ];
						$cell = ( allColumns ) ? row.$row.children() : row.$row.children().eq( column );
						if ( hasCallback ) {
							result = callback({
								tbodyIndex : tbodyIndex,
								rowIndex : rowIndex,
								parsed : parsed,
								raw : raw,
								$row : row.$row,
								$cell : $cell
							});
						}
						if ( result !== false ) {
							data.parsed[ data.parsed.length ] = parsed;
							data.raw[ data.raw.length ] = raw;
							data.$cell[ data.$cell.length ] = $cell;
						}
					}
				}
				// return everything
				return data;
			}
		},

		/*
		██  ██ █████▄ █████▄ ▄████▄ ██████ ██████
		██  ██ ██▄▄██ ██  ██ ██▄▄██   ██   ██▄▄
		██  ██ ██▀▀▀  ██  ██ ██▀▀██   ██   ██▀▀
		▀████▀ ██     █████▀ ██  ██   ██   ██████
		*/
		setHeadersCss : function( c ) {
			var indx, column,
				list = c.sortList,
				len = list.length,
				none = ts.css.sortNone + ' ' + c.cssNone,
				css = [ ts.css.sortAsc + ' ' + c.cssAsc, ts.css.sortDesc + ' ' + c.cssDesc ],
				cssIcon = [ c.cssIconAsc, c.cssIconDesc, c.cssIconNone ],
				aria = [ 'ascending', 'descending' ],
				updateColumnSort = function($el, index) {
					$el
						.removeClass( none )
						.addClass( css[ index ] )
						.attr( 'aria-sort', aria[ index ] )
						.find( '.' + ts.css.icon )
						.removeClass( cssIcon[ 2 ] )
						.addClass( cssIcon[ index ] );
				},
				// find the footer
				$extras = c.$table
					.find( 'tfoot tr' )
					.children( 'td, th' )
					.add( $( c.namespace + '_extra_headers' ) )
					.removeClass( css.join( ' ' ) ),
				// remove all header information
				$sorted = c.$headers
					.add( $( 'thead ' + c.namespace + '_extra_headers' ) )
					.removeClass( css.join( ' ' ) )
					.addClass( none )
					.attr( 'aria-sort', 'none' )
					.find( '.' + ts.css.icon )
					.removeClass( cssIcon.join( ' ' ) )
					.end();
			// add css none to all sortable headers
			$sorted
				.not( '.sorter-false' )
				.find( '.' + ts.css.icon )
				.addClass( cssIcon[ 2 ] );
			// add disabled css icon class
			if ( c.cssIconDisabled ) {
				$sorted
					.filter( '.sorter-false' )
					.find( '.' + ts.css.icon )
					.addClass( c.cssIconDisabled );
			}
			for ( indx = 0; indx < len; indx++ ) {
				// direction = 2 means reset!
				if ( list[ indx ][ 1 ] !== 2 ) {
					// multicolumn sorting updating - see #1005
					// .not(function() {}) needs jQuery 1.4
					// filter(function(i, el) {}) <- el is undefined in jQuery v1.2.6
					$sorted = c.$headers.filter( function( i ) {
						// only include headers that are in the sortList (this includes colspans)
						var include = true,
							$el = c.$headers.eq( i ),
							col = parseInt( $el.attr( 'data-column' ), 10 ),
							end = col + ts.getClosest( $el, 'th, td' )[0].colSpan;
						for ( ; col < end; col++ ) {
							include = include ? include || ts.isValueInArray( col, c.sortList ) > -1 : false;
						}
						return include;
					});

					// choose the :last in case there are nested columns
					$sorted = $sorted
						.not( '.sorter-false' )
						.filter( '[data-column="' + list[ indx ][ 0 ] + '"]' + ( len === 1 ? ':last' : '' ) );
					if ( $sorted.length ) {
						for ( column = 0; column < $sorted.length; column++ ) {
							if ( !$sorted[ column ].sortDisabled ) {
								updateColumnSort( $sorted.eq( column ), list[ indx ][ 1 ] );
							}
						}
					}
					// add sorted class to footer & extra headers, if they exist
					if ( $extras.length ) {
						updateColumnSort( $extras.filter( '[data-column="' + list[ indx ][ 0 ] + '"]' ), list[ indx ][ 1 ] );
					}
				}
			}
			// add verbose aria labels
			len = c.$headers.length;
			for ( indx = 0; indx < len; indx++ ) {
				ts.setColumnAriaLabel( c, c.$headers.eq( indx ) );
			}
		},

		getClosest : function( $el, selector ) {
			// jQuery v1.2.6 doesn't have closest()
			if ( $.fn.closest ) {
				return $el.closest( selector );
			}
			return $el.is( selector ) ?
				$el :
				$el.parents( selector ).filter( ':first' );
		},

		// nextSort (optional), lets you disable next sort text
		setColumnAriaLabel : function( c, $header, nextSort ) {
			if ( $header.length ) {
				var column = parseInt( $header.attr( 'data-column' ), 10 ),
					vars = c.sortVars[ column ],
					tmp = $header.hasClass( ts.css.sortAsc ) ?
						'sortAsc' :
						$header.hasClass( ts.css.sortDesc ) ? 'sortDesc' : 'sortNone',
					txt = $.trim( $header.text() ) + ': ' + ts.language[ tmp ];
				if ( $header.hasClass( 'sorter-false' ) || nextSort === false ) {
					txt += ts.language.sortDisabled;
				} else {
					tmp = ( vars.count + 1 ) % vars.order.length;
					nextSort = vars.order[ tmp ];
					// if nextSort
					txt += ts.language[ nextSort === 0 ? 'nextAsc' : nextSort === 1 ? 'nextDesc' : 'nextNone' ];
				}
				$header.attr( 'aria-label', txt );
				if (vars.sortedBy) {
					$header.attr( 'data-sortedBy', vars.sortedBy );
				} else {
					$header.removeAttr('data-sortedBy');
				}
			}
		},

		updateHeader : function( c ) {
			var index, isDisabled, $header, col,
				table = c.table,
				len = c.$headers.length;
			for ( index = 0; index < len; index++ ) {
				$header = c.$headers.eq( index );
				col = ts.getColumnData( table, c.headers, index, true );
				// add 'sorter-false' class if 'parser-false' is set
				isDisabled = ts.getData( $header, col, 'sorter' ) === 'false' || ts.getData( $header, col, 'parser' ) === 'false';
				ts.setColumnSort( c, $header, isDisabled );
			}
		},

		setColumnSort : function( c, $header, isDisabled ) {
			var id = c.table.id;
			$header[ 0 ].sortDisabled = isDisabled;
			$header[ isDisabled ? 'addClass' : 'removeClass' ]( 'sorter-false' )
				.attr( 'aria-disabled', '' + isDisabled );
			// disable tab index on disabled cells
			if ( c.tabIndex ) {
				if ( isDisabled ) {
					$header.removeAttr( 'tabindex' );
				} else {
					$header.attr( 'tabindex', '0' );
				}
			}
			// aria-controls - requires table ID
			if ( id ) {
				if ( isDisabled ) {
					$header.removeAttr( 'aria-controls' );
				} else {
					$header.attr( 'aria-controls', id );
				}
			}
		},

		updateHeaderSortCount : function( c, list ) {
			var col, dir, group, indx, primary, temp, val, order,
				sortList = list || c.sortList,
				len = sortList.length;
			c.sortList = [];
			for ( indx = 0; indx < len; indx++ ) {
				val = sortList[ indx ];
				// ensure all sortList values are numeric - fixes #127
				col = parseInt( val[ 0 ], 10 );
				// prevents error if sorton array is wrong
				if ( col < c.columns ) {

					// set order if not already defined - due to colspan header without associated header cell
					// adding this check prevents a javascript error
					if ( !c.sortVars[ col ].order ) {
						if ( ts.getOrder( c.sortInitialOrder ) ) {
							order = c.sortReset ? [ 1, 0, 2 ] : [ 1, 0 ];
						} else {
							order = c.sortReset ? [ 0, 1, 2 ] : [ 0, 1 ];
						}
						c.sortVars[ col ].order = order;
						c.sortVars[ col ].count = 0;
					}

					order = c.sortVars[ col ].order;
					dir = ( '' + val[ 1 ] ).match( /^(1|d|s|o|n)/ );
					dir = dir ? dir[ 0 ] : '';
					// 0/(a)sc (default), 1/(d)esc, (s)ame, (o)pposite, (n)ext
					switch ( dir ) {
						case '1' : case 'd' : // descending
							dir = 1;
							break;
						case 's' : // same direction (as primary column)
							// if primary sort is set to 's', make it ascending
							dir = primary || 0;
							break;
						case 'o' :
							temp = order[ ( primary || 0 ) % order.length ];
							// opposite of primary column; but resets if primary resets
							dir = temp === 0 ? 1 : temp === 1 ? 0 : 2;
							break;
						case 'n' :
							dir = order[ ( ++c.sortVars[ col ].count ) % order.length ];
							break;
						default : // ascending
							dir = 0;
							break;
					}
					primary = indx === 0 ? dir : primary;
					group = [ col, parseInt( dir, 10 ) || 0 ];
					c.sortList[ c.sortList.length ] = group;
					dir = $.inArray( group[ 1 ], order ); // fixes issue #167
					c.sortVars[ col ].count = dir >= 0 ? dir : group[ 1 ] % order.length;
				}
			}
		},

		updateAll : function( c, resort, callback ) {
			var table = c.table;
			table.isUpdating = true;
			ts.refreshWidgets( table, true, true );
			ts.buildHeaders( c );
			ts.bindEvents( table, c.$headers, true );
			ts.bindMethods( c );
			ts.commonUpdate( c, resort, callback );
		},

		update : function( c, resort, callback ) {
			var table = c.table;
			table.isUpdating = true;
			// update sorting (if enabled/disabled)
			ts.updateHeader( c );
			ts.commonUpdate( c, resort, callback );
		},

		// simple header update - see #989
		updateHeaders : function( c, callback ) {
			c.table.isUpdating = true;
			ts.buildHeaders( c );
			ts.bindEvents( c.table, c.$headers, true );
			ts.resortComplete( c, callback );
		},

		updateCell : function( c, cell, resort, callback ) {
			// updateCell for child rows is a mess - we'll ignore them for now
			// eventually I'll break out the "update" row cache code to make everything consistent
			if ( $( cell ).closest( 'tr' ).hasClass( c.cssChildRow ) ) {
				console.warn('Tablesorter Warning! "updateCell" for child row content has been disabled, use "update" instead');
				return;
			}
			if ( ts.isEmptyObject( c.cache ) ) {
				// empty table, do an update instead - fixes #1099
				ts.updateHeader( c );
				ts.commonUpdate( c, resort, callback );
				return;
			}
			c.table.isUpdating = true;
			c.$table.find( c.selectorRemove ).remove();
			// get position from the dom
			var tmp, indx, row, icell, cache, len,
				$tbodies = c.$tbodies,
				$cell = $( cell ),
				// update cache - format: function( s, table, cell, cellIndex )
				// no closest in jQuery v1.2.6
				tbodyIndex = $tbodies.index( ts.getClosest( $cell, 'tbody' ) ),
				tbcache = c.cache[ tbodyIndex ],
				$row = ts.getClosest( $cell, 'tr' );
			cell = $cell[ 0 ]; // in case cell is a jQuery object
			// tbody may not exist if update is initialized while tbody is removed for processing
			if ( $tbodies.length && tbodyIndex >= 0 ) {
				row = $tbodies.eq( tbodyIndex ).find( 'tr' ).not( '.' + c.cssChildRow ).index( $row );
				cache = tbcache.normalized[ row ];
				len = $row[ 0 ].cells.length;
				if ( len !== c.columns ) {
					// colspan in here somewhere!
					icell = 0;
					tmp = false;
					for ( indx = 0; indx < len; indx++ ) {
						if ( !tmp && $row[ 0 ].cells[ indx ] !== cell ) {
							icell += $row[ 0 ].cells[ indx ].colSpan;
						} else {
							tmp = true;
						}
					}
				} else {
					icell = $cell.index();
				}
				tmp = ts.getElementText( c, cell, icell ); // raw
				cache[ c.columns ].raw[ icell ] = tmp;
				tmp = ts.getParsedText( c, cell, icell, tmp );
				cache[ icell ] = tmp; // parsed
				if ( ( c.parsers[ icell ].type || '' ).toLowerCase() === 'numeric' ) {
					// update column max value (ignore sign)
					tbcache.colMax[ icell ] = Math.max( Math.abs( tmp ) || 0, tbcache.colMax[ icell ] || 0 );
				}
				tmp = resort !== 'undefined' ? resort : c.resort;
				if ( tmp !== false ) {
					// widgets will be reapplied
					ts.checkResort( c, tmp, callback );
				} else {
					// don't reapply widgets is resort is false, just in case it causes
					// problems with element focus
					ts.resortComplete( c, callback );
				}
			} else {
				if ( ts.debug(c, 'core') ) {
					console.error( 'updateCell aborted, tbody missing or not within the indicated table' );
				}
				c.table.isUpdating = false;
			}
		},

		addRows : function( c, $row, resort, callback ) {
			var txt, val, tbodyIndex, rowIndex, rows, cellIndex, len, order,
				cacheIndex, rowData, cells, cell, span,
				// allow passing a row string if only one non-info tbody exists in the table
				valid = typeof $row === 'string' && c.$tbodies.length === 1 && /<tr/.test( $row || '' ),
				table = c.table;
			if ( valid ) {
				$row = $( $row );
				c.$tbodies.append( $row );
			} else if (
				!$row ||
				// row is a jQuery object?
				!( $row instanceof $ ) ||
				// row contained in the table?
				( ts.getClosest( $row, 'table' )[ 0 ] !== c.table )
			) {
				if ( ts.debug(c, 'core') ) {
					console.error( 'addRows method requires (1) a jQuery selector reference to rows that have already ' +
						'been added to the table, or (2) row HTML string to be added to a table with only one tbody' );
				}
				return false;
			}
			table.isUpdating = true;
			if ( ts.isEmptyObject( c.cache ) ) {
				// empty table, do an update instead - fixes #450
				ts.updateHeader( c );
				ts.commonUpdate( c, resort, callback );
			} else {
				rows = $row.filter( 'tr' ).attr( 'role', 'row' ).length;
				tbodyIndex = c.$tbodies.index( $row.parents( 'tbody' ).filter( ':first' ) );
				// fixes adding rows to an empty table - see issue #179
				if ( !( c.parsers && c.parsers.length ) ) {
					ts.setupParsers( c );
				}
				// add each row
				for ( rowIndex = 0; rowIndex < rows; rowIndex++ ) {
					cacheIndex = 0;
					len = $row[ rowIndex ].cells.length;
					order = c.cache[ tbodyIndex ].normalized.length;
					cells = [];
					rowData = {
						child : [],
						raw : [],
						$row : $row.eq( rowIndex ),
						order : order
					};
					// add each cell
					for ( cellIndex = 0; cellIndex < len; cellIndex++ ) {
						cell = $row[ rowIndex ].cells[ cellIndex ];
						txt = ts.getElementText( c, cell, cacheIndex );
						rowData.raw[ cacheIndex ] = txt;
						val = ts.getParsedText( c, cell, cacheIndex, txt );
						cells[ cacheIndex ] = val;
						if ( ( c.parsers[ cacheIndex ].type || '' ).toLowerCase() === 'numeric' ) {
							// update column max value (ignore sign)
							c.cache[ tbodyIndex ].colMax[ cacheIndex ] =
								Math.max( Math.abs( val ) || 0, c.cache[ tbodyIndex ].colMax[ cacheIndex ] || 0 );
						}
						span = cell.colSpan - 1;
						if ( span > 0 ) {
							cacheIndex += span;
						}
						cacheIndex++;
					}
					// add the row data to the end
					cells[ c.columns ] = rowData;
					// update cache
					c.cache[ tbodyIndex ].normalized[ order ] = cells;
				}
				// resort using current settings
				ts.checkResort( c, resort, callback );
			}
		},

		updateCache : function( c, callback, $tbodies ) {
			// rebuild parsers
			if ( !( c.parsers && c.parsers.length ) ) {
				ts.setupParsers( c, $tbodies );
			}
			// rebuild the cache map
			ts.buildCache( c, callback, $tbodies );
		},

		// init flag (true) used by pager plugin to prevent widget application
		// renamed from appendToTable
		appendCache : function( c, init ) {
			var parsed, totalRows, $tbody, $curTbody, rowIndex, tbodyIndex, appendTime,
				table = c.table,
				$tbodies = c.$tbodies,
				rows = [],
				cache = c.cache;
			// empty table - fixes #206/#346
			if ( ts.isEmptyObject( cache ) ) {
				// run pager appender in case the table was just emptied
				return c.appender ? c.appender( table, rows ) :
					table.isUpdating ? c.$table.triggerHandler( 'updateComplete', table ) : ''; // Fixes #532
			}
			if ( ts.debug(c, 'core') ) {
				appendTime = new Date();
			}
			for ( tbodyIndex = 0; tbodyIndex < $tbodies.length; tbodyIndex++ ) {
				$tbody = $tbodies.eq( tbodyIndex );
				if ( $tbody.length ) {
					// detach tbody for manipulation
					$curTbody = ts.processTbody( table, $tbody, true );
					parsed = cache[ tbodyIndex ].normalized;
					totalRows = parsed.length;
					for ( rowIndex = 0; rowIndex < totalRows; rowIndex++ ) {
						rows[rows.length] = parsed[ rowIndex ][ c.columns ].$row;
						// removeRows used by the pager plugin; don't render if using ajax - fixes #411
						if ( !c.appender || ( c.pager && !c.pager.removeRows && !c.pager.ajax ) ) {
							$curTbody.append( parsed[ rowIndex ][ c.columns ].$row );
						}
					}
					// restore tbody
					ts.processTbody( table, $curTbody, false );
				}
			}
			if ( c.appender ) {
				c.appender( table, rows );
			}
			if ( ts.debug(c, 'core') ) {
				console.log( 'Rebuilt table' + ts.benchmark( appendTime ) );
			}
			// apply table widgets; but not before ajax completes
			if ( !init && !c.appender ) {
				ts.applyWidget( table );
			}
			if ( table.isUpdating ) {
				c.$table.triggerHandler( 'updateComplete', table );
			}
		},

		commonUpdate : function( c, resort, callback ) {
			// remove rows/elements before update
			c.$table.find( c.selectorRemove ).remove();
			// rebuild parsers
			ts.setupParsers( c );
			// rebuild the cache map
			ts.buildCache( c );
			ts.checkResort( c, resort, callback );
		},

		/*
		▄█████ ▄████▄ █████▄ ██████ ██ █████▄ ▄████▄
		▀█▄    ██  ██ ██▄▄██   ██   ██ ██  ██ ██ ▄▄▄
		   ▀█▄ ██  ██ ██▀██    ██   ██ ██  ██ ██ ▀██
		█████▀ ▀████▀ ██  ██   ██   ██ ██  ██ ▀████▀
		*/
		initSort : function( c, cell, event ) {
			if ( c.table.isUpdating ) {
				// let any updates complete before initializing a sort
				return setTimeout( function() {
					ts.initSort( c, cell, event );
				}, 50 );
			}

			var arry, indx, headerIndx, dir, temp, tmp, $header,
				notMultiSort = !event[ c.sortMultiSortKey ],
				table = c.table,
				len = c.$headers.length,
				th = ts.getClosest( $( cell ), 'th, td' ),
				col = parseInt( th.attr( 'data-column' ), 10 ),
				sortedBy = event.type === 'mouseup' ? 'user' : event.type,
				order = c.sortVars[ col ].order;
			th = th[0];
			// Only call sortStart if sorting is enabled
			c.$table.triggerHandler( 'sortStart', table );
			// get current column sort order
			tmp = ( c.sortVars[ col ].count + 1 ) % order.length;
			c.sortVars[ col ].count = event[ c.sortResetKey ] ? 2 : tmp;
			// reset all sorts on non-current column - issue #30
			if ( c.sortRestart ) {
				for ( headerIndx = 0; headerIndx < len; headerIndx++ ) {
					$header = c.$headers.eq( headerIndx );
					tmp = parseInt( $header.attr( 'data-column' ), 10 );
					// only reset counts on columns that weren't just clicked on and if not included in a multisort
					if ( col !== tmp && ( notMultiSort || $header.hasClass( ts.css.sortNone ) ) ) {
						c.sortVars[ tmp ].count = -1;
					}
				}
			}
			// user only wants to sort on one column
			if ( notMultiSort ) {
				$.each( c.sortVars, function( i ) {
					c.sortVars[ i ].sortedBy = '';
				});
				// flush the sort list
				c.sortList = [];
				c.last.sortList = [];
				if ( c.sortForce !== null ) {
					arry = c.sortForce;
					for ( indx = 0; indx < arry.length; indx++ ) {
						if ( arry[ indx ][ 0 ] !== col ) {
							c.sortList[ c.sortList.length ] = arry[ indx ];
							c.sortVars[ arry[ indx ][ 0 ] ].sortedBy = 'sortForce';
						}
					}
				}
				// add column to sort list
				dir = order[ c.sortVars[ col ].count ];
				if ( dir < 2 ) {
					c.sortList[ c.sortList.length ] = [ col, dir ];
					c.sortVars[ col ].sortedBy = sortedBy;
					// add other columns if header spans across multiple
					if ( th.colSpan > 1 ) {
						for ( indx = 1; indx < th.colSpan; indx++ ) {
							c.sortList[ c.sortList.length ] = [ col + indx, dir ];
							// update count on columns in colSpan
							c.sortVars[ col + indx ].count = $.inArray( dir, order );
							c.sortVars[ col + indx ].sortedBy = sortedBy;
						}
					}
				}
				// multi column sorting
			} else {
				// get rid of the sortAppend before adding more - fixes issue #115 & #523
				c.sortList = $.extend( [], c.last.sortList );

				// the user has clicked on an already sorted column
				if ( ts.isValueInArray( col, c.sortList ) >= 0 ) {
					// reverse the sorting direction
					c.sortVars[ col ].sortedBy = sortedBy;
					for ( indx = 0; indx < c.sortList.length; indx++ ) {
						tmp = c.sortList[ indx ];
						if ( tmp[ 0 ] === col ) {
							// order.count seems to be incorrect when compared to cell.count
							tmp[ 1 ] = order[ c.sortVars[ col ].count ];
							if ( tmp[1] === 2 ) {
								c.sortList.splice( indx, 1 );
								c.sortVars[ col ].count = -1;
							}
						}
					}
				} else {
					// add column to sort list array
					dir = order[ c.sortVars[ col ].count ];
					c.sortVars[ col ].sortedBy = sortedBy;
					if ( dir < 2 ) {
						c.sortList[ c.sortList.length ] = [ col, dir ];
						// add other columns if header spans across multiple
						if ( th.colSpan > 1 ) {
							for ( indx = 1; indx < th.colSpan; indx++ ) {
								c.sortList[ c.sortList.length ] = [ col + indx, dir ];
								// update count on columns in colSpan
								c.sortVars[ col + indx ].count = $.inArray( dir, order );
								c.sortVars[ col + indx ].sortedBy = sortedBy;
							}
						}
					}
				}
			}
			// save sort before applying sortAppend
			c.last.sortList = $.extend( [], c.sortList );
			if ( c.sortList.length && c.sortAppend ) {
				arry = $.isArray( c.sortAppend ) ? c.sortAppend : c.sortAppend[ c.sortList[ 0 ][ 0 ] ];
				if ( !ts.isEmptyObject( arry ) ) {
					for ( indx = 0; indx < arry.length; indx++ ) {
						if ( arry[ indx ][ 0 ] !== col && ts.isValueInArray( arry[ indx ][ 0 ], c.sortList ) < 0 ) {
							dir = arry[ indx ][ 1 ];
							temp = ( '' + dir ).match( /^(a|d|s|o|n)/ );
							if ( temp ) {
								tmp = c.sortList[ 0 ][ 1 ];
								switch ( temp[ 0 ] ) {
									case 'd' :
										dir = 1;
										break;
									case 's' :
										dir = tmp;
										break;
									case 'o' :
										dir = tmp === 0 ? 1 : 0;
										break;
									case 'n' :
										dir = ( tmp + 1 ) % order.length;
										break;
									default:
										dir = 0;
										break;
								}
							}
							c.sortList[ c.sortList.length ] = [ arry[ indx ][ 0 ], dir ];
							c.sortVars[ arry[ indx ][ 0 ] ].sortedBy = 'sortAppend';
						}
					}
				}
			}
			// sortBegin event triggered immediately before the sort
			c.$table.triggerHandler( 'sortBegin', table );
			// setTimeout needed so the processing icon shows up
			setTimeout( function() {
				// set css for headers
				ts.setHeadersCss( c );
				ts.multisort( c );
				ts.appendCache( c );
				c.$table.triggerHandler( 'sortBeforeEnd', table );
				c.$table.triggerHandler( 'sortEnd', table );
			}, 1 );
		},

		// sort multiple columns
		multisort : function( c ) { /*jshint loopfunc:true */
			var tbodyIndex, sortTime, colMax, rows, tmp,
				table = c.table,
				sorter = [],
				dir = 0,
				textSorter = c.textSorter || '',
				sortList = c.sortList,
				sortLen = sortList.length,
				len = c.$tbodies.length;
			if ( c.serverSideSorting || ts.isEmptyObject( c.cache ) ) {
				// empty table - fixes #206/#346
				return;
			}
			if ( ts.debug(c, 'core') ) { sortTime = new Date(); }
			// cache textSorter to optimize speed
			if ( typeof textSorter === 'object' ) {
				colMax = c.columns;
				while ( colMax-- ) {
					tmp = ts.getColumnData( table, textSorter, colMax );
					if ( typeof tmp === 'function' ) {
						sorter[ colMax ] = tmp;
					}
				}
			}
			for ( tbodyIndex = 0; tbodyIndex < len; tbodyIndex++ ) {
				colMax = c.cache[ tbodyIndex ].colMax;
				rows = c.cache[ tbodyIndex ].normalized;

				rows.sort( function( a, b ) {
					var sortIndex, num, col, order, sort, x, y;
					// rows is undefined here in IE, so don't use it!
					for ( sortIndex = 0; sortIndex < sortLen; sortIndex++ ) {
						col = sortList[ sortIndex ][ 0 ];
						order = sortList[ sortIndex ][ 1 ];
						// sort direction, true = asc, false = desc
						dir = order === 0;

						if ( c.sortStable && a[ col ] === b[ col ] && sortLen === 1 ) {
							return a[ c.columns ].order - b[ c.columns ].order;
						}

						// fallback to natural sort since it is more robust
						num = /n/i.test( ts.getSortType( c.parsers, col ) );
						if ( num && c.strings[ col ] ) {
							// sort strings in numerical columns
							if ( typeof ( ts.string[ c.strings[ col ] ] ) === 'boolean' ) {
								num = ( dir ? 1 : -1 ) * ( ts.string[ c.strings[ col ] ] ? -1 : 1 );
							} else {
								num = ( c.strings[ col ] ) ? ts.string[ c.strings[ col ] ] || 0 : 0;
							}
							// fall back to built-in numeric sort
							// var sort = $.tablesorter['sort' + s]( a[col], b[col], dir, colMax[col], table );
							sort = c.numberSorter ? c.numberSorter( a[ col ], b[ col ], dir, colMax[ col ], table ) :
								ts[ 'sortNumeric' + ( dir ? 'Asc' : 'Desc' ) ]( a[ col ], b[ col ], num, colMax[ col ], col, c );
						} else {
							// set a & b depending on sort direction
							x = dir ? a : b;
							y = dir ? b : a;
							// text sort function
							if ( typeof textSorter === 'function' ) {
								// custom OVERALL text sorter
								sort = textSorter( x[ col ], y[ col ], dir, col, table );
							} else if ( typeof sorter[ col ] === 'function' ) {
								// custom text sorter for a SPECIFIC COLUMN
								sort = sorter[ col ]( x[ col ], y[ col ], dir, col, table );
							} else {
								// fall back to natural sort
								sort = ts[ 'sortNatural' + ( dir ? 'Asc' : 'Desc' ) ]( a[ col ] || '', b[ col ] || '', col, c );
							}
						}
						if ( sort ) { return sort; }
					}
					return a[ c.columns ].order - b[ c.columns ].order;
				});
			}
			if ( ts.debug(c, 'core') ) {
				console.log( 'Applying sort ' + sortList.toString() + ts.benchmark( sortTime ) );
			}
		},

		resortComplete : function( c, callback ) {
			if ( c.table.isUpdating ) {
				c.$table.triggerHandler( 'updateComplete', c.table );
			}
			if ( $.isFunction( callback ) ) {
				callback( c.table );
			}
		},

		checkResort : function( c, resort, callback ) {
			var sortList = $.isArray( resort ) ? resort : c.sortList,
				// if no resort parameter is passed, fallback to config.resort (true by default)
				resrt = typeof resort === 'undefined' ? c.resort : resort;
			// don't try to resort if the table is still processing
			// this will catch spamming of the updateCell method
			if ( resrt !== false && !c.serverSideSorting && !c.table.isProcessing ) {
				if ( sortList.length ) {
					ts.sortOn( c, sortList, function() {
						ts.resortComplete( c, callback );
					}, true );
				} else {
					ts.sortReset( c, function() {
						ts.resortComplete( c, callback );
						ts.applyWidget( c.table, false );
					} );
				}
			} else {
				ts.resortComplete( c, callback );
				ts.applyWidget( c.table, false );
			}
		},

		sortOn : function( c, list, callback, init ) {
			var indx,
				table = c.table;
			c.$table.triggerHandler( 'sortStart', table );
			for (indx = 0; indx < c.columns; indx++) {
				c.sortVars[ indx ].sortedBy = ts.isValueInArray( indx, list ) > -1 ? 'sorton' : '';
			}
			// update header count index
			ts.updateHeaderSortCount( c, list );
			// set css for headers
			ts.setHeadersCss( c );
			// fixes #346
			if ( c.delayInit && ts.isEmptyObject( c.cache ) ) {
				ts.buildCache( c );
			}
			c.$table.triggerHandler( 'sortBegin', table );
			// sort the table and append it to the dom
			ts.multisort( c );
			ts.appendCache( c, init );
			c.$table.triggerHandler( 'sortBeforeEnd', table );
			c.$table.triggerHandler( 'sortEnd', table );
			ts.applyWidget( table );
			if ( $.isFunction( callback ) ) {
				callback( table );
			}
		},

		sortReset : function( c, callback ) {
			c.sortList = [];
			var indx;
			for (indx = 0; indx < c.columns; indx++) {
				c.sortVars[ indx ].count = -1;
				c.sortVars[ indx ].sortedBy = '';
			}
			ts.setHeadersCss( c );
			ts.multisort( c );
			ts.appendCache( c );
			if ( $.isFunction( callback ) ) {
				callback( c.table );
			}
		},

		getSortType : function( parsers, column ) {
			return ( parsers && parsers[ column ] ) ? parsers[ column ].type || '' : '';
		},

		getOrder : function( val ) {
			// look for 'd' in 'desc' order; return true
			return ( /^d/i.test( val ) || val === 1 );
		},

		// Natural sort - https://github.com/overset/javascript-natural-sort (date sorting removed)
		sortNatural : function( a, b ) {
			if ( a === b ) { return 0; }
			a = ( a || '' ).toString();
			b = ( b || '' ).toString();
			var aNum, bNum, aFloat, bFloat, indx, max,
				regex = ts.regex;
			// first try and sort Hex codes
			if ( regex.hex.test( b ) ) {
				aNum = parseInt( a.match( regex.hex ), 16 );
				bNum = parseInt( b.match( regex.hex ), 16 );
				if ( aNum < bNum ) { return -1; }
				if ( aNum > bNum ) { return 1; }
			}
			// chunk/tokenize
			aNum = a.replace( regex.chunk, '\\0$1\\0' ).replace( regex.chunks, '' ).split( '\\0' );
			bNum = b.replace( regex.chunk, '\\0$1\\0' ).replace( regex.chunks, '' ).split( '\\0' );
			max = Math.max( aNum.length, bNum.length );
			// natural sorting through split numeric strings and default strings
			for ( indx = 0; indx < max; indx++ ) {
				// find floats not starting with '0', string or 0 if not defined
				aFloat = isNaN( aNum[ indx ] ) ? aNum[ indx ] || 0 : parseFloat( aNum[ indx ] ) || 0;
				bFloat = isNaN( bNum[ indx ] ) ? bNum[ indx ] || 0 : parseFloat( bNum[ indx ] ) || 0;
				// handle numeric vs string comparison - number < string - (Kyle Adams)
				if ( isNaN( aFloat ) !== isNaN( bFloat ) ) { return isNaN( aFloat ) ? 1 : -1; }
				// rely on string comparison if different types - i.e. '02' < 2 != '02' < '2'
				if ( typeof aFloat !== typeof bFloat ) {
					aFloat += '';
					bFloat += '';
				}
				if ( aFloat < bFloat ) { return -1; }
				if ( aFloat > bFloat ) { return 1; }
			}
			return 0;
		},

		sortNaturalAsc : function( a, b, col, c ) {
			if ( a === b ) { return 0; }
			var empty = ts.string[ ( c.empties[ col ] || c.emptyTo ) ];
			if ( a === '' && empty !== 0 ) { return typeof empty === 'boolean' ? ( empty ? -1 : 1 ) : -empty || -1; }
			if ( b === '' && empty !== 0 ) { return typeof empty === 'boolean' ? ( empty ? 1 : -1 ) : empty || 1; }
			return ts.sortNatural( a, b );
		},

		sortNaturalDesc : function( a, b, col, c ) {
			if ( a === b ) { return 0; }
			var empty = ts.string[ ( c.empties[ col ] || c.emptyTo ) ];
			if ( a === '' && empty !== 0 ) { return typeof empty === 'boolean' ? ( empty ? -1 : 1 ) : empty || 1; }
			if ( b === '' && empty !== 0 ) { return typeof empty === 'boolean' ? ( empty ? 1 : -1 ) : -empty || -1; }
			return ts.sortNatural( b, a );
		},

		// basic alphabetical sort
		sortText : function( a, b ) {
			return a > b ? 1 : ( a < b ? -1 : 0 );
		},

		// return text string value by adding up ascii value
		// so the text is somewhat sorted when using a digital sort
		// this is NOT an alphanumeric sort
		getTextValue : function( val, num, max ) {
			if ( max ) {
				// make sure the text value is greater than the max numerical value (max)
				var indx,
					len = val ? val.length : 0,
					n = max + num;
				for ( indx = 0; indx < len; indx++ ) {
					n += val.charCodeAt( indx );
				}
				return num * n;
			}
			return 0;
		},

		sortNumericAsc : function( a, b, num, max, col, c ) {
			if ( a === b ) { return 0; }
			var empty = ts.string[ ( c.empties[ col ] || c.emptyTo ) ];
			if ( a === '' && empty !== 0 ) { return typeof empty === 'boolean' ? ( empty ? -1 : 1 ) : -empty || -1; }
			if ( b === '' && empty !== 0 ) { return typeof empty === 'boolean' ? ( empty ? 1 : -1 ) : empty || 1; }
			if ( isNaN( a ) ) { a = ts.getTextValue( a, num, max ); }
			if ( isNaN( b ) ) { b = ts.getTextValue( b, num, max ); }
			return a - b;
		},

		sortNumericDesc : function( a, b, num, max, col, c ) {
			if ( a === b ) { return 0; }
			var empty = ts.string[ ( c.empties[ col ] || c.emptyTo ) ];
			if ( a === '' && empty !== 0 ) { return typeof empty === 'boolean' ? ( empty ? -1 : 1 ) : empty || 1; }
			if ( b === '' && empty !== 0 ) { return typeof empty === 'boolean' ? ( empty ? 1 : -1 ) : -empty || -1; }
			if ( isNaN( a ) ) { a = ts.getTextValue( a, num, max ); }
			if ( isNaN( b ) ) { b = ts.getTextValue( b, num, max ); }
			return b - a;
		},

		sortNumeric : function( a, b ) {
			return a - b;
		},

		/*
		██ ██ ██ ██ █████▄ ▄████▄ ██████ ██████ ▄█████
		██ ██ ██ ██ ██  ██ ██ ▄▄▄ ██▄▄     ██   ▀█▄
		██ ██ ██ ██ ██  ██ ██ ▀██ ██▀▀     ██      ▀█▄
		███████▀ ██ █████▀ ▀████▀ ██████   ██   █████▀
		*/
		addWidget : function( widget ) {
			if ( widget.id && !ts.isEmptyObject( ts.getWidgetById( widget.id ) ) ) {
				console.warn( '"' + widget.id + '" widget was loaded more than once!' );
			}
			ts.widgets[ ts.widgets.length ] = widget;
		},

		hasWidget : function( $table, name ) {
			$table = $( $table );
			return $table.length && $table[ 0 ].config && $table[ 0 ].config.widgetInit[ name ] || false;
		},

		getWidgetById : function( name ) {
			var indx, widget,
				len = ts.widgets.length;
			for ( indx = 0; indx < len; indx++ ) {
				widget = ts.widgets[ indx ];
				if ( widget && widget.id && widget.id.toLowerCase() === name.toLowerCase() ) {
					return widget;
				}
			}
		},

		applyWidgetOptions : function( table ) {
			var indx, widget, wo,
				c = table.config,
				len = c.widgets.length;
			if ( len ) {
				for ( indx = 0; indx < len; indx++ ) {
					widget = ts.getWidgetById( c.widgets[ indx ] );
					if ( widget && widget.options ) {
						wo = $.extend( true, {}, widget.options );
						c.widgetOptions = $.extend( true, wo, c.widgetOptions );
						// add widgetOptions to defaults for option validator
						$.extend( true, ts.defaults.widgetOptions, widget.options );
					}
				}
			}
		},

		addWidgetFromClass : function( table ) {
			var len, indx,
				c = table.config,
				// look for widgets to apply from table class
				// don't match from 'ui-widget-content'; use \S instead of \w to include widgets
				// with dashes in the name, e.g. "widget-test-2" extracts out "test-2"
				regex = '^' + c.widgetClass.replace( ts.regex.templateName, '(\\S+)+' ) + '$',
				widgetClass = new RegExp( regex, 'g' ),
				// split up table class (widget id's can include dashes) - stop using match
				// otherwise only one widget gets extracted, see #1109
				widgets = ( table.className || '' ).split( ts.regex.spaces );
			if ( widgets.length ) {
				len = widgets.length;
				for ( indx = 0; indx < len; indx++ ) {
					if ( widgets[ indx ].match( widgetClass ) ) {
						c.widgets[ c.widgets.length ] = widgets[ indx ].replace( widgetClass, '$1' );
					}
				}
			}
		},

		applyWidgetId : function( table, id, init ) {
			table = $(table)[0];
			var applied, time, name,
				c = table.config,
				wo = c.widgetOptions,
				debug = ts.debug(c, 'core'),
				widget = ts.getWidgetById( id );
			if ( widget ) {
				name = widget.id;
				applied = false;
				// add widget name to option list so it gets reapplied after sorting, filtering, etc
				if ( $.inArray( name, c.widgets ) < 0 ) {
					c.widgets[ c.widgets.length ] = name;
				}
				if ( debug ) { time = new Date(); }

				if ( init || !( c.widgetInit[ name ] ) ) {
					// set init flag first to prevent calling init more than once (e.g. pager)
					c.widgetInit[ name ] = true;
					if ( table.hasInitialized ) {
						// don't reapply widget options on tablesorter init
						ts.applyWidgetOptions( table );
					}
					if ( typeof widget.init === 'function' ) {
						applied = true;
						if ( debug ) {
							console[ console.group ? 'group' : 'log' ]( 'Initializing ' + name + ' widget' );
						}
						widget.init( table, widget, c, wo );
					}
				}
				if ( !init && typeof widget.format === 'function' ) {
					applied = true;
					if ( debug ) {
						console[ console.group ? 'group' : 'log' ]( 'Updating ' + name + ' widget' );
					}
					widget.format( table, c, wo, false );
				}
				if ( debug ) {
					if ( applied ) {
						console.log( 'Completed ' + ( init ? 'initializing ' : 'applying ' ) + name + ' widget' + ts.benchmark( time ) );
						if ( console.groupEnd ) { console.groupEnd(); }
					}
				}
			}
		},

		applyWidget : function( table, init, callback ) {
			table = $( table )[ 0 ]; // in case this is called externally
			var indx, len, names, widget, time,
				c = table.config,
				debug = ts.debug(c, 'core'),
				widgets = [];
			// prevent numerous consecutive widget applications
			if ( init !== false && table.hasInitialized && ( table.isApplyingWidgets || table.isUpdating ) ) {
				return;
			}
			if ( debug ) { time = new Date(); }
			ts.addWidgetFromClass( table );
			// prevent "tablesorter-ready" from firing multiple times in a row
			clearTimeout( c.timerReady );
			if ( c.widgets.length ) {
				table.isApplyingWidgets = true;
				// ensure unique widget ids
				c.widgets = $.grep( c.widgets, function( val, index ) {
					return $.inArray( val, c.widgets ) === index;
				});
				names = c.widgets || [];
				len = names.length;
				// build widget array & add priority as needed
				for ( indx = 0; indx < len; indx++ ) {
					widget = ts.getWidgetById( names[ indx ] );
					if ( widget && widget.id ) {
						// set priority to 10 if not defined
						if ( !widget.priority ) { widget.priority = 10; }
						widgets[ indx ] = widget;
					} else if ( debug ) {
						console.warn( '"' + names[ indx ] + '" was enabled, but the widget code has not been loaded!' );
					}
				}
				// sort widgets by priority
				widgets.sort( function( a, b ) {
					return a.priority < b.priority ? -1 : a.priority === b.priority ? 0 : 1;
				});
				// add/update selected widgets
				len = widgets.length;
				if ( debug ) {
					console[ console.group ? 'group' : 'log' ]( 'Start ' + ( init ? 'initializing' : 'applying' ) + ' widgets' );
				}
				for ( indx = 0; indx < len; indx++ ) {
					widget = widgets[ indx ];
					if ( widget && widget.id ) {
						ts.applyWidgetId( table, widget.id, init );
					}
				}
				if ( debug && console.groupEnd ) { console.groupEnd(); }
			}
			c.timerReady = setTimeout( function() {
				table.isApplyingWidgets = false;
				$.data( table, 'lastWidgetApplication', new Date() );
				c.$table.triggerHandler( 'tablesorter-ready' );
				// callback executed on init only
				if ( !init && typeof callback === 'function' ) {
					callback( table );
				}
				if ( debug ) {
					widget = c.widgets.length;
					console.log( 'Completed ' +
						( init === true ? 'initializing ' : 'applying ' ) + widget +
						' widget' + ( widget !== 1 ? 's' : '' ) + ts.benchmark( time ) );
				}
			}, 10 );
		},

		removeWidget : function( table, name, refreshing ) {
			table = $( table )[ 0 ];
			var index, widget, indx, len,
				c = table.config;
			// if name === true, add all widgets from $.tablesorter.widgets
			if ( name === true ) {
				name = [];
				len = ts.widgets.length;
				for ( indx = 0; indx < len; indx++ ) {
					widget = ts.widgets[ indx ];
					if ( widget && widget.id ) {
						name[ name.length ] = widget.id;
					}
				}
			} else {
				// name can be either an array of widgets names,
				// or a space/comma separated list of widget names
				name = ( $.isArray( name ) ? name.join( ',' ) : name || '' ).toLowerCase().split( /[\s,]+/ );
			}
			len = name.length;
			for ( index = 0; index < len; index++ ) {
				widget = ts.getWidgetById( name[ index ] );
				indx = $.inArray( name[ index ], c.widgets );
				// don't remove the widget from config.widget if refreshing
				if ( indx >= 0 && refreshing !== true ) {
					c.widgets.splice( indx, 1 );
				}
				if ( widget && widget.remove ) {
					if ( ts.debug(c, 'core') ) {
						console.log( ( refreshing ? 'Refreshing' : 'Removing' ) + ' "' + name[ index ] + '" widget' );
					}
					widget.remove( table, c, c.widgetOptions, refreshing );
					c.widgetInit[ name[ index ] ] = false;
				}
			}
			c.$table.triggerHandler( 'widgetRemoveEnd', table );
		},

		refreshWidgets : function( table, doAll, dontapply ) {
			table = $( table )[ 0 ]; // see issue #243
			var indx, widget,
				c = table.config,
				curWidgets = c.widgets,
				widgets = ts.widgets,
				len = widgets.length,
				list = [],
				callback = function( table ) {
					$( table ).triggerHandler( 'refreshComplete' );
				};
			// remove widgets not defined in config.widgets, unless doAll is true
			for ( indx = 0; indx < len; indx++ ) {
				widget = widgets[ indx ];
				if ( widget && widget.id && ( doAll || $.inArray( widget.id, curWidgets ) < 0 ) ) {
					list[ list.length ] = widget.id;
				}
			}
			ts.removeWidget( table, list.join( ',' ), true );
			if ( dontapply !== true ) {
				// call widget init if
				ts.applyWidget( table, doAll || false, callback );
				if ( doAll ) {
					// apply widget format
					ts.applyWidget( table, false, callback );
				}
			} else {
				callback( table );
			}
		},

		/*
		██  ██ ██████ ██ ██     ██ ██████ ██ ██████ ▄█████
		██  ██   ██   ██ ██     ██   ██   ██ ██▄▄   ▀█▄
		██  ██   ██   ██ ██     ██   ██   ██ ██▀▀      ▀█▄
		▀████▀   ██   ██ ██████ ██   ██   ██ ██████ █████▀
		*/
		benchmark : function( diff ) {
			return ( ' (' + ( new Date().getTime() - diff.getTime() ) + ' ms)' );
		},
		// deprecated ts.log
		log : function() {
			console.log( arguments );
		},
		debug : function(c, name) {
			return c && (
				c.debug === true ||
				typeof c.debug === 'string' && c.debug.indexOf(name) > -1
			);
		},

		// $.isEmptyObject from jQuery v1.4
		isEmptyObject : function( obj ) {
			/*jshint forin: false */
			for ( var name in obj ) {
				return false;
			}
			return true;
		},

		isValueInArray : function( column, arry ) {
			var indx,
				len = arry && arry.length || 0;
			for ( indx = 0; indx < len; indx++ ) {
				if ( arry[ indx ][ 0 ] === column ) {
					return indx;
				}
			}
			return -1;
		},

		formatFloat : function( str, table ) {
			if ( typeof str !== 'string' || str === '' ) { return str; }
			// allow using formatFloat without a table; defaults to US number format
			var num,
				usFormat = table && table.config ? table.config.usNumberFormat !== false :
					typeof table !== 'undefined' ? table : true;
			if ( usFormat ) {
				// US Format - 1,234,567.89 -> 1234567.89
				str = str.replace( ts.regex.comma, '' );
			} else {
				// German Format = 1.234.567,89 -> 1234567.89
				// French Format = 1 234 567,89 -> 1234567.89
				str = str.replace( ts.regex.digitNonUS, '' ).replace( ts.regex.comma, '.' );
			}
			if ( ts.regex.digitNegativeTest.test( str ) ) {
				// make (#) into a negative number -> (10) = -10
				str = str.replace( ts.regex.digitNegativeReplace, '-$1' );
			}
			num = parseFloat( str );
			// return the text instead of zero
			return isNaN( num ) ? $.trim( str ) : num;
		},

		isDigit : function( str ) {
			// replace all unwanted chars and match
			return isNaN( str ) ?
				ts.regex.digitTest.test( str.toString().replace( ts.regex.digitReplace, '' ) ) :
				str !== '';
		},

		// computeTableHeaderCellIndexes from:
		// http://www.javascripttoolbox.com/lib/table/examples.php
		// http://www.javascripttoolbox.com/temp/table_cellindex.html
		computeColumnIndex : function( $rows, c ) {
			var i, j, k, l, cell, cells, rowIndex, rowSpan, colSpan, firstAvailCol,
				// total columns has been calculated, use it to set the matrixrow
				columns = c && c.columns || 0,
				matrix = [],
				matrixrow = new Array( columns );
			for ( i = 0; i < $rows.length; i++ ) {
				cells = $rows[ i ].cells;
				for ( j = 0; j < cells.length; j++ ) {
					cell = cells[ j ];
					rowIndex = i;
					rowSpan = cell.rowSpan || 1;
					colSpan = cell.colSpan || 1;
					if ( typeof matrix[ rowIndex ] === 'undefined' ) {
						matrix[ rowIndex ] = [];
					}
					// Find first available column in the first row
					for ( k = 0; k < matrix[ rowIndex ].length + 1; k++ ) {
						if ( typeof matrix[ rowIndex ][ k ] === 'undefined' ) {
							firstAvailCol = k;
							break;
						}
					}
					// jscs:disable disallowEmptyBlocks
					if ( columns && cell.cellIndex === firstAvailCol ) {
						// don't to anything
					} else if ( cell.setAttribute ) {
						// jscs:enable disallowEmptyBlocks
						// add data-column (setAttribute = IE8+)
						cell.setAttribute( 'data-column', firstAvailCol );
					} else {
						// remove once we drop support for IE7 - 1/12/2016
						$( cell ).attr( 'data-column', firstAvailCol );
					}
					for ( k = rowIndex; k < rowIndex + rowSpan; k++ ) {
						if ( typeof matrix[ k ] === 'undefined' ) {
							matrix[ k ] = [];
						}
						matrixrow = matrix[ k ];
						for ( l = firstAvailCol; l < firstAvailCol + colSpan; l++ ) {
							matrixrow[ l ] = 'x';
						}
					}
				}
			}
			ts.checkColumnCount($rows, matrix, matrixrow.length);
			return matrixrow.length;
		},

		checkColumnCount : function($rows, matrix, columns) {
			// this DOES NOT report any tbody column issues, except for the math and
			// and column selector widgets
			var i, len,
				valid = true,
				cells = [];
			for ( i = 0; i < matrix.length; i++ ) {
				// some matrix entries are undefined when testing the footer because
				// it is using the rowIndex property
				if ( matrix[i] ) {
					len = matrix[i].length;
					if ( matrix[i].length !== columns ) {
						valid = false;
						break;
					}
				}
			}
			if ( !valid ) {
				$rows.each( function( indx, el ) {
					var cell = el.parentElement.nodeName;
					if ( cells.indexOf( cell ) < 0 ) {
						cells.push( cell );
					}
				});
				console.error(
					'Invalid or incorrect number of columns in the ' +
					cells.join( ' or ' ) + '; expected ' + columns +
					', but found ' + len + ' columns'
				);
			}
		},

		// automatically add a colgroup with col elements set to a percentage width
		fixColumnWidth : function( table ) {
			table = $( table )[ 0 ];
			var overallWidth, percent, $tbodies, len, index,
				c = table.config,
				$colgroup = c.$table.children( 'colgroup' );
			// remove plugin-added colgroup, in case we need to refresh the widths
			if ( $colgroup.length && $colgroup.hasClass( ts.css.colgroup ) ) {
				$colgroup.remove();
			}
			if ( c.widthFixed && c.$table.children( 'colgroup' ).length === 0 ) {
				$colgroup = $( '<colgroup class="' + ts.css.colgroup + '">' );
				overallWidth = c.$table.width();
				// only add col for visible columns - fixes #371
				$tbodies = c.$tbodies.find( 'tr:first' ).children( ':visible' );
				len = $tbodies.length;
				for ( index = 0; index < len; index++ ) {
					percent = parseInt( ( $tbodies.eq( index ).width() / overallWidth ) * 1000, 10 ) / 10 + '%';
					$colgroup.append( $( '<col>' ).css( 'width', percent ) );
				}
				c.$table.prepend( $colgroup );
			}
		},

		// get sorter, string, empty, etc options for each column from
		// jQuery data, metadata, header option or header class name ('sorter-false')
		// priority = jQuery data > meta > headers option > header class name
		getData : function( header, configHeader, key ) {
			var meta, cl4ss,
				val = '',
				$header = $( header );
			if ( !$header.length ) { return ''; }
			meta = $.metadata ? $header.metadata() : false;
			cl4ss = ' ' + ( $header.attr( 'class' ) || '' );
			if ( typeof $header.data( key ) !== 'undefined' ||
				typeof $header.data( key.toLowerCase() ) !== 'undefined' ) {
				// 'data-lockedOrder' is assigned to 'lockedorder'; but 'data-locked-order' is assigned to 'lockedOrder'
				// 'data-sort-initial-order' is assigned to 'sortInitialOrder'
				val += $header.data( key ) || $header.data( key.toLowerCase() );
			} else if ( meta && typeof meta[ key ] !== 'undefined' ) {
				val += meta[ key ];
			} else if ( configHeader && typeof configHeader[ key ] !== 'undefined' ) {
				val += configHeader[ key ];
			} else if ( cl4ss !== ' ' && cl4ss.match( ' ' + key + '-' ) ) {
				// include sorter class name 'sorter-text', etc; now works with 'sorter-my-custom-parser'
				val = cl4ss.match( new RegExp( '\\s' + key + '-([\\w-]+)' ) )[ 1 ] || '';
			}
			return $.trim( val );
		},

		getColumnData : function( table, obj, indx, getCell, $headers ) {
			if ( typeof obj !== 'object' || obj === null ) {
				return obj;
			}
			table = $( table )[ 0 ];
			var $header, key,
				c = table.config,
				$cells = ( $headers || c.$headers ),
				// c.$headerIndexed is not defined initially
				$cell = c.$headerIndexed && c.$headerIndexed[ indx ] ||
					$cells.find( '[data-column="' + indx + '"]:last' );
			if ( typeof obj[ indx ] !== 'undefined' ) {
				return getCell ? obj[ indx ] : obj[ $cells.index( $cell ) ];
			}
			for ( key in obj ) {
				if ( typeof key === 'string' ) {
					$header = $cell
						// header cell with class/id
						.filter( key )
						// find elements within the header cell with cell/id
						.add( $cell.find( key ) );
					if ( $header.length ) {
						return obj[ key ];
					}
				}
			}
			return;
		},

		// *** Process table ***
		// add processing indicator
		isProcessing : function( $table, toggle, $headers ) {
			$table = $( $table );
			var c = $table[ 0 ].config,
				// default to all headers
				$header = $headers || $table.find( '.' + ts.css.header );
			if ( toggle ) {
				// don't use sortList if custom $headers used
				if ( typeof $headers !== 'undefined' && c.sortList.length > 0 ) {
					// get headers from the sortList
					$header = $header.filter( function() {
						// get data-column from attr to keep compatibility with jQuery 1.2.6
						return this.sortDisabled ?
							false :
							ts.isValueInArray( parseFloat( $( this ).attr( 'data-column' ) ), c.sortList ) >= 0;
					});
				}
				$table.add( $header ).addClass( ts.css.processing + ' ' + c.cssProcessing );
			} else {
				$table.add( $header ).removeClass( ts.css.processing + ' ' + c.cssProcessing );
			}
		},

		// detach tbody but save the position
		// don't use tbody because there are portions that look for a tbody index (updateCell)
		processTbody : function( table, $tb, getIt ) {
			table = $( table )[ 0 ];
			if ( getIt ) {
				table.isProcessing = true;
				$tb.before( '<colgroup class="tablesorter-savemyplace"/>' );
				return $.fn.detach ? $tb.detach() : $tb.remove();
			}
			var holdr = $( table ).find( 'colgroup.tablesorter-savemyplace' );
			$tb.insertAfter( holdr );
			holdr.remove();
			table.isProcessing = false;
		},

		clearTableBody : function( table ) {
			$( table )[ 0 ].config.$tbodies.children().detach();
		},

		// used when replacing accented characters during sorting
		characterEquivalents : {
			'a' : '\u00e1\u00e0\u00e2\u00e3\u00e4\u0105\u00e5', // áàâãäąå
			'A' : '\u00c1\u00c0\u00c2\u00c3\u00c4\u0104\u00c5', // ÁÀÂÃÄĄÅ
			'c' : '\u00e7\u0107\u010d', // çćč
			'C' : '\u00c7\u0106\u010c', // ÇĆČ
			'e' : '\u00e9\u00e8\u00ea\u00eb\u011b\u0119', // éèêëěę
			'E' : '\u00c9\u00c8\u00ca\u00cb\u011a\u0118', // ÉÈÊËĚĘ
			'i' : '\u00ed\u00ec\u0130\u00ee\u00ef\u0131', // íìİîïı
			'I' : '\u00cd\u00cc\u0130\u00ce\u00cf', // ÍÌİÎÏ
			'o' : '\u00f3\u00f2\u00f4\u00f5\u00f6\u014d', // óòôõöō
			'O' : '\u00d3\u00d2\u00d4\u00d5\u00d6\u014c', // ÓÒÔÕÖŌ
			'ss': '\u00df', // ß (s sharp)
			'SS': '\u1e9e', // ẞ (Capital sharp s)
			'u' : '\u00fa\u00f9\u00fb\u00fc\u016f', // úùûüů
			'U' : '\u00da\u00d9\u00db\u00dc\u016e' // ÚÙÛÜŮ
		},

		replaceAccents : function( str ) {
			var chr,
				acc = '[',
				eq = ts.characterEquivalents;
			if ( !ts.characterRegex ) {
				ts.characterRegexArray = {};
				for ( chr in eq ) {
					if ( typeof chr === 'string' ) {
						acc += eq[ chr ];
						ts.characterRegexArray[ chr ] = new RegExp( '[' + eq[ chr ] + ']', 'g' );
					}
				}
				ts.characterRegex = new RegExp( acc + ']' );
			}
			if ( ts.characterRegex.test( str ) ) {
				for ( chr in eq ) {
					if ( typeof chr === 'string' ) {
						str = str.replace( ts.characterRegexArray[ chr ], chr );
					}
				}
			}
			return str;
		},

		validateOptions : function( c ) {
			var setting, setting2, typ, timer,
				// ignore options containing an array
				ignore = 'headers sortForce sortList sortAppend widgets'.split( ' ' ),
				orig = c.originalSettings;
			if ( orig ) {
				if ( ts.debug(c, 'core') ) {
					timer = new Date();
				}
				for ( setting in orig ) {
					typ = typeof ts.defaults[setting];
					if ( typ === 'undefined' ) {
						console.warn( 'Tablesorter Warning! "table.config.' + setting + '" option not recognized' );
					} else if ( typ === 'object' ) {
						for ( setting2 in orig[setting] ) {
							typ = ts.defaults[setting] && typeof ts.defaults[setting][setting2];
							if ( $.inArray( setting, ignore ) < 0 && typ === 'undefined' ) {
								console.warn( 'Tablesorter Warning! "table.config.' + setting + '.' + setting2 + '" option not recognized' );
							}
						}
					}
				}
				if ( ts.debug(c, 'core') ) {
					console.log( 'validate options time:' + ts.benchmark( timer ) );
				}
			}
		},

		// restore headers
		restoreHeaders : function( table ) {
			var index, $cell,
				c = $( table )[ 0 ].config,
				$headers = c.$table.find( c.selectorHeaders ),
				len = $headers.length;
			// don't use c.$headers here in case header cells were swapped
			for ( index = 0; index < len; index++ ) {
				$cell = $headers.eq( index );
				// only restore header cells if it is wrapped
				// because this is also used by the updateAll method
				if ( $cell.find( '.' + ts.css.headerIn ).length ) {
					$cell.html( c.headerContent[ index ] );
				}
			}
		},

		destroy : function( table, removeClasses, callback ) {
			table = $( table )[ 0 ];
			if ( !table.hasInitialized ) { return; }
			// remove all widgets
			ts.removeWidget( table, true, false );
			var events,
				$t = $( table ),
				c = table.config,
				$h = $t.find( 'thead:first' ),
				$r = $h.find( 'tr.' + ts.css.headerRow ).removeClass( ts.css.headerRow + ' ' + c.cssHeaderRow ),
				$f = $t.find( 'tfoot:first > tr' ).children( 'th, td' );
			if ( removeClasses === false && $.inArray( 'uitheme', c.widgets ) >= 0 ) {
				// reapply uitheme classes, in case we want to maintain appearance
				$t.triggerHandler( 'applyWidgetId', [ 'uitheme' ] );
				$t.triggerHandler( 'applyWidgetId', [ 'zebra' ] );
			}
			// remove widget added rows, just in case
			$h.find( 'tr' ).not( $r ).remove();
			// disable tablesorter - not using .unbind( namespace ) because namespacing was
			// added in jQuery v1.4.3 - see http://api.jquery.com/event.namespace/
			events = 'sortReset update updateRows updateAll updateHeaders updateCell addRows updateComplete sorton ' +
				'appendCache updateCache applyWidgetId applyWidgets refreshWidgets removeWidget destroy mouseup mouseleave ' +
				'keypress sortBegin sortEnd resetToLoadState '.split( ' ' )
				.join( c.namespace + ' ' );
			$t
				.removeData( 'tablesorter' )
				.unbind( events.replace( ts.regex.spaces, ' ' ) );
			c.$headers
				.add( $f )
				.removeClass( [ ts.css.header, c.cssHeader, c.cssAsc, c.cssDesc, ts.css.sortAsc, ts.css.sortDesc, ts.css.sortNone ].join( ' ' ) )
				.removeAttr( 'data-column' )
				.removeAttr( 'aria-label' )
				.attr( 'aria-disabled', 'true' );
			$r
				.find( c.selectorSort )
				.unbind( ( 'mousedown mouseup keypress '.split( ' ' ).join( c.namespace + ' ' ) ).replace( ts.regex.spaces, ' ' ) );
			ts.restoreHeaders( table );
			$t.toggleClass( ts.css.table + ' ' + c.tableClass + ' tablesorter-' + c.theme, removeClasses === false );
			$t.removeClass(c.namespace.slice(1));
			// clear flag in case the plugin is initialized again
			table.hasInitialized = false;
			delete table.config.cache;
			if ( typeof callback === 'function' ) {
				callback( table );
			}
			if ( ts.debug(c, 'core') ) {
				console.log( 'tablesorter has been removed' );
			}
		}

	};

	$.fn.tablesorter = function( settings ) {
		return this.each( function() {
			var table = this,
			// merge & extend config options
			c = $.extend( true, {}, ts.defaults, settings, ts.instanceMethods );
			// save initial settings
			c.originalSettings = settings;
			// create a table from data (build table widget)
			if ( !table.hasInitialized && ts.buildTable && this.nodeName !== 'TABLE' ) {
				// return the table (in case the original target is the table's container)
				ts.buildTable( table, c );
			} else {
				ts.setup( table, c );
			}
		});
	};

	// set up debug logs
	if ( !( window.console && window.console.log ) ) {
		// access $.tablesorter.logs for browsers that don't have a console...
		ts.logs = [];
		/*jshint -W020 */
		console = {};
		console.log = console.warn = console.error = console.table = function() {
			var arg = arguments.length > 1 ? arguments : arguments[0];
			ts.logs[ ts.logs.length ] = { date: Date.now(), log: arg };
		};
	}

	// add default parsers
	ts.addParser({
		id : 'no-parser',
		is : function() {
			return false;
		},
		format : function() {
			return '';
		},
		type : 'text'
	});

	ts.addParser({
		id : 'text',
		is : function() {
			return true;
		},
		format : function( str, table ) {
			var c = table.config;
			if ( str ) {
				str = $.trim( c.ignoreCase ? str.toLocaleLowerCase() : str );
				str = c.sortLocaleCompare ? ts.replaceAccents( str ) : str;
			}
			return str;
		},
		type : 'text'
	});

	ts.regex.nondigit = /[^\w,. \-()]/g;
	ts.addParser({
		id : 'digit',
		is : function( str ) {
			return ts.isDigit( str );
		},
		format : function( str, table ) {
			var num = ts.formatFloat( ( str || '' ).replace( ts.regex.nondigit, '' ), table );
			return str && typeof num === 'number' ? num :
				str ? $.trim( str && table.config.ignoreCase ? str.toLocaleLowerCase() : str ) : str;
		},
		type : 'numeric'
	});

	ts.regex.currencyReplace = /[+\-,. ]/g;
	ts.regex.currencyTest = /^\(?\d+[\u00a3$\u20ac\u00a4\u00a5\u00a2?.]|[\u00a3$\u20ac\u00a4\u00a5\u00a2?.]\d+\)?$/;
	ts.addParser({
		id : 'currency',
		is : function( str ) {
			str = ( str || '' ).replace( ts.regex.currencyReplace, '' );
			// test for £$€¤¥¢
			return ts.regex.currencyTest.test( str );
		},
		format : function( str, table ) {
			var num = ts.formatFloat( ( str || '' ).replace( ts.regex.nondigit, '' ), table );
			return str && typeof num === 'number' ? num :
				str ? $.trim( str && table.config.ignoreCase ? str.toLocaleLowerCase() : str ) : str;
		},
		type : 'numeric'
	});

	// too many protocols to add them all https://en.wikipedia.org/wiki/URI_scheme
	// now, this regex can be updated before initialization
	ts.regex.urlProtocolTest = /^(https?|ftp|file):\/\//;
	ts.regex.urlProtocolReplace = /(https?|ftp|file):\/\/(www\.)?/;
	ts.addParser({
		id : 'url',
		is : function( str ) {
			return ts.regex.urlProtocolTest.test( str );
		},
		format : function( str ) {
			return str ? $.trim( str.replace( ts.regex.urlProtocolReplace, '' ) ) : str;
		},
		type : 'text'
	});

	ts.regex.dash = /-/g;
	ts.regex.isoDate = /^\d{4}[\/\-]\d{1,2}[\/\-]\d{1,2}/;
	ts.addParser({
		id : 'isoDate',
		is : function( str ) {
			return ts.regex.isoDate.test( str );
		},
		format : function( str ) {
			var date = str ? new Date( str.replace( ts.regex.dash, '/' ) ) : str;
			return date instanceof Date && isFinite( date ) ? date.getTime() : str;
		},
		type : 'numeric'
	});

	ts.regex.percent = /%/g;
	ts.regex.percentTest = /(\d\s*?%|%\s*?\d)/;
	ts.addParser({
		id : 'percent',
		is : function( str ) {
			return ts.regex.percentTest.test( str ) && str.length < 15;
		},
		format : function( str, table ) {
			return str ? ts.formatFloat( str.replace( ts.regex.percent, '' ), table ) : str;
		},
		type : 'numeric'
	});

	// added image parser to core v2.17.9
	ts.addParser({
		id : 'image',
		is : function( str, table, node, $node ) {
			return $node.find( 'img' ).length > 0;
		},
		format : function( str, table, cell ) {
			return $( cell ).find( 'img' ).attr( table.config.imgAttr || 'alt' ) || str;
		},
		parsed : true, // filter widget flag
		type : 'text'
	});

	ts.regex.dateReplace = /(\S)([AP]M)$/i; // used by usLongDate & time parser
	ts.regex.usLongDateTest1 = /^[A-Z]{3,10}\.?\s+\d{1,2},?\s+(\d{4})(\s+\d{1,2}:\d{2}(:\d{2})?(\s+[AP]M)?)?$/i;
	ts.regex.usLongDateTest2 = /^\d{1,2}\s+[A-Z]{3,10}\s+\d{4}/i;
	ts.addParser({
		id : 'usLongDate',
		is : function( str ) {
			// two digit years are not allowed cross-browser
			// Jan 01, 2013 12:34:56 PM or 01 Jan 2013
			return ts.regex.usLongDateTest1.test( str ) || ts.regex.usLongDateTest2.test( str );
		},
		format : function( str ) {
			var date = str ? new Date( str.replace( ts.regex.dateReplace, '$1 $2' ) ) : str;
			return date instanceof Date && isFinite( date ) ? date.getTime() : str;
		},
		type : 'numeric'
	});

	// testing for ##-##-#### or ####-##-##, so it's not perfect; time can be included
	ts.regex.shortDateTest = /(^\d{1,2}[\/\s]\d{1,2}[\/\s]\d{4})|(^\d{4}[\/\s]\d{1,2}[\/\s]\d{1,2})/;
	// escaped "-" because JSHint in Firefox was showing it as an error
	ts.regex.shortDateReplace = /[\-.,]/g;
	// XXY covers MDY & DMY formats
	ts.regex.shortDateXXY = /(\d{1,2})[\/\s](\d{1,2})[\/\s](\d{4})/;
	ts.regex.shortDateYMD = /(\d{4})[\/\s](\d{1,2})[\/\s](\d{1,2})/;
	ts.convertFormat = function( dateString, format ) {
		dateString = ( dateString || '' )
			.replace( ts.regex.spaces, ' ' )
			.replace( ts.regex.shortDateReplace, '/' );
		if ( format === 'mmddyyyy' ) {
			dateString = dateString.replace( ts.regex.shortDateXXY, '$3/$1/$2' );
		} else if ( format === 'ddmmyyyy' ) {
			dateString = dateString.replace( ts.regex.shortDateXXY, '$3/$2/$1' );
		} else if ( format === 'yyyymmdd' ) {
			dateString = dateString.replace( ts.regex.shortDateYMD, '$1/$2/$3' );
		}
		var date = new Date( dateString );
		return date instanceof Date && isFinite( date ) ? date.getTime() : '';
	};

	ts.addParser({
		id : 'shortDate', // 'mmddyyyy', 'ddmmyyyy' or 'yyyymmdd'
		is : function( str ) {
			str = ( str || '' ).replace( ts.regex.spaces, ' ' ).replace( ts.regex.shortDateReplace, '/' );
			return ts.regex.shortDateTest.test( str );
		},
		format : function( str, table, cell, cellIndex ) {
			if ( str ) {
				var c = table.config,
					$header = c.$headerIndexed[ cellIndex ],
					format = $header.length && $header.data( 'dateFormat' ) ||
						ts.getData( $header, ts.getColumnData( table, c.headers, cellIndex ), 'dateFormat' ) ||
						c.dateFormat;
				// save format because getData can be slow...
				if ( $header.length ) {
					$header.data( 'dateFormat', format );
				}
				return ts.convertFormat( str, format ) || str;
			}
			return str;
		},
		type : 'numeric'
	});

	// match 24 hour time & 12 hours time + am/pm - see http://regexr.com/3c3tk
	ts.regex.timeTest = /^(0?[1-9]|1[0-2]):([0-5]\d)(\s[AP]M)$|^((?:[01]\d|[2][0-4]):[0-5]\d)$/i;
	ts.regex.timeMatch = /(0?[1-9]|1[0-2]):([0-5]\d)(\s[AP]M)|((?:[01]\d|[2][0-4]):[0-5]\d)/i;
	ts.addParser({
		id : 'time',
		is : function( str ) {
			return ts.regex.timeTest.test( str );
		},
		format : function( str ) {
			// isolate time... ignore month, day and year
			var temp,
				timePart = ( str || '' ).match( ts.regex.timeMatch ),
				orig = new Date( str ),
				// no time component? default to 00:00 by leaving it out, but only if str is defined
				time = str && ( timePart !== null ? timePart[ 0 ] : '00:00 AM' ),
				date = time ? new Date( '2000/01/01 ' + time.replace( ts.regex.dateReplace, '$1 $2' ) ) : time;
			if ( date instanceof Date && isFinite( date ) ) {
				temp = orig instanceof Date && isFinite( orig ) ? orig.getTime() : 0;
				// if original string was a valid date, add it to the decimal so the column sorts in some kind of order
				// luckily new Date() ignores the decimals
				return temp ? parseFloat( date.getTime() + '.' + orig.getTime() ) : date.getTime();
			}
			return str;
		},
		type : 'numeric'
	});

	ts.addParser({
		id : 'metadata',
		is : function() {
			return false;
		},
		format : function( str, table, cell ) {
			var c = table.config,
			p = ( !c.parserMetadataName ) ? 'sortValue' : c.parserMetadataName;
			return $( cell ).metadata()[ p ];
		},
		type : 'numeric'
	});

	/*
		██████ ██████ █████▄ █████▄ ▄████▄
		  ▄█▀  ██▄▄   ██▄▄██ ██▄▄██ ██▄▄██
		▄█▀    ██▀▀   ██▀▀██ ██▀▀█  ██▀▀██
		██████ ██████ █████▀ ██  ██ ██  ██
		*/
	// add default widgets
	ts.addWidget({
		id : 'zebra',
		priority : 90,
		format : function( table, c, wo ) {
			var $visibleRows, $row, count, isEven, tbodyIndex, rowIndex, len,
				child = new RegExp( c.cssChildRow, 'i' ),
				$tbodies = c.$tbodies.add( $( c.namespace + '_extra_table' ).children( 'tbody:not(.' + c.cssInfoBlock + ')' ) );
			for ( tbodyIndex = 0; tbodyIndex < $tbodies.length; tbodyIndex++ ) {
				// loop through the visible rows
				count = 0;
				$visibleRows = $tbodies.eq( tbodyIndex ).children( 'tr:visible' ).not( c.selectorRemove );
				len = $visibleRows.length;
				for ( rowIndex = 0; rowIndex < len; rowIndex++ ) {
					$row = $visibleRows.eq( rowIndex );
					// style child rows the same way the parent row was styled
					if ( !child.test( $row[ 0 ].className ) ) { count++; }
					isEven = ( count % 2 === 0 );
					$row
						.removeClass( wo.zebra[ isEven ? 1 : 0 ] )
						.addClass( wo.zebra[ isEven ? 0 : 1 ] );
				}
			}
		},
		remove : function( table, c, wo, refreshing ) {
			if ( refreshing ) { return; }
			var tbodyIndex, $tbody,
				$tbodies = c.$tbodies,
				toRemove = ( wo.zebra || [ 'even', 'odd' ] ).join( ' ' );
			for ( tbodyIndex = 0; tbodyIndex < $tbodies.length; tbodyIndex++ ) {
				$tbody = ts.processTbody( table, $tbodies.eq( tbodyIndex ), true ); // remove tbody
				$tbody.children().removeClass( toRemove );
				ts.processTbody( table, $tbody, false ); // restore tbody
			}
		}
	});

})( jQuery );

/*! Widget: storage - updated 2018-03-18 (v2.30.0) */
/*global JSON:false */
;(function ($, window, document) {
	'use strict';

	var ts = $.tablesorter || {};

	// update defaults for validator; these values must be falsy!
	$.extend(true, ts.defaults, {
		fixedUrl: '',
		widgetOptions: {
			storage_fixedUrl: '',
			storage_group: '',
			storage_page: '',
			storage_storageType: '',
			storage_tableId: '',
			storage_useSessionStorage: ''
		}
	});

	// *** Store data in local storage, with a cookie fallback ***
	/* IE7 needs JSON library for JSON.stringify - (http://caniuse.com/#search=json)
	   if you need it, then include https://github.com/douglascrockford/JSON-js

	   $.parseJSON is not available is jQuery versions older than 1.4.1, using older
	   versions will only allow storing information for one page at a time

	   // *** Save data (JSON format only) ***
	   // val must be valid JSON... use http://jsonlint.com/ to ensure it is valid
	   var val = { "mywidget" : "data1" }; // valid JSON uses double quotes
	   // $.tablesorter.storage(table, key, val);
	   $.tablesorter.storage(table, 'tablesorter-mywidget', val);

	   // *** Get data: $.tablesorter.storage(table, key); ***
	   v = $.tablesorter.storage(table, 'tablesorter-mywidget');
	   // val may be empty, so also check for your data
	   val = (v && v.hasOwnProperty('mywidget')) ? v.mywidget : '';
	   alert(val); // 'data1' if saved, or '' if not
	*/
	ts.storage = function(table, key, value, options) {
		table = $(table)[0];
		var cookieIndex, cookies, date,
			hasStorage = false,
			values = {},
			c = table.config,
			wo = c && c.widgetOptions,
			debug = ts.debug(c, 'storage'),
			storageType = (
				( options && options.storageType ) || ( wo && wo.storage_storageType )
			).toString().charAt(0).toLowerCase(),
			// deprecating "useSessionStorage"; any storageType setting overrides it
			session = storageType ? '' :
				( options && options.useSessionStorage ) || ( wo && wo.storage_useSessionStorage ),
			$table = $(table),
			// id from (1) options ID, (2) table 'data-table-group' attribute, (3) widgetOptions.storage_tableId,
			// (4) table ID, then (5) table index
			id = options && options.id ||
				$table.attr( options && options.group || wo && wo.storage_group || 'data-table-group') ||
				wo && wo.storage_tableId || table.id || $('.tablesorter').index( $table ),
			// url from (1) options url, (2) table 'data-table-page' attribute, (3) widgetOptions.storage_fixedUrl,
			// (4) table.config.fixedUrl (deprecated), then (5) window location path
			url = options && options.url ||
				$table.attr(options && options.page || wo && wo.storage_page || 'data-table-page') ||
				wo && wo.storage_fixedUrl || c && c.fixedUrl || window.location.pathname;

		// skip if using cookies
		if (storageType !== 'c') {
			storageType = (storageType === 's' || session) ? 'sessionStorage' : 'localStorage';
			// https://gist.github.com/paulirish/5558557
			if (storageType in window) {
				try {
					window[storageType].setItem('_tmptest', 'temp');
					hasStorage = true;
					window[storageType].removeItem('_tmptest');
				} catch (error) {
					console.warn( storageType + ' is not supported in this browser' );
				}
			}
		}
		if (debug) {
			console.log('Storage >> Using', hasStorage ? storageType : 'cookies');
		}
		// *** get value ***
		if ($.parseJSON) {
			if (hasStorage) {
				values = $.parseJSON( window[storageType][key] || 'null' ) || {};
			} else {
				// old browser, using cookies
				cookies = document.cookie.split(/[;\s|=]/);
				// add one to get from the key to the value
				cookieIndex = $.inArray(key, cookies) + 1;
				values = (cookieIndex !== 0) ? $.parseJSON(cookies[cookieIndex] || 'null') || {} : {};
			}
		}
		// allow value to be an empty string too
		if (typeof value !== 'undefined' && window.JSON && JSON.hasOwnProperty('stringify')) {
			// add unique identifiers = url pathname > table ID/index on page > data
			if (!values[url]) {
				values[url] = {};
			}
			values[url][id] = value;
			// *** set value ***
			if (hasStorage) {
				window[storageType][key] = JSON.stringify(values);
			} else {
				date = new Date();
				date.setTime(date.getTime() + (31536e+6)); // 365 days
				document.cookie = key + '=' + (JSON.stringify(values)).replace(/\"/g, '\"') + '; expires=' + date.toGMTString() + '; path=/';
			}
		} else {
			return values && values[url] ? values[url][id] : '';
		}
	};

})(jQuery, window, document);

/*! Widget: uitheme - updated 2018-03-18 (v2.30.0) */
;(function ($) {
	'use strict';
	var ts = $.tablesorter || {};

	ts.themes = {
		'bootstrap' : {
			table        : 'table table-bordered table-striped',
			caption      : 'caption',
			// header class names
			header       : 'bootstrap-header', // give the header a gradient background (theme.bootstrap_2.css)
			sortNone     : '',
			sortAsc      : '',
			sortDesc     : '',
			active       : '', // applied when column is sorted
			hover        : '', // custom css required - a defined bootstrap style may not override other classes
			// icon class names
			icons        : '', // add 'bootstrap-icon-white' to make them white; this icon class is added to the <i> in the header
			iconSortNone : 'bootstrap-icon-unsorted', // class name added to icon when column is not sorted
			iconSortAsc  : 'glyphicon glyphicon-chevron-up', // class name added to icon when column has ascending sort
			iconSortDesc : 'glyphicon glyphicon-chevron-down', // class name added to icon when column has descending sort
			filterRow    : '', // filter row class
			footerRow    : '',
			footerCells  : '',
			even         : '', // even row zebra striping
			odd          : ''  // odd row zebra striping
		},
		'jui' : {
			table        : 'ui-widget ui-widget-content ui-corner-all', // table classes
			caption      : 'ui-widget-content',
			// header class names
			header       : 'ui-widget-header ui-corner-all ui-state-default', // header classes
			sortNone     : '',
			sortAsc      : '',
			sortDesc     : '',
			active       : 'ui-state-active', // applied when column is sorted
			hover        : 'ui-state-hover',  // hover class
			// icon class names
			icons        : 'ui-icon', // icon class added to the <i> in the header
			iconSortNone : 'ui-icon-carat-2-n-s ui-icon-caret-2-n-s', // class name added to icon when column is not sorted
			iconSortAsc  : 'ui-icon-carat-1-n ui-icon-caret-1-n', // class name added to icon when column has ascending sort
			iconSortDesc : 'ui-icon-carat-1-s ui-icon-caret-1-s', // class name added to icon when column has descending sort
			filterRow    : '',
			footerRow    : '',
			footerCells  : '',
			even         : 'ui-widget-content', // even row zebra striping
			odd          : 'ui-state-default'   // odd row zebra striping
		}
	};

	$.extend(ts.css, {
		wrapper : 'tablesorter-wrapper' // ui theme & resizable
	});

	ts.addWidget({
		id: 'uitheme',
		priority: 10,
		format: function(table, c, wo) {
			var i, tmp, hdr, icon, time, $header, $icon, $tfoot, $h, oldtheme, oldremove, oldIconRmv, hasOldTheme,
				themesAll = ts.themes,
				$table = c.$table.add( $( c.namespace + '_extra_table' ) ),
				$headers = c.$headers.add( $( c.namespace + '_extra_headers' ) ),
				theme = c.theme || 'jui',
				themes = themesAll[theme] || {},
				remove = $.trim( [ themes.sortNone, themes.sortDesc, themes.sortAsc, themes.active ].join( ' ' ) ),
				iconRmv = $.trim( [ themes.iconSortNone, themes.iconSortDesc, themes.iconSortAsc ].join( ' ' ) ),
				debug = ts.debug(c, 'uitheme');
			if (debug) { time = new Date(); }
			// initialization code - run once
			if (!$table.hasClass('tablesorter-' + theme) || c.theme !== c.appliedTheme || !wo.uitheme_applied) {
				wo.uitheme_applied = true;
				oldtheme = themesAll[c.appliedTheme] || {};
				hasOldTheme = !$.isEmptyObject(oldtheme);
				oldremove =  hasOldTheme ? [ oldtheme.sortNone, oldtheme.sortDesc, oldtheme.sortAsc, oldtheme.active ].join( ' ' ) : '';
				oldIconRmv = hasOldTheme ? [ oldtheme.iconSortNone, oldtheme.iconSortDesc, oldtheme.iconSortAsc ].join( ' ' ) : '';
				if (hasOldTheme) {
					wo.zebra[0] = $.trim( ' ' + wo.zebra[0].replace(' ' + oldtheme.even, '') );
					wo.zebra[1] = $.trim( ' ' + wo.zebra[1].replace(' ' + oldtheme.odd, '') );
					c.$tbodies.children().removeClass( [ oldtheme.even, oldtheme.odd ].join(' ') );
				}
				// update zebra stripes
				if (themes.even) { wo.zebra[0] += ' ' + themes.even; }
				if (themes.odd) { wo.zebra[1] += ' ' + themes.odd; }
				// add caption style
				$table.children('caption')
					.removeClass(oldtheme.caption || '')
					.addClass(themes.caption);
				// add table/footer class names
				$tfoot = $table
					// remove other selected themes
					.removeClass( (c.appliedTheme ? 'tablesorter-' + (c.appliedTheme || '') : '') + ' ' + (oldtheme.table || '') )
					.addClass('tablesorter-' + theme + ' ' + (themes.table || '')) // add theme widget class name
					.children('tfoot');
				c.appliedTheme = c.theme;

				if ($tfoot.length) {
					$tfoot
						// if oldtheme.footerRow or oldtheme.footerCells are undefined, all class names are removed
						.children('tr').removeClass(oldtheme.footerRow || '').addClass(themes.footerRow)
						.children('th, td').removeClass(oldtheme.footerCells || '').addClass(themes.footerCells);
				}
				// update header classes
				$headers
					.removeClass( (hasOldTheme ? [ oldtheme.header, oldtheme.hover, oldremove ].join(' ') : '') || '' )
					.addClass(themes.header)
					.not('.sorter-false')
					.unbind('mouseenter.tsuitheme mouseleave.tsuitheme')
					.bind('mouseenter.tsuitheme mouseleave.tsuitheme', function(event) {
						// toggleClass with switch added in jQuery 1.3
						$(this)[ event.type === 'mouseenter' ? 'addClass' : 'removeClass' ](themes.hover || '');
					});

				$headers.each(function() {
					var $this = $(this);
					if (!$this.find('.' + ts.css.wrapper).length) {
						// Firefox needs this inner div to position the icon & resizer correctly
						$this.wrapInner('<div class="' + ts.css.wrapper + '" style="position:relative;height:100%;width:100%"></div>');
					}
				});
				if (c.cssIcon) {
					// if c.cssIcon is '', then no <i> is added to the header
					$headers
						.find('.' + ts.css.icon)
						.removeClass(hasOldTheme ? [ oldtheme.icons, oldIconRmv ].join(' ') : '')
						.addClass(themes.icons || '');
				}
				// filter widget initializes after uitheme
				if (ts.hasWidget( c.table, 'filter' )) {
					tmp = function() {
						$table.children('thead').children('.' + ts.css.filterRow)
							.removeClass(hasOldTheme ? oldtheme.filterRow || '' : '')
							.addClass(themes.filterRow || '');
					};
					if (wo.filter_initialized) {
						tmp();
					} else {
						$table.one('filterInit', function() {
							tmp();
						});
					}
				}
			}
			for (i = 0; i < c.columns; i++) {
				$header = c.$headers
					.add($(c.namespace + '_extra_headers'))
					.not('.sorter-false')
					.filter('[data-column="' + i + '"]');
				$icon = (ts.css.icon) ? $header.find('.' + ts.css.icon) : $();
				$h = $headers.not('.sorter-false').filter('[data-column="' + i + '"]:last');
				if ($h.length) {
					$header.removeClass(remove);
					$icon.removeClass(iconRmv);
					if ($h[0].sortDisabled) {
						// no sort arrows for disabled columns!
						$icon.removeClass(themes.icons || '');
					} else {
						hdr = themes.sortNone;
						icon = themes.iconSortNone;
						if ($h.hasClass(ts.css.sortAsc)) {
							hdr = [ themes.sortAsc, themes.active ].join(' ');
							icon = themes.iconSortAsc;
						} else if ($h.hasClass(ts.css.sortDesc)) {
							hdr = [ themes.sortDesc, themes.active ].join(' ');
							icon = themes.iconSortDesc;
						}
						$header.addClass(hdr);
						$icon.addClass(icon || '');
					}
				}
			}
			if (debug) {
				console.log('uitheme >> Applied ' + theme + ' theme' + ts.benchmark(time));
			}
		},
		remove: function(table, c, wo, refreshing) {
			if (!wo.uitheme_applied) { return; }
			var $table = c.$table,
				theme = c.appliedTheme || 'jui',
				themes = ts.themes[ theme ] || ts.themes.jui,
				$headers = $table.children('thead').children(),
				remove = themes.sortNone + ' ' + themes.sortDesc + ' ' + themes.sortAsc,
				iconRmv = themes.iconSortNone + ' ' + themes.iconSortDesc + ' ' + themes.iconSortAsc;
			$table.removeClass('tablesorter-' + theme + ' ' + themes.table);
			wo.uitheme_applied = false;
			if (refreshing) { return; }
			$table.find(ts.css.header).removeClass(themes.header);
			$headers
				.unbind('mouseenter.tsuitheme mouseleave.tsuitheme') // remove hover
				.removeClass(themes.hover + ' ' + remove + ' ' + themes.active)
				.filter('.' + ts.css.filterRow)
				.removeClass(themes.filterRow);
			$headers.find('.' + ts.css.icon).removeClass(themes.icons + ' ' + iconRmv);
		}
	});

})(jQuery);

/*! Widget: columns - updated 5/24/2017 (v2.28.11) */
;(function ($) {
	'use strict';
	var ts = $.tablesorter || {};

	ts.addWidget({
		id: 'columns',
		priority: 65,
		options : {
			columns : [ 'primary', 'secondary', 'tertiary' ]
		},
		format: function(table, c, wo) {
			var $tbody, tbodyIndex, $rows, rows, $row, $cells, remove, indx,
			$table = c.$table,
			$tbodies = c.$tbodies,
			sortList = c.sortList,
			len = sortList.length,
			// removed c.widgetColumns support
			css = wo && wo.columns || [ 'primary', 'secondary', 'tertiary' ],
			last = css.length - 1;
			remove = css.join(' ');
			// check if there is a sort (on initialization there may not be one)
			for (tbodyIndex = 0; tbodyIndex < $tbodies.length; tbodyIndex++ ) {
				$tbody = ts.processTbody(table, $tbodies.eq(tbodyIndex), true); // detach tbody
				$rows = $tbody.children('tr');
				// loop through the visible rows
				$rows.each(function() {
					$row = $(this);
					if (this.style.display !== 'none') {
						// remove all columns class names
						$cells = $row.children().removeClass(remove);
						// add appropriate column class names
						if (sortList && sortList[0]) {
							// primary sort column class
							$cells.eq(sortList[0][0]).addClass(css[0]);
							if (len > 1) {
								for (indx = 1; indx < len; indx++) {
									// secondary, tertiary, etc sort column classes
									$cells.eq(sortList[indx][0]).addClass( css[indx] || css[last] );
								}
							}
						}
					}
				});
				ts.processTbody(table, $tbody, false);
			}
			// add classes to thead and tfoot
			rows = wo.columns_thead !== false ? [ 'thead tr' ] : [];
			if (wo.columns_tfoot !== false) {
				rows.push('tfoot tr');
			}
			if (rows.length) {
				$rows = $table.find( rows.join(',') ).children().removeClass(remove);
				if (len) {
					for (indx = 0; indx < len; indx++) {
						// add primary. secondary, tertiary, etc sort column classes
						$rows.filter('[data-column="' + sortList[indx][0] + '"]').addClass(css[indx] || css[last]);
					}
				}
			}
		},
		remove: function(table, c, wo) {
			var tbodyIndex, $tbody,
				$tbodies = c.$tbodies,
				remove = (wo.columns || [ 'primary', 'secondary', 'tertiary' ]).join(' ');
			c.$headers.removeClass(remove);
			c.$table.children('tfoot').children('tr').children('th, td').removeClass(remove);
			for (tbodyIndex = 0; tbodyIndex < $tbodies.length; tbodyIndex++ ) {
				$tbody = ts.processTbody(table, $tbodies.eq(tbodyIndex), true); // remove tbody
				$tbody.children('tr').each(function() {
					$(this).children().removeClass(remove);
				});
				ts.processTbody(table, $tbody, false); // restore tbody
			}
		}
	});

})(jQuery);

/*! Widget: filter - updated 2018-03-18 (v2.30.0) *//*
 * Requires tablesorter v2.8+ and jQuery 1.7+
 * by Rob Garrison
 */
;( function ( $ ) {
	'use strict';
	var tsf, tsfRegex,
		ts = $.tablesorter || {},
		tscss = ts.css,
		tskeyCodes = ts.keyCodes;

	$.extend( tscss, {
		filterRow      : 'tablesorter-filter-row',
		filter         : 'tablesorter-filter',
		filterDisabled : 'disabled',
		filterRowHide  : 'hideme'
	});

	$.extend( tskeyCodes, {
		backSpace : 8,
		escape : 27,
		space : 32,
		left : 37,
		down : 40
	});

	ts.addWidget({
		id: 'filter',
		priority: 50,
		options : {
			filter_cellFilter    : '',    // css class name added to the filter cell ( string or array )
			filter_childRows     : false, // if true, filter includes child row content in the search
			filter_childByColumn : false, // ( filter_childRows must be true ) if true = search child rows by column; false = search all child row text grouped
			filter_childWithSibs : true,  // if true, include matching child row siblings
			filter_columnAnyMatch: true,  // if true, allows using '#:{query}' in AnyMatch searches ( column:query )
			filter_columnFilters : true,  // if true, a filter will be added to the top of each table column
			filter_cssFilter     : '',    // css class name added to the filter row & each input in the row ( tablesorter-filter is ALWAYS added )
			filter_defaultAttrib : 'data-value', // data attribute in the header cell that contains the default filter value
			filter_defaultFilter : {},    // add a default column filter type '~{query}' to make fuzzy searches default; '{q1} AND {q2}' to make all searches use a logical AND.
			filter_excludeFilter : {},    // filters to exclude, per column
			filter_external      : '',    // jQuery selector string ( or jQuery object ) of external filters
			filter_filteredRow   : 'filtered', // class added to filtered rows; define in css with "display:none" to hide the filtered-out rows
			filter_filterLabel   : 'Filter "{{label}}" column by...', // Aria-label added to filter input/select; see #1495
			filter_formatter     : null,  // add custom filter elements to the filter row
			filter_functions     : null,  // add custom filter functions using this option
			filter_hideEmpty     : true,  // hide filter row when table is empty
			filter_hideFilters   : false, // collapse filter row when mouse leaves the area
			filter_ignoreCase    : true,  // if true, make all searches case-insensitive
			filter_liveSearch    : true,  // if true, search column content while the user types ( with a delay )
			filter_matchType     : { 'input': 'exact', 'select': 'exact' }, // global query settings ('exact' or 'match'); overridden by "filter-match" or "filter-exact" class
			filter_onlyAvail     : 'filter-onlyAvail', // a header with a select dropdown & this class name will only show available ( visible ) options within the drop down
			filter_placeholder   : { search : '', select : '' }, // default placeholder text ( overridden by any header 'data-placeholder' setting )
			filter_reset         : null,  // jQuery selector string of an element used to reset the filters
			filter_resetOnEsc    : true,  // Reset filter input when the user presses escape - normalized across browsers
			filter_saveFilters   : false, // Use the $.tablesorter.storage utility to save the most recent filters
			filter_searchDelay   : 300,   // typing delay in milliseconds before starting a search
			filter_searchFiltered: true,  // allow searching through already filtered rows in special circumstances; will speed up searching in large tables if true
			filter_selectSource  : null,  // include a function to return an array of values to be added to the column filter select
			filter_selectSourceSeparator : '|', // filter_selectSource array text left of the separator is added to the option value, right into the option text
			filter_serversideFiltering : false, // if true, must perform server-side filtering b/c client-side filtering is disabled, but the ui and events will still be used.
			filter_startsWith    : false, // if true, filter start from the beginning of the cell contents
			filter_useParsedData : false  // filter all data using parsed content
		},
		format: function( table, c, wo ) {
			if ( !c.$table.hasClass( 'hasFilters' ) ) {
				tsf.init( table, c, wo );
			}
		},
		remove: function( table, c, wo, refreshing ) {
			var tbodyIndex, $tbody,
				$table = c.$table,
				$tbodies = c.$tbodies,
				events = (
					'addRows updateCell update updateRows updateComplete appendCache filterReset ' +
					'filterAndSortReset filterFomatterUpdate filterEnd search stickyHeadersInit '
				).split( ' ' ).join( c.namespace + 'filter ' );
			$table
				.removeClass( 'hasFilters' )
				// add filter namespace to all BUT search
				.unbind( events.replace( ts.regex.spaces, ' ' ) )
				// remove the filter row even if refreshing, because the column might have been moved
				.find( '.' + tscss.filterRow ).remove();
			wo.filter_initialized = false;
			if ( refreshing ) { return; }
			for ( tbodyIndex = 0; tbodyIndex < $tbodies.length; tbodyIndex++ ) {
				$tbody = ts.processTbody( table, $tbodies.eq( tbodyIndex ), true ); // remove tbody
				$tbody.children().removeClass( wo.filter_filteredRow ).show();
				ts.processTbody( table, $tbody, false ); // restore tbody
			}
			if ( wo.filter_reset ) {
				$( document ).undelegate( wo.filter_reset, 'click' + c.namespace + 'filter' );
			}
		}
	});

	tsf = ts.filter = {

		// regex used in filter 'check' functions - not for general use and not documented
		regex: {
			regex     : /^\/((?:\\\/|[^\/])+)\/([migyu]{0,5})?$/, // regex to test for regex
			child     : /tablesorter-childRow/, // child row class name; this gets updated in the script
			filtered  : /filtered/, // filtered (hidden) row class name; updated in the script
			type      : /undefined|number/, // check type
			exact     : /(^[\"\'=]+)|([\"\'=]+$)/g, // exact match (allow '==')
			operators : /[<>=]/g, // replace operators
			query     : '(q|query)', // replace filter queries
			wild01    : /\?/g, // wild card match 0 or 1
			wild0More : /\*/g, // wild care match 0 or more
			quote     : /\"/g,
			isNeg1    : /(>=?\s*-\d)/,
			isNeg2    : /(<=?\s*\d)/
		},
		// function( c, data ) { }
		// c = table.config
		// data.$row = jQuery object of the row currently being processed
		// data.$cells = jQuery object of all cells within the current row
		// data.filters = array of filters for all columns ( some may be undefined )
		// data.filter = filter for the current column
		// data.iFilter = same as data.filter, except lowercase ( if wo.filter_ignoreCase is true )
		// data.exact = table cell text ( or parsed data if column parser enabled; may be a number & not a string )
		// data.iExact = same as data.exact, except lowercase ( if wo.filter_ignoreCase is true; may be a number & not a string )
		// data.cache = table cell text from cache, so it has been parsed ( & in all lower case if c.ignoreCase is true )
		// data.cacheArray = An array of parsed content from each table cell in the row being processed
		// data.index = column index; table = table element ( DOM )
		// data.parsed = array ( by column ) of boolean values ( from filter_useParsedData or 'filter-parsed' class )
		types: {
			or : function( c, data, vars ) {
				// look for "|", but not if it is inside of a regular expression
				if ( ( tsfRegex.orTest.test( data.iFilter ) || tsfRegex.orSplit.test( data.filter ) ) &&
					// this test for regex has potential to slow down the overall search
					!tsfRegex.regex.test( data.filter ) ) {
					var indx, filterMatched, query, regex,
						// duplicate data but split filter
						data2 = $.extend( {}, data ),
						filter = data.filter.split( tsfRegex.orSplit ),
						iFilter = data.iFilter.split( tsfRegex.orSplit ),
						len = filter.length;
					for ( indx = 0; indx < len; indx++ ) {
						data2.nestedFilters = true;
						data2.filter = '' + ( tsf.parseFilter( c, filter[ indx ], data ) || '' );
						data2.iFilter = '' + ( tsf.parseFilter( c, iFilter[ indx ], data ) || '' );
						query = '(' + ( tsf.parseFilter( c, data2.filter, data ) || '' ) + ')';
						try {
							// use try/catch, because query may not be a valid regex if "|" is contained within a partial regex search,
							// e.g "/(Alex|Aar" -> Uncaught SyntaxError: Invalid regular expression: /(/(Alex)/: Unterminated group
							regex = new RegExp( data.isMatch ? query : '^' + query + '$', c.widgetOptions.filter_ignoreCase ? 'i' : '' );
							// filterMatched = data2.filter === '' && indx > 0 ? true
							// look for an exact match with the 'or' unless the 'filter-match' class is found
							filterMatched = regex.test( data2.exact ) || tsf.processTypes( c, data2, vars );
							if ( filterMatched ) {
								return filterMatched;
							}
						} catch ( error ) {
							return null;
						}
					}
					// may be null from processing types
					return filterMatched || false;
				}
				return null;
			},
			// Look for an AND or && operator ( logical and )
			and : function( c, data, vars ) {
				if ( tsfRegex.andTest.test( data.filter ) ) {
					var indx, filterMatched, result, query, regex,
						// duplicate data but split filter
						data2 = $.extend( {}, data ),
						filter = data.filter.split( tsfRegex.andSplit ),
						iFilter = data.iFilter.split( tsfRegex.andSplit ),
						len = filter.length;
					for ( indx = 0; indx < len; indx++ ) {
						data2.nestedFilters = true;
						data2.filter = '' + ( tsf.parseFilter( c, filter[ indx ], data ) || '' );
						data2.iFilter = '' + ( tsf.parseFilter( c, iFilter[ indx ], data ) || '' );
						query = ( '(' + ( tsf.parseFilter( c, data2.filter, data ) || '' ) + ')' )
							// replace wild cards since /(a*)/i will match anything
							.replace( tsfRegex.wild01, '\\S{1}' ).replace( tsfRegex.wild0More, '\\S*' );
						try {
							// use try/catch just in case RegExp is invalid
							regex = new RegExp( data.isMatch ? query : '^' + query + '$', c.widgetOptions.filter_ignoreCase ? 'i' : '' );
							// look for an exact match with the 'and' unless the 'filter-match' class is found
							result = ( regex.test( data2.exact ) || tsf.processTypes( c, data2, vars ) );
							if ( indx === 0 ) {
								filterMatched = result;
							} else {
								filterMatched = filterMatched && result;
							}
						} catch ( error ) {
							return null;
						}
					}
					// may be null from processing types
					return filterMatched || false;
				}
				return null;
			},
			// Look for regex
			regex: function( c, data ) {
				if ( tsfRegex.regex.test( data.filter ) ) {
					var matches,
						// cache regex per column for optimal speed
						regex = data.filter_regexCache[ data.index ] || tsfRegex.regex.exec( data.filter ),
						isRegex = regex instanceof RegExp;
					try {
						if ( !isRegex ) {
							// force case insensitive search if ignoreCase option set?
							// if ( c.ignoreCase && !regex[2] ) { regex[2] = 'i'; }
							data.filter_regexCache[ data.index ] = regex = new RegExp( regex[1], regex[2] );
						}
						matches = regex.test( data.exact );
					} catch ( error ) {
						matches = false;
					}
					return matches;
				}
				return null;
			},
			// Look for operators >, >=, < or <=
			operators: function( c, data ) {
				// ignore empty strings... because '' < 10 is true
				if ( tsfRegex.operTest.test( data.iFilter ) && data.iExact !== '' ) {
					var cachedValue, result, txt,
						table = c.table,
						parsed = data.parsed[ data.index ],
						query = ts.formatFloat( data.iFilter.replace( tsfRegex.operators, '' ), table ),
						parser = c.parsers[ data.index ] || {},
						savedSearch = query;
					// parse filter value in case we're comparing numbers ( dates )
					if ( parsed || parser.type === 'numeric' ) {
						txt = $.trim( '' + data.iFilter.replace( tsfRegex.operators, '' ) );
						result = tsf.parseFilter( c, txt, data, true );
						query = ( typeof result === 'number' && result !== '' && !isNaN( result ) ) ? result : query;
					}
					// iExact may be numeric - see issue #149;
					// check if cached is defined, because sometimes j goes out of range? ( numeric columns )
					if ( ( parsed || parser.type === 'numeric' ) && !isNaN( query ) &&
						typeof data.cache !== 'undefined' ) {
						cachedValue = data.cache;
					} else {
						txt = isNaN( data.iExact ) ? data.iExact.replace( ts.regex.nondigit, '' ) : data.iExact;
						cachedValue = ts.formatFloat( txt, table );
					}
					if ( tsfRegex.gtTest.test( data.iFilter ) ) {
						result = tsfRegex.gteTest.test( data.iFilter ) ? cachedValue >= query : cachedValue > query;
					} else if ( tsfRegex.ltTest.test( data.iFilter ) ) {
						result = tsfRegex.lteTest.test( data.iFilter ) ? cachedValue <= query : cachedValue < query;
					}
					// keep showing all rows if nothing follows the operator
					if ( !result && savedSearch === '' ) {
						result = true;
					}
					return result;
				}
				return null;
			},
			// Look for a not match
			notMatch: function( c, data ) {
				if ( tsfRegex.notTest.test( data.iFilter ) ) {
					var indx,
						txt = data.iFilter.replace( '!', '' ),
						filter = tsf.parseFilter( c, txt, data ) || '';
					if ( tsfRegex.exact.test( filter ) ) {
						// look for exact not matches - see #628
						filter = filter.replace( tsfRegex.exact, '' );
						return filter === '' ? true : $.trim( filter ) !== data.iExact;
					} else {
						indx = data.iExact.search( $.trim( filter ) );
						return filter === '' ? true :
							// return true if not found
							data.anyMatch ? indx < 0 :
							// return false if found
							!( c.widgetOptions.filter_startsWith ? indx === 0 : indx >= 0 );
					}
				}
				return null;
			},
			// Look for quotes or equals to get an exact match; ignore type since iExact could be numeric
			exact: function( c, data ) {
				/*jshint eqeqeq:false */
				if ( tsfRegex.exact.test( data.iFilter ) ) {
					var txt = data.iFilter.replace( tsfRegex.exact, '' ),
						filter = tsf.parseFilter( c, txt, data ) || '';
					// eslint-disable-next-line eqeqeq
					return data.anyMatch ? $.inArray( filter, data.rowArray ) >= 0 : filter == data.iExact;
				}
				return null;
			},
			// Look for a range ( using ' to ' or ' - ' ) - see issue #166; thanks matzhu!
			range : function( c, data ) {
				if ( tsfRegex.toTest.test( data.iFilter ) ) {
					var result, tmp, range1, range2,
						table = c.table,
						index = data.index,
						parsed = data.parsed[index],
						// make sure the dash is for a range and not indicating a negative number
						query = data.iFilter.split( tsfRegex.toSplit );

					tmp = query[0].replace( ts.regex.nondigit, '' ) || '';
					range1 = ts.formatFloat( tsf.parseFilter( c, tmp, data ), table );
					tmp = query[1].replace( ts.regex.nondigit, '' ) || '';
					range2 = ts.formatFloat( tsf.parseFilter( c, tmp, data ), table );
					// parse filter value in case we're comparing numbers ( dates )
					if ( parsed || c.parsers[ index ].type === 'numeric' ) {
						result = c.parsers[ index ].format( '' + query[0], table, c.$headers.eq( index ), index );
						range1 = ( result !== '' && !isNaN( result ) ) ? result : range1;
						result = c.parsers[ index ].format( '' + query[1], table, c.$headers.eq( index ), index );
						range2 = ( result !== '' && !isNaN( result ) ) ? result : range2;
					}
					if ( ( parsed || c.parsers[ index ].type === 'numeric' ) && !isNaN( range1 ) && !isNaN( range2 ) ) {
						result = data.cache;
					} else {
						tmp = isNaN( data.iExact ) ? data.iExact.replace( ts.regex.nondigit, '' ) : data.iExact;
						result = ts.formatFloat( tmp, table );
					}
					if ( range1 > range2 ) {
						tmp = range1; range1 = range2; range2 = tmp; // swap
					}
					return ( result >= range1 && result <= range2 ) || ( range1 === '' || range2 === '' );
				}
				return null;
			},
			// Look for wild card: ? = single, * = multiple, or | = logical OR
			wild : function( c, data ) {
				if ( tsfRegex.wildOrTest.test( data.iFilter ) ) {
					var query = '' + ( tsf.parseFilter( c, data.iFilter, data ) || '' );
					// look for an exact match with the 'or' unless the 'filter-match' class is found
					if ( !tsfRegex.wildTest.test( query ) && data.nestedFilters ) {
						query = data.isMatch ? query : '^(' + query + ')$';
					}
					// parsing the filter may not work properly when using wildcards =/
					try {
						return new RegExp(
							query.replace( tsfRegex.wild01, '\\S{1}' ).replace( tsfRegex.wild0More, '\\S*' ),
							c.widgetOptions.filter_ignoreCase ? 'i' : ''
						)
						.test( data.exact );
					} catch ( error ) {
						return null;
					}
				}
				return null;
			},
			// fuzzy text search; modified from https://github.com/mattyork/fuzzy ( MIT license )
			fuzzy: function( c, data ) {
				if ( tsfRegex.fuzzyTest.test( data.iFilter ) ) {
					var indx,
						patternIndx = 0,
						len = data.iExact.length,
						txt = data.iFilter.slice( 1 ),
						pattern = tsf.parseFilter( c, txt, data ) || '';
					for ( indx = 0; indx < len; indx++ ) {
						if ( data.iExact[ indx ] === pattern[ patternIndx ] ) {
							patternIndx += 1;
						}
					}
					return patternIndx === pattern.length;
				}
				return null;
			}
		},
		init: function( table ) {
			// filter language options
			ts.language = $.extend( true, {}, {
				to  : 'to',
				or  : 'or',
				and : 'and'
			}, ts.language );

			var options, string, txt, $header, column, val, fxn, noSelect,
				c = table.config,
				wo = c.widgetOptions,
				processStr = function(prefix, str, suffix) {
					str = str.trim();
					// don't include prefix/suffix if str is empty
					return str === '' ? '' : (prefix || '') + str + (suffix || '');
				};
			c.$table.addClass( 'hasFilters' );
			c.lastSearch = [];

			// define timers so using clearTimeout won't cause an undefined error
			wo.filter_searchTimer = null;
			wo.filter_initTimer = null;
			wo.filter_formatterCount = 0;
			wo.filter_formatterInit = [];
			wo.filter_anyColumnSelector = '[data-column="all"],[data-column="any"]';
			wo.filter_multipleColumnSelector = '[data-column*="-"],[data-column*=","]';

			val = '\\{' + tsfRegex.query + '\\}';
			$.extend( tsfRegex, {
				child : new RegExp( c.cssChildRow ),
				filtered : new RegExp( wo.filter_filteredRow ),
				alreadyFiltered : new RegExp( '(\\s+(-' + processStr('|', ts.language.or) + processStr('|', ts.language.to) + ')\\s+)', 'i' ),
				toTest : new RegExp( '\\s+(-' + processStr('|', ts.language.to) + ')\\s+', 'i' ),
				toSplit : new RegExp( '(?:\\s+(?:-' + processStr('|', ts.language.to) + ')\\s+)', 'gi' ),
				andTest : new RegExp( '\\s+(' + processStr('', ts.language.and, '|') + '&&)\\s+', 'i' ),
				andSplit : new RegExp( '(?:\\s+(?:' + processStr('', ts.language.and, '|') + '&&)\\s+)', 'gi' ),
				orTest : new RegExp( '(\\|' + processStr('|\\s+', ts.language.or, '\\s+') + ')', 'i' ),
				orSplit : new RegExp( '(?:\\|' + processStr('|\\s+(?:', ts.language.or, ')\\s+') + ')', 'gi' ),
				iQuery : new RegExp( val, 'i' ),
				igQuery : new RegExp( val, 'ig' ),
				operTest : /^[<>]=?/,
				gtTest  : />/,
				gteTest : />=/,
				ltTest  : /</,
				lteTest : /<=/,
				notTest : /^\!/,
				wildOrTest : /[\?\*\|]/,
				wildTest : /\?\*/,
				fuzzyTest : /^~/,
				exactTest : /[=\"\|!]/
			});

			// don't build filter row if columnFilters is false or all columns are set to 'filter-false'
			// see issue #156
			val = c.$headers.filter( '.filter-false, .parser-false' ).length;
			if ( wo.filter_columnFilters !== false && val !== c.$headers.length ) {
				// build filter row
				tsf.buildRow( table, c, wo );
			}

			txt = 'addRows updateCell update updateRows updateComplete appendCache filterReset ' +
				'filterAndSortReset filterResetSaved filterEnd search '.split( ' ' ).join( c.namespace + 'filter ' );
			c.$table.bind( txt, function( event, filter ) {
				val = wo.filter_hideEmpty &&
					$.isEmptyObject( c.cache ) &&
					!( c.delayInit && event.type === 'appendCache' );
				// hide filter row using the 'filtered' class name
				c.$table.find( '.' + tscss.filterRow ).toggleClass( wo.filter_filteredRow, val ); // fixes #450
				if ( !/(search|filter)/.test( event.type ) ) {
					event.stopPropagation();
					tsf.buildDefault( table, true );
				}
				// Add filterAndSortReset - see #1361
				if ( event.type === 'filterReset' || event.type === 'filterAndSortReset' ) {
					c.$table.find( '.' + tscss.filter ).add( wo.filter_$externalFilters ).val( '' );
					if ( event.type === 'filterAndSortReset' ) {
						ts.sortReset( this.config, function() {
							tsf.searching( table, [] );
						});
					} else {
						tsf.searching( table, [] );
					}
				} else if ( event.type === 'filterResetSaved' ) {
					ts.storage( table, 'tablesorter-filters', '' );
				} else if ( event.type === 'filterEnd' ) {
					tsf.buildDefault( table, true );
				} else {
					// send false argument to force a new search; otherwise if the filter hasn't changed,
					// it will return
					filter = event.type === 'search' ? filter :
						event.type === 'updateComplete' ? c.$table.data( 'lastSearch' ) : '';
					if ( /(update|add)/.test( event.type ) && event.type !== 'updateComplete' ) {
						// force a new search since content has changed
						c.lastCombinedFilter = null;
						c.lastSearch = [];
						// update filterFormatters after update (& small delay) - Fixes #1237
						setTimeout(function() {
							c.$table.triggerHandler( 'filterFomatterUpdate' );
						}, 100);
					}
					// pass true ( skipFirst ) to prevent the tablesorter.setFilters function from skipping the first
					// input ensures all inputs are updated when a search is triggered on the table
					// $( 'table' ).trigger( 'search', [...] );
					tsf.searching( table, filter, true );
				}
				return false;
			});

			// reset button/link
			if ( wo.filter_reset ) {
				if ( wo.filter_reset instanceof $ ) {
					// reset contains a jQuery object, bind to it
					wo.filter_reset.click( function() {
						c.$table.triggerHandler( 'filterReset' );
					});
				} else if ( $( wo.filter_reset ).length ) {
					// reset is a jQuery selector, use event delegation
					$( document )
						.undelegate( wo.filter_reset, 'click' + c.namespace + 'filter' )
						.delegate( wo.filter_reset, 'click' + c.namespace + 'filter', function() {
							// trigger a reset event, so other functions ( filter_formatter ) know when to reset
							c.$table.triggerHandler( 'filterReset' );
						});
				}
			}
			if ( wo.filter_functions ) {
				for ( column = 0; column < c.columns; column++ ) {
					fxn = ts.getColumnData( table, wo.filter_functions, column );
					if ( fxn ) {
						// remove 'filter-select' from header otherwise the options added here are replaced with
						// all options
						$header = c.$headerIndexed[ column ].removeClass( 'filter-select' );
						// don't build select if 'filter-false' or 'parser-false' set
						noSelect = !( $header.hasClass( 'filter-false' ) || $header.hasClass( 'parser-false' ) );
						options = '';
						if ( fxn === true && noSelect ) {
							tsf.buildSelect( table, column );
						} else if ( typeof fxn === 'object' && noSelect ) {
							// add custom drop down list
							for ( string in fxn ) {
								if ( typeof string === 'string' ) {
									options += options === '' ?
										'<option value="">' +
											( $header.data( 'placeholder' ) ||
												$header.attr( 'data-placeholder' ) ||
												wo.filter_placeholder.select ||
												''
											) +
										'</option>' : '';
									val = string;
									txt = string;
									if ( string.indexOf( wo.filter_selectSourceSeparator ) >= 0 ) {
										val = string.split( wo.filter_selectSourceSeparator );
										txt = val[1];
										val = val[0];
									}
									options += '<option ' +
										( txt === val ? '' : 'data-function-name="' + string + '" ' ) +
										'value="' + val + '">' + txt + '</option>';
								}
							}
							c.$table
								.find( 'thead' )
								.find( 'select.' + tscss.filter + '[data-column="' + column + '"]' )
								.append( options );
							txt = wo.filter_selectSource;
							fxn = typeof txt === 'function' ? true : ts.getColumnData( table, txt, column );
							if ( fxn ) {
								// updating so the extra options are appended
								tsf.buildSelect( c.table, column, '', true, $header.hasClass( wo.filter_onlyAvail ) );
							}
						}
					}
				}
			}
			// not really updating, but if the column has both the 'filter-select' class &
			// filter_functions set to true, it would append the same options twice.
			tsf.buildDefault( table, true );

			tsf.bindSearch( table, c.$table.find( '.' + tscss.filter ), true );
			if ( wo.filter_external ) {
				tsf.bindSearch( table, wo.filter_external );
			}

			if ( wo.filter_hideFilters ) {
				tsf.hideFilters( c );
			}

			// show processing icon
			if ( c.showProcessing ) {
				txt = 'filterStart filterEnd '.split( ' ' ).join( c.namespace + 'filter-sp ' );
				c.$table
					.unbind( txt.replace( ts.regex.spaces, ' ' ) )
					.bind( txt, function( event, columns ) {
					// only add processing to certain columns to all columns
					$header = ( columns ) ?
						c.$table
							.find( '.' + tscss.header )
							.filter( '[data-column]' )
							.filter( function() {
								return columns[ $( this ).data( 'column' ) ] !== '';
							}) : '';
					ts.isProcessing( table, event.type === 'filterStart', columns ? $header : '' );
				});
			}

			// set filtered rows count ( intially unfiltered )
			c.filteredRows = c.totalRows;

			// add default values
			txt = 'tablesorter-initialized pagerBeforeInitialized '.split( ' ' ).join( c.namespace + 'filter ' );
			c.$table
			.unbind( txt.replace( ts.regex.spaces, ' ' ) )
			.bind( txt, function() {
				tsf.completeInit( this );
			});
			// if filter widget is added after pager has initialized; then set filter init flag
			if ( c.pager && c.pager.initialized && !wo.filter_initialized ) {
				c.$table.triggerHandler( 'filterFomatterUpdate' );
				setTimeout( function() {
					tsf.filterInitComplete( c );
				}, 100 );
			} else if ( !wo.filter_initialized ) {
				tsf.completeInit( table );
			}
		},
		completeInit: function( table ) {
			// redefine 'c' & 'wo' so they update properly inside this callback
			var c = table.config,
				wo = c.widgetOptions,
				filters = tsf.setDefaults( table, c, wo ) || [];
			if ( filters.length ) {
				// prevent delayInit from triggering a cache build if filters are empty
				if ( !( c.delayInit && filters.join( '' ) === '' ) ) {
					ts.setFilters( table, filters, true );
				}
			}
			c.$table.triggerHandler( 'filterFomatterUpdate' );
			// trigger init after setTimeout to prevent multiple filterStart/End/Init triggers
			setTimeout( function() {
				if ( !wo.filter_initialized ) {
					tsf.filterInitComplete( c );
				}
			}, 100 );
		},

		// $cell parameter, but not the config, is passed to the filter_formatters,
		// so we have to work with it instead
		formatterUpdated: function( $cell, column ) {
			// prevent error if $cell is undefined - see #1056
			var $table = $cell && $cell.closest( 'table' );
			var config = $table.length && $table[0].config,
				wo = config && config.widgetOptions;
			if ( wo && !wo.filter_initialized ) {
				// add updates by column since this function
				// may be called numerous times before initialization
				wo.filter_formatterInit[ column ] = 1;
			}
		},
		filterInitComplete: function( c ) {
			var indx, len,
				wo = c.widgetOptions,
				count = 0,
				completed = function() {
					wo.filter_initialized = true;
					// update lastSearch - it gets cleared often
					c.lastSearch = c.$table.data( 'lastSearch' );
					c.$table.triggerHandler( 'filterInit', c );
					tsf.findRows( c.table, c.lastSearch || [] );
					if (ts.debug(c, 'filter')) {
						console.log('Filter >> Widget initialized');
					}
				};
			if ( $.isEmptyObject( wo.filter_formatter ) ) {
				completed();
			} else {
				len = wo.filter_formatterInit.length;
				for ( indx = 0; indx < len; indx++ ) {
					if ( wo.filter_formatterInit[ indx ] === 1 ) {
						count++;
					}
				}
				clearTimeout( wo.filter_initTimer );
				if ( !wo.filter_initialized && count === wo.filter_formatterCount ) {
					// filter widget initialized
					completed();
				} else if ( !wo.filter_initialized ) {
					// fall back in case a filter_formatter doesn't call
					// $.tablesorter.filter.formatterUpdated( $cell, column ), and the count is off
					wo.filter_initTimer = setTimeout( function() {
						completed();
					}, 500 );
				}
			}
		},
		// encode or decode filters for storage; see #1026
		processFilters: function( filters, encode ) {
			var indx,
				// fixes #1237; previously returning an encoded "filters" value
				result = [],
				mode = encode ? encodeURIComponent : decodeURIComponent,
				len = filters.length;
			for ( indx = 0; indx < len; indx++ ) {
				if ( filters[ indx ] ) {
					result[ indx ] = mode( filters[ indx ] );
				}
			}
			return result;
		},
		setDefaults: function( table, c, wo ) {
			var isArray, saved, indx, col, $filters,
				// get current ( default ) filters
				filters = ts.getFilters( table ) || [];
			if ( wo.filter_saveFilters && ts.storage ) {
				saved = ts.storage( table, 'tablesorter-filters' ) || [];
				isArray = $.isArray( saved );
				// make sure we're not just getting an empty array
				if ( !( isArray && saved.join( '' ) === '' || !isArray ) ) {
					filters = tsf.processFilters( saved );
				}
			}
			// if no filters saved, then check default settings
			if ( filters.join( '' ) === '' ) {
				// allow adding default setting to external filters
				$filters = c.$headers.add( wo.filter_$externalFilters )
					.filter( '[' + wo.filter_defaultAttrib + ']' );
				for ( indx = 0; indx <= c.columns; indx++ ) {
					// include data-column='all' external filters
					col = indx === c.columns ? 'all' : indx;
					filters[ indx ] = $filters
						.filter( '[data-column="' + col + '"]' )
						.attr( wo.filter_defaultAttrib ) || filters[indx] || '';
				}
			}
			c.$table.data( 'lastSearch', filters );
			return filters;
		},
		parseFilter: function( c, filter, data, parsed ) {
			return parsed || data.parsed[ data.index ] ?
				c.parsers[ data.index ].format( filter, c.table, [], data.index ) :
				filter;
		},
		buildRow: function( table, c, wo ) {
			var $filter, col, column, $header, makeSelect, disabled, name, ffxn, tmp,
				// c.columns defined in computeThIndexes()
				cellFilter = wo.filter_cellFilter,
				columns = c.columns,
				arry = $.isArray( cellFilter ),
				buildFilter = '<tr role="search" class="' + tscss.filterRow + ' ' + c.cssIgnoreRow + '">';
			for ( column = 0; column < columns; column++ ) {
				if ( c.$headerIndexed[ column ].length ) {
					// account for entire column set with colspan. See #1047
					tmp = c.$headerIndexed[ column ] && c.$headerIndexed[ column ][0].colSpan || 0;
					if ( tmp > 1 ) {
						buildFilter += '<td data-column="' + column + '-' + ( column + tmp - 1 ) + '" colspan="' + tmp + '"';
					} else {
						buildFilter += '<td data-column="' + column + '"';
					}
					if ( arry ) {
						buildFilter += ( cellFilter[ column ] ? ' class="' + cellFilter[ column ] + '"' : '' );
					} else {
						buildFilter += ( cellFilter !== '' ? ' class="' + cellFilter + '"' : '' );
					}
					buildFilter += '></td>';
				}
			}
			c.$filters = $( buildFilter += '</tr>' )
				.appendTo( c.$table.children( 'thead' ).eq( 0 ) )
				.children( 'td' );
			// build each filter input
			for ( column = 0; column < columns; column++ ) {
				disabled = false;
				// assuming last cell of a column is the main column
				$header = c.$headerIndexed[ column ];
				if ( $header && $header.length ) {
					// $filter = c.$filters.filter( '[data-column="' + column + '"]' );
					$filter = tsf.getColumnElm( c, c.$filters, column );
					ffxn = ts.getColumnData( table, wo.filter_functions, column );
					makeSelect = ( wo.filter_functions && ffxn && typeof ffxn !== 'function' ) ||
						$header.hasClass( 'filter-select' );
					// get data from jQuery data, metadata, headers option or header class name
					col = ts.getColumnData( table, c.headers, column );
					disabled = ts.getData( $header[0], col, 'filter' ) === 'false' ||
						ts.getData( $header[0], col, 'parser' ) === 'false';

					if ( makeSelect ) {
						buildFilter = $( '<select>' ).appendTo( $filter );
					} else {
						ffxn = ts.getColumnData( table, wo.filter_formatter, column );
						if ( ffxn ) {
							wo.filter_formatterCount++;
							buildFilter = ffxn( $filter, column );
							// no element returned, so lets go find it
							if ( buildFilter && buildFilter.length === 0 ) {
								buildFilter = $filter.children( 'input' );
							}
							// element not in DOM, so lets attach it
							if ( buildFilter && ( buildFilter.parent().length === 0 ||
								( buildFilter.parent().length && buildFilter.parent()[0] !== $filter[0] ) ) ) {
								$filter.append( buildFilter );
							}
						} else {
							buildFilter = $( '<input type="search">' ).appendTo( $filter );
						}
						if ( buildFilter ) {
							tmp = $header.data( 'placeholder' ) ||
								$header.attr( 'data-placeholder' ) ||
								wo.filter_placeholder.search || '';
							buildFilter.attr( 'placeholder', tmp );
						}
					}
					if ( buildFilter ) {
						// add filter class name
						name = ( $.isArray( wo.filter_cssFilter ) ?
							( typeof wo.filter_cssFilter[column] !== 'undefined' ? wo.filter_cssFilter[column] || '' : '' ) :
							wo.filter_cssFilter ) || '';
						// copy data-column from table cell (it will include colspan)
						buildFilter.addClass( tscss.filter + ' ' + name );
						name = wo.filter_filterLabel;
						tmp = name.match(/{{([^}]+?)}}/g);
						if (!tmp) {
							tmp = [ '{{label}}' ];
						}
						$.each(tmp, function(indx, attr) {
							var regex = new RegExp(attr, 'g'),
								data = $header.attr('data-' + attr.replace(/{{|}}/g, '')),
								text = typeof data === 'undefined' ? $header.text() : data;
							name = name.replace( regex, $.trim( text ) );
						});
						buildFilter.attr({
							'data-column': $filter.attr( 'data-column' ),
							'aria-label': name
						});
						if ( disabled ) {
							buildFilter.attr( 'placeholder', '' ).addClass( tscss.filterDisabled )[0].disabled = true;
						}
					}
				}
			}
		},
		bindSearch: function( table, $el, internal ) {
			table = $( table )[0];
			$el = $( $el ); // allow passing a selector string
			if ( !$el.length ) { return; }
			var tmp,
				c = table.config,
				wo = c.widgetOptions,
				namespace = c.namespace + 'filter',
				$ext = wo.filter_$externalFilters;
			if ( internal !== true ) {
				// save anyMatch element
				tmp = wo.filter_anyColumnSelector + ',' + wo.filter_multipleColumnSelector;
				wo.filter_$anyMatch = $el.filter( tmp );
				if ( $ext && $ext.length ) {
					wo.filter_$externalFilters = wo.filter_$externalFilters.add( $el );
				} else {
					wo.filter_$externalFilters = $el;
				}
				// update values ( external filters added after table initialization )
				ts.setFilters( table, c.$table.data( 'lastSearch' ) || [], internal === false );
			}
			// unbind events
			tmp = ( 'keypress keyup keydown search change input '.split( ' ' ).join( namespace + ' ' ) );
			$el
			// use data attribute instead of jQuery data since the head is cloned without including
			// the data/binding
			.attr( 'data-lastSearchTime', new Date().getTime() )
			.unbind( tmp.replace( ts.regex.spaces, ' ' ) )
			.bind( 'keydown' + namespace, function( event ) {
				if ( event.which === tskeyCodes.escape && !table.config.widgetOptions.filter_resetOnEsc ) {
					// prevent keypress event
					return false;
				}
			})
			.bind( 'keyup' + namespace, function( event ) {
				wo = table.config.widgetOptions; // make sure "wo" isn't cached
				var column = parseInt( $( this ).attr( 'data-column' ), 10 ),
					liveSearch = typeof wo.filter_liveSearch === 'boolean' ? wo.filter_liveSearch :
						ts.getColumnData( table, wo.filter_liveSearch, column );
				if ( typeof liveSearch === 'undefined' ) {
					liveSearch = wo.filter_liveSearch.fallback || false;
				}
				$( this ).attr( 'data-lastSearchTime', new Date().getTime() );
				// emulate what webkit does.... escape clears the filter
				if ( event.which === tskeyCodes.escape ) {
					// make sure to restore the last value on escape
					this.value = wo.filter_resetOnEsc ? '' : c.lastSearch[column];
					// don't return if the search value is empty ( all rows need to be revealed )
				} else if ( this.value !== '' && (
					// liveSearch can contain a min value length; ignore arrow and meta keys, but allow backspace
					( typeof liveSearch === 'number' && this.value.length < liveSearch ) ||
					// let return & backspace continue on, but ignore arrows & non-valid characters
					( event.which !== tskeyCodes.enter && event.which !== tskeyCodes.backSpace &&
						( event.which < tskeyCodes.space || ( event.which >= tskeyCodes.left && event.which <= tskeyCodes.down ) ) ) ) ) {
					return;
					// live search
				} else if ( liveSearch === false ) {
					if ( this.value !== '' && event.which !== tskeyCodes.enter ) {
						return;
					}
				}
				// change event = no delay; last true flag tells getFilters to skip newest timed input
				tsf.searching( table, true, true, column );
			})
			// include change for select - fixes #473
			.bind( 'search change keypress input blur '.split( ' ' ).join( namespace + ' ' ), function( event ) {
				// don't get cached data, in case data-column changes dynamically
				var column = parseInt( $( this ).attr( 'data-column' ), 10 ),
					eventType = event.type,
					liveSearch = typeof wo.filter_liveSearch === 'boolean' ?
						wo.filter_liveSearch :
						ts.getColumnData( table, wo.filter_liveSearch, column );
				if ( table.config.widgetOptions.filter_initialized &&
					// immediate search if user presses enter
					( event.which === tskeyCodes.enter ||
						// immediate search if a "search" or "blur" is triggered on the input
						( eventType === 'search' || eventType === 'blur' ) ||
						// change & input events must be ignored if liveSearch !== true
						( eventType === 'change' || eventType === 'input' ) &&
						// prevent search if liveSearch is a number
						( liveSearch === true || liveSearch !== true && event.target.nodeName !== 'INPUT' ) &&
						// don't allow 'change' or 'input' event to process if the input value
						// is the same - fixes #685
						this.value !== c.lastSearch[column]
					)
				) {
					event.preventDefault();
					// init search with no delay
					$( this ).attr( 'data-lastSearchTime', new Date().getTime() );
					tsf.searching( table, eventType !== 'keypress' || event.which === tskeyCodes.enter, true, column );
				}
			});
		},
		searching: function( table, filter, skipFirst, column ) {
			var liveSearch,
				wo = table.config.widgetOptions;
			if (typeof column === 'undefined') {
				// no delay
				liveSearch = false;
			} else {
				liveSearch = typeof wo.filter_liveSearch === 'boolean' ?
					wo.filter_liveSearch :
					// get column setting, or set to fallback value, or default to false
					ts.getColumnData( table, wo.filter_liveSearch, column );
				if ( typeof liveSearch === 'undefined' ) {
					liveSearch = wo.filter_liveSearch.fallback || false;
				}
			}
			clearTimeout( wo.filter_searchTimer );
			if ( typeof filter === 'undefined' || filter === true ) {
				// delay filtering
				wo.filter_searchTimer = setTimeout( function() {
					tsf.checkFilters( table, filter, skipFirst );
				}, liveSearch ? wo.filter_searchDelay : 10 );
			} else {
				// skip delay
				tsf.checkFilters( table, filter, skipFirst );
			}
		},
		equalFilters: function (c, filter1, filter2) {
			var indx,
				f1 = [],
				f2 = [],
				len = c.columns + 1; // add one to include anyMatch filter
			filter1 = $.isArray(filter1) ? filter1 : [];
			filter2 = $.isArray(filter2) ? filter2 : [];
			for (indx = 0; indx < len; indx++) {
				f1[indx] = filter1[indx] || '';
				f2[indx] = filter2[indx] || '';
			}
			return f1.join(',') === f2.join(',');
		},
		checkFilters: function( table, filter, skipFirst ) {
			var c = table.config,
				wo = c.widgetOptions,
				filterArray = $.isArray( filter ),
				filters = ( filterArray ) ? filter : ts.getFilters( table, true ),
				currentFilters = filters || []; // current filter values
			// prevent errors if delay init is set
			if ( $.isEmptyObject( c.cache ) ) {
				// update cache if delayInit set & pager has initialized ( after user initiates a search )
				if ( c.delayInit && ( !c.pager || c.pager && c.pager.initialized ) ) {
					ts.updateCache( c, function() {
						tsf.checkFilters( table, false, skipFirst );
					});
				}
				return;
			}
			// add filter array back into inputs
			if ( filterArray ) {
				ts.setFilters( table, filters, false, skipFirst !== true );
				if ( !wo.filter_initialized ) {
					c.lastSearch = [];
					c.lastCombinedFilter = '';
				}
			}
			if ( wo.filter_hideFilters ) {
				// show/hide filter row as needed
				c.$table
					.find( '.' + tscss.filterRow )
					.triggerHandler( tsf.hideFiltersCheck( c ) ? 'mouseleave' : 'mouseenter' );
			}
			// return if the last search is the same; but filter === false when updating the search
			// see example-widget-filter.html filter toggle buttons
			if ( tsf.equalFilters(c, c.lastSearch, currentFilters) ) {
				if ( filter !== false ) {
					return;
				} else {
					// force filter refresh
					c.lastCombinedFilter = '';
					c.lastSearch = [];
				}
			}
			// define filter inside it is false
			filters = filters || [];
			// convert filters to strings - see #1070
			filters = Array.prototype.map ?
				filters.map( String ) :
				// for IE8 & older browsers - maybe not the best method
				filters.join( '\ufffd' ).split( '\ufffd' );

			if ( wo.filter_initialized ) {
				c.$table.triggerHandler( 'filterStart', [ filters ] );
			}
			if ( c.showProcessing ) {
				// give it time for the processing icon to kick in
				setTimeout( function() {
					tsf.findRows( table, filters, currentFilters );
					return false;
				}, 30 );
			} else {
				tsf.findRows( table, filters, currentFilters );
				return false;
			}
		},
		hideFiltersCheck: function( c ) {
			if (typeof c.widgetOptions.filter_hideFilters === 'function') {
				var val = c.widgetOptions.filter_hideFilters( c );
				if (typeof val === 'boolean') {
					return val;
				}
			}
			return ts.getFilters( c.$table ).join( '' ) === '';
		},
		hideFilters: function( c, $table ) {
			var timer;
			( $table || c.$table )
				.find( '.' + tscss.filterRow )
				.addClass( tscss.filterRowHide )
				.bind( 'mouseenter mouseleave', function( e ) {
					// save event object - http://bugs.jquery.com/ticket/12140
					var event = e,
						$row = $( this );
					clearTimeout( timer );
					timer = setTimeout( function() {
						if ( /enter|over/.test( event.type ) ) {
							$row.removeClass( tscss.filterRowHide );
						} else {
							// don't hide if input has focus
							// $( ':focus' ) needs jQuery 1.6+
							if ( $( document.activeElement ).closest( 'tr' )[0] !== $row[0] ) {
								// don't hide row if any filter has a value
								$row.toggleClass( tscss.filterRowHide, tsf.hideFiltersCheck( c ) );
							}
						}
					}, 200 );
				})
				.find( 'input, select' ).bind( 'focus blur', function( e ) {
					var event = e,
						$row = $( this ).closest( 'tr' );
					clearTimeout( timer );
					timer = setTimeout( function() {
						clearTimeout( timer );
						// don't hide row if any filter has a value
						$row.toggleClass( tscss.filterRowHide, tsf.hideFiltersCheck( c ) && event.type !== 'focus' );
					}, 200 );
				});
		},
		defaultFilter: function( filter, mask ) {
			if ( filter === '' ) { return filter; }
			var regex = tsfRegex.iQuery,
				maskLen = mask.match( tsfRegex.igQuery ).length,
				query = maskLen > 1 ? $.trim( filter ).split( /\s/ ) : [ $.trim( filter ) ],
				len = query.length - 1,
				indx = 0,
				val = mask;
			if ( len < 1 && maskLen > 1 ) {
				// only one 'word' in query but mask has >1 slots
				query[1] = query[0];
			}
			// replace all {query} with query words...
			// if query = 'Bob', then convert mask from '!{query}' to '!Bob'
			// if query = 'Bob Joe Frank', then convert mask '{q} OR {q}' to 'Bob OR Joe OR Frank'
			while ( regex.test( val ) ) {
				val = val.replace( regex, query[indx++] || '' );
				if ( regex.test( val ) && indx < len && ( query[indx] || '' ) !== '' ) {
					val = mask.replace( regex, val );
				}
			}
			return val;
		},
		getLatestSearch: function( $input ) {
			if ( $input ) {
				return $input.sort( function( a, b ) {
					return $( b ).attr( 'data-lastSearchTime' ) - $( a ).attr( 'data-lastSearchTime' );
				});
			}
			return $input || $();
		},
		findRange: function( c, val, ignoreRanges ) {
			// look for multiple columns '1-3,4-6,8' in data-column
			var temp, ranges, range, start, end, singles, i, indx, len,
				columns = [];
			if ( /^[0-9]+$/.test( val ) ) {
				// always return an array
				return [ parseInt( val, 10 ) ];
			}
			// process column range
			if ( !ignoreRanges && /-/.test( val ) ) {
				ranges = val.match( /(\d+)\s*-\s*(\d+)/g );
				len = ranges ? ranges.length : 0;
				for ( indx = 0; indx < len; indx++ ) {
					range = ranges[indx].split( /\s*-\s*/ );
					start = parseInt( range[0], 10 ) || 0;
					end = parseInt( range[1], 10 ) || ( c.columns - 1 );
					if ( start > end ) {
						temp = start; start = end; end = temp; // swap
					}
					if ( end >= c.columns ) {
						end = c.columns - 1;
					}
					for ( ; start <= end; start++ ) {
						columns[ columns.length ] = start;
					}
					// remove processed range from val
					val = val.replace( ranges[ indx ], '' );
				}
			}
			// process single columns
			if ( !ignoreRanges && /,/.test( val ) ) {
				singles = val.split( /\s*,\s*/ );
				len = singles.length;
				for ( i = 0; i < len; i++ ) {
					if ( singles[ i ] !== '' ) {
						indx = parseInt( singles[ i ], 10 );
						if ( indx < c.columns ) {
							columns[ columns.length ] = indx;
						}
					}
				}
			}
			// return all columns
			if ( !columns.length ) {
				for ( indx = 0; indx < c.columns; indx++ ) {
					columns[ columns.length ] = indx;
				}
			}
			return columns;
		},
		getColumnElm: function( c, $elements, column ) {
			// data-column may contain multiple columns '1-3,5-6,8'
			// replaces: c.$filters.filter( '[data-column="' + column + '"]' );
			return $elements.filter( function() {
				var cols = tsf.findRange( c, $( this ).attr( 'data-column' ) );
				return $.inArray( column, cols ) > -1;
			});
		},
		multipleColumns: function( c, $input ) {
			// look for multiple columns '1-3,4-6,8' in data-column
			var wo = c.widgetOptions,
				// only target 'all' column inputs on initialization
				// & don't target 'all' column inputs if they don't exist
				targets = wo.filter_initialized || !$input.filter( wo.filter_anyColumnSelector ).length,
				val = $.trim( tsf.getLatestSearch( $input ).attr( 'data-column' ) || '' );
			return tsf.findRange( c, val, !targets );
		},
		processTypes: function( c, data, vars ) {
			var ffxn,
				filterMatched = null,
				matches = null;
			for ( ffxn in tsf.types ) {
				if ( $.inArray( ffxn, vars.excludeMatch ) < 0 && matches === null ) {
					matches = tsf.types[ffxn]( c, data, vars );
					if ( matches !== null ) {
						data.matchedOn = ffxn;
						filterMatched = matches;
					}
				}
			}
			return filterMatched;
		},
		matchType: function( c, columnIndex ) {
			var isMatch,
				wo = c.widgetOptions,
				$el = c.$headerIndexed[ columnIndex ];
			// filter-exact > filter-match > filter_matchType for type
			if ( $el.hasClass( 'filter-exact' ) ) {
				isMatch = false;
			} else if ( $el.hasClass( 'filter-match' ) ) {
				isMatch = true;
			} else {
				// filter-select is not applied when filter_functions are used, so look for a select
				if ( wo.filter_columnFilters ) {
					$el = c.$filters
						.find( '.' + tscss.filter )
						.add( wo.filter_$externalFilters )
						.filter( '[data-column="' + columnIndex + '"]' );
				} else if ( wo.filter_$externalFilters ) {
					$el = wo.filter_$externalFilters.filter( '[data-column="' + columnIndex + '"]' );
				}
				isMatch = $el.length ?
					c.widgetOptions.filter_matchType[ ( $el[ 0 ].nodeName || '' ).toLowerCase() ] === 'match' :
					// default to exact, if no inputs found
					false;
			}
			return isMatch;
		},
		processRow: function( c, data, vars ) {
			var result, filterMatched,
				fxn, ffxn, txt,
				wo = c.widgetOptions,
				showRow = true,
				hasAnyMatchInput = wo.filter_$anyMatch && wo.filter_$anyMatch.length,

				// if wo.filter_$anyMatch data-column attribute is changed dynamically
				// we don't want to do an "anyMatch" search on one column using data
				// for the entire row - see #998
				columnIndex = wo.filter_$anyMatch && wo.filter_$anyMatch.length ?
					// look for multiple columns '1-3,4-6,8'
					tsf.multipleColumns( c, wo.filter_$anyMatch ) :
					[];
			data.$cells = data.$row.children();
			data.matchedOn = null;
			if ( data.anyMatchFlag && columnIndex.length > 1 || ( data.anyMatchFilter && !hasAnyMatchInput ) ) {
				data.anyMatch = true;
				data.isMatch = true;
				data.rowArray = data.$cells.map( function( i ) {
					if ( $.inArray( i, columnIndex ) > -1 || ( data.anyMatchFilter && !hasAnyMatchInput ) ) {
						if ( data.parsed[ i ] ) {
							txt = data.cacheArray[ i ];
						} else {
							txt = data.rawArray[ i ];
							txt = $.trim( wo.filter_ignoreCase ? txt.toLowerCase() : txt );
							if ( c.sortLocaleCompare ) {
								txt = ts.replaceAccents( txt );
							}
						}
						return txt;
					}
				}).get();
				data.filter = data.anyMatchFilter;
				data.iFilter = data.iAnyMatchFilter;
				data.exact = data.rowArray.join( ' ' );
				data.iExact = wo.filter_ignoreCase ? data.exact.toLowerCase() : data.exact;
				data.cache = data.cacheArray.slice( 0, -1 ).join( ' ' );
				vars.excludeMatch = vars.noAnyMatch;
				filterMatched = tsf.processTypes( c, data, vars );
				if ( filterMatched !== null ) {
					showRow = filterMatched;
				} else {
					if ( wo.filter_startsWith ) {
						showRow = false;
						// data.rowArray may not contain all columns
						columnIndex = Math.min( c.columns, data.rowArray.length );
						while ( !showRow && columnIndex > 0 ) {
							columnIndex--;
							showRow = showRow || data.rowArray[ columnIndex ].indexOf( data.iFilter ) === 0;
						}
					} else {
						showRow = ( data.iExact + data.childRowText ).indexOf( data.iFilter ) >= 0;
					}
				}
				data.anyMatch = false;
				// no other filters to process
				if ( data.filters.join( '' ) === data.filter ) {
					return showRow;
				}
			}

			for ( columnIndex = 0; columnIndex < c.columns; columnIndex++ ) {
				data.filter = data.filters[ columnIndex ];
				data.index = columnIndex;

				// filter types to exclude, per column
				vars.excludeMatch = vars.excludeFilter[ columnIndex ];

				// ignore if filter is empty or disabled
				if ( data.filter ) {
					data.cache = data.cacheArray[ columnIndex ];
					result = data.parsed[ columnIndex ] ? data.cache : data.rawArray[ columnIndex ] || '';
					data.exact = c.sortLocaleCompare ? ts.replaceAccents( result ) : result; // issue #405
					data.iExact = !tsfRegex.type.test( typeof data.exact ) && wo.filter_ignoreCase ?
						data.exact.toLowerCase() : data.exact;
					data.isMatch = tsf.matchType( c, columnIndex );

					result = showRow; // if showRow is true, show that row

					// in case select filter option has a different value vs text 'a - z|A through Z'
					ffxn = wo.filter_columnFilters ?
						c.$filters.add( wo.filter_$externalFilters )
							.filter( '[data-column="' + columnIndex + '"]' )
							.find( 'select option:selected' )
							.attr( 'data-function-name' ) || '' : '';
					// replace accents - see #357
					if ( c.sortLocaleCompare ) {
						data.filter = ts.replaceAccents( data.filter );
					}

					// replace column specific default filters - see #1088
					if ( wo.filter_defaultFilter && tsfRegex.iQuery.test( vars.defaultColFilter[ columnIndex ] ) ) {
						data.filter = tsf.defaultFilter( data.filter, vars.defaultColFilter[ columnIndex ] );
					}

					// data.iFilter = case insensitive ( if wo.filter_ignoreCase is true ),
					// data.filter = case sensitive
					data.iFilter = wo.filter_ignoreCase ? ( data.filter || '' ).toLowerCase() : data.filter;
					fxn = vars.functions[ columnIndex ];
					filterMatched = null;
					if ( fxn ) {
						if ( typeof fxn === 'function' ) {
							// filter callback( exact cell content, parser normalized content,
							// filter input value, column index, jQuery row object )
							filterMatched = fxn( data.exact, data.cache, data.filter, columnIndex, data.$row, c, data );
						} else if ( typeof fxn[ ffxn || data.filter ] === 'function' ) {
							// selector option function
							txt = ffxn || data.filter;
							filterMatched =
								fxn[ txt ]( data.exact, data.cache, data.filter, columnIndex, data.$row, c, data );
						}
					}
					if ( filterMatched === null ) {
						// cycle through the different filters
						// filters return a boolean or null if nothing matches
						filterMatched = tsf.processTypes( c, data, vars );
						// select with exact match; ignore "and" or "or" within the text; fixes #1486
						txt = fxn === true && (data.matchedOn === 'and' || data.matchedOn === 'or');
						if ( filterMatched !== null && !txt) {
							result = filterMatched;
						// Look for match, and add child row data for matching
						} else {
							// check fxn (filter-select in header) after filter types are checked
							// without this, the filter + jQuery UI selectmenu demo was breaking
							if ( fxn === true ) {
								// default selector uses exact match unless 'filter-match' class is found
								result = data.isMatch ?
									// data.iExact may be a number
									( '' + data.iExact ).search( data.iFilter ) >= 0 :
									data.filter === data.exact;
							} else {
								txt = ( data.iExact + data.childRowText ).indexOf( tsf.parseFilter( c, data.iFilter, data ) );
								result = ( ( !wo.filter_startsWith && txt >= 0 ) || ( wo.filter_startsWith && txt === 0 ) );
							}
						}
					} else {
						result = filterMatched;
					}
					showRow = ( result ) ? showRow : false;
				}
			}
			return showRow;
		},
		findRows: function( table, filters, currentFilters ) {
			if (
				tsf.equalFilters(table.config, table.config.lastSearch, currentFilters) ||
				!table.config.widgetOptions.filter_initialized
			) {
				return;
			}
			var len, norm_rows, rowData, $rows, $row, rowIndex, tbodyIndex, $tbody, columnIndex,
				isChild, childRow, lastSearch, showRow, showParent, time, val, indx,
				notFiltered, searchFiltered, query, injected, res, id, txt,
				storedFilters = $.extend( [], filters ),
				c = table.config,
				wo = c.widgetOptions,
				debug = ts.debug(c, 'filter'),
				// data object passed to filters; anyMatch is a flag for the filters
				data = {
					anyMatch: false,
					filters: filters,
					// regex filter type cache
					filter_regexCache : []
				},
				vars = {
					// anyMatch really screws up with these types of filters
					noAnyMatch: [ 'range',  'operators' ],
					// cache filter variables that use ts.getColumnData in the main loop
					functions : [],
					excludeFilter : [],
					defaultColFilter : [],
					defaultAnyFilter : ts.getColumnData( table, wo.filter_defaultFilter, c.columns, true ) || ''
				};
			// parse columns after formatter, in case the class is added at that point
			data.parsed = [];
			for ( columnIndex = 0; columnIndex < c.columns; columnIndex++ ) {
				data.parsed[ columnIndex ] = wo.filter_useParsedData ||
					// parser has a "parsed" parameter
					( c.parsers && c.parsers[ columnIndex ] && c.parsers[ columnIndex ].parsed ||
					// getData may not return 'parsed' if other 'filter-' class names exist
					// ( e.g. <th class="filter-select filter-parsed"> )
					ts.getData && ts.getData( c.$headerIndexed[ columnIndex ],
						ts.getColumnData( table, c.headers, columnIndex ), 'filter' ) === 'parsed' ||
					c.$headerIndexed[ columnIndex ].hasClass( 'filter-parsed' ) );

				vars.functions[ columnIndex ] =
					ts.getColumnData( table, wo.filter_functions, columnIndex ) ||
					c.$headerIndexed[ columnIndex ].hasClass( 'filter-select' );
				vars.defaultColFilter[ columnIndex ] =
					ts.getColumnData( table, wo.filter_defaultFilter, columnIndex ) || '';
				vars.excludeFilter[ columnIndex ] =
					( ts.getColumnData( table, wo.filter_excludeFilter, columnIndex, true ) || '' ).split( /\s+/ );
			}

			if ( debug ) {
				console.log( 'Filter >> Starting filter widget search', filters );
				time = new Date();
			}
			// filtered rows count
			c.filteredRows = 0;
			c.totalRows = 0;
			currentFilters = ( storedFilters || [] );

			for ( tbodyIndex = 0; tbodyIndex < c.$tbodies.length; tbodyIndex++ ) {
				$tbody = ts.processTbody( table, c.$tbodies.eq( tbodyIndex ), true );
				// skip child rows & widget added ( removable ) rows - fixes #448 thanks to @hempel!
				// $rows = $tbody.children( 'tr' ).not( c.selectorRemove );
				columnIndex = c.columns;
				// convert stored rows into a jQuery object
				norm_rows = c.cache[ tbodyIndex ].normalized;
				$rows = $( $.map( norm_rows, function( el ) {
					return el[ columnIndex ].$row.get();
				}) );

				if ( currentFilters.join('') === '' || wo.filter_serversideFiltering ) {
					$rows
						.removeClass( wo.filter_filteredRow )
						.not( '.' + c.cssChildRow )
						.css( 'display', '' );
				} else {
					// filter out child rows
					$rows = $rows.not( '.' + c.cssChildRow );
					len = $rows.length;

					if ( ( wo.filter_$anyMatch && wo.filter_$anyMatch.length ) ||
						typeof filters[c.columns] !== 'undefined' ) {
						data.anyMatchFlag = true;
						data.anyMatchFilter = '' + (
							filters[ c.columns ] ||
							wo.filter_$anyMatch && tsf.getLatestSearch( wo.filter_$anyMatch ).val() ||
							''
						);
						if ( wo.filter_columnAnyMatch ) {
							// specific columns search
							query = data.anyMatchFilter.split( tsfRegex.andSplit );
							injected = false;
							for ( indx = 0; indx < query.length; indx++ ) {
								res = query[ indx ].split( ':' );
								if ( res.length > 1 ) {
									// make the column a one-based index ( non-developers start counting from one :P )
									if ( isNaN( res[0] ) ) {
										$.each( c.headerContent, function( i, txt ) {
											// multiple matches are possible
											if ( txt.toLowerCase().indexOf( res[0] ) > -1 ) {
												id = i;
												filters[ id ] = res[1];
											}
										});
									} else {
										id = parseInt( res[0], 10 ) - 1;
									}
									if ( id >= 0 && id < c.columns ) { // if id is an integer
										filters[ id ] = res[1];
										query.splice( indx, 1 );
										indx--;
										injected = true;
									}
								}
							}
							if ( injected ) {
								data.anyMatchFilter = query.join( ' && ' );
							}
						}
					}

					// optimize searching only through already filtered rows - see #313
					searchFiltered = wo.filter_searchFiltered;
					lastSearch = c.lastSearch || c.$table.data( 'lastSearch' ) || [];
					if ( searchFiltered ) {
						// cycle through all filters; include last ( columnIndex + 1 = match any column ). Fixes #669
						for ( indx = 0; indx < columnIndex + 1; indx++ ) {
							val = filters[indx] || '';
							// break out of loop if we've already determined not to search filtered rows
							if ( !searchFiltered ) { indx = columnIndex; }
							// search already filtered rows if...
							searchFiltered = searchFiltered && lastSearch.length &&
								// there are no changes from beginning of filter
								val.indexOf( lastSearch[indx] || '' ) === 0 &&
								// if there is NOT a logical 'or', or range ( 'to' or '-' ) in the string
								!tsfRegex.alreadyFiltered.test( val ) &&
								// if we are not doing exact matches, using '|' ( logical or ) or not '!'
								!tsfRegex.exactTest.test( val ) &&
								// don't search only filtered if the value is negative
								// ( '> -10' => '> -100' will ignore hidden rows )
								!( tsfRegex.isNeg1.test( val ) || tsfRegex.isNeg2.test( val ) ) &&
								// if filtering using a select without a 'filter-match' class ( exact match ) - fixes #593
								!( val !== '' && c.$filters && c.$filters.filter( '[data-column="' + indx + '"]' ).find( 'select' ).length &&
									!tsf.matchType( c, indx ) );
						}
					}
					notFiltered = $rows.not( '.' + wo.filter_filteredRow ).length;
					// can't search when all rows are hidden - this happens when looking for exact matches
					if ( searchFiltered && notFiltered === 0 ) { searchFiltered = false; }
					if ( debug ) {
						console.log( 'Filter >> Searching through ' +
							( searchFiltered && notFiltered < len ? notFiltered : 'all' ) + ' rows' );
					}
					if ( data.anyMatchFlag ) {
						if ( c.sortLocaleCompare ) {
							// replace accents
							data.anyMatchFilter = ts.replaceAccents( data.anyMatchFilter );
						}
						if ( wo.filter_defaultFilter && tsfRegex.iQuery.test( vars.defaultAnyFilter ) ) {
							data.anyMatchFilter = tsf.defaultFilter( data.anyMatchFilter, vars.defaultAnyFilter );
							// clear search filtered flag because default filters are not saved to the last search
							searchFiltered = false;
						}
						// make iAnyMatchFilter lowercase unless both filter widget & core ignoreCase options are true
						// when c.ignoreCase is true, the cache contains all lower case data
						data.iAnyMatchFilter = !( wo.filter_ignoreCase && c.ignoreCase ) ?
							data.anyMatchFilter :
							data.anyMatchFilter.toLowerCase();
					}

					// loop through the rows
					for ( rowIndex = 0; rowIndex < len; rowIndex++ ) {

						txt = $rows[ rowIndex ].className;
						// the first row can never be a child row
						isChild = rowIndex && tsfRegex.child.test( txt );
						// skip child rows & already filtered rows
						if ( isChild || ( searchFiltered && tsfRegex.filtered.test( txt ) ) ) {
							continue;
						}

						data.$row = $rows.eq( rowIndex );
						data.rowIndex = rowIndex;
						data.cacheArray = norm_rows[ rowIndex ];
						rowData = data.cacheArray[ c.columns ];
						data.rawArray = rowData.raw;
						data.childRowText = '';

						if ( !wo.filter_childByColumn ) {
							txt = '';
							// child row cached text
							childRow = rowData.child;
							// so, if 'table.config.widgetOptions.filter_childRows' is true and there is
							// a match anywhere in the child row, then it will make the row visible
							// checked here so the option can be changed dynamically
							for ( indx = 0; indx < childRow.length; indx++ ) {
								txt += ' ' + childRow[indx].join( ' ' ) || '';
							}
							data.childRowText = wo.filter_childRows ?
								( wo.filter_ignoreCase ? txt.toLowerCase() : txt ) :
								'';
						}

						showRow = false;
						showParent = tsf.processRow( c, data, vars );
						$row = rowData.$row;

						// don't pass reference to val
						val = showParent ? true : false;
						childRow = rowData.$row.filter( ':gt(0)' );
						if ( wo.filter_childRows && childRow.length ) {
							if ( wo.filter_childByColumn ) {
								if ( !wo.filter_childWithSibs ) {
									// hide all child rows
									childRow.addClass( wo.filter_filteredRow );
									// if only showing resulting child row, only include parent
									$row = $row.eq( 0 );
								}
								// cycle through each child row
								for ( indx = 0; indx < childRow.length; indx++ ) {
									data.$row = childRow.eq( indx );
									data.cacheArray = rowData.child[ indx ];
									data.rawArray = data.cacheArray;
									val = tsf.processRow( c, data, vars );
									// use OR comparison on child rows
									showRow = showRow || val;
									if ( !wo.filter_childWithSibs && val ) {
										childRow.eq( indx ).removeClass( wo.filter_filteredRow );
									}
								}
							}
							// keep parent row match even if no child matches... see #1020
							showRow = showRow || showParent;
						} else {
							showRow = val;
						}
						$row
							.toggleClass( wo.filter_filteredRow, !showRow )[0]
							.display = showRow ? '' : 'none';
					}
				}
				c.filteredRows += $rows.not( '.' + wo.filter_filteredRow ).length;
				c.totalRows += $rows.length;
				ts.processTbody( table, $tbody, false );
			}
			// lastCombinedFilter is no longer used internally
			c.lastCombinedFilter = storedFilters.join(''); // save last search
			// don't save 'filters' directly since it may have altered ( AnyMatch column searches )
			c.lastSearch = storedFilters;
			c.$table.data( 'lastSearch', storedFilters );
			if ( wo.filter_saveFilters && ts.storage ) {
				ts.storage( table, 'tablesorter-filters', tsf.processFilters( storedFilters, true ) );
			}
			if ( debug ) {
				console.log( 'Filter >> Completed search' + ts.benchmark(time) );
			}
			if ( wo.filter_initialized ) {
				c.$table.triggerHandler( 'filterBeforeEnd', c );
				c.$table.triggerHandler( 'filterEnd', c );
			}
			setTimeout( function() {
				ts.applyWidget( c.table ); // make sure zebra widget is applied
			}, 0 );
		},
		getOptionSource: function( table, column, onlyAvail ) {
			table = $( table )[0];
			var c = table.config,
				wo = c.widgetOptions,
				arry = false,
				source = wo.filter_selectSource,
				last = c.$table.data( 'lastSearch' ) || [],
				fxn = typeof source === 'function' ? true : ts.getColumnData( table, source, column );

			if ( onlyAvail && last[column] !== '' ) {
				onlyAvail = false;
			}

			// filter select source option
			if ( fxn === true ) {
				// OVERALL source
				arry = source( table, column, onlyAvail );
			} else if ( fxn instanceof $ || ( $.type( fxn ) === 'string' && fxn.indexOf( '</option>' ) >= 0 ) ) {
				// selectSource is a jQuery object or string of options
				return fxn;
			} else if ( $.isArray( fxn ) ) {
				arry = fxn;
			} else if ( $.type( source ) === 'object' && fxn ) {
				// custom select source function for a SPECIFIC COLUMN
				arry = fxn( table, column, onlyAvail );
				// abort - updating the selects from an external method
				if (arry === null) {
					return null;
				}
			}
			if ( arry === false ) {
				// fall back to original method
				arry = tsf.getOptions( table, column, onlyAvail );
			}

			return tsf.processOptions( table, column, arry );

		},
		processOptions: function( table, column, arry ) {
			if ( !$.isArray( arry ) ) {
				return false;
			}
			table = $( table )[0];
			var cts, txt, indx, len, parsedTxt, str,
				c = table.config,
				validColumn = typeof column !== 'undefined' && column !== null && column >= 0 && column < c.columns,
				direction = validColumn ? c.$headerIndexed[ column ].hasClass( 'filter-select-sort-desc' ) : false,
				parsed = [];
			// get unique elements and sort the list
			// if $.tablesorter.sortText exists ( not in the original tablesorter ),
			// then natural sort the list otherwise use a basic sort
			arry = $.grep( arry, function( value, indx ) {
				if ( value.text ) {
					return true;
				}
				return $.inArray( value, arry ) === indx;
			});
			if ( validColumn && c.$headerIndexed[ column ].hasClass( 'filter-select-nosort' ) ) {
				// unsorted select options
				return arry;
			} else {
				len = arry.length;
				// parse select option values
				for ( indx = 0; indx < len; indx++ ) {
					txt = arry[ indx ];
					// check for object
					str = txt.text ? txt.text : txt;
					// sortNatural breaks if you don't pass it strings
					parsedTxt = ( validColumn && c.parsers && c.parsers.length &&
						c.parsers[ column ].format( str, table, [], column ) || str ).toString();
					parsedTxt = c.widgetOptions.filter_ignoreCase ? parsedTxt.toLowerCase() : parsedTxt;
					// parse array data using set column parser; this DOES NOT pass the original
					// table cell to the parser format function
					if ( txt.text ) {
						txt.parsed = parsedTxt;
						parsed[ parsed.length ] = txt;
					} else {
						parsed[ parsed.length ] = {
							text : txt,
							// check parser length - fixes #934
							parsed : parsedTxt
						};
					}
				}
				// sort parsed select options
				cts = c.textSorter || '';
				parsed.sort( function( a, b ) {
					var x = direction ? b.parsed : a.parsed,
						y = direction ? a.parsed : b.parsed;
					if ( validColumn && typeof cts === 'function' ) {
						// custom OVERALL text sorter
						return cts( x, y, true, column, table );
					} else if ( validColumn && typeof cts === 'object' && cts.hasOwnProperty( column ) ) {
						// custom text sorter for a SPECIFIC COLUMN
						return cts[column]( x, y, true, column, table );
					} else if ( ts.sortNatural ) {
						// fall back to natural sort
						return ts.sortNatural( x, y );
					}
					// using an older version! do a basic sort
					return true;
				});
				// rebuild arry from sorted parsed data
				arry = [];
				len = parsed.length;
				for ( indx = 0; indx < len; indx++ ) {
					arry[ arry.length ] = parsed[indx];
				}
				return arry;
			}
		},
		getOptions: function( table, column, onlyAvail ) {
			table = $( table )[0];
			var rowIndex, tbodyIndex, len, row, cache, indx, child, childLen,
				c = table.config,
				wo = c.widgetOptions,
				arry = [];
			for ( tbodyIndex = 0; tbodyIndex < c.$tbodies.length; tbodyIndex++ ) {
				cache = c.cache[tbodyIndex];
				len = c.cache[tbodyIndex].normalized.length;
				// loop through the rows
				for ( rowIndex = 0; rowIndex < len; rowIndex++ ) {
					// get cached row from cache.row ( old ) or row data object
					// ( new; last item in normalized array )
					row = cache.row ?
						cache.row[ rowIndex ] :
						cache.normalized[ rowIndex ][ c.columns ].$row[0];
					// check if has class filtered
					if ( onlyAvail && row.className.match( wo.filter_filteredRow ) ) {
						continue;
					}
					// get non-normalized cell content
					if ( wo.filter_useParsedData ||
						c.parsers[column].parsed ||
						c.$headerIndexed[column].hasClass( 'filter-parsed' ) ) {
						arry[ arry.length ] = '' + cache.normalized[ rowIndex ][ column ];
						// child row parsed data
						if ( wo.filter_childRows && wo.filter_childByColumn ) {
							childLen = cache.normalized[ rowIndex ][ c.columns ].$row.length - 1;
							for ( indx = 0; indx < childLen; indx++ ) {
								arry[ arry.length ] = '' + cache.normalized[ rowIndex ][ c.columns ].child[ indx ][ column ];
							}
						}
					} else {
						// get raw cached data instead of content directly from the cells
						arry[ arry.length ] = cache.normalized[ rowIndex ][ c.columns ].raw[ column ];
						// child row unparsed data
						if ( wo.filter_childRows && wo.filter_childByColumn ) {
							childLen = cache.normalized[ rowIndex ][ c.columns ].$row.length;
							for ( indx = 1; indx < childLen; indx++ ) {
								child =  cache.normalized[ rowIndex ][ c.columns ].$row.eq( indx ).children().eq( column );
								arry[ arry.length ] = '' + ts.getElementText( c, child, column );
							}
						}
					}
				}
			}
			return arry;
		},
		buildSelect: function( table, column, arry, updating, onlyAvail ) {
			table = $( table )[0];
			column = parseInt( column, 10 );
			if ( !table.config.cache || $.isEmptyObject( table.config.cache ) ) {
				return;
			}

			var indx, val, txt, t, $filters, $filter, option,
				c = table.config,
				wo = c.widgetOptions,
				node = c.$headerIndexed[ column ],
				// t.data( 'placeholder' ) won't work in jQuery older than 1.4.3
				options = '<option value="">' +
					( node.data( 'placeholder' ) ||
						node.attr( 'data-placeholder' ) ||
						wo.filter_placeholder.select || ''
					) + '</option>',
				// Get curent filter value
				currentValue = c.$table
					.find( 'thead' )
					.find( 'select.' + tscss.filter + '[data-column="' + column + '"]' )
					.val();

			// nothing included in arry ( external source ), so get the options from
			// filter_selectSource or column data
			if ( typeof arry === 'undefined' || arry === '' ) {
				arry = tsf.getOptionSource( table, column, onlyAvail );
				// abort, selects are updated by an external method
				if (arry === null) {
					return;
				}
			}

			if ( $.isArray( arry ) ) {
				// build option list
				for ( indx = 0; indx < arry.length; indx++ ) {
					option = arry[ indx ];
					if ( option.text ) {
						// OBJECT!! add data-function-name in case the value is set in filter_functions
						option['data-function-name'] = typeof option.value === 'undefined' ? option.text : option.value;

						// support jQuery < v1.8, otherwise the below code could be shortened to
						// options += $( '<option>', option )[ 0 ].outerHTML;
						options += '<option';
						for ( val in option ) {
							if ( option.hasOwnProperty( val ) && val !== 'text' ) {
								options += ' ' + val + '="' + option[ val ].replace( tsfRegex.quote, '&quot;' ) + '"';
							}
						}
						if ( !option.value ) {
							options += ' value="' + option.text.replace( tsfRegex.quote, '&quot;' ) + '"';
						}
						options += '>' + option.text.replace( tsfRegex.quote, '&quot;' ) + '</option>';
						// above code is needed in jQuery < v1.8

						// make sure we don't turn an object into a string (objects without a "text" property)
					} else if ( '' + option !== '[object Object]' ) {
						txt = option = ( '' + option ).replace( tsfRegex.quote, '&quot;' );
						val = txt;
						// allow including a symbol in the selectSource array
						// 'a-z|A through Z' so that 'a-z' becomes the option value
						// and 'A through Z' becomes the option text
						if ( txt.indexOf( wo.filter_selectSourceSeparator ) >= 0 ) {
							t = txt.split( wo.filter_selectSourceSeparator );
							val = t[0];
							txt = t[1];
						}
						// replace quotes - fixes #242 & ignore empty strings
						// see http://stackoverflow.com/q/14990971/145346
						options += option !== '' ?
							'<option ' +
								( val === txt ? '' : 'data-function-name="' + option + '" ' ) +
								'value="' + val + '">' + txt +
							'</option>' : '';
					}
				}
				// clear arry so it doesn't get appended twice
				arry = [];
			}

			// update all selects in the same column ( clone thead in sticky headers &
			// any external selects ) - fixes 473
			$filters = ( c.$filters ? c.$filters : c.$table.children( 'thead' ) )
				.find( '.' + tscss.filter );
			if ( wo.filter_$externalFilters ) {
				$filters = $filters && $filters.length ?
					$filters.add( wo.filter_$externalFilters ) :
					wo.filter_$externalFilters;
			}
			$filter = $filters.filter( 'select[data-column="' + column + '"]' );

			// make sure there is a select there!
			if ( $filter.length ) {
				$filter[ updating ? 'html' : 'append' ]( options );
				if ( !$.isArray( arry ) ) {
					// append options if arry is provided externally as a string or jQuery object
					// options ( default value ) was already added
					$filter.append( arry ).val( currentValue );
				}
				$filter.val( currentValue );
			}
		},
		buildDefault: function( table, updating ) {
			var columnIndex, $header, noSelect,
				c = table.config,
				wo = c.widgetOptions,
				columns = c.columns;
			// build default select dropdown
			for ( columnIndex = 0; columnIndex < columns; columnIndex++ ) {
				$header = c.$headerIndexed[columnIndex];
				noSelect = !( $header.hasClass( 'filter-false' ) || $header.hasClass( 'parser-false' ) );
				// look for the filter-select class; build/update it if found
				if ( ( $header.hasClass( 'filter-select' ) ||
					ts.getColumnData( table, wo.filter_functions, columnIndex ) === true ) && noSelect ) {
					tsf.buildSelect( table, columnIndex, '', updating, $header.hasClass( wo.filter_onlyAvail ) );
				}
			}
		}
	};

	// filter regex variable
	tsfRegex = tsf.regex;

	ts.getFilters = function( table, getRaw, setFilters, skipFirst ) {
		var i, $filters, $column, cols,
			filters = [],
			c = table ? $( table )[0].config : '',
			wo = c ? c.widgetOptions : '';
		if ( ( getRaw !== true && wo && !wo.filter_columnFilters ) ||
			// setFilters called, but last search is exactly the same as the current
			// fixes issue #733 & #903 where calling update causes the input values to reset
			( $.isArray(setFilters) && tsf.equalFilters(c, setFilters, c.lastSearch) )
		) {
			return $( table ).data( 'lastSearch' ) || [];
		}
		if ( c ) {
			if ( c.$filters ) {
				$filters = c.$filters.find( '.' + tscss.filter );
			}
			if ( wo.filter_$externalFilters ) {
				$filters = $filters && $filters.length ?
					$filters.add( wo.filter_$externalFilters ) :
					wo.filter_$externalFilters;
			}
			if ( $filters && $filters.length ) {
				filters = setFilters || [];
				for ( i = 0; i < c.columns + 1; i++ ) {
					cols = ( i === c.columns ?
						// 'all' columns can now include a range or set of columms ( data-column='0-2,4,6-7' )
						wo.filter_anyColumnSelector + ',' + wo.filter_multipleColumnSelector :
						'[data-column="' + i + '"]' );
					$column = $filters.filter( cols );
					if ( $column.length ) {
						// move the latest search to the first slot in the array
						$column = tsf.getLatestSearch( $column );
						if ( $.isArray( setFilters ) ) {
							// skip first ( latest input ) to maintain cursor position while typing
							if ( skipFirst && $column.length > 1 ) {
								$column = $column.slice( 1 );
							}
							if ( i === c.columns ) {
								// prevent data-column='all' from filling data-column='0,1' ( etc )
								cols = $column.filter( wo.filter_anyColumnSelector );
								$column = cols.length ? cols : $column;
							}
							$column
								.val( setFilters[ i ] )
								// must include a namespace here; but not c.namespace + 'filter'?
								.trigger( 'change' + c.namespace );
						} else {
							filters[i] = $column.val() || '';
							// don't change the first... it will move the cursor
							if ( i === c.columns ) {
								// don't update range columns from 'all' setting
								$column
									.slice( 1 )
									.filter( '[data-column*="' + $column.attr( 'data-column' ) + '"]' )
									.val( filters[ i ] );
							} else {
								$column
									.slice( 1 )
									.val( filters[ i ] );
							}
						}
						// save any match input dynamically
						if ( i === c.columns && $column.length ) {
							wo.filter_$anyMatch = $column;
						}
					}
				}
			}
		}
		return filters;
	};

	ts.setFilters = function( table, filter, apply, skipFirst ) {
		var c = table ? $( table )[0].config : '',
			valid = ts.getFilters( table, true, filter, skipFirst );
		// default apply to "true"
		if ( typeof apply === 'undefined' ) {
			apply = true;
		}
		if ( c && apply ) {
			// ensure new set filters are applied, even if the search is the same
			c.lastCombinedFilter = null;
			c.lastSearch = [];
			tsf.searching( c.table, filter, skipFirst );
			c.$table.triggerHandler( 'filterFomatterUpdate' );
		}
		return valid.length !== 0;
	};

})( jQuery );

/*! Widget: stickyHeaders - updated 9/27/2017 (v2.29.0) *//*
 * Requires tablesorter v2.8+ and jQuery 1.4.3+
 * by Rob Garrison
 */
;(function ($, window) {
	'use strict';
	var ts = $.tablesorter || {};

	$.extend(ts.css, {
		sticky    : 'tablesorter-stickyHeader', // stickyHeader
		stickyVis : 'tablesorter-sticky-visible',
		stickyHide: 'tablesorter-sticky-hidden',
		stickyWrap: 'tablesorter-sticky-wrapper'
	});

	// Add a resize event to table headers
	ts.addHeaderResizeEvent = function(table, disable, settings) {
		table = $(table)[0]; // make sure we're using a dom element
		if ( !table.config ) { return; }
		var defaults = {
				timer : 250
			},
			options = $.extend({}, defaults, settings),
			c = table.config,
			wo = c.widgetOptions,
			checkSizes = function( triggerEvent ) {
				var index, headers, $header, sizes, width, height,
					len = c.$headers.length;
				wo.resize_flag = true;
				headers = [];
				for ( index = 0; index < len; index++ ) {
					$header = c.$headers.eq( index );
					sizes = $header.data( 'savedSizes' ) || [ 0, 0 ]; // fixes #394
					width = $header[0].offsetWidth;
					height = $header[0].offsetHeight;
					if ( width !== sizes[0] || height !== sizes[1] ) {
						$header.data( 'savedSizes', [ width, height ] );
						headers.push( $header[0] );
					}
				}
				if ( headers.length && triggerEvent !== false ) {
					c.$table.triggerHandler( 'resize', [ headers ] );
				}
				wo.resize_flag = false;
			};
		clearInterval(wo.resize_timer);
		if (disable) {
			wo.resize_flag = false;
			return false;
		}
		checkSizes( false );
		wo.resize_timer = setInterval(function() {
			if (wo.resize_flag) { return; }
			checkSizes();
		}, options.timer);
	};

	function getStickyOffset(c, wo) {
		var $el = isNaN(wo.stickyHeaders_offset) ? $(wo.stickyHeaders_offset) : [];
		return $el.length ?
			$el.height() || 0 :
			parseInt(wo.stickyHeaders_offset, 10) || 0;
	}

	// Sticky headers based on this awesome article:
	// http://css-tricks.com/13465-persistent-headers/
	// and https://github.com/jmosbech/StickyTableHeaders by Jonas Mosbech
	// **************************
	ts.addWidget({
		id: 'stickyHeaders',
		priority: 54, // sticky widget must be initialized after the filter & before pager widget!
		options: {
			stickyHeaders : '',       // extra class name added to the sticky header row
			stickyHeaders_appendTo : null, // jQuery selector or object to phycially attach the sticky headers
			stickyHeaders_attachTo : null, // jQuery selector or object to attach scroll listener to (overridden by xScroll & yScroll settings)
			stickyHeaders_xScroll : null, // jQuery selector or object to monitor horizontal scroll position (defaults: xScroll > attachTo > window)
			stickyHeaders_yScroll : null, // jQuery selector or object to monitor vertical scroll position (defaults: yScroll > attachTo > window)
			stickyHeaders_offset : 0, // number or jquery selector targeting the position:fixed element
			stickyHeaders_filteredToTop: true, // scroll table top into view after filtering
			stickyHeaders_cloneId : '-sticky', // added to table ID, if it exists
			stickyHeaders_addResizeEvent : true, // trigger 'resize' event on headers
			stickyHeaders_includeCaption : true, // if false and a caption exist, it won't be included in the sticky header
			stickyHeaders_zIndex : 2 // The zIndex of the stickyHeaders, allows the user to adjust this to their needs
		},
		format: function(table, c, wo) {
			// filter widget doesn't initialize on an empty table. Fixes #449
			if ( c.$table.hasClass('hasStickyHeaders') || ($.inArray('filter', c.widgets) >= 0 && !c.$table.hasClass('hasFilters')) ) {
				return;
			}
			var index, len, $t,
				$table = c.$table,
				// add position: relative to attach element, hopefully it won't cause trouble.
				$attach = $(wo.stickyHeaders_attachTo || wo.stickyHeaders_appendTo),
				namespace = c.namespace + 'stickyheaders ',
				// element to watch for the scroll event
				$yScroll = $(wo.stickyHeaders_yScroll || wo.stickyHeaders_attachTo || window),
				$xScroll = $(wo.stickyHeaders_xScroll || wo.stickyHeaders_attachTo || window),
				$thead = $table.children('thead:first'),
				$header = $thead.children('tr').not('.sticky-false').children(),
				$tfoot = $table.children('tfoot'),
				stickyOffset = getStickyOffset(c, wo),
				// is this table nested? If so, find parent sticky header wrapper (div, not table)
				$nestedSticky = $table.parent().closest('.' + ts.css.table).hasClass('hasStickyHeaders') ?
					$table.parent().closest('table.tablesorter')[0].config.widgetOptions.$sticky.parent() : [],
				nestedStickyTop = $nestedSticky.length ? $nestedSticky.height() : 0,
				// clone table, then wrap to make sticky header
				$stickyTable = wo.$sticky = $table.clone()
					.addClass('containsStickyHeaders ' + ts.css.sticky + ' ' + wo.stickyHeaders + ' ' + c.namespace.slice(1) + '_extra_table' )
					.wrap('<div class="' + ts.css.stickyWrap + '">'),
				$stickyWrap = $stickyTable.parent()
					.addClass(ts.css.stickyHide)
					.css({
						position   : $attach.length ? 'absolute' : 'fixed',
						padding    : parseInt( $stickyTable.parent().parent().css('padding-left'), 10 ),
						top        : stickyOffset + nestedStickyTop,
						left       : 0,
						visibility : 'hidden',
						zIndex     : wo.stickyHeaders_zIndex || 2
					}),
				$stickyThead = $stickyTable.children('thead:first'),
				$stickyCells,
				laststate = '',
				setWidth = function($orig, $clone) {
					var index, width, border, $cell, $this,
						$cells = $orig.filter(':visible'),
						len = $cells.length;
					for ( index = 0; index < len; index++ ) {
						$cell = $clone.filter(':visible').eq(index);
						$this = $cells.eq(index);
						// code from https://github.com/jmosbech/StickyTableHeaders
						if ($this.css('box-sizing') === 'border-box') {
							width = $this.outerWidth();
						} else {
							if ($cell.css('border-collapse') === 'collapse') {
								if (window.getComputedStyle) {
									width = parseFloat( window.getComputedStyle($this[0], null).width );
								} else {
									// ie8 only
									border = parseFloat( $this.css('border-width') );
									width = $this.outerWidth() - parseFloat( $this.css('padding-left') ) - parseFloat( $this.css('padding-right') ) - border;
								}
							} else {
								width = $this.width();
							}
						}
						$cell.css({
							'width': width,
							'min-width': width,
							'max-width': width
						});
					}
				},
				getLeftPosition = function(yWindow) {
					if (yWindow === false && $nestedSticky.length) {
						return $table.position().left;
					}
					return $attach.length ?
						parseInt($attach.css('padding-left'), 10) || 0 :
						$table.offset().left - parseInt($table.css('margin-left'), 10) - $(window).scrollLeft();
				},
				resizeHeader = function() {
					$stickyWrap.css({
						left : getLeftPosition(),
						width: $table.outerWidth()
					});
					setWidth( $table, $stickyTable );
					setWidth( $header, $stickyCells );
				},
				scrollSticky = function( resizing ) {
					if (!$table.is(':visible')) { return; } // fixes #278
					// Detect nested tables - fixes #724
					nestedStickyTop = $nestedSticky.length ? $nestedSticky.offset().top - $yScroll.scrollTop() + $nestedSticky.height() : 0;
					var tmp,
						offset = $table.offset(),
						stickyOffset = getStickyOffset(c, wo),
						yWindow = $.isWindow( $yScroll[0] ), // $.isWindow needs jQuery 1.4.3
						yScroll = yWindow ?
							$yScroll.scrollTop() :
							// use parent sticky position if nested AND inside of a scrollable element - see #1512
							$nestedSticky.length ? parseInt($nestedSticky[0].style.top, 10) : $yScroll.offset().top,
						attachTop = $attach.length ? yScroll : $yScroll.scrollTop(),
						captionHeight = wo.stickyHeaders_includeCaption ? 0 : $table.children( 'caption' ).height() || 0,
						scrollTop = attachTop + stickyOffset + nestedStickyTop - captionHeight,
						tableHeight = $table.height() - ($stickyWrap.height() + ($tfoot.height() || 0)) - captionHeight,
						isVisible = ( scrollTop > offset.top ) && ( scrollTop < offset.top + tableHeight ) ? 'visible' : 'hidden',
						state = isVisible === 'visible' ? ts.css.stickyVis : ts.css.stickyHide,
						needsUpdating = !$stickyWrap.hasClass( state ),
						cssSettings = { visibility : isVisible };
					if ($attach.length) {
						// attached sticky headers always need updating
						needsUpdating = true;
						cssSettings.top = yWindow ? scrollTop - $attach.offset().top : $attach.scrollTop();
					}
					// adjust when scrolling horizontally - fixes issue #143
					tmp = getLeftPosition(yWindow);
					if (tmp !== parseInt($stickyWrap.css('left'), 10)) {
						needsUpdating = true;
						cssSettings.left = tmp;
					}
					cssSettings.top = ( cssSettings.top || 0 ) +
						// If nested AND inside of a scrollable element, only add parent sticky height
						(!yWindow && $nestedSticky.length ? $nestedSticky.height() : stickyOffset + nestedStickyTop);
					if (needsUpdating) {
						$stickyWrap
							.removeClass( ts.css.stickyVis + ' ' + ts.css.stickyHide )
							.addClass( state )
							.css(cssSettings);
					}
					if (isVisible !== laststate || resizing) {
						// make sure the column widths match
						resizeHeader();
						laststate = isVisible;
					}
				};
			// only add a position relative if a position isn't already defined
			if ($attach.length && !$attach.css('position')) {
				$attach.css('position', 'relative');
			}
			// fix clone ID, if it exists - fixes #271
			if ($stickyTable.attr('id')) { $stickyTable[0].id += wo.stickyHeaders_cloneId; }
			// clear out cloned table, except for sticky header
			// include caption & filter row (fixes #126 & #249) - don't remove cells to get correct cell indexing
			$stickyTable.find('> thead:gt(0), tr.sticky-false').hide();
			$stickyTable.find('> tbody, > tfoot').remove();
			$stickyTable.find('caption').toggle(wo.stickyHeaders_includeCaption);
			// issue #172 - find td/th in sticky header
			$stickyCells = $stickyThead.children().children();
			$stickyTable.css({ height:0, width:0, margin: 0 });
			// remove resizable block
			$stickyCells.find('.' + ts.css.resizer).remove();
			// update sticky header class names to match real header after sorting
			$table
				.addClass('hasStickyHeaders')
				.bind('pagerComplete' + namespace, function() {
					resizeHeader();
				});

			ts.bindEvents(table, $stickyThead.children().children('.' + ts.css.header));

			if (wo.stickyHeaders_appendTo) {
				$(wo.stickyHeaders_appendTo).append( $stickyWrap );
			} else {
				// add stickyheaders AFTER the table. If the table is selected by ID, the original one (first) will be returned.
				$table.after( $stickyWrap );
			}

			// onRenderHeader is defined, we need to do something about it (fixes #641)
			if (c.onRenderHeader) {
				$t = $stickyThead.children('tr').children();
				len = $t.length;
				for ( index = 0; index < len; index++ ) {
					// send second parameter
					c.onRenderHeader.apply( $t.eq( index ), [ index, c, $stickyTable ] );
				}
			}
			// make it sticky!
			$xScroll.add($yScroll)
				.unbind( ('scroll resize '.split(' ').join( namespace )).replace(/\s+/g, ' ') )
				.bind('scroll resize '.split(' ').join( namespace ), function( event ) {
					scrollSticky( event.type === 'resize' );
				});
			c.$table
				.unbind('stickyHeadersUpdate' + namespace)
				.bind('stickyHeadersUpdate' + namespace, function() {
					scrollSticky( true );
				});

			if (wo.stickyHeaders_addResizeEvent) {
				ts.addHeaderResizeEvent(table);
			}

			// look for filter widget
			if ($table.hasClass('hasFilters') && wo.filter_columnFilters) {
				// scroll table into view after filtering, if sticky header is active - #482
				$table.bind('filterEnd' + namespace, function() {
					// $(':focus') needs jQuery 1.6+
					var $td = $(document.activeElement).closest('td'),
						column = $td.parent().children().index($td);
					// only scroll if sticky header is active
					if ($stickyWrap.hasClass(ts.css.stickyVis) && wo.stickyHeaders_filteredToTop) {
						// scroll to original table (not sticky clone)
						window.scrollTo(0, $table.position().top);
						// give same input/select focus; check if c.$filters exists; fixes #594
						if (column >= 0 && c.$filters) {
							c.$filters.eq(column).find('a, select, input').filter(':visible').focus();
						}
					}
				});
				ts.filter.bindSearch( $table, $stickyCells.find('.' + ts.css.filter) );
				// support hideFilters
				if (wo.filter_hideFilters) {
					ts.filter.hideFilters(c, $stickyTable);
				}
			}

			// resize table (Firefox)
			if (wo.stickyHeaders_addResizeEvent) {
				$table.bind('resize' + c.namespace + 'stickyheaders', function() {
					resizeHeader();
				});
			}

			// make sure sticky is visible if page is partially scrolled
			scrollSticky( true );
			$table.triggerHandler('stickyHeadersInit');

		},
		remove: function(table, c, wo) {
			var namespace = c.namespace + 'stickyheaders ';
			c.$table
				.removeClass('hasStickyHeaders')
				.unbind( ('pagerComplete resize filterEnd stickyHeadersUpdate '.split(' ').join(namespace)).replace(/\s+/g, ' ') )
				.next('.' + ts.css.stickyWrap).remove();
			if (wo.$sticky && wo.$sticky.length) { wo.$sticky.remove(); } // remove cloned table
			$(window)
				.add(wo.stickyHeaders_xScroll)
				.add(wo.stickyHeaders_yScroll)
				.add(wo.stickyHeaders_attachTo)
				.unbind( ('scroll resize '.split(' ').join(namespace)).replace(/\s+/g, ' ') );
			ts.addHeaderResizeEvent(table, true);
		}
	});

})(jQuery, window);

/*! Widget: resizable - updated 2018-03-26 (v2.30.2) */
/*jshint browser:true, jquery:true, unused:false */
;(function ($, window) {
	'use strict';
	var ts = $.tablesorter || {};

	$.extend(ts.css, {
		resizableContainer : 'tablesorter-resizable-container',
		resizableHandle    : 'tablesorter-resizable-handle',
		resizableNoSelect  : 'tablesorter-disableSelection',
		resizableStorage   : 'tablesorter-resizable'
	});

	// Add extra scroller css
	$(function() {
		var s = '<style>' +
			'body.' + ts.css.resizableNoSelect + ' { -ms-user-select: none; -moz-user-select: -moz-none;' +
				'-khtml-user-select: none; -webkit-user-select: none; user-select: none; }' +
			'.' + ts.css.resizableContainer + ' { position: relative; height: 1px; }' +
			// make handle z-index > than stickyHeader z-index, so the handle stays above sticky header
			'.' + ts.css.resizableHandle + ' { position: absolute; display: inline-block; width: 8px;' +
				'top: 1px; cursor: ew-resize; z-index: 3; user-select: none; -moz-user-select: none; }' +
			'</style>';
		$('head').append(s);
	});

	ts.resizable = {
		init : function( c, wo ) {
			if ( c.$table.hasClass( 'hasResizable' ) ) { return; }
			c.$table.addClass( 'hasResizable' );

			var noResize, $header, column, storedSizes, tmp,
				$table = c.$table,
				$parent = $table.parent(),
				marginTop = parseInt( $table.css( 'margin-top' ), 10 ),

			// internal variables
			vars = wo.resizable_vars = {
				useStorage : ts.storage && wo.resizable !== false,
				$wrap : $parent,
				mouseXPosition : 0,
				$target : null,
				$next : null,
				overflow : $parent.css('overflow') === 'auto' ||
					$parent.css('overflow') === 'scroll' ||
					$parent.css('overflow-x') === 'auto' ||
					$parent.css('overflow-x') === 'scroll',
				storedSizes : []
			};

			// set default widths
			ts.resizableReset( c.table, true );

			// now get measurements!
			vars.tableWidth = $table.width();
			// attempt to autodetect
			vars.fullWidth = Math.abs( $parent.width() - vars.tableWidth ) < 20;

			/*
			// Hacky method to determine if table width is set to 'auto'
			// http://stackoverflow.com/a/20892048/145346
			if ( !vars.fullWidth ) {
				tmp = $table.width();
				$header = $table.wrap('<span>').parent(); // temp variable
				storedSizes = parseInt( $table.css( 'margin-left' ), 10 ) || 0;
				$table.css( 'margin-left', storedSizes + 50 );
				vars.tableWidth = $header.width() > tmp ? 'auto' : tmp;
				$table.css( 'margin-left', storedSizes ? storedSizes : '' );
				$header = null;
				$table.unwrap('<span>');
			}
			*/

			if ( vars.useStorage && vars.overflow ) {
				// save table width
				ts.storage( c.table, 'tablesorter-table-original-css-width', vars.tableWidth );
				tmp = ts.storage( c.table, 'tablesorter-table-resized-width' ) || 'auto';
				ts.resizable.setWidth( $table, tmp, true );
			}
			wo.resizable_vars.storedSizes = storedSizes = ( vars.useStorage ?
				ts.storage( c.table, ts.css.resizableStorage ) :
				[] ) || [];
			ts.resizable.setWidths( c, wo, storedSizes );
			ts.resizable.updateStoredSizes( c, wo );

			wo.$resizable_container = $( '<div class="' + ts.css.resizableContainer + '">' )
				.css({ top : marginTop })
				.insertBefore( $table );
			// add container
			for ( column = 0; column < c.columns; column++ ) {
				$header = c.$headerIndexed[ column ];
				tmp = ts.getColumnData( c.table, c.headers, column );
				noResize = ts.getData( $header, tmp, 'resizable' ) === 'false';
				if ( !noResize ) {
					$( '<div class="' + ts.css.resizableHandle + '">' )
						.appendTo( wo.$resizable_container )
						.attr({
							'data-column' : column,
							'unselectable' : 'on'
						})
						.data( 'header', $header )
						.bind( 'selectstart', false );
				}
			}
			ts.resizable.bindings( c, wo );
		},

		updateStoredSizes : function( c, wo ) {
			var column, $header,
				len = c.columns,
				vars = wo.resizable_vars;
			vars.storedSizes = [];
			for ( column = 0; column < len; column++ ) {
				$header = c.$headerIndexed[ column ];
				vars.storedSizes[ column ] = $header.is(':visible') ? $header.width() : 0;
			}
		},

		setWidth : function( $el, width, overflow ) {
			// overflow tables need min & max width set as well
			$el.css({
				'width' : width,
				'min-width' : overflow ? width : '',
				'max-width' : overflow ? width : ''
			});
		},

		setWidths : function( c, wo, storedSizes ) {
			var column, $temp,
				vars = wo.resizable_vars,
				$extra = $( c.namespace + '_extra_headers' ),
				$col = c.$table.children( 'colgroup' ).children( 'col' );
			storedSizes = storedSizes || vars.storedSizes || [];
			// process only if table ID or url match
			if ( storedSizes.length ) {
				for ( column = 0; column < c.columns; column++ ) {
					// set saved resizable widths
					ts.resizable.setWidth( c.$headerIndexed[ column ], storedSizes[ column ], vars.overflow );
					if ( $extra.length ) {
						// stickyHeaders needs to modify min & max width as well
						$temp = $extra.eq( column ).add( $col.eq( column ) );
						ts.resizable.setWidth( $temp, storedSizes[ column ], vars.overflow );
					}
				}
				$temp = $( c.namespace + '_extra_table' );
				if ( $temp.length && !ts.hasWidget( c.table, 'scroller' ) ) {
					ts.resizable.setWidth( $temp, c.$table.outerWidth(), vars.overflow );
				}
			}
		},

		setHandlePosition : function( c, wo ) {
			var startPosition,
				tableHeight = c.$table.height(),
				$handles = wo.$resizable_container.children(),
				handleCenter = Math.floor( $handles.width() / 2 );

			if ( ts.hasWidget( c.table, 'scroller' ) ) {
				tableHeight = 0;
				c.$table.closest( '.' + ts.css.scrollerWrap ).children().each(function() {
					var $this = $(this);
					// center table has a max-height set
					tableHeight += $this.filter('[style*="height"]').length ? $this.height() : $this.children('table').height();
				});
			}

			if ( !wo.resizable_includeFooter && c.$table.children('tfoot').length ) {
				tableHeight -= c.$table.children('tfoot').height();
			}
			// subtract out table left position from resizable handles. Fixes #864
			// jQuery v3.3.0+ appears to include the start position with the $header.position().left; see #1544
			startPosition = parseFloat($.fn.jquery) >= 3.3 ? 0 : c.$table.position().left;
			$handles.each( function() {
				var $this = $(this),
					column = parseInt( $this.attr( 'data-column' ), 10 ),
					columns = c.columns - 1,
					$header = $this.data( 'header' );
				if ( !$header ) { return; } // see #859
				if (
					!$header.is(':visible') ||
					( !wo.resizable_addLastColumn && ts.resizable.checkVisibleColumns(c, column) )
				) {
					$this.hide();
				} else if ( column < columns || column === columns && wo.resizable_addLastColumn ) {
					$this.css({
						display: 'inline-block',
						height : tableHeight,
						left : $header.position().left - startPosition + $header.outerWidth() - handleCenter
					});
				}
			});
		},

		// Fixes #1485
		checkVisibleColumns: function( c, column ) {
			var i,
				len = 0;
			for ( i = column + 1; i < c.columns; i++ ) {
				len += c.$headerIndexed[i].is( ':visible' ) ? 1 : 0;
			}
			return len === 0;
		},

		// prevent text selection while dragging resize bar
		toggleTextSelection : function( c, wo, toggle ) {
			var namespace = c.namespace + 'tsresize';
			wo.resizable_vars.disabled = toggle;
			$( 'body' ).toggleClass( ts.css.resizableNoSelect, toggle );
			if ( toggle ) {
				$( 'body' )
					.attr( 'unselectable', 'on' )
					.bind( 'selectstart' + namespace, false );
			} else {
				$( 'body' )
					.removeAttr( 'unselectable' )
					.unbind( 'selectstart' + namespace );
			}
		},

		bindings : function( c, wo ) {
			var namespace = c.namespace + 'tsresize';
			wo.$resizable_container.children().bind( 'mousedown', function( event ) {
				// save header cell and mouse position
				var column,
					vars = wo.resizable_vars,
					$extras = $( c.namespace + '_extra_headers' ),
					$header = $( event.target ).data( 'header' );

				column = parseInt( $header.attr( 'data-column' ), 10 );
				vars.$target = $header = $header.add( $extras.filter('[data-column="' + column + '"]') );
				vars.target = column;

				// if table is not as wide as it's parent, then resize the table
				vars.$next = event.shiftKey || wo.resizable_targetLast ?
					$header.parent().children().not( '.resizable-false' ).filter( ':last' ) :
					$header.nextAll( ':not(.resizable-false)' ).eq( 0 );

				column = parseInt( vars.$next.attr( 'data-column' ), 10 );
				vars.$next = vars.$next.add( $extras.filter('[data-column="' + column + '"]') );
				vars.next = column;

				vars.mouseXPosition = event.pageX;
				ts.resizable.updateStoredSizes( c, wo );
				ts.resizable.toggleTextSelection(c, wo, true );
			});

			$( document )
				.bind( 'mousemove' + namespace, function( event ) {
					var vars = wo.resizable_vars;
					// ignore mousemove if no mousedown
					if ( !vars.disabled || vars.mouseXPosition === 0 || !vars.$target ) { return; }
					if ( wo.resizable_throttle ) {
						clearTimeout( vars.timer );
						vars.timer = setTimeout( function() {
							ts.resizable.mouseMove( c, wo, event );
						}, isNaN( wo.resizable_throttle ) ? 5 : wo.resizable_throttle );
					} else {
						ts.resizable.mouseMove( c, wo, event );
					}
				})
				.bind( 'mouseup' + namespace, function() {
					if (!wo.resizable_vars.disabled) { return; }
					ts.resizable.toggleTextSelection( c, wo, false );
					ts.resizable.stopResize( c, wo );
					ts.resizable.setHandlePosition( c, wo );
				});

			// resizeEnd event triggered by scroller widget
			$( window ).bind( 'resize' + namespace + ' resizeEnd' + namespace, function() {
				ts.resizable.setHandlePosition( c, wo );
			});

			// right click to reset columns to default widths
			c.$table
				.bind( 'columnUpdate pagerComplete resizableUpdate '.split( ' ' ).join( namespace + ' ' ), function() {
					ts.resizable.setHandlePosition( c, wo );
				})
				.bind( 'resizableReset' + namespace, function() {
					ts.resizableReset( c.table );
				})
				.find( 'thead:first' )
				.add( $( c.namespace + '_extra_table' ).find( 'thead:first' ) )
				.bind( 'contextmenu' + namespace, function() {
					// $.isEmptyObject() needs jQuery 1.4+; allow right click if already reset
					var allowClick = wo.resizable_vars.storedSizes.length === 0;
					ts.resizableReset( c.table );
					ts.resizable.setHandlePosition( c, wo );
					wo.resizable_vars.storedSizes = [];
					return allowClick;
				});

		},

		mouseMove : function( c, wo, event ) {
			if ( wo.resizable_vars.mouseXPosition === 0 || !wo.resizable_vars.$target ) { return; }
			// resize columns
			var column,
				total = 0,
				vars = wo.resizable_vars,
				$next = vars.$next,
				tar = vars.storedSizes[ vars.target ],
				leftEdge = event.pageX - vars.mouseXPosition;
			if ( vars.overflow ) {
				if ( tar + leftEdge > 0 ) {
					vars.storedSizes[ vars.target ] += leftEdge;
					ts.resizable.setWidth( vars.$target, vars.storedSizes[ vars.target ], true );
					// update the entire table width
					for ( column = 0; column < c.columns; column++ ) {
						total += vars.storedSizes[ column ];
					}
					ts.resizable.setWidth( c.$table.add( $( c.namespace + '_extra_table' ) ), total );
				}
				if ( !$next.length ) {
					// if expanding right-most column, scroll the wrapper
					vars.$wrap[0].scrollLeft = c.$table.width();
				}
			} else if ( vars.fullWidth ) {
				vars.storedSizes[ vars.target ] += leftEdge;
				vars.storedSizes[ vars.next ] -= leftEdge;
				ts.resizable.setWidths( c, wo );
			} else {
				vars.storedSizes[ vars.target ] += leftEdge;
				ts.resizable.setWidths( c, wo );
			}
			vars.mouseXPosition = event.pageX;
			// dynamically update sticky header widths
			c.$table.triggerHandler('stickyHeadersUpdate');
		},

		stopResize : function( c, wo ) {
			var vars = wo.resizable_vars;
			ts.resizable.updateStoredSizes( c, wo );
			if ( vars.useStorage ) {
				// save all column widths
				ts.storage( c.table, ts.css.resizableStorage, vars.storedSizes );
				ts.storage( c.table, 'tablesorter-table-resized-width', c.$table.width() );
			}
			vars.mouseXPosition = 0;
			vars.$target = vars.$next = null;
			// will update stickyHeaders, just in case, see #912
			c.$table.triggerHandler('stickyHeadersUpdate');
			c.$table.triggerHandler('resizableComplete');
		}
	};

	// this widget saves the column widths if
	// $.tablesorter.storage function is included
	// **************************
	ts.addWidget({
		id: 'resizable',
		priority: 40,
		options: {
			resizable : true, // save column widths to storage
			resizable_addLastColumn : false,
			resizable_includeFooter: true,
			resizable_widths : [],
			resizable_throttle : false, // set to true (5ms) or any number 0-10 range
			resizable_targetLast : false
		},
		init: function(table, thisWidget, c, wo) {
			ts.resizable.init( c, wo );
		},
		format: function( table, c, wo ) {
			ts.resizable.setHandlePosition( c, wo );
		},
		remove: function( table, c, wo, refreshing ) {
			if (wo.$resizable_container) {
				var namespace = c.namespace + 'tsresize';
				c.$table.add( $( c.namespace + '_extra_table' ) )
					.removeClass('hasResizable')
					.children( 'thead' )
					.unbind( 'contextmenu' + namespace );

				wo.$resizable_container.remove();
				ts.resizable.toggleTextSelection( c, wo, false );
				ts.resizableReset( table, refreshing );
				$( document ).unbind( 'mousemove' + namespace + ' mouseup' + namespace );
			}
		}
	});

	ts.resizableReset = function( table, refreshing ) {
		$( table ).each(function() {
			var index, $t,
				c = this.config,
				wo = c && c.widgetOptions,
				vars = wo.resizable_vars;
			if ( table && c && c.$headerIndexed.length ) {
				// restore the initial table width
				if ( vars.overflow && vars.tableWidth ) {
					ts.resizable.setWidth( c.$table, vars.tableWidth, true );
					if ( vars.useStorage ) {
						ts.storage( table, 'tablesorter-table-resized-width', vars.tableWidth );
					}
				}
				for ( index = 0; index < c.columns; index++ ) {
					$t = c.$headerIndexed[ index ];
					if ( wo.resizable_widths && wo.resizable_widths[ index ] ) {
						ts.resizable.setWidth( $t, wo.resizable_widths[ index ], vars.overflow );
					} else if ( !$t.hasClass( 'resizable-false' ) ) {
						// don't clear the width of any column that is not resizable
						ts.resizable.setWidth( $t, '', vars.overflow );
					}
				}

				// reset stickyHeader widths
				c.$table.triggerHandler( 'stickyHeadersUpdate' );
				if ( ts.storage && !refreshing ) {
					ts.storage( this, ts.css.resizableStorage, [] );
				}
			}
		});
	};

})( jQuery, window );

/*! Widget: saveSort - updated 2018-03-19 (v2.30.1) *//*
* Requires tablesorter v2.16+
* by Rob Garrison
*/
;(function ($) {
	'use strict';
	var ts = $.tablesorter || {};

	function getStoredSortList(c) {
		var stored = ts.storage( c.table, 'tablesorter-savesort' );
		return (stored && stored.hasOwnProperty('sortList') && $.isArray(stored.sortList)) ? stored.sortList : [];
	}

	function sortListChanged(c, sortList) {
		return (sortList || getStoredSortList(c)).join(',') !== c.sortList.join(',');
	}

	// this widget saves the last sort only if the
	// saveSort widget option is true AND the
	// $.tablesorter.storage function is included
	// **************************
	ts.addWidget({
		id: 'saveSort',
		priority: 20,
		options: {
			saveSort : true
		},
		init: function(table, thisWidget, c, wo) {
			// run widget format before all other widgets are applied to the table
			thisWidget.format(table, c, wo, true);
		},
		format: function(table, c, wo, init) {
			var time,
				$table = c.$table,
				saveSort = wo.saveSort !== false, // make saveSort active/inactive; default to true
				sortList = { 'sortList' : c.sortList },
				debug = ts.debug(c, 'saveSort');
			if (debug) {
				time = new Date();
			}
			if ($table.hasClass('hasSaveSort')) {
				if (saveSort && table.hasInitialized && ts.storage && sortListChanged(c)) {
					ts.storage( table, 'tablesorter-savesort', sortList );
					if (debug) {
						console.log('saveSort >> Saving last sort: ' + c.sortList + ts.benchmark(time));
					}
				}
			} else {
				// set table sort on initial run of the widget
				$table.addClass('hasSaveSort');
				sortList = '';
				// get data
				if (ts.storage) {
					sortList = getStoredSortList(c);
					if (debug) {
						console.log('saveSort >> Last sort loaded: "' + sortList + '"' + ts.benchmark(time));
					}
					$table.bind('saveSortReset', function(event) {
						event.stopPropagation();
						ts.storage( table, 'tablesorter-savesort', '' );
					});
				}
				// init is true when widget init is run, this will run this widget before all other widgets have initialized
				// this method allows using this widget in the original tablesorter plugin; but then it will run all widgets twice.
				if (init && sortList && sortList.length > 0) {
					c.sortList = sortList;
				} else if (table.hasInitialized && sortList && sortList.length > 0) {
					// update sort change
					if (sortListChanged(c, sortList)) {
						ts.sortOn(c, sortList);
					}
				}
			}
		},
		remove: function(table, c) {
			c.$table.removeClass('hasSaveSort');
			// clear storage
			if (ts.storage) { ts.storage( table, 'tablesorter-savesort', '' ); }
		}
	});

})(jQuery);
return jQuery.tablesorter;}));
```
