## Supplementary Data SD1 for "Inferring binding specificities of human transcription factors with the wisdom of crowds": jquery.tablesorter.combined.min.js

```
(function(factory){if (typeof define === 'function' && define.amd){define(['jquery'], factory);} else if (typeof module === 'object' && typeof module.exports === 'object'){module.exports = factory(require('jquery'));} else {factory(jQuery);}}(function(jQuery){
/*! tablesorter (FORK) - updated 2024-08-13 (v2.32.0)*/
!function(e){"function"==typeof define&&define.amd?define(["jquery"],e):"object"==typeof module&&"object"==typeof module.exports?module.exports=e(require("jquery")):e(jQuery)}(function(e){return function(R){"use strict";var T=R.tablesorter={version:"2.32.0",parsers:[],widgets:[],defaults:{theme:"default",widthFixed:!1,showProcessing:!1,headerTemplate:"{content}",onRenderTemplate:null,onRenderHeader:null,cancelSelection:!0,tabIndex:!0,dateFormat:"mmddyyyy",sortMultiSortKey:"shiftKey",sortResetKey:"ctrlKey",usNumberFormat:!0,delayInit:!1,serverSideSorting:!1,resort:!0,headers:null,ignoreCase:!0,sortForce:null,sortList:[],sortAppend:null,sortStable:!1,sortInitialOrder:"asc",sortLocaleCompare:!1,sortReset:!1,sortRestart:!1,emptyTo:"bottom",stringTo:"max",duplicateSpan:!0,textExtraction:"basic",textAttribute:"data-text",textSorter:null,numberSorter:null,initWidgets:!0,widgetClass:"widget-{name}",widgets:[],widgetOptions:{zebra:["even","odd"]},initialized:null,tableClass:"",cssAsc:"",cssDesc:"",cssNone:"",cssHeader:"",cssHeaderRow:"",cssProcessing:"",cssChildRow:"tablesorter-childRow",cssInfoBlock:"tablesorter-infoOnly",cssNoSort:"tablesorter-noSort",cssIgnoreRow:"tablesorter-ignoreRow",cssIcon:"tablesorter-icon",cssIconNone:"",cssIconAsc:"",cssIconDesc:"",cssIconDisabled:"",pointerClick:"click",pointerDown:"mousedown",pointerUp:"mouseup",selectorHeaders:"> thead th, > thead td",selectorSort:"th, td",selectorRemove:".remove-me",debug:!1,headerList:[],empties:{},strings:{},parsers:[],globalize:0,imgAttr:0},css:{table:"tablesorter",cssHasChild:"tablesorter-hasChildRow",childRow:"tablesorter-childRow",colgroup:"tablesorter-colgroup",header:"tablesorter-header",headerRow:"tablesorter-headerRow",headerIn:"tablesorter-header-inner",icon:"tablesorter-icon",processing:"tablesorter-processing",sortAsc:"tablesorter-headerAsc",sortDesc:"tablesorter-headerDesc",sortNone:"tablesorter-headerUnSorted"},language:{sortAsc:"Ascending sort applied, ",sortDesc:"Descending sort applied, ",sortNone:"No sort applied, ",sortDisabled:"sorting is disabled",nextAsc:"activate to apply an ascending sort",nextDesc:"activate to apply a descending sort",nextNone:"activate to remove the sort"},regex:{templateContent:/\{content\}/g,templateIcon:/\{icon\}/g,templateName:/\{name\}/i,spaces:/\s+/g,nonWord:/\W/g,formElements:/(input|select|button|textarea)/i,chunk:/(^([+\-]?(?:\d*)(?:\.\d*)?(?:[eE][+\-]?\d+)?)?$|^0x[0-9a-f]+$|\d+)/gi,chunks:/(^\\0|\\0$)/,hex:/^0x[0-9a-f]+$/i,comma:/,/g,digitNonUS:/[\s|\.]/g,digitNegativeTest:/^\s*\([.\d]+\)/,digitNegativeReplace:/^\s*\(([.\d]+)\)/,digitTest:/^[\-+(]?\d+[)]?$/,digitReplace:/[,.'"\s]/g},string:{max:1,min:-1,emptymin:1,emptymax:-1,zero:0,none:0,null:0,top:!0,bottom:!1},keyCodes:{enter:13},dates:{},instanceMethods:{},setup:function(t,r){var e,a,s,i;t&&t.tHead&&0!==t.tBodies.length&&!0!==t.hasInitialized?(e="",a=R(t),s=R.metadata,t.hasInitialized=!1,t.isProcessing=!0,t.config=r,R.data(t,"tablesorter",r),T.debug(r,"core")&&(console[console.group?"group":"log"]("Initializing tablesorter v"+T.version),R.data(t,"startoveralltimer",new Date)),r.supportsDataObject=((i=R.fn.jquery.split("."))[0]=parseInt(i[0],10),1<i[0]||1===i[0]&&4<=parseInt(i[1],10)),r.emptyTo=r.emptyTo.toLowerCase(),r.stringTo=r.stringTo.toLowerCase(),r.last={sortList:[],clickedIndex:-1},/tablesorter\-/.test(a.attr("class"))||(e=""!==r.theme?" tablesorter-"+r.theme:""),r.namespace?r.namespace="."+r.namespace.replace(T.regex.nonWord,""):r.namespace=".tablesorter"+Math.random().toString(16).slice(2),r.table=t,r.$table=a.addClass(T.css.table+" "+r.tableClass+e+" "+r.namespace.slice(1)).attr("role","grid"),r.$headers=a.find(r.selectorHeaders),r.$table.children().children("tr").attr("role","row"),r.$tbodies=a.children("tbody:not(."+r.cssInfoBlock+")").attr({"aria-live":"polite","aria-relevant":"all"}),r.$table.children("caption").length&&((e=r.$table.children("caption")[0]).id||(e.id=r.namespace.slice(1)+"caption"),r.$table.attr("aria-labelledby",e.id)),r.widgetInit={},r.textExtraction=r.$table.attr("data-text-extraction")||r.textExtraction||"basic",T.buildHeaders(r),T.fixColumnWidth(t),T.addWidgetFromClass(t),T.applyWidgetOptions(t),T.setupParsers(r),r.totalRows=0,r.debug&&T.validateOptions(r),r.delayInit||T.buildCache(r),T.bindEvents(t,r.$headers,!0),T.bindMethods(r),r.supportsDataObject&&void 0!==a.data().sortlist?r.sortList=a.data().sortlist:s&&a.metadata()&&a.metadata().sortlist&&(r.sortList=a.metadata().sortlist),T.applyWidget(t,!0),0<r.sortList.length?(r.last.sortList=r.sortList,T.sortOn(r,r.sortList,{},!r.initWidgets)):(T.setHeadersCss(r),r.initWidgets&&T.applyWidget(t,!1)),r.showProcessing&&a.unbind("sortBegin"+r.namespace+" sortEnd"+r.namespace).bind("sortBegin"+r.namespace+" sortEnd"+r.namespace,function(e){clearTimeout(r.timerProcessing),T.isProcessing(t),"sortBegin"===e.type&&(r.timerProcessing=setTimeout(function(){T.isProcessing(t,!0)},500))}),t.hasInitialized=!0,t.isProcessing=!1,T.debug(r,"core")&&(console.log("Overall initialization time:"+T.benchmark(R.data(t,"startoveralltimer"))),T.debug(r,"core"))&&console.groupEnd&&console.groupEnd(),a.triggerHandler("tablesorter-initialized",t),"function"==typeof r.initialized&&r.initialized(t)):T.debug(r,"core")&&(t.hasInitialized?console.warn("Stopping initialization. Tablesorter has already been initialized"):console.error("Stopping initialization! No table, thead or tbody",t))},bindMethods:function(t){var e=t.$table,r=t.namespace,a="sortReset update updateRows updateAll updateHeaders addRows updateCell updateComplete sorton appendCache updateCache applyWidgetId applyWidgets refreshWidgets destroy mouseup mouseleave ".split(" ").join(r+" ");e.unbind(a.replace(T.regex.spaces," ")).bind("sortReset"+r,function(e,t){e.stopPropagation(),T.sortReset(this.config,function(e){e.isApplyingWidgets?setTimeout(function(){T.applyWidget(e,"",t)},100):T.applyWidget(e,"",t)})}).bind("updateAll"+r,function(e,t,r){e.stopPropagation(),T.updateAll(this.config,t,r)}).bind("update"+r+" updateRows"+r,function(e,t,r){e.stopPropagation(),T.update(this.config,t,r)}).bind("updateHeaders"+r,function(e,t){e.stopPropagation(),T.updateHeaders(this.config,t)}).bind("updateCell"+r,function(e,t,r,a){e.stopPropagation(),T.updateCell(this.config,t,r,a)}).bind("addRows"+r,function(e,t,r,a){e.stopPropagation(),T.addRows(this.config,t,r,a)}).bind("updateComplete"+r,function(){this.isUpdating=!1}).bind("sorton"+r,function(e,t,r,a){e.stopPropagation(),T.sortOn(this.config,t,r,a)}).bind("appendCache"+r,function(e,t,r){e.stopPropagation(),T.appendCache(this.config,r),R.isFunction(t)&&t(this)}).bind("updateCache"+r,function(e,t,r){e.stopPropagation(),T.updateCache(this.config,t,r)}).bind("applyWidgetId"+r,function(e,t){e.stopPropagation(),T.applyWidgetId(this,t)}).bind("applyWidgets"+r,function(e,t){e.stopPropagation(),T.applyWidget(this,!1,t)}).bind("refreshWidgets"+r,function(e,t,r){e.stopPropagation(),T.refreshWidgets(this,t,r)}).bind("removeWidget"+r,function(e,t,r){e.stopPropagation(),T.removeWidget(this,t,r)}).bind("destroy"+r,function(e,t,r){e.stopPropagation(),T.destroy(this,t,r)}).bind("resetToLoadState"+r,function(e){e.stopPropagation(),T.removeWidget(this,!0,!1);e=R.extend(!0,{},t.originalSettings);(t=R.extend(!0,{},T.defaults,e)).originalSettings=e,this.hasInitialized=!1,T.setup(this,t)})},bindEvents:function(e,t,r){var a,s=(e=R(e)[0]).config,i=s.namespace,o=null;!0!==r&&(t.addClass(i.slice(1)+"_extra_headers"),(a=T.getClosest(t,"table")).length)&&"TABLE"===a[0].nodeName&&a[0]!==e&&R(a[0]).addClass(i.slice(1)+"_extra_table"),a=(s.pointerDown+" "+s.pointerUp+" "+s.pointerClick+" sort keyup ").replace(T.regex.spaces," ").split(" ").join(i+" "),t.find(s.selectorSort).add(t.filter(s.selectorSort)).unbind(a).bind(a,function(e,t){var r=R(e.target),a=" "+e.type+" ";if(!(1!==(e.which||e.button)&&!a.match(" "+s.pointerClick+" | sort | keyup ")||" keyup "===a&&e.which!==T.keyCodes.enter||a.match(" "+s.pointerClick+" ")&&void 0!==e.which||a.match(" "+s.pointerUp+" ")&&o!==e.target&&!0!==t))if(a.match(" "+s.pointerDown+" "))o=e.target,"1"===(t=r.jquery.split("."))[0]&&t[1]<4&&e.preventDefault();else{if(o=null,a=T.getClosest(R(this),"."+T.css.header),T.regex.formElements.test(e.target.nodeName)||r.hasClass(s.cssNoSort)||0<r.parents("."+s.cssNoSort).length||a.hasClass("sorter-false")||0<r.parents("button").length)return!s.cancelSelection;s.delayInit&&T.isEmptyObject(s.cache)&&T.buildCache(s),s.last.clickedIndex=a.attr("data-column")||a.index(),(t=s.$headerIndexed[s.last.clickedIndex][0])&&!t.sortDisabled&&T.initSort(s,t,e)}}),s.cancelSelection&&t.attr("unselectable","on").bind("selectstart",!1).css({"user-select":"none",MozUserSelect:"none"})},buildHeaders:function(n){var e,l,t,r;for(n.headerList=[],n.headerContent=[],n.sortVars=[],T.debug(n,"core")&&(t=new Date),n.columns=T.computeColumnIndex(n.$table.children("thead, tfoot").children("tr")),l=n.cssIcon?'<i class="'+(n.cssIcon===T.css.icon?T.css.icon:n.cssIcon+" "+T.css.icon)+'"></i>':"",n.$headers=R(R.map(n.$table.find(n.selectorHeaders),function(e,t){var r,a,s,i,o=R(e);if(!T.getClosest(o,"tr").hasClass(n.cssIgnoreRow))return/(th|td)/i.test(e.nodeName)||(i=T.getClosest(o,"th, td"),o.attr("data-column",i.attr("data-column"))),r=T.getColumnData(n.table,n.headers,t,!0),n.headerContent[t]=o.html(),""===n.headerTemplate||o.find("."+T.css.headerIn).length||(s=n.headerTemplate.replace(T.regex.templateContent,o.html()).replace(T.regex.templateIcon,o.find("."+T.css.icon).length?"":l),n.onRenderTemplate&&(a=n.onRenderTemplate.apply(o,[t,s]))&&"string"==typeof a&&(s=a),o.html('<div class="'+T.css.headerIn+'">'+s+"</div>")),n.onRenderHeader&&n.onRenderHeader.apply(o,[t,n,n.$table]),a=parseInt(o.attr("data-column"),10),e.column=a,i=T.getOrder(T.getData(o,r,"sortInitialOrder")||n.sortInitialOrder),n.sortVars[a]={count:-1,order:i?n.sortReset?[1,0,2]:[1,0]:n.sortReset?[0,1,2]:[0,1],lockedOrder:!1,sortedBy:""},void 0!==(i=T.getData(o,r,"lockedOrder")||!1)&&!1!==i&&(n.sortVars[a].lockedOrder=!0,n.sortVars[a].order=T.getOrder(i)?[1,1]:[0,0]),n.headerList[t]=e,o.addClass(T.css.header+" "+n.cssHeader),T.getClosest(o,"tr").addClass(T.css.headerRow+" "+n.cssHeaderRow).attr("role","row"),n.tabIndex&&o.attr("tabindex",0),e})),n.$headerIndexed=[],r=0;r<n.columns;r++)T.isEmptyObject(n.sortVars[r])&&(n.sortVars[r]={}),e=n.$headers.filter('[data-column="'+r+'"]'),n.$headerIndexed[r]=e.length?(e.not(".sorter-false").length?e.not(".sorter-false"):e).filter(":last"):R();n.$table.find(n.selectorHeaders).attr({scope:"col",role:"columnheader"}),T.updateHeader(n),T.debug(n,"core")&&(console.log("Built headers:"+T.benchmark(t)),console.log(n.$headers))},addInstanceMethods:function(e){R.extend(T.instanceMethods,e)},setupParsers:function(e,t){var r,a,s,i,o,n,l,c,d,f,u,g,p,h,m=e.table,b=0,y=T.debug(e,"core"),w={};if(e.$tbodies=e.$table.children("tbody:not(."+e.cssInfoBlock+")"),0===(h=(p=void 0===t?e.$tbodies:t).length))return y?console.warn("Warning: *Empty table!* Not building a parser cache"):"";for(y&&(g=new Date,console[console.group?"group":"log"]("Detecting parsers for each column")),a={extractors:[],parsers:[]};b<h;){if((r=p[b].rows).length)for(i=e.columns,n=o=0;n<i;n++){if((l=e.$headerIndexed[o])&&l.length&&(c=T.getColumnData(m,e.headers,o),u=T.getParserById(T.getData(l,c,"extractor")),f=T.getParserById(T.getData(l,c,"sorter")),d="false"===T.getData(l,c,"parser"),e.empties[o]=(T.getData(l,c,"empty")||e.emptyTo||(e.emptyToBottom?"bottom":"top")).toLowerCase(),e.strings[o]=(T.getData(l,c,"string")||e.stringTo||"max").toLowerCase(),u=u||!1,f=(f=d?T.getParserById("no-parser"):f)||T.detectParserForColumn(e,r,-1,o),y&&(w["("+o+") "+l.text()]={parser:f.id,extractor:u?u.id:"none",string:e.strings[o],empty:e.empties[o]}),a.parsers[o]=f,a.extractors[o]=u,0<(s=l[0].colSpan-1)))for(o+=s,i+=s;0<s+1;)a.parsers[o-s]=f,a.extractors[o-s]=u,s--;o++}b+=a.parsers.length?h:1}y&&(T.isEmptyObject(w)?console.warn("  No parsers detected!"):console[console.table?"table":"log"](w),console.log("Completed detecting parsers"+T.benchmark(g)),console.groupEnd)&&console.groupEnd(),e.parsers=a.parsers,e.extractors=a.extractors},addParser:function(e){for(var t=T.parsers.length,r=!0,a=0;a<t;a++)T.parsers[a].id.toLowerCase()===e.id.toLowerCase()&&(r=!1);r&&(T.parsers[T.parsers.length]=e)},getParserById:function(e){if("false"!=e)for(var t=T.parsers.length,r=0;r<t;r++)if(T.parsers[r].id.toLowerCase()===e.toString().toLowerCase())return T.parsers[r];return!1},detectParserForColumn:function(e,t,r,a){for(var s,i,o,n=T.parsers.length,l=!1,c="",d=T.debug(e,"core"),f=!0;""===c&&f;)(o=t[++r])&&r<50?o.className.indexOf(T.cssIgnoreRow)<0&&(l=t[r].cells[a],c=T.getElementText(e,l,a),i=R(l),d)&&console.log("Checking if value was empty on row "+r+", column: "+a+': "'+c+'"'):f=!1;for(;0<=--n;)if((s=T.parsers[n])&&"text"!==s.id&&s.is&&s.is(c,e.table,l,i))return s;return T.getParserById("text")},getElementText:function(e,t,r){var a,s,i;return t?(s=e.textExtraction||"",i=t.jquery?t:R(t),"string"==typeof s?"basic"===s&&void 0!==(a=i.attr(e.textAttribute))?R.trim(a):R.trim(t.textContent||i.text()):"function"==typeof s?R.trim(s(i[0],e.table,r)):"function"==typeof(a=T.getColumnData(e.table,s,r))?R.trim(a(i[0],e.table,r)):R.trim(i[0].textContent||i.text())):""},getParsedText:function(e,t,r,a){var s=""+(a=void 0===a?T.getElementText(e,t,r):a),i=e.parsers[r],o=e.extractors[r];return s=i&&(o&&"function"==typeof o.format&&(a=o.format(a,e.table,t,r)),s="no-parser"===i.id?"":i.format(""+a,e.table,t,r),e.ignoreCase)&&"string"==typeof s?s.toLowerCase():s},buildCache:function(e,t,r){var a,s,i,o,n,l,c,d,f,u,g,p,h,m,b,y,w,v,x,C,_,$,S=e.table,z=e.parsers,F=T.debug(e,"core");if(e.$tbodies=e.$table.children("tbody:not(."+e.cssInfoBlock+")"),c=void 0===r?e.$tbodies:r,e.cache={},e.totalRows=0,!z)return F?console.warn("Warning: *Empty table!* Not building a cache"):"";for(F&&(p=new Date),e.showProcessing&&T.isProcessing(S,!0),l=0;l<c.length;l++){for(y=[],a=e.cache[l]={normalized:[]},h=c[l]&&c[l].rows.length||0,o=0;o<h;++o)if(m={child:[],raw:[]},f=[],!(d=R(c[l].rows[o])).hasClass(e.selectorRemove.slice(1)))if(d.hasClass(e.cssChildRow)&&0!==o)for(_=a.normalized.length-1,(b=a.normalized[_][e.columns]).$row=b.$row.add(d),d.prev().hasClass(e.cssChildRow)||d.prev().addClass(T.css.cssHasChild),u=d.children("th, td"),_=b.child.length,b.child[_]=[],C=e.columns,n=v=0;n<C;n++)(g=u[n])&&(b.child[_][n]=T.getParsedText(e,g,n),0<(w=u[n].colSpan-1))&&(v+=w,C+=w),v++;else{for(m.$row=d,m.order=o,C=e.columns,n=v=0;n<C;++n){if((g=d[0].cells[n])&&v<e.columns&&(!(x=void 0!==z[v])&&F&&console.warn("No parser found for row: "+o+", column: "+n+'; cell containing: "'+R(g).text()+'"; does it have a header?'),s=T.getElementText(e,g,v),m.raw[v]=s,i=T.getParsedText(e,g,v,s),f[v]=i,x&&"numeric"===(z[v].type||"").toLowerCase()&&(y[v]=Math.max(Math.abs(i)||0,y[v]||0)),0<(w=g.colSpan-1))){for($=0;$<=w;)i=e.duplicateSpan||0===$?i:"string"!=typeof e.textExtraction&&T.getElementText(e,g,v+$)||"",m.raw[v+$]=i,f[v+$]=i,$++;v+=w,C+=w}v++}f[e.columns]=m,a.normalized[a.normalized.length]=f}a.colMax=y,e.totalRows+=a.normalized.length}if(e.showProcessing&&T.isProcessing(S),F){for(_=Math.min(5,e.cache[0].normalized.length),console[console.group?"group":"log"]("Building cache for "+e.totalRows+" rows (showing "+_+" rows in log) and "+e.columns+" columns"+T.benchmark(p)),s={},n=0;n<e.columns;n++)for(v=0;v<_;v++)s["row: "+v]||(s["row: "+v]={}),s["row: "+v][e.$headerIndexed[n].text()]=e.cache[0].normalized[v][n];console[console.table?"table":"log"](s),console.groupEnd&&console.groupEnd()}R.isFunction(t)&&t(S)},getColumnText:function(e,t,r,a){var s,i,o,n,l,c,d,f,u,g,p="function"==typeof r,h="all"===t,m={raw:[],parsed:[],$cell:[]},b=(e=R(e)[0]).config;if(!T.isEmptyObject(b)){for(l=b.$tbodies.length,s=0;s<l;s++)for(c=(o=b.cache[s].normalized).length,i=0;i<c;i++)n=o[i],a&&!n[b.columns].$row.is(a)||(g=!0,f=h?n.slice(0,b.columns):n[t],n=n[b.columns],d=h?n.raw:n.raw[t],u=h?n.$row.children():n.$row.children().eq(t),!1!==(g=p?r({tbodyIndex:s,rowIndex:i,parsed:f,raw:d,$row:n.$row,$cell:u}):g)&&(m.parsed[m.parsed.length]=f,m.raw[m.raw.length]=d,m.$cell[m.$cell.length]=u));return m}T.debug(b,"core")&&console.warn("No cache found - aborting getColumnText function!")},setHeadersCss:function(s){function e(e,t){e.removeClass(o).addClass(n[t]).attr("aria-sort",c[t]).find("."+T.css.icon).removeClass(l[2]).addClass(l[t])}var t,r,a=s.sortList,i=a.length,o=T.css.sortNone+" "+s.cssNone,n=[T.css.sortAsc+" "+s.cssAsc,T.css.sortDesc+" "+s.cssDesc],l=[s.cssIconAsc,s.cssIconDesc,s.cssIconNone],c=["ascending","descending"],d=s.$table.find("tfoot tr").children("td, th").add(R(s.namespace+"_extra_headers")).removeClass(n.join(" ")),f=s.$headers.add(R("thead "+s.namespace+"_extra_headers")).removeClass(n.join(" ")).addClass(o).attr("aria-sort","none").find("."+T.css.icon).removeClass(l.join(" ")).end();for(f.not(".sorter-false").find("."+T.css.icon).addClass(l[2]),s.cssIconDisabled&&f.filter(".sorter-false").find("."+T.css.icon).addClass(s.cssIconDisabled),t=0;t<i;t++)if(2!==a[t][1]){if((f=(f=s.$headers.filter(function(e){for(var t=!0,e=s.$headers.eq(e),r=parseInt(e.attr("data-column"),10),a=r+T.getClosest(e,"th, td")[0].colSpan;r<a;r++)t=t||!1;return t})).not(".sorter-false").filter('[data-column="'+a[t][0]+'"]'+(1===i?":last":""))).length)for(r=0;r<f.length;r++)f[r].sortDisabled||e(f.eq(r),a[t][1]);d.length&&e(d.filter('[data-column="'+a[t][0]+'"]'),a[t][1])}for(i=s.$headers.length,t=0;t<i;t++)T.setColumnAriaLabel(s,s.$headers.eq(t))},getClosest:function(e,t){return R.fn.closest?e.closest(t):e.is(t)?e:e.parents(t).filter(":first")},setColumnAriaLabel:function(e,t,r){var a,s;t.length&&(a=parseInt(t.attr("data-column"),10),e=e.sortVars[a],a=t.hasClass(T.css.sortAsc)?"sortAsc":t.hasClass(T.css.sortDesc)?"sortDesc":"sortNone",s=R.trim(t.text())+": "+T.language[a],t.hasClass("sorter-false")||!1===r?s+=T.language.sortDisabled:(a=(e.count+1)%e.order.length,r=e.order[a],s+=T.language[0===r?"nextAsc":1===r?"nextDesc":"nextNone"]),t.attr("aria-label",s),e.sortedBy?t.attr("data-sortedBy",e.sortedBy):t.removeAttr("data-sortedBy"))},updateHeader:function(e){for(var t,r,a=e.table,s=e.$headers.length,i=0;i<s;i++)t=e.$headers.eq(i),r=T.getColumnData(a,e.headers,i,!0),r="false"===T.getData(t,r,"sorter")||"false"===T.getData(t,r,"parser"),T.setColumnSort(e,t,r)},setColumnSort:function(e,t,r){var a=e.table.id;t[0].sortDisabled=r,t[r?"addClass":"removeClass"]("sorter-false").attr("aria-disabled",""+r),e.tabIndex&&(r?t.removeAttr("tabindex"):t.attr("tabindex","0")),a&&(r?t.removeAttr("aria-controls"):t.attr("aria-controls",a))},updateHeaderSortCount:function(e,t){var r,a,s,i,o,n,l,c=t||e.sortList,d=c.length;for(e.sortList=[],s=0;s<d;s++)if(n=c[s],(r=parseInt(n[0],10))<e.columns){switch(e.sortVars[r].order||(l=T.getOrder(e.sortInitialOrder)?e.sortReset?[1,0,2]:[1,0]:e.sortReset?[0,1,2]:[0,1],e.sortVars[r].order=l,e.sortVars[r].count=0),l=e.sortVars[r].order,a=(a=(""+n[1]).match(/^(1|d|s|o|n)/))?a[0]:""){case"1":case"d":a=1;break;case"s":a=i||0;break;case"o":a=0===(o=l[(i||0)%l.length])?1:1===o?0:2;break;case"n":a=l[++e.sortVars[r].count%l.length];break;default:a=0}i=0===s?a:i,n=[r,parseInt(a,10)||0],e.sortList[e.sortList.length]=n,a=R.inArray(n[1],l),e.sortVars[r].count=0<=a?a:n[1]%l.length}},updateAll:function(e,t,r){var a=e.table;a.isUpdating=!0,T.refreshWidgets(a,!0,!0),T.buildHeaders(e),T.bindEvents(a,e.$headers,!0),T.bindMethods(e),T.commonUpdate(e,t,r)},update:function(e,t,r){e.table.isUpdating=!0,T.updateHeader(e),T.commonUpdate(e,t,r)},updateHeaders:function(e,t){e.table.isUpdating=!0,T.buildHeaders(e),T.bindEvents(e.table,e.$headers,!0),T.resortComplete(e,t)},updateCell:function(e,t,r,a){if(R(t).closest("tr").hasClass(e.cssChildRow))console.warn('Tablesorter Warning! "updateCell" for child row content has been disabled, use "update" instead');else if(T.isEmptyObject(e.cache))T.updateHeader(e),T.commonUpdate(e,r,a);else{e.table.isUpdating=!0,e.$table.find(e.selectorRemove).remove();var s,i,o,n,l=e.$tbodies,c=R(t),d=l.index(T.getClosest(c,"tbody")),f=e.cache[d],u=T.getClosest(c,"tr");if(t=c[0],l.length&&0<=d){if(l=l.eq(d).find("tr").not("."+e.cssChildRow).index(u),d=f.normalized[l],(n=u[0].cells.length)!==e.columns)for(s=!1,i=o=0;i<n;i++)s||u[0].cells[i]===t?s=!0:o+=u[0].cells[i].colSpan;else o=c.index();s=T.getElementText(e,t,o),d[e.columns].raw[o]=s,s=T.getParsedText(e,t,o,s),d[o]=s,"numeric"===(e.parsers[o].type||"").toLowerCase()&&(f.colMax[o]=Math.max(Math.abs(s)||0,f.colMax[o]||0)),!1!==(s="undefined"!==r?r:e.resort)?T.checkResort(e,s,a):T.resortComplete(e,a)}else T.debug(e,"core")&&console.error("updateCell aborted, tbody missing or not within the indicated table"),e.table.isUpdating=!1}},addRows:function(e,t,r,a){var s,i,o,n,l,c,d,f,u,g,p,h="string"==typeof t&&1===e.$tbodies.length&&/<tr/.test(t||""),m=e.table;if(h)t=R(t),e.$tbodies.append(t);else if(!(t&&t instanceof R&&T.getClosest(t,"table")[0]===e.table))return T.debug(e,"core")&&console.error("addRows method requires (1) a jQuery selector reference to rows that have already been added to the table, or (2) row HTML string to be added to a table with only one tbody"),!1;if(m.isUpdating=!0,T.isEmptyObject(e.cache))T.updateHeader(e),T.commonUpdate(e,r,a);else{for(o=t.filter("tr").attr("role","row").length,s=e.$tbodies.index(t.parents("tbody").filter(":first")),e.parsers&&e.parsers.length||T.setupParsers(e),i=0;i<o;i++){for(d=0,l=t[i].cells.length,c=e.cache[s].normalized.length,u=[],f={child:[],raw:[],$row:t.eq(i),order:c},n=0;n<l;n++)g=t[i].cells[n],p=T.getElementText(e,g,d),f.raw[d]=p,p=T.getParsedText(e,g,d,p),u[d]=p,"numeric"===(e.parsers[d].type||"").toLowerCase()&&(e.cache[s].colMax[d]=Math.max(Math.abs(p)||0,e.cache[s].colMax[d]||0)),0<(p=g.colSpan-1)&&(d+=p),d++;u[e.columns]=f,e.cache[s].normalized[c]=u}T.checkResort(e,r,a)}},updateCache:function(e,t,r){e.parsers&&e.parsers.length||T.setupParsers(e,r),T.buildCache(e,t,r)},appendCache:function(e,t){var r,a,s,i,o,n,l,c=e.table,d=e.$tbodies,f=[],u=e.cache;if(T.isEmptyObject(u))return e.appender?e.appender(c,f):c.isUpdating?e.$table.triggerHandler("updateComplete",c):"";for(T.debug(e,"core")&&(l=new Date),n=0;n<d.length;n++)if((s=d.eq(n)).length){for(i=T.processTbody(c,s,!0),a=(r=u[n].normalized).length,o=0;o<a;o++)f[f.length]=r[o][e.columns].$row,e.appender&&(!e.pager||e.pager.removeRows||e.pager.ajax)||i.append(r[o][e.columns].$row);T.processTbody(c,i,!1)}e.appender&&e.appender(c,f),T.debug(e,"core")&&console.log("Rebuilt table"+T.benchmark(l)),t||e.appender||T.applyWidget(c),c.isUpdating&&e.$table.triggerHandler("updateComplete",c)},commonUpdate:function(e,t,r){e.$table.find(e.selectorRemove).remove(),T.setupParsers(e),T.buildCache(e),T.checkResort(e,t,r)},initSort:function(t,e,r){if(t.table.isUpdating)return setTimeout(function(){T.initSort(t,e,r)},50);var a,s,i,o,n,l,c,d=!r[t.sortMultiSortKey],f=t.table,u=t.$headers.length,g=T.getClosest(R(e),"th, td"),p=parseInt(g.attr("data-column"),10),h="mouseup"===r.type?"user":r.type,m=t.sortVars[p].order,g=g[0];if(t.$table.triggerHandler("sortStart",f),l=(t.sortVars[p].count+1)%m.length,t.sortVars[p].count=r[t.sortResetKey]?2:l,t.sortRestart)for(i=0;i<u;i++)c=t.$headers.eq(i),p!==(l=parseInt(c.attr("data-column"),10))&&(d||c.hasClass(T.css.sortNone))&&(t.sortVars[l].count=-1);if(d){if(R.each(t.sortVars,function(e){t.sortVars[e].sortedBy=""}),t.sortList=[],t.last.sortList=[],null!==t.sortForce)for(a=t.sortForce,s=0;s<a.length;s++)a[s][0]!==p&&(t.sortList[t.sortList.length]=a[s],t.sortVars[a[s][0]].sortedBy="sortForce");if((o=m[t.sortVars[p].count])<2&&(t.sortList[t.sortList.length]=[p,o],t.sortVars[p].sortedBy=h,1<g.colSpan))for(s=1;s<g.colSpan;s++)t.sortList[t.sortList.length]=[p+s,o],t.sortVars[p+s].count=R.inArray(o,m),t.sortVars[p+s].sortedBy=h}else if(t.sortList=R.extend([],t.last.sortList),0<=T.isValueInArray(p,t.sortList))for(t.sortVars[p].sortedBy=h,s=0;s<t.sortList.length;s++)(l=t.sortList[s])[0]===p&&(l[1]=m[t.sortVars[p].count],2===l[1])&&(t.sortList.splice(s,1),t.sortVars[p].count=-1);else if(o=m[t.sortVars[p].count],t.sortVars[p].sortedBy=h,o<2&&(t.sortList[t.sortList.length]=[p,o],1<g.colSpan))for(s=1;s<g.colSpan;s++)t.sortList[t.sortList.length]=[p+s,o],t.sortVars[p+s].count=R.inArray(o,m),t.sortVars[p+s].sortedBy=h;if(t.last.sortList=R.extend([],t.sortList),t.sortList.length&&t.sortAppend&&(a=R.isArray(t.sortAppend)?t.sortAppend:t.sortAppend[t.sortList[0][0]],!T.isEmptyObject(a)))for(s=0;s<a.length;s++)if(a[s][0]!==p&&T.isValueInArray(a[s][0],t.sortList)<0){if(n=(""+(o=a[s][1])).match(/^(a|d|s|o|n)/))switch(l=t.sortList[0][1],n[0]){case"d":o=1;break;case"s":o=l;break;case"o":o=0===l?1:0;break;case"n":o=(l+1)%m.length;break;default:o=0}t.sortList[t.sortList.length]=[a[s][0],o],t.sortVars[a[s][0]].sortedBy="sortAppend"}t.$table.triggerHandler("sortBegin",f),setTimeout(function(){T.setHeadersCss(t),T.multisort(t),T.appendCache(t),t.$table.triggerHandler("sortBeforeEnd",f),t.$table.triggerHandler("sortEnd",f)},1)},multisort:function(o){var e,t,n,r,l=o.table,c=[],d=0,f=o.textSorter||"",u=o.sortList,g=u.length,a=o.$tbodies.length;if(!o.serverSideSorting&&!T.isEmptyObject(o.cache)){if(T.debug(o,"core")&&(t=new Date),"object"==typeof f)for(n=o.columns;n--;)"function"==typeof(r=T.getColumnData(l,f,n))&&(c[n]=r);for(e=0;e<a;e++)n=o.cache[e].colMax,o.cache[e].normalized.sort(function(e,t){for(var r,a,s,i=0;i<g;i++){if(r=u[i][0],a=u[i][1],d=0===a,o.sortStable&&e[r]===t[r]&&1===g)return e[o.columns].order-t[o.columns].order;if(a=(a=/n/i.test(T.getSortType(o.parsers,r)))&&o.strings[r]?(a="boolean"==typeof T.string[o.strings[r]]?(d?1:-1)*(T.string[o.strings[r]]?-1:1):o.strings[r]&&T.string[o.strings[r]]||0,o.numberSorter?o.numberSorter(e[r],t[r],d,n[r],l):T["sortNumeric"+(d?"Asc":"Desc")](e[r],t[r],a,n[r],r,o)):(a=d?e:t,s=d?t:e,"function"==typeof f?f(a[r],s[r],d,r,l):"function"==typeof c[r]?c[r](a[r],s[r],d,r,l):T["sortNatural"+(d?"Asc":"Desc")](e[r]||"",t[r]||"",r,o)))return a}return e[o.columns].order-t[o.columns].order});T.debug(o,"core")&&console.log("Applying sort "+u.toString()+T.benchmark(t))}},resortComplete:function(e,t){e.table.isUpdating&&e.$table.triggerHandler("updateComplete",e.table),R.isFunction(t)&&t(e.table)},checkResort:function(e,t,r){var a=R.isArray(t)?t:e.sortList;!1===(void 0===t?e.resort:t)||e.serverSideSorting||e.table.isProcessing?(T.resortComplete(e,r),T.applyWidget(e.table,!1)):a.length?T.sortOn(e,a,function(){T.resortComplete(e,r)},!0):T.sortReset(e,function(){T.resortComplete(e,r),T.applyWidget(e.table,!1)})},sortOn:function(e,t,r,a){var s,i=e.table;for(e.$table.triggerHandler("sortStart",i),s=0;s<e.columns;s++)e.sortVars[s].sortedBy=-1<T.isValueInArray(s,t)?"sorton":"";T.updateHeaderSortCount(e,t),T.setHeadersCss(e),e.delayInit&&T.isEmptyObject(e.cache)&&T.buildCache(e),e.$table.triggerHandler("sortBegin",i),T.multisort(e),T.appendCache(e,a),e.$table.triggerHandler("sortBeforeEnd",i),e.$table.triggerHandler("sortEnd",i),T.applyWidget(i),R.isFunction(r)&&r(i)},sortReset:function(e,t){var r;for(e.sortList=[],r=0;r<e.columns;r++)e.sortVars[r].count=-1,e.sortVars[r].sortedBy="";T.setHeadersCss(e),T.multisort(e),T.appendCache(e),R.isFunction(t)&&t(e.table)},getSortType:function(e,t){return e&&e[t]&&e[t].type||""},getOrder:function(e){return/^d/i.test(e)||1===e},sortNatural:function(e,t){if(e!==t){e=(e||"").toString(),t=(t||"").toString();var r,a,s,i,o,n,l=T.regex;if(l.hex.test(t)){if((r=parseInt(e.match(l.hex),16))<(a=parseInt(t.match(l.hex),16)))return-1;if(a<r)return 1}for(r=e.replace(l.chunk,"\\0$1\\0").replace(l.chunks,"").split("\\0"),a=t.replace(l.chunk,"\\0$1\\0").replace(l.chunks,"").split("\\0"),n=Math.max(r.length,a.length),o=0;o<n;o++){if(s=isNaN(r[o])?r[o]||0:parseFloat(r[o])||0,i=isNaN(a[o])?a[o]||0:parseFloat(a[o])||0,isNaN(s)!==isNaN(i))return isNaN(s)?1:-1;if(typeof s!=typeof i&&(s+="",i+=""),s<i)return-1;if(i<s)return 1}}return 0},sortNaturalAsc:function(e,t,r,a){return e===t?0:(r=T.string[a.empties[r]||a.emptyTo],""===e&&0!==r?"boolean"==typeof r?r?-1:1:-r||-1:""===t&&0!==r?"boolean"==typeof r?r?1:-1:r||1:T.sortNatural(e,t))},sortNaturalDesc:function(e,t,r,a){return e===t?0:(r=T.string[a.empties[r]||a.emptyTo],""===e&&0!==r?"boolean"==typeof r?r?-1:1:r||1:""===t&&0!==r?"boolean"==typeof r?r?1:-1:-r||-1:T.sortNatural(t,e))},sortText:function(e,t){return t<e?1:e<t?-1:0},getTextValue:function(e,t,r){if(r){for(var a=e?e.length:0,s=r+t,i=0;i<a;i++)s+=e.charCodeAt(i);return t*s}return 0},sortNumericAsc:function(e,t,r,a,s,i){return e===t?0:(s=T.string[i.empties[s]||i.emptyTo],""===e&&0!==s?"boolean"==typeof s?s?-1:1:-s||-1:""===t&&0!==s?"boolean"==typeof s?s?1:-1:s||1:(e=isNaN(e)?T.getTextValue(e,r,a):e)-(t=isNaN(t)?T.getTextValue(t,r,a):t))},sortNumericDesc:function(e,t,r,a,s,i){return e===t?0:(s=T.string[i.empties[s]||i.emptyTo],""===e&&0!==s?"boolean"==typeof s?s?-1:1:s||1:""===t&&0!==s?"boolean"==typeof s?s?1:-1:-s||-1:(isNaN(e)&&(e=T.getTextValue(e,r,a)),(t=isNaN(t)?T.getTextValue(t,r,a):t)-e))},sortNumeric:function(e,t){return e-t},addWidget:function(e){e.id&&!T.isEmptyObject(T.getWidgetById(e.id))&&console.warn('"'+e.id+'" widget was loaded more than once!'),T.widgets[T.widgets.length]=e},hasWidget:function(e,t){return(e=R(e)).length&&e[0].config&&e[0].config.widgetInit[t]||!1},getWidgetById:function(e){for(var t,r=T.widgets.length,a=0;a<r;a++)if((t=T.widgets[a])&&t.id&&t.id.toLowerCase()===e.toLowerCase())return t},applyWidgetOptions:function(e){var t,r,a,s=e.config,i=s.widgets.length;if(i)for(t=0;t<i;t++)(r=T.getWidgetById(s.widgets[t]))&&r.options&&(a=R.extend(!0,{},r.options),s.widgetOptions=R.extend(!0,a,s.widgetOptions),R.extend(!0,T.defaults.widgetOptions,r.options))},addWidgetFromClass:function(e){var t,r,a=e.config,s="^"+a.widgetClass.replace(T.regex.templateName,"(\\S+)+")+"$",i=new RegExp(s,"g"),o=(e.className||"").split(T.regex.spaces);if(o.length)for(t=o.length,r=0;r<t;r++)o[r].match(i)&&(a.widgets[a.widgets.length]=o[r].replace(i,"$1"))},applyWidgetId:function(e,t,r){var a,s,i,o=(e=R(e)[0]).config,n=o.widgetOptions,l=T.debug(o,"core"),t=T.getWidgetById(t);t&&(i=t.id,a=!1,R.inArray(i,o.widgets)<0&&(o.widgets[o.widgets.length]=i),l&&(s=new Date),!r&&o.widgetInit[i]||(o.widgetInit[i]=!0,e.hasInitialized&&T.applyWidgetOptions(e),"function"==typeof t.init&&(a=!0,l&&console[console.group?"group":"log"]("Initializing "+i+" widget"),t.init(e,t,o,n))),r||"function"!=typeof t.format||(a=!0,l&&console[console.group?"group":"log"]("Updating "+i+" widget"),t.format(e,o,n,!1)),l)&&a&&(console.log("Completed "+(r?"initializing ":"applying ")+i+" widget"+T.benchmark(s)),console.groupEnd)&&console.groupEnd()},applyWidget:function(e,t,r){var a,s,i,o,n,l=(e=R(e)[0]).config,c=T.debug(l,"core"),d=[];if(!1===t||!e.hasInitialized||!e.isApplyingWidgets&&!e.isUpdating){if(c&&(n=new Date),T.addWidgetFromClass(e),clearTimeout(l.timerReady),l.widgets.length){for(e.isApplyingWidgets=!0,l.widgets=R.grep(l.widgets,function(e,t){return R.inArray(e,l.widgets)===t}),s=(i=l.widgets||[]).length,a=0;a<s;a++)(o=T.getWidgetById(i[a]))&&o.id?(o.priority||(o.priority=10),d[a]=o):c&&console.warn('"'+i[a]+'" was enabled, but the widget code has not been loaded!');for(d.sort(function(e,t){return e.priority<t.priority?-1:e.priority===t.priority?0:1}),s=d.length,c&&console[console.group?"group":"log"]("Start "+(t?"initializing":"applying")+" widgets"),a=0;a<s;a++)(o=d[a])&&o.id&&T.applyWidgetId(e,o.id,t);c&&console.groupEnd&&console.groupEnd()}l.timerReady=setTimeout(function(){e.isApplyingWidgets=!1,R.data(e,"lastWidgetApplication",new Date),l.$table.triggerHandler("tablesorter-ready"),t||"function"!=typeof r||r(e),c&&(o=l.widgets.length,console.log("Completed "+(!0===t?"initializing ":"applying ")+o+" widget"+(1!==o?"s":"")+T.benchmark(n)))},10)}},removeWidget:function(e,t,r){var a,s,i,o,n=(e=R(e)[0]).config;if(!0===t)for(t=[],o=T.widgets.length,i=0;i<o;i++)(s=T.widgets[i])&&s.id&&(t[t.length]=s.id);else t=(R.isArray(t)?t.join(","):t||"").toLowerCase().split(/[\s,]+/);for(o=t.length,a=0;a<o;a++)s=T.getWidgetById(t[a]),0<=(i=R.inArray(t[a],n.widgets))&&!0!==r&&n.widgets.splice(i,1),s&&s.remove&&(T.debug(n,"core")&&console.log((r?"Refreshing":"Removing")+' "'+t[a]+'" widget'),s.remove(e,n,n.widgetOptions,r),n.widgetInit[t[a]]=!1);n.$table.triggerHandler("widgetRemoveEnd",e)},refreshWidgets:function(e,t,r){for(var a,s=(e=R(e)[0]).config.widgets,i=T.widgets,o=i.length,n=[],l=function(e){R(e).triggerHandler("refreshComplete")},c=0;c<o;c++)(a=i[c])&&a.id&&(t||R.inArray(a.id,s)<0)&&(n[n.length]=a.id);T.removeWidget(e,n.join(","),!0),!0!==r?(T.applyWidget(e,t||!1,l),t&&T.applyWidget(e,!1,l)):l(e)},benchmark:function(e){return" ("+((new Date).getTime()-e.getTime())+" ms)"},log:function(){console.log(arguments)},debug:function(e,t){return e&&(!0===e.debug||"string"==typeof e.debug&&-1<e.debug.indexOf(t))},isEmptyObject:function(e){for(var t in e)return!1;return!0},isValueInArray:function(e,t){for(var r=t&&t.length||0,a=0;a<r;a++)if(t[a][0]===e)return a;return-1},formatFloat:function(e,t){return"string"!=typeof e||""===e?e:(e=(t&&t.config?!1!==t.config.usNumberFormat:void 0===t||t)?e.replace(T.regex.comma,""):e.replace(T.regex.digitNonUS,"").replace(T.regex.comma,"."),T.regex.digitNegativeTest.test(e)&&(e=e.replace(T.regex.digitNegativeReplace,"-$1")),t=parseFloat(e),isNaN(t)?R.trim(e):t)},isDigit:function(e){return isNaN(e)?T.regex.digitTest.test(e.toString().replace(T.regex.digitReplace,"")):""!==e},computeColumnIndex:function(e,t){for(var r,a,s,i,o,n,l,c,d,f=t&&t.columns||0,u=[],g=new Array(f),p=0;p<e.length;p++)for(o=e[p].cells,r=0;r<o.length;r++){for(n=p,l=(i=o[r]).rowSpan||1,c=i.colSpan||1,void 0===u[n]&&(u[n]=[]),a=0;a<u[n].length+1;a++)if(void 0===u[n][a]){d=a;break}for(f&&i.cellIndex===d||(i.setAttribute?i.setAttribute("data-column",d):R(i).attr("data-column",d)),a=n;a<n+l;a++)for(void 0===u[a]&&(u[a]=[]),g=u[a],s=d;s<d+c;s++)g[s]="x"}return T.checkColumnCount(e,u,g.length),g.length},checkColumnCount:function(e,t,r){for(var a,s=!0,i=[],o=0;o<t.length;o++)if(t[o]&&(a=t[o].length,t[o].length!==r)){s=!1;break}s||(e.each(function(e,t){t=t.parentElement.nodeName;i.indexOf(t)<0&&i.push(t)}),console.error("Invalid or incorrect number of columns in the "+i.join(" or ")+"; expected "+r+", but found "+a+" columns"))},fixColumnWidth:function(e){var t,r,a,s,i,e=(e=R(e)[0]).config,o=e.$table.children("colgroup");if(o.length&&o.hasClass(T.css.colgroup)&&o.remove(),e.widthFixed&&0===e.$table.children("colgroup").length){for(o=R('<colgroup class="'+T.css.colgroup+'">'),t=e.$table.width(),s=(a=e.$tbodies.find("tr:first").children(":visible")).length,i=0;i<s;i++)r=parseInt(a.eq(i).width()/t*1e3,10)/10+"%",o.append(R("<col>").css("width",r));e.$table.prepend(o)}},getData:function(e,t,r){var a,s,i="",e=R(e);return e.length?(a=!!R.metadata&&e.metadata(),s=" "+(e.attr("class")||""),void 0!==e.data(r)||void 0!==e.data(r.toLowerCase())?i+=e.data(r)||e.data(r.toLowerCase()):a&&void 0!==a[r]?i+=a[r]:t&&void 0!==t[r]?i+=t[r]:" "!==s&&s.match(" "+r+"-")&&(i=s.match(new RegExp("\\s"+r+"-([\\w-]+)"))[1]||""),R.trim(i)):""},getColumnData:function(e,t,r,a,s){if("object"!=typeof t||null===t)return t;var i,e=(e=R(e)[0]).config,s=s||e.$headers,o=e.$headerIndexed&&e.$headerIndexed[r]||s.find('[data-column="'+r+'"]:last');if(void 0!==t[r])return a?t[r]:t[s.index(o)];for(i in t)if("string"==typeof i&&o.filter(i).add(o.find(i)).length)return t[i]},isProcessing:function(e,t,r){var a=(e=R(e))[0].config,s=r||e.find("."+T.css.header);t?(void 0!==r&&0<a.sortList.length&&(s=s.filter(function(){return!this.sortDisabled&&0<=T.isValueInArray(parseFloat(R(this).attr("data-column")),a.sortList)})),e.add(s).addClass(T.css.processing+" "+a.cssProcessing)):e.add(s).removeClass(T.css.processing+" "+a.cssProcessing)},processTbody:function(e,t,r){if(e=R(e)[0],r)return e.isProcessing=!0,t.before('<colgroup class="tablesorter-savemyplace"/>'),R.fn.detach?t.detach():t.remove();r=R(e).find("colgroup.tablesorter-savemyplace");t.insertAfter(r),r.remove(),e.isProcessing=!1},clearTableBody:function(e){R(e)[0].config.$tbodies.children().detach()},characterEquivalents:{a:"áàâãäąå",A:"ÁÀÂÃÄĄÅ",c:"çćč",C:"ÇĆČ",e:"éèêëěę",E:"ÉÈÊËĚĘ",i:"íìİîïı",I:"ÍÌİÎÏ",o:"óòôõöō",O:"ÓÒÔÕÖŌ",ss:"ß",SS:"ẞ",u:"úùûüů",U:"ÚÙÛÜŮ"},replaceAccents:function(e){var t,r="[",a=T.characterEquivalents;if(!T.characterRegex){for(t in T.characterRegexArray={},a)"string"==typeof t&&(r+=a[t],T.characterRegexArray[t]=new RegExp("["+a[t]+"]","g"));T.characterRegex=new RegExp(r+"]")}if(T.characterRegex.test(e))for(t in a)"string"==typeof t&&(e=e.replace(T.characterRegexArray[t],t));return e},validateOptions:function(e){var t,r,a,s,i="headers sortForce sortList sortAppend widgets".split(" "),o=e.originalSettings;if(o){for(t in T.debug(e,"core")&&(s=new Date),o)if("undefined"===(a=typeof T.defaults[t]))console.warn('Tablesorter Warning! "table.config.'+t+'" option not recognized');else if("object"===a)for(r in o[t])a=T.defaults[t]&&typeof T.defaults[t][r],R.inArray(t,i)<0&&"undefined"===a&&console.warn('Tablesorter Warning! "table.config.'+t+"."+r+'" option not recognized');T.debug(e,"core")&&console.log("validate options time:"+T.benchmark(s))}},restoreHeaders:function(e){for(var t,r=R(e)[0].config,a=r.$table.find(r.selectorHeaders),s=a.length,i=0;i<s;i++)(t=a.eq(i)).find("."+T.css.headerIn).length&&t.html(r.headerContent[i])},destroy:function(e,t,r){var a,s,i,o,n;(e=R(e)[0]).hasInitialized&&(T.removeWidget(e,!0,!1),a=R(e),s=e.config,o=(i=a.find("thead:first")).find("tr."+T.css.headerRow).removeClass(T.css.headerRow+" "+s.cssHeaderRow),n=a.find("tfoot:first > tr").children("th, td"),!1===t&&0<=R.inArray("uitheme",s.widgets)&&(a.triggerHandler("applyWidgetId",["uitheme"]),a.triggerHandler("applyWidgetId",["zebra"])),i.find("tr").not(o).remove(),i="sortReset update updateRows updateAll updateHeaders updateCell addRows updateComplete sorton appendCache updateCache applyWidgetId applyWidgets refreshWidgets removeWidget destroy mouseup mouseleave "+"keypress sortBegin sortEnd resetToLoadState ".split(" ").join(s.namespace+" "),a.removeData("tablesorter").unbind(i.replace(T.regex.spaces," ")),s.$headers.add(n).removeClass([T.css.header,s.cssHeader,s.cssAsc,s.cssDesc,T.css.sortAsc,T.css.sortDesc,T.css.sortNone].join(" ")).removeAttr("data-column").removeAttr("aria-label").attr("aria-disabled","true"),o.find(s.selectorSort).unbind("mousedown mouseup keypress ".split(" ").join(s.namespace+" ").replace(T.regex.spaces," ")),T.restoreHeaders(e),a.toggleClass(T.css.table+" "+s.tableClass+" tablesorter-"+s.theme,!1===t),a.removeClass(s.namespace.slice(1)),e.hasInitialized=!1,delete e.config.cache,"function"==typeof r&&r(e),T.debug(s,"core"))&&console.log("tablesorter has been removed")}};R.fn.tablesorter=function(t){return this.each(function(){var e=R.extend(!0,{},T.defaults,t,T.instanceMethods);e.originalSettings=t,!this.hasInitialized&&T.buildTable&&"TABLE"!==this.nodeName?T.buildTable(this,e):T.setup(this,e)})},window.console&&window.console.log||(T.logs=[],(console={}).log=console.warn=console.error=console.table=function(){var e=1<arguments.length?arguments:arguments[0];T.logs[T.logs.length]={date:Date.now(),log:e}}),T.addParser({id:"no-parser",is:function(){return!1},format:function(){return""},type:"text"}),T.addParser({id:"text",is:function(){return!0},format:function(e,t){t=t.config;return e&&(e=R.trim(t.ignoreCase?e.toLocaleLowerCase():e),e=t.sortLocaleCompare?T.replaceAccents(e):e),e},type:"text"}),T.regex.nondigit=/[^\w,. \-()]/g,T.addParser({id:"digit",is:function(e){return T.isDigit(e)},format:function(e,t){var r=T.formatFloat((e||"").replace(T.regex.nondigit,""),t);return e&&"number"==typeof r?r:e&&R.trim(e&&t.config.ignoreCase?e.toLocaleLowerCase():e)},type:"numeric"}),T.regex.currencyReplace=/[+\-,. ]/g,T.regex.currencyTest=/^\(?\d+[\u00a3$\u20ac\u00a4\u00a5\u00a2?.]|[\u00a3$\u20ac\u00a4\u00a5\u00a2?.]\d+\)?$/,T.addParser({id:"currency",is:function(e){return e=(e||"").replace(T.regex.currencyReplace,""),T.regex.currencyTest.test(e)},format:function(e,t){var r=T.formatFloat((e||"").replace(T.regex.nondigit,""),t);return e&&"number"==typeof r?r:e&&R.trim(e&&t.config.ignoreCase?e.toLocaleLowerCase():e)},type:"numeric"}),T.regex.urlProtocolTest=/^(https?|ftp|file):\/\//,T.regex.urlProtocolReplace=/(https?|ftp|file):\/\/(www\.)?/,T.addParser({id:"url",is:function(e){return T.regex.urlProtocolTest.test(e)},format:function(e){return e&&R.trim(e.replace(T.regex.urlProtocolReplace,""))},type:"text"}),T.regex.dash=/-/g,T.regex.isoDate=/^\d{4}[\/\-]\d{1,2}[\/\-]\d{1,2}/,T.addParser({id:"isoDate",is:function(e){return T.regex.isoDate.test(e)},format:function(e){var t=e&&new Date(e.replace(T.regex.dash,"/"));return t instanceof Date&&isFinite(t)?t.getTime():e},type:"numeric"}),T.regex.percent=/%/g,T.regex.percentTest=/(\d\s*?%|%\s*?\d)/,T.addParser({id:"percent",is:function(e){return T.regex.percentTest.test(e)&&e.length<15},format:function(e,t){return e&&T.formatFloat(e.replace(T.regex.percent,""),t)},type:"numeric"}),T.addParser({id:"image",is:function(e,t,r,a){return 0<a.find("img").length},format:function(e,t,r){return R(r).find("img").attr(t.config.imgAttr||"alt")||e},parsed:!0,type:"text"}),T.regex.dateReplace=/(\S)([AP]M)$/i,T.regex.usLongDateTest1=/^[A-Z]{3,10}\.?\s+\d{1,2},?\s+(\d{4})(\s+\d{1,2}:\d{2}(:\d{2})?(\s+[AP]M)?)?$/i,T.regex.usLongDateTest2=/^\d{1,2}\s+[A-Z]{3,10}\s+\d{4}/i,T.addParser({id:"usLongDate",is:function(e){return T.regex.usLongDateTest1.test(e)||T.regex.usLongDateTest2.test(e)},format:function(e){var t=e&&new Date(e.replace(T.regex.dateReplace,"$1 $2"));return t instanceof Date&&isFinite(t)?t.getTime():e},type:"numeric"}),T.regex.shortDateTest=/(^\d{1,2}[\/\s]\d{1,2}[\/\s]\d{4})|(^\d{4}[\/\s]\d{1,2}[\/\s]\d{1,2})/,T.regex.shortDateReplace=/[\-.,]/g,T.regex.shortDateXXY=/(\d{1,2})[\/\s](\d{1,2})[\/\s](\d{4})/,T.regex.shortDateYMD=/(\d{4})[\/\s](\d{1,2})[\/\s](\d{1,2})/,T.convertFormat=function(e,t){e=(e||"").replace(T.regex.spaces," ").replace(T.regex.shortDateReplace,"/"),"mmddyyyy"===t?e=e.replace(T.regex.shortDateXXY,"$3/$1/$2"):"ddmmyyyy"===t?e=e.replace(T.regex.shortDateXXY,"$3/$2/$1"):"yyyymmdd"===t&&(e=e.replace(T.regex.shortDateYMD,"$1/$2/$3"));t=new Date(e);return t instanceof Date&&isFinite(t)?t.getTime():""},T.addParser({id:"shortDate",is:function(e){return e=(e||"").replace(T.regex.spaces," ").replace(T.regex.shortDateReplace,"/"),T.regex.shortDateTest.test(e)},format:function(e,t,r,a){var s,i;return e&&(t=(i=(s=t.config).$headerIndexed[a]).length&&i.data("dateFormat")||T.getData(i,T.getColumnData(t,s.headers,a),"dateFormat")||s.dateFormat,i.length&&i.data("dateFormat",t),T.convertFormat(e,t)||e)},type:"numeric"}),T.regex.timeTest=/^(0?[1-9]|1[0-2]):([0-5]\d)(\s[AP]M)$|^((?:[01]\d|[2][0-4]):[0-5]\d)$/i,T.regex.timeMatch=/(0?[1-9]|1[0-2]):([0-5]\d)(\s[AP]M)|((?:[01]\d|[2][0-4]):[0-5]\d)/i,T.addParser({id:"time",is:function(e){return T.regex.timeTest.test(e)},format:function(e){var t=(e||"").match(T.regex.timeMatch),r=new Date(e),t=e&&(null!==t?t[0]:"00:00 AM"),t=t&&new Date("2000/01/01 "+t.replace(T.regex.dateReplace,"$1 $2"));return t instanceof Date&&isFinite(t)?(r instanceof Date&&isFinite(r)?r.getTime():0)?parseFloat(t.getTime()+"."+r.getTime()):t.getTime():e},type:"numeric"}),T.addParser({id:"metadata",is:function(){return!1},format:function(e,t,r){t=t.config,t=t.parserMetadataName||"sortValue";return R(r).metadata()[t]},type:"numeric"}),T.addWidget({id:"zebra",priority:90,format:function(e,t,r){for(var a,s,i,o,n,l=new RegExp(t.cssChildRow,"i"),c=t.$tbodies.add(R(t.namespace+"_extra_table").children("tbody:not(."+t.cssInfoBlock+")")),d=0;d<c.length;d++)for(s=0,n=(a=c.eq(d).children("tr:visible").not(t.selectorRemove)).length,o=0;o<n;o++)i=a.eq(o),l.test(i[0].className)||s++,i.removeClass(r.zebra[(i=s%2==0)?1:0]).addClass(r.zebra[i?0:1])},remove:function(e,t,r,a){if(!a)for(var s,i=t.$tbodies,o=(r.zebra||["even","odd"]).join(" "),n=0;n<i.length;n++)(s=T.processTbody(e,i.eq(n),!0)).children().removeClass(o),T.processTbody(e,s,!1)}})}(e),function(u,g,p){"use strict";var h=u.tablesorter||{};u.extend(!0,h.defaults,{fixedUrl:"",widgetOptions:{storage_fixedUrl:"",storage_group:"",storage_page:"",storage_storageType:"",storage_tableId:"",storage_useSessionStorage:""}}),h.storage=function(e,t,r,a){var s=!1,i={},o=(e=u(e)[0]).config,n=o&&o.widgetOptions,l=h.debug(o,"storage"),c=(a&&a.storageType||n&&n.storage_storageType).toString().charAt(0).toLowerCase(),d=c?"":a&&a.useSessionStorage||n&&n.storage_useSessionStorage,f=u(e),e=a&&a.id||f.attr(a&&a.group||n&&n.storage_group||"data-table-group")||n&&n.storage_tableId||e.id||u(".tablesorter").index(f),f=a&&a.url||f.attr(a&&a.page||n&&n.storage_page||"data-table-page")||n&&n.storage_fixedUrl||o&&o.fixedUrl||g.location.pathname;if("c"!==c&&(c="s"===c||d?"sessionStorage":"localStorage")in g)try{g[c].setItem("_tmptest","temp"),s=!0,g[c].removeItem("_tmptest")}catch(e){console.warn(c+" is not supported in this browser")}if(l&&console.log("Storage >> Using",s?c:"cookies"),u.parseJSON&&(i=s?u.parseJSON(g[c][t]||"null")||{}:(a=p.cookie.split(/[;\s|=]/),0!==(n=u.inArray(t,a)+1)&&u.parseJSON(a[n]||"null")||{})),void 0===r||!g.JSON||!JSON.hasOwnProperty("stringify"))return i&&i[f]?i[f][e]:"";i[f]||(i[f]={}),i[f][e]=r,s?g[c][t]=JSON.stringify(i):((o=new Date).setTime(o.getTime()+31536e6),p.cookie=t+"="+JSON.stringify(i).replace(/\"/g,'"')+"; expires="+o.toGMTString()+"; path=/")}}(e,window,document),function(_){"use strict";var $=_.tablesorter||{};$.themes={bootstrap:{table:"table table-bordered table-striped",caption:"caption",header:"bootstrap-header",sortNone:"",sortAsc:"",sortDesc:"",active:"",hover:"",icons:"",iconSortNone:"bootstrap-icon-unsorted",iconSortAsc:"glyphicon glyphicon-chevron-up",iconSortDesc:"glyphicon glyphicon-chevron-down",filterRow:"",footerRow:"",footerCells:"",even:"",odd:""},jui:{table:"ui-widget ui-widget-content ui-corner-all",caption:"ui-widget-content",header:"ui-widget-header ui-corner-all ui-state-default",sortNone:"",sortAsc:"",sortDesc:"",active:"ui-state-active",hover:"ui-state-hover",icons:"ui-icon",iconSortNone:"ui-icon-carat-2-n-s ui-icon-caret-2-n-s",iconSortAsc:"ui-icon-carat-1-n ui-icon-caret-1-n",iconSortDesc:"ui-icon-carat-1-s ui-icon-caret-1-s",filterRow:"",footerRow:"",footerCells:"",even:"ui-widget-content",odd:"ui-state-default"}},_.extend($.css,{wrapper:"tablesorter-wrapper"}),$.addWidget({id:"uitheme",priority:10,format:function(e,t,r){var a,s,i,o,n,l,c,d,f,u,g,p,h=$.themes,m=t.$table.add(_(t.namespace+"_extra_table")),b=t.$headers.add(_(t.namespace+"_extra_headers")),y=t.theme||"jui",w=h[y]||{},v=_.trim([w.sortNone,w.sortDesc,w.sortAsc,w.active].join(" ")),x=_.trim([w.iconSortNone,w.iconSortDesc,w.iconSortAsc].join(" ")),C=$.debug(t,"uitheme");for(C&&(n=new Date),m.hasClass("tablesorter-"+y)&&t.theme===t.appliedTheme&&r.uitheme_applied||(r.uitheme_applied=!0,u=h[t.appliedTheme]||{},h=(p=!_.isEmptyObject(u))?[u.sortNone,u.sortDesc,u.sortAsc,u.active].join(" "):"",g=p?[u.iconSortNone,u.iconSortDesc,u.iconSortAsc].join(" "):"",p&&(r.zebra[0]=_.trim(" "+r.zebra[0].replace(" "+u.even,"")),r.zebra[1]=_.trim(" "+r.zebra[1].replace(" "+u.odd,"")),t.$tbodies.children().removeClass([u.even,u.odd].join(" "))),w.even&&(r.zebra[0]+=" "+w.even),w.odd&&(r.zebra[1]+=" "+w.odd),m.children("caption").removeClass(u.caption||"").addClass(w.caption),d=m.removeClass((t.appliedTheme?"tablesorter-"+(t.appliedTheme||""):"")+" "+(u.table||"")).addClass("tablesorter-"+y+" "+(w.table||"")).children("tfoot"),t.appliedTheme=t.theme,d.length&&d.children("tr").removeClass(u.footerRow||"").addClass(w.footerRow).children("th, td").removeClass(u.footerCells||"").addClass(w.footerCells),b.removeClass((p?[u.header,u.hover,h].join(" "):"")||"").addClass(w.header).not(".sorter-false").unbind("mouseenter.tsuitheme mouseleave.tsuitheme").bind("mouseenter.tsuitheme mouseleave.tsuitheme",function(e){_(this)["mouseenter"===e.type?"addClass":"removeClass"](w.hover||"")}),b.each(function(){var e=_(this);e.find("."+$.css.wrapper).length||e.wrapInner('<div class="'+$.css.wrapper+'" style="position:relative;height:100%;width:100%"></div>')}),t.cssIcon&&b.find("."+$.css.icon).removeClass(p?[u.icons,g].join(" "):"").addClass(w.icons||""),$.hasWidget(t.table,"filter")&&(s=function(){m.children("thead").children("."+$.css.filterRow).removeClass(p&&u.filterRow||"").addClass(w.filterRow||"")},r.filter_initialized?s():m.one("filterInit",function(){s()}))),a=0;a<t.columns;a++)l=t.$headers.add(_(t.namespace+"_extra_headers")).not(".sorter-false").filter('[data-column="'+a+'"]'),c=$.css.icon?l.find("."+$.css.icon):_(),(f=b.not(".sorter-false").filter('[data-column="'+a+'"]:last')).length&&(l.removeClass(v),c.removeClass(x),f[0].sortDisabled?c.removeClass(w.icons||""):(i=w.sortNone,o=w.iconSortNone,f.hasClass($.css.sortAsc)?(i=[w.sortAsc,w.active].join(" "),o=w.iconSortAsc):f.hasClass($.css.sortDesc)&&(i=[w.sortDesc,w.active].join(" "),o=w.iconSortDesc),l.addClass(i),c.addClass(o||"")));C&&console.log("uitheme >> Applied "+y+" theme"+$.benchmark(n))},remove:function(e,t,r,a){var s,i,o,n,l;r.uitheme_applied&&(s=t.$table,t=t.appliedTheme||"jui",i=$.themes[t]||$.themes.jui,o=s.children("thead").children(),n=i.sortNone+" "+i.sortDesc+" "+i.sortAsc,l=i.iconSortNone+" "+i.iconSortDesc+" "+i.iconSortAsc,s.removeClass("tablesorter-"+t+" "+i.table),r.uitheme_applied=!1,a||(s.find($.css.header).removeClass(i.header),o.unbind("mouseenter.tsuitheme mouseleave.tsuitheme").removeClass(i.hover+" "+n+" "+i.active).filter("."+$.css.filterRow).removeClass(i.filterRow),o.find("."+$.css.icon).removeClass(i.icons+" "+l)))}})}(e),function(m){"use strict";var b=m.tablesorter||{};b.addWidget({id:"columns",priority:65,options:{columns:["primary","secondary","tertiary"]},format:function(e,t,r){for(var a,s,i,o,n,l=t.$table,c=t.$tbodies,d=t.sortList,f=d.length,u=r&&r.columns||["primary","secondary","tertiary"],g=u.length-1,p=u.join(" "),h=0;h<c.length;h++)(s=(a=b.processTbody(e,c.eq(h),!0)).children("tr")).each(function(){if(i=m(this),"none"!==this.style.display&&(o=i.children().removeClass(p),d)&&d[0]&&(o.eq(d[0][0]).addClass(u[0]),1<f))for(n=1;n<f;n++)o.eq(d[n][0]).addClass(u[n]||u[g])}),b.processTbody(e,a,!1);if(t=!1!==r.columns_thead?["thead tr"]:[],!1!==r.columns_tfoot&&t.push("tfoot tr"),t.length&&(s=l.find(t.join(",")).children().removeClass(p),f))for(n=0;n<f;n++)s.filter('[data-column="'+d[n][0]+'"]').addClass(u[n]||u[g])},remove:function(e,t,r){var a,s,i=t.$tbodies,o=(r.columns||["primary","secondary","tertiary"]).join(" ");for(t.$headers.removeClass(o),t.$table.children("tfoot").children("tr").children("th, td").removeClass(o),a=0;a<i.length;a++)(s=b.processTbody(e,i.eq(a),!0)).children("tr").each(function(){m(this).children().removeClass(o)}),b.processTbody(e,s,!1)}})}(e),function(A){"use strict";var D,H,N=A.tablesorter||{},h=N.css,l=N.keyCodes;A.extend(h,{filterRow:"tablesorter-filter-row",filter:"tablesorter-filter",filterDisabled:"disabled",filterRowHide:"hideme"}),A.extend(l,{backSpace:8,escape:27,space:32,left:37,down:40}),N.addWidget({id:"filter",priority:50,options:{filter_cellFilter:"",filter_childRows:!1,filter_childByColumn:!1,filter_childWithSibs:!0,filter_columnAnyMatch:!0,filter_columnFilters:!0,filter_cssFilter:"",filter_defaultAttrib:"data-value",filter_defaultFilter:{},filter_excludeFilter:{},filter_external:"",filter_filteredRow:"filtered",filter_filterLabel:'Filter "{{label}}" column by...',filter_formatter:null,filter_functions:null,filter_hideEmpty:!0,filter_hideFilters:!1,filter_ignoreCase:!0,filter_liveSearch:!0,filter_matchType:{input:"exact",select:"exact"},filter_onlyAvail:"filter-onlyAvail",filter_placeholder:{search:"",select:""},filter_reset:null,filter_resetOnEsc:!0,filter_saveFilters:!1,filter_searchDelay:300,filter_searchFiltered:!0,filter_selectSource:null,filter_selectSourceSeparator:"|",filter_serversideFiltering:!1,filter_startsWith:!1,filter_useParsedData:!1},format:function(e,t,r){t.$table.hasClass("hasFilters")||D.init(e,t,r)},remove:function(e,t,r,a){var s,i,o=t.$table,n=t.$tbodies,l="addRows updateCell update updateRows updateComplete appendCache filterReset filterAndSortReset filterFomatterUpdate filterEnd search stickyHeadersInit ".split(" ").join(t.namespace+"filter ");if(o.removeClass("hasFilters").unbind(l.replace(N.regex.spaces," ")).find("."+h.filterRow).remove(),r.filter_initialized=!1,!a){for(s=0;s<n.length;s++)(i=N.processTbody(e,n.eq(s),!0)).children().removeClass(r.filter_filteredRow).show(),N.processTbody(e,i,!1);r.filter_reset&&A(document).undelegate(r.filter_reset,"click"+t.namespace+"filter")}}}),D=N.filter={regex:{regex:/^\/((?:\\\/|[^\/])+)\/([migyu]{0,5})?$/,child:/tablesorter-childRow/,filtered:/filtered/,type:/undefined|number/,exact:/(^[\"\'=]+)|([\"\'=]+$)/g,operators:/[<>=]/g,query:"(q|query)",wild01:/\?/g,wild0More:/\*/g,quote:/\"/g,isNeg1:/(>=?\s*-\d)/,isNeg2:/(<=?\s*\d)/},types:{or:function(e,t,r){if(!H.orTest.test(t.iFilter)&&!H.orSplit.test(t.filter)||H.regex.test(t.filter))return null;for(var a,s,i=A.extend({},t),o=t.filter.split(H.orSplit),n=t.iFilter.split(H.orSplit),l=o.length,c=0;c<l;c++){i.nestedFilters=!0,i.filter=""+(D.parseFilter(e,o[c],t)||""),i.iFilter=""+(D.parseFilter(e,n[c],t)||""),s="("+(D.parseFilter(e,i.filter,t)||"")+")";try{if(a=new RegExp(t.isMatch?s:"^"+s+"$",e.widgetOptions.filter_ignoreCase?"i":"").test(i.exact)||D.processTypes(e,i,r))return a}catch(e){return null}}return a||!1},and:function(e,t,r){if(H.andTest.test(t.filter)){for(var a,s,i,o=A.extend({},t),n=t.filter.split(H.andSplit),l=t.iFilter.split(H.andSplit),c=n.length,d=0;d<c;d++){o.nestedFilters=!0,o.filter=""+(D.parseFilter(e,n[d],t)||""),o.iFilter=""+(D.parseFilter(e,l[d],t)||""),i=("("+(D.parseFilter(e,o.filter,t)||"")+")").replace(H.wild01,"\\S{1}").replace(H.wild0More,"\\S*");try{s=new RegExp(t.isMatch?i:"^"+i+"$",e.widgetOptions.filter_ignoreCase?"i":"").test(o.exact)||D.processTypes(e,o,r),a=(0===d||a)&&s}catch(e){return null}}return a||!1}return null},regex:function(e,t){if(H.regex.test(t.filter)){var r,a=t.filter_regexCache[t.index]||H.regex.exec(t.filter),s=a instanceof RegExp;try{s||(t.filter_regexCache[t.index]=a=new RegExp(a[1],a[2])),r=a.test(t.exact)}catch(e){r=!1}return r}return null},operators:function(e,t){var r,a,s,i,o,n,l;return H.operTest.test(t.iFilter)&&""!==t.iExact?(s=e.table,i=t.parsed[t.index],o=N.formatFloat(t.iFilter.replace(H.operators,""),s),n=e.parsers[t.index]||{},l=o,!i&&"numeric"!==n.type||(a=A.trim(""+t.iFilter.replace(H.operators,"")),o="number"!=typeof(r=D.parseFilter(e,a,t,!0))||""===r||isNaN(r)?o:r),e=!i&&"numeric"!==n.type||isNaN(o)||void 0===t.cache?(a=isNaN(t.iExact)?t.iExact.replace(N.regex.nondigit,""):t.iExact,N.formatFloat(a,s)):t.cache,H.gtTest.test(t.iFilter)?r=H.gteTest.test(t.iFilter)?o<=e:o<e:H.ltTest.test(t.iFilter)&&(r=H.lteTest.test(t.iFilter)?e<=o:e<o),!r&&""===l||r):null},notMatch:function(e,t){var r,a;return H.notTest.test(t.iFilter)?(a=t.iFilter.replace("!",""),a=D.parseFilter(e,a,t)||"",H.exact.test(a)?""===(a=a.replace(H.exact,""))||A.trim(a)!==t.iExact:(r=t.iExact.search(A.trim(a)),""===a||(t.anyMatch?r<0:!(e.widgetOptions.filter_startsWith?0===r:0<=r)))):null},exact:function(e,t){var r;return H.exact.test(t.iFilter)?(r=t.iFilter.replace(H.exact,""),e=D.parseFilter(e,r,t)||"",t.anyMatch?0<=A.inArray(e,t.rowArray):e==t.iExact):null},range:function(e,t){var r,a,s,i,o,n,l,c;return H.toTest.test(t.iFilter)?(s=e.table,i=t.index,o=t.parsed[i],l=(n=t.iFilter.split(H.toSplit))[0].replace(N.regex.nondigit,"")||"",c=N.formatFloat(D.parseFilter(e,l,t),s),l=n[1].replace(N.regex.nondigit,"")||"",a=N.formatFloat(D.parseFilter(e,l,t),s),!o&&"numeric"!==e.parsers[i].type||(c=""===(r=e.parsers[i].format(""+n[0],s,e.$headers.eq(i),i))||isNaN(r)?c:r,a=""===(r=e.parsers[i].format(""+n[1],s,e.$headers.eq(i),i))||isNaN(r)?a:r),r=!o&&"numeric"!==e.parsers[i].type||isNaN(c)||isNaN(a)?(l=isNaN(t.iExact)?t.iExact.replace(N.regex.nondigit,""):t.iExact,N.formatFloat(l,s)):t.cache,a<c&&(l=c,c=a,a=l),c<=r&&r<=a||""===c||""===a):null},wild:function(e,t){if(H.wildOrTest.test(t.iFilter)){var r=""+(D.parseFilter(e,t.iFilter,t)||"");!H.wildTest.test(r)&&t.nestedFilters&&(r=t.isMatch?r:"^("+r+")$");try{return new RegExp(r.replace(H.wild01,"\\S{1}").replace(H.wild0More,"\\S*"),e.widgetOptions.filter_ignoreCase?"i":"").test(t.exact)}catch(e){}}return null},fuzzy:function(e,t){if(H.fuzzyTest.test(t.iFilter)){for(var r=0,a=t.iExact.length,s=t.iFilter.slice(1),i=D.parseFilter(e,s,t)||"",o=0;o<a;o++)t.iExact[o]===i[r]&&(r+=1);return r===i.length}return null}},init:function(r){N.language=A.extend(!0,{},{to:"to",or:"or",and:"and"},N.language);function e(e,t,r){return""===(t=t.trim())?"":(e||"")+t+(r||"")}var t,a,s,i,o,n,l,c,d=r.config,f=d.widgetOptions;if(d.$table.addClass("hasFilters"),d.lastSearch=[],f.filter_searchTimer=null,f.filter_initTimer=null,f.filter_formatterCount=0,f.filter_formatterInit=[],f.filter_anyColumnSelector='[data-column="all"],[data-column="any"]',f.filter_multipleColumnSelector='[data-column*="-"],[data-column*=","]',n="\\{"+H.query+"\\}",A.extend(H,{child:new RegExp(d.cssChildRow),filtered:new RegExp(f.filter_filteredRow),alreadyFiltered:new RegExp("(\\s+(-"+e("|",N.language.or)+e("|",N.language.to)+")\\s+)","i"),toTest:new RegExp("\\s+(-"+e("|",N.language.to)+")\\s+","i"),toSplit:new RegExp("(?:\\s+(?:-"+e("|",N.language.to)+")\\s+)","gi"),andTest:new RegExp("\\s+("+e("",N.language.and,"|")+"&&)\\s+","i"),andSplit:new RegExp("(?:\\s+(?:"+e("",N.language.and,"|")+"&&)\\s+)","gi"),orTest:new RegExp("(\\|"+e("|\\s+",N.language.or,"\\s+")+")","i"),orSplit:new RegExp("(?:\\|"+e("|\\s+(?:",N.language.or,")\\s+")+")","gi"),iQuery:new RegExp(n,"i"),igQuery:new RegExp(n,"ig"),operTest:/^[<>]=?/,gtTest:/>/,gteTest:/>=/,ltTest:/</,lteTest:/<=/,notTest:/^\!/,wildOrTest:/[\?\*\|]/,wildTest:/\?\*/,fuzzyTest:/^~/,exactTest:/[=\"\|!]/}),n=d.$headers.filter(".filter-false, .parser-false").length,!1!==f.filter_columnFilters&&n!==d.$headers.length&&D.buildRow(r,d,f),s="addRows updateCell update updateRows updateComplete appendCache filterReset "+"filterAndSortReset filterResetSaved filterEnd search ".split(" ").join(d.namespace+"filter "),d.$table.bind(s,function(e,t){return n=f.filter_hideEmpty&&A.isEmptyObject(d.cache)&&!(d.delayInit&&"appendCache"===e.type),d.$table.find("."+h.filterRow).toggleClass(f.filter_filteredRow,n),/(search|filter)/.test(e.type)||(e.stopPropagation(),D.buildDefault(r,!0)),"filterReset"===e.type||"filterAndSortReset"===e.type?(d.$table.find("."+h.filter).add(f.filter_$externalFilters).val(""),"filterAndSortReset"===e.type?N.sortReset(this.config,function(){D.searching(r,[])}):D.searching(r,[])):"filterResetSaved"===e.type?N.storage(r,"tablesorter-filters",""):"filterEnd"===e.type?D.buildDefault(r,!0):(t="search"===e.type?t:"updateComplete"===e.type?d.$table.data("lastSearch"):"",/(update|add)/.test(e.type)&&"updateComplete"!==e.type&&(d.lastCombinedFilter=null,d.lastSearch=[],setTimeout(function(){d.$table.triggerHandler("filterFomatterUpdate")},100)),D.searching(r,t,!0)),!1}),f.filter_reset&&(f.filter_reset instanceof A?f.filter_reset.click(function(){d.$table.triggerHandler("filterReset")}):A(f.filter_reset).length&&A(document).undelegate(f.filter_reset,"click"+d.namespace+"filter").delegate(f.filter_reset,"click"+d.namespace+"filter",function(){d.$table.triggerHandler("filterReset")})),f.filter_functions)for(o=0;o<d.columns;o++)if(l=N.getColumnData(r,f.filter_functions,o))if(c=!((i=d.$headerIndexed[o].removeClass("filter-select")).hasClass("filter-false")||i.hasClass("parser-false")),!(t="")===l&&c)D.buildSelect(r,o);else if("object"==typeof l&&c){for(a in l)"string"==typeof a&&(t+=""===t?'<option value="">'+(i.data("placeholder")||i.attr("data-placeholder")||f.filter_placeholder.select||"")+"</option>":"",0<=(s=n=a).indexOf(f.filter_selectSourceSeparator)&&(s=(n=a.split(f.filter_selectSourceSeparator))[1],n=n[0]),t+="<option "+(s===n?"":'data-function-name="'+a+'" ')+'value="'+n+'">'+s+"</option>");d.$table.find("thead").find("select."+h.filter+'[data-column="'+o+'"]').append(t),("function"==typeof(s=f.filter_selectSource)||N.getColumnData(r,s,o))&&D.buildSelect(d.table,o,"",!0,i.hasClass(f.filter_onlyAvail))}D.buildDefault(r,!0),D.bindSearch(r,d.$table.find("."+h.filter),!0),f.filter_external&&D.bindSearch(r,f.filter_external),f.filter_hideFilters&&D.hideFilters(d),d.showProcessing&&(s="filterStart filterEnd ".split(" ").join(d.namespace+"filter-sp "),d.$table.unbind(s.replace(N.regex.spaces," ")).bind(s,function(e,t){i=t?d.$table.find("."+h.header).filter("[data-column]").filter(function(){return""!==t[A(this).data("column")]}):"",N.isProcessing(r,"filterStart"===e.type,t?i:"")})),d.filteredRows=d.totalRows,s="tablesorter-initialized pagerBeforeInitialized ".split(" ").join(d.namespace+"filter "),d.$table.unbind(s.replace(N.regex.spaces," ")).bind(s,function(){D.completeInit(this)}),d.pager&&d.pager.initialized&&!f.filter_initialized?(d.$table.triggerHandler("filterFomatterUpdate"),setTimeout(function(){D.filterInitComplete(d)},100)):f.filter_initialized||D.completeInit(r)},completeInit:function(e){var t=e.config,r=t.widgetOptions,a=D.setDefaults(e,t,r)||[];!a.length||t.delayInit&&""===a.join("")||N.setFilters(e,a,!0),t.$table.triggerHandler("filterFomatterUpdate"),setTimeout(function(){r.filter_initialized||D.filterInitComplete(t)},100)},formatterUpdated:function(e,t){e=e&&e.closest("table"),e=e.length&&e[0].config,e=e&&e.widgetOptions;e&&!e.filter_initialized&&(e.filter_formatterInit[t]=1)},filterInitComplete:function(e){function t(){s.filter_initialized=!0,e.lastSearch=e.$table.data("lastSearch"),e.$table.triggerHandler("filterInit",e),D.findRows(e.table,e.lastSearch||[]),N.debug(e,"filter")&&console.log("Filter >> Widget initialized")}var r,a,s=e.widgetOptions,i=0;if(A.isEmptyObject(s.filter_formatter))t();else{for(a=s.filter_formatterInit.length,r=0;r<a;r++)1===s.filter_formatterInit[r]&&i++;clearTimeout(s.filter_initTimer),s.filter_initialized||i!==s.filter_formatterCount?s.filter_initialized||(s.filter_initTimer=setTimeout(function(){t()},500)):t()}},processFilters:function(e,t){for(var r=[],a=t?encodeURIComponent:decodeURIComponent,s=e.length,i=0;i<s;i++)e[i]&&(r[i]=a(e[i]));return r},setDefaults:function(e,t,r){var a,s,i,o,n=N.getFilters(e)||[];if(r.filter_saveFilters&&N.storage&&(e=N.storage(e,"tablesorter-filters")||[],(a=A.isArray(e))&&""===e.join("")||!a||(n=D.processFilters(e))),""===n.join(""))for(o=t.$headers.add(r.filter_$externalFilters).filter("["+r.filter_defaultAttrib+"]"),s=0;s<=t.columns;s++)i=s===t.columns?"all":s,n[s]=o.filter('[data-column="'+i+'"]').attr(r.filter_defaultAttrib)||n[s]||"";return t.$table.data("lastSearch",n),n},parseFilter:function(e,t,r,a){return a||r.parsed[r.index]?e.parsers[r.index].format(t,e.table,[],r.index):t},buildRow:function(e,t,r){for(var a,s,i,o,n,l,c,d=r.filter_cellFilter,f=t.columns,u=A.isArray(d),g='<tr role="search" class="'+h.filterRow+" "+t.cssIgnoreRow+'">',p=0;p<f;p++)t.$headerIndexed[p].length&&(g=(g+=1<(c=t.$headerIndexed[p]&&t.$headerIndexed[p][0].colSpan||0)?'<td data-column="'+p+"-"+(p+c-1)+'" colspan="'+c+'"':'<td data-column="'+p+'"')+(u?d[p]?' class="'+d[p]+'"':"":""!==d?' class="'+d+'"':"")+"></td>");for(t.$filters=A(g+="</tr>").appendTo(t.$table.children("thead").eq(0)).children("td"),p=0;p<f;p++)(s=t.$headerIndexed[p])&&s.length&&(a=D.getColumnElm(t,t.$filters,p),l=N.getColumnData(e,r.filter_functions,p),i=r.filter_functions&&l&&"function"!=typeof l||s.hasClass("filter-select"),o=N.getColumnData(e,t.headers,p),o="false"===N.getData(s[0],o,"filter")||"false"===N.getData(s[0],o,"parser"),i?g=A("<select>").appendTo(a):((l=N.getColumnData(e,r.filter_formatter,p))?(r.filter_formatterCount++,(g=(g=l(a,p))&&0===g.length?a.children("input"):g)&&(0===g.parent().length||g.parent().length&&g.parent()[0]!==a[0])&&a.append(g)):g=A('<input type="search">').appendTo(a),g&&(c=s.data("placeholder")||s.attr("data-placeholder")||r.filter_placeholder.search||"",g.attr("placeholder",c))),g)&&(n=(A.isArray(r.filter_cssFilter)?void 0!==r.filter_cssFilter[p]&&r.filter_cssFilter[p]||"":r.filter_cssFilter)||"",g.addClass(h.filter+" "+n),c=(n=r.filter_filterLabel).match(/{{([^}]+?)}}/g),A.each(c=c||["{{label}}"],function(e,t){var r=new RegExp(t,"g"),t=s.attr("data-"+t.replace(/{{|}}/g,"")),t=void 0===t?s.text():t;n=n.replace(r,A.trim(t))}),g.attr({"data-column":a.attr("data-column"),"aria-label":n}),o)&&(g.attr("placeholder","").addClass(h.filterDisabled)[0].disabled=!0)},bindSearch:function(s,e,t){var r,i,o,a,n;s=A(s)[0],(e=A(e)).length&&(i=s.config,o=i.widgetOptions,a=i.namespace+"filter",n=o.filter_$externalFilters,!0!==t&&(r=o.filter_anyColumnSelector+","+o.filter_multipleColumnSelector,o.filter_$anyMatch=e.filter(r),n&&n.length?o.filter_$externalFilters=o.filter_$externalFilters.add(e):o.filter_$externalFilters=e,N.setFilters(s,i.$table.data("lastSearch")||[],!1===t)),r="keypress keyup keydown search change input ".split(" ").join(a+" "),e.attr("data-lastSearchTime",(new Date).getTime()).unbind(r.replace(N.regex.spaces," ")).bind("keydown"+a,function(e){if(e.which===l.escape&&!s.config.widgetOptions.filter_resetOnEsc)return!1}).bind("keyup"+a,function(e){o=s.config.widgetOptions;var t=parseInt(A(this).attr("data-column"),10),r="boolean"==typeof o.filter_liveSearch?o.filter_liveSearch:N.getColumnData(s,o.filter_liveSearch,t);if(void 0===r&&(r=o.filter_liveSearch.fallback||!1),A(this).attr("data-lastSearchTime",(new Date).getTime()),e.which===l.escape)this.value=o.filter_resetOnEsc?"":i.lastSearch[t];else{if(""!==this.value&&("number"==typeof r&&this.value.length<r||e.which!==l.enter&&e.which!==l.backSpace&&(e.which<l.space||e.which>=l.left&&e.which<=l.down)))return;if(!1===r&&""!==this.value&&e.which!==l.enter)return}D.searching(s,!0,!0,t)}).bind("search change keypress input blur ".split(" ").join(a+" "),function(e){var t=parseInt(A(this).attr("data-column"),10),r=e.type,a="boolean"==typeof o.filter_liveSearch?o.filter_liveSearch:N.getColumnData(s,o.filter_liveSearch,t);!s.config.widgetOptions.filter_initialized||e.which!==l.enter&&"search"!==r&&"blur"!==r&&("change"!==r&&"input"!==r||!0!==a&&(!0===a||"INPUT"===e.target.nodeName)||this.value===i.lastSearch[t])||(e.preventDefault(),A(this).attr("data-lastSearchTime",(new Date).getTime()),D.searching(s,"keypress"!==r||e.which===l.enter,!0,t))}))},searching:function(e,t,r,a){var s,i=e.config.widgetOptions;void 0===a?s=!1:void 0===(s="boolean"==typeof i.filter_liveSearch?i.filter_liveSearch:N.getColumnData(e,i.filter_liveSearch,a))&&(s=i.filter_liveSearch.fallback||!1),clearTimeout(i.filter_searchTimer),void 0===t||!0===t?i.filter_searchTimer=setTimeout(function(){D.checkFilters(e,t,r)},s?i.filter_searchDelay:10):D.checkFilters(e,t,r)},equalFilters:function(e,t,r){var a,s=[],i=[],o=e.columns+1;for(t=A.isArray(t)?t:[],r=A.isArray(r)?r:[],a=0;a<o;a++)s[a]=t[a]||"",i[a]=r[a]||"";return s.join(",")===i.join(",")},checkFilters:function(e,t,r){var a=e.config,s=a.widgetOptions,i=A.isArray(t),o=i?t:N.getFilters(e,!0),n=o||[];if(A.isEmptyObject(a.cache))a.delayInit&&(!a.pager||a.pager&&a.pager.initialized)&&N.updateCache(a,function(){D.checkFilters(e,!1,r)});else{if(i&&(N.setFilters(e,o,!1,!0!==r),s.filter_initialized||(a.lastSearch=[],a.lastCombinedFilter="")),s.filter_hideFilters&&a.$table.find("."+h.filterRow).triggerHandler(D.hideFiltersCheck(a)?"mouseleave":"mouseenter"),D.equalFilters(a,a.lastSearch,n)){if(!1!==t)return;a.lastCombinedFilter="",a.lastSearch=[]}if(o=o||[],o=Array.prototype.map?o.map(String):o.join("�").split("�"),s.filter_initialized&&a.$table.triggerHandler("filterStart",[o]),!a.showProcessing)return D.findRows(e,o,n),!1;setTimeout(function(){return D.findRows(e,o,n),!1},30)}},hideFiltersCheck:function(e){if("function"==typeof e.widgetOptions.filter_hideFilters){var t=e.widgetOptions.filter_hideFilters(e);if("boolean"==typeof t)return t}return""===N.getFilters(e.$table).join("")},hideFilters:function(a,e){var s;(e||a.$table).find("."+h.filterRow).addClass(h.filterRowHide).bind("mouseenter mouseleave",function(e){var t=e,r=A(this);clearTimeout(s),s=setTimeout(function(){/enter|over/.test(t.type)?r.removeClass(h.filterRowHide):A(document.activeElement).closest("tr")[0]!==r[0]&&r.toggleClass(h.filterRowHide,D.hideFiltersCheck(a))},200)}).find("input, select").bind("focus blur",function(e){var t=e,r=A(this).closest("tr");clearTimeout(s),s=setTimeout(function(){clearTimeout(s),r.toggleClass(h.filterRowHide,D.hideFiltersCheck(a)&&"focus"!==t.type)},200)})},defaultFilter:function(e,t){if(""===e)return e;var r=H.iQuery,a=t.match(H.igQuery).length,s=1<a?A.trim(e).split(/\s/):[A.trim(e)],i=s.length-1,o=0,n=t;for(i<1&&1<a&&(s[1]=s[0]);r.test(n);)n=n.replace(r,s[o++]||""),r.test(n)&&o<i&&""!==(s[o]||"")&&(n=t.replace(r,n));return n},getLatestSearch:function(e){return e?e.sort(function(e,t){return A(t).attr("data-lastSearchTime")-A(e).attr("data-lastSearchTime")}):e||A()},findRange:function(e,t,r){var a,s,i,o,n,l,c,d,f=[];if(/^[0-9]+$/.test(t))return[parseInt(t,10)];if(!r&&/-/.test(t))for(d=(a=t.match(/(\d+)\s*-\s*(\d+)/g))?a.length:0,c=0;c<d;c++){for(s=a[c].split(/\s*-\s*/),i=parseInt(s[0],10)||0,(o=parseInt(s[1],10)||e.columns-1)<i&&(s=i,i=o,o=s),o>=e.columns&&(o=e.columns-1);i<=o;i++)f[f.length]=i;t=t.replace(a[c],"")}if(!r&&/,/.test(t))for(d=(n=t.split(/\s*,\s*/)).length,l=0;l<d;l++)""!==n[l]&&(c=parseInt(n[l],10))<e.columns&&(f[f.length]=c);if(!f.length)for(c=0;c<e.columns;c++)f[f.length]=c;return f},getColumnElm:function(t,e,r){return e.filter(function(){var e=D.findRange(t,A(this).attr("data-column"));return-1<A.inArray(r,e)})},multipleColumns:function(e,t){var r=e.widgetOptions,r=r.filter_initialized||!t.filter(r.filter_anyColumnSelector).length,t=A.trim(D.getLatestSearch(t).attr("data-column")||"");return D.findRange(e,t,!r)},processTypes:function(e,t,r){var a,s=null,i=null;for(a in D.types)A.inArray(a,r.excludeMatch)<0&&null===i&&null!==(i=D.types[a](e,t,r))&&(t.matchedOn=a,s=i);return s},matchType:function(e,t){var r=e.widgetOptions,a=e.$headerIndexed[t],r=!(a.hasClass("filter-exact")||!a.hasClass("filter-match")&&(r.filter_columnFilters?a=e.$filters.find("."+h.filter).add(r.filter_$externalFilters).filter('[data-column="'+t+'"]'):r.filter_$externalFilters&&(a=r.filter_$externalFilters.filter('[data-column="'+t+'"]')),!a.length||"match"!==e.widgetOptions.filter_matchType[(a[0].nodeName||"").toLowerCase()]));return r},processRow:function(t,r,e){var a,s,i,o,n,l=t.widgetOptions,c=!0,d=l.filter_$anyMatch&&l.filter_$anyMatch.length,f=l.filter_$anyMatch&&l.filter_$anyMatch.length?D.multipleColumns(t,l.filter_$anyMatch):[];if(r.$cells=r.$row.children(),r.matchedOn=null,r.anyMatchFlag&&1<f.length||r.anyMatchFilter&&!d){if(r.anyMatch=!0,r.isMatch=!0,r.rowArray=r.$cells.map(function(e){if(-1<A.inArray(e,f)||r.anyMatchFilter&&!d)return r.parsed[e]?n=r.cacheArray[e]:(n=r.rawArray[e],n=A.trim(l.filter_ignoreCase?n.toLowerCase():n),t.sortLocaleCompare&&(n=N.replaceAccents(n))),n}).get(),r.filter=r.anyMatchFilter,r.iFilter=r.iAnyMatchFilter,r.exact=r.rowArray.join(" "),r.iExact=l.filter_ignoreCase?r.exact.toLowerCase():r.exact,r.cache=r.cacheArray.slice(0,-1).join(" "),e.excludeMatch=e.noAnyMatch,null!==(s=D.processTypes(t,r,e)))c=s;else if(l.filter_startsWith)for(c=!1,f=Math.min(t.columns,r.rowArray.length);!c&&0<f;)f--,c=c||0===r.rowArray[f].indexOf(r.iFilter);else c=0<=(r.iExact+r.childRowText).indexOf(r.iFilter);if(r.anyMatch=!1,r.filters.join("")===r.filter)return c}for(f=0;f<t.columns;f++)r.filter=r.filters[f],r.index=f,e.excludeMatch=e.excludeFilter[f],r.filter&&(r.cache=r.cacheArray[f],a=r.parsed[f]?r.cache:r.rawArray[f]||"",r.exact=t.sortLocaleCompare?N.replaceAccents(a):a,r.iExact=!H.type.test(typeof r.exact)&&l.filter_ignoreCase?r.exact.toLowerCase():r.exact,r.isMatch=D.matchType(t,f),a=c,o=l.filter_columnFilters&&t.$filters.add(l.filter_$externalFilters).filter('[data-column="'+f+'"]').find("select option:selected").attr("data-function-name")||"",t.sortLocaleCompare&&(r.filter=N.replaceAccents(r.filter)),l.filter_defaultFilter&&H.iQuery.test(e.defaultColFilter[f])&&(r.filter=D.defaultFilter(r.filter,e.defaultColFilter[f])),r.iFilter=l.filter_ignoreCase?(r.filter||"").toLowerCase():r.filter,s=null,(i=e.functions[f])&&("function"==typeof i?s=i(r.exact,r.cache,r.filter,f,r.$row,t,r):"function"==typeof i[o||r.filter]&&(s=i[n=o||r.filter](r.exact,r.cache,r.filter,f,r.$row,t,r))),c=!!(a=null===s&&(s=D.processTypes(t,r,e),n=!0===i&&("and"===r.matchedOn||"or"===r.matchedOn),null===s||n)?!0===i?r.isMatch?0<=(""+r.iExact).search(r.iFilter):r.filter===r.exact:(n=(r.iExact+r.childRowText).indexOf(D.parseFilter(t,r.iFilter,r)),!l.filter_startsWith&&0<=n||l.filter_startsWith&&0===n):s)&&c);return c},findRows:function(e,r,t){if(!D.equalFilters(e.config,e.config.lastSearch,t)&&e.config.widgetOptions.filter_initialized){var a,s,i,o,n,l,c,d,f,u,g,p,h,m,b,y,w,v,x,C,_,$,S,z=A.extend([],r),F=e.config,R=F.widgetOptions,T=N.debug(F,"filter"),I={anyMatch:!1,filters:r,filter_regexCache:[]},k={noAnyMatch:["range","operators"],functions:[],excludeFilter:[],defaultColFilter:[],defaultAnyFilter:N.getColumnData(e,R.filter_defaultFilter,F.columns,!0)||""};for(I.parsed=[],f=0;f<F.columns;f++)I.parsed[f]=R.filter_useParsedData||F.parsers&&F.parsers[f]&&F.parsers[f].parsed||N.getData&&"parsed"===N.getData(F.$headerIndexed[f],N.getColumnData(e,F.headers,f),"filter")||F.$headerIndexed[f].hasClass("filter-parsed"),k.functions[f]=N.getColumnData(e,R.filter_functions,f)||F.$headerIndexed[f].hasClass("filter-select"),k.defaultColFilter[f]=N.getColumnData(e,R.filter_defaultFilter,f)||"",k.excludeFilter[f]=(N.getColumnData(e,R.filter_excludeFilter,f,!0)||"").split(/\s+/);for(T&&(console.log("Filter >> Starting filter widget search",r),m=new Date),F.filteredRows=0,t=z||[],c=F.totalRows=0;c<F.$tbodies.length;c++){if(d=N.processTbody(e,F.$tbodies.eq(c),!0),f=F.columns,s=F.cache[c].normalized,o=A(A.map(s,function(e){return e[f].$row.get()})),""===t.join("")||R.filter_serversideFiltering)o.removeClass(R.filter_filteredRow).not("."+F.cssChildRow).css("display","");else{if(a=(o=o.not("."+F.cssChildRow)).length,(R.filter_$anyMatch&&R.filter_$anyMatch.length||void 0!==r[F.columns])&&(I.anyMatchFlag=!0,I.anyMatchFilter=""+(r[F.columns]||R.filter_$anyMatch&&D.getLatestSearch(R.filter_$anyMatch).val()||""),R.filter_columnAnyMatch)){for(x=I.anyMatchFilter.split(H.andSplit),C=!1,y=0;y<x.length;y++)1<(_=x[y].split(":")).length&&(isNaN(_[0])?A.each(F.headerContent,function(e,t){-1<t.toLowerCase().indexOf(_[0])&&(r[$=e]=_[1])}):$=parseInt(_[0],10)-1,0<=$)&&$<F.columns&&(r[$]=_[1],x.splice(y,1),y--,C=!0);C&&(I.anyMatchFilter=x.join(" && "))}if(v=R.filter_searchFiltered,g=F.lastSearch||F.$table.data("lastSearch")||[],v)for(y=0;y<f+1;y++)b=r[y]||"",v||(y=f),v=v&&g.length&&0===b.indexOf(g[y]||"")&&!H.alreadyFiltered.test(b)&&!H.exactTest.test(b)&&!(H.isNeg1.test(b)||H.isNeg2.test(b))&&!(""!==b&&F.$filters&&F.$filters.filter('[data-column="'+y+'"]').find("select").length&&!D.matchType(F,y));for(w=o.not("."+R.filter_filteredRow).length,v&&0===w&&(v=!1),T&&console.log("Filter >> Searching through "+(v&&w<a?w:"all")+" rows"),I.anyMatchFlag&&(F.sortLocaleCompare&&(I.anyMatchFilter=N.replaceAccents(I.anyMatchFilter)),R.filter_defaultFilter&&H.iQuery.test(k.defaultAnyFilter)&&(I.anyMatchFilter=D.defaultFilter(I.anyMatchFilter,k.defaultAnyFilter),v=!1),I.iAnyMatchFilter=R.filter_ignoreCase&&F.ignoreCase?I.anyMatchFilter.toLowerCase():I.anyMatchFilter),l=0;l<a;l++)if(S=o[l].className,!(l&&H.child.test(S)||v&&H.filtered.test(S))){if(I.$row=o.eq(l),I.rowIndex=l,I.cacheArray=s[l],i=I.cacheArray[F.columns],I.rawArray=i.raw,I.childRowText="",!R.filter_childByColumn){for(S="",u=i.child,y=0;y<u.length;y++)S+=" "+u[y].join(" ")||"";I.childRowText=R.filter_childRows?R.filter_ignoreCase?S.toLowerCase():S:""}if(p=!1,h=D.processRow(F,I,k),n=i.$row,b=!!h,u=i.$row.filter(":gt(0)"),R.filter_childRows&&u.length){if(R.filter_childByColumn)for(R.filter_childWithSibs||(u.addClass(R.filter_filteredRow),n=n.eq(0)),y=0;y<u.length;y++)I.$row=u.eq(y),I.cacheArray=i.child[y],I.rawArray=I.cacheArray,b=D.processRow(F,I,k),p=p||b,!R.filter_childWithSibs&&b&&u.eq(y).removeClass(R.filter_filteredRow);p=p||h}else p=b;n.toggleClass(R.filter_filteredRow,!p)[0].display=p?"":"none"}}F.filteredRows+=o.not("."+R.filter_filteredRow).length,F.totalRows+=o.length,N.processTbody(e,d,!1)}F.lastCombinedFilter=z.join(""),F.lastSearch=z,F.$table.data("lastSearch",z),R.filter_saveFilters&&N.storage&&N.storage(e,"tablesorter-filters",D.processFilters(z,!0)),T&&console.log("Filter >> Completed search"+N.benchmark(m)),R.filter_initialized&&(F.$table.triggerHandler("filterBeforeEnd",F),F.$table.triggerHandler("filterEnd",F)),setTimeout(function(){N.applyWidget(F.table)},0)}},getOptionSource:function(e,t,r){var a=(e=A(e)[0]).config,s=!1,i=a.widgetOptions.filter_selectSource,a=a.$table.data("lastSearch")||[],o="function"==typeof i||N.getColumnData(e,i,t);if(r&&""!==a[t]&&(r=!1),!0===o)s=i(e,t,r);else{if(o instanceof A||"string"===A.type(o)&&0<=o.indexOf("</option>"))return o;if(A.isArray(o))s=o;else if("object"===A.type(i)&&o&&null===(s=o(e,t,r)))return null}return!1===s&&(s=D.getOptions(e,t,r)),D.processOptions(e,t,s)},processOptions:function(a,s,r){if(!A.isArray(r))return!1;var i,e,t,o,n,l=(a=A(a)[0]).config,c=null!=s&&0<=s&&s<l.columns,d=!!c&&l.$headerIndexed[s].hasClass("filter-select-sort-desc"),f=[];if(r=A.grep(r,function(e,t){return!!e.text||A.inArray(e,r)===t}),!c||!l.$headerIndexed[s].hasClass("filter-select-nosort")){for(o=r.length,t=0;t<o;t++)n=(e=r[t]).text||e,n=(c&&l.parsers&&l.parsers.length&&l.parsers[s].format(n,a,[],s)||n).toString(),n=l.widgetOptions.filter_ignoreCase?n.toLowerCase():n,e.text?(e.parsed=n,f[f.length]=e):f[f.length]={text:e,parsed:n};for(i=l.textSorter||"",f.sort(function(e,t){var r=(d?t:e).parsed,e=(d?e:t).parsed;return c&&"function"==typeof i?i(r,e,!0,s,a):c&&"object"==typeof i&&i.hasOwnProperty(s)?i[s](r,e,!0,s,a):!N.sortNatural||N.sortNatural(r,e)}),r=[],o=f.length,t=0;t<o;t++)r[r.length]=f[t]}return r},getOptions:function(e,t,r){for(var a,s,i,o,n,l,c,d=(e=A(e)[0]).config,f=d.widgetOptions,u=[],g=0;g<d.$tbodies.length;g++)for(o=d.cache[g],s=d.cache[g].normalized.length,a=0;a<s;a++)if(i=o.row?o.row[a]:o.normalized[a][d.columns].$row[0],!r||!i.className.match(f.filter_filteredRow))if(f.filter_useParsedData||d.parsers[t].parsed||d.$headerIndexed[t].hasClass("filter-parsed")){if(u[u.length]=""+o.normalized[a][t],f.filter_childRows&&f.filter_childByColumn)for(c=o.normalized[a][d.columns].$row.length-1,n=0;n<c;n++)u[u.length]=""+o.normalized[a][d.columns].child[n][t]}else if(u[u.length]=o.normalized[a][d.columns].raw[t],f.filter_childRows&&f.filter_childByColumn)for(c=o.normalized[a][d.columns].$row.length,n=1;n<c;n++)l=o.normalized[a][d.columns].$row.eq(n).children().eq(t),u[u.length]=""+N.getElementText(d,l,t);return u},buildSelect:function(e,t,r,a,s){if(e=A(e)[0],t=parseInt(t,10),e.config.cache&&!A.isEmptyObject(e.config.cache)){var i,o,n,l,c,d=e.config,f=d.widgetOptions,u=d.$headerIndexed[t],g='<option value="">'+(u.data("placeholder")||u.attr("data-placeholder")||f.filter_placeholder.select||"")+"</option>",u=d.$table.find("thead").find("select."+h.filter+'[data-column="'+t+'"]').val();if(void 0!==r&&""!==r||null!==(r=D.getOptionSource(e,t,s))){if(A.isArray(r)){for(i=0;i<r.length;i++)if((c=r[i]).text){for(o in c["data-function-name"]=void 0===c.value?c.text:c.value,g+="<option",c)c.hasOwnProperty(o)&&"text"!==o&&(g+=" "+o+'="'+c[o].replace(H.quote,"&quot;")+'"');c.value||(g+=' value="'+c.text.replace(H.quote,"&quot;")+'"'),g+=">"+c.text.replace(H.quote,"&quot;")+"</option>"}else""+c!="[object Object]"&&(0<=(o=n=c=(""+c).replace(H.quote,"&quot;")).indexOf(f.filter_selectSourceSeparator)&&(o=(l=n.split(f.filter_selectSourceSeparator))[0],n=l[1]),g+=""!==c?"<option "+(o===n?"":'data-function-name="'+c+'" ')+'value="'+o+'">'+n+"</option>":"");r=[]}e=(d.$filters||d.$table.children("thead")).find("."+h.filter),(s=(e=f.filter_$externalFilters?e&&e.length?e.add(f.filter_$externalFilters):f.filter_$externalFilters:e).filter('select[data-column="'+t+'"]')).length&&(s[a?"html":"append"](g),A.isArray(r)||s.append(r).val(u),s.val(u))}}},buildDefault:function(e,t){for(var r,a,s=e.config,i=s.widgetOptions,o=s.columns,n=0;n<o;n++)a=!((r=s.$headerIndexed[n]).hasClass("filter-false")||r.hasClass("parser-false")),(r.hasClass("filter-select")||!0===N.getColumnData(e,i.filter_functions,n))&&a&&D.buildSelect(e,n,"",t,r.hasClass(i.filter_onlyAvail))}},H=D.regex,N.getFilters=function(e,t,r,a){var s,i,o,n,l=[],c=e?A(e)[0].config:"",d=c?c.widgetOptions:"";if(!0!==t&&d&&!d.filter_columnFilters||A.isArray(r)&&D.equalFilters(c,r,c.lastSearch))return A(e).data("lastSearch")||[];if(c&&(c.$filters&&(i=c.$filters.find("."+h.filter)),i=d.filter_$externalFilters?i&&i.length?i.add(d.filter_$externalFilters):d.filter_$externalFilters:i)&&i.length)for(l=r||[],s=0;s<c.columns+1;s++)n=s===c.columns?d.filter_anyColumnSelector+","+d.filter_multipleColumnSelector:'[data-column="'+s+'"]',(o=i.filter(n)).length&&(o=D.getLatestSearch(o),A.isArray(r)?(a&&1<o.length&&(o=o.slice(1)),(o=s===c.columns?(n=o.filter(d.filter_anyColumnSelector)).length?n:o:o).val(r[s]).trigger("change"+c.namespace)):(l[s]=o.val()||"",(s===c.columns?o.slice(1).filter('[data-column*="'+o.attr("data-column")+'"]'):o.slice(1)).val(l[s])),s===c.columns)&&o.length&&(d.filter_$anyMatch=o);return l},N.setFilters=function(e,t,r,a){var s=e?A(e)[0].config:"",e=N.getFilters(e,!0,t,a);return void 0===r&&(r=!0),s&&r&&(s.lastCombinedFilter=null,s.lastSearch=[],D.searching(s.table,t,a),s.$table.triggerHandler("filterFomatterUpdate")),0!==e.length}}(e),function($,S){"use strict";var z=$.tablesorter||{};function F(e,t){var r=isNaN(t.stickyHeaders_offset)?$(t.stickyHeaders_offset):[];return r.length?r.height()||0:parseInt(t.stickyHeaders_offset,10)||0}$.extend(z.css,{sticky:"tablesorter-stickyHeader",stickyVis:"tablesorter-sticky-visible",stickyHide:"tablesorter-sticky-hidden",stickyWrap:"tablesorter-sticky-wrapper"}),z.addHeaderResizeEvent=function(e,t,r){if((e=$(e)[0]).config){var r=$.extend({},{timer:250},r),l=e.config,c=l.widgetOptions,a=function(e){var t,r,a,s,i,o,n=l.$headers.length;for(c.resize_flag=!0,r=[],t=0;t<n;t++)s=(a=l.$headers.eq(t)).data("savedSizes")||[0,0],i=a[0].offsetWidth,o=a[0].offsetHeight,i===s[0]&&o===s[1]||(a.data("savedSizes",[i,o]),r.push(a[0]));r.length&&!1!==e&&l.$table.triggerHandler("resize",[r]),c.resize_flag=!1};if(clearInterval(c.resize_timer),t)return c.resize_flag=!1;a(!1),c.resize_timer=setInterval(function(){c.resize_flag||a()},r.timer)}},z.addWidget({id:"stickyHeaders",priority:54,options:{stickyHeaders:"",stickyHeaders_appendTo:null,stickyHeaders_attachTo:null,stickyHeaders_xScroll:null,stickyHeaders_yScroll:null,stickyHeaders_offset:0,stickyHeaders_filteredToTop:!0,stickyHeaders_cloneId:"-sticky",stickyHeaders_addResizeEvent:!0,stickyHeaders_includeCaption:!0,stickyHeaders_zIndex:2},format:function(e,t,l){if(!(t.$table.hasClass("hasStickyHeaders")||0<=$.inArray("filter",t.widgets)&&!t.$table.hasClass("hasFilters"))){var r,a,s,i,c=t.$table,d=$(l.stickyHeaders_attachTo||l.stickyHeaders_appendTo),o=t.namespace+"stickyheaders ",f=$(l.stickyHeaders_yScroll||l.stickyHeaders_attachTo||S),n=$(l.stickyHeaders_xScroll||l.stickyHeaders_attachTo||S),u=c.children("thead:first").children("tr").not(".sticky-false").children(),g=c.children("tfoot"),p=F(0,l),h=c.parent().closest("."+z.css.table).hasClass("hasStickyHeaders")?c.parent().closest("table.tablesorter")[0].config.widgetOptions.$sticky.parent():[],m=h.length?h.height():0,b=l.$sticky=c.clone().addClass("containsStickyHeaders "+z.css.sticky+" "+l.stickyHeaders+" "+t.namespace.slice(1)+"_extra_table").wrap('<div class="'+z.css.stickyWrap+'">'),y=b.parent().addClass(z.css.stickyHide).css({position:d.length?"absolute":"fixed",padding:parseInt(b.parent().parent().css("padding-left"),10),top:p+m,left:0,visibility:"hidden",zIndex:l.stickyHeaders_zIndex||2}),p=b.children("thead:first"),w="",v=function(e,t){for(var r,a,s,i=e.filter(":visible"),o=i.length,n=0;n<o;n++)a=t.filter(":visible").eq(n),r="border-box"===(s=i.eq(n)).css("box-sizing")?s.outerWidth():"collapse"===a.css("border-collapse")?S.getComputedStyle?parseFloat(S.getComputedStyle(s[0],null).width):(r=parseFloat(s.css("border-width")),s.outerWidth()-parseFloat(s.css("padding-left"))-parseFloat(s.css("padding-right"))-r):s.width(),a.css({width:r,"min-width":r,"max-width":r})},x=function(e){return!1===e&&h.length?c.position().left:d.length?parseInt(d.css("padding-left"),10)||0:c.offset().left-parseInt(c.css("margin-left"),10)-$(S).scrollLeft()},C=function(){y.css({left:x(),width:c.outerWidth()}),v(c,b),v(u,i)},_=function(e){var t,r,a,s,i,o,n;c.is(":visible")&&(m=h.length?h.offset().top-f.scrollTop()+h.height():0,s=c.offset(),t=F(0,l),a=(r=$.isWindow(f[0]))?f.scrollTop():h.length?parseInt(h[0].style.top,10):f.offset().top,a=d.length?a:f.scrollTop(),i=!l.stickyHeaders_includeCaption&&c.children("caption").height()||0,a=a+t+m-i,i=c.height()-(y.height()+(g.height()||0))-i,i="visible"==(s=a>s.top&&a<s.top+i?"visible":"hidden")?z.css.stickyVis:z.css.stickyHide,o=!y.hasClass(i),n={visibility:s},d.length&&(o=!0,n.top=r?a-d.offset().top:d.scrollTop()),(a=x(r))!==parseInt(y.css("left"),10)&&(o=!0,n.left=a),n.top=(n.top||0)+(!r&&h.length?h.height():t+m),o&&y.removeClass(z.css.stickyVis+" "+z.css.stickyHide).addClass(i).css(n),s===w&&!e||(C(),w=s))};if(d.length&&!d.css("position")&&d.css("position","relative"),b.attr("id")&&(b[0].id+=l.stickyHeaders_cloneId),b.find("> thead:gt(0), tr.sticky-false").hide(),b.find("> tbody, > tfoot").remove(),b.find("caption").toggle(l.stickyHeaders_includeCaption),i=p.children().children(),b.css({height:0,width:0,margin:0}),i.find("."+z.css.resizer).remove(),c.addClass("hasStickyHeaders").bind("pagerComplete"+o,function(){C()}),z.bindEvents(e,p.children().children("."+z.css.header)),l.stickyHeaders_appendTo?$(l.stickyHeaders_appendTo).append(y):c.after(y),t.onRenderHeader)for(a=(s=p.children("tr").children()).length,r=0;r<a;r++)t.onRenderHeader.apply(s.eq(r),[r,t,b]);n.add(f).unbind("scroll resize ".split(" ").join(o).replace(/\s+/g," ")).bind("scroll resize ".split(" ").join(o),function(e){_("resize"===e.type)}),t.$table.unbind("stickyHeadersUpdate"+o).bind("stickyHeadersUpdate"+o,function(){_(!0)}),l.stickyHeaders_addResizeEvent&&z.addHeaderResizeEvent(e),c.hasClass("hasFilters")&&l.filter_columnFilters&&(c.bind("filterEnd"+o,function(){var e=$(document.activeElement).closest("td"),e=e.parent().children().index(e);y.hasClass(z.css.stickyVis)&&l.stickyHeaders_filteredToTop&&(S.scrollTo(0,c.position().top),0<=e)&&t.$filters&&t.$filters.eq(e).find("a, select, input").filter(":visible").focus()}),z.filter.bindSearch(c,i.find("."+z.css.filter)),l.filter_hideFilters)&&z.filter.hideFilters(t,b),l.stickyHeaders_addResizeEvent&&c.bind("resize"+t.namespace+"stickyheaders",function(){C()}),_(!0),c.triggerHandler("stickyHeadersInit")}},remove:function(e,t,r){var a=t.namespace+"stickyheaders ";t.$table.removeClass("hasStickyHeaders").unbind("pagerComplete resize filterEnd stickyHeadersUpdate ".split(" ").join(a).replace(/\s+/g," ")).next("."+z.css.stickyWrap).remove(),r.$sticky&&r.$sticky.length&&r.$sticky.remove(),$(S).add(r.stickyHeaders_xScroll).add(r.stickyHeaders_yScroll).add(r.stickyHeaders_attachTo).unbind("scroll resize ".split(" ").join(a).replace(/\s+/g," ")),z.addHeaderResizeEvent(e,!0)}})}(e,window),function(c,t){"use strict";var d=c.tablesorter||{};c.extend(d.css,{resizableContainer:"tablesorter-resizable-container",resizableHandle:"tablesorter-resizable-handle",resizableNoSelect:"tablesorter-disableSelection",resizableStorage:"tablesorter-resizable"}),c(function(){var e="<style>body."+d.css.resizableNoSelect+" { -ms-user-select: none; -moz-user-select: -moz-none;-khtml-user-select: none; -webkit-user-select: none; user-select: none; }."+d.css.resizableContainer+" { position: relative; height: 1px; }."+d.css.resizableHandle+" { position: absolute; display: inline-block; width: 8px;top: 1px; cursor: ew-resize; z-index: 3; user-select: none; -moz-user-select: none; }</style>";c("head").append(e)}),d.resizable={init:function(e,t){if(!e.$table.hasClass("hasResizable")){e.$table.addClass("hasResizable");var r,a,s,i=e.$table,o=i.parent(),n=parseInt(i.css("margin-top"),10),l=t.resizable_vars={useStorage:d.storage&&!1!==t.resizable,$wrap:o,mouseXPosition:0,$target:null,$next:null,overflow:"auto"===o.css("overflow")||"scroll"===o.css("overflow")||"auto"===o.css("overflow-x")||"scroll"===o.css("overflow-x"),storedSizes:[]};for(d.resizableReset(e.table,!0),l.tableWidth=i.width(),l.fullWidth=Math.abs(o.width()-l.tableWidth)<20,l.useStorage&&l.overflow&&(d.storage(e.table,"tablesorter-table-original-css-width",l.tableWidth),s=d.storage(e.table,"tablesorter-table-resized-width")||"auto",d.resizable.setWidth(i,s,!0)),t.resizable_vars.storedSizes=o=(l.useStorage?d.storage(e.table,d.css.resizableStorage):[])||[],d.resizable.setWidths(e,t,o),d.resizable.updateStoredSizes(e,t),t.$resizable_container=c('<div class="'+d.css.resizableContainer+'">').css({top:n}).insertBefore(i),a=0;a<e.columns;a++)r=e.$headerIndexed[a],s=d.getColumnData(e.table,e.headers,a),"false"===d.getData(r,s,"resizable")||c('<div class="'+d.css.resizableHandle+'">').appendTo(t.$resizable_container).attr({"data-column":a,unselectable:"on"}).data("header",r).bind("selectstart",!1);d.resizable.bindings(e,t)}},updateStoredSizes:function(e,t){var r,a,s=e.columns,i=t.resizable_vars;for(i.storedSizes=[],r=0;r<s;r++)a=e.$headerIndexed[r],i.storedSizes[r]=a.is(":visible")?a.width():0},setWidth:function(e,t,r){e.css({width:t,"min-width":r?t:"","max-width":r?t:""})},setWidths:function(e,t,r){var a,s,i=t.resizable_vars,o=c(e.namespace+"_extra_headers"),n=e.$table.children("colgroup").children("col");if((r=r||i.storedSizes||[]).length){for(a=0;a<e.columns;a++)d.resizable.setWidth(e.$headerIndexed[a],r[a],i.overflow),o.length&&(s=o.eq(a).add(n.eq(a)),d.resizable.setWidth(s,r[a],i.overflow));(s=c(e.namespace+"_extra_table")).length&&!d.hasWidget(e.table,"scroller")&&d.resizable.setWidth(s,e.$table.outerWidth(),i.overflow)}},setHandlePosition:function(s,i){var o,n=s.$table.height(),e=i.$resizable_container.children(),l=Math.floor(e.width()/2);d.hasWidget(s.table,"scroller")&&(n=0,s.$table.closest("."+d.css.scrollerWrap).children().each(function(){var e=c(this);n+=(e.filter('[style*="height"]').length?e:e.children("table")).height()})),!i.resizable_includeFooter&&s.$table.children("tfoot").length&&(n-=s.$table.children("tfoot").height()),o=3.3<=parseFloat(c.fn.jquery)?0:s.$table.position().left,e.each(function(){var e=c(this),t=parseInt(e.attr("data-column"),10),r=s.columns-1,a=e.data("header");a&&(!a.is(":visible")||!i.resizable_addLastColumn&&d.resizable.checkVisibleColumns(s,t)?e.hide():(t<r||t===r&&i.resizable_addLastColumn)&&e.css({display:"inline-block",height:n,left:a.position().left-o+a.outerWidth()-l}))})},checkVisibleColumns:function(e,t){for(var r=0,a=t+1;a<e.columns;a++)r+=e.$headerIndexed[a].is(":visible")?1:0;return 0===r},toggleTextSelection:function(e,t,r){e=e.namespace+"tsresize";t.resizable_vars.disabled=r,c("body").toggleClass(d.css.resizableNoSelect,r),r?c("body").attr("unselectable","on").bind("selectstart"+e,!1):c("body").removeAttr("unselectable").unbind("selectstart"+e)},bindings:function(i,o){var e=i.namespace+"tsresize";o.$resizable_container.children().bind("mousedown",function(e){var t=o.resizable_vars,r=c(i.namespace+"_extra_headers"),a=c(e.target).data("header"),s=parseInt(a.attr("data-column"),10);t.$target=a=a.add(r.filter('[data-column="'+s+'"]')),t.target=s,t.$next=e.shiftKey||o.resizable_targetLast?a.parent().children().not(".resizable-false").filter(":last"):a.nextAll(":not(.resizable-false)").eq(0),s=parseInt(t.$next.attr("data-column"),10),t.$next=t.$next.add(r.filter('[data-column="'+s+'"]')),t.next=s,t.mouseXPosition=e.pageX,d.resizable.updateStoredSizes(i,o),d.resizable.toggleTextSelection(i,o,!0)}),c(document).bind("mousemove"+e,function(e){var t=o.resizable_vars;t.disabled&&0!==t.mouseXPosition&&t.$target&&(o.resizable_throttle?(clearTimeout(t.timer),t.timer=setTimeout(function(){d.resizable.mouseMove(i,o,e)},isNaN(o.resizable_throttle)?5:o.resizable_throttle)):d.resizable.mouseMove(i,o,e))}).bind("mouseup"+e,function(){o.resizable_vars.disabled&&(d.resizable.toggleTextSelection(i,o,!1),d.resizable.stopResize(i,o),d.resizable.setHandlePosition(i,o))}),c(t).bind("resize"+e+" resizeEnd"+e,function(){d.resizable.setHandlePosition(i,o)}),i.$table.bind("columnUpdate pagerComplete resizableUpdate ".split(" ").join(e+" "),function(){d.resizable.setHandlePosition(i,o)}).bind("resizableReset"+e,function(){d.resizableReset(i.table)}).find("thead:first").add(c(i.namespace+"_extra_table").find("thead:first")).bind("contextmenu"+e,function(){var e=0===o.resizable_vars.storedSizes.length;return d.resizableReset(i.table),d.resizable.setHandlePosition(i,o),o.resizable_vars.storedSizes=[],e})},mouseMove:function(e,t,r){if(0!==t.resizable_vars.mouseXPosition&&t.resizable_vars.$target){var a,s=0,i=t.resizable_vars,o=i.$next,n=i.storedSizes[i.target],l=r.pageX-i.mouseXPosition;if(i.overflow){if(0<n+l){for(i.storedSizes[i.target]+=l,d.resizable.setWidth(i.$target,i.storedSizes[i.target],!0),a=0;a<e.columns;a++)s+=i.storedSizes[a];d.resizable.setWidth(e.$table.add(c(e.namespace+"_extra_table")),s)}o.length||(i.$wrap[0].scrollLeft=e.$table.width())}else i.fullWidth?(i.storedSizes[i.target]+=l,i.storedSizes[i.next]-=l):i.storedSizes[i.target]+=l,d.resizable.setWidths(e,t);i.mouseXPosition=r.pageX,e.$table.triggerHandler("stickyHeadersUpdate")}},stopResize:function(e,t){var r=t.resizable_vars;d.resizable.updateStoredSizes(e,t),r.useStorage&&(d.storage(e.table,d.css.resizableStorage,r.storedSizes),d.storage(e.table,"tablesorter-table-resized-width",e.$table.width())),r.mouseXPosition=0,r.$target=r.$next=null,e.$table.triggerHandler("stickyHeadersUpdate"),e.$table.triggerHandler("resizableComplete")}},d.addWidget({id:"resizable",priority:40,options:{resizable:!0,resizable_addLastColumn:!1,resizable_includeFooter:!0,resizable_widths:[],resizable_throttle:!1,resizable_targetLast:!1},init:function(e,t,r,a){d.resizable.init(r,a)},format:function(e,t,r){d.resizable.setHandlePosition(t,r)},remove:function(e,t,r,a){var s;r.$resizable_container&&(s=t.namespace+"tsresize",t.$table.add(c(t.namespace+"_extra_table")).removeClass("hasResizable").children("thead").unbind("contextmenu"+s),r.$resizable_container.remove(),d.resizable.toggleTextSelection(t,r,!1),d.resizableReset(e,a),c(document).unbind("mousemove"+s+" mouseup"+s))}}),d.resizableReset=function(i,o){c(i).each(function(){var e,t,r=this.config,a=r&&r.widgetOptions,s=a.resizable_vars;if(i&&r&&r.$headerIndexed.length){for(s.overflow&&s.tableWidth&&(d.resizable.setWidth(r.$table,s.tableWidth,!0),s.useStorage)&&d.storage(i,"tablesorter-table-resized-width",s.tableWidth),e=0;e<r.columns;e++)t=r.$headerIndexed[e],a.resizable_widths&&a.resizable_widths[e]?d.resizable.setWidth(t,a.resizable_widths[e],s.overflow):t.hasClass("resizable-false")||d.resizable.setWidth(t,"",s.overflow);r.$table.triggerHandler("stickyHeadersUpdate"),d.storage&&!o&&d.storage(this,d.css.resizableStorage,[])}})}}(e,window),function(t){"use strict";var l=t.tablesorter||{};function c(e){e=l.storage(e.table,"tablesorter-savesort");return e&&e.hasOwnProperty("sortList")&&t.isArray(e.sortList)?e.sortList:[]}function d(e,t){return(t||c(e)).join(",")!==e.sortList.join(",")}l.addWidget({id:"saveSort",priority:20,options:{saveSort:!0},init:function(e,t,r,a){t.format(e,r,a,!0)},format:function(t,e,r,a){var s,i=e.$table,r=!1!==r.saveSort,o={sortList:e.sortList},n=l.debug(e,"saveSort");n&&(s=new Date),i.hasClass("hasSaveSort")?r&&t.hasInitialized&&l.storage&&d(e)&&(l.storage(t,"tablesorter-savesort",o),n)&&console.log("saveSort >> Saving last sort: "+e.sortList+l.benchmark(s)):(i.addClass("hasSaveSort"),o="",l.storage&&(o=c(e),n&&console.log('saveSort >> Last sort loaded: "'+o+'"'+l.benchmark(s)),i.bind("saveSortReset",function(e){e.stopPropagation(),l.storage(t,"tablesorter-savesort","")})),a&&o&&0<o.length?e.sortList=o:t.hasInitialized&&o&&0<o.length&&d(e,o)&&l.sortOn(e,o))},remove:function(e,t){t.$table.removeClass("hasSaveSort"),l.storage&&l.storage(e,"tablesorter-savesort","")}})}(e),e.tablesorter});return jQuery;}));
```
