## Supplementary Data SD1 for "Inferring binding specificities of human transcription factors with the wisdom of crowds": jquery.tablesorter.min.js

```
!function(e){"function"==typeof define&&define.amd?define(["jquery"],e):"object"==typeof module&&"object"==typeof module.exports?module.exports=e(require("jquery")):e(jQuery)}(function(e){return function(A){"use strict";var L=A.tablesorter={version:"2.32.0",parsers:[],widgets:[],defaults:{theme:"default",widthFixed:!1,showProcessing:!1,headerTemplate:"{content}",onRenderTemplate:null,onRenderHeader:null,cancelSelection:!0,tabIndex:!0,dateFormat:"mmddyyyy",sortMultiSortKey:"shiftKey",sortResetKey:"ctrlKey",usNumberFormat:!0,delayInit:!1,serverSideSorting:!1,resort:!0,headers:null,ignoreCase:!0,sortForce:null,sortList:[],sortAppend:null,sortStable:!1,sortInitialOrder:"asc",sortLocaleCompare:!1,sortReset:!1,sortRestart:!1,emptyTo:"bottom",stringTo:"max",duplicateSpan:!0,textExtraction:"basic",textAttribute:"data-text",textSorter:null,numberSorter:null,initWidgets:!0,widgetClass:"widget-{name}",widgets:[],widgetOptions:{zebra:["even","odd"]},initialized:null,tableClass:"",cssAsc:"",cssDesc:"",cssNone:"",cssHeader:"",cssHeaderRow:"",cssProcessing:"",cssChildRow:"tablesorter-childRow",cssInfoBlock:"tablesorter-infoOnly",cssNoSort:"tablesorter-noSort",cssIgnoreRow:"tablesorter-ignoreRow",cssIcon:"tablesorter-icon",cssIconNone:"",cssIconAsc:"",cssIconDesc:"",cssIconDisabled:"",pointerClick:"click",pointerDown:"mousedown",pointerUp:"mouseup",selectorHeaders:"> thead th, > thead td",selectorSort:"th, td",selectorRemove:".remove-me",debug:!1,headerList:[],empties:{},strings:{},parsers:[],globalize:0,imgAttr:0},css:{table:"tablesorter",cssHasChild:"tablesorter-hasChildRow",childRow:"tablesorter-childRow",colgroup:"tablesorter-colgroup",header:"tablesorter-header",headerRow:"tablesorter-headerRow",headerIn:"tablesorter-header-inner",icon:"tablesorter-icon",processing:"tablesorter-processing",sortAsc:"tablesorter-headerAsc",sortDesc:"tablesorter-headerDesc",sortNone:"tablesorter-headerUnSorted"},language:{sortAsc:"Ascending sort applied, ",sortDesc:"Descending sort applied, ",sortNone:"No sort applied, ",sortDisabled:"sorting is disabled",nextAsc:"activate to apply an ascending sort",nextDesc:"activate to apply a descending sort",nextNone:"activate to remove the sort"},regex:{templateContent:/\{content\}/g,templateIcon:/\{icon\}/g,templateName:/\{name\}/i,spaces:/\s+/g,nonWord:/\W/g,formElements:/(input|select|button|textarea)/i,chunk:/(^([+\-]?(?:\d*)(?:\.\d*)?(?:[eE][+\-]?\d+)?)?$|^0x[0-9a-f]+$|\d+)/gi,chunks:/(^\\0|\\0$)/,hex:/^0x[0-9a-f]+$/i,comma:/,/g,digitNonUS:/[\s|\.]/g,digitNegativeTest:/^\s*\([.\d]+\)/,digitNegativeReplace:/^\s*\(([.\d]+)\)/,digitTest:/^[\-+(]?\d+[)]?$/,digitReplace:/[,.'"\s]/g},string:{max:1,min:-1,emptymin:1,emptymax:-1,zero:0,none:0,null:0,top:!0,bottom:!1},keyCodes:{enter:13},dates:{},instanceMethods:{},setup:function(t,r){var e,o,s,a;t&&t.tHead&&0!==t.tBodies.length&&!0!==t.hasInitialized?(e="",o=A(t),s=A.metadata,t.hasInitialized=!1,t.isProcessing=!0,t.config=r,A.data(t,"tablesorter",r),L.debug(r,"core")&&(console[console.group?"group":"log"]("Initializing tablesorter v"+L.version),A.data(t,"startoveralltimer",new Date)),r.supportsDataObject=((a=A.fn.jquery.split("."))[0]=parseInt(a[0],10),1<a[0]||1===a[0]&&4<=parseInt(a[1],10)),r.emptyTo=r.emptyTo.toLowerCase(),r.stringTo=r.stringTo.toLowerCase(),r.last={sortList:[],clickedIndex:-1},/tablesorter\-/.test(o.attr("class"))||(e=""!==r.theme?" tablesorter-"+r.theme:""),r.namespace?r.namespace="."+r.namespace.replace(L.regex.nonWord,""):r.namespace=".tablesorter"+Math.random().toString(16).slice(2),r.table=t,r.$table=o.addClass(L.css.table+" "+r.tableClass+e+" "+r.namespace.slice(1)).attr("role","grid"),r.$headers=o.find(r.selectorHeaders),r.$table.children().children("tr").attr("role","row"),r.$tbodies=o.children("tbody:not(."+r.cssInfoBlock+")").attr({"aria-live":"polite","aria-relevant":"all"}),r.$table.children("caption").length&&((e=r.$table.children("caption")[0]).id||(e.id=r.namespace.slice(1)+"caption"),r.$table.attr("aria-labelledby",e.id)),r.widgetInit={},r.textExtraction=r.$table.attr("data-text-extraction")||r.textExtraction||"basic",L.buildHeaders(r),L.fixColumnWidth(t),L.addWidgetFromClass(t),L.applyWidgetOptions(t),L.setupParsers(r),r.totalRows=0,r.debug&&L.validateOptions(r),r.delayInit||L.buildCache(r),L.bindEvents(t,r.$headers,!0),L.bindMethods(r),r.supportsDataObject&&void 0!==o.data().sortlist?r.sortList=o.data().sortlist:s&&o.metadata()&&o.metadata().sortlist&&(r.sortList=o.metadata().sortlist),L.applyWidget(t,!0),0<r.sortList.length?(r.last.sortList=r.sortList,L.sortOn(r,r.sortList,{},!r.initWidgets)):(L.setHeadersCss(r),r.initWidgets&&L.applyWidget(t,!1)),r.showProcessing&&o.unbind("sortBegin"+r.namespace+" sortEnd"+r.namespace).bind("sortBegin"+r.namespace+" sortEnd"+r.namespace,function(e){clearTimeout(r.timerProcessing),L.isProcessing(t),"sortBegin"===e.type&&(r.timerProcessing=setTimeout(function(){L.isProcessing(t,!0)},500))}),t.hasInitialized=!0,t.isProcessing=!1,L.debug(r,"core")&&(console.log("Overall initialization time:"+L.benchmark(A.data(t,"startoveralltimer"))),L.debug(r,"core"))&&console.groupEnd&&console.groupEnd(),o.triggerHandler("tablesorter-initialized",t),"function"==typeof r.initialized&&r.initialized(t)):L.debug(r,"core")&&(t.hasInitialized?console.warn("Stopping initialization. Tablesorter has already been initialized"):console.error("Stopping initialization! No table, thead or tbody",t))},bindMethods:function(t){var e=t.$table,r=t.namespace,o="sortReset update updateRows updateAll updateHeaders addRows updateCell updateComplete sorton appendCache updateCache applyWidgetId applyWidgets refreshWidgets destroy mouseup mouseleave ".split(" ").join(r+" ");e.unbind(o.replace(L.regex.spaces," ")).bind("sortReset"+r,function(e,t){e.stopPropagation(),L.sortReset(this.config,function(e){e.isApplyingWidgets?setTimeout(function(){L.applyWidget(e,"",t)},100):L.applyWidget(e,"",t)})}).bind("updateAll"+r,function(e,t,r){e.stopPropagation(),L.updateAll(this.config,t,r)}).bind("update"+r+" updateRows"+r,function(e,t,r){e.stopPropagation(),L.update(this.config,t,r)}).bind("updateHeaders"+r,function(e,t){e.stopPropagation(),L.updateHeaders(this.config,t)}).bind("updateCell"+r,function(e,t,r,o){e.stopPropagation(),L.updateCell(this.config,t,r,o)}).bind("addRows"+r,function(e,t,r,o){e.stopPropagation(),L.addRows(this.config,t,r,o)}).bind("updateComplete"+r,function(){this.isUpdating=!1}).bind("sorton"+r,function(e,t,r,o){e.stopPropagation(),L.sortOn(this.config,t,r,o)}).bind("appendCache"+r,function(e,t,r){e.stopPropagation(),L.appendCache(this.config,r),A.isFunction(t)&&t(this)}).bind("updateCache"+r,function(e,t,r){e.stopPropagation(),L.updateCache(this.config,t,r)}).bind("applyWidgetId"+r,function(e,t){e.stopPropagation(),L.applyWidgetId(this,t)}).bind("applyWidgets"+r,function(e,t){e.stopPropagation(),L.applyWidget(this,!1,t)}).bind("refreshWidgets"+r,function(e,t,r){e.stopPropagation(),L.refreshWidgets(this,t,r)}).bind("removeWidget"+r,function(e,t,r){e.stopPropagation(),L.removeWidget(this,t,r)}).bind("destroy"+r,function(e,t,r){e.stopPropagation(),L.destroy(this,t,r)}).bind("resetToLoadState"+r,function(e){e.stopPropagation(),L.removeWidget(this,!0,!1);e=A.extend(!0,{},t.originalSettings);(t=A.extend(!0,{},L.defaults,e)).originalSettings=e,this.hasInitialized=!1,L.setup(this,t)})},bindEvents:function(e,t,r){var o,s=(e=A(e)[0]).config,a=s.namespace,n=null;!0!==r&&(t.addClass(a.slice(1)+"_extra_headers"),(o=L.getClosest(t,"table")).length)&&"TABLE"===o[0].nodeName&&o[0]!==e&&A(o[0]).addClass(a.slice(1)+"_extra_table"),o=(s.pointerDown+" "+s.pointerUp+" "+s.pointerClick+" sort keyup ").replace(L.regex.spaces," ").split(" ").join(a+" "),t.find(s.selectorSort).add(t.filter(s.selectorSort)).unbind(o).bind(o,function(e,t){var r=A(e.target),o=" "+e.type+" ";if(!(1!==(e.which||e.button)&&!o.match(" "+s.pointerClick+" | sort | keyup ")||" keyup "===o&&e.which!==L.keyCodes.enter||o.match(" "+s.pointerClick+" ")&&void 0!==e.which||o.match(" "+s.pointerUp+" ")&&n!==e.target&&!0!==t))if(o.match(" "+s.pointerDown+" "))n=e.target,"1"===(t=r.jquery.split("."))[0]&&t[1]<4&&e.preventDefault();else{if(n=null,o=L.getClosest(A(this),"."+L.css.header),L.regex.formElements.test(e.target.nodeName)||r.hasClass(s.cssNoSort)||0<r.parents("."+s.cssNoSort).length||o.hasClass("sorter-false")||0<r.parents("button").length)return!s.cancelSelection;s.delayInit&&L.isEmptyObject(s.cache)&&L.buildCache(s),s.last.clickedIndex=o.attr("data-column")||o.index(),(t=s.$headerIndexed[s.last.clickedIndex][0])&&!t.sortDisabled&&L.initSort(s,t,e)}}),s.cancelSelection&&t.attr("unselectable","on").bind("selectstart",!1).css({"user-select":"none",MozUserSelect:"none"})},buildHeaders:function(i){var e,l,t,r;for(i.headerList=[],i.headerContent=[],i.sortVars=[],L.debug(i,"core")&&(t=new Date),i.columns=L.computeColumnIndex(i.$table.children("thead, tfoot").children("tr")),l=i.cssIcon?'<i class="'+(i.cssIcon===L.css.icon?L.css.icon:i.cssIcon+" "+L.css.icon)+'"></i>':"",i.$headers=A(A.map(i.$table.find(i.selectorHeaders),function(e,t){var r,o,s,a,n=A(e);if(!L.getClosest(n,"tr").hasClass(i.cssIgnoreRow))return/(th|td)/i.test(e.nodeName)||(a=L.getClosest(n,"th, td"),n.attr("data-column",a.attr("data-column"))),r=L.getColumnData(i.table,i.headers,t,!0),i.headerContent[t]=n.html(),""===i.headerTemplate||n.find("."+L.css.headerIn).length||(s=i.headerTemplate.replace(L.regex.templateContent,n.html()).replace(L.regex.templateIcon,n.find("."+L.css.icon).length?"":l),i.onRenderTemplate&&(o=i.onRenderTemplate.apply(n,[t,s]))&&"string"==typeof o&&(s=o),n.html('<div class="'+L.css.headerIn+'">'+s+"</div>")),i.onRenderHeader&&i.onRenderHeader.apply(n,[t,i,i.$table]),o=parseInt(n.attr("data-column"),10),e.column=o,a=L.getOrder(L.getData(n,r,"sortInitialOrder")||i.sortInitialOrder),i.sortVars[o]={count:-1,order:a?i.sortReset?[1,0,2]:[1,0]:i.sortReset?[0,1,2]:[0,1],lockedOrder:!1,sortedBy:""},void 0!==(a=L.getData(n,r,"lockedOrder")||!1)&&!1!==a&&(i.sortVars[o].lockedOrder=!0,i.sortVars[o].order=L.getOrder(a)?[1,1]:[0,0]),i.headerList[t]=e,n.addClass(L.css.header+" "+i.cssHeader),L.getClosest(n,"tr").addClass(L.css.headerRow+" "+i.cssHeaderRow).attr("role","row"),i.tabIndex&&n.attr("tabindex",0),e})),i.$headerIndexed=[],r=0;r<i.columns;r++)L.isEmptyObject(i.sortVars[r])&&(i.sortVars[r]={}),e=i.$headers.filter('[data-column="'+r+'"]'),i.$headerIndexed[r]=e.length?(e.not(".sorter-false").length?e.not(".sorter-false"):e).filter(":last"):A();i.$table.find(i.selectorHeaders).attr({scope:"col",role:"columnheader"}),L.updateHeader(i),L.debug(i,"core")&&(console.log("Built headers:"+L.benchmark(t)),console.log(i.$headers))},addInstanceMethods:function(e){A.extend(L.instanceMethods,e)},setupParsers:function(e,t){var r,o,s,a,n,i,l,d,c,g,p,u,f,h,m=e.table,b=0,y=L.debug(e,"core"),w={};if(e.$tbodies=e.$table.children("tbody:not(."+e.cssInfoBlock+")"),0===(h=(f=void 0===t?e.$tbodies:t).length))return y?console.warn("Warning: *Empty table!* Not building a parser cache"):"";for(y&&(u=new Date,console[console.group?"group":"log"]("Detecting parsers for each column")),o={extractors:[],parsers:[]};b<h;){if((r=f[b].rows).length)for(a=e.columns,i=n=0;i<a;i++){if((l=e.$headerIndexed[n])&&l.length&&(d=L.getColumnData(m,e.headers,n),p=L.getParserById(L.getData(l,d,"extractor")),g=L.getParserById(L.getData(l,d,"sorter")),c="false"===L.getData(l,d,"parser"),e.empties[n]=(L.getData(l,d,"empty")||e.emptyTo||(e.emptyToBottom?"bottom":"top")).toLowerCase(),e.strings[n]=(L.getData(l,d,"string")||e.stringTo||"max").toLowerCase(),p=p||!1,g=(g=c?L.getParserById("no-parser"):g)||L.detectParserForColumn(e,r,-1,n),y&&(w["("+n+") "+l.text()]={parser:g.id,extractor:p?p.id:"none",string:e.strings[n],empty:e.empties[n]}),o.parsers[n]=g,o.extractors[n]=p,0<(s=l[0].colSpan-1)))for(n+=s,a+=s;0<s+1;)o.parsers[n-s]=g,o.extractors[n-s]=p,s--;n++}b+=o.parsers.length?h:1}y&&(L.isEmptyObject(w)?console.warn("  No parsers detected!"):console[console.table?"table":"log"](w),console.log("Completed detecting parsers"+L.benchmark(u)),console.groupEnd)&&console.groupEnd(),e.parsers=o.parsers,e.extractors=o.extractors},addParser:function(e){for(var t=L.parsers.length,r=!0,o=0;o<t;o++)L.parsers[o].id.toLowerCase()===e.id.toLowerCase()&&(r=!1);r&&(L.parsers[L.parsers.length]=e)},getParserById:function(e){if("false"!=e)for(var t=L.parsers.length,r=0;r<t;r++)if(L.parsers[r].id.toLowerCase()===e.toString().toLowerCase())return L.parsers[r];return!1},detectParserForColumn:function(e,t,r,o){for(var s,a,n,i=L.parsers.length,l=!1,d="",c=L.debug(e,"core"),g=!0;""===d&&g;)(n=t[++r])&&r<50?n.className.indexOf(L.cssIgnoreRow)<0&&(l=t[r].cells[o],d=L.getElementText(e,l,o),a=A(l),c)&&console.log("Checking if value was empty on row "+r+", column: "+o+': "'+d+'"'):g=!1;for(;0<=--i;)if((s=L.parsers[i])&&"text"!==s.id&&s.is&&s.is(d,e.table,l,a))return s;return L.getParserById("text")},getElementText:function(e,t,r){var o,s,a;return t?(s=e.textExtraction||"",a=t.jquery?t:A(t),"string"==typeof s?"basic"===s&&void 0!==(o=a.attr(e.textAttribute))?A.trim(o):A.trim(t.textContent||a.text()):"function"==typeof s?A.trim(s(a[0],e.table,r)):"function"==typeof(o=L.getColumnData(e.table,s,r))?A.trim(o(a[0],e.table,r)):A.trim(a[0].textContent||a.text())):""},getParsedText:function(e,t,r,o){var s=""+(o=void 0===o?L.getElementText(e,t,r):o),a=e.parsers[r],n=e.extractors[r];return s=a&&(n&&"function"==typeof n.format&&(o=n.format(o,e.table,t,r)),s="no-parser"===a.id?"":a.format(""+o,e.table,t,r),e.ignoreCase)&&"string"==typeof s?s.toLowerCase():s},buildCache:function(e,t,r){var o,s,a,n,i,l,d,c,g,p,u,f,h,m,b,y,w,x,C,v,$,I,D=e.table,R=e.parsers,T=L.debug(e,"core");if(e.$tbodies=e.$table.children("tbody:not(."+e.cssInfoBlock+")"),d=void 0===r?e.$tbodies:r,e.cache={},e.totalRows=0,!R)return T?console.warn("Warning: *Empty table!* Not building a cache"):"";for(T&&(f=new Date),e.showProcessing&&L.isProcessing(D,!0),l=0;l<d.length;l++){for(y=[],o=e.cache[l]={normalized:[]},h=d[l]&&d[l].rows.length||0,n=0;n<h;++n)if(m={child:[],raw:[]},g=[],!(c=A(d[l].rows[n])).hasClass(e.selectorRemove.slice(1)))if(c.hasClass(e.cssChildRow)&&0!==n)for($=o.normalized.length-1,(b=o.normalized[$][e.columns]).$row=b.$row.add(c),c.prev().hasClass(e.cssChildRow)||c.prev().addClass(L.css.cssHasChild),p=c.children("th, td"),$=b.child.length,b.child[$]=[],v=e.columns,i=x=0;i<v;i++)(u=p[i])&&(b.child[$][i]=L.getParsedText(e,u,i),0<(w=p[i].colSpan-1))&&(x+=w,v+=w),x++;else{for(m.$row=c,m.order=n,v=e.columns,i=x=0;i<v;++i){if((u=c[0].cells[i])&&x<e.columns&&(!(C=void 0!==R[x])&&T&&console.warn("No parser found for row: "+n+", column: "+i+'; cell containing: "'+A(u).text()+'"; does it have a header?'),s=L.getElementText(e,u,x),m.raw[x]=s,a=L.getParsedText(e,u,x,s),g[x]=a,C&&"numeric"===(R[x].type||"").toLowerCase()&&(y[x]=Math.max(Math.abs(a)||0,y[x]||0)),0<(w=u.colSpan-1))){for(I=0;I<=w;)a=e.duplicateSpan||0===I?a:"string"!=typeof e.textExtraction&&L.getElementText(e,u,x+I)||"",m.raw[x+I]=a,g[x+I]=a,I++;x+=w,v+=w}x++}g[e.columns]=m,o.normalized[o.normalized.length]=g}o.colMax=y,e.totalRows+=o.normalized.length}if(e.showProcessing&&L.isProcessing(D),T){for($=Math.min(5,e.cache[0].normalized.length),console[console.group?"group":"log"]("Building cache for "+e.totalRows+" rows (showing "+$+" rows in log) and "+e.columns+" columns"+L.benchmark(f)),s={},i=0;i<e.columns;i++)for(x=0;x<$;x++)s["row: "+x]||(s["row: "+x]={}),s["row: "+x][e.$headerIndexed[i].text()]=e.cache[0].normalized[x][i];console[console.table?"table":"log"](s),console.groupEnd&&console.groupEnd()}A.isFunction(t)&&t(D)},getColumnText:function(e,t,r,o){var s,a,n,i,l,d,c,g,p,u,f="function"==typeof r,h="all"===t,m={raw:[],parsed:[],$cell:[]},b=(e=A(e)[0]).config;if(!L.isEmptyObject(b)){for(l=b.$tbodies.length,s=0;s<l;s++)for(d=(n=b.cache[s].normalized).length,a=0;a<d;a++)i=n[a],o&&!i[b.columns].$row.is(o)||(u=!0,g=h?i.slice(0,b.columns):i[t],i=i[b.columns],c=h?i.raw:i.raw[t],p=h?i.$row.children():i.$row.children().eq(t),!1!==(u=f?r({tbodyIndex:s,rowIndex:a,parsed:g,raw:c,$row:i.$row,$cell:p}):u)&&(m.parsed[m.parsed.length]=g,m.raw[m.raw.length]=c,m.$cell[m.$cell.length]=p));return m}L.debug(b,"core")&&console.warn("No cache found - aborting getColumnText function!")},setHeadersCss:function(s){function e(e,t){e.removeClass(n).addClass(i[t]).attr("aria-sort",d[t]).find("."+L.css.icon).removeClass(l[2]).addClass(l[t])}var t,r,o=s.sortList,a=o.length,n=L.css.sortNone+" "+s.cssNone,i=[L.css.sortAsc+" "+s.cssAsc,L.css.sortDesc+" "+s.cssDesc],l=[s.cssIconAsc,s.cssIconDesc,s.cssIconNone],d=["ascending","descending"],c=s.$table.find("tfoot tr").children("td, th").add(A(s.namespace+"_extra_headers")).removeClass(i.join(" ")),g=s.$headers.add(A("thead "+s.namespace+"_extra_headers")).removeClass(i.join(" ")).addClass(n).attr("aria-sort","none").find("."+L.css.icon).removeClass(l.join(" ")).end();for(g.not(".sorter-false").find("."+L.css.icon).addClass(l[2]),s.cssIconDisabled&&g.filter(".sorter-false").find("."+L.css.icon).addClass(s.cssIconDisabled),t=0;t<a;t++)if(2!==o[t][1]){if((g=(g=s.$headers.filter(function(e){for(var t=!0,e=s.$headers.eq(e),r=parseInt(e.attr("data-column"),10),o=r+L.getClosest(e,"th, td")[0].colSpan;r<o;r++)t=t||!1;return t})).not(".sorter-false").filter('[data-column="'+o[t][0]+'"]'+(1===a?":last":""))).length)for(r=0;r<g.length;r++)g[r].sortDisabled||e(g.eq(r),o[t][1]);c.length&&e(c.filter('[data-column="'+o[t][0]+'"]'),o[t][1])}for(a=s.$headers.length,t=0;t<a;t++)L.setColumnAriaLabel(s,s.$headers.eq(t))},getClosest:function(e,t){return A.fn.closest?e.closest(t):e.is(t)?e:e.parents(t).filter(":first")},setColumnAriaLabel:function(e,t,r){var o,s;t.length&&(o=parseInt(t.attr("data-column"),10),e=e.sortVars[o],o=t.hasClass(L.css.sortAsc)?"sortAsc":t.hasClass(L.css.sortDesc)?"sortDesc":"sortNone",s=A.trim(t.text())+": "+L.language[o],t.hasClass("sorter-false")||!1===r?s+=L.language.sortDisabled:(o=(e.count+1)%e.order.length,r=e.order[o],s+=L.language[0===r?"nextAsc":1===r?"nextDesc":"nextNone"]),t.attr("aria-label",s),e.sortedBy?t.attr("data-sortedBy",e.sortedBy):t.removeAttr("data-sortedBy"))},updateHeader:function(e){for(var t,r,o=e.table,s=e.$headers.length,a=0;a<s;a++)t=e.$headers.eq(a),r=L.getColumnData(o,e.headers,a,!0),r="false"===L.getData(t,r,"sorter")||"false"===L.getData(t,r,"parser"),L.setColumnSort(e,t,r)},setColumnSort:function(e,t,r){var o=e.table.id;t[0].sortDisabled=r,t[r?"addClass":"removeClass"]("sorter-false").attr("aria-disabled",""+r),e.tabIndex&&(r?t.removeAttr("tabindex"):t.attr("tabindex","0")),o&&(r?t.removeAttr("aria-controls"):t.attr("aria-controls",o))},updateHeaderSortCount:function(e,t){var r,o,s,a,n,i,l,d=t||e.sortList,c=d.length;for(e.sortList=[],s=0;s<c;s++)if(i=d[s],(r=parseInt(i[0],10))<e.columns){switch(e.sortVars[r].order||(l=L.getOrder(e.sortInitialOrder)?e.sortReset?[1,0,2]:[1,0]:e.sortReset?[0,1,2]:[0,1],e.sortVars[r].order=l,e.sortVars[r].count=0),l=e.sortVars[r].order,o=(o=(""+i[1]).match(/^(1|d|s|o|n)/))?o[0]:""){case"1":case"d":o=1;break;case"s":o=a||0;break;case"o":o=0===(n=l[(a||0)%l.length])?1:1===n?0:2;break;case"n":o=l[++e.sortVars[r].count%l.length];break;default:o=0}a=0===s?o:a,i=[r,parseInt(o,10)||0],e.sortList[e.sortList.length]=i,o=A.inArray(i[1],l),e.sortVars[r].count=0<=o?o:i[1]%l.length}},updateAll:function(e,t,r){var o=e.table;o.isUpdating=!0,L.refreshWidgets(o,!0,!0),L.buildHeaders(e),L.bindEvents(o,e.$headers,!0),L.bindMethods(e),L.commonUpdate(e,t,r)},update:function(e,t,r){e.table.isUpdating=!0,L.updateHeader(e),L.commonUpdate(e,t,r)},updateHeaders:function(e,t){e.table.isUpdating=!0,L.buildHeaders(e),L.bindEvents(e.table,e.$headers,!0),L.resortComplete(e,t)},updateCell:function(e,t,r,o){if(A(t).closest("tr").hasClass(e.cssChildRow))console.warn('Tablesorter Warning! "updateCell" for child row content has been disabled, use "update" instead');else if(L.isEmptyObject(e.cache))L.updateHeader(e),L.commonUpdate(e,r,o);else{e.table.isUpdating=!0,e.$table.find(e.selectorRemove).remove();var s,a,n,i,l=e.$tbodies,d=A(t),c=l.index(L.getClosest(d,"tbody")),g=e.cache[c],p=L.getClosest(d,"tr");if(t=d[0],l.length&&0<=c){if(l=l.eq(c).find("tr").not("."+e.cssChildRow).index(p),c=g.normalized[l],(i=p[0].cells.length)!==e.columns)for(s=!1,a=n=0;a<i;a++)s||p[0].cells[a]===t?s=!0:n+=p[0].cells[a].colSpan;else n=d.index();s=L.getElementText(e,t,n),c[e.columns].raw[n]=s,s=L.getParsedText(e,t,n,s),c[n]=s,"numeric"===(e.parsers[n].type||"").toLowerCase()&&(g.colMax[n]=Math.max(Math.abs(s)||0,g.colMax[n]||0)),!1!==(s="undefined"!==r?r:e.resort)?L.checkResort(e,s,o):L.resortComplete(e,o)}else L.debug(e,"core")&&console.error("updateCell aborted, tbody missing or not within the indicated table"),e.table.isUpdating=!1}},addRows:function(e,t,r,o){var s,a,n,i,l,d,c,g,p,u,f,h="string"==typeof t&&1===e.$tbodies.length&&/<tr/.test(t||""),m=e.table;if(h)t=A(t),e.$tbodies.append(t);else if(!(t&&t instanceof A&&L.getClosest(t,"table")[0]===e.table))return L.debug(e,"core")&&console.error("addRows method requires (1) a jQuery selector reference to rows that have already been added to the table, or (2) row HTML string to be added to a table with only one tbody"),!1;if(m.isUpdating=!0,L.isEmptyObject(e.cache))L.updateHeader(e),L.commonUpdate(e,r,o);else{for(n=t.filter("tr").attr("role","row").length,s=e.$tbodies.index(t.parents("tbody").filter(":first")),e.parsers&&e.parsers.length||L.setupParsers(e),a=0;a<n;a++){for(c=0,l=t[a].cells.length,d=e.cache[s].normalized.length,p=[],g={child:[],raw:[],$row:t.eq(a),order:d},i=0;i<l;i++)u=t[a].cells[i],f=L.getElementText(e,u,c),g.raw[c]=f,f=L.getParsedText(e,u,c,f),p[c]=f,"numeric"===(e.parsers[c].type||"").toLowerCase()&&(e.cache[s].colMax[c]=Math.max(Math.abs(f)||0,e.cache[s].colMax[c]||0)),0<(f=u.colSpan-1)&&(c+=f),c++;p[e.columns]=g,e.cache[s].normalized[d]=p}L.checkResort(e,r,o)}},updateCache:function(e,t,r){e.parsers&&e.parsers.length||L.setupParsers(e,r),L.buildCache(e,t,r)},appendCache:function(e,t){var r,o,s,a,n,i,l,d=e.table,c=e.$tbodies,g=[],p=e.cache;if(L.isEmptyObject(p))return e.appender?e.appender(d,g):d.isUpdating?e.$table.triggerHandler("updateComplete",d):"";for(L.debug(e,"core")&&(l=new Date),i=0;i<c.length;i++)if((s=c.eq(i)).length){for(a=L.processTbody(d,s,!0),o=(r=p[i].normalized).length,n=0;n<o;n++)g[g.length]=r[n][e.columns].$row,e.appender&&(!e.pager||e.pager.removeRows||e.pager.ajax)||a.append(r[n][e.columns].$row);L.processTbody(d,a,!1)}e.appender&&e.appender(d,g),L.debug(e,"core")&&console.log("Rebuilt table"+L.benchmark(l)),t||e.appender||L.applyWidget(d),d.isUpdating&&e.$table.triggerHandler("updateComplete",d)},commonUpdate:function(e,t,r){e.$table.find(e.selectorRemove).remove(),L.setupParsers(e),L.buildCache(e),L.checkResort(e,t,r)},initSort:function(t,e,r){if(t.table.isUpdating)return setTimeout(function(){L.initSort(t,e,r)},50);var o,s,a,n,i,l,d,c=!r[t.sortMultiSortKey],g=t.table,p=t.$headers.length,u=L.getClosest(A(e),"th, td"),f=parseInt(u.attr("data-column"),10),h="mouseup"===r.type?"user":r.type,m=t.sortVars[f].order,u=u[0];if(t.$table.triggerHandler("sortStart",g),l=(t.sortVars[f].count+1)%m.length,t.sortVars[f].count=r[t.sortResetKey]?2:l,t.sortRestart)for(a=0;a<p;a++)d=t.$headers.eq(a),f!==(l=parseInt(d.attr("data-column"),10))&&(c||d.hasClass(L.css.sortNone))&&(t.sortVars[l].count=-1);if(c){if(A.each(t.sortVars,function(e){t.sortVars[e].sortedBy=""}),t.sortList=[],t.last.sortList=[],null!==t.sortForce)for(o=t.sortForce,s=0;s<o.length;s++)o[s][0]!==f&&(t.sortList[t.sortList.length]=o[s],t.sortVars[o[s][0]].sortedBy="sortForce");if((n=m[t.sortVars[f].count])<2&&(t.sortList[t.sortList.length]=[f,n],t.sortVars[f].sortedBy=h,1<u.colSpan))for(s=1;s<u.colSpan;s++)t.sortList[t.sortList.length]=[f+s,n],t.sortVars[f+s].count=A.inArray(n,m),t.sortVars[f+s].sortedBy=h}else if(t.sortList=A.extend([],t.last.sortList),0<=L.isValueInArray(f,t.sortList))for(t.sortVars[f].sortedBy=h,s=0;s<t.sortList.length;s++)(l=t.sortList[s])[0]===f&&(l[1]=m[t.sortVars[f].count],2===l[1])&&(t.sortList.splice(s,1),t.sortVars[f].count=-1);else if(n=m[t.sortVars[f].count],t.sortVars[f].sortedBy=h,n<2&&(t.sortList[t.sortList.length]=[f,n],1<u.colSpan))for(s=1;s<u.colSpan;s++)t.sortList[t.sortList.length]=[f+s,n],t.sortVars[f+s].count=A.inArray(n,m),t.sortVars[f+s].sortedBy=h;if(t.last.sortList=A.extend([],t.sortList),t.sortList.length&&t.sortAppend&&(o=A.isArray(t.sortAppend)?t.sortAppend:t.sortAppend[t.sortList[0][0]],!L.isEmptyObject(o)))for(s=0;s<o.length;s++)if(o[s][0]!==f&&L.isValueInArray(o[s][0],t.sortList)<0){if(i=(""+(n=o[s][1])).match(/^(a|d|s|o|n)/))switch(l=t.sortList[0][1],i[0]){case"d":n=1;break;case"s":n=l;break;case"o":n=0===l?1:0;break;case"n":n=(l+1)%m.length;break;default:n=0}t.sortList[t.sortList.length]=[o[s][0],n],t.sortVars[o[s][0]].sortedBy="sortAppend"}t.$table.triggerHandler("sortBegin",g),setTimeout(function(){L.setHeadersCss(t),L.multisort(t),L.appendCache(t),t.$table.triggerHandler("sortBeforeEnd",g),t.$table.triggerHandler("sortEnd",g)},1)},multisort:function(n){var e,t,i,r,l=n.table,d=[],c=0,g=n.textSorter||"",p=n.sortList,u=p.length,o=n.$tbodies.length;if(!n.serverSideSorting&&!L.isEmptyObject(n.cache)){if(L.debug(n,"core")&&(t=new Date),"object"==typeof g)for(i=n.columns;i--;)"function"==typeof(r=L.getColumnData(l,g,i))&&(d[i]=r);for(e=0;e<o;e++)i=n.cache[e].colMax,n.cache[e].normalized.sort(function(e,t){for(var r,o,s,a=0;a<u;a++){if(r=p[a][0],o=p[a][1],c=0===o,n.sortStable&&e[r]===t[r]&&1===u)return e[n.columns].order-t[n.columns].order;if(o=(o=/n/i.test(L.getSortType(n.parsers,r)))&&n.strings[r]?(o="boolean"==typeof L.string[n.strings[r]]?(c?1:-1)*(L.string[n.strings[r]]?-1:1):n.strings[r]&&L.string[n.strings[r]]||0,n.numberSorter?n.numberSorter(e[r],t[r],c,i[r],l):L["sortNumeric"+(c?"Asc":"Desc")](e[r],t[r],o,i[r],r,n)):(o=c?e:t,s=c?t:e,"function"==typeof g?g(o[r],s[r],c,r,l):"function"==typeof d[r]?d[r](o[r],s[r],c,r,l):L["sortNatural"+(c?"Asc":"Desc")](e[r]||"",t[r]||"",r,n)))return o}return e[n.columns].order-t[n.columns].order});L.debug(n,"core")&&console.log("Applying sort "+p.toString()+L.benchmark(t))}},resortComplete:function(e,t){e.table.isUpdating&&e.$table.triggerHandler("updateComplete",e.table),A.isFunction(t)&&t(e.table)},checkResort:function(e,t,r){var o=A.isArray(t)?t:e.sortList;!1===(void 0===t?e.resort:t)||e.serverSideSorting||e.table.isProcessing?(L.resortComplete(e,r),L.applyWidget(e.table,!1)):o.length?L.sortOn(e,o,function(){L.resortComplete(e,r)},!0):L.sortReset(e,function(){L.resortComplete(e,r),L.applyWidget(e.table,!1)})},sortOn:function(e,t,r,o){var s,a=e.table;for(e.$table.triggerHandler("sortStart",a),s=0;s<e.columns;s++)e.sortVars[s].sortedBy=-1<L.isValueInArray(s,t)?"sorton":"";L.updateHeaderSortCount(e,t),L.setHeadersCss(e),e.delayInit&&L.isEmptyObject(e.cache)&&L.buildCache(e),e.$table.triggerHandler("sortBegin",a),L.multisort(e),L.appendCache(e,o),e.$table.triggerHandler("sortBeforeEnd",a),e.$table.triggerHandler("sortEnd",a),L.applyWidget(a),A.isFunction(r)&&r(a)},sortReset:function(e,t){var r;for(e.sortList=[],r=0;r<e.columns;r++)e.sortVars[r].count=-1,e.sortVars[r].sortedBy="";L.setHeadersCss(e),L.multisort(e),L.appendCache(e),A.isFunction(t)&&t(e.table)},getSortType:function(e,t){return e&&e[t]&&e[t].type||""},getOrder:function(e){return/^d/i.test(e)||1===e},sortNatural:function(e,t){if(e!==t){e=(e||"").toString(),t=(t||"").toString();var r,o,s,a,n,i,l=L.regex;if(l.hex.test(t)){if((r=parseInt(e.match(l.hex),16))<(o=parseInt(t.match(l.hex),16)))return-1;if(o<r)return 1}for(r=e.replace(l.chunk,"\\0$1\\0").replace(l.chunks,"").split("\\0"),o=t.replace(l.chunk,"\\0$1\\0").replace(l.chunks,"").split("\\0"),i=Math.max(r.length,o.length),n=0;n<i;n++){if(s=isNaN(r[n])?r[n]||0:parseFloat(r[n])||0,a=isNaN(o[n])?o[n]||0:parseFloat(o[n])||0,isNaN(s)!==isNaN(a))return isNaN(s)?1:-1;if(typeof s!=typeof a&&(s+="",a+=""),s<a)return-1;if(a<s)return 1}}return 0},sortNaturalAsc:function(e,t,r,o){return e===t?0:(r=L.string[o.empties[r]||o.emptyTo],""===e&&0!==r?"boolean"==typeof r?r?-1:1:-r||-1:""===t&&0!==r?"boolean"==typeof r?r?1:-1:r||1:L.sortNatural(e,t))},sortNaturalDesc:function(e,t,r,o){return e===t?0:(r=L.string[o.empties[r]||o.emptyTo],""===e&&0!==r?"boolean"==typeof r?r?-1:1:r||1:""===t&&0!==r?"boolean"==typeof r?r?1:-1:-r||-1:L.sortNatural(t,e))},sortText:function(e,t){return t<e?1:e<t?-1:0},getTextValue:function(e,t,r){if(r){for(var o=e?e.length:0,s=r+t,a=0;a<o;a++)s+=e.charCodeAt(a);return t*s}return 0},sortNumericAsc:function(e,t,r,o,s,a){return e===t?0:(s=L.string[a.empties[s]||a.emptyTo],""===e&&0!==s?"boolean"==typeof s?s?-1:1:-s||-1:""===t&&0!==s?"boolean"==typeof s?s?1:-1:s||1:(e=isNaN(e)?L.getTextValue(e,r,o):e)-(t=isNaN(t)?L.getTextValue(t,r,o):t))},sortNumericDesc:function(e,t,r,o,s,a){return e===t?0:(s=L.string[a.empties[s]||a.emptyTo],""===e&&0!==s?"boolean"==typeof s?s?-1:1:s||1:""===t&&0!==s?"boolean"==typeof s?s?1:-1:-s||-1:(isNaN(e)&&(e=L.getTextValue(e,r,o)),(t=isNaN(t)?L.getTextValue(t,r,o):t)-e))},sortNumeric:function(e,t){return e-t},addWidget:function(e){e.id&&!L.isEmptyObject(L.getWidgetById(e.id))&&console.warn('"'+e.id+'" widget was loaded more than once!'),L.widgets[L.widgets.length]=e},hasWidget:function(e,t){return(e=A(e)).length&&e[0].config&&e[0].config.widgetInit[t]||!1},getWidgetById:function(e){for(var t,r=L.widgets.length,o=0;o<r;o++)if((t=L.widgets[o])&&t.id&&t.id.toLowerCase()===e.toLowerCase())return t},applyWidgetOptions:function(e){var t,r,o,s=e.config,a=s.widgets.length;if(a)for(t=0;t<a;t++)(r=L.getWidgetById(s.widgets[t]))&&r.options&&(o=A.extend(!0,{},r.options),s.widgetOptions=A.extend(!0,o,s.widgetOptions),A.extend(!0,L.defaults.widgetOptions,r.options))},addWidgetFromClass:function(e){var t,r,o=e.config,s="^"+o.widgetClass.replace(L.regex.templateName,"(\\S+)+")+"$",a=new RegExp(s,"g"),n=(e.className||"").split(L.regex.spaces);if(n.length)for(t=n.length,r=0;r<t;r++)n[r].match(a)&&(o.widgets[o.widgets.length]=n[r].replace(a,"$1"))},applyWidgetId:function(e,t,r){var o,s,a,n=(e=A(e)[0]).config,i=n.widgetOptions,l=L.debug(n,"core"),t=L.getWidgetById(t);t&&(a=t.id,o=!1,A.inArray(a,n.widgets)<0&&(n.widgets[n.widgets.length]=a),l&&(s=new Date),!r&&n.widgetInit[a]||(n.widgetInit[a]=!0,e.hasInitialized&&L.applyWidgetOptions(e),"function"==typeof t.init&&(o=!0,l&&console[console.group?"group":"log"]("Initializing "+a+" widget"),t.init(e,t,n,i))),r||"function"!=typeof t.format||(o=!0,l&&console[console.group?"group":"log"]("Updating "+a+" widget"),t.format(e,n,i,!1)),l)&&o&&(console.log("Completed "+(r?"initializing ":"applying ")+a+" widget"+L.benchmark(s)),console.groupEnd)&&console.groupEnd()},applyWidget:function(e,t,r){var o,s,a,n,i,l=(e=A(e)[0]).config,d=L.debug(l,"core"),c=[];if(!1===t||!e.hasInitialized||!e.isApplyingWidgets&&!e.isUpdating){if(d&&(i=new Date),L.addWidgetFromClass(e),clearTimeout(l.timerReady),l.widgets.length){for(e.isApplyingWidgets=!0,l.widgets=A.grep(l.widgets,function(e,t){return A.inArray(e,l.widgets)===t}),s=(a=l.widgets||[]).length,o=0;o<s;o++)(n=L.getWidgetById(a[o]))&&n.id?(n.priority||(n.priority=10),c[o]=n):d&&console.warn('"'+a[o]+'" was enabled, but the widget code has not been loaded!');for(c.sort(function(e,t){return e.priority<t.priority?-1:e.priority===t.priority?0:1}),s=c.length,d&&console[console.group?"group":"log"]("Start "+(t?"initializing":"applying")+" widgets"),o=0;o<s;o++)(n=c[o])&&n.id&&L.applyWidgetId(e,n.id,t);d&&console.groupEnd&&console.groupEnd()}l.timerReady=setTimeout(function(){e.isApplyingWidgets=!1,A.data(e,"lastWidgetApplication",new Date),l.$table.triggerHandler("tablesorter-ready"),t||"function"!=typeof r||r(e),d&&(n=l.widgets.length,console.log("Completed "+(!0===t?"initializing ":"applying ")+n+" widget"+(1!==n?"s":"")+L.benchmark(i)))},10)}},removeWidget:function(e,t,r){var o,s,a,n,i=(e=A(e)[0]).config;if(!0===t)for(t=[],n=L.widgets.length,a=0;a<n;a++)(s=L.widgets[a])&&s.id&&(t[t.length]=s.id);else t=(A.isArray(t)?t.join(","):t||"").toLowerCase().split(/[\s,]+/);for(n=t.length,o=0;o<n;o++)s=L.getWidgetById(t[o]),0<=(a=A.inArray(t[o],i.widgets))&&!0!==r&&i.widgets.splice(a,1),s&&s.remove&&(L.debug(i,"core")&&console.log((r?"Refreshing":"Removing")+' "'+t[o]+'" widget'),s.remove(e,i,i.widgetOptions,r),i.widgetInit[t[o]]=!1);i.$table.triggerHandler("widgetRemoveEnd",e)},refreshWidgets:function(e,t,r){for(var o,s=(e=A(e)[0]).config.widgets,a=L.widgets,n=a.length,i=[],l=function(e){A(e).triggerHandler("refreshComplete")},d=0;d<n;d++)(o=a[d])&&o.id&&(t||A.inArray(o.id,s)<0)&&(i[i.length]=o.id);L.removeWidget(e,i.join(","),!0),!0!==r?(L.applyWidget(e,t||!1,l),t&&L.applyWidget(e,!1,l)):l(e)},benchmark:function(e){return" ("+((new Date).getTime()-e.getTime())+" ms)"},log:function(){console.log(arguments)},debug:function(e,t){return e&&(!0===e.debug||"string"==typeof e.debug&&-1<e.debug.indexOf(t))},isEmptyObject:function(e){for(var t in e)return!1;return!0},isValueInArray:function(e,t){for(var r=t&&t.length||0,o=0;o<r;o++)if(t[o][0]===e)return o;return-1},formatFloat:function(e,t){return"string"!=typeof e||""===e?e:(e=(t&&t.config?!1!==t.config.usNumberFormat:void 0===t||t)?e.replace(L.regex.comma,""):e.replace(L.regex.digitNonUS,"").replace(L.regex.comma,"."),L.regex.digitNegativeTest.test(e)&&(e=e.replace(L.regex.digitNegativeReplace,"-$1")),t=parseFloat(e),isNaN(t)?A.trim(e):t)},isDigit:function(e){return isNaN(e)?L.regex.digitTest.test(e.toString().replace(L.regex.digitReplace,"")):""!==e},computeColumnIndex:function(e,t){for(var r,o,s,a,n,i,l,d,c,g=t&&t.columns||0,p=[],u=new Array(g),f=0;f<e.length;f++)for(n=e[f].cells,r=0;r<n.length;r++){for(i=f,l=(a=n[r]).rowSpan||1,d=a.colSpan||1,void 0===p[i]&&(p[i]=[]),o=0;o<p[i].length+1;o++)if(void 0===p[i][o]){c=o;break}for(g&&a.cellIndex===c||(a.setAttribute?a.setAttribute("data-column",c):A(a).attr("data-column",c)),o=i;o<i+l;o++)for(void 0===p[o]&&(p[o]=[]),u=p[o],s=c;s<c+d;s++)u[s]="x"}return L.checkColumnCount(e,p,u.length),u.length},checkColumnCount:function(e,t,r){for(var o,s=!0,a=[],n=0;n<t.length;n++)if(t[n]&&(o=t[n].length,t[n].length!==r)){s=!1;break}s||(e.each(function(e,t){t=t.parentElement.nodeName;a.indexOf(t)<0&&a.push(t)}),console.error("Invalid or incorrect number of columns in the "+a.join(" or ")+"; expected "+r+", but found "+o+" columns"))},fixColumnWidth:function(e){var t,r,o,s,a,e=(e=A(e)[0]).config,n=e.$table.children("colgroup");if(n.length&&n.hasClass(L.css.colgroup)&&n.remove(),e.widthFixed&&0===e.$table.children("colgroup").length){for(n=A('<colgroup class="'+L.css.colgroup+'">'),t=e.$table.width(),s=(o=e.$tbodies.find("tr:first").children(":visible")).length,a=0;a<s;a++)r=parseInt(o.eq(a).width()/t*1e3,10)/10+"%",n.append(A("<col>").css("width",r));e.$table.prepend(n)}},getData:function(e,t,r){var o,s,a="",e=A(e);return e.length?(o=!!A.metadata&&e.metadata(),s=" "+(e.attr("class")||""),void 0!==e.data(r)||void 0!==e.data(r.toLowerCase())?a+=e.data(r)||e.data(r.toLowerCase()):o&&void 0!==o[r]?a+=o[r]:t&&void 0!==t[r]?a+=t[r]:" "!==s&&s.match(" "+r+"-")&&(a=s.match(new RegExp("\\s"+r+"-([\\w-]+)"))[1]||""),A.trim(a)):""},getColumnData:function(e,t,r,o,s){if("object"!=typeof t||null===t)return t;var a,e=(e=A(e)[0]).config,s=s||e.$headers,n=e.$headerIndexed&&e.$headerIndexed[r]||s.find('[data-column="'+r+'"]:last');if(void 0!==t[r])return o?t[r]:t[s.index(n)];for(a in t)if("string"==typeof a&&n.filter(a).add(n.find(a)).length)return t[a]},isProcessing:function(e,t,r){var o=(e=A(e))[0].config,s=r||e.find("."+L.css.header);t?(void 0!==r&&0<o.sortList.length&&(s=s.filter(function(){return!this.sortDisabled&&0<=L.isValueInArray(parseFloat(A(this).attr("data-column")),o.sortList)})),e.add(s).addClass(L.css.processing+" "+o.cssProcessing)):e.add(s).removeClass(L.css.processing+" "+o.cssProcessing)},processTbody:function(e,t,r){if(e=A(e)[0],r)return e.isProcessing=!0,t.before('<colgroup class="tablesorter-savemyplace"/>'),A.fn.detach?t.detach():t.remove();r=A(e).find("colgroup.tablesorter-savemyplace");t.insertAfter(r),r.remove(),e.isProcessing=!1},clearTableBody:function(e){A(e)[0].config.$tbodies.children().detach()},characterEquivalents:{a:"áàâãäąå",A:"ÁÀÂÃÄĄÅ",c:"çćč",C:"ÇĆČ",e:"éèêëěę",E:"ÉÈÊËĚĘ",i:"íìİîïı",I:"ÍÌİÎÏ",o:"óòôõöō",O:"ÓÒÔÕÖŌ",ss:"ß",SS:"ẞ",u:"úùûüů",U:"ÚÙÛÜŮ"},replaceAccents:function(e){var t,r="[",o=L.characterEquivalents;if(!L.characterRegex){for(t in L.characterRegexArray={},o)"string"==typeof t&&(r+=o[t],L.characterRegexArray[t]=new RegExp("["+o[t]+"]","g"));L.characterRegex=new RegExp(r+"]")}if(L.characterRegex.test(e))for(t in o)"string"==typeof t&&(e=e.replace(L.characterRegexArray[t],t));return e},validateOptions:function(e){var t,r,o,s,a="headers sortForce sortList sortAppend widgets".split(" "),n=e.originalSettings;if(n){for(t in L.debug(e,"core")&&(s=new Date),n)if("undefined"===(o=typeof L.defaults[t]))console.warn('Tablesorter Warning! "table.config.'+t+'" option not recognized');else if("object"===o)for(r in n[t])o=L.defaults[t]&&typeof L.defaults[t][r],A.inArray(t,a)<0&&"undefined"===o&&console.warn('Tablesorter Warning! "table.config.'+t+"."+r+'" option not recognized');L.debug(e,"core")&&console.log("validate options time:"+L.benchmark(s))}},restoreHeaders:function(e){for(var t,r=A(e)[0].config,o=r.$table.find(r.selectorHeaders),s=o.length,a=0;a<s;a++)(t=o.eq(a)).find("."+L.css.headerIn).length&&t.html(r.headerContent[a])},destroy:function(e,t,r){var o,s,a,n,i;(e=A(e)[0]).hasInitialized&&(L.removeWidget(e,!0,!1),o=A(e),s=e.config,n=(a=o.find("thead:first")).find("tr."+L.css.headerRow).removeClass(L.css.headerRow+" "+s.cssHeaderRow),i=o.find("tfoot:first > tr").children("th, td"),!1===t&&0<=A.inArray("uitheme",s.widgets)&&(o.triggerHandler("applyWidgetId",["uitheme"]),o.triggerHandler("applyWidgetId",["zebra"])),a.find("tr").not(n).remove(),a="sortReset update updateRows updateAll updateHeaders updateCell addRows updateComplete sorton appendCache updateCache applyWidgetId applyWidgets refreshWidgets removeWidget destroy mouseup mouseleave "+"keypress sortBegin sortEnd resetToLoadState ".split(" ").join(s.namespace+" "),o.removeData("tablesorter").unbind(a.replace(L.regex.spaces," ")),s.$headers.add(i).removeClass([L.css.header,s.cssHeader,s.cssAsc,s.cssDesc,L.css.sortAsc,L.css.sortDesc,L.css.sortNone].join(" ")).removeAttr("data-column").removeAttr("aria-label").attr("aria-disabled","true"),n.find(s.selectorSort).unbind("mousedown mouseup keypress ".split(" ").join(s.namespace+" ").replace(L.regex.spaces," ")),L.restoreHeaders(e),o.toggleClass(L.css.table+" "+s.tableClass+" tablesorter-"+s.theme,!1===t),o.removeClass(s.namespace.slice(1)),e.hasInitialized=!1,delete e.config.cache,"function"==typeof r&&r(e),L.debug(s,"core"))&&console.log("tablesorter has been removed")}};A.fn.tablesorter=function(t){return this.each(function(){var e=A.extend(!0,{},L.defaults,t,L.instanceMethods);e.originalSettings=t,!this.hasInitialized&&L.buildTable&&"TABLE"!==this.nodeName?L.buildTable(this,e):L.setup(this,e)})},window.console&&window.console.log||(L.logs=[],(console={}).log=console.warn=console.error=console.table=function(){var e=1<arguments.length?arguments:arguments[0];L.logs[L.logs.length]={date:Date.now(),log:e}}),L.addParser({id:"no-parser",is:function(){return!1},format:function(){return""},type:"text"}),L.addParser({id:"text",is:function(){return!0},format:function(e,t){t=t.config;return e&&(e=A.trim(t.ignoreCase?e.toLocaleLowerCase():e),e=t.sortLocaleCompare?L.replaceAccents(e):e),e},type:"text"}),L.regex.nondigit=/[^\w,. \-()]/g,L.addParser({id:"digit",is:function(e){return L.isDigit(e)},format:function(e,t){var r=L.formatFloat((e||"").replace(L.regex.nondigit,""),t);return e&&"number"==typeof r?r:e&&A.trim(e&&t.config.ignoreCase?e.toLocaleLowerCase():e)},type:"numeric"}),L.regex.currencyReplace=/[+\-,. ]/g,L.regex.currencyTest=/^\(?\d+[\u00a3$\u20ac\u00a4\u00a5\u00a2?.]|[\u00a3$\u20ac\u00a4\u00a5\u00a2?.]\d+\)?$/,L.addParser({id:"currency",is:function(e){return e=(e||"").replace(L.regex.currencyReplace,""),L.regex.currencyTest.test(e)},format:function(e,t){var r=L.formatFloat((e||"").replace(L.regex.nondigit,""),t);return e&&"number"==typeof r?r:e&&A.trim(e&&t.config.ignoreCase?e.toLocaleLowerCase():e)},type:"numeric"}),L.regex.urlProtocolTest=/^(https?|ftp|file):\/\//,L.regex.urlProtocolReplace=/(https?|ftp|file):\/\/(www\.)?/,L.addParser({id:"url",is:function(e){return L.regex.urlProtocolTest.test(e)},format:function(e){return e&&A.trim(e.replace(L.regex.urlProtocolReplace,""))},type:"text"}),L.regex.dash=/-/g,L.regex.isoDate=/^\d{4}[\/\-]\d{1,2}[\/\-]\d{1,2}/,L.addParser({id:"isoDate",is:function(e){return L.regex.isoDate.test(e)},format:function(e){var t=e&&new Date(e.replace(L.regex.dash,"/"));return t instanceof Date&&isFinite(t)?t.getTime():e},type:"numeric"}),L.regex.percent=/%/g,L.regex.percentTest=/(\d\s*?%|%\s*?\d)/,L.addParser({id:"percent",is:function(e){return L.regex.percentTest.test(e)&&e.length<15},format:function(e,t){return e&&L.formatFloat(e.replace(L.regex.percent,""),t)},type:"numeric"}),L.addParser({id:"image",is:function(e,t,r,o){return 0<o.find("img").length},format:function(e,t,r){return A(r).find("img").attr(t.config.imgAttr||"alt")||e},parsed:!0,type:"text"}),L.regex.dateReplace=/(\S)([AP]M)$/i,L.regex.usLongDateTest1=/^[A-Z]{3,10}\.?\s+\d{1,2},?\s+(\d{4})(\s+\d{1,2}:\d{2}(:\d{2})?(\s+[AP]M)?)?$/i,L.regex.usLongDateTest2=/^\d{1,2}\s+[A-Z]{3,10}\s+\d{4}/i,L.addParser({id:"usLongDate",is:function(e){return L.regex.usLongDateTest1.test(e)||L.regex.usLongDateTest2.test(e)},format:function(e){var t=e&&new Date(e.replace(L.regex.dateReplace,"$1 $2"));return t instanceof Date&&isFinite(t)?t.getTime():e},type:"numeric"}),L.regex.shortDateTest=/(^\d{1,2}[\/\s]\d{1,2}[\/\s]\d{4})|(^\d{4}[\/\s]\d{1,2}[\/\s]\d{1,2})/,L.regex.shortDateReplace=/[\-.,]/g,L.regex.shortDateXXY=/(\d{1,2})[\/\s](\d{1,2})[\/\s](\d{4})/,L.regex.shortDateYMD=/(\d{4})[\/\s](\d{1,2})[\/\s](\d{1,2})/,L.convertFormat=function(e,t){e=(e||"").replace(L.regex.spaces," ").replace(L.regex.shortDateReplace,"/"),"mmddyyyy"===t?e=e.replace(L.regex.shortDateXXY,"$3/$1/$2"):"ddmmyyyy"===t?e=e.replace(L.regex.shortDateXXY,"$3/$2/$1"):"yyyymmdd"===t&&(e=e.replace(L.regex.shortDateYMD,"$1/$2/$3"));t=new Date(e);return t instanceof Date&&isFinite(t)?t.getTime():""},L.addParser({id:"shortDate",is:function(e){return e=(e||"").replace(L.regex.spaces," ").replace(L.regex.shortDateReplace,"/"),L.regex.shortDateTest.test(e)},format:function(e,t,r,o){var s,a;return e&&(t=(a=(s=t.config).$headerIndexed[o]).length&&a.data("dateFormat")||L.getData(a,L.getColumnData(t,s.headers,o),"dateFormat")||s.dateFormat,a.length&&a.data("dateFormat",t),L.convertFormat(e,t)||e)},type:"numeric"}),L.regex.timeTest=/^(0?[1-9]|1[0-2]):([0-5]\d)(\s[AP]M)$|^((?:[01]\d|[2][0-4]):[0-5]\d)$/i,L.regex.timeMatch=/(0?[1-9]|1[0-2]):([0-5]\d)(\s[AP]M)|((?:[01]\d|[2][0-4]):[0-5]\d)/i,L.addParser({id:"time",is:function(e){return L.regex.timeTest.test(e)},format:function(e){var t=(e||"").match(L.regex.timeMatch),r=new Date(e),t=e&&(null!==t?t[0]:"00:00 AM"),t=t&&new Date("2000/01/01 "+t.replace(L.regex.dateReplace,"$1 $2"));return t instanceof Date&&isFinite(t)?(r instanceof Date&&isFinite(r)?r.getTime():0)?parseFloat(t.getTime()+"."+r.getTime()):t.getTime():e},type:"numeric"}),L.addParser({id:"metadata",is:function(){return!1},format:function(e,t,r){t=t.config,t=t.parserMetadataName||"sortValue";return A(r).metadata()[t]},type:"numeric"}),L.addWidget({id:"zebra",priority:90,format:function(e,t,r){for(var o,s,a,n,i,l=new RegExp(t.cssChildRow,"i"),d=t.$tbodies.add(A(t.namespace+"_extra_table").children("tbody:not(."+t.cssInfoBlock+")")),c=0;c<d.length;c++)for(s=0,i=(o=d.eq(c).children("tr:visible").not(t.selectorRemove)).length,n=0;n<i;n++)a=o.eq(n),l.test(a[0].className)||s++,a.removeClass(r.zebra[(a=s%2==0)?1:0]).addClass(r.zebra[a?0:1])},remove:function(e,t,r,o){if(!o)for(var s,a=t.$tbodies,n=(r.zebra||["even","odd"]).join(" "),i=0;i<a.length;i++)(s=L.processTbody(e,a.eq(i),!0)).children().removeClass(n),L.processTbody(e,s,!1)}})}(e),e.tablesorter});
```
