## Supplementary Data SD1 for "Inferring binding specificities of human transcription factors with the wisdom of crowds": jquery.tablesorter.widgets.min.js

```
(function(factory){if (typeof define === 'function' && define.amd){define(['jquery'], factory);} else if (typeof module === 'object' && typeof module.exports === 'object'){module.exports = factory(require('jquery'));} else {factory(jQuery);}}(function(jQuery){
/*! tablesorter (FORK) - updated 2024-08-13 (v2.32.0)*/
!function(e){"function"==typeof define&&define.amd?define(["jquery"],e):"object"==typeof module&&"object"==typeof module.exports?module.exports=e(require("jquery")):e(jQuery)}(function(e){return function(h,u,p){"use strict";var g=h.tablesorter||{};h.extend(!0,g.defaults,{fixedUrl:"",widgetOptions:{storage_fixedUrl:"",storage_group:"",storage_page:"",storage_storageType:"",storage_tableId:"",storage_useSessionStorage:""}}),g.storage=function(e,t,r,i){var a=!1,l={},s=(e=h(e)[0]).config,n=s&&s.widgetOptions,o=g.debug(s,"storage"),c=(i&&i.storageType||n&&n.storage_storageType).toString().charAt(0).toLowerCase(),d=c?"":i&&i.useSessionStorage||n&&n.storage_useSessionStorage,f=h(e),e=i&&i.id||f.attr(i&&i.group||n&&n.storage_group||"data-table-group")||n&&n.storage_tableId||e.id||h(".tablesorter").index(f),f=i&&i.url||f.attr(i&&i.page||n&&n.storage_page||"data-table-page")||n&&n.storage_fixedUrl||s&&s.fixedUrl||u.location.pathname;if("c"!==c&&(c="s"===c||d?"sessionStorage":"localStorage")in u)try{u[c].setItem("_tmptest","temp"),a=!0,u[c].removeItem("_tmptest")}catch(e){console.warn(c+" is not supported in this browser")}if(o&&console.log("Storage >> Using",a?c:"cookies"),h.parseJSON&&(l=a?h.parseJSON(u[c][t]||"null")||{}:(i=p.cookie.split(/[;\s|=]/),0!==(n=h.inArray(t,i)+1)&&h.parseJSON(i[n]||"null")||{})),void 0===r||!u.JSON||!JSON.hasOwnProperty("stringify"))return l&&l[f]?l[f][e]:"";l[f]||(l[f]={}),l[f][e]=r,a?u[c][t]=JSON.stringify(l):((s=new Date).setTime(s.getTime()+31536e6),p.cookie=t+"="+JSON.stringify(l).replace(/\"/g,'"')+"; expires="+s.toGMTString()+"; path=/")}}(e,window,document),function(S){"use strict";var C=S.tablesorter||{};C.themes={bootstrap:{table:"table table-bordered table-striped",caption:"caption",header:"bootstrap-header",sortNone:"",sortAsc:"",sortDesc:"",active:"",hover:"",icons:"",iconSortNone:"bootstrap-icon-unsorted",iconSortAsc:"glyphicon glyphicon-chevron-up",iconSortDesc:"glyphicon glyphicon-chevron-down",filterRow:"",footerRow:"",footerCells:"",even:"",odd:""},jui:{table:"ui-widget ui-widget-content ui-corner-all",caption:"ui-widget-content",header:"ui-widget-header ui-corner-all ui-state-default",sortNone:"",sortAsc:"",sortDesc:"",active:"ui-state-active",hover:"ui-state-hover",icons:"ui-icon",iconSortNone:"ui-icon-carat-2-n-s ui-icon-caret-2-n-s",iconSortAsc:"ui-icon-carat-1-n ui-icon-caret-1-n",iconSortDesc:"ui-icon-carat-1-s ui-icon-caret-1-s",filterRow:"",footerRow:"",footerCells:"",even:"ui-widget-content",odd:"ui-state-default"}},S.extend(C.css,{wrapper:"tablesorter-wrapper"}),C.addWidget({id:"uitheme",priority:10,format:function(e,t,r){var i,a,l,s,n,o,c,d,f,h,u,p,g=C.themes,m=t.$table.add(S(t.namespace+"_extra_table")),b=t.$headers.add(S(t.namespace+"_extra_headers")),y=t.theme||"jui",_=g[y]||{},v=S.trim([_.sortNone,_.sortDesc,_.sortAsc,_.active].join(" ")),w=S.trim([_.iconSortNone,_.iconSortDesc,_.iconSortAsc].join(" ")),x=C.debug(t,"uitheme");for(x&&(n=new Date),m.hasClass("tablesorter-"+y)&&t.theme===t.appliedTheme&&r.uitheme_applied||(r.uitheme_applied=!0,h=g[t.appliedTheme]||{},g=(p=!S.isEmptyObject(h))?[h.sortNone,h.sortDesc,h.sortAsc,h.active].join(" "):"",u=p?[h.iconSortNone,h.iconSortDesc,h.iconSortAsc].join(" "):"",p&&(r.zebra[0]=S.trim(" "+r.zebra[0].replace(" "+h.even,"")),r.zebra[1]=S.trim(" "+r.zebra[1].replace(" "+h.odd,"")),t.$tbodies.children().removeClass([h.even,h.odd].join(" "))),_.even&&(r.zebra[0]+=" "+_.even),_.odd&&(r.zebra[1]+=" "+_.odd),m.children("caption").removeClass(h.caption||"").addClass(_.caption),d=m.removeClass((t.appliedTheme?"tablesorter-"+(t.appliedTheme||""):"")+" "+(h.table||"")).addClass("tablesorter-"+y+" "+(_.table||"")).children("tfoot"),t.appliedTheme=t.theme,d.length&&d.children("tr").removeClass(h.footerRow||"").addClass(_.footerRow).children("th, td").removeClass(h.footerCells||"").addClass(_.footerCells),b.removeClass((p?[h.header,h.hover,g].join(" "):"")||"").addClass(_.header).not(".sorter-false").unbind("mouseenter.tsuitheme mouseleave.tsuitheme").bind("mouseenter.tsuitheme mouseleave.tsuitheme",function(e){S(this)["mouseenter"===e.type?"addClass":"removeClass"](_.hover||"")}),b.each(function(){var e=S(this);e.find("."+C.css.wrapper).length||e.wrapInner('<div class="'+C.css.wrapper+'" style="position:relative;height:100%;width:100%"></div>')}),t.cssIcon&&b.find("."+C.css.icon).removeClass(p?[h.icons,u].join(" "):"").addClass(_.icons||""),C.hasWidget(t.table,"filter")&&(a=function(){m.children("thead").children("."+C.css.filterRow).removeClass(p&&h.filterRow||"").addClass(_.filterRow||"")},r.filter_initialized?a():m.one("filterInit",function(){a()}))),i=0;i<t.columns;i++)o=t.$headers.add(S(t.namespace+"_extra_headers")).not(".sorter-false").filter('[data-column="'+i+'"]'),c=C.css.icon?o.find("."+C.css.icon):S(),(f=b.not(".sorter-false").filter('[data-column="'+i+'"]:last')).length&&(o.removeClass(v),c.removeClass(w),f[0].sortDisabled?c.removeClass(_.icons||""):(l=_.sortNone,s=_.iconSortNone,f.hasClass(C.css.sortAsc)?(l=[_.sortAsc,_.active].join(" "),s=_.iconSortAsc):f.hasClass(C.css.sortDesc)&&(l=[_.sortDesc,_.active].join(" "),s=_.iconSortDesc),o.addClass(l),c.addClass(s||"")));x&&console.log("uitheme >> Applied "+y+" theme"+C.benchmark(n))},remove:function(e,t,r,i){var a,l,s,n,o;r.uitheme_applied&&(a=t.$table,t=t.appliedTheme||"jui",l=C.themes[t]||C.themes.jui,s=a.children("thead").children(),n=l.sortNone+" "+l.sortDesc+" "+l.sortAsc,o=l.iconSortNone+" "+l.iconSortDesc+" "+l.iconSortAsc,a.removeClass("tablesorter-"+t+" "+l.table),r.uitheme_applied=!1,i||(a.find(C.css.header).removeClass(l.header),s.unbind("mouseenter.tsuitheme mouseleave.tsuitheme").removeClass(l.hover+" "+n+" "+l.active).filter("."+C.css.filterRow).removeClass(l.filterRow),s.find("."+C.css.icon).removeClass(l.icons+" "+o)))}})}(e),function(m){"use strict";var b=m.tablesorter||{};b.addWidget({id:"columns",priority:65,options:{columns:["primary","secondary","tertiary"]},format:function(e,t,r){for(var i,a,l,s,n,o=t.$table,c=t.$tbodies,d=t.sortList,f=d.length,h=r&&r.columns||["primary","secondary","tertiary"],u=h.length-1,p=h.join(" "),g=0;g<c.length;g++)(a=(i=b.processTbody(e,c.eq(g),!0)).children("tr")).each(function(){if(l=m(this),"none"!==this.style.display&&(s=l.children().removeClass(p),d)&&d[0]&&(s.eq(d[0][0]).addClass(h[0]),1<f))for(n=1;n<f;n++)s.eq(d[n][0]).addClass(h[n]||h[u])}),b.processTbody(e,i,!1);if(t=!1!==r.columns_thead?["thead tr"]:[],!1!==r.columns_tfoot&&t.push("tfoot tr"),t.length&&(a=o.find(t.join(",")).children().removeClass(p),f))for(n=0;n<f;n++)a.filter('[data-column="'+d[n][0]+'"]').addClass(h[n]||h[u])},remove:function(e,t,r){var i,a,l=t.$tbodies,s=(r.columns||["primary","secondary","tertiary"]).join(" ");for(t.$headers.removeClass(s),t.$table.children("tfoot").children("tr").children("th, td").removeClass(s),i=0;i<l.length;i++)(a=b.processTbody(e,l.eq(i),!0)).children("tr").each(function(){m(this).children().removeClass(s)}),b.processTbody(e,a,!1)}})}(e),function(A){"use strict";var I,O,E=A.tablesorter||{},g=E.css,o=E.keyCodes;A.extend(g,{filterRow:"tablesorter-filter-row",filter:"tablesorter-filter",filterDisabled:"disabled",filterRowHide:"hideme"}),A.extend(o,{backSpace:8,escape:27,space:32,left:37,down:40}),E.addWidget({id:"filter",priority:50,options:{filter_cellFilter:"",filter_childRows:!1,filter_childByColumn:!1,filter_childWithSibs:!0,filter_columnAnyMatch:!0,filter_columnFilters:!0,filter_cssFilter:"",filter_defaultAttrib:"data-value",filter_defaultFilter:{},filter_excludeFilter:{},filter_external:"",filter_filteredRow:"filtered",filter_filterLabel:'Filter "{{label}}" column by...',filter_formatter:null,filter_functions:null,filter_hideEmpty:!0,filter_hideFilters:!1,filter_ignoreCase:!0,filter_liveSearch:!0,filter_matchType:{input:"exact",select:"exact"},filter_onlyAvail:"filter-onlyAvail",filter_placeholder:{search:"",select:""},filter_reset:null,filter_resetOnEsc:!0,filter_saveFilters:!1,filter_searchDelay:300,filter_searchFiltered:!0,filter_selectSource:null,filter_selectSourceSeparator:"|",filter_serversideFiltering:!1,filter_startsWith:!1,filter_useParsedData:!1},format:function(e,t,r){t.$table.hasClass("hasFilters")||I.init(e,t,r)},remove:function(e,t,r,i){var a,l,s=t.$table,n=t.$tbodies,o="addRows updateCell update updateRows updateComplete appendCache filterReset filterAndSortReset filterFomatterUpdate filterEnd search stickyHeadersInit ".split(" ").join(t.namespace+"filter ");if(s.removeClass("hasFilters").unbind(o.replace(E.regex.spaces," ")).find("."+g.filterRow).remove(),r.filter_initialized=!1,!i){for(a=0;a<n.length;a++)(l=E.processTbody(e,n.eq(a),!0)).children().removeClass(r.filter_filteredRow).show(),E.processTbody(e,l,!1);r.filter_reset&&A(document).undelegate(r.filter_reset,"click"+t.namespace+"filter")}}}),I=E.filter={regex:{regex:/^\/((?:\\\/|[^\/])+)\/([migyu]{0,5})?$/,child:/tablesorter-childRow/,filtered:/filtered/,type:/undefined|number/,exact:/(^[\"\'=]+)|([\"\'=]+$)/g,operators:/[<>=]/g,query:"(q|query)",wild01:/\?/g,wild0More:/\*/g,quote:/\"/g,isNeg1:/(>=?\s*-\d)/,isNeg2:/(<=?\s*\d)/},types:{or:function(e,t,r){if(!O.orTest.test(t.iFilter)&&!O.orSplit.test(t.filter)||O.regex.test(t.filter))return null;for(var i,a,l=A.extend({},t),s=t.filter.split(O.orSplit),n=t.iFilter.split(O.orSplit),o=s.length,c=0;c<o;c++){l.nestedFilters=!0,l.filter=""+(I.parseFilter(e,s[c],t)||""),l.iFilter=""+(I.parseFilter(e,n[c],t)||""),a="("+(I.parseFilter(e,l.filter,t)||"")+")";try{if(i=new RegExp(t.isMatch?a:"^"+a+"$",e.widgetOptions.filter_ignoreCase?"i":"").test(l.exact)||I.processTypes(e,l,r))return i}catch(e){return null}}return i||!1},and:function(e,t,r){if(O.andTest.test(t.filter)){for(var i,a,l,s=A.extend({},t),n=t.filter.split(O.andSplit),o=t.iFilter.split(O.andSplit),c=n.length,d=0;d<c;d++){s.nestedFilters=!0,s.filter=""+(I.parseFilter(e,n[d],t)||""),s.iFilter=""+(I.parseFilter(e,o[d],t)||""),l=("("+(I.parseFilter(e,s.filter,t)||"")+")").replace(O.wild01,"\\S{1}").replace(O.wild0More,"\\S*");try{a=new RegExp(t.isMatch?l:"^"+l+"$",e.widgetOptions.filter_ignoreCase?"i":"").test(s.exact)||I.processTypes(e,s,r),i=(0===d||i)&&a}catch(e){return null}}return i||!1}return null},regex:function(e,t){if(O.regex.test(t.filter)){var r,i=t.filter_regexCache[t.index]||O.regex.exec(t.filter),a=i instanceof RegExp;try{a||(t.filter_regexCache[t.index]=i=new RegExp(i[1],i[2])),r=i.test(t.exact)}catch(e){r=!1}return r}return null},operators:function(e,t){var r,i,a,l,s,n,o;return O.operTest.test(t.iFilter)&&""!==t.iExact?(a=e.table,l=t.parsed[t.index],s=E.formatFloat(t.iFilter.replace(O.operators,""),a),n=e.parsers[t.index]||{},o=s,!l&&"numeric"!==n.type||(i=A.trim(""+t.iFilter.replace(O.operators,"")),s="number"!=typeof(r=I.parseFilter(e,i,t,!0))||""===r||isNaN(r)?s:r),e=!l&&"numeric"!==n.type||isNaN(s)||void 0===t.cache?(i=isNaN(t.iExact)?t.iExact.replace(E.regex.nondigit,""):t.iExact,E.formatFloat(i,a)):t.cache,O.gtTest.test(t.iFilter)?r=O.gteTest.test(t.iFilter)?s<=e:s<e:O.ltTest.test(t.iFilter)&&(r=O.lteTest.test(t.iFilter)?e<=s:e<s),!r&&""===o||r):null},notMatch:function(e,t){var r,i;return O.notTest.test(t.iFilter)?(i=t.iFilter.replace("!",""),i=I.parseFilter(e,i,t)||"",O.exact.test(i)?""===(i=i.replace(O.exact,""))||A.trim(i)!==t.iExact:(r=t.iExact.search(A.trim(i)),""===i||(t.anyMatch?r<0:!(e.widgetOptions.filter_startsWith?0===r:0<=r)))):null},exact:function(e,t){var r;return O.exact.test(t.iFilter)?(r=t.iFilter.replace(O.exact,""),e=I.parseFilter(e,r,t)||"",t.anyMatch?0<=A.inArray(e,t.rowArray):e==t.iExact):null},range:function(e,t){var r,i,a,l,s,n,o,c;return O.toTest.test(t.iFilter)?(a=e.table,l=t.index,s=t.parsed[l],o=(n=t.iFilter.split(O.toSplit))[0].replace(E.regex.nondigit,"")||"",c=E.formatFloat(I.parseFilter(e,o,t),a),o=n[1].replace(E.regex.nondigit,"")||"",i=E.formatFloat(I.parseFilter(e,o,t),a),!s&&"numeric"!==e.parsers[l].type||(c=""===(r=e.parsers[l].format(""+n[0],a,e.$headers.eq(l),l))||isNaN(r)?c:r,i=""===(r=e.parsers[l].format(""+n[1],a,e.$headers.eq(l),l))||isNaN(r)?i:r),r=!s&&"numeric"!==e.parsers[l].type||isNaN(c)||isNaN(i)?(o=isNaN(t.iExact)?t.iExact.replace(E.regex.nondigit,""):t.iExact,E.formatFloat(o,a)):t.cache,i<c&&(o=c,c=i,i=o),c<=r&&r<=i||""===c||""===i):null},wild:function(e,t){if(O.wildOrTest.test(t.iFilter)){var r=""+(I.parseFilter(e,t.iFilter,t)||"");!O.wildTest.test(r)&&t.nestedFilters&&(r=t.isMatch?r:"^("+r+")$");try{return new RegExp(r.replace(O.wild01,"\\S{1}").replace(O.wild0More,"\\S*"),e.widgetOptions.filter_ignoreCase?"i":"").test(t.exact)}catch(e){}}return null},fuzzy:function(e,t){if(O.fuzzyTest.test(t.iFilter)){for(var r=0,i=t.iExact.length,a=t.iFilter.slice(1),l=I.parseFilter(e,a,t)||"",s=0;s<i;s++)t.iExact[s]===l[r]&&(r+=1);return r===l.length}return null}},init:function(r){E.language=A.extend(!0,{},{to:"to",or:"or",and:"and"},E.language);function e(e,t,r){return""===(t=t.trim())?"":(e||"")+t+(r||"")}var t,i,a,l,s,n,o,c,d=r.config,f=d.widgetOptions;if(d.$table.addClass("hasFilters"),d.lastSearch=[],f.filter_searchTimer=null,f.filter_initTimer=null,f.filter_formatterCount=0,f.filter_formatterInit=[],f.filter_anyColumnSelector='[data-column="all"],[data-column="any"]',f.filter_multipleColumnSelector='[data-column*="-"],[data-column*=","]',n="\\{"+O.query+"\\}",A.extend(O,{child:new RegExp(d.cssChildRow),filtered:new RegExp(f.filter_filteredRow),alreadyFiltered:new RegExp("(\\s+(-"+e("|",E.language.or)+e("|",E.language.to)+")\\s+)","i"),toTest:new RegExp("\\s+(-"+e("|",E.language.to)+")\\s+","i"),toSplit:new RegExp("(?:\\s+(?:-"+e("|",E.language.to)+")\\s+)","gi"),andTest:new RegExp("\\s+("+e("",E.language.and,"|")+"&&)\\s+","i"),andSplit:new RegExp("(?:\\s+(?:"+e("",E.language.and,"|")+"&&)\\s+)","gi"),orTest:new RegExp("(\\|"+e("|\\s+",E.language.or,"\\s+")+")","i"),orSplit:new RegExp("(?:\\|"+e("|\\s+(?:",E.language.or,")\\s+")+")","gi"),iQuery:new RegExp(n,"i"),igQuery:new RegExp(n,"ig"),operTest:/^[<>]=?/,gtTest:/>/,gteTest:/>=/,ltTest:/</,lteTest:/<=/,notTest:/^\!/,wildOrTest:/[\?\*\|]/,wildTest:/\?\*/,fuzzyTest:/^~/,exactTest:/[=\"\|!]/}),n=d.$headers.filter(".filter-false, .parser-false").length,!1!==f.filter_columnFilters&&n!==d.$headers.length&&I.buildRow(r,d,f),a="addRows updateCell update updateRows updateComplete appendCache filterReset "+"filterAndSortReset filterResetSaved filterEnd search ".split(" ").join(d.namespace+"filter "),d.$table.bind(a,function(e,t){return n=f.filter_hideEmpty&&A.isEmptyObject(d.cache)&&!(d.delayInit&&"appendCache"===e.type),d.$table.find("."+g.filterRow).toggleClass(f.filter_filteredRow,n),/(search|filter)/.test(e.type)||(e.stopPropagation(),I.buildDefault(r,!0)),"filterReset"===e.type||"filterAndSortReset"===e.type?(d.$table.find("."+g.filter).add(f.filter_$externalFilters).val(""),"filterAndSortReset"===e.type?E.sortReset(this.config,function(){I.searching(r,[])}):I.searching(r,[])):"filterResetSaved"===e.type?E.storage(r,"tablesorter-filters",""):"filterEnd"===e.type?I.buildDefault(r,!0):(t="search"===e.type?t:"updateComplete"===e.type?d.$table.data("lastSearch"):"",/(update|add)/.test(e.type)&&"updateComplete"!==e.type&&(d.lastCombinedFilter=null,d.lastSearch=[],setTimeout(function(){d.$table.triggerHandler("filterFomatterUpdate")},100)),I.searching(r,t,!0)),!1}),f.filter_reset&&(f.filter_reset instanceof A?f.filter_reset.click(function(){d.$table.triggerHandler("filterReset")}):A(f.filter_reset).length&&A(document).undelegate(f.filter_reset,"click"+d.namespace+"filter").delegate(f.filter_reset,"click"+d.namespace+"filter",function(){d.$table.triggerHandler("filterReset")})),f.filter_functions)for(s=0;s<d.columns;s++)if(o=E.getColumnData(r,f.filter_functions,s))if(c=!((l=d.$headerIndexed[s].removeClass("filter-select")).hasClass("filter-false")||l.hasClass("parser-false")),!(t="")===o&&c)I.buildSelect(r,s);else if("object"==typeof o&&c){for(i in o)"string"==typeof i&&(t+=""===t?'<option value="">'+(l.data("placeholder")||l.attr("data-placeholder")||f.filter_placeholder.select||"")+"</option>":"",0<=(a=n=i).indexOf(f.filter_selectSourceSeparator)&&(a=(n=i.split(f.filter_selectSourceSeparator))[1],n=n[0]),t+="<option "+(a===n?"":'data-function-name="'+i+'" ')+'value="'+n+'">'+a+"</option>");d.$table.find("thead").find("select."+g.filter+'[data-column="'+s+'"]').append(t),("function"==typeof(a=f.filter_selectSource)||E.getColumnData(r,a,s))&&I.buildSelect(d.table,s,"",!0,l.hasClass(f.filter_onlyAvail))}I.buildDefault(r,!0),I.bindSearch(r,d.$table.find("."+g.filter),!0),f.filter_external&&I.bindSearch(r,f.filter_external),f.filter_hideFilters&&I.hideFilters(d),d.showProcessing&&(a="filterStart filterEnd ".split(" ").join(d.namespace+"filter-sp "),d.$table.unbind(a.replace(E.regex.spaces," ")).bind(a,function(e,t){l=t?d.$table.find("."+g.header).filter("[data-column]").filter(function(){return""!==t[A(this).data("column")]}):"",E.isProcessing(r,"filterStart"===e.type,t?l:"")})),d.filteredRows=d.totalRows,a="tablesorter-initialized pagerBeforeInitialized ".split(" ").join(d.namespace+"filter "),d.$table.unbind(a.replace(E.regex.spaces," ")).bind(a,function(){I.completeInit(this)}),d.pager&&d.pager.initialized&&!f.filter_initialized?(d.$table.triggerHandler("filterFomatterUpdate"),setTimeout(function(){I.filterInitComplete(d)},100)):f.filter_initialized||I.completeInit(r)},completeInit:function(e){var t=e.config,r=t.widgetOptions,i=I.setDefaults(e,t,r)||[];!i.length||t.delayInit&&""===i.join("")||E.setFilters(e,i,!0),t.$table.triggerHandler("filterFomatterUpdate"),setTimeout(function(){r.filter_initialized||I.filterInitComplete(t)},100)},formatterUpdated:function(e,t){e=e&&e.closest("table"),e=e.length&&e[0].config,e=e&&e.widgetOptions;e&&!e.filter_initialized&&(e.filter_formatterInit[t]=1)},filterInitComplete:function(e){function t(){a.filter_initialized=!0,e.lastSearch=e.$table.data("lastSearch"),e.$table.triggerHandler("filterInit",e),I.findRows(e.table,e.lastSearch||[]),E.debug(e,"filter")&&console.log("Filter >> Widget initialized")}var r,i,a=e.widgetOptions,l=0;if(A.isEmptyObject(a.filter_formatter))t();else{for(i=a.filter_formatterInit.length,r=0;r<i;r++)1===a.filter_formatterInit[r]&&l++;clearTimeout(a.filter_initTimer),a.filter_initialized||l!==a.filter_formatterCount?a.filter_initialized||(a.filter_initTimer=setTimeout(function(){t()},500)):t()}},processFilters:function(e,t){for(var r=[],i=t?encodeURIComponent:decodeURIComponent,a=e.length,l=0;l<a;l++)e[l]&&(r[l]=i(e[l]));return r},setDefaults:function(e,t,r){var i,a,l,s,n=E.getFilters(e)||[];if(r.filter_saveFilters&&E.storage&&(e=E.storage(e,"tablesorter-filters")||[],(i=A.isArray(e))&&""===e.join("")||!i||(n=I.processFilters(e))),""===n.join(""))for(s=t.$headers.add(r.filter_$externalFilters).filter("["+r.filter_defaultAttrib+"]"),a=0;a<=t.columns;a++)l=a===t.columns?"all":a,n[a]=s.filter('[data-column="'+l+'"]').attr(r.filter_defaultAttrib)||n[a]||"";return t.$table.data("lastSearch",n),n},parseFilter:function(e,t,r,i){return i||r.parsed[r.index]?e.parsers[r.index].format(t,e.table,[],r.index):t},buildRow:function(e,t,r){for(var i,a,l,s,n,o,c,d=r.filter_cellFilter,f=t.columns,h=A.isArray(d),u='<tr role="search" class="'+g.filterRow+" "+t.cssIgnoreRow+'">',p=0;p<f;p++)t.$headerIndexed[p].length&&(u=(u+=1<(c=t.$headerIndexed[p]&&t.$headerIndexed[p][0].colSpan||0)?'<td data-column="'+p+"-"+(p+c-1)+'" colspan="'+c+'"':'<td data-column="'+p+'"')+(h?d[p]?' class="'+d[p]+'"':"":""!==d?' class="'+d+'"':"")+"></td>");for(t.$filters=A(u+="</tr>").appendTo(t.$table.children("thead").eq(0)).children("td"),p=0;p<f;p++)(a=t.$headerIndexed[p])&&a.length&&(i=I.getColumnElm(t,t.$filters,p),o=E.getColumnData(e,r.filter_functions,p),l=r.filter_functions&&o&&"function"!=typeof o||a.hasClass("filter-select"),s=E.getColumnData(e,t.headers,p),s="false"===E.getData(a[0],s,"filter")||"false"===E.getData(a[0],s,"parser"),l?u=A("<select>").appendTo(i):((o=E.getColumnData(e,r.filter_formatter,p))?(r.filter_formatterCount++,(u=(u=o(i,p))&&0===u.length?i.children("input"):u)&&(0===u.parent().length||u.parent().length&&u.parent()[0]!==i[0])&&i.append(u)):u=A('<input type="search">').appendTo(i),u&&(c=a.data("placeholder")||a.attr("data-placeholder")||r.filter_placeholder.search||"",u.attr("placeholder",c))),u)&&(n=(A.isArray(r.filter_cssFilter)?void 0!==r.filter_cssFilter[p]&&r.filter_cssFilter[p]||"":r.filter_cssFilter)||"",u.addClass(g.filter+" "+n),c=(n=r.filter_filterLabel).match(/{{([^}]+?)}}/g),A.each(c=c||["{{label}}"],function(e,t){var r=new RegExp(t,"g"),t=a.attr("data-"+t.replace(/{{|}}/g,"")),t=void 0===t?a.text():t;n=n.replace(r,A.trim(t))}),u.attr({"data-column":i.attr("data-column"),"aria-label":n}),s)&&(u.attr("placeholder","").addClass(g.filterDisabled)[0].disabled=!0)},bindSearch:function(a,e,t){var r,l,s,i,n;a=A(a)[0],(e=A(e)).length&&(l=a.config,s=l.widgetOptions,i=l.namespace+"filter",n=s.filter_$externalFilters,!0!==t&&(r=s.filter_anyColumnSelector+","+s.filter_multipleColumnSelector,s.filter_$anyMatch=e.filter(r),n&&n.length?s.filter_$externalFilters=s.filter_$externalFilters.add(e):s.filter_$externalFilters=e,E.setFilters(a,l.$table.data("lastSearch")||[],!1===t)),r="keypress keyup keydown search change input ".split(" ").join(i+" "),e.attr("data-lastSearchTime",(new Date).getTime()).unbind(r.replace(E.regex.spaces," ")).bind("keydown"+i,function(e){if(e.which===o.escape&&!a.config.widgetOptions.filter_resetOnEsc)return!1}).bind("keyup"+i,function(e){s=a.config.widgetOptions;var t=parseInt(A(this).attr("data-column"),10),r="boolean"==typeof s.filter_liveSearch?s.filter_liveSearch:E.getColumnData(a,s.filter_liveSearch,t);if(void 0===r&&(r=s.filter_liveSearch.fallback||!1),A(this).attr("data-lastSearchTime",(new Date).getTime()),e.which===o.escape)this.value=s.filter_resetOnEsc?"":l.lastSearch[t];else{if(""!==this.value&&("number"==typeof r&&this.value.length<r||e.which!==o.enter&&e.which!==o.backSpace&&(e.which<o.space||e.which>=o.left&&e.which<=o.down)))return;if(!1===r&&""!==this.value&&e.which!==o.enter)return}I.searching(a,!0,!0,t)}).bind("search change keypress input blur ".split(" ").join(i+" "),function(e){var t=parseInt(A(this).attr("data-column"),10),r=e.type,i="boolean"==typeof s.filter_liveSearch?s.filter_liveSearch:E.getColumnData(a,s.filter_liveSearch,t);!a.config.widgetOptions.filter_initialized||e.which!==o.enter&&"search"!==r&&"blur"!==r&&("change"!==r&&"input"!==r||!0!==i&&(!0===i||"INPUT"===e.target.nodeName)||this.value===l.lastSearch[t])||(e.preventDefault(),A(this).attr("data-lastSearchTime",(new Date).getTime()),I.searching(a,"keypress"!==r||e.which===o.enter,!0,t))}))},searching:function(e,t,r,i){var a,l=e.config.widgetOptions;void 0===i?a=!1:void 0===(a="boolean"==typeof l.filter_liveSearch?l.filter_liveSearch:E.getColumnData(e,l.filter_liveSearch,i))&&(a=l.filter_liveSearch.fallback||!1),clearTimeout(l.filter_searchTimer),void 0===t||!0===t?l.filter_searchTimer=setTimeout(function(){I.checkFilters(e,t,r)},a?l.filter_searchDelay:10):I.checkFilters(e,t,r)},equalFilters:function(e,t,r){var i,a=[],l=[],s=e.columns+1;for(t=A.isArray(t)?t:[],r=A.isArray(r)?r:[],i=0;i<s;i++)a[i]=t[i]||"",l[i]=r[i]||"";return a.join(",")===l.join(",")},checkFilters:function(e,t,r){var i=e.config,a=i.widgetOptions,l=A.isArray(t),s=l?t:E.getFilters(e,!0),n=s||[];if(A.isEmptyObject(i.cache))i.delayInit&&(!i.pager||i.pager&&i.pager.initialized)&&E.updateCache(i,function(){I.checkFilters(e,!1,r)});else{if(l&&(E.setFilters(e,s,!1,!0!==r),a.filter_initialized||(i.lastSearch=[],i.lastCombinedFilter="")),a.filter_hideFilters&&i.$table.find("."+g.filterRow).triggerHandler(I.hideFiltersCheck(i)?"mouseleave":"mouseenter"),I.equalFilters(i,i.lastSearch,n)){if(!1!==t)return;i.lastCombinedFilter="",i.lastSearch=[]}if(s=s||[],s=Array.prototype.map?s.map(String):s.join("�").split("�"),a.filter_initialized&&i.$table.triggerHandler("filterStart",[s]),!i.showProcessing)return I.findRows(e,s,n),!1;setTimeout(function(){return I.findRows(e,s,n),!1},30)}},hideFiltersCheck:function(e){if("function"==typeof e.widgetOptions.filter_hideFilters){var t=e.widgetOptions.filter_hideFilters(e);if("boolean"==typeof t)return t}return""===E.getFilters(e.$table).join("")},hideFilters:function(i,e){var a;(e||i.$table).find("."+g.filterRow).addClass(g.filterRowHide).bind("mouseenter mouseleave",function(e){var t=e,r=A(this);clearTimeout(a),a=setTimeout(function(){/enter|over/.test(t.type)?r.removeClass(g.filterRowHide):A(document.activeElement).closest("tr")[0]!==r[0]&&r.toggleClass(g.filterRowHide,I.hideFiltersCheck(i))},200)}).find("input, select").bind("focus blur",function(e){var t=e,r=A(this).closest("tr");clearTimeout(a),a=setTimeout(function(){clearTimeout(a),r.toggleClass(g.filterRowHide,I.hideFiltersCheck(i)&&"focus"!==t.type)},200)})},defaultFilter:function(e,t){if(""===e)return e;var r=O.iQuery,i=t.match(O.igQuery).length,a=1<i?A.trim(e).split(/\s/):[A.trim(e)],l=a.length-1,s=0,n=t;for(l<1&&1<i&&(a[1]=a[0]);r.test(n);)n=n.replace(r,a[s++]||""),r.test(n)&&s<l&&""!==(a[s]||"")&&(n=t.replace(r,n));return n},getLatestSearch:function(e){return e?e.sort(function(e,t){return A(t).attr("data-lastSearchTime")-A(e).attr("data-lastSearchTime")}):e||A()},findRange:function(e,t,r){var i,a,l,s,n,o,c,d,f=[];if(/^[0-9]+$/.test(t))return[parseInt(t,10)];if(!r&&/-/.test(t))for(d=(i=t.match(/(\d+)\s*-\s*(\d+)/g))?i.length:0,c=0;c<d;c++){for(a=i[c].split(/\s*-\s*/),l=parseInt(a[0],10)||0,(s=parseInt(a[1],10)||e.columns-1)<l&&(a=l,l=s,s=a),s>=e.columns&&(s=e.columns-1);l<=s;l++)f[f.length]=l;t=t.replace(i[c],"")}if(!r&&/,/.test(t))for(d=(n=t.split(/\s*,\s*/)).length,o=0;o<d;o++)""!==n[o]&&(c=parseInt(n[o],10))<e.columns&&(f[f.length]=c);if(!f.length)for(c=0;c<e.columns;c++)f[f.length]=c;return f},getColumnElm:function(t,e,r){return e.filter(function(){var e=I.findRange(t,A(this).attr("data-column"));return-1<A.inArray(r,e)})},multipleColumns:function(e,t){var r=e.widgetOptions,r=r.filter_initialized||!t.filter(r.filter_anyColumnSelector).length,t=A.trim(I.getLatestSearch(t).attr("data-column")||"");return I.findRange(e,t,!r)},processTypes:function(e,t,r){var i,a=null,l=null;for(i in I.types)A.inArray(i,r.excludeMatch)<0&&null===l&&null!==(l=I.types[i](e,t,r))&&(t.matchedOn=i,a=l);return a},matchType:function(e,t){var r=e.widgetOptions,i=e.$headerIndexed[t],r=!(i.hasClass("filter-exact")||!i.hasClass("filter-match")&&(r.filter_columnFilters?i=e.$filters.find("."+g.filter).add(r.filter_$externalFilters).filter('[data-column="'+t+'"]'):r.filter_$externalFilters&&(i=r.filter_$externalFilters.filter('[data-column="'+t+'"]')),!i.length||"match"!==e.widgetOptions.filter_matchType[(i[0].nodeName||"").toLowerCase()]));return r},processRow:function(t,r,e){var i,a,l,s,n,o=t.widgetOptions,c=!0,d=o.filter_$anyMatch&&o.filter_$anyMatch.length,f=o.filter_$anyMatch&&o.filter_$anyMatch.length?I.multipleColumns(t,o.filter_$anyMatch):[];if(r.$cells=r.$row.children(),r.matchedOn=null,r.anyMatchFlag&&1<f.length||r.anyMatchFilter&&!d){if(r.anyMatch=!0,r.isMatch=!0,r.rowArray=r.$cells.map(function(e){if(-1<A.inArray(e,f)||r.anyMatchFilter&&!d)return r.parsed[e]?n=r.cacheArray[e]:(n=r.rawArray[e],n=A.trim(o.filter_ignoreCase?n.toLowerCase():n),t.sortLocaleCompare&&(n=E.replaceAccents(n))),n}).get(),r.filter=r.anyMatchFilter,r.iFilter=r.iAnyMatchFilter,r.exact=r.rowArray.join(" "),r.iExact=o.filter_ignoreCase?r.exact.toLowerCase():r.exact,r.cache=r.cacheArray.slice(0,-1).join(" "),e.excludeMatch=e.noAnyMatch,null!==(a=I.processTypes(t,r,e)))c=a;else if(o.filter_startsWith)for(c=!1,f=Math.min(t.columns,r.rowArray.length);!c&&0<f;)f--,c=c||0===r.rowArray[f].indexOf(r.iFilter);else c=0<=(r.iExact+r.childRowText).indexOf(r.iFilter);if(r.anyMatch=!1,r.filters.join("")===r.filter)return c}for(f=0;f<t.columns;f++)r.filter=r.filters[f],r.index=f,e.excludeMatch=e.excludeFilter[f],r.filter&&(r.cache=r.cacheArray[f],i=r.parsed[f]?r.cache:r.rawArray[f]||"",r.exact=t.sortLocaleCompare?E.replaceAccents(i):i,r.iExact=!O.type.test(typeof r.exact)&&o.filter_ignoreCase?r.exact.toLowerCase():r.exact,r.isMatch=I.matchType(t,f),i=c,s=o.filter_columnFilters&&t.$filters.add(o.filter_$externalFilters).filter('[data-column="'+f+'"]').find("select option:selected").attr("data-function-name")||"",t.sortLocaleCompare&&(r.filter=E.replaceAccents(r.filter)),o.filter_defaultFilter&&O.iQuery.test(e.defaultColFilter[f])&&(r.filter=I.defaultFilter(r.filter,e.defaultColFilter[f])),r.iFilter=o.filter_ignoreCase?(r.filter||"").toLowerCase():r.filter,a=null,(l=e.functions[f])&&("function"==typeof l?a=l(r.exact,r.cache,r.filter,f,r.$row,t,r):"function"==typeof l[s||r.filter]&&(a=l[n=s||r.filter](r.exact,r.cache,r.filter,f,r.$row,t,r))),c=!!(i=null===a&&(a=I.processTypes(t,r,e),n=!0===l&&("and"===r.matchedOn||"or"===r.matchedOn),null===a||n)?!0===l?r.isMatch?0<=(""+r.iExact).search(r.iFilter):r.filter===r.exact:(n=(r.iExact+r.childRowText).indexOf(I.parseFilter(t,r.iFilter,r)),!o.filter_startsWith&&0<=n||o.filter_startsWith&&0===n):a)&&c);return c},findRows:function(e,r,t){if(!I.equalFilters(e.config,e.config.lastSearch,t)&&e.config.widgetOptions.filter_initialized){var i,a,l,s,n,o,c,d,f,h,u,p,g,m,b,y,_,v,w,x,S,C,z,$=A.extend([],r),F=e.config,R=F.widgetOptions,T=E.debug(F,"filter"),k={anyMatch:!1,filters:r,filter_regexCache:[]},H={noAnyMatch:["range","operators"],functions:[],excludeFilter:[],defaultColFilter:[],defaultAnyFilter:E.getColumnData(e,R.filter_defaultFilter,F.columns,!0)||""};for(k.parsed=[],f=0;f<F.columns;f++)k.parsed[f]=R.filter_useParsedData||F.parsers&&F.parsers[f]&&F.parsers[f].parsed||E.getData&&"parsed"===E.getData(F.$headerIndexed[f],E.getColumnData(e,F.headers,f),"filter")||F.$headerIndexed[f].hasClass("filter-parsed"),H.functions[f]=E.getColumnData(e,R.filter_functions,f)||F.$headerIndexed[f].hasClass("filter-select"),H.defaultColFilter[f]=E.getColumnData(e,R.filter_defaultFilter,f)||"",H.excludeFilter[f]=(E.getColumnData(e,R.filter_excludeFilter,f,!0)||"").split(/\s+/);for(T&&(console.log("Filter >> Starting filter widget search",r),m=new Date),F.filteredRows=0,t=$||[],c=F.totalRows=0;c<F.$tbodies.length;c++){if(d=E.processTbody(e,F.$tbodies.eq(c),!0),f=F.columns,a=F.cache[c].normalized,s=A(A.map(a,function(e){return e[f].$row.get()})),""===t.join("")||R.filter_serversideFiltering)s.removeClass(R.filter_filteredRow).not("."+F.cssChildRow).css("display","");else{if(i=(s=s.not("."+F.cssChildRow)).length,(R.filter_$anyMatch&&R.filter_$anyMatch.length||void 0!==r[F.columns])&&(k.anyMatchFlag=!0,k.anyMatchFilter=""+(r[F.columns]||R.filter_$anyMatch&&I.getLatestSearch(R.filter_$anyMatch).val()||""),R.filter_columnAnyMatch)){for(w=k.anyMatchFilter.split(O.andSplit),x=!1,y=0;y<w.length;y++)1<(S=w[y].split(":")).length&&(isNaN(S[0])?A.each(F.headerContent,function(e,t){-1<t.toLowerCase().indexOf(S[0])&&(r[C=e]=S[1])}):C=parseInt(S[0],10)-1,0<=C)&&C<F.columns&&(r[C]=S[1],w.splice(y,1),y--,x=!0);x&&(k.anyMatchFilter=w.join(" && "))}if(v=R.filter_searchFiltered,u=F.lastSearch||F.$table.data("lastSearch")||[],v)for(y=0;y<f+1;y++)b=r[y]||"",v||(y=f),v=v&&u.length&&0===b.indexOf(u[y]||"")&&!O.alreadyFiltered.test(b)&&!O.exactTest.test(b)&&!(O.isNeg1.test(b)||O.isNeg2.test(b))&&!(""!==b&&F.$filters&&F.$filters.filter('[data-column="'+y+'"]').find("select").length&&!I.matchType(F,y));for(_=s.not("."+R.filter_filteredRow).length,v&&0===_&&(v=!1),T&&console.log("Filter >> Searching through "+(v&&_<i?_:"all")+" rows"),k.anyMatchFlag&&(F.sortLocaleCompare&&(k.anyMatchFilter=E.replaceAccents(k.anyMatchFilter)),R.filter_defaultFilter&&O.iQuery.test(H.defaultAnyFilter)&&(k.anyMatchFilter=I.defaultFilter(k.anyMatchFilter,H.defaultAnyFilter),v=!1),k.iAnyMatchFilter=R.filter_ignoreCase&&F.ignoreCase?k.anyMatchFilter.toLowerCase():k.anyMatchFilter),o=0;o<i;o++)if(z=s[o].className,!(o&&O.child.test(z)||v&&O.filtered.test(z))){if(k.$row=s.eq(o),k.rowIndex=o,k.cacheArray=a[o],l=k.cacheArray[F.columns],k.rawArray=l.raw,k.childRowText="",!R.filter_childByColumn){for(z="",h=l.child,y=0;y<h.length;y++)z+=" "+h[y].join(" ")||"";k.childRowText=R.filter_childRows?R.filter_ignoreCase?z.toLowerCase():z:""}if(p=!1,g=I.processRow(F,k,H),n=l.$row,b=!!g,h=l.$row.filter(":gt(0)"),R.filter_childRows&&h.length){if(R.filter_childByColumn)for(R.filter_childWithSibs||(h.addClass(R.filter_filteredRow),n=n.eq(0)),y=0;y<h.length;y++)k.$row=h.eq(y),k.cacheArray=l.child[y],k.rawArray=k.cacheArray,b=I.processRow(F,k,H),p=p||b,!R.filter_childWithSibs&&b&&h.eq(y).removeClass(R.filter_filteredRow);p=p||g}else p=b;n.toggleClass(R.filter_filteredRow,!p)[0].display=p?"":"none"}}F.filteredRows+=s.not("."+R.filter_filteredRow).length,F.totalRows+=s.length,E.processTbody(e,d,!1)}F.lastCombinedFilter=$.join(""),F.lastSearch=$,F.$table.data("lastSearch",$),R.filter_saveFilters&&E.storage&&E.storage(e,"tablesorter-filters",I.processFilters($,!0)),T&&console.log("Filter >> Completed search"+E.benchmark(m)),R.filter_initialized&&(F.$table.triggerHandler("filterBeforeEnd",F),F.$table.triggerHandler("filterEnd",F)),setTimeout(function(){E.applyWidget(F.table)},0)}},getOptionSource:function(e,t,r){var i=(e=A(e)[0]).config,a=!1,l=i.widgetOptions.filter_selectSource,i=i.$table.data("lastSearch")||[],s="function"==typeof l||E.getColumnData(e,l,t);if(r&&""!==i[t]&&(r=!1),!0===s)a=l(e,t,r);else{if(s instanceof A||"string"===A.type(s)&&0<=s.indexOf("</option>"))return s;if(A.isArray(s))a=s;else if("object"===A.type(l)&&s&&null===(a=s(e,t,r)))return null}return!1===a&&(a=I.getOptions(e,t,r)),I.processOptions(e,t,a)},processOptions:function(i,a,r){if(!A.isArray(r))return!1;var l,e,t,s,n,o=(i=A(i)[0]).config,c=null!=a&&0<=a&&a<o.columns,d=!!c&&o.$headerIndexed[a].hasClass("filter-select-sort-desc"),f=[];if(r=A.grep(r,function(e,t){return!!e.text||A.inArray(e,r)===t}),!c||!o.$headerIndexed[a].hasClass("filter-select-nosort")){for(s=r.length,t=0;t<s;t++)n=(e=r[t]).text||e,n=(c&&o.parsers&&o.parsers.length&&o.parsers[a].format(n,i,[],a)||n).toString(),n=o.widgetOptions.filter_ignoreCase?n.toLowerCase():n,e.text?(e.parsed=n,f[f.length]=e):f[f.length]={text:e,parsed:n};for(l=o.textSorter||"",f.sort(function(e,t){var r=(d?t:e).parsed,e=(d?e:t).parsed;return c&&"function"==typeof l?l(r,e,!0,a,i):c&&"object"==typeof l&&l.hasOwnProperty(a)?l[a](r,e,!0,a,i):!E.sortNatural||E.sortNatural(r,e)}),r=[],s=f.length,t=0;t<s;t++)r[r.length]=f[t]}return r},getOptions:function(e,t,r){for(var i,a,l,s,n,o,c,d=(e=A(e)[0]).config,f=d.widgetOptions,h=[],u=0;u<d.$tbodies.length;u++)for(s=d.cache[u],a=d.cache[u].normalized.length,i=0;i<a;i++)if(l=s.row?s.row[i]:s.normalized[i][d.columns].$row[0],!r||!l.className.match(f.filter_filteredRow))if(f.filter_useParsedData||d.parsers[t].parsed||d.$headerIndexed[t].hasClass("filter-parsed")){if(h[h.length]=""+s.normalized[i][t],f.filter_childRows&&f.filter_childByColumn)for(c=s.normalized[i][d.columns].$row.length-1,n=0;n<c;n++)h[h.length]=""+s.normalized[i][d.columns].child[n][t]}else if(h[h.length]=s.normalized[i][d.columns].raw[t],f.filter_childRows&&f.filter_childByColumn)for(c=s.normalized[i][d.columns].$row.length,n=1;n<c;n++)o=s.normalized[i][d.columns].$row.eq(n).children().eq(t),h[h.length]=""+E.getElementText(d,o,t);return h},buildSelect:function(e,t,r,i,a){if(e=A(e)[0],t=parseInt(t,10),e.config.cache&&!A.isEmptyObject(e.config.cache)){var l,s,n,o,c,d=e.config,f=d.widgetOptions,h=d.$headerIndexed[t],u='<option value="">'+(h.data("placeholder")||h.attr("data-placeholder")||f.filter_placeholder.select||"")+"</option>",h=d.$table.find("thead").find("select."+g.filter+'[data-column="'+t+'"]').val();if(void 0!==r&&""!==r||null!==(r=I.getOptionSource(e,t,a))){if(A.isArray(r)){for(l=0;l<r.length;l++)if((c=r[l]).text){for(s in c["data-function-name"]=void 0===c.value?c.text:c.value,u+="<option",c)c.hasOwnProperty(s)&&"text"!==s&&(u+=" "+s+'="'+c[s].replace(O.quote,"&quot;")+'"');c.value||(u+=' value="'+c.text.replace(O.quote,"&quot;")+'"'),u+=">"+c.text.replace(O.quote,"&quot;")+"</option>"}else""+c!="[object Object]"&&(0<=(s=n=c=(""+c).replace(O.quote,"&quot;")).indexOf(f.filter_selectSourceSeparator)&&(s=(o=n.split(f.filter_selectSourceSeparator))[0],n=o[1]),u+=""!==c?"<option "+(s===n?"":'data-function-name="'+c+'" ')+'value="'+s+'">'+n+"</option>":"");r=[]}e=(d.$filters||d.$table.children("thead")).find("."+g.filter),(a=(e=f.filter_$externalFilters?e&&e.length?e.add(f.filter_$externalFilters):f.filter_$externalFilters:e).filter('select[data-column="'+t+'"]')).length&&(a[i?"html":"append"](u),A.isArray(r)||a.append(r).val(h),a.val(h))}}},buildDefault:function(e,t){for(var r,i,a=e.config,l=a.widgetOptions,s=a.columns,n=0;n<s;n++)i=!((r=a.$headerIndexed[n]).hasClass("filter-false")||r.hasClass("parser-false")),(r.hasClass("filter-select")||!0===E.getColumnData(e,l.filter_functions,n))&&i&&I.buildSelect(e,n,"",t,r.hasClass(l.filter_onlyAvail))}},O=I.regex,E.getFilters=function(e,t,r,i){var a,l,s,n,o=[],c=e?A(e)[0].config:"",d=c?c.widgetOptions:"";if(!0!==t&&d&&!d.filter_columnFilters||A.isArray(r)&&I.equalFilters(c,r,c.lastSearch))return A(e).data("lastSearch")||[];if(c&&(c.$filters&&(l=c.$filters.find("."+g.filter)),l=d.filter_$externalFilters?l&&l.length?l.add(d.filter_$externalFilters):d.filter_$externalFilters:l)&&l.length)for(o=r||[],a=0;a<c.columns+1;a++)n=a===c.columns?d.filter_anyColumnSelector+","+d.filter_multipleColumnSelector:'[data-column="'+a+'"]',(s=l.filter(n)).length&&(s=I.getLatestSearch(s),A.isArray(r)?(i&&1<s.length&&(s=s.slice(1)),(s=a===c.columns?(n=s.filter(d.filter_anyColumnSelector)).length?n:s:s).val(r[a]).trigger("change"+c.namespace)):(o[a]=s.val()||"",(a===c.columns?s.slice(1).filter('[data-column*="'+s.attr("data-column")+'"]'):s.slice(1)).val(o[a])),a===c.columns)&&s.length&&(d.filter_$anyMatch=s);return o},E.setFilters=function(e,t,r,i){var a=e?A(e)[0].config:"",e=E.getFilters(e,!0,t,i);return void 0===r&&(r=!0),a&&r&&(a.lastCombinedFilter=null,a.lastSearch=[],I.searching(a.table,t,i),a.$table.triggerHandler("filterFomatterUpdate")),0!==e.length}}(e),function(C,z){"use strict";var $=C.tablesorter||{};function F(e,t){var r=isNaN(t.stickyHeaders_offset)?C(t.stickyHeaders_offset):[];return r.length?r.height()||0:parseInt(t.stickyHeaders_offset,10)||0}C.extend($.css,{sticky:"tablesorter-stickyHeader",stickyVis:"tablesorter-sticky-visible",stickyHide:"tablesorter-sticky-hidden",stickyWrap:"tablesorter-sticky-wrapper"}),$.addHeaderResizeEvent=function(e,t,r){if((e=C(e)[0]).config){var r=C.extend({},{timer:250},r),o=e.config,c=o.widgetOptions,i=function(e){var t,r,i,a,l,s,n=o.$headers.length;for(c.resize_flag=!0,r=[],t=0;t<n;t++)a=(i=o.$headers.eq(t)).data("savedSizes")||[0,0],l=i[0].offsetWidth,s=i[0].offsetHeight,l===a[0]&&s===a[1]||(i.data("savedSizes",[l,s]),r.push(i[0]));r.length&&!1!==e&&o.$table.triggerHandler("resize",[r]),c.resize_flag=!1};if(clearInterval(c.resize_timer),t)return c.resize_flag=!1;i(!1),c.resize_timer=setInterval(function(){c.resize_flag||i()},r.timer)}},$.addWidget({id:"stickyHeaders",priority:54,options:{stickyHeaders:"",stickyHeaders_appendTo:null,stickyHeaders_attachTo:null,stickyHeaders_xScroll:null,stickyHeaders_yScroll:null,stickyHeaders_offset:0,stickyHeaders_filteredToTop:!0,stickyHeaders_cloneId:"-sticky",stickyHeaders_addResizeEvent:!0,stickyHeaders_includeCaption:!0,stickyHeaders_zIndex:2},format:function(e,t,o){if(!(t.$table.hasClass("hasStickyHeaders")||0<=C.inArray("filter",t.widgets)&&!t.$table.hasClass("hasFilters"))){var r,i,a,l,c=t.$table,d=C(o.stickyHeaders_attachTo||o.stickyHeaders_appendTo),s=t.namespace+"stickyheaders ",f=C(o.stickyHeaders_yScroll||o.stickyHeaders_attachTo||z),n=C(o.stickyHeaders_xScroll||o.stickyHeaders_attachTo||z),h=c.children("thead:first").children("tr").not(".sticky-false").children(),u=c.children("tfoot"),p=F(0,o),g=c.parent().closest("."+$.css.table).hasClass("hasStickyHeaders")?c.parent().closest("table.tablesorter")[0].config.widgetOptions.$sticky.parent():[],m=g.length?g.height():0,b=o.$sticky=c.clone().addClass("containsStickyHeaders "+$.css.sticky+" "+o.stickyHeaders+" "+t.namespace.slice(1)+"_extra_table").wrap('<div class="'+$.css.stickyWrap+'">'),y=b.parent().addClass($.css.stickyHide).css({position:d.length?"absolute":"fixed",padding:parseInt(b.parent().parent().css("padding-left"),10),top:p+m,left:0,visibility:"hidden",zIndex:o.stickyHeaders_zIndex||2}),p=b.children("thead:first"),_="",v=function(e,t){for(var r,i,a,l=e.filter(":visible"),s=l.length,n=0;n<s;n++)i=t.filter(":visible").eq(n),r="border-box"===(a=l.eq(n)).css("box-sizing")?a.outerWidth():"collapse"===i.css("border-collapse")?z.getComputedStyle?parseFloat(z.getComputedStyle(a[0],null).width):(r=parseFloat(a.css("border-width")),a.outerWidth()-parseFloat(a.css("padding-left"))-parseFloat(a.css("padding-right"))-r):a.width(),i.css({width:r,"min-width":r,"max-width":r})},w=function(e){return!1===e&&g.length?c.position().left:d.length?parseInt(d.css("padding-left"),10)||0:c.offset().left-parseInt(c.css("margin-left"),10)-C(z).scrollLeft()},x=function(){y.css({left:w(),width:c.outerWidth()}),v(c,b),v(h,l)},S=function(e){var t,r,i,a,l,s,n;c.is(":visible")&&(m=g.length?g.offset().top-f.scrollTop()+g.height():0,a=c.offset(),t=F(0,o),i=(r=C.isWindow(f[0]))?f.scrollTop():g.length?parseInt(g[0].style.top,10):f.offset().top,i=d.length?i:f.scrollTop(),l=!o.stickyHeaders_includeCaption&&c.children("caption").height()||0,i=i+t+m-l,l=c.height()-(y.height()+(u.height()||0))-l,l="visible"==(a=i>a.top&&i<a.top+l?"visible":"hidden")?$.css.stickyVis:$.css.stickyHide,s=!y.hasClass(l),n={visibility:a},d.length&&(s=!0,n.top=r?i-d.offset().top:d.scrollTop()),(i=w(r))!==parseInt(y.css("left"),10)&&(s=!0,n.left=i),n.top=(n.top||0)+(!r&&g.length?g.height():t+m),s&&y.removeClass($.css.stickyVis+" "+$.css.stickyHide).addClass(l).css(n),a===_&&!e||(x(),_=a))};if(d.length&&!d.css("position")&&d.css("position","relative"),b.attr("id")&&(b[0].id+=o.stickyHeaders_cloneId),b.find("> thead:gt(0), tr.sticky-false").hide(),b.find("> tbody, > tfoot").remove(),b.find("caption").toggle(o.stickyHeaders_includeCaption),l=p.children().children(),b.css({height:0,width:0,margin:0}),l.find("."+$.css.resizer).remove(),c.addClass("hasStickyHeaders").bind("pagerComplete"+s,function(){x()}),$.bindEvents(e,p.children().children("."+$.css.header)),o.stickyHeaders_appendTo?C(o.stickyHeaders_appendTo).append(y):c.after(y),t.onRenderHeader)for(i=(a=p.children("tr").children()).length,r=0;r<i;r++)t.onRenderHeader.apply(a.eq(r),[r,t,b]);n.add(f).unbind("scroll resize ".split(" ").join(s).replace(/\s+/g," ")).bind("scroll resize ".split(" ").join(s),function(e){S("resize"===e.type)}),t.$table.unbind("stickyHeadersUpdate"+s).bind("stickyHeadersUpdate"+s,function(){S(!0)}),o.stickyHeaders_addResizeEvent&&$.addHeaderResizeEvent(e),c.hasClass("hasFilters")&&o.filter_columnFilters&&(c.bind("filterEnd"+s,function(){var e=C(document.activeElement).closest("td"),e=e.parent().children().index(e);y.hasClass($.css.stickyVis)&&o.stickyHeaders_filteredToTop&&(z.scrollTo(0,c.position().top),0<=e)&&t.$filters&&t.$filters.eq(e).find("a, select, input").filter(":visible").focus()}),$.filter.bindSearch(c,l.find("."+$.css.filter)),o.filter_hideFilters)&&$.filter.hideFilters(t,b),o.stickyHeaders_addResizeEvent&&c.bind("resize"+t.namespace+"stickyheaders",function(){x()}),S(!0),c.triggerHandler("stickyHeadersInit")}},remove:function(e,t,r){var i=t.namespace+"stickyheaders ";t.$table.removeClass("hasStickyHeaders").unbind("pagerComplete resize filterEnd stickyHeadersUpdate ".split(" ").join(i).replace(/\s+/g," ")).next("."+$.css.stickyWrap).remove(),r.$sticky&&r.$sticky.length&&r.$sticky.remove(),C(z).add(r.stickyHeaders_xScroll).add(r.stickyHeaders_yScroll).add(r.stickyHeaders_attachTo).unbind("scroll resize ".split(" ").join(i).replace(/\s+/g," ")),$.addHeaderResizeEvent(e,!0)}})}(e,window),function(c,t){"use strict";var d=c.tablesorter||{};c.extend(d.css,{resizableContainer:"tablesorter-resizable-container",resizableHandle:"tablesorter-resizable-handle",resizableNoSelect:"tablesorter-disableSelection",resizableStorage:"tablesorter-resizable"}),c(function(){var e="<style>body."+d.css.resizableNoSelect+" { -ms-user-select: none; -moz-user-select: -moz-none;-khtml-user-select: none; -webkit-user-select: none; user-select: none; }."+d.css.resizableContainer+" { position: relative; height: 1px; }."+d.css.resizableHandle+" { position: absolute; display: inline-block; width: 8px;top: 1px; cursor: ew-resize; z-index: 3; user-select: none; -moz-user-select: none; }</style>";c("head").append(e)}),d.resizable={init:function(e,t){if(!e.$table.hasClass("hasResizable")){e.$table.addClass("hasResizable");var r,i,a,l=e.$table,s=l.parent(),n=parseInt(l.css("margin-top"),10),o=t.resizable_vars={useStorage:d.storage&&!1!==t.resizable,$wrap:s,mouseXPosition:0,$target:null,$next:null,overflow:"auto"===s.css("overflow")||"scroll"===s.css("overflow")||"auto"===s.css("overflow-x")||"scroll"===s.css("overflow-x"),storedSizes:[]};for(d.resizableReset(e.table,!0),o.tableWidth=l.width(),o.fullWidth=Math.abs(s.width()-o.tableWidth)<20,o.useStorage&&o.overflow&&(d.storage(e.table,"tablesorter-table-original-css-width",o.tableWidth),a=d.storage(e.table,"tablesorter-table-resized-width")||"auto",d.resizable.setWidth(l,a,!0)),t.resizable_vars.storedSizes=s=(o.useStorage?d.storage(e.table,d.css.resizableStorage):[])||[],d.resizable.setWidths(e,t,s),d.resizable.updateStoredSizes(e,t),t.$resizable_container=c('<div class="'+d.css.resizableContainer+'">').css({top:n}).insertBefore(l),i=0;i<e.columns;i++)r=e.$headerIndexed[i],a=d.getColumnData(e.table,e.headers,i),"false"===d.getData(r,a,"resizable")||c('<div class="'+d.css.resizableHandle+'">').appendTo(t.$resizable_container).attr({"data-column":i,unselectable:"on"}).data("header",r).bind("selectstart",!1);d.resizable.bindings(e,t)}},updateStoredSizes:function(e,t){var r,i,a=e.columns,l=t.resizable_vars;for(l.storedSizes=[],r=0;r<a;r++)i=e.$headerIndexed[r],l.storedSizes[r]=i.is(":visible")?i.width():0},setWidth:function(e,t,r){e.css({width:t,"min-width":r?t:"","max-width":r?t:""})},setWidths:function(e,t,r){var i,a,l=t.resizable_vars,s=c(e.namespace+"_extra_headers"),n=e.$table.children("colgroup").children("col");if((r=r||l.storedSizes||[]).length){for(i=0;i<e.columns;i++)d.resizable.setWidth(e.$headerIndexed[i],r[i],l.overflow),s.length&&(a=s.eq(i).add(n.eq(i)),d.resizable.setWidth(a,r[i],l.overflow));(a=c(e.namespace+"_extra_table")).length&&!d.hasWidget(e.table,"scroller")&&d.resizable.setWidth(a,e.$table.outerWidth(),l.overflow)}},setHandlePosition:function(a,l){var s,n=a.$table.height(),e=l.$resizable_container.children(),o=Math.floor(e.width()/2);d.hasWidget(a.table,"scroller")&&(n=0,a.$table.closest("."+d.css.scrollerWrap).children().each(function(){var e=c(this);n+=(e.filter('[style*="height"]').length?e:e.children("table")).height()})),!l.resizable_includeFooter&&a.$table.children("tfoot").length&&(n-=a.$table.children("tfoot").height()),s=3.3<=parseFloat(c.fn.jquery)?0:a.$table.position().left,e.each(function(){var e=c(this),t=parseInt(e.attr("data-column"),10),r=a.columns-1,i=e.data("header");i&&(!i.is(":visible")||!l.resizable_addLastColumn&&d.resizable.checkVisibleColumns(a,t)?e.hide():(t<r||t===r&&l.resizable_addLastColumn)&&e.css({display:"inline-block",height:n,left:i.position().left-s+i.outerWidth()-o}))})},checkVisibleColumns:function(e,t){for(var r=0,i=t+1;i<e.columns;i++)r+=e.$headerIndexed[i].is(":visible")?1:0;return 0===r},toggleTextSelection:function(e,t,r){e=e.namespace+"tsresize";t.resizable_vars.disabled=r,c("body").toggleClass(d.css.resizableNoSelect,r),r?c("body").attr("unselectable","on").bind("selectstart"+e,!1):c("body").removeAttr("unselectable").unbind("selectstart"+e)},bindings:function(l,s){var e=l.namespace+"tsresize";s.$resizable_container.children().bind("mousedown",function(e){var t=s.resizable_vars,r=c(l.namespace+"_extra_headers"),i=c(e.target).data("header"),a=parseInt(i.attr("data-column"),10);t.$target=i=i.add(r.filter('[data-column="'+a+'"]')),t.target=a,t.$next=e.shiftKey||s.resizable_targetLast?i.parent().children().not(".resizable-false").filter(":last"):i.nextAll(":not(.resizable-false)").eq(0),a=parseInt(t.$next.attr("data-column"),10),t.$next=t.$next.add(r.filter('[data-column="'+a+'"]')),t.next=a,t.mouseXPosition=e.pageX,d.resizable.updateStoredSizes(l,s),d.resizable.toggleTextSelection(l,s,!0)}),c(document).bind("mousemove"+e,function(e){var t=s.resizable_vars;t.disabled&&0!==t.mouseXPosition&&t.$target&&(s.resizable_throttle?(clearTimeout(t.timer),t.timer=setTimeout(function(){d.resizable.mouseMove(l,s,e)},isNaN(s.resizable_throttle)?5:s.resizable_throttle)):d.resizable.mouseMove(l,s,e))}).bind("mouseup"+e,function(){s.resizable_vars.disabled&&(d.resizable.toggleTextSelection(l,s,!1),d.resizable.stopResize(l,s),d.resizable.setHandlePosition(l,s))}),c(t).bind("resize"+e+" resizeEnd"+e,function(){d.resizable.setHandlePosition(l,s)}),l.$table.bind("columnUpdate pagerComplete resizableUpdate ".split(" ").join(e+" "),function(){d.resizable.setHandlePosition(l,s)}).bind("resizableReset"+e,function(){d.resizableReset(l.table)}).find("thead:first").add(c(l.namespace+"_extra_table").find("thead:first")).bind("contextmenu"+e,function(){var e=0===s.resizable_vars.storedSizes.length;return d.resizableReset(l.table),d.resizable.setHandlePosition(l,s),s.resizable_vars.storedSizes=[],e})},mouseMove:function(e,t,r){if(0!==t.resizable_vars.mouseXPosition&&t.resizable_vars.$target){var i,a=0,l=t.resizable_vars,s=l.$next,n=l.storedSizes[l.target],o=r.pageX-l.mouseXPosition;if(l.overflow){if(0<n+o){for(l.storedSizes[l.target]+=o,d.resizable.setWidth(l.$target,l.storedSizes[l.target],!0),i=0;i<e.columns;i++)a+=l.storedSizes[i];d.resizable.setWidth(e.$table.add(c(e.namespace+"_extra_table")),a)}s.length||(l.$wrap[0].scrollLeft=e.$table.width())}else l.fullWidth?(l.storedSizes[l.target]+=o,l.storedSizes[l.next]-=o):l.storedSizes[l.target]+=o,d.resizable.setWidths(e,t);l.mouseXPosition=r.pageX,e.$table.triggerHandler("stickyHeadersUpdate")}},stopResize:function(e,t){var r=t.resizable_vars;d.resizable.updateStoredSizes(e,t),r.useStorage&&(d.storage(e.table,d.css.resizableStorage,r.storedSizes),d.storage(e.table,"tablesorter-table-resized-width",e.$table.width())),r.mouseXPosition=0,r.$target=r.$next=null,e.$table.triggerHandler("stickyHeadersUpdate"),e.$table.triggerHandler("resizableComplete")}},d.addWidget({id:"resizable",priority:40,options:{resizable:!0,resizable_addLastColumn:!1,resizable_includeFooter:!0,resizable_widths:[],resizable_throttle:!1,resizable_targetLast:!1},init:function(e,t,r,i){d.resizable.init(r,i)},format:function(e,t,r){d.resizable.setHandlePosition(t,r)},remove:function(e,t,r,i){var a;r.$resizable_container&&(a=t.namespace+"tsresize",t.$table.add(c(t.namespace+"_extra_table")).removeClass("hasResizable").children("thead").unbind("contextmenu"+a),r.$resizable_container.remove(),d.resizable.toggleTextSelection(t,r,!1),d.resizableReset(e,i),c(document).unbind("mousemove"+a+" mouseup"+a))}}),d.resizableReset=function(l,s){c(l).each(function(){var e,t,r=this.config,i=r&&r.widgetOptions,a=i.resizable_vars;if(l&&r&&r.$headerIndexed.length){for(a.overflow&&a.tableWidth&&(d.resizable.setWidth(r.$table,a.tableWidth,!0),a.useStorage)&&d.storage(l,"tablesorter-table-resized-width",a.tableWidth),e=0;e<r.columns;e++)t=r.$headerIndexed[e],i.resizable_widths&&i.resizable_widths[e]?d.resizable.setWidth(t,i.resizable_widths[e],a.overflow):t.hasClass("resizable-false")||d.resizable.setWidth(t,"",a.overflow);r.$table.triggerHandler("stickyHeadersUpdate"),d.storage&&!s&&d.storage(this,d.css.resizableStorage,[])}})}}(e,window),function(t){"use strict";var o=t.tablesorter||{};function c(e){e=o.storage(e.table,"tablesorter-savesort");return e&&e.hasOwnProperty("sortList")&&t.isArray(e.sortList)?e.sortList:[]}function d(e,t){return(t||c(e)).join(",")!==e.sortList.join(",")}o.addWidget({id:"saveSort",priority:20,options:{saveSort:!0},init:function(e,t,r,i){t.format(e,r,i,!0)},format:function(t,e,r,i){var a,l=e.$table,r=!1!==r.saveSort,s={sortList:e.sortList},n=o.debug(e,"saveSort");n&&(a=new Date),l.hasClass("hasSaveSort")?r&&t.hasInitialized&&o.storage&&d(e)&&(o.storage(t,"tablesorter-savesort",s),n)&&console.log("saveSort >> Saving last sort: "+e.sortList+o.benchmark(a)):(l.addClass("hasSaveSort"),s="",o.storage&&(s=c(e),n&&console.log('saveSort >> Last sort loaded: "'+s+'"'+o.benchmark(a)),l.bind("saveSortReset",function(e){e.stopPropagation(),o.storage(t,"tablesorter-savesort","")})),i&&s&&0<s.length?e.sortList=s:t.hasInitialized&&s&&0<s.length&&d(e,s)&&o.sortOn(e,s))},remove:function(e,t){t.$table.removeClass("hasSaveSort"),o.storage&&o.storage(e,"tablesorter-savesort","")}})}(e),e.tablesorter});return jQuery;}));
```
