## Supplementary Data SD1 for "Inferring binding specificities of human transcription factors with the wisdom of crowds": parser-date-extract.min.js

```
(function(factory){if (typeof define === 'function' && define.amd){define(['jquery'], factory);} else if (typeof module === 'object' && typeof module.exports === 'object'){module.exports = factory(require('jquery'));} else {factory(jQuery);}}(function(jQuery){
/*! Parser: Extract out date - updated 10/26/2014 (v2.18.0) */
!function(e){"use strict";var r=/[A-Z]{3,10}\.?\s+\d{1,2},?\s+(?:\d{4})(?:\s+\d{1,2}:\d{2}(?::\d{2})?(?:\s+[AP]M)?)?/i,a=/(\d{1,2}[\/\s]\d{1,2}[\/\s]\d{4}(\s+\d{1,2}:\d{2}(:\d{2})?(\s+[AP]M)?)?)/i,n=/(\d{1,2}[\/\s]\d{1,2}[\/\s]\d{4}(\s+\d{1,2}:\d{2}(:\d{2})?(\s+[AP]M)?)?)/i,i=/(\d{1,2})[\/\s](\d{1,2})[\/\s](\d{4})/,s=/(\d{4}[\/\s]\d{1,2}[\/\s]\d{1,2}(\s+\d{1,2}:\d{2}(:\d{2})?(\s+[AP]M)?)?)/i,d=/(\d{4})[\/\s](\d{1,2})[\/\s](\d{1,2})/;
/*! extract US Long Date */e.tablesorter.addParser({id:"extractUSLongDate",is:function(){return!1},format:function(e){var t=e&&e.match(r);return t&&(t=new Date(t[0]))instanceof Date&&isFinite(t)?t.getTime():e},type:"numeric"}),
/*! extract MMDDYYYY */
e.tablesorter.addParser({id:"extractMMDDYYYY",is:function(){return!1},format:function(e){var t=e&&e.replace(/\s+/g," ").replace(/[\-.,]/g,"/").match(a);return t&&(t=new Date(t[0]))instanceof Date&&isFinite(t)?t.getTime():e},type:"numeric"}),
/*! extract DDMMYYYY */
e.tablesorter.addParser({id:"extractDDMMYYYY",is:function(){return!1},format:function(e){var t=e&&e.replace(/\s+/g," ").replace(/[\-.,]/g,"/").match(n);return t&&(t=new Date(t[0].replace(i,"$2/$1/$3")))instanceof Date&&isFinite(t)?t.getTime():e},type:"numeric"}),
/*! extract YYYYMMDD */
e.tablesorter.addParser({id:"extractYYYYMMDD",is:function(){return!1},format:function(e){var t=e&&e.replace(/\s+/g," ").replace(/[\-.,]/g,"/").match(s);return t&&(t=new Date(t[0].replace(d,"$2/$3/$1")))instanceof Date&&isFinite(t)?t.getTime():e},type:"numeric"})}(jQuery);return jQuery;}));
```
