## Supplementary Data SD1 for "Inferring binding specificities of human transcription factors with the wisdom of crowds": parser-date-iso8601.min.js

```
(function(factory){if (typeof define === 'function' && define.amd){define(['jquery'], factory);} else if (typeof module === 'object' && typeof module.exports === 'object'){module.exports = factory(require('jquery'));} else {factory(jQuery);}}(function(jQuery){
/*! Parser: ISO-8601 date - updated 10/26/2014 (v2.18.0) */
!function(e){"use strict";var r=/^([0-9]{4})(-([0-9]{2})(-([0-9]{2})(T([0-9]{2}):([0-9]{2})(:([0-9]{2})(\.([0-9]+))?)?(Z|(([-+])([0-9]{2}):([0-9]{2})))?)?)?)?$/;e.tablesorter.addParser({id:"iso8601date",is:function(e){return!!e&&e.match(r)},format:function(e){var t,s=e&&e.match(r);return s?(t=new Date(s[1],0,1),s[3]&&t.setMonth(s[3]-1),s[5]&&t.setDate(s[5]),s[7]&&t.setHours(s[7]),s[8]&&t.setMinutes(s[8]),s[10]&&t.setSeconds(s[10]),s[12]&&t.setMilliseconds(1e3*Number("0."+s[12])),t.getTime()):e},type:"numeric"})}(jQuery);return jQuery;}));
```
