## Supplementary Data SD1 for "Inferring binding specificities of human transcription factors with the wisdom of crowds": parser-date-month.min.js

```
(function(factory){if (typeof define === 'function' && define.amd){define(['jquery'], factory);} else if (typeof module === 'object' && typeof module.exports === 'object'){module.exports = factory(require('jquery'));} else {factory(jQuery);}}(function(jQuery){
/*! Parser: Month - updated 11/22/2015 (v2.24.6) */
!function(){"use strict";var u=jQuery.tablesorter;u.dates||(u.dates={}),u.dates.months||(u.dates.months={}),u.dates.months.en={1:"Jan",2:"Feb",3:"Mar",4:"Apr",5:"May",6:"Jun",7:"Jul",8:"Aug",9:"Sep",10:"Oct",11:"Nov",12:"Dec"},u.addParser({id:"month",is:function(){return!1},format:function(e,t,n,a){if(e){var r,o,s=t.config,t=s.globalize&&(s.globalize[a]||s.globalize)||{},i=u.dates.months[t.lang||"en"];for(o in s.ignoreCase&&(e=e.toLowerCase()),i)if("string"==typeof o&&(r=i[o],s.ignoreCase&&(r=r.toLowerCase()),e.match(r)))return parseInt(o,10)}return e},type:"numeric"})}();return jQuery;}));
```
