## Supplementary Data SD1 for "Inferring binding specificities of human transcription factors with the wisdom of crowds": parser-date-range.min.js

```
(function(factory){if (typeof define === 'function' && define.amd){define(['jquery'], factory);} else if (typeof module === 'object' && typeof module.exports === 'object'){module.exports = factory(require('jquery'));} else {factory(jQuery);}}(function(jQuery){
/*! Parser: date ranges -updated 11/22/2015 (v2.24.6) */
!function(e){"use strict";var u,f=e.tablesorter,i=/(\d{1,2}[-\s]\d{1,2}[-\s]\d{4}(\s+\d{1,2}:\d{2}(:\d{2})?(\s+[AP]M)?)?)/gi,d=/(\d{1,2}[-\s]\d{1,2}[-\s]\d{4}(\s+\d{1,2}:\d{2}(:\d{2})?(\s+[AP]M)?)?)/gi,o=/(\d{1,2})[-\s](\d{1,2})[-\s](\d{4})/,c=/(\d{4}[-\s]\d{1,2}[-\s]\d{1,2}(\s+\d{1,2}:\d{2}(:\d{2})?(\s+[AP]M)?)?)/gi,g=/(\d{4})[-\s](\d{1,2})[-\s](\d{1,2})/,l=/(\d{1,2}\s+\w+\s+\d{4}(\s+\d{1,2}:\d{2}(:\d{2})?(\s\w+)?)?)/g,p=/(\d{1,2})\s+(\w+)\s+(\d{4})/;
/*! date-range MMDDYYYY */e.tablesorter.addParser({id:"date-range-mdy",is:function(){return!1},format:function(e){var t,r,a=[],n=e.replace(/\s+/g," ").replace(/[\/\-.,]/g,"-").match(i),s=n&&n.length;if(s){for(r=0;r<s;r++)t=new Date(n[r]),a.push(t instanceof Date&&isFinite(t)?t.getTime():n[r]);return a.sort().join(" - ")}return e},type:"text"}),
/*! date-range DDMMYYYY */
e.tablesorter.addParser({id:"date-range-dmy",is:function(){return!1},format:function(e){var t,r,a=[],n=e.replace(/\s+/g," ").replace(/[\/\-.,]/g,"-").match(d),s=n&&n.length;if(s){for(r=0;r<s;r++)t=new Date((""+n[r]).replace(o,"$2/$1/$3")),a.push(t instanceof Date&&isFinite(t)?t.getTime():n[r]);return a.sort().join(" - ")}return e},type:"text"}),
/*! date-range DDMMYYYY */
e.tablesorter.addParser({id:"date-range-ymd",is:function(){return!1},format:function(e){var t,r,a=[],n=e.replace(/\s+/g," ").replace(/[\/\-.,]/g,"-").match(c),s=n&&n.length;if(s){for(r=0;r<s;r++)t=new Date((""+n[r]).replace(g,"$2/$3/$1")),a.push(t instanceof Date&&isFinite(t)?t.getTime():n[r]);return a.sort().join(" - ")}return e},type:"text"}),f.dates||(f.dates={}),f.dates.months||(f.dates.months={}),f.dates.months.en={1:"Jan",2:"Feb",3:"Mar",4:"Apr",5:"May",6:"Jun",7:"Jul",8:"Aug",9:"Sep",10:"Oct",11:"Nov",12:"Dec"},u=function(e,t,r){var a,n,r=t.globalize&&(t.globalize[r]||t.globalize)||{},s=f.dates.months[r.lang||"en"];for(n in t.ignoreCase&&(e=e.toLowerCase()),s)if("string"==typeof n&&(a=s[n],t.ignoreCase&&(a=a.toLowerCase()),e.match(a)))return parseInt(n,10);return e},
/*! date-range "dd MMM yyyy - dd MMM yyyy" */
f.addParser({id:"date-range-dMMMyyyy",is:function(){return!1},format:function(e,t,r,a){var n,s,i,d,o=[],c=e.replace(/\s+/g," ").match(l),g=c&&c.length;if(g){for(d=0;d<g;d++)n="",(i=c[d].match(p))&&4<=i.length&&(i.shift(),s=u(i[1],t.config,a),isNaN(s)||(c[d]=c[d].replace(i[1],s)),n=new Date((""+c[d]).replace(f.regex.shortDateXXY,"$3/$2/$1"))),o.push(n instanceof Date&&isFinite(n)?n.getTime():c[d]);return o.sort().join(" - ")}return e},type:"text"})}(jQuery);return jQuery;}));
```
