## Supplementary Data SD1 for "Inferring binding specificities of human transcription factors with the wisdom of crowds": parser-date-two-digit-year.min.js

```
(function(factory){if (typeof define === 'function' && define.amd){define(['jquery'], factory);} else if (typeof module === 'object' && typeof module.exports === 'object'){module.exports = factory(require('jquery'));} else {factory(jQuery);}}(function(jQuery){
/*! Parser: two digit year - updated 11/26/2016 (v2.28.0) */
!function(e){"use strict";var r=e.tablesorter,s=(new Date).getFullYear();r.defaults.dataRange="",r.dates||(r.dates={}),r.dates.regxxxxyy=/(\d{1,2})[\/\s](\d{1,2})[\/\s](\d{2})/,r.dates.regyyxxxx=/(\d{2})[\/\s](\d{1,2})[\/\s](\d{1,2})/,r.formatDate=function(e,t,r,a){if(e){var n,d,t=e.replace(/\s+/g," ").replace(/[-.,]/g,"/").replace(t,r),r=new Date(t);if(r instanceof Date&&isFinite(r)){for(n=r.getFullYear(),d=a&&a.config.dateRange||50;d<s-n;)n+=100;return r.setFullYear(n)}}return e},e.tablesorter.addParser({id:"ddmmyy",is:function(){return!1},format:function(e,t){return r.formatDate(e,r.dates.regxxxxyy,"$2/$1/19$3",t)},type:"numeric"}),e.tablesorter.addParser({id:"mmddyy",is:function(){return!1},format:function(e,t){return r.formatDate(e,r.dates.regxxxxyy,"$1/$2/19$3",t)},type:"numeric"}),e.tablesorter.addParser({id:"yymmdd",is:function(){return!1},format:function(e,t){return r.formatDate(e,r.dates.regyyxxxx,"$2/$3/19$1",t)},type:"numeric"})}(jQuery);return jQuery;}));
```
