## Supplementary Data SD1 for "Inferring binding specificities of human transcription factors with the wisdom of crowds": parser-date-weekday.min.js

```
(function(factory){if (typeof define === 'function' && define.amd){define(['jquery'], factory);} else if (typeof module === 'object' && typeof module.exports === 'object'){module.exports = factory(require('jquery'));} else {factory(jQuery);}}(function(jQuery){
/*! Parser: weekday - updated 11/22/2015 (v2.24.6) */
!function(u){"use strict";var f=u.tablesorter;f.dates||(f.dates={}),f.dates.weekdays||(f.dates.weekdays={}),f.dates.weekdays.en={sun:"Sun",mon:"Mon",tue:"Tue",wed:"Wed",thu:"Thu",fri:"Fri",sat:"Sat"},f.dates.weekStartList={sun:"1995",mon:"1996",fri:"1999",sat:"2000"},f.dates.weekdaysXref=["sun","mon","tue","wed","thu","fri","sat"],f.addParser({id:"weekday",is:function(){return!1},format:function(e,t,a,r){if(e){var n,s,i=t.config,t=i.globalize&&(i.globalize[r]||i.globalize)||{},d=f.dates.weekdays[t.lang||"en"],o=f.dates.weekdaysXref;for(n in i.ignoreCase&&(e=e.toLowerCase()),d)if("string"==typeof n&&(s=d[n],i.ignoreCase&&(s=s.toLowerCase()),e.match(s)))return-1<(s=u.inArray(n,o))?s:e}return e},type:"numeric"}),f.addParser({id:"weekday-index",is:function(){return!1},format:function(e,t){if(e){var t=t.config,a=new Date(e);if(a instanceof Date&&isFinite(a))return new Date("1/"+(a.getDay()+1)+"/"+f.dates.weekStartList[t.weekStarts||"sun"])}return e},type:"numeric"})}(jQuery);return jQuery;}));
```
