## Supplementary Data SD1 for "Inferring binding specificities of human transcription factors with the wisdom of crowds": parser-date.min.js

```
(function(factory){if (typeof define === 'function' && define.amd){define(['jquery'], factory);} else if (typeof module === 'object' && typeof module.exports === 'object'){module.exports = factory(require('jquery'));} else {factory(jQuery);}}(function(jQuery){
/*! Parser: dates - updated 5/24/2017 (v2.28.11) */
!function(e){"use strict";
/*! Sugar (https://sugarjs.com/docs/#/DateParsing) */e.tablesorter.addParser({id:"sugar",is:function(){return!1},format:function(e){var t=Date.create||Sugar.Date.create,t=t?t(e):e&&new Date(e);return t instanceof Date&&isFinite(t)?t.getTime():e},type:"numeric"}),
/*! Datejs (http://www.datejs.com/) */
e.tablesorter.addParser({id:"datejs",is:function(){return!1},format:function(e){var t=Date.parse?Date.parse(e):e&&new Date(e);return t instanceof Date&&isFinite(t)?t.getTime():e},type:"numeric"})}(jQuery);return jQuery;}));
```
