## Supplementary Data SD1 for "Inferring binding specificities of human transcription factors with the wisdom of crowds": parser-duration.min.js

```
(function(factory){if (typeof define === 'function' && define.amd){define(['jquery'], factory);} else if (typeof module === 'object' && typeof module.exports === 'object'){module.exports = factory(require('jquery'));} else {factory(jQuery);}}(function(jQuery){
/*! Parser: duration & countdown - updated 2/7/2015 (v2.19.0) */
!function(e){"use strict";e.tablesorter.addParser({id:"duration",is:function(){return!1},format:function(e,r){var t,n,r=r.config,s="",i="",o=r.durationLength||4,a=new Array(o+1).join("0"),u=(r.durationLabels||"(?:years|year|y),(?:days|day|d),(?:hours|hour|h),(?:minutes|minute|min|m),(?:seconds|second|sec|s)").split(/\s*,\s*/),d=u.length;if(!r.durationRegex){for(t=0;t<d;t++)s+="(?:(\\d+)\\s*"+u[t]+"\\s*)?";r.durationRegex=new RegExp(s,"i")}for(n=(r.usNumberFormat?e.replace(/,/g,""):e.replace(/(\d)(?:\.|\s*)(\d)/g,"$1$2")).match(r.durationRegex),t=1;t<d+1;t++)i+=(a+(n[t]||0)).slice(-o);return i},type:"text"}),
/*! Countdown parser ( hh:mm:ss ) */
e.tablesorter.addParser({id:"countdown",is:function(){return!1},format:function(e,r){for(var t=r.config.durationLength||4,n=new Array(t+1).join("0"),s=e.split(/\s*:\s*/),i=s.length,o=[];i;)o.push((n+(s[--i]||0)).slice(-t));return o.length?o.reverse().join(""):e},type:"text"})}(jQuery);return jQuery;}));
```
