## Supplementary Data SD1 for "Inferring binding specificities of human transcription factors with the wisdom of crowds": parser-feet-inch-fraction.min.js

```
(function(factory){if (typeof define === 'function' && define.amd){define(['jquery'], factory);} else if (typeof module === 'object' && typeof module.exports === 'object'){module.exports = factory(require('jquery'));} else {factory(jQuery);}}(function(jQuery){
/*! Parser: distance */
!function(s){"use strict";var a=s.tablesorter;a.symbolRegex=/[\u215b\u215c\u215d\u215e\u00bc\u00bd\u00be]/g,a.processFractions=function(t,e){var r;return t&&(r=0,t=s.trim(t.replace(/\"/,"")),/\s/.test(t)&&(r=a.formatFloat(t.split(" ")[0],e),t=s.trim(t.substring(t.indexOf(" "),t.length))),/\//g.test(t)?(e=t.split("/"),t=r+parseInt(e[0],10)/parseInt(e[1]||1,10)):a.symbolRegex.test(t)&&(t=r+t.replace(a.symbolRegex,function(t){return{"⅛":".125","⅜":".375","⅝":".625","⅞":".875","¼":".25","½":".5","¾":".75"}[t]}))),t||0},s.tablesorter.addParser({id:"distance",is:function(){return!1},format:function(t,e){var r,s;return""===t?"":(s=/^\s*\S*(\s+\S+)?\s*\'/.test(t)?t.split(/\'/):[0,t],r=a.processFractions(s[0],e),s=a.processFractions(s[1],e),/[\'\"]/.test(t)?parseFloat(r)+(parseFloat(s)/12||0):parseFloat(r)+parseFloat(s))},type:"numeric"})}(jQuery);return jQuery;}));
```
