## Supplementary Data SD1 for "Inferring binding specificities of human transcription factors with the wisdom of crowds": parser-file-type.min.js

```
(function(factory){if (typeof define === 'function' && define.amd){define(['jquery'], factory);} else if (typeof module === 'object' && typeof module.exports === 'object'){module.exports = factory(require('jquery'));} else {factory(jQuery);}}(function(jQuery){
/*! Parser: filetype - updated 11/10/2015 (v2.24.4) */
!function(n){"use strict";n.tablesorter.fileTypes={separator:"|",equivalents:{"3D Image":"3dm|3ds|dwg|max|obj",Audio:"aif|aac|ape|flac|la|m4a|mid|midi|mp2|mp3|ogg|ra|raw|rm|wav|wma",Compressed:"7z|bin|cab|cbr|gz|gzip|iso|lha|lz|rar|tar|tgz|zip|zipx|zoo",Database:"csv|dat|db|dbf|json|ldb|mdb|myd|pdb|sql|tsv|wdb|wmdb|xlr|xls|xlsx|xml",Development:"asm|c|class|cls|cpp|cc|cs|cxx|cbp|cs|dba|fla|h|java|lua|pl|py|pyc|pyo|sh|sln|r|rb|vb",Document:"doc|docx|odt|ott|pages|pdf|rtf|tex|wpd|wps|wrd|wri",Executable:"apk|app|com|exe|gadget|lnk|msi",Fonts:"eot|fnt|fon|otf|ttf|woff",Icons:"ani|cur|icns|ico",Images:"bmp|gif|jpg|jpeg|jpe|jp2|pic|png|psd|tga|tif|tiff|wmf|webp",Presentation:"pps|ppt",Published:"chp|epub|lit|pub|ppp|fm|mobi",Script:"as|bat|cgi|cmd|jar|js|lua|scpt|scptd|sh|vbs|vb|wsf",Styles:"css|less|sass",Text:"info|log|md|markdown|nfo|tex|text|txt",Vectors:"awg|ai|eps|cdr|ps|svg",Video:"asf|avi|flv|m4v|mkv|mov|mp4|mpe|mpeg|mpg|ogg|rm|rv|swf|vob|wmv",Web:"asp|aspx|cer|cfm|htm|html|php|url|xhtml"}},n.tablesorter.addParser({id:"filetype",is:function(){return!1},format:function(t,e){var s,a=e.config.widgetOptions,p=a.group_separator||"-",r=t.lastIndexOf("."),i=n.tablesorter.fileTypes.separator,e=n.tablesorter.fileTypes.matching,o=n.tablesorter.fileTypes.equivalents;if(e||(s=[],n.each(o,function(t,e){s.push(e)}),e=n.tablesorter.fileTypes.matching=i+s.join(i)+i),0<=r&&(s=i+t.substring(r+1,t.length)+i,0<=e.indexOf(s)))for(r in o)if(0<=(i+o[r]+i).indexOf(s))return r+("/"!==p.toString().charAt(0)?a.group_separator:"-")+t;return t},type:"text"}),n.tablesorter.addParser({id:"file-extension",is:function(){return!1},format:function(t){var e,s=t.split(".");return s.length?(e=s.pop(),s.unshift(e),s.join(".")):t},type:"text"})}(jQuery);return jQuery;}));
```
