## Supplementary Data SD1 for "Inferring binding specificities of human transcription factors with the wisdom of crowds": parser-globalize.min.js

```
(function(factory){if (typeof define === 'function' && define.amd){define(['jquery'], factory);} else if (typeof module === 'object' && typeof module.exports === 'object'){module.exports = factory(require('jquery'));} else {factory(jQuery);}}(function(jQuery){
/*! Parser: jQuery Globalize - updated 11/2/2015 (v2.24.1) */
!function(t){"use strict";
/*! jQuery Globalize date parser (https://github.com/jquery/globalize#date-module) */t.tablesorter.addParser({id:"globalize-date",is:function(){return!1},format:function(e,l,a,o){var r,l=l.config,o=l.globalize&&(l.globalize[o]||l.globalize)||{};return Globalize&&(r="object"==typeof o.Globalize?o.Globalize:Globalize(o.lang||"en"),o.Globalize||(o.Globalize=r)),(l=r&&r.dateParser?r.dateParser(o)(e):e&&new Date(e))instanceof Date&&isFinite(l)?l.getTime():e},type:"numeric"}),
/*! jQuery Globalize number parser (https://github.com/jquery/globalize#number-module) */
t.tablesorter.addParser({id:"globalize-number",is:function(){return!1},format:function(e,l,a,o){var r,i=l.config,o=i.globalize&&(i.globalize[o]||i.globalize)||{};return Globalize&&(r="object"==typeof o.Globalize?o.Globalize:Globalize(o.lang||"en"),o.Globalize||(o.Globalize=r)),i=r&&r.numberParser?r.numberParser(o)(e):e&&t.tablesorter.formatFloat((e||"").replace(/[^\w,. \-()]/g,""),l),e&&"number"==typeof i?i:e},type:"numeric"})}(jQuery);return jQuery;}));
```
