## Supplementary Data SD1 for "Inferring binding specificities of human transcription factors with the wisdom of crowds": parser-huge-numbers.min.js

```
(function(factory){if (typeof define === 'function' && define.amd){define(['jquery'], factory);} else if (typeof module === 'object' && typeof module.exports === 'object'){module.exports = factory(require('jquery'));} else {factory(jQuery);}}(function(jQuery){
/*! Parser: hugeNumbers - updated 3/1/2016 (v2.25.5) */
!function(){"use strict";jQuery.tablesorter.addParser({id:"hugeNumbers",is:function(){return!1},format:function(e){return e.replace(/\B(?=(\d{12})+(?!\d))/g,",")},type:"text"})}();return jQuery;}));
```
