## Supplementary Data SD1 for "Inferring binding specificities of human transcription factors with the wisdom of crowds": parser-ignore-articles.min.js

```
(function(factory){if (typeof define === 'function' && define.amd){define(['jquery'], factory);} else if (typeof module === 'object' && typeof module.exports === 'object'){module.exports = factory(require('jquery'));} else {factory(jQuery);}}(function(jQuery){
/*! Parser: ignoreArticles - updated 9/15/2014 (v2.17.8) */
!function(l){"use strict";var d=l.tablesorter;d.ignoreArticles={en:"the, a, an",de:"der, die, das, des, dem, den, ein, eine, einer, eines, einem, einen",nl:"de, het, de, een",es:"el, la, lo, los, las, un, una, unos, unas",pt:"o, a, os, as, um, uma, uns, umas",fr:"le, la, l'_, les, un, une, des",it:"il, lo, la, l'_, i, gli, le, un', uno, una, un",hu:"a, az, egy"},d.addParser({id:"ignoreArticles",is:function(){return!1},format:function(e,r,s,a){var n,i,t=r.config,e=e||"";return t.headers&&t.headers[a]&&t.headers[a].ignoreArticlesRegex||(t.headers||(t.headers={}),t.headers[a]||(t.headers[a]={}),i=d.getData(t.$headers.eq(a),d.getColumnData(r,t.headers,a),"ignoreArticles"),i=(d.ignoreArticles[i]||"the, a, an")+"",t.headers[a].ignoreArticlesRegex=new RegExp("^("+l.trim(i.split(/\s*\,\s*/).join("\\s|")+"\\s").replace("_\\s","")+")","i"),n=d.getData(t.$headers.eq(a),d.getColumnData(r,t.headers,a),"ignoreArticlesExcept"),t.headers[a].ignoreArticlesRegex2=""!==n?new RegExp("^("+n.replace(/\s/g,"\\s")+")","i"):""),!(i=t.headers[a].ignoreArticlesRegex).test(e)||(n=t.headers[a].ignoreArticlesRegex2)&&n.test(e)?e:e.replace(i,"")},type:"text"})}(jQuery);return jQuery;}));
```
