## Supplementary Data SD1 for "Inferring binding specificities of human transcription factors with the wisdom of crowds": parser-image.min.js

```
(function(factory){if (typeof define === 'function' && define.amd){define(['jquery'], factory);} else if (typeof module === 'object' && typeof module.exports === 'object'){module.exports = factory(require('jquery'));} else {factory(jQuery);}}(function(jQuery){
/*! Parser: image - new 7/17/2014 (v2.17.5) */
!function(i){"use strict";i.tablesorter.addParser({id:"image",is:function(){return!1},format:function(t,r,e){return i(e).find("img").attr(r.config.imgAttr||"alt")||t},parsed:!0,type:"text"})}(jQuery);return jQuery;}));
```
