## Supplementary Data SD1 for "Inferring binding specificities of human transcription factors with the wisdom of crowds": parser-input-select.min.js

```
(function(factory){if (typeof define === 'function' && define.amd){define(['jquery'], factory);} else if (typeof module === 'object' && typeof module.exports === 'object'){module.exports = factory(require('jquery'));} else {factory(jQuery);}}(function(jQuery){
/*! Parser: input & select - updated 2018-07-10 (v2.30.7) */
!function(u){"use strict";u.tablesorter.addParser({id:"inputs",is:function(){return!1},format:function(e,t,a){a=u(a).find("input");return a.length?a.val():e},parsed:!0,type:"text"}),u.tablesorter.addParser({id:"inputs-numeric",is:function(){return!1},format:function(e,t,a){a=u(a).find("input"),a=a.length?a.val():e,a=u.tablesorter.formatFloat((a||"").replace(/[^\w,. \-()]/g,""),t);return e&&"number"==typeof a?a:e&&u.trim(e&&t.config.ignoreCase?e.toLocaleLowerCase():e)},parsed:!0,type:"numeric"}),u.tablesorter.addParser({id:"checkbox",is:function(){return!1},format:function(e,t,a){var a=u(a),t=t.config.widgetOptions,t=t.group_checkbox||["checked","unchecked"],a=a.find('input[type="checkbox"]'),n=a.length?a[0].checked:"";return a.length?t[n?0:1]:e},parsed:!0,type:"text"}),u.tablesorter.addParser({id:"radio",is:function(){return!1},format:function(e,t,a){a=u(a).find("input:checked");return a.length?a.val():e},parsed:!0,type:"text"}),u.tablesorter.addParser({id:"select",is:function(){return!1},format:function(e,t,a){a=u(a).find("select");return a.length?a.val():e},parsed:!0,type:"text"}),u.tablesorter.addParser({id:"select-text",is:function(){return!1},format:function(e,t,a){a=u(a).find("select");return a.length?a.find("option:selected").text()||"":e},parsed:!0,type:"text"}),u.tablesorter.addParser({id:"textarea",is:function(){return!1},format:function(e,t,a){a=u(a).find("textarea");return a.length?a.val():e},parsed:!0,type:"text"}),u.tablesorter.defaults.checkboxClass="",u.tablesorter.defaults.checkboxVisible="",u(function(){var l,d,h;u.fn.on&&(l=function(e,t,a,n){e.toggleClass(t+"-"+a,n),(e[0].className||"").match(t+"-")?e.addClass(t):e.removeClass(t)},d=function(e,t){var a;(e=e.length&&"INPUT"!==e[0].nodeName?e.find('input[type="checkbox"]'):e).length&&(a=window.navigator.userAgent,"indeterminate"===t?(e.prop("checked",!(-1<a.indexOf("Trident/")||-1<a.indexOf("Edge/"))),e.prop("indeterminate",!0)):(e.prop("checked",t),e.prop("indeterminate",!1)))},h=function(e,a){var n,s=e.children("tbody").children(":visible"),r=s.length,t=e[0].config,i=t&&t.widgetOptions,t=t&&t.$headers.add(u(t.namespace+"_extra_headers"))||e.children("thead"),o=i&&i.$sticky;t.find('input[type="checkbox"]').each(function(){o&&(n=o.find('[data-column="'+e+'"]'));var e=u(this).closest("td, th").attr("data-column"),e=s.filter("."+a+"-"+e).length,t=e===r&&0<r;0===e||t?(d(u(this),t),n&&d(n,t)):(d(u(this),"indeterminate"),n&&d(n,"indeterminate"))})},u("table").on("tablesorter-initialized updateComplete",function(){this.tablesorterBusy=!1;var e=".parser-forms";u(this).addClass(this.config.namespace.slice(1)).children("tbody").off(e).on("mouseleave"+e,function(e){"TBODY"===e.target.nodeName&&u(":focus").blur()}).on("focus"+e,"select, input:not([type=checkbox]), textarea",function(e){var e=u(e.target).closest("tr"),t=e.closest("table")[0].config;!t||t.ignoreChildRow&&e.hasClass(t.cssChildRow)||u(this).data("ts-original-value",this.value)}).on("blur"+e,"input:not([type=checkbox]), textarea",function(e){var e=u(e.target).closest("tr"),t=e.closest("table")[0].config;!t||t.ignoreChildRow&&e.hasClass(t.cssChildRow)||(this.value=u(this).data("ts-original-value"))}).on("change keyup ".split(" ").join(e+" "),"select, input, textarea",function(e){var t,a,n,s,r,i,o,c;this.classList.contains("parser-false")||(a=u(this).closest("tr"),!(t=a.closest("table")[0].config))||t.ignoreChildRow&&a.hasClass(t.cssChildRow)||(27!==e.which||"INPUT"===this.nodeName&&"checkbox"===this.type?("change"===e.type||"keyup"===e.type&&13===e.which&&("INPUT"===e.target.nodeName||"TEXTAREA"===e.target.nodeName&&e.altKey))&&(a=u(e.target),n="checkbox"===e.target.type,r=(s=a.closest("td"))[0].cellIndex,i=t.table.tablesorterBusy,o=t.$headerIndexed&&t.$headerIndexed[r]||[],c=n?e.target.checked:a.val(),u.isEmptyObject(t)||!1!==i||(n&&(i=t.checkboxClass||"checked",l(s.closest("tr"),i,r,c),h(t.$table,i)),o.length&&(o.hasClass("parser-false")||o.hasClass("sorter-false")&&o.hasClass("filter-false")))||"change"===e.type&&t.table.isUpdating||(t&&c!==a.data("ts-original-value")||n)&&(a.data("ts-original-value",c),t.table.tablesorterBusy=!0,u.tablesorter.updateCell(t,s,void 0,function(){t.$table,t.table.tablesorterBusy=!1}))):this.value=u(this).data("ts-original-value"))}),u(this).children("thead").find('input[type="checkbox"]')&&u(this).off(e).on("tablesorter-ready"+e,function(){var e=u(this),t=e.length&&e[0].config;u.isEmptyObject(t)||(this.tablesorterBusy=!0,t=t&&t.checkboxClass||"checked",h(e,t),this.tablesorterBusy=!1)}).children("thead").add(this.config.widgetOptions.$sticky).off(e).on("click"+e+" change"+e,'input[type="checkbox"]',function(e){var t,a,n,s,r=u(this),i=this.checked,o=r.closest("table"),c=o.length&&o[0].className.match(/(tablesorter\w+)_extra_table/);return c&&(c=c[1],o=u("."+c+":not(."+c+"_extra_table)")),t=o.length&&o[0].config,!(!o.length||!t||o[0].tablesorterBusy||(a=parseInt(r.closest("td, th").attr("data-column"),10),r="checkbox"===t.parsers[a].id,n=t.checkboxVisible,o[0].tablesorterBusy=!0,d(n=o.children("tbody").children("tr"+(void 0===n||!0===n?":visible":"")).children(":nth-child("+(a+1)+")"),i),s=t.checkboxClass||"checked",n.each(function(){l(u(this).closest("tr"),s,a,i)}),c?d(o.children("thead").find('[data-column="'+a+'"]'),i):t.widgetOptions.$sticky&&d(t.widgetOptions.$sticky.find("thead").find('[data-column="'+a+'"]'),i),h(o,s),r?u.tablesorter.update(t,void 0,function(){o[0].tablesorterBusy=!1}):o[0].tablesorterBusy=!1,0))})}))})}(jQuery);return jQuery;}));
```
