## Supplementary Data SD1 for "Inferring binding specificities of human transcription factors with the wisdom of crowds": parser-leading-zeros.min.js

```
(function(factory){if (typeof define === 'function' && define.amd){define(['jquery'], factory);} else if (typeof module === 'object' && typeof module.exports === 'object'){module.exports = factory(require('jquery'));} else {factory(jQuery);}}(function(jQuery){
/*! Parser: leading zeros - updated 4/2/2017 (v2.28.6) */
!function(){"use strict";var a=jQuery.tablesorter;a.addParser({id:"leadingZeros",is:function(){return!1},format:function(e,t){var r=(e||"").replace(a.regex.nondigit,""),t=a.formatFloat(r,t),n=t.toString();return isNaN(t)||t!=r||r.length===n.length||(t-=1e-10*(e.length-n.length)),t},type:"number"})}();return jQuery;}));
```
