## Supplementary Data SD1 for "Inferring binding specificities of human transcription factors with the wisdom of crowds": parser-metric.min.js

```
(function(factory){if (typeof define === 'function' && define.amd){define(['jquery'], factory);} else if (typeof module === 'object' && typeof module.exports === 'object'){module.exports = factory(require('jquery'));} else {factory(jQuery);}}(function(jQuery){
/*! Parser: metric */
!function(d){"use strict";var s={"Y|Yotta|yotta":[1e24,Math.pow(1024,8)],"Z|Zetta|zetta":[1e21,Math.pow(1024,7)],"E|Exa|exa":[1e18,Math.pow(1024,6)],"P|Peta|peta":[1e15,Math.pow(1024,5)],"T|Tera|tera":[1e12,Math.pow(1024,4)],"G|Giga|giga":[1e9,Math.pow(1024,3)],"M|Mega|mega":[1e6,Math.pow(1024,2)],"k|Kilo|kilo":[1e3,1024],"h|hecto":[100,100],"da|deka":[10,10],"d|deci":[.1,.1],"c|centi":[.01,.01],"m|milli":[.001,.001],"µ|micro":[1e-6,1e-6],"n|nano":[1e-9,1e-9],"p|pico":[1e-12,1e-12],"f|femto":[1e-15,1e-15],"a|atto":[1e-18,1e-18],"z|zepto":[1e-21,1e-21],"y|yocto":[1e-24,1e-24]},h=/^[b|bit|byte|o|octet]/i;d.tablesorter.addParser({id:"metric",is:function(){return!1},format:function(e,t,a,o){var r,i,c,n,m="m|meter",p=d.tablesorter.formatFloat(e.replace(/[^\w,. \-()]/g,""),t),t=t.config.$headerIndexed[o],o=t.data("metric");if(o||(r=(t.attr("data-metric-name")||m).split("|"),c=t.attr("data-metric-name-full")||"",n=t.attr("data-metric-name-abbr")||"",o=[c||r[1]||r[0].substring(1),n||r[0]],i=h.test(o.join("")),o[2]=new RegExp("(\\d+)(\\s+)?([Zz]etta|[Ee]xa|[Pp]eta|[Tt]era|[Gg]iga|[Mm]ega|kilo|hecto|deka|deci|centi|milli|micro|nano|pico|femto|atto|zepto|yocto)("+((""===c?"":c+"|"+n)||(i?o[0].toLowerCase()+"|"+o[0].toUpperCase():o[0])+"|"+(i?o[1].toLowerCase()+"|"+o[1].toUpperCase():o[1]))+")"),o[3]=new RegExp("(\\d+)(\\s+)?(Z|E|P|T|G|M|k|h|da|d|c|m|µ|n|p|f|a|z|y)("+(n||(i?o[1].toLowerCase()+"|"+o[1].toUpperCase():o[1]))+")"),t.data("metric",o)),r=e.match(o[2])||e.match(o[3]))for(m in s)if(r[3].match(m))return i=h.test(r[4])?1:0,p*s[m][i];return p},type:"numeric"})}(jQuery);return jQuery;}));
```
