## Supplementary Data SD1 for "Inferring binding specificities of human transcription factors with the wisdom of crowds": parser-named-numbers.min.js

```
(function(factory){if (typeof define === 'function' && define.amd){define(['jquery'], factory);} else if (typeof module === 'object' && typeof module.exports === 'object'){module.exports = factory(require('jquery'));} else {factory(jQuery);}}(function(jQuery){
/*! Parser: namedNumbers - updated 10/26/2014 (v2.18.0) */
!function(s){"use strict";var a,d,g={negative:["negative","minus"],numbers:{zero:0,one:1,two:2,three:3,four:4,five:5,six:6,seven:7,eight:8,nine:9,ten:10,eleven:11,twelve:12,thirteen:13,fourteen:14,fifteen:15,sixteen:16,seventeen:17,eighteen:18,nineteen:19,twenty:20,thirty:30,forty:40,fourty:40,fifty:50,sixty:60,seventy:70,eighty:80,ninety:90},hundred:"hundred",powers:{thousand:1e3,million:1e6,billion:1e9,trillion:1e12,quadrillion:1e15,quintillion:1e18,sextillion:1e21,septillion:1e24,octillion:1e27,nonillion:1e30,decillion:1e33,undecillion:1e36,duodecillion:1e39,tredecillion:1e42,quattuordecillion:1e45,quindecillion:1e48,sexdecillion:1e51,septendecillion:1e54,octodecillion:1e57,novemdecillion:1e60,vigintillion:1e63,unvigintillion:1e66,duovigintillion:1e69,trevigintillion:1e72,quattuorvigintillion:1e75,quinvigintillion:1e78,sexvigintillion:1e81,septenvigintillion:1e84,octovigintillion:1e87,novemvigintillion:1e90,trigintillion:1e93,untrigintillion:1e96,duotrigintillion:1e99,googl:1e100}},v=new RegExp("("+g.negative.join("|")+")");s.tablesorter.addParser({id:"namedNumbers",is:function(){return!1},format:function(e,i){d=a=0;for(var n,t,l,o=(e||"").split(/[\s-]+/),r=o.length,u=0;u<r;u++)n=o[u].toLowerCase(),t=i,l=void 0,l=n.replace(/[,."']/g,""),n=s.tablesorter.formatFloat(n||"",t),t=g.powers.hasOwnProperty(l)?g.powers[l]:null,null!==(n="number"==typeof n?n:g.numbers.hasOwnProperty(l)?g.numbers[l]:null)?d+=n:l===g.hundred?d*=100:null!==t&&(a+=d*t,d=0);return(a=(a+d)*(e.match(v)?-1:1))||g.numbers.hasOwnProperty(e)?a:s.tablesorter.formatFloat(e||"",i)},type:"numeric"})}(jQuery);return jQuery;}));
```
