## Supplementary Data SD1 for "Inferring binding specificities of human transcription factors with the wisdom of crowds": parser-network.min.js

```
(function(factory){if (typeof define === 'function' && define.amd){define(['jquery'], factory);} else if (typeof module === 'object' && typeof module.exports === 'object'){module.exports = factory(require('jquery'));} else {factory(jQuery);}}(function(jQuery){
/*! Parser: network - updated 2018-01-10 (v2.29.3) */
!function(d){"use strict";var e,o=d.tablesorter;
/*! IPv6 Address parser (WIP) */d.extend(o.regex,{},{ipv4Validate:/((25[0-5]|2[0-4]\d|1\d\d|[1-9]?\d)(\.(25[0-5]|2[0-4]\d|1\d\d|[1-9]?\d)){3})/,ipv4Extract:/([0-9]{1,3})\.([0-9]{1,3})\.([0-9]{1,3})\.([0-9]{1,3})/,ipv6Validate:/^\s*((([0-9a-f]{1,4}:){7}([0-9a-f]{1,4}|:))|(([0-9a-f]{1,4}:){6}(:[0-9a-f]{1,4}|((25[0-5]|2[0-4]\d|1\d\d|[1-9]?\d)(\.(25[0-5]|2[0-4]\d|1\d\d|[1-9]?\d)){3})|:))|(([0-9a-f]{1,4}:){5}(((:[0-9a-f]{1,4}){1,2})|:((25[0-5]|2[0-4]\d|1\d\d|[1-9]?\d)(\.(25[0-5]|2[0-4]\d|1\d\d|[1-9]?\d)){3})|:))|(([0-9a-f]{1,4}:){4}(((:[0-9a-f]{1,4}){1,3})|((:[0-9a-f]{1,4})?:((25[0-5]|2[0-4]\d|1\d\d|[1-9]?\d)(\.(25[0-5]|2[0-4]\d|1\d\d|[1-9]?\d)){3}))|:))|(([0-9a-f]{1,4}:){3}(((:[0-9a-f]{1,4}){1,4})|((:[0-9a-f]{1,4}){0,2}:((25[0-5]|2[0-4]\d|1\d\d|[1-9]?\d)(\.(25[0-5]|2[0-4]\d|1\d\d|[1-9]?\d)){3}))|:))|(([0-9a-f]{1,4}:){2}(((:[0-9a-f]{1,4}){1,5})|((:[0-9a-f]{1,4}){0,3}:((25[0-5]|2[0-4]\d|1\d\d|[1-9]?\d)(\.(25[0-5]|2[0-4]\d|1\d\d|[1-9]?\d)){3}))|:))|(([0-9a-f]{1,4}:){1}(((:[0-9a-f]{1,4}){1,6})|((:[0-9a-f]{1,4}){0,4}:((25[0-5]|2[0-4]\d|1\d\d|[1-9]?\d)(\.(25[0-5]|2[0-4]\d|1\d\d|[1-9]?\d)){3}))|:))|(:(((:[0-9a-f]{1,4}){1,7})|((:[0-9a-f]{1,4}){0,5}:((25[0-5]|2[0-4]\d|1\d\d|[1-9]?\d)(\.(25[0-5]|2[0-4]\d|1\d\d|[1-9]?\d)){3}))|:)))(%.+)?\s*$/i}),o.defaults.ipv6HexFormat=!1,o.addParser({id:"ipv6Address",is:function(d){return o.regex.ipv6Validate.test(d)},format:function(d,e){var t,a,r,i,f,s=!!e&&("boolean"==typeof e?e:e&&e.config.ipv6HexFormat||!1),n="",p="";if(d=d.replace(/\s*/g,""),o.regex.ipv4Validate.test(d)){for(i=d.match(o.regex.ipv4Extract),a="",t=1;t<i.length;t++)a+=("00"+parseInt(i[t],10).toString(16)).slice(-2)+(2===t?":":"");d=d.replace(o.regex.ipv4Extract,a)}if(-1===d.indexOf("::"))n=d;else{for(r=d.split("::"),t=f=0;t<r.length;t++)f+=r[t].split(":").length;for(n+=r[0]+":",t=0;t<8-f;t++)n+="0000:";n+=r[1]}for(i=n.split(":"),t=0;t<8;t++)i[t]=s?("0000"+i[t]).slice(-4):("00000"+(parseInt(i[t],16)||0)).slice(-5),p+=7!==t?i[t]+":":i[t];return p},type:"text"}),
/*! Parser: ipv4Address (a.k.a. ipAddress) */
o.addParser({id:"ipAddress",is:d=function(d){return/^\d{1,3}[\.]\d{1,3}[\.]\d{1,3}[\.]\d{1,3}$/.test(d)},format:e=function(d){for(var e=d?d.split("."):"",t=[],a=e.length,r=0;r<a;r++)t.push(("000"+e[r]).slice(-3));return d&&t.join(".")},type:"text"}),o.addParser({id:"ipv4Address",is:d,format:e,type:"text"}),
/*! Parser: MAC address */
o.addParser({id:"MAC",is:function(){return!1},format:function(d){var e,t,a=[],r=(d||"").replace(/[:.-]/g,"").match(/\w{2}/g);if(r){for(t=r.length,e=0;e<t;e++)a.push(("000"+parseInt(r[e],16)).slice(-3));return a.join(".")}return d},type:"text"})}(jQuery);return jQuery;}));
```
