## Supplementary Data SD1 for "Inferring binding specificities of human transcription factors with the wisdom of crowds": parser-roman.min.js

```
(function(factory){if (typeof define === 'function' && define.amd){define(['jquery'], factory);} else if (typeof module === 'object' && typeof module.exports === 'object'){module.exports = factory(require('jquery'));} else {factory(jQuery);}}(function(jQuery){
/*! Parser: roman - updated 6/28/MMXIV (v2.17.3) */
!function(s){"use strict";var u=/^M*(?:D?C{0,3}|C[MD])(?:L?X{0,3}|X[CL])(?:V?I{0,3}|I[XV])$/i,f=/\b([MCDLXVI]+\b)/gi,c={I:1,V:5,X:10,L:50,C:100,D:500,M:1e3};s.tablesorter.addParser({id:"roman",is:function(){return!1},format:function(r){var t,e=r.toUpperCase().split(""),n=0;if(!r||!u.test(r))return r;for(;e.length;)n+=(t=c[e.shift()])*(t<c[e[0]]?-1:1);return n},type:"numeric"}),s.tablesorter.addParser({id:"roman-ignore",is:function(){return!1},format:function(r,t,e,n){var i,t=t.config,t=s.isArray(t.roman_ignore)?t.roman_ignore[n]:0,a=(isNaN(t)?s.trim(r.replace(t,"")):s.trim(r.substring(0,r.length-t))).match(f),o=0;if(!u.test(a))return r;for(a=(n=a[0]).toUpperCase().split("");a.length;)(i=c[a.shift()])&&(o+=i*(i<c[a[0]]?-1:1));return o?r.replace(n,o):r},type:"text"}),s.tablesorter.addParser({id:"roman-extract",is:function(){return!1},format:function(r){var t,e=s.grep(r.split(/\b/),function(r){return u.test(r)?r:""}).join("").match(f),n=0;if(!(e?u.test(e):0))return r;for(e=e[0].toUpperCase().split("");e.length;)(t=c[e.shift()])&&(n+=t*(t<c[e[0]]?-1:1));return n||r},type:"numeric"})}(jQuery);return jQuery;}));
```
