## Supplementary Data SD1 for "Inferring binding specificities of human transcription factors with the wisdom of crowds": widget-alignChar.min.js

```
(function(factory){if (typeof define === 'function' && define.amd){define(['jquery'], factory);} else if (typeof module === 'object' && typeof module.exports === 'object'){module.exports = factory(require('jquery'));} else {factory(jQuery);}}(function(jQuery){
/*! Widget: alignChar - updated 2/7/2015 (v2.19.0) */
!function(_){"use strict";var r=_.tablesorter;r.alignChar={init:function(a,i,t){i.$headers.filter("["+t.alignChar_charAttrib+"]").each(function(){var n=_(this),n={column:this.column,align:n.attr(t.alignChar_charAttrib),alignIndex:parseInt(n.attr(t.alignChar_indexAttrib)||0,10),adjust:parseFloat(n.attr(t.alignChar_adjustAttrib))||0};n.regex=new RegExp("\\"+n.align,"g"),void 0!==n.align&&(t.alignChar_savedVars[this.column]=n,r.alignChar.setup(a,i,t,n))})},setup:function(n,a,i,t){if(!_.isEmptyObject(a.cache)){for(var r,e,l,h,g,o,s,d,c,u,f,m,p,C=[],w=[],b=0;b<a.$tbodies.length;b++)for(c=(o=a.cache[b]).normalized.length,r=0;r<c;r++){if(0<(d=((s=(p=o.row?o.row[r]:o.normalized[r][a.columns].$row).find("td").eq(t.column).text().replace(/[ ]/g," ")).match(t.regex)||[]).length)&&0<t.alignIndex)for(l=Math.min(t.alignIndex,d),h=g=e=0;e++<l;)g=(h=s.indexOf(t.align,h+1))<0?g:h;else g=s.indexOf(t.align);0<=g?(C.push(s.substring(0,g)||""),w.push(s.substring(g,s.length)||"")):(C.push(!(1<=d&&t.alignIndex>=d)&&s||""),w.push(1<=d&&t.alignIndex>=d&&s||""))}for(u=_.extend([],C).sort(function(n,a){return a.length-n.length})[0],f=_.extend([],w).sort(function(n,a){return a.length-n.length})[0],t.width=t.width||Math.floor(u.length/(u.length+f.length)*100)+t.adjust,u="min-width:"+t.width+"%",f="min-width:"+(100-t.width)+"%",b=0;b<a.$tbodies.length;b++)for(c=(o=a.cache[b]).normalized.length,r=0;r<c;r++)m=_(i.alignChar_wrap).length?_(i.alignChar_wrap).html(t.align)[0].outerHTML:t.align,p=o.row?o.row[r]:o.normalized[r][a.columns].$row,h=w[r].slice(t.align.length),p.find("td").eq(t.column).html('<span class="ts-align-wrap"><span class="ts-align-left" style="'+u+'">'+C[r]+'</span><span class="ts-align-right" style="'+f+'">'+(h.length?m+h:"")+"</span></span>");i.alignChar_initialized=!0}},remove:function(n,a,i){if(!_.isEmptyObject(a.cache))for(var t,r,e,l,h=0;h<a.$tbodies.length;h++)for(r=(e=a.cache[h]).normalized.length,t=0;t<r;t++)(l=(e.row?e.row[t]:e.normalized[t][a.columns].$row).find("td").eq(i)).html(l.text().replace(/\s/g," "))}},r.addWidget({id:"alignChar",priority:100,options:{alignChar_wrap:"",alignChar_charAttrib:"data-align-char",alignChar_indexAttrib:"data-align-index",alignChar_adjustAttrib:"data-align-adjust"},init:function(n,a,i,t){t.alignChar_initialized=!1,t.alignChar_savedVars=[],r.alignChar.init(n,i,t),i.$table.on("pagerEnd refreshAlign",function(){i.$headers.filter("["+t.alignChar_charAttrib+"]").each(function(){r.alignChar.remove(n,i,this.column)}),r.alignChar.init(n,i,t)})},format:function(n,a,i){i.alignChar_initialized||a.$table.triggerHandler("refreshAlign")},remove:function(n,a,i,t){t||(a.$headers.filter("["+i.alignChar_charAttrib+"]").each(function(){r.alignChar.remove(n,a,this.column)}),i.alignChar_initialized=!1)}})}(jQuery);return jQuery;}));
```
