## Supplementary Data SD1 for "Inferring binding specificities of human transcription factors with the wisdom of crowds": widget-build-table.min.js

```
(function(factory){if (typeof define === 'function' && define.amd){define(['jquery'], factory);} else if (typeof module === 'object' && typeof module.exports === 'object'){module.exports = factory(require('jquery'));} else {factory(jQuery);}}(function(jQuery){
/*! Widget: Build Table - updated 2018-03-26 (v2.30.2) */
!function(f){"use strict";var h=f.tablesorter=f.tablesorter||{},g=h.buildTable=function(e,o){f.extend(!0,h.defaults.widgetOptions,g.defaults);function t(e){var t=f.type(e),l=e instanceof f;if("function"==typeof i&&(e=i(e,n)),o.data=e,l||"string"===t){if(l||/<\s*\/tr\s*>/.test(e))return g.html(r,e,n);try{if(e=f.parseJSON(e||"null"))return g.object(r,e,n)}catch(e){}}"array"===t||"string"===t||"array"===d||"csv"===d?g.csv(r,e,n):g.object(r,e,n)}var l="TABLE"===e.nodeName?f(e):f("<table>").appendTo(e),r=l[0],n=o.widgetOptions=f.extend(!0,{},g.defaults,o.widgetOptions),i=n.build_processing,d=n.build_type,e=n.build_source||o.data,s=h.debug(o,"build");if(r.config=o,!h.buildTable.hasOwnProperty(d)&&""!==d)return s&&console.error("Build >> ERROR: Aborting build table widget, incorrect build type"),!1;e instanceof f?t(f.trim(e.html())):e&&(e.hasOwnProperty("url")||"json"===d)?f.ajax(n.build_source).done(function(e){t(e)}).fail(function(e,t){s&&console.error("Build >> ERROR: Aborting build table widget, failed ajax load"),l.html('<tr><td class="error">'+e.status+" "+t+"</td></tr>")}):t(e)};h.defaults.data="",g.defaults={build_type:"",build_source:"",build_processing:null,build_complete:"tablesorter-build-complete",build_headers:{rows:1,classes:[],text:[],widths:[]},build_footers:{rows:1,classes:[],text:[]},build_numbers:{addColumn:!1,sortable:!1},build_csvStartLine:0,build_csvSeparator:",",build_objectRowKey:"rows",build_objectCellKey:"cells",build_objectHeaderKey:"headers",build_objectFooterKey:"footers"},g.build={colgroup:function(e){var l="";return e&&e.length&&(l+="<colgroup>",f.each(e,function(e,t){l+="<col"+(t?' style="width:'+t+'"':"")+">"}),l+="</colgroup>"),l},cell:function(e,t,l,o,r){var n,i,d=r?f("<col>"):"",s=t.build_headers.classes,t=t.build_headers.widths;if(/string|number/.test(typeof e))i=f("<"+l+(s&&s[o]?' class="'+s[o]+'"':"")+">"+e+"</"+l+">"),r&&t&&t[o]&&d.width(t[o]||"");else for(n in i=f("<"+l+">"),e)e.hasOwnProperty(n)&&("text"===n||"html"===n?i[n](e[n]):r&&"width"===n?d.width(e[n]||""):i.attr(n,e[n]));return[i,d]},header:function(e,t){var l=t.build_headers.text,o=t.build_headers.classes,r="<tr>"+(t.build_numbers.addColumn?"<th"+(t.build_numbers.sortable?"":' class="sorter-false"')+">"+t.build_numbers.addColumn+"</th>":"");return f.each(e,function(e,t){/<\s*\/t(d|h)\s*>/.test(t)?r+=t:r+="<th"+(o&&o[e]?' class="'+o[e]+'"':"")+">"+(l&&l[e]?l[e]:t)+"</th>"}),r+"</tr>"},rows:function(e,l,o,t,r,n){var i=n?"th":"td",d="<tr>"+(t.build_numbers.addColumn?"<"+i+">"+(n?"":r)+"</"+i+">":"");return f.each(e,function(e,t){/<\s*\/t(d|h)\s*>/.test(t)?d+=t:d+="<"+(n?i+(o&&o[e]?' class="'+o[e]+'"':""):i)+">"+(n&&l&&l.length&&l[e]?l[e]:t)+"</"+i+">"}),d+"</tr>"}},g.buildComplete=function(e,t){f(e).triggerHandler(t.build_complete),e.config&&h.debug(e.config,"build")&&console.log("Build >> Table build complete"),h.setup(e,e.config)},g.array=function(e,t,l){return g.csv(e,t,l)},g.csv=function(e,t,l){var o,r,n="csv"===l.build_type||"string"==typeof t,i=f(e),t=n?t.replace("\r","").split("\n"):t,d=t.length,s=0,a=!1,u=l.build_headers.rows+(n?l.build_csvStartLine:0),c=l.build_footers.rows,b=0,h="",p=g.build.colgroup(l.build_headers.widths)+"<thead>";f.each(t,function(e,t){d-c<=e&&(a=!0),(!n||e>=l.build_csvStartLine)&&e<u?(r=n?g.splitCSV(t,l.build_csvSeparator):t,b=r.length,p+=g.build.header(r,l)):u<=e&&(e===u&&(p+="</thead><tbody>"),r=n?g.splitCSV(t,l.build_csvSeparator):t,a&&0<c&&(p+=(e===d-c?"</tbody><tfoot>":"")+(e===d?"</tfoot>":"")),1<r.length)&&(s++,r.length!==b&&(h+="error on line "+e+": Item count ("+r.length+") does not match header count ("+b+") \n"),o=a?l.build_footers.classes:"",p+=g.build.rows(r,l.build_footers.text,o,l,s,a))}),p+=0<c?"":"</tbody>",h?i.html(h):(i.html(p),g.buildComplete(e,l))},g.splitCSV=function(e,t){for(var l,o=f.trim(e).split(t=t||","),r=o.length-1;0<=r;r--)'"'===o[r].replace(/\"\s+$/,'"').charAt(o[r].length-1)?1<(l=o[r].replace(/^\s+\"/,'"')).length&&'"'===l.charAt(0)?o[r]=o[r].replace(/^\s*"|"\s*$/g,"").replace(/""/g,'"'):r?o.splice(r-1,2,[o[r-1],o[r]].join(t)):o=o.shift().split(t).concat(o):o[r].replace(/""/g,'"');return o},g.html=function(e,t,l){var o=f(e);t instanceof f?o.empty().append(t):o.html(t),g.buildComplete(e,l)},g.object=function(e,t,o){var l,r,n,i,d,s,a,u=e.config,c=o.build_objectHeaderKey,b=o.build_objectRowKey,c=t.hasOwnProperty(c)&&!f.isEmptyObject(t.kh)?t.kh:!!t.hasOwnProperty("headers")&&t.headers,b=t.hasOwnProperty(b)&&!f.isEmptyObject(t.kr)?t.kr:!!t.hasOwnProperty("rows")&&t.rows;if(!c||!b||0===c.length||0===b.length)return h.debug(u,"build")&&console.error("Build >> ERROR: Aborting build table widget, missing data for object build"),!1;i=f("<colgroup>"),d=f("<table><thead/></table>"),f.each(c,function(e,t){for(a=f("<tr>").appendTo(d.find("thead")),r=t.length,l=0;l<r;l++)(n=g.build.cell(t[l],o,"th",l,0===e))[0]&&n[0].length&&n[0].appendTo(a),0===e&&n[1]&&n[1].appendTo(i)}),i.find("col[style]").length&&d.prepend(i),s=f("<tbody>"),f.each(b,function(e,t){if((n="object"===f.type(t))&&t.newTbody)for(var l in s=f("<tbody>").appendTo(d),t)t.hasOwnProperty(l)&&"newTbody"!==l&&s.attr(l,t[l]);else{if(0===e&&s.appendTo(d),a=f("<tr>").appendTo(s),n){for(l in t)t.hasOwnProperty(l)&&l!==o.build_objectCellKey&&a.attr(l,t[l]);t.hasOwnProperty(o.build_objectCellKey)&&(t=t.cells)}for(r=t.length,l=0;l<r;l++)(i=g.build.cell(t[l],o,"td",l))[0]&&i[0].length&&i[0].appendTo(a)}}),t.hasOwnProperty(o.build_objectFooterKey)&&("clone"===(n=t[o.build_objectFooterKey])?(i=d.find("thead").html(),d.append("<tfoot>"+i+"</tfoot>")):(i=f("<tfoot>").appendTo(d),f.each(n,function(e,t){for(a=f("<tr>").appendTo(i),r=t.length,l=0;l<r;l++)(s=g.build.cell(t[l],o,"th",l))[0]&&s[0].length&&s[0].appendTo(a)}))),f(e).html(d.html()),g.buildComplete(e,o)},g.ajax=g.json=function(e,t,l){return g.object(e,t,l)}}(jQuery);return jQuery;}));
```
