## Supplementary Data SD1 for "Inferring binding specificities of human transcription factors with the wisdom of crowds": widget-chart.min.js

```
(function(factory){if (typeof define === 'function' && define.amd){define(['jquery'], factory);} else if (typeof module === 'object' && typeof module.exports === 'object'){module.exports = factory(require('jquery'));} else {factory(jQuery);}}(function(jQuery){
!function(l){"use strict";var n=l.tablesorter,s=[],c=[],u=[],a=[],_=[],f=[],d=[],g=[],p=n.chart={nonDigit:/[^\d,.\-()]/g,init:function(t,e){t.$table.off(e.chart_event).on(e.chart_event,function(){var t;this.hasInitialized&&(t=this.config,p.getCols(t,t.widgetOptions),p.getData(t,t.widgetOptions))})},getCols:function(t,e){var a;for(s=[],f=[],g=[],a=0;a<t.columns;a++)e.chart_useSelector&&n.hasWidget(t.table,"columnSelector")&&!t.selector.auto?(t.selector.states[a]&&l.inArray(a,e.chart_ignoreColumns)<0||a===e.chart_labelCol||a===e.chart_sort[0][0])&&s.push(a):(l.inArray(a,e.chart_ignoreColumns)<0||a===e.chart_labelCol||a===e.chart_sort[0][0])&&s.push(a)},getData:function(t,e){p.getHeaders(t,e),p.getRows(t,e),a=[c],l.each(u,function(t,e){a.push(e)}),t.chart={data:a,categories:_,series:f,category:d,dataset:g}},getHeaders:function(a,r){var o;f=[],g=[],(c=[]).push(a.headerContent[r.chart_labelCol]),l.each(s,function(t,e){if(e===r.chart_labelCol)return!0;o=a.headerContent[e],c.push(o),f.push({name:o,data:[]}),g.push({seriesname:o,data:[]})})},getRows:function(c,i){var t=c.cache[0].normalized,h=[];u=[],_=[],d=[],l.each(t,function(t,e){var a,r,o=e[c.columns].$row,s=o.children("th,td"),n=[];if(/v/i.test(i.chart_incRows)&&o.is(":visible")||/f/i.test(i.chart_incRows)&&!o.hasClass(i.filter_filteredRow||"filtered")||!/(v|f)/i.test(i.chart_incRows)){for(a=0;a<c.columns;a++)0<=l.inArray(t,i.chart_parsed)?n.push(e[a]):(r=s[a].getAttribute(c.textAttribute)||s[a].textContent||s.eq(a).text(),n.push(l.trim(r)));h.push(n)}}),h.sort(function(t,e){return 1===i.chart_sort[0][1]?n.sortNatural(e[i.chart_sort[0][0]],t[i.chart_sort[0][0]]):n.sortNatural(t[i.chart_sort[0][0]],e[i.chart_sort[0][0]])}),l.each(h,function(t,e){var a,r=0,o=[],s=e[i.chart_labelCol];o.push(""+s),l.each(e,function(t,e){if(t===i.chart_labelCol)return _.push(e),d.push({label:e}),!0;a=!1,i.chart_useSelector&&n.hasWidget(c.table,"columnSelector")&&!c.selector.auto?c.selector.states[t]&&l.inArray(t,i.chart_ignoreColumns)<0&&(a=""+e):l.inArray(t,i.chart_ignoreColumns)<0&&(a=""+e),!1!==a&&(/s/i.test(""+i.chart_layout[t])?(o.push(a),f[r].data.push(a),g[r].data.push(a)):(e=n.formatFloat(a.replace(p.nonDigit,""),c.table),e=isNaN(e)?a:e,o.push(e),f[r].data.push(e),g[r].data.push({value:e})),r++)}),u.push(o)})},remove:function(t,e){t.$table.off(e.chart_event)}};n.addWidget({id:"chart",options:{chart_incRows:"filtered",chart_useSelector:!1,chart_ignoreColumns:[],chart_parsed:[],chart_layout:{0:"string"},chart_labelCol:0,chart_sort:[[0,0]],chart_event:"chartData"},init:function(t,e,a,r){p.init(a,r)},remove:function(t,e,a){p.remove(e,a)}})}(jQuery);return jQuery;}));
```
