## Supplementary Data SD1 for "Inferring binding specificities of human transcription factors with the wisdom of crowds": widget-columns.min.js

```
(function(factory){if (typeof define === 'function' && define.amd){define(['jquery'], factory);} else if (typeof module === 'object' && typeof module.exports === 'object'){module.exports = factory(require('jquery'));} else {factory(jQuery);}}(function(jQuery){
/*! Widget: columns - updated 5/24/2017 (v2.28.11) */
!function(p){"use strict";var b=p.tablesorter||{};b.addWidget({id:"columns",priority:65,options:{columns:["primary","secondary","tertiary"]},format:function(e,r,o){for(var t,s,n,i,a,d=r.$table,l=r.$tbodies,c=r.sortList,h=c.length,f=o&&o.columns||["primary","secondary","tertiary"],m=f.length-1,y=f.join(" "),u=0;u<l.length;u++)(s=(t=b.processTbody(e,l.eq(u),!0)).children("tr")).each(function(){if(n=p(this),"none"!==this.style.display&&(i=n.children().removeClass(y),c)&&c[0]&&(i.eq(c[0][0]).addClass(f[0]),1<h))for(a=1;a<h;a++)i.eq(c[a][0]).addClass(f[a]||f[m])}),b.processTbody(e,t,!1);if(r=!1!==o.columns_thead?["thead tr"]:[],!1!==o.columns_tfoot&&r.push("tfoot tr"),r.length&&(s=d.find(r.join(",")).children().removeClass(y),h))for(a=0;a<h;a++)s.filter('[data-column="'+c[a][0]+'"]').addClass(f[a]||f[m])},remove:function(e,r,o){var t,s,n=r.$tbodies,i=(o.columns||["primary","secondary","tertiary"]).join(" ");for(r.$headers.removeClass(i),r.$table.children("tfoot").children("tr").children("th, td").removeClass(i),t=0;t<n.length;t++)(s=b.processTbody(e,n.eq(t),!0)).children("tr").each(function(){p(this).children().removeClass(i)}),b.processTbody(e,s,!1)}})}(jQuery);return jQuery;}));
```
