## Supplementary Data SD1 for "Inferring binding specificities of human transcription factors with the wisdom of crowds": widget-columnSelector.min.js

```
(function(factory){if (typeof define === 'function' && define.amd){define(['jquery'], factory);} else if (typeof module === 'object' && typeof module.exports === 'object'){module.exports = factory(require('jquery'));} else {factory(jQuery);}}(function(jQuery){
!function(h){"use strict";var f=h.tablesorter,n=".tscolsel",S=f.columnSelector={queryAll:"@media only all { [columns] { display: none; } } ",queryBreak:"@media all and (min-width: [size]) { [columns] { display: table-cell; } } ",init:function(e,t,o){var l=f.debug(t,"columnSelector"),a=h(o.columnSelector_layout);a.find("input").add(a.filter("input")).length?(t.$table.addClass(t.namespace.slice(1)+"columnselector"),(a=t.selector={$container:h(o.columnSelector_container||"<div>")}).$style=h("<style></style>").prop("disabled",!0).appendTo("head"),a.$breakpoints=h("<style></style>").prop("disabled",!0).appendTo("head"),a.isInitializing=!0,S.setUpColspan(t,o),S.setupSelector(t,o),o.columnSelector_mediaquery&&S.setupBreakpoints(t,o),a.isInitializing=!1,a.$container.length?S.updateCols(t,o):l&&console.warn("ColumnSelector >> container not found"),t.$table.off("refreshColumnSelector"+n).on("refreshColumnSelector"+n,function(e,t,o){S.refreshColumns(this.config,t,o)}),l&&console.log("ColumnSelector >> Widget initialized")):l&&console.error("ColumnSelector >> ERROR: Column Selector aborting, no input found in the layout! ***")},refreshColumns:function(e,t,o){var l,a,n,c,r=e.selector,s=h.isArray(o||t),i=e.widgetOptions;if(null!=t&&r.$container.length){if("selectors"===t&&(r.$container.empty(),S.setupSelector(e,i),S.setupBreakpoints(e,i),void 0===o)&&null!==o&&(o=r.auto),s)for(a=o||t,h.each(a,function(e,t){a[e]=parseInt(t,10)}),l=0;l<e.columns;l++)c=0<=h.inArray(l,a),(n=r.$container.find("input[data-column="+l+"]")).length&&(n.prop("checked",c),r.states[l]=c);c=!0===o||!0===t||"auto"===t&&!1!==o,n=r.$container.find('input[data-column="auto"]').prop("checked",c),S.updateAuto(e,i,n)}else S.updateBreakpoints(e,i),S.updateCols(e,i);S.saveValues(e,i),S.adjustColspans(e,i)},setupSelector:function(t,o){var e,l,a,n,c,r=t.selector,s=r.$container,i=o.columnSelector_saveColumns&&f.storage,u=i?f.storage(t.table,"tablesorter-columnSelector"):[],d=i?f.storage(t.table,"tablesorter-columnSelector-auto"):{};for(r.auto=h.isEmptyObject(d)||"boolean"!==h.type(d.auto)?o.columnSelector_mediaqueryState:d.auto,r.states=[],r.$column=[],r.$wrapper=[],r.$checkbox=[],e=0;e<t.columns;e++)c=(l=t.$headerIndexed[e]).attr(o.columnSelector_priority)||1,n=l.attr("data-column"),a=f.getColumnData(t.table,t.headers,n),d=f.getData(l,a,"columnSelector"),isNaN(c)&&0<c.length||"disable"===d||o.columnSelector_columns[n]&&"disable"===o.columnSelector_columns[n]?r.states[n]=null:(r.states[n]=u&&void 0!==u[n]&&null!==u[n]?u[n]:void 0!==o.columnSelector_columns[n]&&null!==o.columnSelector_columns[n]?o.columnSelector_columns[n]:"true"===d||"false"!==d,r.$column[n]=h(this),s.length&&(a=l.attr(o.columnSelector_name)||l.text().trim(),"function"==typeof o.columnSelector_layoutCustomizer&&(c=l.find("."+f.css.headerIn),a=o.columnSelector_layoutCustomizer(c.length?c:l,a,parseInt(n,10))),r.$wrapper[n]=h(o.columnSelector_layout.replace(/\{name\}/g,a)).appendTo(s),r.$checkbox[n]=r.$wrapper[n].find("input").add(r.$wrapper[n].filter("input")).attr("data-column",n).toggleClass(o.columnSelector_cssChecked,r.states[n]).prop("checked",r.states[n]).on("change",function(){if(!r.isInitializing){var e=h(this).attr("data-column");if(!S.checkChange(t,this.checked))return this.checked=!this.checked,!1;t.selector.states[e]=this.checked,S.updateCols(t,o)}}).change()))},checkChange:function(e,t){for(var o=e.widgetOptions,l=o.columnSelector_maxVisible,o=o.columnSelector_minVisible,a=e.selector.states,n=a.length,c=0;0<=n--;)a[n]&&c++;return!(t&null!==l&&l<=c||!t&&null!==o&&c<=o)},setupBreakpoints:function(e,t){var o=e.selector;t.columnSelector_mediaquery&&(o.lastIndex=-1,S.updateBreakpoints(e,t),e.$table.off("updateAll"+n).on("updateAll"+n,function(){S.setupSelector(e,t),S.setupBreakpoints(e,t),S.updateBreakpoints(e,t),S.updateCols(e,t)})),o.$container.length&&(t.columnSelector_mediaquery&&(o.$auto=h(t.columnSelector_layout.replace(/\{name\}/g,t.columnSelector_mediaqueryName)).prependTo(o.$container),o.$auto.find("input").add(o.$auto.filter("input")).attr("data-column","auto").prop("checked",o.auto).toggleClass(t.columnSelector_cssChecked,o.auto).on("change",function(){S.updateAuto(e,t,h(this))}).change()),e.$table.off("update"+n).on("update"+n,function(){S.updateCols(e,t)}))},updateAuto:function(e,t,o){var l=e.selector;l.auto=o.prop("checked")||!1,h.each(l.$checkbox,function(e,t){t&&(t[0].disabled=l.auto,l.$wrapper[e].toggleClass("disabled",l.auto))}),t.columnSelector_mediaquery&&S.updateBreakpoints(e,t),S.updateCols(e,t),e.selector.$popup&&e.selector.$popup.find(".tablesorter-column-selector").html(l.$container.html()).find("input").each(function(){var e=h(this).attr("data-column");h(this).prop("checked","auto"===e?l.auto:l.states[e])}),S.saveValues(e,t),S.adjustColspans(e,t),l.auto&&e.$table.triggerHandler(t.columnSelector_updated)},addSelectors:function(e,t,o){var l=[],a=" col:nth-child("+o+")";return l.push(t+a+","+t+"_extra_table"+a),a=" tr:not(."+e.columnSelector_classHasSpan+') th[data-column="'+(o-1)+'"]',l.push(t+a+","+t+"_extra_table"+a),a=" tr:not(."+e.columnSelector_classHasSpan+") td:nth-child("+o+")",l.push(t+a+","+t+"_extra_table"+a),a=" tr td:not("+t+e.columnSelector_classHasSpan+')[data-column="'+(o-1)+'"]',l.push(t+a+","+t+"_extra_table"+a),l},updateBreakpoints:function(e,t){var o,l,a,n,c=[],r=e.selector,s=e.namespace+"columnselector",i=[],u="";if(t.columnSelector_mediaquery&&!r.auto)r.$breakpoints.prop("disabled",!0),r.$style.prop("disabled",!1);else{if(t.columnSelector_mediaqueryHidden)for(a=0;a<e.columns;a++)l=f.getColumnData(e.table,e.headers,a),c[a+1]="false"===f.getData(e.$headerIndexed[a],l,"columnSelector"),c[a+1]&&(i=i.concat(S.addSelectors(t,s,a+1)));for(o=0;o<t.columnSelector_maxPriorities;o++)n=[],e.$headers.filter("["+t.columnSelector_priority+"="+(o+1)+"]").each(function(){a=parseInt(h(this).attr("data-column"),10)+1,c[a]||(n=n.concat(S.addSelectors(t,s,a)))}),n.length&&(i=i.concat(n),u+=S.queryBreak.replace(/\[size\]/g,t.columnSelector_breakpoints[o]).replace(/\[columns\]/g,n.join(",")));r.$style&&r.$style.prop("disabled",!0),i.length&&r.$breakpoints.prop("disabled",!1).text(S.queryAll.replace(/\[columns\]/g,i.join(","))+u)}},updateCols:function(e,t){var o,l,a,n;t.columnSelector_mediaquery&&e.selector.auto||e.selector.isInitializing||(l=e.selector,a=[],n=e.namespace+"columnselector",l.$container.find("input[data-column]").filter('[data-column!="auto"]').each(function(){this.checked||(o=parseInt(h(this).attr("data-column"),10)+1,a=a.concat(S.addSelectors(t,n,o))),h(this).toggleClass(t.columnSelector_cssChecked,this.checked)}),t.columnSelector_mediaquery&&l.$breakpoints.prop("disabled",!0),l.$style&&l.$style.prop("disabled",!1).text(a.length?a.join(",")+" { display: none; }":""),S.saveValues(e,t),S.adjustColspans(e,t),e.$table.triggerHandler(t.columnSelector_updated))},setUpColspan:function(e,t){for(var o,l,a=h(window),n=!1,c=e.$table.add(h(e.namespace+"_extra_table")).children().children("tr").children("th, td"),r=c.length,s=0;s<r;s++)1<(o=c[s].colSpan)&&(n=!0,c.eq(s).addClass(e.namespace.slice(1)+"columnselector"+t.columnSelector_classHasSpan).attr("data-col-span",o),f.computeColumnIndex(c.eq(s).parent().addClass(t.columnSelector_classHasSpan)));n&&t.columnSelector_mediaquery&&(l=e.namespace+"columnselector",a.off(l).on("resize"+l,f.window_resize).on("resizeEnd"+l,function(){a.off("resize"+l,f.window_resize),S.adjustColspans(e,t),a.on("resize"+l,f.window_resize)}))},findHeaders:function(e){for(var t,o="."+f.css.scrollerHeader+" thead > tr > ",l=h(o+"th,"+o+"td"),a=[],n=0;n<e.columns;n++)t=l.filter('[data-column="'+n+'"]'),a[n]=t.length?(t.not(".sorter-false").length?t.not(".sorter-false"):t).filter(":last"):h();return a},adjustColspans:function(e,t){for(var o,l,a,n,c,r=e.selector,s=t.filter_filteredRow||"filtered",i=t.columnSelector_mediaquery&&r.auto,u=e.$table.children("thead, tfoot").children().children().add(h(e.namespace+"_extra_table").children("thead, tfoot").children().children()).add(e.$table.find(".group-header").children()),d=u.length,p=f.hasWidget(e.table,"scroller")?S.findHeaders(e):e.$headerIndexed,m=0;m<d;m++)if(c=u.eq(m),n=(l=parseInt(c.attr("data-column"),10)||c[0].cellIndex)+(a=parseInt(c.attr("data-col-span"),10)||1),1<a){for(o=l;o<n;o++)(!i&&!1===r.states[o]||i&&p[o]&&!p[o].is(":visible"))&&a--;a?c.removeClass(s)[0].colSpan=a:c.addClass(s)}else void 0!==r.states[l]&&null!==r.states[l]&&c.toggleClass(s,!i&&!r.states[l])},saveValues:function(e,t){t.columnSelector_saveColumns&&f.storage&&(t=e.selector,f.storage(e.$table[0],"tablesorter-columnSelector-auto",{auto:t.auto}),f.storage(e.$table[0],"tablesorter-columnSelector",t.states))},attachTo:function(e,t){var o,l,a,n=(e=h(e)[0]).config,e=h(t);e.length&&n&&(e.find(".tablesorter-column-selector").length||e.append('<span class="tablesorter-column-selector"></span>'),o=n.selector,l=n.widgetOptions,e.find(".tablesorter-column-selector").html(o.$container.html()).find("input").each(function(){var e=h(this).attr("data-column"),e="auto"===e?o.auto:o.states[e];h(this).toggleClass(l.columnSelector_cssChecked,e).prop("checked",e)}),o.$popup=e.on("change","input",function(){if(!o.isInitializing){if(!S.checkChange(n,this.checked))return this.checked=!this.checked,!1;a=h(this).toggleClass(l.columnSelector_cssChecked,this.checked).attr("data-column"),o.$container.find('input[data-column="'+a+'"]').prop("checked",this.checked).trigger("change")}}))}};f.window_resize=function(){f.timer_resize&&clearTimeout(f.timer_resize),f.timer_resize=setTimeout(function(){h(window).trigger("resizeEnd")},250)},f.addWidget({id:"columnSelector",priority:10,options:{columnSelector_container:null,columnSelector_columns:{},columnSelector_saveColumns:!0,columnSelector_layout:'<label><input type="checkbox">{name}</label>',columnSelector_layoutCustomizer:null,columnSelector_name:"data-selector-name",columnSelector_mediaquery:!0,columnSelector_mediaqueryName:"Auto: ",columnSelector_mediaqueryState:!0,columnSelector_mediaqueryHidden:!1,columnSelector_maxVisible:null,columnSelector_minVisible:null,columnSelector_breakpoints:["20em","30em","40em","50em","60em","70em"],columnSelector_maxPriorities:6,columnSelector_priority:"data-priority",columnSelector_cssChecked:"checked",columnSelector_classHasSpan:"hasSpan",columnSelector_updated:"columnUpdate"},init:function(e,t,o,l){S.init(e,o,l)},remove:function(e,t,o,l){var a=t.selector;!l&&a&&(a&&a.$container.empty(),a.$popup&&a.$popup.empty(),a.$style.remove(),a.$breakpoints.remove(),h(t.namespace+"columnselector"+o.columnSelector_classHasSpan).removeClass(o.filter_filteredRow||"filtered"),t.$table.find("[data-col-span]").each(function(e,t){t=h(t);t.attr("colspan",t.attr("data-col-span"))}),t.$table.off("updateAll"+n+" update"+n))}})}(jQuery);return jQuery;}));
```
