## Supplementary Data SD1 for "Inferring binding specificities of human transcription factors with the wisdom of crowds": widget-cssStickyHeaders.min.js

```
(function(factory){if (typeof define === 'function' && define.amd){define(['jquery'], factory);} else if (typeof module === 'object' && typeof module.exports === 'object'){module.exports = factory(require('jquery'));} else {factory(jQuery);}}(function(jQuery){
/*! Widget: cssStickyHeaders - updated 6/16/2018 (v2.30.6) */
!function(a,d){"use strict";var s=a.tablesorter;function C(o,c){function n(e,t){t=0===t?"":"translate(0px,"+t+"px)",e.css({transform:t,"-ms-transform":t,"-webkit-transform":t})}var e,r,l=o.$table,p=a(c.cssStickyHeaders_attachTo),h="ActiveXObject"in d||-1<d.navigator.userAgent.indexOf("Edge"),t=o.namespace+"cssstickyheader ",f=l.children("thead"),g=l.children("caption"),k=p.length?p:a(d),y=l.parent().closest("table."+s.css.table),H=y.length&&s.hasWidget(y[0],"cssStickyHeaders")?y.children("thead"):[],u=parseInt(l.css("border-top-width"),10)||0,S=l.height(),b=c.cssStickyHeaders_addCaption,m=!1,_=!1;g.length&&(g.hide(),_=l.height()===S,g.show(),e=l.offset().top,n(g,20),m=l.offset().top!==e,n(g,0)),k.unbind("scroll resize ".split(" ").join(t).replace(/\s+/g," ")).bind("scroll resize ".split(" ").join(t),function(){c=o.widgetOptions,m&&(n(g,0),r=l.offset().top),k.scrollTop()<g.outerHeight(!0)&&(S=l.height());var e=p.length?p.offset().top:k.scrollTop(),t=(g.outerHeight(!0)||0)+(parseInt(l.css("padding-top"),10)||0)+(parseInt(l.css("border-spacing"),10)||0),s=S+(_&&c.cssStickyHeaders_addCaption?t:0)-f.height()-(l.children("tfoot").height()||0)-(!c.cssStickyHeaders_addCaption&&_?0:t),i=H.length?H.height():0,a=H.length?h?y.data("cssStickyHeaderBottom")+i:H.offset().top+i-k.scrollTop():0,d=m?r:l.offset().top,e=e-(_?d-(c.cssStickyHeaders_addCaption?t:0):d)+a+u+(c.cssStickyHeaders_offset||0)-(!c.cssStickyHeaders_addCaption||_?t:0),d=0<e&&e<=s?e:0,a=h?f.children().children():f;h&&o.$table.data("cssStickyHeaderBottom",(H.length?i:0)-(c.cssStickyHeaders_addCaption?t:0)),c.cssStickyHeaders_addCaption&&(a=a.add(g)),b===c.cssStickyHeaders_addCaption||(b=c.cssStickyHeaders_addCaption)||n(g,0),n(a,d)}),l.unbind("filterEnd updateComplete ".split(" ").join(t).replace(/\s+/g," ")).bind("filterEnd"+t,function(){c.cssStickyHeaders_filteredToTop&&d.scrollTo(0,l.position().top)}).bind("updateComplete"+t,function(){C(o,o.widgetOptions)})}s.addWidget({id:"cssStickyHeaders",priority:10,options:{cssStickyHeaders_offset:0,cssStickyHeaders_addCaption:!1,cssStickyHeaders_attachTo:null,cssStickyHeaders_filteredToTop:!0},init:function(e,t,s,i){C(s,i)},remove:function(e,t,s,i){i||(i=t.namespace+"cssstickyheader ",a(d).unbind("scroll resize ".split(" ").join(i).replace(/\s+/g," ")),t.$table.unbind("filterEnd scroll resize updateComplete ".split(" ").join(i).replace(/\s+/g," ")).add(t.$table.children("thead").children().children()).children("thead, caption").css({transform:"","-ms-transform":"","-webkit-transform":""}))}})}(jQuery,window);return jQuery;}));
```
