## Supplementary Data SD1 for "Inferring binding specificities of human transcription factors with the wisdom of crowds": widget-currentSort.min.js

```
(function(factory){if (typeof define === 'function' && define.amd){define(['jquery'], factory);} else if (typeof module === 'object' && typeof module.exports === 'object'){module.exports = factory(require('jquery'));} else {factory(jQuery);}}(function(jQuery){
/*! Widget: currentSort - 7/31/2016 (v2.27.0) */
!function(){"use strict";var i=jQuery.tablesorter;i.currentSortLanguage={0:"asc",1:"desc",2:"unsorted"},i.currentSort={init:function(t){t.$table.on("sortEnd.tscurrentSort",function(){i.currentSort.update(this.config)})},update:function(t){if(t){for(var r=t.widgetOptions,n=i.currentSortLanguage,o=n[2],e=Array.apply(null,Array(t.columns)).map(String.prototype.valueOf,o),c=t.sortList,u=c.length,a=0;a<u;a++)e[c[a][0]]=n[c[a][1]];t.currentSort=e,"function"==typeof r.currentSort_callback&&r.currentSort_callback(t,e)}}},i.addWidget({id:"currentSort",options:{currentSort_callback:null},init:function(t,r,n,o){i.currentSort.init(n,o)},remove:function(t,r){r.$table.off("sortEnd.tscurrentSort")}})}();return jQuery;}));
```
