## Supplementary Data SD1 for "Inferring binding specificities of human transcription factors with the wisdom of crowds": widget-editable.min.js

```
(function(factory){if (typeof define === 'function' && define.amd){define(['jquery'], factory);} else if (typeof module === 'object' && typeof module.exports === 'object'){module.exports = factory(require('jquery'));} else {factory(jQuery);}}(function(jQuery){
/*! Widget: editable - updated 2018-08-27 (v2.31.0) */
!function(b){"use strict";var p=b.tablesorter.editable={namespace:".tseditable",lastEdited:"tseditable-last-edited-cell",editComplete:function(e,t,n,i){e.$table.find("."+p.lastEdited).removeClass(p.lastEdited).trigger(t.editable_editComplete,[e]),i&&setTimeout(function(){n.focus()},50)},selectAll:function(n){setTimeout(function(){var e,t;document.queryCommandSupported("SelectAll")?document.execCommand("selectAll",!1,null):document.body.createTextRange?((e=document.body.createTextRange()).moveToElementText(n),e.select()):window.getSelection&&(t=window.getSelection(),(e=document.createRange()).selectNodeContents(n),t.removeAllRanges(),t.addRange(e))},100)},getColumns:function(e,t){var n,i,o,a,l,d=t.editable_columns,s=[];if("string"==typeof d)for(a=(n=d.replace(/\s+/,"").split(/,/)).length-1;0<=a;){if(0<=n[a].indexOf("-"))for(o=n[a].split("-"),i=parseInt(o[0],10)||0,(o=parseInt(o[1],10)||e.columns-1)<i&&(l=i,i=o,o=l);i<=o;i++)s.push("td:nth-child("+(i+1)+")");else s.push("td:nth-child("+((parseInt(n[a],10)||0)+1)+")");a--}else if(b.isArray(d))for(a=d.length,i=0;i<a;i++)d[i]<e.columns&&s.push("td:nth-child("+(d[i]+1)+")");return s},trimContent:function(e,t){e.editable_trimContent&&(e=t.html()).trim()!==e&&t.html(""===e?"&nbsp;":e)},update:function(e,t){var n,i,o,a,l,d,s,c=b("<div>").wrapInner(t.editable_wrapContent).children().length||b.isFunction(t.editable_wrapContent),r=p.getColumns(e,t).join(",");for(e.$tbodies.find(r).find("[contenteditable]").prop("contenteditable",!1),a=(i=e.$tbodies.find(r).not("."+t.editable_noEdit)).length,o=0;o<a;o++)if(n=i.eq(o),c&&0===n.children("div, span").length&&(n.wrapInner(t.editable_wrapContent),""===n.children().text().trim())&&n.children().html("&nbsp;"),s=(l=n.children("div, span").not("."+t.editable_noEdit)).length)for(d=0;d<s;d++){var u=l.eq(d);p.trimContent(t,u),u.prop("contenteditable",!0)}else p.trimContent(t,n),n.prop("contenteditable",!0)},bindEvents:function(d,s){var c=p.namespace;d.$table.off("updateComplete pagerComplete ".split(" ").join(c+" ").replace(/\s+/g," ")).on("updateComplete pagerComplete ".split(" ").join(c+" "),function(){p.update(d,d.widgetOptions)}).children("thead").add(b(d.namespace+"_extra_table").children("thead")).off("mouseenter"+c).on("mouseenter"+c,function(){d.$table.data("contentFocused")&&(d.$table.data("contentFocused",!0),b(":focus").trigger("focusout"))}),d.$tbodies.off("focus focusout keydown ".split(" ").join(c+" ").replace(/\s+/g," ")).on("focus"+c,"[contenteditable]",function(e){clearTimeout(b(this).data("timer")),d.$table.data("contentFocused",e.target),d.table.isUpdating=!0;var e=b(this),t=s.editable_selectAll,n=e.closest("td").index(),i=e.html();s.editable_trimContent&&(i=b.trim(""===i?"&nbsp;":i)),e.off("keydown"+c).on("keydown"+c,function(e){s.editable_enterToAccept&&13===e.which&&!e.shiftKey&&e.preventDefault()}),e.data({before:i,original:i}),"function"==typeof s.editable_focused&&s.editable_focused(i,n,e),t&&("function"!=typeof t||t(i,n,e))&&p.selectAll(e[0])}).on("focusout keydown ".split(" ").join(c+" "),"[contenteditable]",function(e){if(d.$table.data("contentFocused")){var t,n,i=!1,o=b(e.target),a=o.html(),l=o.closest("td").index();if(s.editable_trimContent&&(a=b.trim(""===a?"&nbsp;":a)),27===e.which)return o.html(o.data("original")).trigger("blur"+c),d.$table.data("contentFocused",!1),d.table.isUpdating=!1;if((t=13===e.which&&!e.shiftKey&&(s.editable_enterToAccept||e.altKey)||s.editable_autoAccept&&"keydown"!==e.type)&&o.data("before")!==a){if(n=s.editable_validate,i=a,("function"==typeof n||"function"==typeof(n=b.tablesorter.getColumnData(d.table,n,l)))&&(i=n(a,o.data("original"),l,o)),t&&!1!==i)return d.$table.find("."+p.lastEdited).removeClass(p.lastEdited),o.addClass(p.lastEdited).html(i).data("before",i).data("original",i).trigger("change"),d.table.hasInitialized&&b.tablesorter.updateCell(d,o.closest("td"),!1,function(){s.editable_autoResort?setTimeout(function(){b.tablesorter.sortOn(d,d.sortList,function(){p.editComplete(d,s,d.$table.data("contentFocused"),!0)},!0)},10):p.editComplete(d,s,d.$table.data("contentFocused"))}),!1}else i||"keydown"===e.type||(clearTimeout(o.data("timer")),o.data("timer",setTimeout(function(){d.table.isUpdating=!1,b.isFunction(s.editable_blur)&&(a=o.html(),s.editable_blur(s.editable_trimContent?b.trim(a):a,l,o))},100)),o.html(o.data("original")))}}).on("paste"+c,"[contenteditable]",function(){var e,t=b(this);setTimeout(function(){t.is(":focus")&&(e="<div>"+t.html()+"</div>",t.html(b(e).text().trim()))},0)})},destroy:function(e,t){var n=p.namespace,t=p.getColumns(e,t),i="updateComplete pagerComplete ".split(" ").join(n+" ").replace(/\s+/g," ");e.$table.off(i),i="focus focusout keydown paste ".split(" ").join(n+" ").replace(/\s+/g," "),e.$tbodies.off(i).find(t.join(",")).find("[contenteditable]").prop("contenteditable",!1)}};b.tablesorter.addWidget({id:"editable",options:{editable_columns:[],editable_enterToAccept:!0,editable_autoAccept:!0,editable_autoResort:!1,editable_wrapContent:"<div>",editable_trimContent:!0,editable_validate:null,editable_focused:null,editable_blur:null,editable_selectAll:!1,editable_noEdit:"no-edit",editable_editComplete:"editComplete"},init:function(e,t,n,i){i.editable_columns.length&&(p.update(n,i),p.bindEvents(n,i))},remove:function(e,t,n,i){i||p.destroy(t,n)}})}(jQuery);return jQuery;}));
```
