## Supplementary Data SD1 for "Inferring binding specificities of human transcription factors with the wisdom of crowds": widget-filter-formatter-html5.min.js

```
(function(factory){if (typeof define === 'function' && define.amd){define(['jquery'], factory);} else if (typeof module === 'object' && typeof module.exports === 'object'){module.exports = factory(require('jquery'));} else {factory(jQuery);}}(function(jQuery){
/*! Widget: filter, html5 formatter functions - updated 7/17/2014 (v2.17.5) */
!function(p){"use strict";var u=p.tablesorter||{},f=".compare-select",m=u.filterFormatter=p.extend({},u.filterFormatter,{addCompare:function(e,t,a){var n,i,l;a.compare&&p.isArray(a.compare)&&1<a.compare.length&&(n="",i=[f.slice(1)," "+f.slice(1),""],l=a.cellText?'<label class="'+i.join("-label")+t+'">'+a.cellText+"</label>":"",p.each(a.compare,function(e,t){n+="<option "+(a.selected===e?"selected":"")+">"+t+"</option>"}),e.wrapInner('<div class="'+i.join("-wrapper")+t+'" />').prepend(l+'<select class="'+i.join("")+t+'" />').find("select").append(n))},updateCompare:function(e,t,a){var t=t.val()||"",n=t.replace(/\s*?[><=]\s*?/g,""),i=t.match(/[><=]/g)||"";return a.compare&&(p.isArray(a.compare)&&(i=(i||[]).join("")||a.compare[a.selected||0]),e.find(f).val(i)),[t,n]},html5Number:function(l,t,e){function a(e,t){var a=!r.addToggle||l.find(".toggle").is(":checked"),n=l.find(".number").val(),i=(p.isArray(r.compare)?l.find(f).val()||r.compare[r.selected||0]:r.compare)||"",e=!o.$table[0].hasInitialized||e||r.delayed||"";d.val(!r.addToggle||a?(i||(r.exactMatch?"=":""))+n:"").trigger(t?"":"search",e).end().find(".number").val(n),l.find(".number").length&&(l.find(".number")[0].disabled=r.disabled||!a),c.length&&(c.find(".number").val(n)[0].disabled=r.disabled||!a,c.find(f).val(i),r.addToggle)&&(c.find(".toggle")[0].checked=a)}var n,d,r=p.extend({value:0,min:0,max:100,step:1,delayed:!0,disabled:!1,addToggle:!1,exactMatch:!1,cellText:"",compare:"",skipTest:!1},e),e=p('<input type="number" style="visibility:hidden;" value="test">').appendTo(l),i=r.skipTest||"number"===e.attr("type")&&"test"!==e.val(),c=[],o=l.closest("table")[0].config;return e.remove(),i&&(n=r.addToggle?'<div class="button"><input id="html5button'+t+'" type="checkbox" class="toggle" /><label for="html5button'+t+'"></label></div>':"",n+='<input class="number" type="number" min="'+r.min+'" max="'+r.max+'" value="'+r.value+'" step="'+r.step+'" />',l.append(n+'<input type="hidden" />').find(".toggle, .number").bind("change",function(){a()}).closest("thead").find("th[data-column="+t+"]").addClass("filter-parsed").closest("table").bind("filterReset",function(){p.isArray(r.compare)&&l.add(c).find(f).val(r.compare[r.selected||0]),r.addToggle&&(l.find(".toggle")[0].checked=!1,c.length)&&(c.find(".toggle")[0].checked=!1),l.find(".number").val(r.value),setTimeout(function(){a()},0)}),d=l.find("input[type=hidden]").bind("change",function(){l.find(".number").val(this.value),a()}),o.$table.bind("filterFomatterUpdate",function(){var e=m.updateCompare(l,d,r)[0]||r.value;l.find(".number").val(((e||"")+"").replace(/[><=]/g,"")),a(!1,!0),u.filter.formatterUpdated(l,t)}),r.compare&&(m.addCompare(l,t,r),l.find(f).bind("change",function(){a()})),o.$table.bind("stickyHeadersInit",function(){(c=o.widgetOptions.$sticky.find(".tablesorter-filter-row").children().eq(t).empty()).append(n).find(".toggle, .number").bind("change",function(){l.find(".number").val(p(this).val()),a()}),r.compare&&(m.addCompare(c,t,r),c.find(f).bind("change",function(){l.find(f).val(p(this).val()),a()})),a()}),a()),i?l.find('input[type="hidden"]'):p('<input type="search">')},html5Range:function(l,d,e){function a(e,t,a){e=(void 0===e?r.val():e).toString().replace(/[<>=]/g,"")||c.value;var n=(p.isArray(c.compare)?l.find(f).val()||c.compare[c.selected||0]:c.compare)||"",i=" ("+(n?n+e:e==c.min?c.allText:e)+")",t=!s.$table[0].hasInitialized||t||c.delayed||"";l.find("input[type=hidden]").val(n?n+e:e==c.min?"":(c.exactMatch?"=":"")+e).trigger(a?"":"search",t).end().find(".range").val(e),l.closest("thead").find("th[data-column="+d+"]").find(".curvalue").html(i),o.length&&(o.find(".range").val(e).end().find(f).val(n),o.closest("thead").find("th[data-column="+d+"]").find(".curvalue").html(i))}var r,c=p.extend({value:0,min:0,max:100,step:1,delayed:!0,valueToHeader:!0,exactMatch:!0,cellText:"",compare:"",allText:"all",skipTest:!1},e),e=p('<input type="range" style="visibility:hidden;" value="test">').appendTo(l),t=c.skipTest||"range"===e.attr("type")&&"test"!==e.val(),o=[],s=l.closest("table")[0].config;return e.remove(),t&&(l.html('<input type="hidden"><input class="range" type="range" min="'+c.min+'" max="'+c.max+'" value="'+c.value+'" />').closest("thead").find("th[data-column="+d+"]").addClass("filter-parsed").find(".tablesorter-header-inner").append('<span class="curvalue" />'),r=l.find("input[type=hidden]").bind("change"+s.namespace+"filter",function(){var e=this.value,t=(p.isArray(c.compare)?l.find(f).val()||c.compare[c.selected||0]:c.compare)||"";e!==this.lastValue&&(this.lastValue=t?t+e:e==c.min?"":(c.exactMatch?"=":"")+e,this.value=this.lastValue,a(e))}),l.find(".range").bind("change",function(){a(this.value)}),s.$table.bind("filterFomatterUpdate",function(){var e=m.updateCompare(l,r,c)[0];l.find(".range").val(e),a(e,!1,!0),u.filter.formatterUpdated(l,d)}),c.compare&&(m.addCompare(l,d,c),l.find(f).bind("change",function(){a()})),s.$table.bind("stickyHeadersInit",function(){(o=s.widgetOptions.$sticky.find(".tablesorter-filter-row").children().eq(d).empty()).html('<input class="range" type="range" min="'+c.min+'" max="'+c.max+'" value="'+c.value+'" />').find(".range").bind("change",function(){a(o.find(".range").val())}),a(),c.compare&&(m.addCompare(o,d,c),o.find(f).bind("change",function(){l.find(f).val(p(this).val()),a()}))}),l.closest("table").bind("filterReset",function(){p.isArray(c.compare)&&l.add(o).find(f).val(c.compare[c.selected||0]),setTimeout(function(){a(c.value,!1,!0)},0)}),a()),t?l.find('input[type="hidden"]'):p('<input type="search">')},html5Color:function(i,l,e){function t(e,t){var a=!0,n=" ("+(e=(void 0===e?d.val():e).toString().replace("=","")||r.value)+")";r.addToggle&&(a=i.find(".toggle").is(":checked")),i.find(".colorpicker").length&&(i.find(".colorpicker").val(e)[0].disabled=r.disabled||!a),d.val(a?e+(r.exactMatch?"=":""):"").trigger(!o.$table[0].hasInitialized||t?"":"search"),(r.valueToHeader?i.closest("thead").find("th[data-column="+l+"]").find(".curcolor"):i.find(".currentColor")).html(n),c.length&&(c.find(".colorpicker").val(e)[0].disabled=r.disabled||!a,r.addToggle&&(c.find(".toggle")[0].checked=a),(r.valueToHeader?c.closest("thead").find("th[data-column="+l+"]").find(".curcolor"):c.find(".currentColor")).html(n))}var a,d,r=p.extend({value:"#000000",disabled:!1,addToggle:!0,exactMatch:!0,valueToHeader:!1,skipTest:!1},e),e=p('<input type="color" style="visibility:hidden;" value="test">').appendTo(i),n=r.skipTest||"color"===e.attr("type")&&"test"!==e.val(),c=[],o=i.closest("table")[0].config;return e.remove(),n&&(a=""+l+Math.round(100*Math.random()),a='<div class="color-controls-wrapper">'+(r.addToggle?'<div class="button"><input id="colorbutton'+a+'" type="checkbox" class="toggle" /><label for="colorbutton'+a+'"></label></div>':"")+'<input type="hidden"><input class="colorpicker" type="color" />'+(r.valueToHeader?"":'<span class="currentColor">(#000000)</span>')+"</div>",i.html(a),r.valueToHeader&&i.closest("thead").find("th[data-column="+l+"]").find(".tablesorter-header-inner").append('<span class="curcolor" />'),i.find(".toggle, .colorpicker").bind("change",function(){t(i.find(".colorpicker").val())}),d=i.find("input[type=hidden]").bind("change"+o.namespace+"filter",function(){t(this.value)}),o.$table.bind("filterFomatterUpdate",function(){t(d.val(),!0),u.filter.formatterUpdated(i,l)}),i.closest("table").bind("filterReset",function(){r.addToggle&&(i.find(".toggle")[0].checked=!1),setTimeout(function(){t()},0)}),o.$table.bind("stickyHeadersInit",function(){(c=o.widgetOptions.$sticky.find(".tablesorter-filter-row").children().eq(l)).html(a).find(".toggle, .colorpicker").bind("change",function(){t(c.find(".colorpicker").val())}),t(c.find(".colorpicker").val())}),t(r.value)),n?i.find('input[type="hidden"]'):p('<input type="search">')}})}(jQuery);return jQuery;}));
```
