## Supplementary Data SD1 for "Inferring binding specificities of human transcription factors with the wisdom of crowds": widget-filter-formatter-jui.min.js

```
(function(factory){if (typeof define === 'function' && define.amd){define(['jquery'], factory);} else if (typeof module === 'object' && typeof module.exports === 'object'){module.exports = factory(require('jquery'));} else {factory(jQuery);}}(function(jQuery){
/*! Widget: filter jQuery UI formatter functions - updated 7/17/2014 (v2.17.5) */
!function(c){"use strict";var p=c.tablesorter||{},f=".compare-select",u=p.filterFormatter=c.extend({},p.filterFormatter,{addCompare:function(e,a,t){var n,d,i;t.compare&&c.isArray(t.compare)&&1<t.compare.length&&(n="",d=[f.slice(1)," "+f.slice(1),""],i=t.cellText?'<label class="'+d.join("-label")+a+'">'+t.cellText+"</label>":"",c.each(t.compare,function(e,a){n+="<option "+(t.selected===e?"selected":"")+">"+a+"</option>"}),e.wrapInner('<div class="'+d.join("-wrapper")+a+'" />').prepend(i+'<select class="'+d.join("")+a+'" />').find("select").append(n))},updateCompare:function(e,a,t){var a=a.val()||"",n=a.replace(/\s*?[><=]\s*?/g,""),d=a.match(/[><=]/g)||"";return t.compare&&(c.isArray(t.compare)&&(d=(d||[]).join("")||t.compare[t.selected||0]),e.find(f).val(d)),[a,n]},uiSpinner:function(l,a,e){var r=c.extend({delayed:!0,addToggle:!0,exactMatch:!0,value:1,cellText:"",compare:"",min:0,max:100,step:1,disabled:!1},e),o=l.closest("table")[0].config,t=c('<input class="filter" type="hidden">').appendTo(l).bind("change"+o.namespace+"filter",function(){n({value:this.value,delayed:!1})}),s=[],n=function(e,a){var t,n=!0,d=e&&e.value&&p.formatFloat((e.value+"").replace(/[><=]/g,""))||l.find(".spinner").val()||r.value,i=(c.isArray(r.compare)?l.find(f).val()||r.compare[r.selected||0]:r.compare)||"",e=e&&"boolean"==typeof e.delayed?e.delayed:!o.$table[0].hasInitialized||r.delayed||"";r.addToggle&&(n=l.find(".toggle").is(":checked")),t=r.disabled||!n?"disable":"enable",p.isEmptyObject(l.find(".spinner").data())||(l.find(".filter").val(n?(i||(r.exactMatch?"=":""))+d:"").trigger(a?"":"search",e).end().find(".spinner").spinner(t).val(d),s.length&&(s.find(".spinner").spinner(t).val(d).end().find(f).val(i),r.addToggle)&&(s.find(".toggle")[0].checked=n))};return r.oldcreate=r.create,r.oldspin=r.spin,r.create=function(e,a){n(),"function"==typeof r.oldcreate&&r.oldcreate(e,a)},r.spin=function(e,a){n(a),"function"==typeof r.oldspin&&r.oldspin(e,a)},r.addToggle&&c('<div class="button"><input id="uispinnerbutton'+a+'" type="checkbox" class="toggle" /><label for="uispinnerbutton'+a+'"></label></div>').appendTo(l).find(".toggle").bind("change",function(){n()}),l.closest("thead").find("th[data-column="+a+"]").addClass("filter-parsed"),c('<input class="spinner spinner'+a+'" />').val(r.value).appendTo(l).spinner(r).bind("change keyup",function(){n()}),o.$table.bind("filterFomatterUpdate"+o.namespace+"filter",function(){var e=u.updateCompare(l,t,r)[0];l.find(".spinner").val(e),n({value:e},!0),p.filter.formatterUpdated(l,a)}),r.compare&&(u.addCompare(l,a,r),l.find(f).bind("change",function(){n()})),o.$table.bind("stickyHeadersInit"+o.namespace+"filter",function(){s=o.widgetOptions.$sticky.find(".tablesorter-filter-row").children().eq(a).empty(),r.addToggle&&c('<div class="button"><input id="stickyuispinnerbutton'+a+'" type="checkbox" class="toggle" /><label for="stickyuispinnerbutton'+a+'"></label></div>').appendTo(s).find(".toggle").bind("change",function(){l.find(".toggle")[0].checked=this.checked,n()}),c('<input class="spinner spinner'+a+'" />').val(r.value).appendTo(s).spinner(r).bind("change keyup",function(){l.find(".spinner").val(this.value),n()}),r.compare&&(u.addCompare(s,a,r),s.find(f).bind("change",function(){l.find(f).val(c(this).val()),n()}))}),o.$table.bind("filterReset"+o.namespace+"filter",function(){c.isArray(r.compare)&&l.add(s).find(f).val(r.compare[r.selected||0]),r.addToggle&&(l.find(".toggle")[0].checked=!1),l.find(".spinner").spinner("value",r.value),setTimeout(function(){n()},0)}),n(),t},uiSlider:function(i,l,e){var r=c.extend({delayed:!0,valueToHeader:!1,exactMatch:!0,cellText:"",compare:"",allText:"all",value:0,min:0,max:100,step:1,range:"min"},e),o=i.closest("table")[0].config,a=c('<input class="filter" type="hidden">').appendTo(i).bind("change"+o.namespace+"filter",function(){t({value:this.value})}),s=[],t=function(e,a){var t=void 0!==e&&p.formatFloat((e.value+"").replace(/[><=]/g,""))||r.value,n=!r.compare&&t===r.min?r.allText:t,d=(c.isArray(r.compare)?i.find(f).val()||r.compare[r.selected||0]:r.compare)||"",n=d+n,e=e&&"boolean"==typeof e.delayed?e.delayed:!o.$table[0].hasInitialized||r.delayed||"";r.valueToHeader?i.closest("thead").find("th[data-column="+l+"]").find(".curvalue").html(" ("+n+")"):i.find(".ui-slider-handle").addClass("value-popup").attr("data-value",n),p.isEmptyObject(i.find(".slider").data())||(i.find(".filter").val(d?d+t:t===r.min?"":(r.exactMatch?"=":"")+t).trigger(a?"":"search",e).end().find(".slider").slider("value",t),s.length&&(s.find(f).val(d).end().find(".slider").slider("value",t),r.valueToHeader?s.closest("thead").find("th[data-column="+l+"]").find(".curvalue").html(" ("+n+")"):s.find(".ui-slider-handle").addClass("value-popup").attr("data-value",n)))};return i.closest("thead").find("th[data-column="+l+"]").addClass("filter-parsed"),r.valueToHeader&&i.closest("thead").find("th[data-column="+l+"]").find(".tablesorter-header-inner").append('<span class="curvalue" />'),r.oldcreate=r.create,r.oldslide=r.slide,r.create=function(e,a){t(),"function"==typeof r.oldcreate&&r.oldcreate(e,a)},r.slide=function(e,a){t(a),"function"==typeof r.oldslide&&r.oldslide(e,a)},c('<div class="slider slider'+l+'"/>').appendTo(i).slider(r),o.$table.bind("filterFomatterUpdate"+o.namespace+"filter",function(){var e=u.updateCompare(i,a,r)[0];i.find(".slider").slider("value",e),t({value:e},!1),p.filter.formatterUpdated(i,l)}),r.compare&&(u.addCompare(i,l,r),i.find(f).bind("change",function(){t({value:i.find(".slider").slider("value")})})),o.$table.bind("filterReset"+o.namespace+"filter",function(){c.isArray(r.compare)&&i.add(s).find(f).val(r.compare[r.selected||0]),setTimeout(function(){t({value:r.value})},0)}),o.$table.bind("stickyHeadersInit"+o.namespace+"filter",function(){s=o.widgetOptions.$sticky.find(".tablesorter-filter-row").children().eq(l).empty(),c('<div class="slider slider'+l+'"/>').val(r.value).appendTo(s).slider(r).bind("change keyup",function(){i.find(".slider").slider("value",this.value),t()}),r.compare&&(u.addCompare(s,l,r),s.find(f).bind("change",function(){i.find(f).val(c(this).val()),t()}))}),a},uiRange:function(i,l,e){var r=c.extend({delayed:!0,valueToHeader:!1,values:[0,100],min:0,max:100,range:!0},e),o=i.closest("table")[0].config,t=c('<input class="filter" type="hidden">').appendTo(i).bind("change"+o.namespace+"filter",function(){a()}),s=[],a=function(){var e=t.val(),a=e.split(" - ");(a=""===e?[r.min,r.max]:a)&&a[1]&&n({values:a,delay:!1},!0)},n=function(e,a){var t=e&&e.values||r.values,n=t[0]+" - "+t[1],d=t[0]===r.min&&t[1]===r.max?"":n,e=e&&"boolean"==typeof e.delayed?e.delayed:!o.$table[0].hasInitialized||r.delayed||"";r.valueToHeader?i.closest("thead").find("th[data-column="+l+"]").find(".currange").html(" ("+n+")"):i.find(".ui-slider-handle").addClass("value-popup").eq(0).attr("data-value",t[0]).end().eq(1).attr("data-value",t[1]),p.isEmptyObject(i.find(".range").data())||(i.find(".filter").val(d).trigger(a?"":"search",e).end().find(".range").slider("values",t),s.length&&(s.find(".range").slider("values",t),r.valueToHeader?s.closest("thead").find("th[data-column="+l+"]").find(".currange").html(" ("+n+")"):s.find(".ui-slider-handle").addClass("value-popup").eq(0).attr("data-value",t[0]).end().eq(1).attr("data-value",t[1])))};return i.closest("thead").find("th[data-column="+l+"]").addClass("filter-parsed"),r.valueToHeader&&i.closest("thead").find("th[data-column="+l+"]").find(".tablesorter-header-inner").append('<span class="currange"/>'),r.oldcreate=r.create,r.oldslide=r.slide,r.create=function(e,a){n(),"function"==typeof r.oldcreate&&r.oldcreate(e,a)},r.slide=function(e,a){n(a),"function"==typeof r.oldslide&&r.oldslide(e,a)},c('<div class="range range'+l+'"/>').appendTo(i).slider(r),o.$table.bind("filterFomatterUpdate"+o.namespace+"filter",function(){a(),p.filter.formatterUpdated(i,l)}),o.$table.bind("filterReset"+o.namespace+"filter",function(){i.find(".range").slider("values",r.values),setTimeout(function(){n()},0)}),o.$table.bind("stickyHeadersInit"+o.namespace+"filter",function(){s=o.widgetOptions.$sticky.find(".tablesorter-filter-row").children().eq(l).empty(),c('<div class="range range'+l+'"/>').val(r.value).appendTo(s).slider(r).bind("change keyup",function(){i.find(".range").val(this.value),n()})}),t},uiDateCompare:function(i,t,e){function n(e){var a,t=s.datepicker("getDate")||"",n=(c.isArray(l.compare)?i.find(f).val()||l.compare[l.selected||0]:l.compare)||"",d=!r.$table[0].hasInitialized||l.delayed||"";s.datepicker("setDate",(""===t?"":t)||null),""===t&&(e=!1),a=(t=s.datepicker("getDate"))&&(l.endOfDay&&/<=/.test(n)?t.setHours(23,59,59,999):t.getTime())||"",t&&l.endOfDay&&"="===n&&(n="",a+=" - "+t.setHours(23,59,59,999),e=!1),i.find(".dateCompare").val(n+a).trigger(e?"":"search",d).end(),o.length&&o.find(".dateCompare").val(n+a).end().find(f).val(n)}var l=c.extend({cellText:"",compare:"",endOfDay:!0,defaultDate:"",changeMonth:!0,changeYear:!0,numberOfMonths:1},e),r=i.closest("table")[0].config,e=i.closest("thead").find("th[data-column="+t+"]").addClass("filter-parsed"),d=c('<input class="dateCompare" type="hidden">').appendTo(i).bind("change"+r.namespace+"filter",function(){var e=this.value;e&&l.onClose(e)}),o=[],a='<input type="text" class="date date'+t+'" placeholder="'+(e.data("placeholder")||e.attr("data-placeholder")||r.widgetOptions.filter_placeholder.search||"")+'" />',s=c(a).appendTo(i);return l.oldonClose=l.onClose,l.onClose=function(e,a){n(),"function"==typeof l.oldonClose&&l.oldonClose(e,a)},s.datepicker(l),r.$table.bind("filterReset"+r.namespace+"filter",function(){c.isArray(l.compare)&&i.add(o).find(f).val(l.compare[l.selected||0]),i.add(o).find(".date").val(l.defaultDate).datepicker("setDate",l.defaultDate||null),setTimeout(function(){n()},0)}),r.$table.bind("filterFomatterUpdate"+r.namespace+"filter",function(){var e,a=d.val();/\s+-\s+/.test(a)?(i.find(f).val("="),e=a.split(/\s+-\s+/)[0],s.datepicker("setDate",e||null)):e=""!==(e=u.updateCompare(i,d,l)[1].toString()||"")?/\d{5}/g.test(e)?new Date(Number(e)):e||"":"",i.add(o).find(".date").datepicker("setDate",e||null),setTimeout(function(){n(!0),p.filter.formatterUpdated(i,t)},0)}),l.compare&&(u.addCompare(i,t,l),i.find(f).bind("change",function(){n()})),r.$table.bind("stickyHeadersInit"+r.namespace+"filter",function(){(o=r.widgetOptions.$sticky.find(".tablesorter-filter-row").children().eq(t).empty()).append(a).find(".date").datepicker(l),l.compare&&(u.addCompare(o,t,l),o.find(f).bind("change",function(){i.find(f).val(c(this).val()),n()}))}),d.val(l.defaultDate||"")},uiDatepicker:function(i,n,e){function l(e){return e instanceof Date&&isFinite(e)}var d,r=c.extend({endOfDay:!0,textFrom:"from",textTo:"to",from:"",to:"",changeMonth:!0,changeYear:!0,numberOfMonths:1},e),o=[],a=i.closest("table")[0].config,s=c('<input class="dateRange" type="hidden">').appendTo(i).bind("change"+a.namespace+"filter",function(){var e=this.value;e.match(" - ")?(e=e.split(" - "),i.find(".dateTo").val(e[1]),d(e[0])):e.match(">=")?d(e.replace(">=","")):e.match("<=")&&d(e.replace("<=",""))}),e=i.closest("thead").find("th[data-column="+n+"]").addClass("filter-parsed"),t="<label>"+r.textFrom+'</label><input type="text" class="dateFrom" placeholder="'+(e.data("placeholderFrom")||e.attr("data-placeholder-from")||a.widgetOptions.filter_placeholder.from||"")+'" /><label>'+r.textTo+'</label><input type="text" class="dateTo" placeholder="'+(e.data("placeholderTo")||e.attr("data-placeholder-to")||a.widgetOptions.filter_placeholder.to||"")+'" />';return c(t).appendTo(i),r.oldonClose=r.onClose,d=r.onClose=function(e,a){var t=i.find(".dateFrom").datepicker("getDate"),n=i.find(".dateTo").datepicker("getDate"),t=l(t)?t.getTime():"",n=l(n)&&(r.endOfDay?n.setHours(23,59,59,999):n.getTime())||"",d=t?n?t+" - "+n:">="+t:n?"<="+n:"";i.add(o).find(".dateRange").val(d).trigger("search"),t=t?new Date(t):"",n=n?new Date(n):"",/<=/.test(d)?i.add(o).find(".dateFrom").datepicker("option","maxDate",n||null).end().find(".dateTo").datepicker("option","minDate",null).datepicker("setDate",n||null):/>=/.test(d)?i.add(o).find(".dateFrom").datepicker("option","maxDate",null).datepicker("setDate",t||null).end().find(".dateTo").datepicker("option","minDate",t||null):i.add(o).find(".dateFrom").datepicker("option","maxDate",null).datepicker("setDate",t||null).end().find(".dateTo").datepicker("option","minDate",null).datepicker("setDate",n||null),"function"==typeof r.oldonClose&&r.oldonClose(e,a)},r.defaultDate=r.from||"",i.find(".dateFrom").datepicker(r),r.defaultDate=r.to||"+7d",i.find(".dateTo").datepicker(r),a.$table.bind("filterFomatterUpdate"+a.namespace+"filter",function(){var e=s.val()||"",a="",t="";/\s+-\s+/.test(e)?(a=(e=e.split(/\s+-\s+/)||[])[0]||"",t=e[1]||""):/>=/.test(e)?a=e.replace(/>=/,"")||"":/<=/.test(e)&&(t=e.replace(/<=/,"")||""),a=""!==a?/\d{5}/g.test(a)?new Date(Number(a)):a||"":"",t=""!==t?/\d{5}/g.test(t)?new Date(Number(t)):t||"":"",i.add(o).find(".dateFrom").datepicker("setDate",a||null),i.add(o).find(".dateTo").datepicker("setDate",t||null),setTimeout(function(){d(),p.filter.formatterUpdated(i,n)},0)}),a.$table.bind("stickyHeadersInit"+a.namespace+"filter",function(){(o=a.widgetOptions.$sticky.find(".tablesorter-filter-row").children().eq(n).empty()).append(t),r.defaultDate=r.from||"",o.find(".dateFrom").datepicker(r),r.defaultDate=r.to||"+7d",o.find(".dateTo").datepicker(r)}),i.closest("table").bind("filterReset"+a.namespace+"filter",function(){i.add(o).find(".dateFrom").val("").datepicker("setDate",r.from||null),i.add(o).find(".dateTo").val("").datepicker("setDate",r.to||null),setTimeout(function(){d()},0)}),s.val(r.from?r.to?r.from+" - "+r.to:">="+r.from:r.to?"<="+r.to:"")}})}(jQuery);return jQuery;}));
```
