## Supplementary Data SD1 for "Inferring binding specificities of human transcription factors with the wisdom of crowds": widget-filter-formatter-select2.min.js

```
(function(factory){if (typeof define === 'function' && define.amd){define(['jquery'], factory);} else if (typeof module === 'object' && typeof module.exports === 'object'){module.exports = factory(require('jquery'));} else {factory(jQuery);}}(function(jQuery){
/*! Widget: filter, select2 formatter function - updated 12/1/2019 (v2.31.2) */
!function(v){"use strict";var g=v.tablesorter||{};g.filterFormatter=g.filterFormatter||{},g.filterFormatter.select2=function(i,c,e){function t(){a=[],l=g.filter.getOptionSource(s.$table[0],c,o)||[],v.each(l,function(e,t){a.push({id:""+t.parsed,text:t.text})}),n.data=a}var l,a,n=v.extend({cellText:"",match:!0,value:"",multiple:!0,width:"100%"},e),s=i.addClass("select2col"+c).closest("table")[0].config,e=s.widgetOptions,d=v('<input class="filter" type="hidden">').appendTo(i).bind("change"+s.namespace+"filter",function(){var e=b(this.value);s.$table.find(".select2col"+c+" .select2").select2("val",e),$()}),r=s.$headerIndexed[c],o=r.hasClass(e.filter_onlyAvail),f=n.match?"":"^",p=n.match?"":"$",u=e.filter_ignoreCase?"i":"",b=function(e){return e.replace(/^\/\(\^?/,"").replace(/\$\|\^/g,"|").replace(/\$?\)\/i?$/g,"").replace(/\\/g,"").split("|")},$=function(){var e=!1,t=s.$table.find(".select2col"+c+" .select2").select2("val")||n.value||"",l=(v.isArray(t)&&(e=!0,t=t.join("\0")),t.replace(/[-[\]{}()*+?.,/\\^$|#]/g,"\\$&"));e&&(t=t.split("\0"),l=l.split("\0")),g.isEmptyObject(i.find(".select2").data())||(d.val(v.isArray(l)&&l.length&&""!==l.join("")?"/("+f+(l||[]).join(p+"|"+f)+p+")/"+u:"").trigger("search"),i.find(".select2").select2("val",t),s.widgetOptions.$sticky&&s.widgetOptions.$sticky.find(".select2col"+c+" .select2").select2("val",t))};return r.toggleClass("filter-match",n.match),n.cellText&&i.prepend("<label>"+n.cellText+"</label>"),n.ajax&&!v.isEmptyObject(n.ajax)||n.data||(t(),s.$table.bind("filterEnd",function(){t(),s.$table.find(".select2col"+c).add(s.widgetOptions.$sticky&&s.widgetOptions.$sticky.find(".select2col"+c)).find(".select2").select2(n)})),v('<input class="select2 select2-'+c+'" type="hidden" />').val(n.value).appendTo(i).select2(n).bind("change",function(){$()}),s.$table.bind("filterFomatterUpdate",function(){var e=b(s.$table.data("lastSearch")[c]||"");(i=s.$table.find(".select2col"+c)).find(".select2").select2("val",e),$(),g.filter.formatterUpdated(i,c)}),s.$table.bind("stickyHeadersInit",function(){var e=s.widgetOptions.$sticky.find(".select2col"+c).empty();v('<input class="select2 select2-'+c+'" type="hidden">').val(n.value).appendTo(e).select2(n).bind("change",function(){s.$table.find(".select2col"+c).find(".select2").select2("val",s.widgetOptions.$sticky.find(".select2col"+c+" .select2").select2("val")),$()}),n.cellText&&e.prepend("<label>"+n.cellText+"</label>")}),s.$table.bind("filterReset",function(){s.$table.find(".select2col"+c).find(".select2").select2("val",n.value||""),setTimeout(function(){$()},0)}),$(),d}}(jQuery);return jQuery;}));
```
