## Supplementary Data SD1 for "Inferring binding specificities of human transcription factors with the wisdom of crowds": widget-filter-type-insideRange.min.js

```
(function(factory){if (typeof define === 'function' && define.amd){define(['jquery'], factory);} else if (typeof module === 'object' && typeof module.exports === 'object'){module.exports = factory(require('jquery'));} else {factory(jQuery);}}(function(jQuery){
/*! Widget: filter, insideRange filter type - updated 12/10/2015 (v2.25.0) */
!function(){"use strict";function c(t){return isNaN(t)?t:parseFloat(t)}var t=jQuery.tablesorter,u=/\d+/,o=/\s+-\s+/;t.filter.types.insideRange=function(t,e){var i,r,n,a,s,l;return e.anyMatch||!u.test(e.iFilter)||!o.test(e.iExact)||(n=e.index,a=e.$cells[n],s=e.iExact.split(o),l=t.parsers[e.index]&&t.parsers[e.index].format,s&&s.length<2)||"function"!=typeof l?null:(r=c(l(s[0],t.table,a,n)),(s=c(l(s[1],t.table,a,n)))<r&&(i=s,s=r,r=i),r<=(i=c(l(e.iFilter,t.table,a,n)))&&i<=s)}}();return jQuery;}));
```
