## Supplementary Data SD1 for "Inferring binding specificities of human transcription factors with the wisdom of crowds": widget-filter.min.js

```
(function(factory){if (typeof define === 'function' && define.amd){define(['jquery'], factory);} else if (typeof module === 'object' && typeof module.exports === 'object'){module.exports = factory(require('jquery'));} else {factory(jQuery);}}(function(jQuery){
/*! Widget: filter - updated 2018-03-18 (v2.30.0) */
!function(M){"use strict";var I,D,z=M.tablesorter||{},g=z.css,o=z.keyCodes;M.extend(g,{filterRow:"tablesorter-filter-row",filter:"tablesorter-filter",filterDisabled:"disabled",filterRowHide:"hideme"}),M.extend(o,{backSpace:8,escape:27,space:32,left:37,down:40}),z.addWidget({id:"filter",priority:50,options:{filter_cellFilter:"",filter_childRows:!1,filter_childByColumn:!1,filter_childWithSibs:!0,filter_columnAnyMatch:!0,filter_columnFilters:!0,filter_cssFilter:"",filter_defaultAttrib:"data-value",filter_defaultFilter:{},filter_excludeFilter:{},filter_external:"",filter_filteredRow:"filtered",filter_filterLabel:'Filter "{{label}}" column by...',filter_formatter:null,filter_functions:null,filter_hideEmpty:!0,filter_hideFilters:!1,filter_ignoreCase:!0,filter_liveSearch:!0,filter_matchType:{input:"exact",select:"exact"},filter_onlyAvail:"filter-onlyAvail",filter_placeholder:{search:"",select:""},filter_reset:null,filter_resetOnEsc:!0,filter_saveFilters:!1,filter_searchDelay:300,filter_searchFiltered:!0,filter_selectSource:null,filter_selectSourceSeparator:"|",filter_serversideFiltering:!1,filter_startsWith:!1,filter_useParsedData:!1},format:function(e,t,r){t.$table.hasClass("hasFilters")||I.init(e,t,r)},remove:function(e,t,r,i){var l,a,n=t.$table,s=t.$tbodies,o="addRows updateCell update updateRows updateComplete appendCache filterReset filterAndSortReset filterFomatterUpdate filterEnd search stickyHeadersInit ".split(" ").join(t.namespace+"filter ");if(n.removeClass("hasFilters").unbind(o.replace(z.regex.spaces," ")).find("."+g.filterRow).remove(),r.filter_initialized=!1,!i){for(l=0;l<s.length;l++)(a=z.processTbody(e,s.eq(l),!0)).children().removeClass(r.filter_filteredRow).show(),z.processTbody(e,a,!1);r.filter_reset&&M(document).undelegate(r.filter_reset,"click"+t.namespace+"filter")}}}),I=z.filter={regex:{regex:/^\/((?:\\\/|[^\/])+)\/([migyu]{0,5})?$/,child:/tablesorter-childRow/,filtered:/filtered/,type:/undefined|number/,exact:/(^[\"\'=]+)|([\"\'=]+$)/g,operators:/[<>=]/g,query:"(q|query)",wild01:/\?/g,wild0More:/\*/g,quote:/\"/g,isNeg1:/(>=?\s*-\d)/,isNeg2:/(<=?\s*\d)/},types:{or:function(e,t,r){if(!D.orTest.test(t.iFilter)&&!D.orSplit.test(t.filter)||D.regex.test(t.filter))return null;for(var i,l,a=M.extend({},t),n=t.filter.split(D.orSplit),s=t.iFilter.split(D.orSplit),o=n.length,c=0;c<o;c++){a.nestedFilters=!0,a.filter=""+(I.parseFilter(e,n[c],t)||""),a.iFilter=""+(I.parseFilter(e,s[c],t)||""),l="("+(I.parseFilter(e,a.filter,t)||"")+")";try{if(i=new RegExp(t.isMatch?l:"^"+l+"$",e.widgetOptions.filter_ignoreCase?"i":"").test(a.exact)||I.processTypes(e,a,r))return i}catch(e){return null}}return i||!1},and:function(e,t,r){if(D.andTest.test(t.filter)){for(var i,l,a,n=M.extend({},t),s=t.filter.split(D.andSplit),o=t.iFilter.split(D.andSplit),c=s.length,f=0;f<c;f++){n.nestedFilters=!0,n.filter=""+(I.parseFilter(e,s[f],t)||""),n.iFilter=""+(I.parseFilter(e,o[f],t)||""),a=("("+(I.parseFilter(e,n.filter,t)||"")+")").replace(D.wild01,"\\S{1}").replace(D.wild0More,"\\S*");try{l=new RegExp(t.isMatch?a:"^"+a+"$",e.widgetOptions.filter_ignoreCase?"i":"").test(n.exact)||I.processTypes(e,n,r),i=(0===f||i)&&l}catch(e){return null}}return i||!1}return null},regex:function(e,t){if(D.regex.test(t.filter)){var r,i=t.filter_regexCache[t.index]||D.regex.exec(t.filter),l=i instanceof RegExp;try{l||(t.filter_regexCache[t.index]=i=new RegExp(i[1],i[2])),r=i.test(t.exact)}catch(e){r=!1}return r}return null},operators:function(e,t){var r,i,l,a,n,s,o;return D.operTest.test(t.iFilter)&&""!==t.iExact?(l=e.table,a=t.parsed[t.index],n=z.formatFloat(t.iFilter.replace(D.operators,""),l),s=e.parsers[t.index]||{},o=n,!a&&"numeric"!==s.type||(i=M.trim(""+t.iFilter.replace(D.operators,"")),n="number"!=typeof(r=I.parseFilter(e,i,t,!0))||""===r||isNaN(r)?n:r),e=!a&&"numeric"!==s.type||isNaN(n)||void 0===t.cache?(i=isNaN(t.iExact)?t.iExact.replace(z.regex.nondigit,""):t.iExact,z.formatFloat(i,l)):t.cache,D.gtTest.test(t.iFilter)?r=D.gteTest.test(t.iFilter)?n<=e:n<e:D.ltTest.test(t.iFilter)&&(r=D.lteTest.test(t.iFilter)?e<=n:e<n),!r&&""===o||r):null},notMatch:function(e,t){var r,i;return D.notTest.test(t.iFilter)?(i=t.iFilter.replace("!",""),i=I.parseFilter(e,i,t)||"",D.exact.test(i)?""===(i=i.replace(D.exact,""))||M.trim(i)!==t.iExact:(r=t.iExact.search(M.trim(i)),""===i||(t.anyMatch?r<0:!(e.widgetOptions.filter_startsWith?0===r:0<=r)))):null},exact:function(e,t){var r;return D.exact.test(t.iFilter)?(r=t.iFilter.replace(D.exact,""),e=I.parseFilter(e,r,t)||"",t.anyMatch?0<=M.inArray(e,t.rowArray):e==t.iExact):null},range:function(e,t){var r,i,l,a,n,s,o,c;return D.toTest.test(t.iFilter)?(l=e.table,a=t.index,n=t.parsed[a],o=(s=t.iFilter.split(D.toSplit))[0].replace(z.regex.nondigit,"")||"",c=z.formatFloat(I.parseFilter(e,o,t),l),o=s[1].replace(z.regex.nondigit,"")||"",i=z.formatFloat(I.parseFilter(e,o,t),l),!n&&"numeric"!==e.parsers[a].type||(c=""===(r=e.parsers[a].format(""+s[0],l,e.$headers.eq(a),a))||isNaN(r)?c:r,i=""===(r=e.parsers[a].format(""+s[1],l,e.$headers.eq(a),a))||isNaN(r)?i:r),r=!n&&"numeric"!==e.parsers[a].type||isNaN(c)||isNaN(i)?(o=isNaN(t.iExact)?t.iExact.replace(z.regex.nondigit,""):t.iExact,z.formatFloat(o,l)):t.cache,i<c&&(o=c,c=i,i=o),c<=r&&r<=i||""===c||""===i):null},wild:function(e,t){if(D.wildOrTest.test(t.iFilter)){var r=""+(I.parseFilter(e,t.iFilter,t)||"");!D.wildTest.test(r)&&t.nestedFilters&&(r=t.isMatch?r:"^("+r+")$");try{return new RegExp(r.replace(D.wild01,"\\S{1}").replace(D.wild0More,"\\S*"),e.widgetOptions.filter_ignoreCase?"i":"").test(t.exact)}catch(e){}}return null},fuzzy:function(e,t){if(D.fuzzyTest.test(t.iFilter)){for(var r=0,i=t.iExact.length,l=t.iFilter.slice(1),a=I.parseFilter(e,l,t)||"",n=0;n<i;n++)t.iExact[n]===a[r]&&(r+=1);return r===a.length}return null}},init:function(r){z.language=M.extend(!0,{},{to:"to",or:"or",and:"and"},z.language);function e(e,t,r){return""===(t=t.trim())?"":(e||"")+t+(r||"")}var t,i,l,a,n,s,o,c,f=r.config,d=f.widgetOptions;if(f.$table.addClass("hasFilters"),f.lastSearch=[],d.filter_searchTimer=null,d.filter_initTimer=null,d.filter_formatterCount=0,d.filter_formatterInit=[],d.filter_anyColumnSelector='[data-column="all"],[data-column="any"]',d.filter_multipleColumnSelector='[data-column*="-"],[data-column*=","]',s="\\{"+D.query+"\\}",M.extend(D,{child:new RegExp(f.cssChildRow),filtered:new RegExp(d.filter_filteredRow),alreadyFiltered:new RegExp("(\\s+(-"+e("|",z.language.or)+e("|",z.language.to)+")\\s+)","i"),toTest:new RegExp("\\s+(-"+e("|",z.language.to)+")\\s+","i"),toSplit:new RegExp("(?:\\s+(?:-"+e("|",z.language.to)+")\\s+)","gi"),andTest:new RegExp("\\s+("+e("",z.language.and,"|")+"&&)\\s+","i"),andSplit:new RegExp("(?:\\s+(?:"+e("",z.language.and,"|")+"&&)\\s+)","gi"),orTest:new RegExp("(\\|"+e("|\\s+",z.language.or,"\\s+")+")","i"),orSplit:new RegExp("(?:\\|"+e("|\\s+(?:",z.language.or,")\\s+")+")","gi"),iQuery:new RegExp(s,"i"),igQuery:new RegExp(s,"ig"),operTest:/^[<>]=?/,gtTest:/>/,gteTest:/>=/,ltTest:/</,lteTest:/<=/,notTest:/^\!/,wildOrTest:/[\?\*\|]/,wildTest:/\?\*/,fuzzyTest:/^~/,exactTest:/[=\"\|!]/}),s=f.$headers.filter(".filter-false, .parser-false").length,!1!==d.filter_columnFilters&&s!==f.$headers.length&&I.buildRow(r,f,d),l="addRows updateCell update updateRows updateComplete appendCache filterReset "+"filterAndSortReset filterResetSaved filterEnd search ".split(" ").join(f.namespace+"filter "),f.$table.bind(l,function(e,t){return s=d.filter_hideEmpty&&M.isEmptyObject(f.cache)&&!(f.delayInit&&"appendCache"===e.type),f.$table.find("."+g.filterRow).toggleClass(d.filter_filteredRow,s),/(search|filter)/.test(e.type)||(e.stopPropagation(),I.buildDefault(r,!0)),"filterReset"===e.type||"filterAndSortReset"===e.type?(f.$table.find("."+g.filter).add(d.filter_$externalFilters).val(""),"filterAndSortReset"===e.type?z.sortReset(this.config,function(){I.searching(r,[])}):I.searching(r,[])):"filterResetSaved"===e.type?z.storage(r,"tablesorter-filters",""):"filterEnd"===e.type?I.buildDefault(r,!0):(t="search"===e.type?t:"updateComplete"===e.type?f.$table.data("lastSearch"):"",/(update|add)/.test(e.type)&&"updateComplete"!==e.type&&(f.lastCombinedFilter=null,f.lastSearch=[],setTimeout(function(){f.$table.triggerHandler("filterFomatterUpdate")},100)),I.searching(r,t,!0)),!1}),d.filter_reset&&(d.filter_reset instanceof M?d.filter_reset.click(function(){f.$table.triggerHandler("filterReset")}):M(d.filter_reset).length&&M(document).undelegate(d.filter_reset,"click"+f.namespace+"filter").delegate(d.filter_reset,"click"+f.namespace+"filter",function(){f.$table.triggerHandler("filterReset")})),d.filter_functions)for(n=0;n<f.columns;n++)if(o=z.getColumnData(r,d.filter_functions,n))if(c=!((a=f.$headerIndexed[n].removeClass("filter-select")).hasClass("filter-false")||a.hasClass("parser-false")),!(t="")===o&&c)I.buildSelect(r,n);else if("object"==typeof o&&c){for(i in o)"string"==typeof i&&(t+=""===t?'<option value="">'+(a.data("placeholder")||a.attr("data-placeholder")||d.filter_placeholder.select||"")+"</option>":"",0<=(l=s=i).indexOf(d.filter_selectSourceSeparator)&&(l=(s=i.split(d.filter_selectSourceSeparator))[1],s=s[0]),t+="<option "+(l===s?"":'data-function-name="'+i+'" ')+'value="'+s+'">'+l+"</option>");f.$table.find("thead").find("select."+g.filter+'[data-column="'+n+'"]').append(t),("function"==typeof(l=d.filter_selectSource)||z.getColumnData(r,l,n))&&I.buildSelect(f.table,n,"",!0,a.hasClass(d.filter_onlyAvail))}I.buildDefault(r,!0),I.bindSearch(r,f.$table.find("."+g.filter),!0),d.filter_external&&I.bindSearch(r,d.filter_external),d.filter_hideFilters&&I.hideFilters(f),f.showProcessing&&(l="filterStart filterEnd ".split(" ").join(f.namespace+"filter-sp "),f.$table.unbind(l.replace(z.regex.spaces," ")).bind(l,function(e,t){a=t?f.$table.find("."+g.header).filter("[data-column]").filter(function(){return""!==t[M(this).data("column")]}):"",z.isProcessing(r,"filterStart"===e.type,t?a:"")})),f.filteredRows=f.totalRows,l="tablesorter-initialized pagerBeforeInitialized ".split(" ").join(f.namespace+"filter "),f.$table.unbind(l.replace(z.regex.spaces," ")).bind(l,function(){I.completeInit(this)}),f.pager&&f.pager.initialized&&!d.filter_initialized?(f.$table.triggerHandler("filterFomatterUpdate"),setTimeout(function(){I.filterInitComplete(f)},100)):d.filter_initialized||I.completeInit(r)},completeInit:function(e){var t=e.config,r=t.widgetOptions,i=I.setDefaults(e,t,r)||[];!i.length||t.delayInit&&""===i.join("")||z.setFilters(e,i,!0),t.$table.triggerHandler("filterFomatterUpdate"),setTimeout(function(){r.filter_initialized||I.filterInitComplete(t)},100)},formatterUpdated:function(e,t){e=e&&e.closest("table"),e=e.length&&e[0].config,e=e&&e.widgetOptions;e&&!e.filter_initialized&&(e.filter_formatterInit[t]=1)},filterInitComplete:function(e){function t(){l.filter_initialized=!0,e.lastSearch=e.$table.data("lastSearch"),e.$table.triggerHandler("filterInit",e),I.findRows(e.table,e.lastSearch||[]),z.debug(e,"filter")&&console.log("Filter >> Widget initialized")}var r,i,l=e.widgetOptions,a=0;if(M.isEmptyObject(l.filter_formatter))t();else{for(i=l.filter_formatterInit.length,r=0;r<i;r++)1===l.filter_formatterInit[r]&&a++;clearTimeout(l.filter_initTimer),l.filter_initialized||a!==l.filter_formatterCount?l.filter_initialized||(l.filter_initTimer=setTimeout(function(){t()},500)):t()}},processFilters:function(e,t){for(var r=[],i=t?encodeURIComponent:decodeURIComponent,l=e.length,a=0;a<l;a++)e[a]&&(r[a]=i(e[a]));return r},setDefaults:function(e,t,r){var i,l,a,n,s=z.getFilters(e)||[];if(r.filter_saveFilters&&z.storage&&(e=z.storage(e,"tablesorter-filters")||[],(i=M.isArray(e))&&""===e.join("")||!i||(s=I.processFilters(e))),""===s.join(""))for(n=t.$headers.add(r.filter_$externalFilters).filter("["+r.filter_defaultAttrib+"]"),l=0;l<=t.columns;l++)a=l===t.columns?"all":l,s[l]=n.filter('[data-column="'+a+'"]').attr(r.filter_defaultAttrib)||s[l]||"";return t.$table.data("lastSearch",s),s},parseFilter:function(e,t,r,i){return i||r.parsed[r.index]?e.parsers[r.index].format(t,e.table,[],r.index):t},buildRow:function(e,t,r){for(var i,l,a,n,s,o,c,f=r.filter_cellFilter,d=t.columns,u=M.isArray(f),h='<tr role="search" class="'+g.filterRow+" "+t.cssIgnoreRow+'">',p=0;p<d;p++)t.$headerIndexed[p].length&&(h=(h+=1<(c=t.$headerIndexed[p]&&t.$headerIndexed[p][0].colSpan||0)?'<td data-column="'+p+"-"+(p+c-1)+'" colspan="'+c+'"':'<td data-column="'+p+'"')+(u?f[p]?' class="'+f[p]+'"':"":""!==f?' class="'+f+'"':"")+"></td>");for(t.$filters=M(h+="</tr>").appendTo(t.$table.children("thead").eq(0)).children("td"),p=0;p<d;p++)(l=t.$headerIndexed[p])&&l.length&&(i=I.getColumnElm(t,t.$filters,p),o=z.getColumnData(e,r.filter_functions,p),a=r.filter_functions&&o&&"function"!=typeof o||l.hasClass("filter-select"),n=z.getColumnData(e,t.headers,p),n="false"===z.getData(l[0],n,"filter")||"false"===z.getData(l[0],n,"parser"),a?h=M("<select>").appendTo(i):((o=z.getColumnData(e,r.filter_formatter,p))?(r.filter_formatterCount++,(h=(h=o(i,p))&&0===h.length?i.children("input"):h)&&(0===h.parent().length||h.parent().length&&h.parent()[0]!==i[0])&&i.append(h)):h=M('<input type="search">').appendTo(i),h&&(c=l.data("placeholder")||l.attr("data-placeholder")||r.filter_placeholder.search||"",h.attr("placeholder",c))),h)&&(s=(M.isArray(r.filter_cssFilter)?void 0!==r.filter_cssFilter[p]&&r.filter_cssFilter[p]||"":r.filter_cssFilter)||"",h.addClass(g.filter+" "+s),c=(s=r.filter_filterLabel).match(/{{([^}]+?)}}/g),M.each(c=c||["{{label}}"],function(e,t){var r=new RegExp(t,"g"),t=l.attr("data-"+t.replace(/{{|}}/g,"")),t=void 0===t?l.text():t;s=s.replace(r,M.trim(t))}),h.attr({"data-column":i.attr("data-column"),"aria-label":s}),n)&&(h.attr("placeholder","").addClass(g.filterDisabled)[0].disabled=!0)},bindSearch:function(l,e,t){var r,a,n,i,s;l=M(l)[0],(e=M(e)).length&&(a=l.config,n=a.widgetOptions,i=a.namespace+"filter",s=n.filter_$externalFilters,!0!==t&&(r=n.filter_anyColumnSelector+","+n.filter_multipleColumnSelector,n.filter_$anyMatch=e.filter(r),s&&s.length?n.filter_$externalFilters=n.filter_$externalFilters.add(e):n.filter_$externalFilters=e,z.setFilters(l,a.$table.data("lastSearch")||[],!1===t)),r="keypress keyup keydown search change input ".split(" ").join(i+" "),e.attr("data-lastSearchTime",(new Date).getTime()).unbind(r.replace(z.regex.spaces," ")).bind("keydown"+i,function(e){if(e.which===o.escape&&!l.config.widgetOptions.filter_resetOnEsc)return!1}).bind("keyup"+i,function(e){n=l.config.widgetOptions;var t=parseInt(M(this).attr("data-column"),10),r="boolean"==typeof n.filter_liveSearch?n.filter_liveSearch:z.getColumnData(l,n.filter_liveSearch,t);if(void 0===r&&(r=n.filter_liveSearch.fallback||!1),M(this).attr("data-lastSearchTime",(new Date).getTime()),e.which===o.escape)this.value=n.filter_resetOnEsc?"":a.lastSearch[t];else{if(""!==this.value&&("number"==typeof r&&this.value.length<r||e.which!==o.enter&&e.which!==o.backSpace&&(e.which<o.space||e.which>=o.left&&e.which<=o.down)))return;if(!1===r&&""!==this.value&&e.which!==o.enter)return}I.searching(l,!0,!0,t)}).bind("search change keypress input blur ".split(" ").join(i+" "),function(e){var t=parseInt(M(this).attr("data-column"),10),r=e.type,i="boolean"==typeof n.filter_liveSearch?n.filter_liveSearch:z.getColumnData(l,n.filter_liveSearch,t);!l.config.widgetOptions.filter_initialized||e.which!==o.enter&&"search"!==r&&"blur"!==r&&("change"!==r&&"input"!==r||!0!==i&&(!0===i||"INPUT"===e.target.nodeName)||this.value===a.lastSearch[t])||(e.preventDefault(),M(this).attr("data-lastSearchTime",(new Date).getTime()),I.searching(l,"keypress"!==r||e.which===o.enter,!0,t))}))},searching:function(e,t,r,i){var l,a=e.config.widgetOptions;void 0===i?l=!1:void 0===(l="boolean"==typeof a.filter_liveSearch?a.filter_liveSearch:z.getColumnData(e,a.filter_liveSearch,i))&&(l=a.filter_liveSearch.fallback||!1),clearTimeout(a.filter_searchTimer),void 0===t||!0===t?a.filter_searchTimer=setTimeout(function(){I.checkFilters(e,t,r)},l?a.filter_searchDelay:10):I.checkFilters(e,t,r)},equalFilters:function(e,t,r){var i,l=[],a=[],n=e.columns+1;for(t=M.isArray(t)?t:[],r=M.isArray(r)?r:[],i=0;i<n;i++)l[i]=t[i]||"",a[i]=r[i]||"";return l.join(",")===a.join(",")},checkFilters:function(e,t,r){var i=e.config,l=i.widgetOptions,a=M.isArray(t),n=a?t:z.getFilters(e,!0),s=n||[];if(M.isEmptyObject(i.cache))i.delayInit&&(!i.pager||i.pager&&i.pager.initialized)&&z.updateCache(i,function(){I.checkFilters(e,!1,r)});else{if(a&&(z.setFilters(e,n,!1,!0!==r),l.filter_initialized||(i.lastSearch=[],i.lastCombinedFilter="")),l.filter_hideFilters&&i.$table.find("."+g.filterRow).triggerHandler(I.hideFiltersCheck(i)?"mouseleave":"mouseenter"),I.equalFilters(i,i.lastSearch,s)){if(!1!==t)return;i.lastCombinedFilter="",i.lastSearch=[]}if(n=n||[],n=Array.prototype.map?n.map(String):n.join("�").split("�"),l.filter_initialized&&i.$table.triggerHandler("filterStart",[n]),!i.showProcessing)return I.findRows(e,n,s),!1;setTimeout(function(){return I.findRows(e,n,s),!1},30)}},hideFiltersCheck:function(e){if("function"==typeof e.widgetOptions.filter_hideFilters){var t=e.widgetOptions.filter_hideFilters(e);if("boolean"==typeof t)return t}return""===z.getFilters(e.$table).join("")},hideFilters:function(i,e){var l;(e||i.$table).find("."+g.filterRow).addClass(g.filterRowHide).bind("mouseenter mouseleave",function(e){var t=e,r=M(this);clearTimeout(l),l=setTimeout(function(){/enter|over/.test(t.type)?r.removeClass(g.filterRowHide):M(document.activeElement).closest("tr")[0]!==r[0]&&r.toggleClass(g.filterRowHide,I.hideFiltersCheck(i))},200)}).find("input, select").bind("focus blur",function(e){var t=e,r=M(this).closest("tr");clearTimeout(l),l=setTimeout(function(){clearTimeout(l),r.toggleClass(g.filterRowHide,I.hideFiltersCheck(i)&&"focus"!==t.type)},200)})},defaultFilter:function(e,t){if(""===e)return e;var r=D.iQuery,i=t.match(D.igQuery).length,l=1<i?M.trim(e).split(/\s/):[M.trim(e)],a=l.length-1,n=0,s=t;for(a<1&&1<i&&(l[1]=l[0]);r.test(s);)s=s.replace(r,l[n++]||""),r.test(s)&&n<a&&""!==(l[n]||"")&&(s=t.replace(r,s));return s},getLatestSearch:function(e){return e?e.sort(function(e,t){return M(t).attr("data-lastSearchTime")-M(e).attr("data-lastSearchTime")}):e||M()},findRange:function(e,t,r){var i,l,a,n,s,o,c,f,d=[];if(/^[0-9]+$/.test(t))return[parseInt(t,10)];if(!r&&/-/.test(t))for(f=(i=t.match(/(\d+)\s*-\s*(\d+)/g))?i.length:0,c=0;c<f;c++){for(l=i[c].split(/\s*-\s*/),a=parseInt(l[0],10)||0,(n=parseInt(l[1],10)||e.columns-1)<a&&(l=a,a=n,n=l),n>=e.columns&&(n=e.columns-1);a<=n;a++)d[d.length]=a;t=t.replace(i[c],"")}if(!r&&/,/.test(t))for(f=(s=t.split(/\s*,\s*/)).length,o=0;o<f;o++)""!==s[o]&&(c=parseInt(s[o],10))<e.columns&&(d[d.length]=c);if(!d.length)for(c=0;c<e.columns;c++)d[d.length]=c;return d},getColumnElm:function(t,e,r){return e.filter(function(){var e=I.findRange(t,M(this).attr("data-column"));return-1<M.inArray(r,e)})},multipleColumns:function(e,t){var r=e.widgetOptions,r=r.filter_initialized||!t.filter(r.filter_anyColumnSelector).length,t=M.trim(I.getLatestSearch(t).attr("data-column")||"");return I.findRange(e,t,!r)},processTypes:function(e,t,r){var i,l=null,a=null;for(i in I.types)M.inArray(i,r.excludeMatch)<0&&null===a&&null!==(a=I.types[i](e,t,r))&&(t.matchedOn=i,l=a);return l},matchType:function(e,t){var r=e.widgetOptions,i=e.$headerIndexed[t],r=!(i.hasClass("filter-exact")||!i.hasClass("filter-match")&&(r.filter_columnFilters?i=e.$filters.find("."+g.filter).add(r.filter_$externalFilters).filter('[data-column="'+t+'"]'):r.filter_$externalFilters&&(i=r.filter_$externalFilters.filter('[data-column="'+t+'"]')),!i.length||"match"!==e.widgetOptions.filter_matchType[(i[0].nodeName||"").toLowerCase()]));return r},processRow:function(t,r,e){var i,l,a,n,s,o=t.widgetOptions,c=!0,f=o.filter_$anyMatch&&o.filter_$anyMatch.length,d=o.filter_$anyMatch&&o.filter_$anyMatch.length?I.multipleColumns(t,o.filter_$anyMatch):[];if(r.$cells=r.$row.children(),r.matchedOn=null,r.anyMatchFlag&&1<d.length||r.anyMatchFilter&&!f){if(r.anyMatch=!0,r.isMatch=!0,r.rowArray=r.$cells.map(function(e){if(-1<M.inArray(e,d)||r.anyMatchFilter&&!f)return r.parsed[e]?s=r.cacheArray[e]:(s=r.rawArray[e],s=M.trim(o.filter_ignoreCase?s.toLowerCase():s),t.sortLocaleCompare&&(s=z.replaceAccents(s))),s}).get(),r.filter=r.anyMatchFilter,r.iFilter=r.iAnyMatchFilter,r.exact=r.rowArray.join(" "),r.iExact=o.filter_ignoreCase?r.exact.toLowerCase():r.exact,r.cache=r.cacheArray.slice(0,-1).join(" "),e.excludeMatch=e.noAnyMatch,null!==(l=I.processTypes(t,r,e)))c=l;else if(o.filter_startsWith)for(c=!1,d=Math.min(t.columns,r.rowArray.length);!c&&0<d;)d--,c=c||0===r.rowArray[d].indexOf(r.iFilter);else c=0<=(r.iExact+r.childRowText).indexOf(r.iFilter);if(r.anyMatch=!1,r.filters.join("")===r.filter)return c}for(d=0;d<t.columns;d++)r.filter=r.filters[d],r.index=d,e.excludeMatch=e.excludeFilter[d],r.filter&&(r.cache=r.cacheArray[d],i=r.parsed[d]?r.cache:r.rawArray[d]||"",r.exact=t.sortLocaleCompare?z.replaceAccents(i):i,r.iExact=!D.type.test(typeof r.exact)&&o.filter_ignoreCase?r.exact.toLowerCase():r.exact,r.isMatch=I.matchType(t,d),i=c,n=o.filter_columnFilters&&t.$filters.add(o.filter_$externalFilters).filter('[data-column="'+d+'"]').find("select option:selected").attr("data-function-name")||"",t.sortLocaleCompare&&(r.filter=z.replaceAccents(r.filter)),o.filter_defaultFilter&&D.iQuery.test(e.defaultColFilter[d])&&(r.filter=I.defaultFilter(r.filter,e.defaultColFilter[d])),r.iFilter=o.filter_ignoreCase?(r.filter||"").toLowerCase():r.filter,l=null,(a=e.functions[d])&&("function"==typeof a?l=a(r.exact,r.cache,r.filter,d,r.$row,t,r):"function"==typeof a[n||r.filter]&&(l=a[s=n||r.filter](r.exact,r.cache,r.filter,d,r.$row,t,r))),c=!!(i=null===l&&(l=I.processTypes(t,r,e),s=!0===a&&("and"===r.matchedOn||"or"===r.matchedOn),null===l||s)?!0===a?r.isMatch?0<=(""+r.iExact).search(r.iFilter):r.filter===r.exact:(s=(r.iExact+r.childRowText).indexOf(I.parseFilter(t,r.iFilter,r)),!o.filter_startsWith&&0<=s||o.filter_startsWith&&0===s):l)&&c);return c},findRows:function(e,r,t){if(!I.equalFilters(e.config,e.config.lastSearch,t)&&e.config.widgetOptions.filter_initialized){var i,l,a,n,s,o,c,f,d,u,h,p,g,m,_,y,F,w,x,b,C,v,$,S=M.extend([],r),R=e.config,T=R.widgetOptions,A=z.debug(R,"filter"),O={anyMatch:!1,filters:r,filter_regexCache:[]},E={noAnyMatch:["range","operators"],functions:[],excludeFilter:[],defaultColFilter:[],defaultAnyFilter:z.getColumnData(e,T.filter_defaultFilter,R.columns,!0)||""};for(O.parsed=[],d=0;d<R.columns;d++)O.parsed[d]=T.filter_useParsedData||R.parsers&&R.parsers[d]&&R.parsers[d].parsed||z.getData&&"parsed"===z.getData(R.$headerIndexed[d],z.getColumnData(e,R.headers,d),"filter")||R.$headerIndexed[d].hasClass("filter-parsed"),E.functions[d]=z.getColumnData(e,T.filter_functions,d)||R.$headerIndexed[d].hasClass("filter-select"),E.defaultColFilter[d]=z.getColumnData(e,T.filter_defaultFilter,d)||"",E.excludeFilter[d]=(z.getColumnData(e,T.filter_excludeFilter,d,!0)||"").split(/\s+/);for(A&&(console.log("Filter >> Starting filter widget search",r),m=new Date),R.filteredRows=0,t=S||[],c=R.totalRows=0;c<R.$tbodies.length;c++){if(f=z.processTbody(e,R.$tbodies.eq(c),!0),d=R.columns,l=R.cache[c].normalized,n=M(M.map(l,function(e){return e[d].$row.get()})),""===t.join("")||T.filter_serversideFiltering)n.removeClass(T.filter_filteredRow).not("."+R.cssChildRow).css("display","");else{if(i=(n=n.not("."+R.cssChildRow)).length,(T.filter_$anyMatch&&T.filter_$anyMatch.length||void 0!==r[R.columns])&&(O.anyMatchFlag=!0,O.anyMatchFilter=""+(r[R.columns]||T.filter_$anyMatch&&I.getLatestSearch(T.filter_$anyMatch).val()||""),T.filter_columnAnyMatch)){for(x=O.anyMatchFilter.split(D.andSplit),b=!1,y=0;y<x.length;y++)1<(C=x[y].split(":")).length&&(isNaN(C[0])?M.each(R.headerContent,function(e,t){-1<t.toLowerCase().indexOf(C[0])&&(r[v=e]=C[1])}):v=parseInt(C[0],10)-1,0<=v)&&v<R.columns&&(r[v]=C[1],x.splice(y,1),y--,b=!0);b&&(O.anyMatchFilter=x.join(" && "))}if(w=T.filter_searchFiltered,h=R.lastSearch||R.$table.data("lastSearch")||[],w)for(y=0;y<d+1;y++)_=r[y]||"",w||(y=d),w=w&&h.length&&0===_.indexOf(h[y]||"")&&!D.alreadyFiltered.test(_)&&!D.exactTest.test(_)&&!(D.isNeg1.test(_)||D.isNeg2.test(_))&&!(""!==_&&R.$filters&&R.$filters.filter('[data-column="'+y+'"]').find("select").length&&!I.matchType(R,y));for(F=n.not("."+T.filter_filteredRow).length,w&&0===F&&(w=!1),A&&console.log("Filter >> Searching through "+(w&&F<i?F:"all")+" rows"),O.anyMatchFlag&&(R.sortLocaleCompare&&(O.anyMatchFilter=z.replaceAccents(O.anyMatchFilter)),T.filter_defaultFilter&&D.iQuery.test(E.defaultAnyFilter)&&(O.anyMatchFilter=I.defaultFilter(O.anyMatchFilter,E.defaultAnyFilter),w=!1),O.iAnyMatchFilter=T.filter_ignoreCase&&R.ignoreCase?O.anyMatchFilter.toLowerCase():O.anyMatchFilter),o=0;o<i;o++)if($=n[o].className,!(o&&D.child.test($)||w&&D.filtered.test($))){if(O.$row=n.eq(o),O.rowIndex=o,O.cacheArray=l[o],a=O.cacheArray[R.columns],O.rawArray=a.raw,O.childRowText="",!T.filter_childByColumn){for($="",u=a.child,y=0;y<u.length;y++)$+=" "+u[y].join(" ")||"";O.childRowText=T.filter_childRows?T.filter_ignoreCase?$.toLowerCase():$:""}if(p=!1,g=I.processRow(R,O,E),s=a.$row,_=!!g,u=a.$row.filter(":gt(0)"),T.filter_childRows&&u.length){if(T.filter_childByColumn)for(T.filter_childWithSibs||(u.addClass(T.filter_filteredRow),s=s.eq(0)),y=0;y<u.length;y++)O.$row=u.eq(y),O.cacheArray=a.child[y],O.rawArray=O.cacheArray,_=I.processRow(R,O,E),p=p||_,!T.filter_childWithSibs&&_&&u.eq(y).removeClass(T.filter_filteredRow);p=p||g}else p=_;s.toggleClass(T.filter_filteredRow,!p)[0].display=p?"":"none"}}R.filteredRows+=n.not("."+T.filter_filteredRow).length,R.totalRows+=n.length,z.processTbody(e,f,!1)}R.lastCombinedFilter=S.join(""),R.lastSearch=S,R.$table.data("lastSearch",S),T.filter_saveFilters&&z.storage&&z.storage(e,"tablesorter-filters",I.processFilters(S,!0)),A&&console.log("Filter >> Completed search"+z.benchmark(m)),T.filter_initialized&&(R.$table.triggerHandler("filterBeforeEnd",R),R.$table.triggerHandler("filterEnd",R)),setTimeout(function(){z.applyWidget(R.table)},0)}},getOptionSource:function(e,t,r){var i=(e=M(e)[0]).config,l=!1,a=i.widgetOptions.filter_selectSource,i=i.$table.data("lastSearch")||[],n="function"==typeof a||z.getColumnData(e,a,t);if(r&&""!==i[t]&&(r=!1),!0===n)l=a(e,t,r);else{if(n instanceof M||"string"===M.type(n)&&0<=n.indexOf("</option>"))return n;if(M.isArray(n))l=n;else if("object"===M.type(a)&&n&&null===(l=n(e,t,r)))return null}return!1===l&&(l=I.getOptions(e,t,r)),I.processOptions(e,t,l)},processOptions:function(i,l,r){if(!M.isArray(r))return!1;var a,e,t,n,s,o=(i=M(i)[0]).config,c=null!=l&&0<=l&&l<o.columns,f=!!c&&o.$headerIndexed[l].hasClass("filter-select-sort-desc"),d=[];if(r=M.grep(r,function(e,t){return!!e.text||M.inArray(e,r)===t}),!c||!o.$headerIndexed[l].hasClass("filter-select-nosort")){for(n=r.length,t=0;t<n;t++)s=(e=r[t]).text||e,s=(c&&o.parsers&&o.parsers.length&&o.parsers[l].format(s,i,[],l)||s).toString(),s=o.widgetOptions.filter_ignoreCase?s.toLowerCase():s,e.text?(e.parsed=s,d[d.length]=e):d[d.length]={text:e,parsed:s};for(a=o.textSorter||"",d.sort(function(e,t){var r=(f?t:e).parsed,e=(f?e:t).parsed;return c&&"function"==typeof a?a(r,e,!0,l,i):c&&"object"==typeof a&&a.hasOwnProperty(l)?a[l](r,e,!0,l,i):!z.sortNatural||z.sortNatural(r,e)}),r=[],n=d.length,t=0;t<n;t++)r[r.length]=d[t]}return r},getOptions:function(e,t,r){for(var i,l,a,n,s,o,c,f=(e=M(e)[0]).config,d=f.widgetOptions,u=[],h=0;h<f.$tbodies.length;h++)for(n=f.cache[h],l=f.cache[h].normalized.length,i=0;i<l;i++)if(a=n.row?n.row[i]:n.normalized[i][f.columns].$row[0],!r||!a.className.match(d.filter_filteredRow))if(d.filter_useParsedData||f.parsers[t].parsed||f.$headerIndexed[t].hasClass("filter-parsed")){if(u[u.length]=""+n.normalized[i][t],d.filter_childRows&&d.filter_childByColumn)for(c=n.normalized[i][f.columns].$row.length-1,s=0;s<c;s++)u[u.length]=""+n.normalized[i][f.columns].child[s][t]}else if(u[u.length]=n.normalized[i][f.columns].raw[t],d.filter_childRows&&d.filter_childByColumn)for(c=n.normalized[i][f.columns].$row.length,s=1;s<c;s++)o=n.normalized[i][f.columns].$row.eq(s).children().eq(t),u[u.length]=""+z.getElementText(f,o,t);return u},buildSelect:function(e,t,r,i,l){if(e=M(e)[0],t=parseInt(t,10),e.config.cache&&!M.isEmptyObject(e.config.cache)){var a,n,s,o,c,f=e.config,d=f.widgetOptions,u=f.$headerIndexed[t],h='<option value="">'+(u.data("placeholder")||u.attr("data-placeholder")||d.filter_placeholder.select||"")+"</option>",u=f.$table.find("thead").find("select."+g.filter+'[data-column="'+t+'"]').val();if(void 0!==r&&""!==r||null!==(r=I.getOptionSource(e,t,l))){if(M.isArray(r)){for(a=0;a<r.length;a++)if((c=r[a]).text){for(n in c["data-function-name"]=void 0===c.value?c.text:c.value,h+="<option",c)c.hasOwnProperty(n)&&"text"!==n&&(h+=" "+n+'="'+c[n].replace(D.quote,"&quot;")+'"');c.value||(h+=' value="'+c.text.replace(D.quote,"&quot;")+'"'),h+=">"+c.text.replace(D.quote,"&quot;")+"</option>"}else""+c!="[object Object]"&&(0<=(n=s=c=(""+c).replace(D.quote,"&quot;")).indexOf(d.filter_selectSourceSeparator)&&(n=(o=s.split(d.filter_selectSourceSeparator))[0],s=o[1]),h+=""!==c?"<option "+(n===s?"":'data-function-name="'+c+'" ')+'value="'+n+'">'+s+"</option>":"");r=[]}e=(f.$filters||f.$table.children("thead")).find("."+g.filter),(l=(e=d.filter_$externalFilters?e&&e.length?e.add(d.filter_$externalFilters):d.filter_$externalFilters:e).filter('select[data-column="'+t+'"]')).length&&(l[i?"html":"append"](h),M.isArray(r)||l.append(r).val(u),l.val(u))}}},buildDefault:function(e,t){for(var r,i,l=e.config,a=l.widgetOptions,n=l.columns,s=0;s<n;s++)i=!((r=l.$headerIndexed[s]).hasClass("filter-false")||r.hasClass("parser-false")),(r.hasClass("filter-select")||!0===z.getColumnData(e,a.filter_functions,s))&&i&&I.buildSelect(e,s,"",t,r.hasClass(a.filter_onlyAvail))}},D=I.regex,z.getFilters=function(e,t,r,i){var l,a,n,s,o=[],c=e?M(e)[0].config:"",f=c?c.widgetOptions:"";if(!0!==t&&f&&!f.filter_columnFilters||M.isArray(r)&&I.equalFilters(c,r,c.lastSearch))return M(e).data("lastSearch")||[];if(c&&(c.$filters&&(a=c.$filters.find("."+g.filter)),a=f.filter_$externalFilters?a&&a.length?a.add(f.filter_$externalFilters):f.filter_$externalFilters:a)&&a.length)for(o=r||[],l=0;l<c.columns+1;l++)s=l===c.columns?f.filter_anyColumnSelector+","+f.filter_multipleColumnSelector:'[data-column="'+l+'"]',(n=a.filter(s)).length&&(n=I.getLatestSearch(n),M.isArray(r)?(i&&1<n.length&&(n=n.slice(1)),(n=l===c.columns?(s=n.filter(f.filter_anyColumnSelector)).length?s:n:n).val(r[l]).trigger("change"+c.namespace)):(o[l]=n.val()||"",(l===c.columns?n.slice(1).filter('[data-column*="'+n.attr("data-column")+'"]'):n.slice(1)).val(o[l])),l===c.columns)&&n.length&&(f.filter_$anyMatch=n);return o},z.setFilters=function(e,t,r,i){var l=e?M(e)[0].config:"",e=z.getFilters(e,!0,t,i);return void 0===r&&(r=!0),l&&r&&(l.lastCombinedFilter=null,l.lastSearch=[],I.searching(l.table,t,i),l.$table.triggerHandler("filterFomatterUpdate")),0!==e.length}}(jQuery);return jQuery;}));
```
