## Supplementary Data SD1 for "Inferring binding specificities of human transcription factors with the wisdom of crowds": widget-formatter.min.js

```
(function(factory){if (typeof define === 'function' && define.amd){define(['jquery'], factory);} else if (typeof module === 'object' && typeof module.exports === 'object'){module.exports = factory(require('jquery'));} else {factory(jQuery);}}(function(jQuery){
/*! Widget: formatter - 2/9/2015 (v2.19.1) */
!function(d){"use strict";var u=d.tablesorter;u.formatter={init:function(t){var e=t.widgetOptions.formatter_event+" pagerComplete updateComplete ".split(" ").join(".tsformatter ");t.$table.off(e.replace(/\s+/g," ")).on(e,function(){u.formatter.setup(t)}),u.formatter.setup(t)},setup:function(t){if(!d.isEmptyObject(t.cache)){for(var e,o,r,n,i,a,l=t.widgetOptions,c={config:t,wo:l},f=[],s=[],m=0;m<t.columns;m++)s[m]=t.$headerIndexed[m],f[m]=u.getColumnData(t.table,l.formatter_column,m)||!1;for(o=0;o<t.$tbodies.length;o++){for(e=u.processTbody(t.table,t.$tbodies.eq(o),!0),a=(n=t.cache[o]).normalized.length,r=0;r<a;r++)for(c.$row=n.normalized[r][t.columns].$row,c.$cells=c.$row.children("th, td"),m=0;m<t.columns;m++)f[m]&&(c.columnIndex=m,c.$header=s[m],c.$cell=c.$cells.eq(m),i=c.$cell[0],c.text=i.getAttribute(t.textAttribute)||i.textContent||c.$cell.text(),i.innerHTML=f[m](c.text,c));u.processTbody(t.table,e,!1)}}}},u.addWidget({id:"formatter",priority:100,options:{formatter_column:{},formatter_event:"applyFormatter"},init:function(t){u.formatter.init(t.config)}})}(jQuery);return jQuery;}));
```
