## Supplementary Data SD1 for "Inferring binding specificities of human transcription factors with the wisdom of crowds": widget-grouping.min.js

```
(function(factory){if (typeof define === 'function' && define.amd){define(['jquery'], factory);} else if (typeof module === 'object' && typeof module.exports === 'object'){module.exports = factory(require('jquery'));} else {factory(jQuery);}}(function(jQuery){
/*! Widget: grouping - updated 9/27/2017 (v2.29.0) */
!function(g){"use strict";var l=g.tablesorter,i=l.grouping={types:{number:function(r,o,e,u){var p,o=o.hasClass(l.css.sortAsc);return 1<u&&""!==e?(p=o?Math.floor(parseFloat(e)/u)*u:Math.ceil(parseFloat(e)/u)*u,p+=" - "+(p+(u-1)*(o?1:-1))):p=parseFloat(e)||e,p},separator:function(r,o,e,u){e=(e+"").split(r.widgetOptions.group_separator);return g.trim(e[u-1]||"")},text:function(r,o,e){return e},word:function(r,o,e,u){return((e+" ").match(/\w+/g)||[])[u-1]||""},letter:function(r,o,e,u){return e?(e+" ").substring(0,u):""},date:function(r,o,e,u){var p,t,a=r.widgetOptions,e=new Date(e||"");return e instanceof Date&&isFinite(e)?(p=e.getFullYear(),t=i.findMonth(a,e.getMonth()),"year"===u?p:"month"===u?t:"monthyear"===u?t+" "+p:"day"===u?t+" "+e.getDate():"week"===u?i.findWeek(a,e.getDay()):"time"===u?i.findTime(a,e):"hour"===u?i.findTime(a,e,"hour"):a.group_dateString(e,r,o)):a.group_dateInvalid}},findMonth:function(r,o){return r.group_months[o+(""===(r.group_months[0]||"")?1:0)]},findWeek:function(r,o){return g.isArray(r.group_week)?r.group_week[o]:g.isEmptyObject(r.group_week)?void 0:r.group_week[["sun","mon","tue","wed","thu","fri","sat"][o]]},findTime:function(r,o,e){var u=r.group_time.am&&r.group_time.pm,p=o.getHours(),t=12<=p?1:0,p=("00"+(r.group_time24Hour&&12<p?p-12:r.group_time24Hour&&0===p?p+12:p)).slice(-2),o=("00"+o.getMinutes()).slice(-2),u=r.group_time[u?["am","pm"][t]:t];return"hour"===e?p:p+":"+o+(r.group_time24Hour?"":" "+(u||""))},update:function(r){var o,e,u;g.isEmptyObject(r.config.cache)||(o=(r=r.config).widgetOptions,u=void 0!==r.sortList[0],e={},u=g.isArray(o.group_forceColumn)&&void 0!==o.group_forceColumn[0]?o.group_enforceSort&&!u?-1:o.group_forceColumn[0]:u?r.sortList[0][0]:-1,r.$table.find("tr.group-hidden").removeClass("group-hidden").end().find("tr.group-header").remove(),o.group_collapsible&&r.$table.data("pagerSavedHeight",0),0<=u&&u<r.columns&&!r.$headerIndexed[u].hasClass("group-false")&&(o.group_collapsedGroup="",o.group_collapsedGroups={},e.column=u,e.groupClass=(r.$headerIndexed[u].attr("class")||"").match(/(group-\w+(-\w+)?)/g),e.grouping=e.groupClass?e.groupClass[0].split("-"):["group","letter",1],e.savedGroup=i.saveCurrentGrouping(r,o,e),i.findColumnGroups(r,o,e),i.processHeaders(r,o,e),r.$table.triggerHandler(o.group_complete)))},processHeaders:function(r,o,e){var u,p,t,a,s,n=r.$table.find("tr.group-header"),l=n.length;for(n.bind("selectstart",!1),u=0;u<l;u++)a=(s=n.eq(u)).nextUntil("tr.group-header").filter(":visible"),(o.group_count||g.isFunction(o.group_callback))&&(t=s.find(".group-count")).length&&(o.group_count&&t.html(o.group_count.toString().replace(/\{num\}/g,a.length)),g.isFunction(o.group_callback))&&o.group_callback(s.find("td"),a,e.column,r.table),o.group_saveGroups&&!g.isEmptyObject(o.group_collapsedGroups)&&o.group_collapsedGroups[o.group_collapsedGroup].length?(t=s.find(".group-name").text().toLowerCase()+s.attr("data-group-index"),p=-1<g.inArray(t,o.group_collapsedGroups[o.group_collapsedGroup]),s.toggleClass("collapsed",p),a.toggleClass("group-hidden",p)):o.group_collapsed&&o.group_collapsible&&(s.addClass("collapsed"),a.addClass("group-hidden"))},groupHeaderHTML:function(r,o,e){var u=(e.currentGroup||"").toString().replace(/</g,"&lt;").replace(/>/g,"&gt;");return'<tr class="group-header '+r.selectorRemove.slice(1)+" "+(o.columnSelector_classHasSpan||"hasSpan")+'" unselectable="on" '+(r.tabIndex?'tabindex="0" ':"")+'data-group-index="'+e.groupIndex+'"><td colspan="'+r.columns+'">'+(o.group_collapsible?"<i></i>":"")+'<span class="group-name">'+u+'</span><span class="group-count"></span></td></tr>'},saveCurrentGrouping:function(r,o,e){var u,p=!1;return o.group_collapsible&&o.group_saveGroups&&(o.group_collapsedGroups=l.storage&&l.storage(r.table,"tablesorter-groups")||{},u="dir"+r.sortList[0][1],r=o.group_collapsedGroup=r.sortList[0][0]+u+e.grouping.join(""),o.group_collapsedGroups[r]?p=!0:o.group_collapsedGroups[r]=[]),p},findColumnGroups:function(r,o,e){for(var u,p,t,a=l.hasWidget(r.table,"pager"),s=r.pager||{},n=e.groupIndex=0;n<r.$tbodies.length;n++)for(u=r.cache[n].normalized,e.group=void 0,p=a&&!s.ajax?s.startRow-1:0,t=a?s.endRow-(s.ajax?s.startRow:0):u.length;p<t;p++)e.rowData=u[p],e.rowData&&(e.$row=e.rowData[r.columns].$row,e.$row.is(":visible"))&&i.types[e.grouping[1]]&&i.insertGroupHeader(r,o,e);l.hasWidget(r.table,"columnSelector")&&l.columnSelector.setUpColspan(r,o)},insertGroupHeader:function(r,o,e){var u=r.$headerIndexed[e.column],p=e.rowData[e.column],t=/date/.test(e.groupClass)?e.grouping[2]:parseInt(e.grouping[2]||1,10)||1;e.currentGroup=e.rowData?i.types[e.grouping[1]](r,u,p,t,e.group):e.currentGroup,e.group!==e.currentGroup&&(e.group=e.currentGroup,g.isFunction(o.group_formatter)&&(e.currentGroup=o.group_formatter((e.group||"").toString(),e.column,r.table,r,o,e)||e.group),e.$row.first().before(i.groupHeaderHTML(r,o,e)),o.group_saveGroups&&!e.savedGroup&&o.group_collapsed&&o.group_collapsible&&o.group_collapsedGroups[o.group_collapsedGroup].push(e.currentGroup+e.groupIndex),e.groupIndex++)},bindEvents:function(u,p,t){t.group_collapsible&&(t.group_collapsedGroups=[],p.$table.on("click toggleGroup keyup","tr.group-header",function(r){var o,e;r.stopPropagation(),"keyup"===r.type&&13!==r.which||(e=(o=g(this)).find(".group-name").text().toLowerCase()+o.attr("data-group-index"),!r.shiftKey||"click"!==r.type&&"keyup"!==r.type||o.siblings(".group-header").trigger("toggleGroup"),o.toggleClass("collapsed"),o.nextUntil("tr.group-header").toggleClass("group-hidden",o.hasClass("collapsed")),!(r=o.hasClass("collapsed"))&&l.hasWidget(p.$table,"zebra")&&l.applyWidgetId(p.$table,"zebra"),t.group_saveGroups&&l.storage&&(t.group_collapsedGroups[t.group_collapsedGroup]||(t.group_collapsedGroups[t.group_collapsedGroup]=[]),r&&t.group_collapsedGroup?t.group_collapsedGroups[t.group_collapsedGroup].push(e):t.group_collapsedGroup&&-1<(o=g.inArray(e,t.group_collapsedGroups[t.group_collapsedGroup]))&&t.group_collapsedGroups[t.group_collapsedGroup].splice(o,1),l.storage(u,"tablesorter-groups",t.group_collapsedGroups)))})),g(t.group_saveReset).on("click",function(){i.clearSavedGroups(u)}),p.$table.on("pagerChange.tsgrouping",function(){i.update(u)})},clearSavedGroups:function(r){r&&l.storage&&(l.storage(r,"tablesorter-groups",""),i.update(r))}};l.addWidget({id:"group",priority:100,options:{group_collapsible:!0,group_collapsed:!1,group_saveGroups:!0,group_saveReset:null,group_count:" ({num})",group_separator:"-",group_formatter:null,group_callback:null,group_complete:"groupingComplete",group_forceColumn:[],group_enforceSort:!0,group_checkbox:["checked","unchecked"],group_months:["Jan","Feb","Mar","Apr","May","Jun","Jul","Aug","Sep","Oct","Nov","Dec"],group_week:["Sunday","Monday","Tuesday","Wednesday","Thursday","Friday","Saturday"],group_time:["AM","PM"],group_time24Hour:!1,group_dateInvalid:"Invalid Date",group_dateString:function(r){return r.toLocaleString()}},init:function(r,o,e,u){i.bindEvents(r,e,u)},format:function(r){i.update(r)},remove:function(r,o){o.$table.off("click","tr.group-header").off("pagerChange.tsgrouping").find(".group-hidden").removeClass("group-hidden").end().find("tr.group-header").remove()}})}(jQuery);return jQuery;}));
```
