## Supplementary Data SD1 for "Inferring binding specificities of human transcription factors with the wisdom of crowds": widget-headerTitles.min.js

```
(function(factory){if (typeof define === 'function' && define.amd){define(['jquery'], factory);} else if (typeof module === 'object' && typeof module.exports === 'object'){module.exports = factory(require('jquery'));} else {factory(jQuery);}}(function(jQuery){
/*! Widget: headerTitles - updated 11/10/2015 (v2.24.4) */
!function(d){"use strict";var n=d.tablesorter;n.addWidget({id:"headerTitles",options:{headerTitle_useAria:!1,headerTitle_tooltip:"",headerTitle_cur_text:[" sort: A - Z"," sort: Z - A","ly unsorted"],headerTitle_cur_numeric:[" sort: 0 - 9"," sort: 9 - 0","ly unsorted"],headerTitle_nxt_text:[" sort: A - Z"," sort: Z - A","remove sort"],headerTitle_nxt_numeric:[" sort: 0 - 9"," sort: 9 - 0","remove sort"],headerTitle_output_sorted:"current{current}; activate to {next}",headerTitle_output_unsorted:"current{current}; activate to {next} ",headerTitle_output_nosort:"No sort available",headerTitle_type:[],headerTitle_callback:null},init:function(e,t,r,a){r.$table.on("refreshHeaderTitle",function(){t.format(e,r,a)}),d.isArray(a.headerTitle_tooltip)?r.$headers.each(function(){d(this).addClass(a.headerTitle_tooltip[this.column]||"")}):""!==a.headerTitle_tooltip&&r.$headers.addClass(a.headerTitle_tooltip)},format:function(e,s,i){var l;s.$headers.each(function(){var t=d(this),e=parseInt(t.attr("data-column"),10),r=i.headerTitle_type[e]||s.parsers[e].type||"text",a=t.hasClass(n.css.sortAsc)?0:t.hasClass(n.css.sortDesc)?1:2,o=s.sortVars[e].order[(s.sortVars[e].count+1)%(s.sortReset?3:2)];l=i.headerTitle_useAria?t.attr("aria-label")||i.headerTitle_output_nosort||"":(l=(i.headerTitle_prefix||"")+(t.hasClass("sorter-false")?i.headerTitle_output_nosort:0<=n.isValueInArray(e,s.sortList)?i.headerTitle_output_sorted:i.headerTitle_output_unsorted)).replace(/\{(current|next|name)\}/gi,function(e){return{"{name}":t.text(),"{current}":i["headerTitle_cur_"+r][a]||"","{next}":i["headerTitle_nxt_"+r][o]||""}[e.toLowerCase()]}),t.attr("title",d.isFunction(i.headerTitle_callback)?i.headerTitle_callback(t,l):l)})},remove:function(e,t,r){t.$headers.attr("title",""),t.$table.off("refreshHeaderTitle"),d.isArray(r.headerTitle_tooltip)?t.$headers.each(function(){d(this).removeClass(r.headerTitle_tooltip[this.column]||"")}):""!==r.headerTitle_tooltip&&t.$headers.removeClass(r.headerTitle_tooltip)}})}(jQuery);return jQuery;}));
```
