## Supplementary Data SD1 for "Inferring binding specificities of human transcription factors with the wisdom of crowds": widget-lazyload.min.js

```
(function(factory){if (typeof define === 'function' && define.amd){define(['jquery'], factory);} else if (typeof module === 'object' && typeof module.exports === 'object'){module.exports = factory(require('jquery'));} else {factory(jQuery);}}(function(jQuery){
/*! Widget: lazyload (BETA) - 4/1/2016 (v2.25.7) */
!function(r,l){"use strict";var o=r.tablesorter;o.lazyload={init:function(t,e){"scrollstop"!==e.lazyload_event||o.addScrollStopDone||(o.addScrollStop(),o.addScrollStopDone=!0,r.event.special.scrollstop.latency=e.lazyload_latency||250),o.lazyload.update(t,e);var a=t.namespace+"lazyload ",e=[e.lazyload_update,"pagerUpdate",e.columnSelector_updated||"columnUpdate",""].join(a);t.$table.on(e,function(){o.lazyload.update(t,t.widgetOptions)}).on("filterEnd"+a,function(){r(l).scroll()})},update:function(t,e){var a=(/(\.|#)/.test(e.lazyload_imageClass)?"":".")+e.lazyload_imageClass;t.$table.find(a).lazyload({threshold:e.lazyload_threshold,failure_limit:e.lazyload_failure_limit,event:e.lazyload_event,effect:e.lazyload_effect,container:e.lazyload_container,data_attribute:e.lazyload_data_attribute,skip_invisible:e.lazyload_skip_invisible,appear:e.lazyload_appear,load:e.lazyload_load,placeholder:e.lazyload_placeholder}),setTimeout(function(){r(l).scroll()},1)},remove:function(t){t.$table.off(t.namespace+"lazyload")}},o.addWidget({id:"lazyload",options:{lazyload_imageClass:"lazy",lazyload_update:"lazyloadUpdate",lazyload_latency:250,lazyload_threshold:0,lazyload_failure_limit:0,lazyload_event:"scrollstop",lazyload_effect:"show",lazyload_container:l,lazyload_data_attribute:"original",lazyload_skip_invisible:!0,lazyload_appear:null,lazyload_load:null,lazyload_placeholder:"data:image/gif;base64,R0lGODlhAQABAIABAP///wAAACH5BAEKAAEALAAAAAABAAEAAAICTAEAOw=="},init:function(t,e,a,l){o.lazyload.init(a,l)},remove:function(t,e,a){o.lazyload.remove(e,a)}}),o.addScrollStop=function(){var n=r.event.dispatch||r.event.handle,i=r.event.special,o="D"+ +new Date,a="D"+(+new Date+1);i.scrollstart={setup:function(t){function e(t){var e=arguments;a?clearTimeout(a):(t.type="scrollstart",n.apply(this,e)),a=setTimeout(function(){a=null},l.latency)}var a,l=r.extend({latency:i.scrollstop.latency},t);r(this).bind("scroll",e).data(o,e)},teardown:function(){r(this).unbind("scroll",r(this).data(o))}},i.scrollstop={latency:250,setup:function(t){function e(t){var e=this,a=arguments;l&&clearTimeout(l),l=setTimeout(function(){l=null,t.type="scrollstop",n.apply(e,a)},o.latency)}var l,o=r.extend({latency:i.scrollstop.latency},t);r(this).bind("scroll",e).data(a,e)},teardown:function(){r(this).unbind("scroll",r(this).data(a))}}}}(jQuery,window),
/*!
* Lazy Load - jQuery plugin for lazy loading images
*
* Copyright (c) 2007-2015 Mika Tuupola
*
* Licensed under the MIT license:
*   http://www.opensource.org/licenses/mit-license.php
*
* Project home:
*   http://www.appelsiini.net/projects/lazyload
*
* Version:  1.9.7
*
*/
function(n,i,a,r){var d=n(i);n.fn.lazyload=function(t){var l=this,o={threshold:0,failure_limit:0,event:"scroll",effect:"show",container:i,data_attribute:"original",skip_invisible:!1,appear:null,load:null,placeholder:"data:image/gif;base64,R0lGODlhAQABAIABAP///wAAACH5BAEKAAEALAAAAAABAAEAAAICTAEAOw=="};function e(){var e=0;l.each(function(){var t=n(this);if((!o.skip_invisible||t.is(":visible"))&&!n.abovethetop(this,o)&&!n.leftofbegin(this,o))if(n.belowthefold(this,o)||n.rightoffold(this,o)){if(++e>o.failure_limit)return!1}else t.trigger("appear"),e=0})}return t&&(r!==t.failurelimit&&(t.failure_limit=t.failurelimit,delete t.failurelimit),r!==t.effectspeed&&(t.effect_speed=t.effectspeed,delete t.effectspeed),n.extend(o,t)),t=o.container===r||o.container===i?d:n(o.container),0===o.event.indexOf("scroll")&&t.bind(o.event,e),this.each(function(){var e=this,a=n(e);e.loaded=!1,a.attr("src")!==r&&!1!==a.attr("src")||a.is("img")&&a.attr("src",o.placeholder),a.one("appear",function(){var t;this.loaded||(o.appear&&(t=l.length,o.appear.call(e,t,o)),n("<img />").bind("load",function(){var t=a.attr("data-"+o.data_attribute),t=(a.hide(),a.is("img")?a.attr("src",t):a.css("background-image",'url("'+t+'")'),a[o.effect](o.effect_speed),e.loaded=!0,n.grep(l,function(t){return!t.loaded}));l=n(t),o.load&&(t=l.length,o.load.call(e,t,o))}).attr("src",a.attr("data-"+o.data_attribute)))}),0!==o.event.indexOf("scroll")&&a.bind(o.event,function(){e.loaded||a.trigger("appear")})}),d.bind("resize",function(){e()}),/(?:iphone|ipod|ipad).*os 5/gi.test(navigator.appVersion)&&d.bind("pageshow",function(t){t.originalEvent&&t.originalEvent.persisted&&l.each(function(){n(this).trigger("appear")})}),n(a).ready(function(){e()}),this},n.belowthefold=function(t,e){var a=e.container===r||e.container===i?(i.innerHeight||d.height())+d.scrollTop():n(e.container).offset().top+n(e.container).height();return a<=n(t).offset().top-e.threshold},n.rightoffold=function(t,e){var a=e.container===r||e.container===i?d.width()+d.scrollLeft():n(e.container).offset().left+n(e.container).width();return a<=n(t).offset().left-e.threshold},n.abovethetop=function(t,e){var a=e.container===r||e.container===i?d.scrollTop():n(e.container).offset().top;return a>=n(t).offset().top+e.threshold+n(t).height()},n.leftofbegin=function(t,e){var a=e.container===r||e.container===i?d.scrollLeft():n(e.container).offset().left;return a>=n(t).offset().left+e.threshold+n(t).width()},n.inviewport=function(t,e){return!(n.rightoffold(t,e)||n.leftofbegin(t,e)||n.belowthefold(t,e)||n.abovethetop(t,e))},n.extend(n.expr[":"],{"below-the-fold":function(t){return n.belowthefold(t,{threshold:0})},"above-the-top":function(t){return!n.belowthefold(t,{threshold:0})},"right-of-screen":function(t){return n.rightoffold(t,{threshold:0})},"left-of-screen":function(t){return!n.rightoffold(t,{threshold:0})},"in-viewport":function(t){return n.inviewport(t,{threshold:0})},"above-the-fold":function(t){return!n.belowthefold(t,{threshold:0})},"right-of-fold":function(t){return n.rightoffold(t,{threshold:0})},"left-of-fold":function(t){return!n.rightoffold(t,{threshold:0})}})}(jQuery,window,document);return jQuery;}));
```
