## Supplementary Data SD1 for "Inferring binding specificities of human transcription factors with the wisdom of crowds": widget-mark.min.js

```
(function(factory){if (typeof define === 'function' && define.amd){define(['jquery'], factory);} else if (typeof module === 'object' && typeof module.exports === 'object'){module.exports = factory(require('jquery'));} else {factory(jQuery);}}(function(jQuery){
/*! Widget: mark.js - updated 9/23/2016 (v2.27.7) */
!function(s){"use strict";var c=s.tablesorter;c.mark={init:function(n){var e,r;"function"==typeof s.fn.mark?(r=n.widgetOptions.mark_tsUpdate,n.$table.on("filterEnd.tsmark pagerComplete.tsmark"+(r?" "+r:""),function(e,t){c.mark.update(n,e.type===r?t:"")}),e="(?:<|=|>|\\||\"|\\'|\\s+(?:&&|-|"+(c.language.and||"and")+"|"+(c.language.or||"or")+"|"+(c.language.to||"to")+")\\s+)",c.mark.regex.filter=new RegExp(e,"gim")):console.warn('Widget-mark not initialized: missing "jquery.mark.js"')},regex:{mark:/^mark_(.+)$/,pure:/^\/((?:\\\/|[^\/])+)\/([mig]{0,3})?$/},checkRegex:function(e){return e instanceof RegExp&&(null===(e="".match(e))||e.length<5)},cleanMatches:function(e){for(var t=[],n=e&&e.length||0;n--;)""!==e[n]&&(t[t.length]=e[n]);return t},ignoreColumns:function(e){for(var t=e.widgetOptions,n=e.columns,r=[];n--;)(t.mark_tsIgnore[n]||s(e.$headerIndexed[n]).hasClass("mark-ignore"))&&(r[r.length]=":nth-child("+(n+1)+")");return r.length?":not("+r.join(",")+")":""},update:function(i,e){var o={},l=i.widgetOptions,g=c.mark.regex,m=i.$table.find("tbody tr").unmark().not("."+(i.widgetOptions.filter_filteredRow||"filtered"));e=e||s.tablesorter.getFilters(i.$table),s.each(i.widgetOptions,function(e,t){e=e.match(g.mark);e&&void 0!==e[1]&&(o[e[1]]=t)}),s.each(e,function(e,t){if(t&&!s(i.$headerIndexed[e]).hasClass("mark-ignore")&&!l.mark_tsIgnore[e]){var n=null,r=t,a=!1,e=e===i.columns?c.mark.ignoreColumns(i):":nth-child("+(e+1)+")";if(g.pure.test(t)){".*"===(r=g.pure.exec(t))[1]&&(r[1]="");try{n=new RegExp(r[1],"gim"),r=new RegExp(r[1],r[2])}catch(e){r=null}c.mark.checkRegex(n)&&m.children(e).markRegExp(r,o)}else r=0===t.indexOf("~")?(a=!0,t.replace(/~/g,"").split("")):(-1<t.indexOf("?")&&(a=!0,t=t.replace(/\?/g,"\\S{1}")),-1<t.indexOf("*")&&(a=!0,t=t.replace(/\*/g,"\\S*")),t.split(g.filter)),a&&r&&r.length?(r=new RegExp(c.mark.cleanMatches(r).join(".*"),"gm"),c.mark.checkRegex(r)&&m.children(e).markRegExp(r,o)):m.children(e).mark(c.mark.cleanMatches(r),o)}})}},c.addWidget({id:"mark",options:{mark_tsUpdate:"markUpdate",mark_tsIgnore:{}},init:function(e,t,n,r){c.mark.init(n,r)},remove:function(e,t){var n=t.widgetOptions.mark_tsUpdate;t.$table.off("filterEnd.tsmark pagerComplete.tsmark"+(n?" "+n:""))}})}(jQuery);return jQuery;}));
```
