## Supplementary Data SD1 for "Inferring binding specificities of human transcription factors with the wisdom of crowds": widget-math.min.js

```
(function(factory){if (typeof define === 'function' && define.amd){define(['jquery'], factory);} else if (typeof module === 'object' && typeof module.exports === 'object'){module.exports = factory(require('jquery'));} else {factory(jQuery);}}(function(jQuery){
/*! Widget: math - updated 12/1/2019 (v2.31.2) */
!function(v){"use strict";var _=v.tablesorter,x={error:{0:"Infinity result: Divide by zero",1:"Need more than one element to make this calculation",undef:"No elements found"},invalid:function(t,e,n){return console.warn(e,x.error[n]),t&&t.widgetOptions.math_none||""},events:"tablesorter-initialized update updateAll updateRows addRows updateCell filterReset ".split(" ").join(".tsmath "),processText:function(t,e){var n=t.widgetOptions,a=_.getElementText(t,e,x.getCellIndex(e)),r=t.widgetOptions.math_prefix;return n.math_textAttr&&(a=e.attr(n.math_textAttr)||a),/</.test(r)&&(e=v("<div>"+r+"</div>").text().replace(/\{content\}/g,"").trim(),a=a.replace(e,"")),a=_.formatFloat(a.replace(/[^\w,. \-()]/g,""),t.table)||0,isNaN(a)?0:a},getRow:function(t,e,n){var a=t.widgetOptions,r=[],i=e.closest("tr"),l=i.hasClass(a.filter_filteredRow||"filtered");return n&&(i=i.filter(n)),!n&&l||(n=i.children().not("["+a.math_dataAttrib+"=ignore]"),r=(n=a.math_ignore.length?n.filter(function(){return-1===v.inArray(x.getCellIndex(v(this)),a.math_ignore)}):n).not(e).map(function(){return x.processText(t,v(this))}).get()),r},getColumn:function(t,e,n,a){var r,i,l,o,h,s,d=t.widgetOptions,u=[],c=e.closest("tr"),g=d.math_dataAttrib,f="["+g+"=ignore]",m=d.filter_filteredRow||"filtered",p=x.getCellIndex(e),_=t.$table.children("tbody").children(),b=["["+g+"^=above]","["+g+"^=below]","["+g+"^=col]","["+g+"^=all]"];if("above"===n)for(r=o=_.index(c);0<=r;)s=(l=_.eq(r)).children().filter(b[0]).length,i=(l=a?l.filter(a):l).children().filter(function(){return x.getCellIndex(v(this))===p}),((a||!l.hasClass(m))&&l.not(f).length&&r!==o||s&&r!==o)&&(s?r=0:i.length&&i.not(f).length&&(u[u.length]=x.processText(t,i))),r--;else if("below"===n)for(o=_.length,r=_.index(c)+1;r<o&&!(l=_.eq(r)).children().filter(b[1]).length;r++)i=(l=a?l.filter(a):l).children().filter(function(){return x.getCellIndex(v(this))===p}),(a||!l.hasClass(m))&&l.not(f).length&&i.length&&i.not(f)&&(u[u.length]=x.processText(t,i));else for(o=(h=_.not(f)).length,r=0;r<o;r++)l=h.eq(r),i=(l=a?l.filter(a):l).children().filter(function(){return x.getCellIndex(v(this))===p}),!a&&l.hasClass(m)||!i.not(b.join(",")).length||i.is(e)||!i.not(f).length||(u[u.length]=x.processText(t,i));return u},getAll:function(t,e){for(var n,a,r,i,l,o,h=[],s=t.widgetOptions,d=s.math_dataAttrib,u="["+d+"=ignore]",c=s.filter_filteredRow||"filtered",g=t.$table.children("tbody").children().not(u),f=g.length,m=0;m<f;m++)if(r=g.eq(m),e&&(r=r.filter(e)),e||!r.hasClass(c))for(o=(i=r.children().not(u)).length,l=0;l<o;l++)n=i.eq(l),a=x.getCellIndex(n),!n.filter("["+d+"]").length&&v.inArray(a,s.math_ignore)<0&&(h[h.length]=x.processText(t,n));return h},setColumnIndexes:function(t){var e=t.$table,r=1,e=e.children("tbody").children().filter(function(){var t,e,n=v(this),a=0<n.children("[colspan]").length;if(1<r?(r--,a=!0):r<1&&(r=1),0<n.children("[rowspan]").length)for(t=this.cells,e=0;e<t.length;e++)r=Math.max(t[e].rowSpan,r);return a});_.computeColumnIndex(e,t)},getCellIndex:function(t){var e=t.attr("data-column");return void 0===e?t[0].cellIndex:parseInt(e,10)},recalculate:function(n,t,e){if(n&&(!t.math_isUpdating||e)){var a,r,i,l,o,h=!1,s={};for((n.debug||t.math_debug)&&(a=new Date),e&&x.setColumnIndexes(n),t.math_dataAttrib="data-"+(t.math_data||"math"),r=t.math_dataAttrib,i=n.$tbodies.children("tr").children("["+r+"]"),h=x.mathType(n,i,t.math_priority)||h,i=n.$table.children("."+n.cssInfoBlock+", tfoot").children("tr").children("["+r+"]"),x.mathType(n,i,t.math_priority),o=(i=n.$table.children().children("tr").children("["+r+"^=all]")).length,l=0;l<o;l++){var d=i.eq(l),u=d.attr(r+"-filter")||t.math_rowFilter;s[u]=s[u]?s[u].add(d):d}v.each(s,function(t,e){h=x.mathType(n,e,["all"],t)||h}),h?(t.math_isUpdating=!0,(n.debug||t.math_debug)&&console[console.group?"group":"log"]("Math widget updating the cache after recalculation"),_.updateCache(n,function(){x.updateComplete(n),e||"function"!=typeof t.math_completed||t.math_completed(n),(n.debug||t.math_debug)&&console.log("Math widget update completed"+_.benchmark(a))})):(e||"function"!=typeof t.math_completed||t.math_completed(n),(n.debug||t.math_debug)&&console.log("Math widget found no changes in data"+_.benchmark(a)))}},updateComplete:function(t){var e=t.widgetOptions;e.math_isUpdating&&(t.debug||e.math_debug)&&console.groupEnd&&console.groupEnd(),e.math_isUpdating=!1},mathType:function(s,d,t,u){var c,g,f,m,p;return!!d.length&&(g=!1,f=s.widgetOptions,m=f.math_dataAttrib,p=_.equations,"all"===t[0]&&(c=x.getAll(s,u)),(s.debug||f.math_debug)&&console[console.group?"group":"log"]("Tablesorter Math widget recalculation"),v.each(t,function(t,e){var n,a,r,i,l,o=d.filter("["+m+"^="+e+"]"),h=o.length;if(h){for((s.debug||f.math_debug)&&console[console.group?"group":"log"](e),n=0;n<h;n++)(l=o.eq(n)).parent().hasClass(f.filter_filteredRow||"filtered")||(u=l.attr(m+"-filter")||f.math_rowFilter,r=(l.attr(m)||"").replace(e+"-",""),a="row"===e?x.getRow(s,l,u):"all"===e?c:x.getColumn(s,l,e,u),p[r]&&(a.length?(i=p[r](a,s),(s.debug||f.math_debug)&&console.log(l.attr(m),u?'("'+u+'")':"",a,"=",i)):i=x.invalid(s,r,"mean"===r?0:"undef"),g=x.output(l,s,i,a)||g));(s.debug||f.math_debug)&&console.groupEnd&&console.groupEnd()}}),(s.debug||f.math_debug)&&console.groupEnd&&console.groupEnd(),g)},output:function(t,e,n,a){var r,i=e.widgetOptions,l=!1,o=t.html(),h=t.attr("data-"+i.math_data+"-mask")||i.math_mask,s=t.attr("data-"+i.math_data+"-target")||"",h=_.formatMask(h,n,i.math_prefix,i.math_suffix);return s&&(r=t.find(s)).length&&(t=r),!1!==(h="function"==typeof i.math_complete?i.math_complete(t,i,h,n,a):h)&&(l=o!==h,t.html(h)),!(l&&(!(r=t.closest("tbody")).length||r.hasClass(e.cssInfoBlock)||r.parent()[0]!==e.table))&&l}};_.formatMask=function(t,e,n,a){if(!t||isNaN(+e))return e;var r,i,l,o,h,s,d,u,c="",g=t.length,f=t.search(/[0-9\-\+#]/),m=0<f?t.substring(0,f):"",p=m;if(n&&(p=/\{content\}/.test(n||"")?(n||"").replace(/\{content\}/g,m||""):(n||"")+m),u=g-(n=t.split("").reverse().join("").search(/[0-9\-\+#]/)),u+="."===t.substring(u,u+1)?1:0,c=m=0<n?t.substring(u,g):"",a&&(c=/\{content\}/.test(a||"")?(a||"").replace(/\{content\}/g,m||""):m+(a||"")),n=(e="-"===(t=t.substring(f,u)).charAt(0)?-e:+e)<0?e=-e:0,a=(m=t.match(/[^\d\-\+#]/g))&&m[m.length-1]||".",r=m&&m[1]&&m[0]||",",t=t.split(a),e=+(e=e.toFixed(t[1]&&t[1].length))+"",f=t[1]&&t[1].lastIndexOf("0"),(!(o=e.split("."))[1]||o[1]&&o[1].length<=f)&&(e=(+e).toFixed(f+1)),m=t[0].split(r),t[0]=m.join(""),-1<(i=t[0]&&t[0].indexOf("0")))for(;o[0].length<t[0].length-i;)o[0]="0"+o[0];else 0==+o[0]&&(o[0]="");if((e=e.split("."))[0]=o[0],l=m[1]&&m[m.length-1].length){for(s="",d=(h=e[0]).length%l,g=h.length,u=0;u<g;u++)s+=h.charAt(u),!((u-d+1)%l)&&u<g-l&&(s+=r);e[0]=s}return e[1]=t[1]&&e[1]?a+e[1]:"",p+((n?"-":"")+e[0]+e[1])+c},_.equations={count:function(t){return t.length},sum:function(t){for(var e=t.length,n=0,a=0;a<e;a++)n+=t[a];return n},mean:function(t){return _.equations.sum(t)/t.length},median:function(t,e){var n,a=t.length;return 1<a?(t.sort(function(t,e){return t-e}),n=Math.floor(a/2),a%2?t[n]:(t[n-1]+t[n])/2):x.invalid(e,"median",1)},mode:function(t){for(var e,n,a={},r=1,i=[t[0]],l=0;l<t.length;l++)a[e=t[l]]=a[e]?a[e]+1:1,r<(n=a[e])?(i=[e],r=n):n===r&&(i[i.length]=e,r=n);return i.sort(function(t,e){return t-e})},max:function(t){return Math.max.apply(Math,t)},min:function(t){return Math.min.apply(Math,t)},range:function(t){var e=t.sort(function(t,e){return t-e});return e[t.length-1]-e[0]},variance:function(t,e,n){for(var a=_.equations.mean(t),r=0,i=t.length;i--;)r+=Math.pow(t[i]-a,2);return 0==(e=t.length-(e?0:1))?x.invalid(n,"variance",0):r/e},varp:function(t,e){return _.equations.variance(t,!0,e)},vars:function(t,e){return _.equations.variance(t,!1,e)},stdevs:function(t,e){t=_.equations.variance(t,!1,e);return Math.sqrt(t)},stdevp:function(t,e){t=_.equations.variance(t,!0,e);return Math.sqrt(t)}},_.addWidget({id:"math",priority:100,options:{math_data:"math",math_debug:!1,math_ignore:[],math_mask:"#,##0.00",math_complete:null,math_completed:function(){},math_priority:["row","above","below","col"],math_prefix:"",math_suffix:"",math_textAttr:"",math_none:"N/A",math_event:"recalculate",math_rowFilter:""},init:function(t,e,n,a){var r=(_.hasWidget(t,"filter")?"filterEnd":"updateComplete")+".tsmath";x.events+=(_.hasWidget(t,"pager")?"pagerComplete":"filterEnd")+".tsmath ",n.$table.off((x.events+"updateComplete.tsmath "+a.math_event).replace(/\s+/g," ")).on(x.events+a.math_event,function(t){var e;this.hasInitialized&&(e="tablesorter-initialized"===t.type,a.math_isUpdating&&!e||(/filter/.test(t.type)||e||x.setColumnIndexes(n),x.recalculate(n,a,e)))}).on(r,function(){setTimeout(function(){x.updateComplete(n)},40)}),a.math_isUpdating=!1,t.hasInitialized&&x.recalculate(n,a,!0)},remove:function(t,e,n,a){a||e.$table.off((x.events+" updateComplete.tsmath "+n.math_event).replace(/\s+/g," ")).children().children("tr").children("[data-"+n.math_data+"]").empty()}})}(jQuery);return jQuery;}));
```
