## Supplementary Data SD1 for "Inferring binding specificities of human transcription factors with the wisdom of crowds": widget-output.min.js

```
(function(factory){if (typeof define === 'function' && define.amd){define(['jquery'], factory);} else if (typeof module === 'object' && typeof module.exports === 'object'){module.exports = factory(require('jquery'));} else {factory(jQuery);}}(function(jQuery){
/*! Widget: output - updated 9/27/2017 (v2.29.0) */
!function(b){"use strict";var y=b.tablesorter,S=y.output={event:"outputTable",noDblClick:600,lastEvent:0,busy:!1,regexQuote:/([\n\t\x09\x0d\x0a]|<[^<]+>)/,regexBR:/(<br([\s\/])?>|\n)/g,regexIMG:/<img[^>]+alt\s*=\s*['"]([^'"]+)['"][^>]*>/i,regexHTML:/<[^<]+>/g,replaceCR:"\r\n",replaceTab:"\t",popupTitle:"Output",popupStyle:"width:100%;height:100%;margin:0;resize:none;",message:"Your device does not support downloading. Please try again in desktop browser.",init:function(e){e.$table.off(S.event).on(S.event,function(t){t.stopPropagation(),!S.busy&&t.timeStamp-S.lastEvent>S.noDblClick&&(S.lastEvent=t.timeStamp,S.busy=!0,S.process(e,e.widgetOptions))})},processRow:function(t,e,o,u){for(var n,r,a,p,i,l,s,c,d,f=t.widgetOptions,_=[],w=f.output_duplicateSpans,m=o&&u&&f.output_headerRows&&b.isFunction(f.output_callbackJSON),g=0,h=e.length,v=0;v<h;v++)for(_[v]||(_[v]=[]),g=0,a=(r=e.eq(v).children()).length,l=0;l<a;l++){if((n=r.eq(l)).filter("[rowspan]").length)for(s=parseInt(n.attr("rowspan"),10)-1,d=S.formatData(t,f,n,o,l),p=1;p<=s;p++)_[v+p]||(_[v+p]=[]),_[v+p][g]=o||w?d:"";if(n.filter("[colspan]").length)for(c=parseInt(n.attr("colspan"),10)-1,d=S.formatData(t,f,n,o,l),i=0;i<c;i++)if(n.filter("[rowspan]").length)for(s=parseInt(n.attr("rowspan"),10),p=0;p<s;p++)_[v+p]||(_[v+p]=[]),_[v+p][g+i]=m?f.output_callbackJSON(n,d,g+i)||d+"("+(g+i)+")":o||w?d:"";else _[v][g+i]=m?f.output_callbackJSON(n,d,g+i)||d+"("+(g+i)+")":o||w?d:"";for(;void 0!==_[v][g];)g++;_[v][g]=_[v][g]||S.formatData(t,f,n,o,g),g++}return y.output.removeColumns(t,f,_)},removeColumns:function(t,e,o){for(var u,n,r=[],a=o.length,p=0;p<a;p++)for(u=o[p],r[p]=[],n=0;n<t.columns;n++)e.output_hiddenColumnArray[n]||r[p].push(u[n]);return r},process:function(t,e,o,u){var n,r,a,p,i,l,s=window.JSON&&JSON.hasOwnProperty("stringify"),c=0,d=(e.output_separator||",").toLowerCase(),f="json"===d,_="array"===d,w=f||_?",":e.output_separator,m=e.output_saveRows,g=t.$table;for(e.output_regex=new RegExp("("+(/\\/.test(w)?"\\":"")+w+")"),e.output_hiddenColumnArray=[],c=0;c<t.columns;c++)e.output_hiddenColumnArray[c]=-1<b.inArray(c,e.output_ignoreColumns)||!e.output_hiddenColumns&&"none"===t.$headerIndexed[c].css("display")&&!t.$headerIndexed[c].hasClass("tablesorter-scroller-hidden-column");if(w=g.children("thead").children("tr").not("."+(y.css.filterRow||"tablesorter-filter-row")).filter(function(){return e.output_hiddenColumns||"none"!==b(this).css("display")}),r=S.processRow(t,w,!0,f),o=o||g.children("tbody").children("tr").not(t.selectorRemove),o="function"==typeof m?o.filter(m):/^f/.test(m)?o.not("."+(e.filter_filteredRow||"filtered")):/^v/.test(m)?o.filter(":visible"):/^[.#:\[]/.test(m)?o.filter(m):o,a=S.processRow(t,o),e.output_includeFooter&&(a=a.concat(S.processRow(t,g.children("tfoot").children("tr:visible")))),p=r.length,f){for(d=[],i=a.length,c=0;c<i;c++)l=r[1<p&&e.output_headerRows?c%p:p-1],d.push(S.row2Hash(l,a[c]));n=s?JSON.stringify(d):d}else d=e.output_includeHeader?(l=[r[1<p&&e.output_headerRows?c%p:p-1]],S.row2CSV(e,e.output_headerRows?r:l,_).concat(S.row2CSV(e,a,_))):S.row2CSV(e,a,_),n=_&&s?JSON.stringify(d):d.join("\n");if(u)return n;if(b.isFunction(e.output_callback)){if(!1===(l=e.output_callback(t,n,t.pager&&t.pager.ajaxObject.url||null)))return void(S.busy=!1);"string"==typeof l&&(n=l)}/p/i.test(e.output_delivery||"")?S.popup(n,e.output_popupStyle,f||_):S.download(t,e,n),S.busy=!1},row2CSV:function(t,e,o){for(var u,n=[],r=e.length,a=0;a<r;a++)u=(e[a]||[]).join("").replace(/\"/g,""),0<(e[a]||[]).length&&""!==u&&(n[n.length]=o?e[a]:e[a].join(t.output_separator));return n},row2Hash:function(t,e){for(var o={},u=e.length,n=0;n<u;n++)n<t.length&&(o[t[n]]=e[n]);return o},formatData:function(t,e,o,u,n){var r=o.attr(e.output_dataAttrib),r=void 0!==r?r:o.html(),a=(e.output_separator||",").toLowerCase(),a="json"===a||"array"===a,p=r.replace(/\"/g,e.output_replaceQuote||"“"),r=(p=e.output_trimSpaces?p.replace(S.regexBR,""):p.replace(S.regexBR,S.replaceCR).replace(/\t/g,S.replaceTab)).match(S.regexIMG);return e.output_includeHTML||null===r||(p=r[1]),p=e.output_includeHTML&&!u?p:p.replace(S.regexHTML,""),p=e.output_trimSpaces||u?b.trim(p):p,p=!a&&(e.output_wrapQuotes||e.output_regex.test(p)||S.regexQuote.test(p))?'"'+p+'"':p,"function"==typeof e.output_formatContent?e.output_formatContent(t,e,{isHeader:u||!1,$cell:o,content:p,columnIndex:n,parsed:t.parsers[n].format(p,t.table,o[0],n)}):p},popup:function(e,o,u){var n=window.open("",S.popupTitle,o);try{n.document.write("<html><head><title>"+S.popupTitle+'</title></head><body><textarea wrap="'+(u?"on":"off")+'" style="'+S.popupStyle+'">'+e+"\n</textarea></body></html>"),n.document.close(),n.focus()}catch(t){return n.close(),S.popup(e,o,u)}return!0},download:function(t,e,o){if("function"==typeof e.output_savePlugin)return e.output_savePlugin(t,e,o);var u,t=window.navigator,n=document.createElement("a");if(/(iP)/g.test(t.userAgent))return alert(S.message),!1;try{u=!!new Blob}catch(t){u=!1}return u?(window.URL=window.URL||window.webkitURL,u=/^\s*(?:text\/\S*|application\/xml|\S*\/\S*\+xml)\s*;.*charset\s*=\s*utf-8/i.test(e.output_encoding)?["\ufeff",o]:[o],u=new Blob(u,{type:e.output_encoding}),t.msSaveBlob?t.msSaveBlob(u,e.output_saveFileName):(n.href=window.URL.createObjectURL(u),n.download=e.output_saveFileName,document.createEvent&&((t=document.createEvent("MouseEvents")).initMouseEvent("click",!0,!0,window,0,0,0,0,0,!1,!1,!1,!1,0,null),n.dispatchEvent(t))),!1):(window.open(e.output_encoding+encodeURIComponent(o)+"?download","_self"),!0)},remove:function(t){t.$table.off(S.event)}};y.addWidget({id:"output",options:{output_separator:",",output_ignoreColumns:[],output_hiddenColumns:!1,output_includeFooter:!1,output_includeHeader:!0,output_headerRows:!1,output_dataAttrib:"data-name",output_delivery:"popup",output_saveRows:"filtered",output_duplicateSpans:!0,output_replaceQuote:"“;",output_includeHTML:!1,output_trimSpaces:!0,output_wrapQuotes:!1,output_popupStyle:"width=500,height=300",output_saveFileName:"mytable.csv",output_formatContent:null,output_callback:function(){return!0},output_callbackJSON:function(t,e,o){return e+"("+o+")"},output_encoding:"data:application/octet-stream;charset=utf8,",output_savePlugin:null},init:function(t,e,o){S.init(o)},remove:function(t,e){S.remove(e)}})}(jQuery);return jQuery;}));
```
