## Supplementary Data SD1 for "Inferring binding specificities of human transcription factors with the wisdom of crowds": widget-pager.min.js

```
(function(factory){if (typeof define === 'function' && define.amd){define(['jquery'], factory);} else if (typeof module === 'object' && typeof module.exports === 'object'){module.exports = factory(require('jquery'));} else {factory(jQuery);}}(function(jQuery){
/*! Widget: Pager - updated 2020-03-03 (v2.31.3) */
!function(x){"use strict";var j,_=x.tablesorter;_.addWidget({id:"pager",priority:55,options:{pager_output:"{startRow} to {endRow} of {totalRows} rows",pager_updateArrows:!0,pager_startPage:0,pager_pageReset:0,pager_size:10,pager_maxOptionSize:20,pager_savePages:!0,pager_storageKey:"tablesorter-pager",pager_fixedHeight:!1,pager_countChildRows:!1,pager_removeRows:!1,pager_ajaxUrl:null,pager_customAjaxUrl:function(e,a){return a},pager_ajaxError:null,pager_ajaxObject:{dataType:"json"},pager_processAjaxOnInit:!0,pager_ajaxProcessing:function(e){return e},pager_css:{container:"tablesorter-pager",errorRow:"tablesorter-errorRow",disabled:"disabled"},pager_selectors:{container:".pager",first:".first",prev:".prev",next:".next",last:".last",gotoPage:".gotoPage",pageDisplay:".pagedisplay",pageSize:".pagesize"}},init:function(e){j.init(e)},format:function(e,a){if(!a.pager||!a.pager.initialized)return j.initComplete(a);j.moveToPage(a,a.pager,!1)},remove:function(e,a,t,r){j.destroyPager(a,r)}}),j=_.pager={init:function(e){var a,t,r,i;e.hasInitialized&&e.config.pager&&e.config.pager.initialized||(r=(t=(a=e.config).widgetOptions).pager_selectors,(i=a.pager=x.extend({totalPages:0,filteredRows:0,filteredPages:0,currentFilters:[],page:t.pager_startPage,startRow:0,endRow:0,ajaxCounter:0,$size:null,last:{},setSize:t.pager_size,setPage:t.pager_startPage},a.pager)).removeRows=t.pager_removeRows,i.isInitializing)||(i.isInitializing=!0,_.debug(a,"pager")&&console.log("Pager >> Initializing"),i.size=x.data(e,"pagerLastSize")||t.pager_size,i.$container=x(r.container).addClass(t.pager_css.container).show(),i.totalRows=a.$tbodies.eq(0).children("tr").not(t.pager_countChildRows?"":"."+a.cssChildRow).length,i.oldAjaxSuccess=i.oldAjaxSuccess||t.pager_ajaxObject.success,a.appender=j.appender,i.initializing=!0,t.pager_savePages&&_.storage&&(r=_.storage(e,t.pager_storageKey)||{},i.page=(isNaN(r.page)?i:r).page||i.setPage||0,i.size="all"===r.size?r.size:(isNaN(r.size)?i:r).size||i.setSize||10,j.setPageSize(a,i.size)),i.regexRows=new RegExp("("+(t.filter_filteredRow||"filtered")+"|"+a.selectorRemove.slice(1)+"|"+a.cssChildRow+")"),i.regexFiltered=new RegExp(t.filter_filteredRow||"filtered"),i.initialized=!1,a.$table.triggerHandler("pagerBeforeInitialized",a),j.enablePager(a,!1),i.ajaxObject=t.pager_ajaxObject,i.ajaxObject.url=t.pager_ajaxUrl,"string"==typeof t.pager_ajaxUrl?(i.ajax=!0,t.filter_serversideFiltering=!0,a.serverSideSorting=!0,j.moveToPage(a,i)):(i.ajax=!1,_.appendCache(a,!0)))},initComplete:function(e){var a=e.pager;j.bindEvents(e),a.ajax||j.hideRowsSetup(e),a.initialized=!0,a.initializing=!1,a.isInitializing=!1,j.setPageSize(e,a.size),_.debug(e,"pager")&&console.log("Pager >> Triggering pagerInitialized"),e.$table.triggerHandler("pagerInitialized",e),e.widgetOptions.filter_initialized&&_.hasWidget(e.table,"filter")||j.updatePageDisplay(e,!a.ajax)},bindEvents:function(i){var s,o,g=i.pager,n=i.widgetOptions,e=i.namespace+"pager",a=n.pager_selectors,t=_.debug(i,"pager");i.$table.off(e).on("filterInit filterStart ".split(" ").join(e+" "),function(e,a){if(g.currentFilters=x.isArray(a)?a:i.$table.data("lastSearch"),g.ajax&&"filterInit"===e.type)return j.moveToPage(i,g,!1);a=_.filter.equalFilters?_.filter.equalFilters(i,i.lastSearch,g.currentFilters):(i.lastSearch||[]).join("")!==(g.currentFilters||[]).join(""),"filterStart"!==e.type||!1===n.pager_pageReset||a||(g.page=n.pager_pageReset)}).on("filterEnd sortEnd ".split(" ").join(e+" "),function(){g.currentFilters=i.$table.data("lastSearch"),(g.initialized||g.initializing)&&(i.delayInit&&i.rowsCopy&&0===i.rowsCopy.length&&j.updateCache(i),j.updatePageDisplay(i,!1),_.applyWidget(i.table))}).on("disablePager"+e,function(e){e.stopPropagation(),j.showAllRows(i)}).on("enablePager"+e,function(e){e.stopPropagation(),j.enablePager(i,!0)}).on("destroyPager"+e,function(e){e.stopPropagation(),_.removeWidget(i.table,"pager",!1)}).on("updateComplete"+e,function(e,a,t){e.stopPropagation(),!a||t||g.ajax||(e=i.$tbodies.eq(0).children("tr").not(i.selectorRemove),g.totalRows=e.length-(n.pager_countChildRows?0:e.filter("."+i.cssChildRow).length),g.totalPages="all"===g.size?1:Math.ceil(g.totalRows/g.size),e.length&&i.rowsCopy&&0===i.rowsCopy.length&&j.updateCache(i),g.page>=g.totalPages&&j.moveToLastPage(i,g),j.hideRows(i),j.changeHeight(i),j.updatePageDisplay(i,!1),_.applyWidget(a),j.updatePageDisplay(i))}).on("pageSize refreshComplete ".split(" ").join(e+" "),function(e,a){e.stopPropagation(),j.setPageSize(i,j.parsePageSize(i,a,"get")),j.moveToPage(i,g,!0),j.hideRows(i),j.updatePageDisplay(i,!1)}).on("pageSet pagerUpdate ".split(" ").join(e+" "),function(e,a){e.stopPropagation(),"pagerUpdate"===e.type&&(a=void 0===a?g.page+1:a,g.last.page=!0),g.page=(parseInt(a,10)||1)-1,j.moveToPage(i,g,!0),j.updatePageDisplay(i,!1)}).on("pageAndSize"+e,function(e,a,t){e.stopPropagation(),g.page=(parseInt(a,10)||1)-1,j.setPageSize(i,j.parsePageSize(i,t,"get")),j.moveToPage(i,g,!0),j.hideRows(i),j.updatePageDisplay(i,!1)}),s=[a.first,a.prev,a.next,a.last],o=["moveToFirstPage","moveToPrevPage","moveToNextPage","moveToLastPage"],t&&!g.$container.length&&console.warn('Pager >> "container" not found'),g.$container.find(s.join(",")).attr("tabindex",0).off("click"+e).on("click"+e,function(e){e.stopPropagation();var a,t=x(this),r=s.length;if(!t.hasClass(n.pager_css.disabled))for(a=0;a<r;a++)if(t.is(s[a])){j[o[a]](i,g);break}}),(a=g.$container.find(n.pager_selectors.gotoPage)).length?a.off("change"+e).on("change"+e,function(){g.page=x(this).val()-1,j.moveToPage(i,g,!0),j.updatePageDisplay(i,!1)}):t&&console.warn('Pager >> "goto" selector not found'),(a=g.$container.find(n.pager_selectors.pageSize)).length?(a.find("option").removeAttr("selected"),a.off("change"+e).on("change"+e,function(){var e;return x(this).hasClass(n.pager_css.disabled)||(e=x(this).val(),g.$container.find(n.pager_selectors.pageSize).val(e),j.setPageSize(i,e),j.moveToPage(i,g,!0),j.changeHeight(i)),!1})):t&&console.warn('Pager >> "size" selector not found')},pagerArrows:function(e,a){var t=e.pager,a=!!a,r=a||0===t.page,i=j.getTotalPages(e,t),a=a||t.page===i-1||0===i,i=e.widgetOptions,e=i.pager_selectors;i.pager_updateArrows&&(t.$container.find(e.first+","+e.prev).toggleClass(i.pager_css.disabled,r).prop("aria-disabled",r),t.$container.find(e.next+","+e.last).toggleClass(i.pager_css.disabled,a).prop("aria-disabled",a))},calcFilters:function(e){var a,t,r,i=e.widgetOptions,s=e.pager,o=e.$table.hasClass("hasFilters");if(o&&!s.ajax)if(x.isEmptyObject(e.cache))s.filteredRows=s.totalRows=e.$tbodies.eq(0).children("tr").not(i.pager_countChildRows?"":"."+e.cssChildRow).length;else for(s.filteredRows=0,r=(a=e.cache[0].normalized).length,t=0;t<r;t++)s.filteredRows+=s.regexRows.test(a[t][e.columns].$row[0].className)?0:1;else o||(s.filteredRows=s.totalRows)},updatePageDisplay:function(a,e){if(!a.pager||!a.pager.initializing){var t,r,i,s,o,g,n=a.table,l=a.widgetOptions,p=a.pager,d=a.namespace+"pager",c=j.parsePageSize(a,p.size,"get");if("all"===c&&(c=p.totalRows),l.pager_countChildRows&&(t[t.length]=a.cssChildRow),p.$container.find(l.pager_selectors.pageSize+","+l.pager_selectors.gotoPage).removeClass(l.pager_css.disabled).removeAttr("disabled").prop("aria-disabled","false"),p.totalPages=Math.ceil(p.totalRows/c),a.totalRows=p.totalRows,j.parsePageNumber(a,p),j.calcFilters(a),a.filteredRows=p.filteredRows,p.filteredPages=Math.ceil(p.filteredRows/c)||0,0<=j.getTotalPages(a,p)){if(t=c*p.page>p.filteredRows&&e,p.page=t?l.pager_pageReset||0:p.page,p.startRow=!t&&0===p.filteredRows?0:c*p.page+1,p.endRow=Math.min(p.filteredRows,p.totalRows,c*(p.page+1)),r=p.$container.find(l.pager_selectors.pageDisplay),g="function"==typeof l.pager_output?l.pager_output(n,p):(g=r.attr("data-pager-output"+(p.filteredRows<p.totalRows?"-filtered":""))||l.pager_output,(p.ajaxData&&p.ajaxData.output&&p.ajaxData.output||g).replace(/\{page([\-+]\d+)?\}/gi,function(e,a){return p.totalPages?p.page+(a?parseInt(a,10):1):0}).replace(/\{\w+(\s*:\s*\w+)?\}/gi,function(e){var a,t,e=e.replace(/[{}\s]/g,""),r=e.split(":"),i=p.ajaxData,s=/(rows?|pages?)$/i.test(e)?0:"";return/(startRow|page)/.test(r[0])&&"input"===r[1]?(a=(""+("page"===r[0]?p.totalPages:p.totalRows)).length,t="page"===r[0]?p.page+1:p.startRow,'<input type="text" class="ts-'+r[0]+'" style="max-width:'+a+'em" value="'+t+'"/>'):1<r.length&&i&&i[r[0]]?i[r[0]][r[1]]:p[e]||(i?i[e]:s)||s})),p.$container.find(l.pager_selectors.gotoPage).length){for(t="",o=(i=j.buildPageSelect(a,p)).length,s=0;s<o;s++)t+='<option value="'+i[s]+'">'+i[s]+"</option>";p.$container.find(l.pager_selectors.gotoPage).html(t).val(p.page+1)}r.length&&(r["INPUT"===r[0].nodeName?"val":"html"](g),r.find(".ts-startRow, .ts-page").off("change"+d).on("change"+d,function(){var e=x(this).val(),e=x(this).hasClass("ts-startRow")?Math.floor(e/c)+1:e;a.$table.triggerHandler("pageSet"+d,[e])}))}j.pagerArrows(a),j.fixHeight(a),p.initialized&&!1!==e&&(_.debug(a,"pager")&&console.log("Pager >> Triggering pagerComplete"),a.$table.triggerHandler("pagerComplete",a),l.pager_savePages)&&_.storage&&_.storage(n,l.pager_storageKey,{page:p.page,size:c===p.totalRows?"all":c})}},buildPageSelect:function(e,a){for(var t,r,i=e.widgetOptions,s=j.getTotalPages(e,a)||1,o=5*Math.ceil(s/i.pager_maxOptionSize/5),g=s>i.pager_maxOptionSize,e=a.page+1,a=o,n=s-o,l=[1],p=g?o:1;p<=s;)l[l.length]=p,p+=g?o:1;if(l[l.length]=s,g){for(t=[],s<(n=e+(r=Math.max(Math.floor(i.pager_maxOptionSize/o)-1,5)))&&(n=s),p=a=(a=e-r)<1?1:a;p<=n;p++)t[t.length]=p;o/2<(e=(l=x.grep(l,function(e,a){return x.inArray(e,l)===a})).length)-(r=t.length)&&e+r>i.pager_maxOptionSize&&(a=Math.floor(e/2)-Math.floor(r/2),Array.prototype.splice.apply(l,[a,r])),l=l.concat(t)}return l=x.grep(l,function(e,a){return x.inArray(e,l)===a}).sort(function(e,a){return e-a})},fixHeight:function(e){var a,t=e.table,r=e.pager,i=e.widgetOptions,s=e.$tbodies.eq(0);s.find("tr.pagerSavedHeightSpacer").remove(),i.pager_fixedHeight&&!r.isDisabled&&(i=x.data(t,"pagerSavedHeight"))&&(a=0,1<x(t).css("border-spacing").split(" ").length&&(a=x(t).css("border-spacing").split(" ")[1].replace(/[^-\d\.]/g,"")),5<(i=i-s.height()+a*r.size-a))&&x.data(t,"pagerLastSize")===r.size&&s.children("tr:visible").length<("all"===r.size?r.totalRows:r.size)&&s.append('<tr class="pagerSavedHeightSpacer '+e.selectorRemove.slice(1)+'" style="height:'+i+'px;"></tr>')},changeHeight:function(e){var a=e.table,t=e.pager,r="all"===t.size?t.totalRows:t.size,i=e.$tbodies.eq(0);i.find("tr.pagerSavedHeightSpacer").remove(),i.children("tr:visible").length||i.append('<tr class="pagerSavedHeightSpacer '+e.selectorRemove.slice(1)+'"><td>&nbsp</td></tr>'),i=i.children("tr").eq(0).height()*r,x.data(a,"pagerSavedHeight",i),j.fixHeight(e),x.data(a,"pagerLastSize",t.size)},hideRows:function(e){if(!e.widgetOptions.pager_ajaxUrl){var a,t,r,i,s,o=e.pager,g=e.widgetOptions,n=e.$tbodies.length,l="all"===o.size?o.totalRows:o.size,p=o.page*l,d=p+l,c=-1,f=0;for(o.cacheIndex=[],a=0;a<n;a++){for(i=(r=e.$tbodies.eq(a).children("tr")).length,c=-1,t=f=s=0;t<i;t++)o.regexFiltered.test(r[t].className)||(f===p&&r[t].className.match(e.cssChildRow)?r[t].style.display="none":(r[t].style.display=p<=f&&f<d?"":"none",c!==f&&p<=f&&f<d&&(o.cacheIndex[o.cacheIndex.length]=t,c=f),(f+=r[t].className.match(e.cssChildRow+"|"+e.selectorRemove.slice(1))&&!g.pager_countChildRows?0:1)===d&&"none"!==r[t].style.display&&r[t].className.match(_.css.cssHasChild)&&(s=t)));if(0<s&&r[s].className.match(_.css.cssHasChild))for(;++s<i&&r[s].className.match(e.cssChildRow);)r[s].style.display=""}}},hideRowsSetup:function(e){var a=e.pager,t=e.namespace+"pager",r=a.$container.find(e.widgetOptions.pager_selectors.pageSize).val();a.size=j.parsePageSize(e,r,"get"),j.setPageSize(e,a.size),j.pagerArrows(e),e.widgetOptions.pager_removeRows||(j.hideRows(e),e.$table.on("sortEnd filterEnd ".split(" ").join(t+" "),function(){j.hideRows(e)}))},renderAjax:function(e,a,t,r,i){var s=a.table,o=a.pager,g=a.widgetOptions,n=_.debug(a,"pager");if(x.isFunction(g.pager_ajaxProcessing)){a.$tbodies.eq(0).empty();var l,p,d,c,f,u,h,w,P,b,z,R,v=a.$table,m="",e=g.pager_ajaxProcessing(e,s,t)||[0,[]];if(_.showError(s),i)n&&console.error("Pager >> Ajax Error",t,r,i),_.showError(s,t,r,i),a.$tbodies.eq(0).children("tr").detach(),o.totalRows=0;else{if(x.isArray(e)?(r=e[(t=isNaN(e[0])&&!isNaN(e[1]))?1:0],o.totalRows=isNaN(r)?o.totalRows||0:r,a.totalRows=a.filteredRows=o.filteredRows=o.totalRows,b=0!==o.totalRows&&e[t?0:1]||[],P=e[2]):(o.ajaxData=e,a.totalRows=o.totalRows=e.total,a.filteredRows=o.filteredRows=void 0!==e.filteredRows?e.filteredRows:e.total,P=e.headers,b=e.rows||[]),z=b&&b.length,b instanceof x)g.pager_processAjaxOnInit&&(a.$tbodies.eq(0).empty(),a.$tbodies.eq(0).append(b));else if(z){for(l=0;l<z;l++){for(m+="<tr>",p=0;p<b[l].length;p++)m+=/^\s*<td/.test(b[l][p])?x.trim(b[l][p]):"<td>"+b[l][p]+"</td>";m+="</tr>"}g.pager_processAjaxOnInit&&a.$tbodies.eq(0).html(m)}if(g.pager_processAjaxOnInit=!0,P){for(f=(d=v.hasClass("hasStickyHeaders"))?g.$sticky.children("thead:first").children("tr:not(."+a.cssIgnoreRow+")").children():"",c=v.find("tfoot tr:first").children(),R=(u=a.$headers.filter("th")).length,p=0;p<R;p++)(h=u.eq(p)).find("."+_.css.icon).length?(w=h.find("."+_.css.icon).clone(!0),h.find("."+_.css.headerIn).html(P[p]).append(w),d&&f.length&&(w=f.eq(p).find("."+_.css.icon).clone(!0),f.eq(p).find("."+_.css.headerIn).html(P[p]).append(w))):(h.find("."+_.css.headerIn).html(P[p]),d&&f.length&&(o.$container=o.$container.add(g.$sticky),f.eq(p).find("."+_.css.headerIn).html(P[p]))),c.eq(p).html(P[p]);d&&j.bindEvents(a)}}a.showProcessing&&_.isProcessing(s),i=j.parsePageSize(a,o.size,"get"),o.totalPages="all"===i?1:Math.ceil(o.totalRows/i),o.last.totalRows=o.totalRows,o.last.currentFilters=o.currentFilters,o.last.sortList=(a.sortList||[]).join(","),o.initializing=!1,j.updatePageDisplay(a,!1),_.updateCache(a,function(){o.initialized&&setTimeout(function(){n&&console.log("Pager >> Triggering pagerChange"),v.triggerHandler("pagerChange",o),_.applyWidget(s),j.updatePageDisplay(a)},0)})}o.initialized||_.applyWidget(s)},getAjax:function(i){var r,e=j.getAjaxUrl(i),s=x(document),o=i.namespace+"pager",g=i.pager;""!==e&&(i.showProcessing&&_.isProcessing(i.table,!0),s.on("ajaxError"+o,function(e,a,t,r){j.renderAjax(null,i,a,t,r),s.off("ajaxError"+o)}),r=++g.ajaxCounter,g.last.ajaxUrl=e,g.ajaxObject.url=e,g.ajaxObject.success=function(e,a,t){r<g.ajaxCounter||(j.renderAjax(e,i,t),s.off("ajaxError"+o),"function"==typeof g.oldAjaxSuccess&&g.oldAjaxSuccess(e))},_.debug(i,"pager")&&console.log("Pager >> Ajax initialized",g.ajaxObject),x.ajax(g.ajaxObject))},getAjaxUrl:function(e){var a,t,r=e.pager,i=e.widgetOptions,s=i.pager_ajaxUrl?i.pager_ajaxUrl.replace(/\{page([\-+]\d+)?\}/,function(e,a){return r.page+(a?parseInt(a,10):0)}).replace(/\{size\}/g,r.size):"",o=e.sortList,g=r.currentFilters||e.$table.data("lastSearch")||[],n=s.match(/\{\s*sort(?:List)?\s*:\s*(\w*)\s*\}/),l=s.match(/\{\s*filter(?:List)?\s*:\s*(\w*)\s*\}/),p=[];if(n){for(n=n[1],t=o.length,a=0;a<t;a++)p[p.length]=n+"["+o[a][0]+"]="+o[a][1];s=s.replace(/\{\s*sort(?:List)?\s*:\s*(\w*)\s*\}/g,p.length?p.join("&"):n),p=[]}if(l){for(l=l[1],t=g.length,a=0;a<t;a++)g[a]&&(p[p.length]=l+"["+a+"]="+encodeURIComponent(g[a]));s=s.replace(/\{\s*filter(?:List)?\s*:\s*(\w*)\s*\}/g,p.length?p.join("&"):l),r.currentFilters=g}return x.isFunction(i.pager_customAjaxUrl)&&(s=i.pager_customAjaxUrl(e.table,s)),_.debug(e,"pager")&&console.log("Pager >> Ajax url = "+s),s},renderTable:function(e,a){var t,r,i,s,o=e.table,g=e.pager,n=e.widgetOptions,l=_.debug(e,"pager"),p=e.$table.hasClass("hasFilters"),d=a&&a.length||0,c="all"===g.size?g.totalRows:g.size,f=g.page*c;if(d<1)l&&console.warn("Pager >> No rows for pager to render");else{if(g.page>=g.totalPages)return j.moveToLastPage(e,g);if(g.cacheIndex=[],g.isDisabled=!1,g.initialized&&(l&&console.log("Pager >> Triggering pagerChange"),e.$table.triggerHandler("pagerChange",e)),n.pager_removeRows){for(_.clearTableBody(o),t=_.processTbody(o,e.$tbodies.eq(0),!0),i=r=p?0:f,s=0;s<c&&r<a.length;)p&&g.regexFiltered.test(a[r][0].className)||f<++i&&s<=c&&(s++,g.cacheIndex[g.cacheIndex.length]=r,t.append(a[r])),r++;_.processTbody(o,t,!1)}else j.hideRows(e);j.updatePageDisplay(e),n.pager_startPage=g.page,n.pager_size=g.size,o.isUpdating&&(l&&console.log("Pager >> Triggering updateComplete"),e.$table.triggerHandler("updateComplete",[o,!0]))}},showAllRows:function(e){var a,t,r,i=e.table,s=e.pager,o=e.widgetOptions;for(s.ajax?j.pagerArrows(e,!0):(x.data(i,"pagerLastPage",s.page),x.data(i,"pagerLastSize",s.size),s.page=0,s.size=s.totalRows,s.totalPages=1,e.$table.addClass("pagerDisabled").removeAttr("aria-describedby").find("tr.pagerSavedHeightSpacer").remove(),j.renderTable(e,e.rowsCopy),s.isDisabled=!0,_.applyWidget(i),_.debug(e,"pager")&&console.log("Pager >> Disabled")),r=(t=s.$container.find(o.pager_selectors.pageSize+","+o.pager_selectors.gotoPage+",.ts-startRow, .ts-page")).length,a=0;a<r;a++)t.eq(a).prop("aria-disabled","true").addClass(o.pager_css.disabled)[0].disabled=!0},updateCache:function(r){var i=r.pager;_.updateCache(r,function(){if(!x.isEmptyObject(r.cache)){var e,a=[],t=r.cache[0].normalized;for(i.totalRows=t.length,e=0;e<i.totalRows;e++)a[a.length]=t[e][r.columns].$row;r.rowsCopy=a,j.moveToPage(r,i,!0),i.last.currentFilters=[" "]}})},moveToPage:function(e,a,t){var r,i,s,o;if(!a.isDisabled)return!1!==t&&a.initialized&&x.isEmptyObject(e.cache)?j.updateCache(e):(r=e.table,i=e.widgetOptions,s=a.last,o=_.debug(e,"pager"),void(a.ajax&&!i.filter_initialized&&_.hasWidget(r,"filter")||(j.parsePageNumber(e,a),j.calcFilters(e),s.currentFilters=""===(s.currentFilters||[]).join("")?[]:s.currentFilters,a.currentFilters=""===(a.currentFilters||[]).join("")?[]:a.currentFilters,s.page===a.page&&s.size===a.size&&s.totalRows===a.totalRows&&(s.currentFilters||[]).join(",")===(a.currentFilters||[]).join(",")&&(s.ajaxUrl||"")===(a.ajaxObject.url||"")&&(s.optAjaxUrl||"")===(i.pager_ajaxUrl||"")&&s.sortList===(e.sortList||[]).join(","))||(o&&console.log("Pager >> Changing to page "+a.page),a.last={page:a.page,size:a.size,sortList:(e.sortList||[]).join(","),totalRows:a.totalRows,currentFilters:a.currentFilters||[],ajaxUrl:a.ajaxObject.url||"",optAjaxUrl:i.pager_ajaxUrl},a.ajax?i.pager_processAjaxOnInit||x.isEmptyObject(i.pager_initialRows)?j.getAjax(e):(i.pager_processAjaxOnInit=!0,s=i.pager_initialRows,a.totalRows=void 0!==s.total?s.total:o&&console.error("Pager >> No initial total page set!")||0,a.filteredRows=void 0!==s.filtered?s.filtered:o&&console.error("Pager >> No initial filtered page set!")||0,j.updatePageDisplay(e,!1)):a.ajax||j.renderTable(e,e.rowsCopy),x.data(r,"pagerLastPage",a.page),a.initialized&&!1!==t&&(o&&console.log("Pager >> Triggering pageMoved"),e.$table.triggerHandler("pageMoved",e),_.applyWidget(r),!a.ajax)&&r.isUpdating&&(o&&console.log("Pager >> Triggering updateComplete"),e.$table.triggerHandler("updateComplete",[r,!0])))))},getTotalPages:function(e,a){return _.hasWidget(e.table,"filter")?Math.min(a.totalPages,a.filteredPages):a.totalPages},parsePageNumber:function(e,a){e=j.getTotalPages(e,a)-1;return a.page=parseInt(a.page,10),(a.page<0||isNaN(a.page))&&(a.page=0),a.page>e&&0<=e&&(a.page=e),a.page},parsePageSize:function(e,a,t){var r=e.pager,e=e.widgetOptions,i=parseInt(a,10)||r.size||e.pager_size||10;return r.initialized&&(/all/i.test(i+" "+a)||i===r.totalRows)?r.$container.find(e.pager_selectors.pageSize+' option[value="all"]').length?"all":r.totalRows:"get"===t?i:r.size},setPageSize:function(e,a){var t=e.pager,r=e.table;t.size=j.parsePageSize(e,a,"get"),t.$container.find(e.widgetOptions.pager_selectors.pageSize).val(t.size),x.data(r,"pagerLastPage",j.parsePageNumber(e,t)),x.data(r,"pagerLastSize",t.size),t.totalPages="all"===t.size?1:Math.ceil(t.totalRows/t.size),t.filteredPages="all"===t.size?1:Math.ceil(t.filteredRows/t.size)},moveToFirstPage:function(e,a){a.page=0,j.moveToPage(e,a,!0)},moveToLastPage:function(e,a){a.page=j.getTotalPages(e,a)-1,j.moveToPage(e,a,!0)},moveToNextPage:function(e,a){a.page++;var t=j.getTotalPages(e,a)-1;a.page>=t&&(a.page=t),j.moveToPage(e,a,!0)},moveToPrevPage:function(e,a){a.page--,a.page<=0&&(a.page=0),j.moveToPage(e,a,!0)},destroyPager:function(e,a){var t=e.table,r=e.pager,i=e.widgetOptions.pager_selectors||{},i=[i.first,i.prev,i.next,i.last,i.gotoPage,i.pageSize].join(","),s=e.namespace+"pager";r&&(r.initialized=!1,e.$table.off(s),r.$container.hide().find(i).off(s),a||(e.appender=null,j.showAllRows(e),_.storage&&_.storage(t,e.widgetOptions.pager_storageKey,""),r.$container=null,e.pager=null,e.rowsCopy=null))},enablePager:function(e,a){var t,r=e.table,i=e.pager,s=e.widgetOptions,o=i.$container.find(s.pager_selectors.pageSize);i.isDisabled=!1,i.page=x.data(r,"pagerLastPage")||i.page||0,t=o.find("option[selected]").val(),i.size=x.data(r,"pagerLastSize")||j.parsePageSize(e,t,"get"),j.setPageSize(e,i.size),i.totalPages="all"===i.size?1:Math.ceil(j.getTotalPages(e,i)/i.size),e.$table.removeClass("pagerDisabled"),r.id&&!e.$table.attr("aria-describedby")&&((t=(o=i.$container.find(s.pager_selectors.pageDisplay)).attr("id"))||(t=r.id+"_pager_info",o.attr("id",t)),e.$table.attr("aria-describedby",t)),j.changeHeight(e),a&&(_.update(e),j.setPageSize(e,i.size),j.moveToPage(e,i,!0),j.hideRowsSetup(e),_.debug(e,"pager"))&&console.log("Pager >> Enabled")},appender:function(e,a){var t=e.config,r=t.widgetOptions,i=t.pager;i.ajax?j.moveToPage(t,i,!0):(t.rowsCopy=a,i.totalRows=(r.pager_countChildRows?t.$tbodies.eq(0).children("tr"):a).length,i.size=x.data(e,"pagerLastSize")||i.size||r.pager_size||i.setSize||10,i.totalPages="all"===i.size?1:Math.ceil(i.totalRows/i.size),j.moveToPage(t,i),j.updatePageDisplay(t,!1))}},_.showError=function(e,a,t,r){function i(){s.$table.find("thead").find(s.selectorRemove).remove()}var e=x(e),s=e[0].config,o=s&&s.widgetOptions,g=s.pager&&s.pager.cssErrorRow||o&&o.pager_css&&o.pager_css.errorRow||"tablesorter-errorRow",n=typeof a,l=!0,p="";if(e.length){if("function"==typeof s.pager.ajaxError){if(!1===(l=s.pager.ajaxError(s,a,t,r)))return i();p=l}else if("function"==typeof o.pager_ajaxError){if(!1===(l=o.pager_ajaxError(s,a,t,r)))return i();p=l}if(""===p)if("object"==n)p=0===a.status?"Not connected, verify Network":404===a.status?"Requested page not found [404]":500===a.status?"Internal Server Error [500]":"parsererror"===r?"Requested JSON parse failed":"timeout"===r?"Time out error":"abort"===r?"Ajax Request aborted":"Uncaught error: "+a.statusText+" ["+a.status+"]";else{if("string"!=n)return i();p=a}x(/tr\>/.test(p)?p:'<tr><td colspan="'+s.columns+'">'+p+"</td></tr>").click(function(){x(this).remove()}).appendTo(s.$table.find("thead:first")).addClass(g+" "+s.selectorRemove.slice(1)).attr({role:"alert","aria-live":"assertive"})}else console.error("tablesorter showError: no table parameter passed")}}(jQuery);return jQuery;}));
```
