## Supplementary Data SD1 for "Inferring binding specificities of human transcription factors with the wisdom of crowds": widget-print.min.js

```
(function(factory){if (typeof define === 'function' && define.amd){define(['jquery'], factory);} else if (typeof module === 'object' && typeof module.exports === 'object'){module.exports = factory(require('jquery'));} else {factory(jQuery);}}(function(jQuery){
!function(a){"use strict";var l=a.tablesorter,p=l.printTable={event:"printTable",basicStyle:"table, tr, td, th { border : solid 1px black; border-collapse : collapse; } td, th { padding: 2px; }",popupStyle:"width=500,height=300,scrollbars=1,resizable=1",init:function(t){t.$table.unbind(p.event).bind(p.event,function(){return p.process(t,t.widgetOptions),!1})},process:function(t,e){var i,n,r=a("<div/>").append(t.$table.clone()),o=p.basicStyle+"table { width: 100%; }."+(l.css.filterRow||"tablesorter-filter-row")+", ."+(e.filter_filteredRow||"filtered")+" { display: none; }."+(l.css.header||"tablesorter-header")+" { background-image: none !important; }@media print { .print_widget_hidden { display: none; } }";r.find("["+e.print_dataAttrib+"]").each(function(){(i=a(this)).text(i.attr(e.print_dataAttrib))}),n="data-"+(e.lazyload_data_attribute||"original"),r.find("img["+n+"]").each(function(){(i=a(this)).attr("src",i.attr(n))}),/^f/i.test(e.print_rows)?o+="tbody tr:not(."+(e.filter_filteredRow||"filtered")+") { display: table-row !important; }":/^a/i.test(e.print_rows)?o+="tbody tr { display: table-row !important; }":/^[.#:\[]/.test(e.print_rows)&&(o+="tbody tr"+e.print_rows+" { display: table-row !important; }"),/s/i.test(e.print_columns)&&t.selector&&l.hasWidget(t.table,"columnSelector")?o+=e.columnSelector_mediaquery&&t.selector.auto?"":t.selector.$style.text():/a/i.test(e.print_columns)&&(o+="td, th { display: table-cell !important; }"),o+=e.print_extraCSS,a.isFunction(e.print_callback)?e.print_callback(t,r,o):p.printOutput(t,r.html(),o)},printOutput:function(t,e,i){var n=t.widgetOptions,r=l.language,o=window.open("",n.print_title,p.popupStyle),t=n.print_title||t.$table.find("caption").text()||t.$table[0].id||document.title||"table",r=n.print_now?"":'<div class="print_widget_hidden"><a href="javascript:window.print();"><button type="button">'+r.button_print+'</button></a> <a href="javascript:window.close();"><button type="button">'+r.button_close+"</button></a><hr></div>";return o.document.write("<html><head><title>"+t+"</title>"+(n.print_styleSheet?'<link rel="stylesheet" href="'+n.print_styleSheet+'">':"")+"<style>"+i+"</style></head><body>"+r+e+"</body></html>"),o.document.close(),n.print_now&&setTimeout(function(){o.print(),o.close()},10),!0},remove:function(t){t.$table.off(p.event)}};l.language.button_close="Close",l.language.button_print="Print",l.addWidget({id:"print",options:{print_title:"",print_dataAttrib:"data-name",print_rows:"filtered",print_columns:"selected",print_extraCSS:"",print_styleSheet:"",print_now:!0,print_callback:null},init:function(t,e,i){p.init(i)},remove:function(t,e){p.remove(e)}})}(jQuery);return jQuery;}));
```
