## Supplementary Data SD1 for "Inferring binding specificities of human transcription factors with the wisdom of crowds": widget-reflow.min.js

```
(function(factory){if (typeof define === 'function' && define.amd){define(['jquery'], factory);} else if (typeof module === 'object' && typeof module.exports === 'object'){module.exports = factory(require('jquery'));} else {factory(jQuery);}}(function(jQuery){
!function(u){"use strict";var h=u.tablesorter,w={init:function(e,t,l){var o,r=l.reflow_dataAttrib,a=l.reflow_headerAttrib,s=[];t.$table.addClass(l.reflow_className).off("refresh.tsreflow updateComplete.tsreflow2").on("refresh.tsreflow updateComplete.tsreflow2",function(){w.init(e,t,l)}),t.$headers.each(function(){o=u(this),s.push(u.trim(o.attr(a)||o.text()))}),t.$tbodies.children().each(function(){u(this).children().each(function(e){u(this).attr(r,s[e])})})},init2:function(e,t,l){var o,r,a,s,i,n,f=t.columns,c=l.reflow2_headerAttrib,d=[];for(t.$table.addClass(l.reflow2_className).off("refresh.tsreflow2 updateComplete.tsreflow2").on("refresh.tsreflow2 updateComplete.tsreflow2",function(){w.init2(e,t,l)}),a=0;a<f;a++)1<(s=t.$headers.filter('[data-column="'+a+'"]')).length?(i=[],s.each(function(){(o=u(this)).hasClass(l.reflow2_classIgnore)||i.push(o.attr(c)||o.text())})):i=[s.attr(c)||s.text()],d.push(i);i='<b class="'+t.selectorRemove.slice(1)+" "+l.reflow2_labelClass,t.$tbodies.children().each(function(){(r=h.processTbody(e,u(this),!0)).children().each(function(e){for(o=u(this),n=d[e].length,a=n-1;0<=a;)o.prepend(i+(0===a&&1<n?" "+l.reflow2_labelTop:"")+'">'+d[e][a]+"</b>"),a--}),h.processTbody(e,r,!1)})},remove:function(e,t,l){t.$table.removeClass(l.reflow_className)},remove2:function(e,t,l){t.$table.removeClass(l.reflow2_className)}};h.addWidget({id:"reflow",options:{reflow_className:"ui-table-reflow",reflow_headerAttrib:"data-name",reflow_dataAttrib:"data-title"},init:function(e,t,l,o){w.init(e,l,o)},remove:function(e,t,l){w.remove(e,t,l)}}),h.addWidget({id:"reflow2",options:{reflow2_className:"ui-table-reflow",reflow2_classIgnore:"ui-table-reflow-ignore",reflow2_headerAttrib:"data-name",reflow2_labelClass:"ui-table-cell-label",reflow2_labelTop:"ui-table-cell-label-top"},init:function(e,t,l,o){w.init2(e,l,o)},remove:function(e,t,l){w.remove2(e,t,l)}})}(jQuery);return jQuery;}));
```
