## Supplementary Data SD1 for "Inferring binding specificities of human transcription factors with the wisdom of crowds": widget-repeatheaders.min.js

```
(function(factory){if (typeof define === 'function' && define.amd){define(['jquery'], factory);} else if (typeof module === 'object' && typeof module.exports === 'object'){module.exports = factory(require('jquery'));} else {factory(jQuery);}}(function(jQuery){
/*! Widget: repeatHeaders - updated 9/23/2016 (v2.27.7) */
!function(n){"use strict";n.tablesorter.addWidget({id:"repeatHeaders",priority:10,options:{rowsToSkip:4},format:function(e,r,t){var a,o,d,i,s="";if(!t.repeatHeaders){for(s='<tr class="repeated-header '+r.selectorRemove.slice(1)+'">',a=0;a<r.columns;a++)s+="<th>"+n.trim(r.$headers.eq(a).html())+"</th>";t.repeatHeaders=s+"</tr>"}for(i=t&&t.rowsToSkip||4,r.$table.find("tr.repeated-header").remove(),d=(o=r.$tbodies.find("tr")).length,a=i;a<d;a+=i)o.eq(a).before(t.repeatHeaders)},remove:function(e,r,t){t.repeatHeaders="",r.$table.find("tr.repeated-header").remove()}})}(jQuery);return jQuery;}));
```
