## Supplementary Data SD1 for "Inferring binding specificities of human transcription factors with the wisdom of crowds": widget-resizable.min.js

```
(function(factory){if (typeof define === 'function' && define.amd){define(['jquery'], factory);} else if (typeof module === 'object' && typeof module.exports === 'object'){module.exports = factory(require('jquery'));} else {factory(jQuery);}}(function(jQuery){
/*! Widget: resizable - updated 2018-03-26 (v2.30.2) */
!function(d,t){"use strict";var b=d.tablesorter||{};d.extend(b.css,{resizableContainer:"tablesorter-resizable-container",resizableHandle:"tablesorter-resizable-handle",resizableNoSelect:"tablesorter-disableSelection",resizableStorage:"tablesorter-resizable"}),d(function(){var e="<style>body."+b.css.resizableNoSelect+" { -ms-user-select: none; -moz-user-select: -moz-none;-khtml-user-select: none; -webkit-user-select: none; user-select: none; }."+b.css.resizableContainer+" { position: relative; height: 1px; }."+b.css.resizableHandle+" { position: absolute; display: inline-block; width: 8px;top: 1px; cursor: ew-resize; z-index: 3; user-select: none; -moz-user-select: none; }</style>";d("head").append(e)}),b.resizable={init:function(e,t){if(!e.$table.hasClass("hasResizable")){e.$table.addClass("hasResizable");var s,a,i,r=e.$table,l=r.parent(),o=parseInt(r.css("margin-top"),10),n=t.resizable_vars={useStorage:b.storage&&!1!==t.resizable,$wrap:l,mouseXPosition:0,$target:null,$next:null,overflow:"auto"===l.css("overflow")||"scroll"===l.css("overflow")||"auto"===l.css("overflow-x")||"scroll"===l.css("overflow-x"),storedSizes:[]};for(b.resizableReset(e.table,!0),n.tableWidth=r.width(),n.fullWidth=Math.abs(l.width()-n.tableWidth)<20,n.useStorage&&n.overflow&&(b.storage(e.table,"tablesorter-table-original-css-width",n.tableWidth),i=b.storage(e.table,"tablesorter-table-resized-width")||"auto",b.resizable.setWidth(r,i,!0)),t.resizable_vars.storedSizes=l=(n.useStorage?b.storage(e.table,b.css.resizableStorage):[])||[],b.resizable.setWidths(e,t,l),b.resizable.updateStoredSizes(e,t),t.$resizable_container=d('<div class="'+b.css.resizableContainer+'">').css({top:o}).insertBefore(r),a=0;a<e.columns;a++)s=e.$headerIndexed[a],i=b.getColumnData(e.table,e.headers,a),"false"===b.getData(s,i,"resizable")||d('<div class="'+b.css.resizableHandle+'">').appendTo(t.$resizable_container).attr({"data-column":a,unselectable:"on"}).data("header",s).bind("selectstart",!1);b.resizable.bindings(e,t)}},updateStoredSizes:function(e,t){var s,a,i=e.columns,r=t.resizable_vars;for(r.storedSizes=[],s=0;s<i;s++)a=e.$headerIndexed[s],r.storedSizes[s]=a.is(":visible")?a.width():0},setWidth:function(e,t,s){e.css({width:t,"min-width":s?t:"","max-width":s?t:""})},setWidths:function(e,t,s){var a,i,r=t.resizable_vars,l=d(e.namespace+"_extra_headers"),o=e.$table.children("colgroup").children("col");if((s=s||r.storedSizes||[]).length){for(a=0;a<e.columns;a++)b.resizable.setWidth(e.$headerIndexed[a],s[a],r.overflow),l.length&&(i=l.eq(a).add(o.eq(a)),b.resizable.setWidth(i,s[a],r.overflow));(i=d(e.namespace+"_extra_table")).length&&!b.hasWidget(e.table,"scroller")&&b.resizable.setWidth(i,e.$table.outerWidth(),r.overflow)}},setHandlePosition:function(i,r){var l,o=i.$table.height(),e=r.$resizable_container.children(),n=Math.floor(e.width()/2);b.hasWidget(i.table,"scroller")&&(o=0,i.$table.closest("."+b.css.scrollerWrap).children().each(function(){var e=d(this);o+=(e.filter('[style*="height"]').length?e:e.children("table")).height()})),!r.resizable_includeFooter&&i.$table.children("tfoot").length&&(o-=i.$table.children("tfoot").height()),l=3.3<=parseFloat(d.fn.jquery)?0:i.$table.position().left,e.each(function(){var e=d(this),t=parseInt(e.attr("data-column"),10),s=i.columns-1,a=e.data("header");a&&(!a.is(":visible")||!r.resizable_addLastColumn&&b.resizable.checkVisibleColumns(i,t)?e.hide():(t<s||t===s&&r.resizable_addLastColumn)&&e.css({display:"inline-block",height:o,left:a.position().left-l+a.outerWidth()-n}))})},checkVisibleColumns:function(e,t){for(var s=0,a=t+1;a<e.columns;a++)s+=e.$headerIndexed[a].is(":visible")?1:0;return 0===s},toggleTextSelection:function(e,t,s){e=e.namespace+"tsresize";t.resizable_vars.disabled=s,d("body").toggleClass(b.css.resizableNoSelect,s),s?d("body").attr("unselectable","on").bind("selectstart"+e,!1):d("body").removeAttr("unselectable").unbind("selectstart"+e)},bindings:function(r,l){var e=r.namespace+"tsresize";l.$resizable_container.children().bind("mousedown",function(e){var t=l.resizable_vars,s=d(r.namespace+"_extra_headers"),a=d(e.target).data("header"),i=parseInt(a.attr("data-column"),10);t.$target=a=a.add(s.filter('[data-column="'+i+'"]')),t.target=i,t.$next=e.shiftKey||l.resizable_targetLast?a.parent().children().not(".resizable-false").filter(":last"):a.nextAll(":not(.resizable-false)").eq(0),i=parseInt(t.$next.attr("data-column"),10),t.$next=t.$next.add(s.filter('[data-column="'+i+'"]')),t.next=i,t.mouseXPosition=e.pageX,b.resizable.updateStoredSizes(r,l),b.resizable.toggleTextSelection(r,l,!0)}),d(document).bind("mousemove"+e,function(e){var t=l.resizable_vars;t.disabled&&0!==t.mouseXPosition&&t.$target&&(l.resizable_throttle?(clearTimeout(t.timer),t.timer=setTimeout(function(){b.resizable.mouseMove(r,l,e)},isNaN(l.resizable_throttle)?5:l.resizable_throttle)):b.resizable.mouseMove(r,l,e))}).bind("mouseup"+e,function(){l.resizable_vars.disabled&&(b.resizable.toggleTextSelection(r,l,!1),b.resizable.stopResize(r,l),b.resizable.setHandlePosition(r,l))}),d(t).bind("resize"+e+" resizeEnd"+e,function(){b.resizable.setHandlePosition(r,l)}),r.$table.bind("columnUpdate pagerComplete resizableUpdate ".split(" ").join(e+" "),function(){b.resizable.setHandlePosition(r,l)}).bind("resizableReset"+e,function(){b.resizableReset(r.table)}).find("thead:first").add(d(r.namespace+"_extra_table").find("thead:first")).bind("contextmenu"+e,function(){var e=0===l.resizable_vars.storedSizes.length;return b.resizableReset(r.table),b.resizable.setHandlePosition(r,l),l.resizable_vars.storedSizes=[],e})},mouseMove:function(e,t,s){if(0!==t.resizable_vars.mouseXPosition&&t.resizable_vars.$target){var a,i=0,r=t.resizable_vars,l=r.$next,o=r.storedSizes[r.target],n=s.pageX-r.mouseXPosition;if(r.overflow){if(0<o+n){for(r.storedSizes[r.target]+=n,b.resizable.setWidth(r.$target,r.storedSizes[r.target],!0),a=0;a<e.columns;a++)i+=r.storedSizes[a];b.resizable.setWidth(e.$table.add(d(e.namespace+"_extra_table")),i)}l.length||(r.$wrap[0].scrollLeft=e.$table.width())}else r.fullWidth?(r.storedSizes[r.target]+=n,r.storedSizes[r.next]-=n):r.storedSizes[r.target]+=n,b.resizable.setWidths(e,t);r.mouseXPosition=s.pageX,e.$table.triggerHandler("stickyHeadersUpdate")}},stopResize:function(e,t){var s=t.resizable_vars;b.resizable.updateStoredSizes(e,t),s.useStorage&&(b.storage(e.table,b.css.resizableStorage,s.storedSizes),b.storage(e.table,"tablesorter-table-resized-width",e.$table.width())),s.mouseXPosition=0,s.$target=s.$next=null,e.$table.triggerHandler("stickyHeadersUpdate"),e.$table.triggerHandler("resizableComplete")}},b.addWidget({id:"resizable",priority:40,options:{resizable:!0,resizable_addLastColumn:!1,resizable_includeFooter:!0,resizable_widths:[],resizable_throttle:!1,resizable_targetLast:!1},init:function(e,t,s,a){b.resizable.init(s,a)},format:function(e,t,s){b.resizable.setHandlePosition(t,s)},remove:function(e,t,s,a){var i;s.$resizable_container&&(i=t.namespace+"tsresize",t.$table.add(d(t.namespace+"_extra_table")).removeClass("hasResizable").children("thead").unbind("contextmenu"+i),s.$resizable_container.remove(),b.resizable.toggleTextSelection(t,s,!1),b.resizableReset(e,a),d(document).unbind("mousemove"+i+" mouseup"+i))}}),b.resizableReset=function(r,l){d(r).each(function(){var e,t,s=this.config,a=s&&s.widgetOptions,i=a.resizable_vars;if(r&&s&&s.$headerIndexed.length){for(i.overflow&&i.tableWidth&&(b.resizable.setWidth(s.$table,i.tableWidth,!0),i.useStorage)&&b.storage(r,"tablesorter-table-resized-width",i.tableWidth),e=0;e<s.columns;e++)t=s.$headerIndexed[e],a.resizable_widths&&a.resizable_widths[e]?b.resizable.setWidth(t,a.resizable_widths[e],i.overflow):t.hasClass("resizable-false")||b.resizable.setWidth(t,"",i.overflow);s.$table.triggerHandler("stickyHeadersUpdate"),b.storage&&!l&&b.storage(this,b.css.resizableStorage,[])}})}}(jQuery,window);return jQuery;}));
```
