## Supplementary Data SD1 for "Inferring binding specificities of human transcription factors with the wisdom of crowds": widget-saveSort.min.js

```
(function(factory){if (typeof define === 'function' && define.amd){define(['jquery'], factory);} else if (typeof module === 'object' && typeof module.exports === 'object'){module.exports = factory(require('jquery'));} else {factory(jQuery);}}(function(jQuery){
/*! Widget: saveSort - updated 2018-03-19 (v2.30.1) */
!function(s){"use strict";var l=s.tablesorter||{};function v(t){t=l.storage(t.table,"tablesorter-savesort");return t&&t.hasOwnProperty("sortList")&&s.isArray(t.sortList)?t.sortList:[]}function g(t,s){return(s||v(t)).join(",")!==t.sortList.join(",")}l.addWidget({id:"saveSort",priority:20,options:{saveSort:!0},init:function(t,s,o,r){s.format(t,o,r,!0)},format:function(s,t,o,r){var e,a=t.$table,o=!1!==o.saveSort,i={sortList:t.sortList},n=l.debug(t,"saveSort");n&&(e=new Date),a.hasClass("hasSaveSort")?o&&s.hasInitialized&&l.storage&&g(t)&&(l.storage(s,"tablesorter-savesort",i),n)&&console.log("saveSort >> Saving last sort: "+t.sortList+l.benchmark(e)):(a.addClass("hasSaveSort"),i="",l.storage&&(i=v(t),n&&console.log('saveSort >> Last sort loaded: "'+i+'"'+l.benchmark(e)),a.bind("saveSortReset",function(t){t.stopPropagation(),l.storage(s,"tablesorter-savesort","")})),r&&i&&0<i.length?t.sortList=i:s.hasInitialized&&i&&0<i.length&&g(t,i)&&l.sortOn(t,i))},remove:function(t,s){s.$table.removeClass("hasSaveSort"),l.storage&&l.storage(t,"tablesorter-savesort","")}})}(jQuery);return jQuery;}));
```
