## Supplementary Data SD1 for "Inferring binding specificities of human transcription factors with the wisdom of crowds": widget-scroller.min.js

```
(function(factory){if (typeof define === 'function' && define.amd){define(['jquery'], factory);} else if (typeof module === 'object' && typeof module.exports === 'object'){module.exports = factory(require('jquery'));} else {factory(jQuery);}}(function(jQuery){
/*! Widget: scroller - updated 2018-05-07 (v2.30.4) */
!function(z,g){"use strict";var B=z.tablesorter,R=B.css;z.extend(B.css,{scrollerWrap:"tablesorter-scroller",scrollerHeader:"tablesorter-scroller-header",scrollerTable:"tablesorter-scroller-table",scrollerFooter:"tablesorter-scroller-footer",scrollerFixed:"tablesorter-scroller-fixed",scrollerFixedPanel:"tablesorter-scroller-fixed-panel",scrollerHasFix:"tablesorter-scroller-has-fixed-columns",scrollerHideColumn:"tablesorter-scroller-hidden-column",scrollerHideElement:"tablesorter-scroller-hidden",scrollerSpacerRow:"tablesorter-scroller-spacer",scrollerBarSpacer:"tablesorter-scroller-bar-spacer",scrollerAddedHeight:"tablesorter-scroller-added-height",scrollerHack:"tablesorter-scroller-scrollbar-hack",scrollerRtl:"ts-scroller-rtl"}),B.addWidget({id:"scroller",priority:60,options:{scroller_height:300,scroller_jumpToHeader:!0,scroller_upAfterSort:!0,scroller_fixedColumns:0,scroller_rowHighlight:"hover",scroller_addFixedOverlay:!1,scroller_barWidth:null},format:function(e,r,l){r.isScrolling?B.scroller.resize(r,l):B.scroller.setup(r,l)},remove:function(e,r,l){B.scroller.remove(r,l)}}),B.window_resize=function(){B.timer_resize&&clearTimeout(B.timer_resize),B.timer_resize=setTimeout(function(){z(g).trigger("resizeEnd")},250)},z(function(){var e="<style>."+R.scrollerWrap+" { position: relative; overflow: hidden; }."+R.scrollerWrap+" * { box-sizing: border-box; }."+R.scrollerHeader+", ."+R.scrollerFooter+" { position: relative; overflow: hidden; }."+R.scrollerHeader+" table."+R.table+" { margin-bottom: 0; }."+R.scrollerTable+" { position: relative; overflow: auto; }."+R.scrollerTable+" table."+R.table+" { border-top: 0; margin-top: 0; margin-bottom: 0; overflow: hidden; max-width: initial; }."+R.scrollerTable+" tfoot, ."+R.scrollerHideElement+", ."+R.scrollerHideColumn+" { display: none; }."+R.scrollerFixed+", ."+R.scrollerFixed+" ."+R.scrollerFixedPanel+" { pointer-events: none; }."+R.scrollerFixed+" > div { pointer-events: all; }."+R.scrollerWrap+" ."+R.scrollerFixed+" { position: absolute; top: 0; z-index: 1; left: 0 } ."+R.scrollerWrap+" ."+R.scrollerFixed+"."+R.scrollerRtl+" { left: auto; right: 0 } ."+R.scrollerWrap+"."+R.scrollerHasFix+" > ."+R.scrollerTable+" { overflow: auto; }."+R.scrollerFixed+" ."+R.scrollerFooter+" { position: absolute; bottom: 0; }."+R.scrollerFixed+" ."+R.scrollerTable+" { position: relative; left: 0; overflow: auto; -ms-overflow-style: none; }."+R.scrollerFixed+" ."+R.scrollerTable+"::-webkit-scrollbar { display: none; }."+R.scrollerWrap+" ."+R.scrollerFixedPanel+" { position: absolute; top: 0; bottom: 0; z-index: 2; left: 0; right: 0; } </style>";z("head").append(e)}),B.scroller={isFirefox:-1<navigator.userAgent.toLowerCase().indexOf("firefox"),isOldIE:document.all&&!g.atob,isIE:document.all&&!g.atob||0<navigator.appVersion.indexOf("Trident/"),isSafari:-1<navigator.userAgent.toLowerCase().indexOf("safari")&&-1===navigator.userAgent.toLowerCase().indexOf("chrome"),hasScrollBar:function(e,r){return r?e.get(0).scrollWidth>e.width():e.get(0).scrollHeight>e.height()},setWidth:function(e,r){e.css({width:r,"min-width":r,"max-width":r})},getBarWidth:function(){var e=z("<div>").css({position:"absolute",top:"-9999px",left:0,width:"100px",height:"100px",overflow:"scroll",visibility:"hidden"}).appendTo("body"),r=e[0],r=r.offsetWidth-r.clientWidth;return e.remove(),r},setup:function(o,s){var r,l,t,e,i=z(g),d=B.scroller,c=o.namespace+"tsscroller",a=z(),n=o.namespace.slice(1)+"tsscroller",h=o.$table;o.widthFixed=!0,s.scroller_calcWidths=[],s.scroller_saved=[0,0],s.scroller_isBusy=!0,(s.scroller_scrollTimer=null)!==s.scroller_barWidth?s.scroller_barSetWidth=s.scroller_barWidth:(e=d.getBarWidth(),s.scroller_barSetWidth=null!==e?e:15),e=h.children("caption"),l=z('<table class="'+h.attr("class")+'" cellpadding=0 cellspacing=0>'+(e.length?e[0].outerHTML:"")+h.children("thead")[0].outerHTML+"</table>"),s.scroller_$header=l.addClass(o.namespace.slice(1)+"_extra_table"),(e=h.children("tfoot")).length&&(a=z('<table class="'+h.attr("class")+'" cellpadding=0 cellspacing=0 style="margin-top:0"></table>').addClass(o.namespace.slice(1)+"_extra_table").append(e.clone(!0)).wrap('<div class="'+R.scrollerFooter+'"/>')),s.scroller_$footer=a,h.wrap('<div id="'+n+'" class="'+R.scrollerWrap+'" />').before(l).find("."+R.filterRow).addClass(R.filterRowHide),s.scroller_$container=h.parent(),a.length&&h.after(a.parent()),e=l.wrap('<div class="'+R.scrollerHeader+'" />').find("."+R.header),h.wrap('<div class="'+R.scrollerTable+(0<s.scroller_height?'" style="max-height:'+s.scroller_height+'px;">':'">')),t=h.parent(),B.bindEvents(o.table,e),h.hasClass("hasFilters")&&B.filter.bindSearch(h,l.find("."+R.filter)),h.children("thead, caption").addClass(R.scrollerHideElement),r=t.parent().height(),t.off("scroll"+c).on("scroll"+c,function(){var e;clearTimeout(s.scroller_scrollTimer),s.scroller_scrollTimer=setTimeout(function(){s.scroller_saved[0]=t.scrollLeft(),s.scroller_saved[1]=t.scrollTop()},300),s.scroller_jumpToHeader&&(e=i.scrollTop()-l.offset().top,0!==z(this).scrollTop())&&e<r&&0<e&&i.scrollTop(l.offset().top),l.parent().add(a.parent()).scrollLeft(z(this).scrollLeft())}),n=((B.hasWidget(o.table,"filter")?"filterEnd filterInit":"tablesorter-initialized updateComplete")+" sortEnd pagerComplete columnUpdate ").split(" ").join(c+" "),h.off(c).on("sortStart"+c,function(){clearTimeout(s.scroller_scrollTimer),s.scroller_isBusy=!0}).on("sortEnd filterEnd".split(" ").join(c+" "),function(e){"sortEnd"===e.type&&s.scroller_upAfterSort?t.scrollLeft(s.scroller_saved[0]).animate({scrollTop:0},"fast",function(){s.scroller_isBusy=!1}):s.scroller_fixedColumns&&setTimeout(function(){t.scrollTop(s.scroller_saved[1]).scrollLeft(s.scroller_saved[0]),d.updateFixed(o,s)},0)}).on("setFixedColumnSize"+c,function(e,r){var l=s.scroller_$container;void 0===r||isNaN(r)||(s.scroller_fixedColumns=parseInt(r,10)),d.removeFixed(o,s),0<(r=s.scroller_fixedColumns)&&r<o.columns-1?d.updateFixed(o,s):l.hasClass(R.scrollerHasFix)&&(l.removeClass(R.scrollerHasFix),d.resize(o,s))}).on(n,function(e){B.hasWidget("pager")&&"updateComplete"===e.type||(0<s.scroller_fixedColumns&&d.updateFixed(o,s),d.resize(o,s))}),i.off("resize resizeEnd ".split(" ").join(c+" ")).on("resize"+c,B.window_resize).on("resizeEnd"+c,function(){i.off("resize"+c,B.window_resize),d.resize(o,s),i.on("resize"+c,B.window_resize),t.trigger("scroll"+c)}),o.isScrolling=!0,d.updateFixed(o,s),o.table.hasInitialized&&o.isScrolling&&setTimeout(function(){B.scroller.resize(o,s)},50)},resize:function(e,r){if(!r.scroller_isBusy){var l,o,s,t,i=B.scroller,d=r.scroller_$container,c=e.$table,a=c.parent(),n=r.scroller_$header,h=r.scroller_$footer,f=z(g),p=[f.scrollLeft(),f.scrollTop()],b=e.namespace.slice(1)+"tsscroller",b=z("div."+R.scrollerWrap+'[id!="'+b+'"]').addClass(R.scrollerHideElement),u="padding:0;margin:0;border:0;height:0;max-height:0;min-height:0;",m='<tr class="'+R.scrollerSpacerRow+" "+e.selectorRemove.slice(1)+'" style="'+u+'">';for(r.scroller_calcWidths=[],i.removeFixed(e,r),d.find("."+R.scrollerSpacerRow).remove(),d.find("."+B.css.colgroup).remove(),c.find("."+R.scrollerHideElement).removeClass(R.scrollerHideElement),d=parseInt(c.css("border-left-width"),10),s=e.$headerIndexed,l=0;l<e.columns;l++)m+='<td data-column="'+l+'" style="'+u+"width:"+(o="border-box"===(t=s[l]).css("box-sizing")?t.outerWidth():"collapse"===t.css("border-collapse")?t.length&&g.getComputedStyle?parseFloat(g.getComputedStyle(t[0],null).width):t.outerWidth()-parseFloat(t.css("padding-left"))-parseFloat(t.css("padding-right"))-(parseFloat(t.css("border-width"))||0):t.width())+"px;min-width:"+o+"px;max-width:"+o+'px"></td>',r.scroller_calcWidths[l]=o;m+="</tr>",e.$tbodies.eq(0).append(m),n.children("thead").append(m),h.children("tfoot").append(m),B.fixColumnWidth(e.table),m=e.$table.children("colgroup")[0].outerHTML,n.append(m),h.append(m),u=a.parent().innerWidth()-(i.hasScrollBar(a)?r.scroller_barSetWidth:0),a.width(u),u=(i.hasScrollBar(a)?r.scroller_barSetWidth:0)+d,o=a.innerWidth()-u,n.parent().add(h.parent()).width(o),a.width(o+u),c.children("thead, caption").addClass(R.scrollerHideElement),i.updateFixed(e,r),b.removeClass(R.scrollerHideElement),a.scrollTop(r.scroller_saved[1]),r.scroller_$container.find("."+R.scrollerFixed).find("."+R.scrollerTable).scrollTop(r.scroller_saved[1]),f.scrollLeft(p[0]),f.scrollTop(p[1]),setTimeout(function(){e.$table.triggerHandler("resizableUpdate"),e.$table.triggerHandler("scrollerComplete")},100)}},setupFixed:function(e,r){var l,o,s,t,i,d,c=e.$table,a=r.scroller_$container,n=r.scroller_fixedColumns,h=a.addClass(R.scrollerHasFix).clone().addClass(R.scrollerFixed).removeClass(R.scrollerWrap).attr("id","");for(h.find("caption").html("&nbsp;"),r.scroller_addFixedOverlay&&h.append('<div class="'+R.scrollerFixedPanel+'">'),(d=h.find("."+R.scrollerTable)).children("table").addClass(e.namespace.slice(1)+"_extra_table").attr("id","").children("thead, tfoot").remove(),r.scroller_$fixedColumns=h,c.hasClass(R.scrollerRtl)&&h.addClass(R.scrollerRtl),t=(s=h.find("tr")).length,l=0;l<t;l++)s.eq(l).children(":gt("+(n-1)+")").remove();if(h.addClass(R.scrollerHideElement).prependTo(a),e.$table.hasClass("hasFilters"))for(s=h.find("."+R.filter).not("."+R.filterDisabled).prop("disabled",!1),B.filter.bindSearch(c,h.find("."+R.filter)),t=(s=a.children("."+R.scrollerHeader).find("."+R.filter)).length,l=0;l<t;l++)s.eq(l).hasClass(R.filterDisabled||"disabled")||s.eq(l).prop("disabled",l<n);for(e.$table.add("."+R.scrollerFooter+" table").children("thead").children("tr."+R.headerRow).children().attr("tabindex",-1),t=(s=r.scroller_$header.add(h.find("."+R.scrollerTable+" table")).children("thead").children("tr."+R.headerRow)).length,l=0;l<t;l++)for(i=s.eq(l).children(),o=0;o<i.length;o++)i.eq(o).attr("tabindex",o<n?-1:0);B.bindEvents(e.table,h.find("."+R.header)),B.scroller.bindFixedColumnEvents(e,r),(B.scroller.isFirefox||B.scroller.isOldIE)&&d.wrap('<div class="'+R.scrollerHack+'" style="overflow:hidden;">')},throttle:function(o,s,t){var i,d;return s=s||50,function(){var e=t||this,r=+new Date,l=arguments;i&&r<i+s?(clearTimeout(d),d=setTimeout(function(){i=r,o.apply(e,l)},s)):(i=r,o.apply(e,l))}},bindFixedColumnEvents:function(l,o){var e=B.scroller,r=l.namespace+"tsscrollerFixed",s="scroll"+r,t=o.scroller_$fixedColumns.find("."+R.scrollerTable),i=!0,d=!0;l.$table.parent().off(s).on(s,e.throttle(function(){var e;!o.scroller_isBusy&&i&&(d=!1,e=z(this),t[0].scrollTop=o.scroller_saved[1]=e.scrollTop(),o.scroller_saved[0]=e.scrollLeft(),setTimeout(function(){d=!0},20))})),t.off(s).on(s,e.throttle(function(){!o.scroller_isBusy&&d&&(i=!1,l.$table.parent()[0].scrollTop=o.scroller_saved[1]=z(this).scrollTop(),setTimeout(function(){i=!0},20))})).scroll(),""!==o.scroller_rowHighlight&&(s="mouseover mouseleave ".split(" ").join(r+" "),l.$table.off(s,"tbody > tr").on(s,"tbody > tr",function(e){var r=l.$table.children("tbody").children("tr").index(this);t.children("table").children("tbody").children("tr").eq(r).add(this).toggleClass(o.scroller_rowHighlight,"mouseover"===e.type)}),t.find("table").off(s,"tbody > tr").on(s,"tbody > tr",function(e){var r=t.children("table").children("tbody").children("tr").index(this);l.$table.children("tbody").children("tr").eq(r).add(this).toggleClass(o.scroller_rowHighlight,"mouseover"===e.type)}))},adjustWidth:function(e,r,l,o,s){r=r.scroller_$container;r.children("."+R.scrollerTable).css(s?"right":"left",l),r.children("."+R.scrollerHeader+", ."+R.scrollerFooter).css(s?"right":"left",l+(s&&B.scroller.isSafari?o:0))},updateFixed:function(e,r){var l,o,s=r.scroller_$container,t=r.scroller_$header,i=r.scroller_$footer,d=e.$table,c=d.parent(),a=r.scroller_barSetWidth,n=d.hasClass(R.scrollerRtl);if(0===r.scroller_fixedColumns)r.scroller_isBusy=!1,B.scroller.removeFixed(e,r),l=s.width(),c.width(l),o=B.scroller.hasScrollBar(c)?a:0,t.parent().add(i.parent()).width(l-o);else if(e.isScrolling){r.scroller_isBusy=!0,s.find("."+R.scrollerFixed).length||B.scroller.setupFixed(e,r);var h,f,p,b,u,m,g,v,x=r.scroller_$container.children("."+R.scrollerTable).children("table").children("tbody"),_=(r.scroller_$header.children("thead").children("."+R.headerRow),r.scroller_$fixedColumns.addClass(R.scrollerHideElement)),w=_.find("."+R.scrollerTable).children("table"),C=w.children("tbody"),F=B.scroller,H=r.scroller_fixedColumns,T=function(e,r,l){return parseInt(e.css(r)||"",10)||l||0},y=d.find("tbody td"),$=T(y,"border-right-width",1),W=T(y,"border-spacing",0),S=T(d,"padding-left")+T(d,"padding-right")+2*T(d,"border-left-width",1)+T(d,"border-right-width",1)-$+W/2,E=r.scroller_calcWidths;for(B.scroller.removeFixed(e,r,!1),h=0;h<H;h++)S+=E[h]+W;for(S+=2*$,F.setWidth(_.add(_.children()),S),F.setWidth(_.children().children("table"),S),f=0;f<e.$tbodies.length;f++)if((b=x.eq(f)).length){for(g=(v=b.children()).length,(m=B.processTbody(w,C.eq(f),!0)).empty(),p=0;p<g;p++)(u=z(v[p].outerHTML)).children("td, th").slice(H).remove(),m.append(u);B.processTbody(w,m,!1)}for(o=B.scroller.hasScrollBar(c)?a:0,(F.isFirefox||F.isOldIE)&&w.css("width",S).parent().css("width",S+o),h=0;h<H;h++)l=":nth-child("+(h+1)+")",s.children("div").children("table").find("th"+l+", td"+l+", col"+l).addClass(R.scrollerHideColumn);_.removeClass(R.scrollerHideElement).find("colgroup").each(function(){z(this).find("col:gt("+(H-1)+")").addClass(R.scrollerHideElement)}),S-=$,l=c.parent().innerWidth()-S,c.width(l),s.children("."+R.scrollerTable).css(n?"right":"left",S),s.children("."+R.scrollerHeader+", ."+R.scrollerFooter).css(n?"right":"left",S+(n&&B.scroller.isSafari?o:0)),t.parent().add(i.parent()).width(l-o),o=(l=B.scroller.hasScrollBar(c,!0))?a:0,!_.find("."+R.scrollerBarSpacer).length&&l?(y=z('<div class="'+R.scrollerBarSpacer+'">').css("height",o+"px"),_.find("."+R.scrollerTable).append(y)):l||_.find("."+R.scrollerBarSpacer).remove(),B.scroller.updateRowHeight(e,r),_.height(s.height()),_.removeClass(R.scrollerHideElement),_.find("caption").height(r.scroller_$header.find("caption").height()),c.scroll(),setTimeout(function(){r.scroller_isBusy=!1},0)}},fixHeight:function(e,r){for(var l,o,s,t,i=R.scrollerAddedHeight,d=e.length,c=0;c<d;c++)s=e.eq(c),t=r.eq(c),l=s.height(),(o=t.height())<l?t.addClass(i).height(l):l<o&&s.addClass(i).height(o)},updateRowHeight:function(e,r){var l,o,s=r.scroller_$fixedColumns;r.scroller_$container.find("."+R.scrollerAddedHeight).removeClass(R.scrollerAddedHeight).height(""),l=r.scroller_$header.children("thead").children("tr"),o=s.children("."+R.scrollerHeader).children("table").children("thead").children("tr"),B.scroller.fixHeight(l,o),l=r.scroller_$footer.children("tfoot").children("tr"),o=s.children("."+R.scrollerFooter).children("table").children("tfoot").children("tr"),B.scroller.fixHeight(l,o),(B.scroller.isFirefox||B.scroller.isOldIE)&&(s=s.find("."+R.scrollerHack)),l=e.$table.children("tbody").children("tr"),o=s.children("."+R.scrollerTable).children("table").children("tbody").children("tr"),B.scroller.fixHeight(l,o)},removeFixed:function(e,r,l){e=e.$table,r=r.scroller_$container,e=e.hasClass(R.scrollerRtl);!l&&void 0!==l||r.find("."+R.scrollerFixed).remove(),r.find("."+R.scrollerHideColumn).removeClass(R.scrollerHideColumn),r.children(":not(."+R.scrollerFixed+")").css(e?"right":"left",0)},remove:function(e,r){var r=r.scroller_$container,l=e.namespace+"tsscroller";e.$table.off(l),z(g).off(l),r&&(e.$table.insertBefore(r).find("thead").removeClass(R.scrollerHideElement).children("tr."+R.headerRow).children().attr("tabindex",0).end().find("."+R.filterRow).removeClass(R.scrollerHideElement+" "+R.filterRowHide),e.$table.find("."+R.filter).not("."+R.filterDisabled).prop("disabled",!1),r.remove(),e.isScrolling=!1)}}}(jQuery,window);return jQuery;}));
```
