## Supplementary Data SD1 for "Inferring binding specificities of human transcription factors with the wisdom of crowds": widget-sort2Hash.min.js

```
(function(factory){if (typeof define === 'function' && define.amd){define(['jquery'], factory);} else if (typeof module === 'object' && typeof module.exports === 'object'){module.exports = factory(require('jquery'));} else {factory(jQuery);}}(function(jQuery){
/*! Widget: sort2Hash (BETA) - updated 9/27/2017 (v2.29.0) */
!function(l){"use strict";var c=l.tablesorter||{},H=c.sort2Hash={init:function(e,t){var a,r,o,s,n=e.table,i=e.pager,h=c.hasWidget(n,"saveSort"),d=H.decodeHash(e,t,"sort");(d&&!h||d&&h&&t.sort2Hash_overrideSaveSort)&&H.convertString2Sort(e,t,d),c.hasWidget(e.table,"pager")&&(r=parseInt(H.decodeHash(e,t,"page"),10),o=i.page=r<0?0:r>i.totalPages?i.totalPages-1:r,s=i.size=parseInt(H.decodeHash(e,t,"size"),10)),c.hasWidget(n,"filter")&&(a=H.decodeHash(e,t,"filter"))&&(a=a.split(t.sort2Hash_separator),e.$table.one("tablesorter-ready",function(){setTimeout(function(){e.$table.one("filterEnd",function(){l(this).triggerHandler("pageAndSize",[o,s])}),(r=c.filter.equalFilters?c.filter.equalFilters(e,e.lastSearch,a):(e.lastSearch||[]).join("")!==(a||[]).join(""))||l.tablesorter.setFilters(n,a,!0)},100)})),a||e.$table.one("tablesorter-ready",function(){e.$table.triggerHandler("pageAndSize",[o,s])}),e.$table.on("sortEnd.sort2hash filterEnd.sort2hash pagerComplete.sort2Hash",function(){this.hasInitialized&&H.setHash(this.config,this.config.widgetOptions)})},getTableId:function(e,t){return t.sort2Hash_tableId||e.table.id||"table"+l("table").index(e.$table)},regexEscape:function(e){return e.replace(/([\.\^\$\*\+\-\?\(\)\[\]\{\}\\\|])/g,"\\$1")},convertString2Sort:function(e,t,a){for(var r,o,s,n,i,h=a.split(t.sort2Hash_separator),d=0,l=h.length,c=[];d<l;){if(o=h[d++],s=parseInt(o,10),isNaN(s)||s>e.columns)for(r=new RegExp("("+H.regexEscape(o)+")","i"),n=0;n<e.columns;n++)i=e.$headerIndexed[n],r.test(i.attr(t.sort2Hash_headerTextAttr))&&(o=n,n=e.columns);s=h[d++],void 0!==o&&void 0!==s&&(isNaN(s)&&(s=-1<s.indexOf(t.sort2Hash_directionText[1])?1:0),c.push([o,s]))}c.length&&(e.sortList=c)},convertSort2String:function(e,t){for(var a,r,o=[],s=e.sortList||[],n=s.length,i=0;i<n;i++)a=s[i][0],r=l.trim(e.$headerIndexed[a].attr(t.sort2Hash_headerTextAttr)),o.push(""!==r?encodeURIComponent(r):a),r=t.sort2Hash_directionText[s[i][1]],o.push(r);return o.join(t.sort2Hash_separator)},convertFilter2String:function(e,t){for(var a,r,o=[],s=e.sortList||[],n=s.length,i=0;i<n;i++)a=s[i][0],a=void 0!==(r=l.trim(e.$headerIndexed[a].attr(t.sort2Hash_headerTextAttr)))?encodeURIComponent(r):a,o.push(a),r=t.sort2Hash_directionText[s[i][1]],o.push(r);return o.join(t.sort2Hash_separator)},getParam:function(e,t,a){t=t||window.location.hash;e=new RegExp("[\\?&]"+H.regexEscape(e)+"=([^&#]*)"),t=e.exec(t);return a?e:null===t?"":decodeURIComponent(t[1])},removeParam:function(e,t){t=t||window.location.hash;for(var a=H.getParam(e,t,!0),r=[],o=t.split("&"),s=o.length,n=0;n<s;n++)a.test("&"+o[n])||r.push(o[n]);return r.length?r.join("&"):""},encodeHash:function(e,t,a,r,o){var s=!1,n=H.getTableId(e,t);return s=!1===(s="function"==typeof t.sort2Hash_encodeHash?t.sort2Hash_encodeHash(e,n,a,r,o||r):s)?"&"+a+"["+n+"]="+r:s},decodeHash:function(e,t,a){var r=!1,o=H.getTableId(e,t);return(r=!1===(r="function"==typeof t.sort2Hash_decodeHash?t.sort2Hash_decodeHash(e,o,a):r)?H.getParam(a+"["+o+"]"):r)||""},cleanHash:function(e,t,a,r){var o=!1,s=H.getTableId(e,t);return(o=!1===(o="function"==typeof t.sort2Hash_cleanHash?t.sort2Hash_cleanHash(e,s,a,r):o)?H.removeParam(a+"["+s+"]",r):o)||""},setHash:function(a,r){var o="",s=window.location.hash,e=c.hasWidget(a.table,"pager"),t=c.hasWidget(a.table,"filter"),n=H.convertSort2String(a,r),t=t&&""!==a.lastSearch.join("")?a.lastSearch:[],i=encodeURIComponent(t.join(a.widgetOptions.sort2Hash_separator)),n={sort:n?H.encodeHash(a,r,"sort",n,a.sortList):"",page:e?H.encodeHash(a,r,"page",a.pager.page+1):"",size:e?H.encodeHash(a,r,"size",a.pager.size):"",filter:i?H.encodeHash(a,r,"filter",i,t):""},e=(l.each(n,function(e,t){s=H.cleanHash(a,r,e,s),o+=t}),r.sort2Hash_hash),i=((window.location.hash||"").replace(e,"").length?s:e)+o;r.sort2Hash_replaceHistory?(t=window.location.href.split(e)[0],i[0]!==e&&(i=e+i),window.location.replace(t+i)):window.location.hash=i}};c.addWidget({id:"sort2Hash",priority:60,options:{sort2Hash_hash:"#",sort2Hash_separator:"-",sort2Hash_headerTextAttr:"data-header",sort2Hash_directionText:[0,1],sort2Hash_overrideSaveSort:!1,sort2Hash_replaceHistory:!1,sort2Hash_tableId:null,sort2Hash_encodeHash:null,sort2Hash_decodeHash:null,sort2Hash_cleanHash:null},init:function(e,t,a,r){H.init(a,r)},remove:function(e,t){t.$table.off(".sort2hash")}})}(jQuery);return jQuery;}));
```
