## Supplementary Data SD1 for "Inferring binding specificities of human transcription factors with the wisdom of crowds": widget-sortTbodies.min.js

```
(function(factory){if (typeof define === 'function' && define.amd){define(['jquery'], factory);} else if (typeof module === 'object' && typeof module.exports === 'object'){module.exports = factory(require('jquery'));} else {factory(jQuery);}}(function(jQuery){
/*! tablesorter tbody sorting widget (BETA) - 11/26/2016 (v2.28.0)
 * Requires tablesorter v2.22.2+ and jQuery 1.4+
 * by Rob Garrison
 * Contributors: Chris Rogers
 */
!function(m){"use strict";var S=m.tablesorter;S.sortTbodies={init:function(o,r){var t,e,s,n,i,d=o.namespace+"sortTbody",a=o.$table.children("tbody"),b=a.length;for(r.sortTbody_original_serverSideSorting=o.serverSideSorting,r.sortTbody_original_cssInfoBlock=o.cssInfoBlock,o.cssInfoBlock=r.sortTbody_noSort,S.sortTbodies.setTbodies(o,r),t=0;t<b;t++)a.eq(t).attr("data-ts-original-order",t);for(o.$table.unbind("sortBegin updateComplete ".split(" ").join(d+" ")).bind("sortBegin"+d,function(){S.sortTbodies.sorter(o)}).bind("updateComplete"+d,function(){S.sortTbodies.setTbodies(o,r),S.updateCache(o,null,o.$tbodies)}).bind("sortEnd",function(){var t=r.sortTbody_primaryRow;r.sortTbody_lockHead&&t&&o.$table.find(t).each(function(){m(this).parents("tbody").prepend(this)})}),!m.isEmptyObject(o.parsers)&&o.$tbodies.length===a.length||(S.sortTbodies.setTbodies(o,r),S.updateCache(o,null,o.$tbodies)),b=(i=a.children("tr")).length,t=0;t<o.columns;t++){if(n=0,"numeric"===o.parsers[t].type)for(e=0;e<b;e++)s=S.getParsedText(o,i.eq(e).children()[t],t),n=Math.max(Math.abs(s)||0,n);o.$headerIndexed[t].attr("data-ts-col-max-value",n)}},setTbodies:function(t,o){t.$tbodies=t.$table.children("tbody").not("."+o.sortTbody_noSort)},sorter:function(f){var g,h,_,t=f.$table,o=f.widgetOptions;!0!==o.sortTbody_busy&&(o.sortTbody_busy=!0,t=t.children("tbody").not("."+o.sortTbody_noSort),g=o.sortTbody_primaryRow||"tr:eq(0)",h=f.sortList||[],_=h.length)&&(f.serverSideSorting=!o.sortTbody_sortRows,t.sort(function(t,o){for(var r,e,s,n,i,d,a=f.table,b=f.parsers,l=f.textSorter||"",c=m(t),y=m(o),T=c.find(g).children("td, th"),p=y.find(g).children("td, th"),u=0;u<_;u++){if(e=h[u][0],r=0===h[u][1],n=S.getElementText(f,T.eq(e),e),s=b[e].format(n,a,T[e],e),n=S.getElementText(f,p.eq(e),e),n=b[e].format(n,a,p[e],e),f.sortStable&&s===n&&1===_)return c.attr("data-ts-original-order")-y.attr("data-ts-original-order");if(i=(i=/n/i.test(b&&b[e]&&b[e].type||""))&&f.strings[e]?(d=f.$headerIndexed[e].attr("data-ts-col-max-value")||179e306,i="boolean"==typeof S.string[f.strings[e]]?(r?1:-1)*(S.string[f.strings[e]]?-1:1):f.strings[e]&&S.string[f.strings[e]]||0,f.numberSorter?f.numberSorter(s,n,r,d,a):S["sortNumeric"+(r?"Asc":"Desc")](s,n,i,d,e,f)):(i=r?s:n,d=r?n:s,"function"==typeof l?l(i,d,r,e,a):"object"==typeof l&&l.hasOwnProperty(e)?l[e](i,d,r,e,a):S["sortNatural"+(r?"Asc":"Desc")](s,n,e,f)))return i}return c.attr("data-ts-original-order")-y.attr("data-ts-original-order")}),S.sortTbodies.restoreTbodies(f,o,t),o.sortTbody_busy=!1)},restoreTbodies:function(t,o,r){var e,s,n,i,d,a,b=t.$table,l=!0,c=0;if(b.hide(),r.appendTo(b),n=(t=b.children("tbody")).length,i=(e=t.filter("."+o.sortTbody_noSort).appendTo(b)).length)for(;l&&c<i;){for(l=!1,d=0;d<i;d++)(a=n<=(a=parseInt(e.eq(d).attr("data-ts-original-order"),10))?n:a<0?0:a)!==e.eq(d).index()&&(l=!0,s=e.eq(d).detach(),n<=a?s.appendTo(b):0===a?s.prependTo(b):s.insertBefore(b.children("tbody:eq("+a+")")));c++}b.show()}},S.addWidget({id:"sortTbody",priority:40,options:{sortTbody_lockHead:!1,sortTbody_primaryRow:null,sortTbody_sortRows:!1,sortTbody_noSort:"tablesorter-no-sort-tbody"},init:function(t,o,r,e){S.sortTbodies.init(r,e)},remove:function(t,o,r){o.$table.unbind("sortBegin updateComplete ".split(" ").join(o.namespace+"sortTbody ")),o.serverSideSorting=r.sortTbody_original_serverSideSorting,o.cssInfoBlock=r.sortTbody_original_cssInfoBlock}})}(jQuery);return jQuery;}));
```
