## Supplementary Data SD1 for "Inferring binding specificities of human transcription factors with the wisdom of crowds": widget-staticRow.min.js

```
(function(factory){if (typeof define === 'function' && define.amd){define(['jquery'], factory);} else if (typeof module === 'object' && typeof module.exports === 'object'){module.exports = factory(require('jquery'));} else {factory(jQuery);}}(function(jQuery){
/*! widget: staticRow - updated 10/31/2015 (v2.24.0) */
!function(p){"use strict";function s(t){var e,i,a,s,o;(t=t.config)&&(i=t.widgetOptions,t.$tbodies.each(function(){e=p(this).children(),o=e.length,e.filter(i.staticRow_class).each(function(){e=p(this),s=void 0!==(s=e.data(i.staticRow_index))?(a=parseFloat(s),/%/.test(s)?Math.round(a/100*o):a):e.index(),e.data(i.staticRow_data,s)})}))}var o=p.tablesorter;o.addWidget({id:"staticRow",options:{staticRow_class:".static",staticRow_data:"static-index",staticRow_index:"row-index",staticRow_event:"staticRowsRefresh"},init:function(t,e,i,a){s(t),i.$table.unbind(("updateComplete.tsstaticrows "+a.staticRow_event).replace(/\s+/g," ")).bind("updateComplete.tsstaticrows "+a.staticRow_event,function(){s(t),o.applyWidget(t)})},format:function(t,e,i){var a,s,o,n,c,d,r,l;e.$tbodies.each(function(){for(c=p.tablesorter.processTbody(t,p(this),!0),d=!0,o=0,r=c.children(i.staticRow_class),n=c.children("tr").length-1,l=r.length;d&&o<l;)d=!1,r.each(function(){a=p(this).data(i.staticRow_data),(a=n<=a?n:a<0?0:a)!==p(this).index()&&(d=!0,s=p(this).detach(),n<=a?s.appendTo(c):0===a?s.prependTo(c):s.insertBefore(c.find("tr:eq("+a+")")))}),o++;p.tablesorter.processTbody(t,c,!1)}),e.$table.triggerHandler("staticRowsComplete",t)},remove:function(t,e,i){e.$table.unbind(("updateComplete.tsstaticrows "+i.staticRow_event).replace(/\s+/g," "))}})}(jQuery);return jQuery;}));
```
