## Supplementary Data SD1 for "Inferring binding specificities of human transcription factors with the wisdom of crowds": widget-stickyHeaders.min.js

```
(function(factory){if (typeof define === 'function' && define.amd){define(['jquery'], factory);} else if (typeof module === 'object' && typeof module.exports === 'object'){module.exports = factory(require('jquery'));} else {factory(jQuery);}}(function(jQuery){
/*! Widget: stickyHeaders - updated 9/27/2017 (v2.29.0) */
!function(C,T){"use strict";var w=C.tablesorter||{};function S(e,s){var t=isNaN(s.stickyHeaders_offset)?C(s.stickyHeaders_offset):[];return t.length?t.height()||0:parseInt(s.stickyHeaders_offset,10)||0}C.extend(w.css,{sticky:"tablesorter-stickyHeader",stickyVis:"tablesorter-sticky-visible",stickyHide:"tablesorter-sticky-hidden",stickyWrap:"tablesorter-sticky-wrapper"}),w.addHeaderResizeEvent=function(e,s,t){if((e=C(e)[0]).config){var t=C.extend({},{timer:250},t),c=e.config,l=c.widgetOptions,i=function(e){var s,t,i,r,a,d,n=c.$headers.length;for(l.resize_flag=!0,t=[],s=0;s<n;s++)r=(i=c.$headers.eq(s)).data("savedSizes")||[0,0],a=i[0].offsetWidth,d=i[0].offsetHeight,a===r[0]&&d===r[1]||(i.data("savedSizes",[a,d]),t.push(i[0]));t.length&&!1!==e&&c.$table.triggerHandler("resize",[t]),l.resize_flag=!1};if(clearInterval(l.resize_timer),s)return l.resize_flag=!1;i(!1),l.resize_timer=setInterval(function(){l.resize_flag||i()},t.timer)}},w.addWidget({id:"stickyHeaders",priority:54,options:{stickyHeaders:"",stickyHeaders_appendTo:null,stickyHeaders_attachTo:null,stickyHeaders_xScroll:null,stickyHeaders_yScroll:null,stickyHeaders_offset:0,stickyHeaders_filteredToTop:!0,stickyHeaders_cloneId:"-sticky",stickyHeaders_addResizeEvent:!0,stickyHeaders_includeCaption:!0,stickyHeaders_zIndex:2},format:function(e,s,c){if(!(s.$table.hasClass("hasStickyHeaders")||0<=C.inArray("filter",s.widgets)&&!s.$table.hasClass("hasFilters"))){var t,i,r,a,l=s.$table,o=C(c.stickyHeaders_attachTo||c.stickyHeaders_appendTo),d=s.namespace+"stickyheaders ",p=C(c.stickyHeaders_yScroll||c.stickyHeaders_attachTo||T),n=C(c.stickyHeaders_xScroll||c.stickyHeaders_attachTo||T),h=l.children("thead:first").children("tr").not(".sticky-false").children(),f=l.children("tfoot"),y=S(0,c),k=l.parent().closest("."+w.css.table).hasClass("hasStickyHeaders")?l.parent().closest("table.tablesorter")[0].config.widgetOptions.$sticky.parent():[],g=k.length?k.height():0,H=c.$sticky=l.clone().addClass("containsStickyHeaders "+w.css.sticky+" "+c.stickyHeaders+" "+s.namespace.slice(1)+"_extra_table").wrap('<div class="'+w.css.stickyWrap+'">'),u=H.parent().addClass(w.css.stickyHide).css({position:o.length?"absolute":"fixed",padding:parseInt(H.parent().parent().css("padding-left"),10),top:y+g,left:0,visibility:"hidden",zIndex:c.stickyHeaders_zIndex||2}),y=H.children("thead:first"),b="",_=function(e,s){for(var t,i,r,a=e.filter(":visible"),d=a.length,n=0;n<d;n++)i=s.filter(":visible").eq(n),t="border-box"===(r=a.eq(n)).css("box-sizing")?r.outerWidth():"collapse"===i.css("border-collapse")?T.getComputedStyle?parseFloat(T.getComputedStyle(r[0],null).width):(t=parseFloat(r.css("border-width")),r.outerWidth()-parseFloat(r.css("padding-left"))-parseFloat(r.css("padding-right"))-t):r.width(),i.css({width:t,"min-width":t,"max-width":t})},v=function(e){return!1===e&&k.length?l.position().left:o.length?parseInt(o.css("padding-left"),10)||0:l.offset().left-parseInt(l.css("margin-left"),10)-C(T).scrollLeft()},m=function(){u.css({left:v(),width:l.outerWidth()}),_(l,H),_(h,a)},z=function(e){var s,t,i,r,a,d,n;l.is(":visible")&&(g=k.length?k.offset().top-p.scrollTop()+k.height():0,r=l.offset(),s=S(0,c),i=(t=C.isWindow(p[0]))?p.scrollTop():k.length?parseInt(k[0].style.top,10):p.offset().top,i=o.length?i:p.scrollTop(),a=!c.stickyHeaders_includeCaption&&l.children("caption").height()||0,i=i+s+g-a,a=l.height()-(u.height()+(f.height()||0))-a,a="visible"==(r=i>r.top&&i<r.top+a?"visible":"hidden")?w.css.stickyVis:w.css.stickyHide,d=!u.hasClass(a),n={visibility:r},o.length&&(d=!0,n.top=t?i-o.offset().top:o.scrollTop()),(i=v(t))!==parseInt(u.css("left"),10)&&(d=!0,n.left=i),n.top=(n.top||0)+(!t&&k.length?k.height():s+g),d&&u.removeClass(w.css.stickyVis+" "+w.css.stickyHide).addClass(a).css(n),r===b&&!e||(m(),b=r))};if(o.length&&!o.css("position")&&o.css("position","relative"),H.attr("id")&&(H[0].id+=c.stickyHeaders_cloneId),H.find("> thead:gt(0), tr.sticky-false").hide(),H.find("> tbody, > tfoot").remove(),H.find("caption").toggle(c.stickyHeaders_includeCaption),a=y.children().children(),H.css({height:0,width:0,margin:0}),a.find("."+w.css.resizer).remove(),l.addClass("hasStickyHeaders").bind("pagerComplete"+d,function(){m()}),w.bindEvents(e,y.children().children("."+w.css.header)),c.stickyHeaders_appendTo?C(c.stickyHeaders_appendTo).append(u):l.after(u),s.onRenderHeader)for(i=(r=y.children("tr").children()).length,t=0;t<i;t++)s.onRenderHeader.apply(r.eq(t),[t,s,H]);n.add(p).unbind("scroll resize ".split(" ").join(d).replace(/\s+/g," ")).bind("scroll resize ".split(" ").join(d),function(e){z("resize"===e.type)}),s.$table.unbind("stickyHeadersUpdate"+d).bind("stickyHeadersUpdate"+d,function(){z(!0)}),c.stickyHeaders_addResizeEvent&&w.addHeaderResizeEvent(e),l.hasClass("hasFilters")&&c.filter_columnFilters&&(l.bind("filterEnd"+d,function(){var e=C(document.activeElement).closest("td"),e=e.parent().children().index(e);u.hasClass(w.css.stickyVis)&&c.stickyHeaders_filteredToTop&&(T.scrollTo(0,l.position().top),0<=e)&&s.$filters&&s.$filters.eq(e).find("a, select, input").filter(":visible").focus()}),w.filter.bindSearch(l,a.find("."+w.css.filter)),c.filter_hideFilters)&&w.filter.hideFilters(s,H),c.stickyHeaders_addResizeEvent&&l.bind("resize"+s.namespace+"stickyheaders",function(){m()}),z(!0),l.triggerHandler("stickyHeadersInit")}},remove:function(e,s,t){var i=s.namespace+"stickyheaders ";s.$table.removeClass("hasStickyHeaders").unbind("pagerComplete resize filterEnd stickyHeadersUpdate ".split(" ").join(i).replace(/\s+/g," ")).next("."+w.css.stickyWrap).remove(),t.$sticky&&t.$sticky.length&&t.$sticky.remove(),C(T).add(t.stickyHeaders_xScroll).add(t.stickyHeaders_yScroll).add(t.stickyHeaders_attachTo).unbind("scroll resize ".split(" ").join(i).replace(/\s+/g," ")),w.addHeaderResizeEvent(e,!0)}})}(jQuery,window);return jQuery;}));
```
