## Supplementary Data SD1 for "Inferring binding specificities of human transcription factors with the wisdom of crowds": widget-storage.min.js

```
(function(factory){if (typeof define === 'function' && define.amd){define(['jquery'], factory);} else if (typeof module === 'object' && typeof module.exports === 'object'){module.exports = factory(require('jquery'));} else {factory(jQuery);}}(function(jQuery){
/*! Widget: storage - updated 2018-03-18 (v2.30.0) */
!function(u,S,c){"use strict";var _=u.tablesorter||{};u.extend(!0,_.defaults,{fixedUrl:"",widgetOptions:{storage_fixedUrl:"",storage_group:"",storage_page:"",storage_storageType:"",storage_tableId:"",storage_useSessionStorage:""}}),_.storage=function(e,t,r,o){var s=!1,a={},i=(e=u(e)[0]).config,g=i&&i.widgetOptions,n=_.debug(i,"storage"),p=(o&&o.storageType||g&&g.storage_storageType).toString().charAt(0).toLowerCase(),l=p?"":o&&o.useSessionStorage||g&&g.storage_useSessionStorage,d=u(e),e=o&&o.id||d.attr(o&&o.group||g&&g.storage_group||"data-table-group")||g&&g.storage_tableId||e.id||u(".tablesorter").index(d),d=o&&o.url||d.attr(o&&o.page||g&&g.storage_page||"data-table-page")||g&&g.storage_fixedUrl||i&&i.fixedUrl||S.location.pathname;if("c"!==p&&(p="s"===p||l?"sessionStorage":"localStorage")in S)try{S[p].setItem("_tmptest","temp"),s=!0,S[p].removeItem("_tmptest")}catch(e){console.warn(p+" is not supported in this browser")}if(n&&console.log("Storage >> Using",s?p:"cookies"),u.parseJSON&&(a=s?u.parseJSON(S[p][t]||"null")||{}:(o=c.cookie.split(/[;\s|=]/),0!==(g=u.inArray(t,o)+1)&&u.parseJSON(o[g]||"null")||{})),void 0===r||!S.JSON||!JSON.hasOwnProperty("stringify"))return a&&a[d]?a[d][e]:"";a[d]||(a[d]={}),a[d][e]=r,s?S[p][t]=JSON.stringify(a):((i=new Date).setTime(i.getTime()+31536e6),c.cookie=t+"="+JSON.stringify(a).replace(/\"/g,'"')+"; expires="+i.toGMTString()+"; path=/")}}(jQuery,window,document);return jQuery;}));
```
