## Supplementary Data SD1 for "Inferring binding specificities of human transcription factors with the wisdom of crowds": widget-toggle.min.js

```
(function(factory){if (typeof define === 'function' && define.amd){define(['jquery'], factory);} else if (typeof module === 'object' && typeof module.exports === 'object'){module.exports = factory(require('jquery'));} else {factory(jQuery);}}(function(jQuery){
/*! tablesorter enable/disable sort & filter (BETA) - 11/10/2015 (v2.24.4)
 * Requires tablesorter v2.24.4+ & jQuery 1.7+
 * by Rob Garrison
 */
!function(){"use strict";var g=jQuery.tablesorter,s=g.toggleTS={init:function(e,l){l.toggleTS_isEnabled=!0,l.toggleTS_areDisabled={headers:[],filters:[]},e.$table.on("enable.toggleTS disable.toggleTS",function(e){s.toggle(this.config,this.config.widgetOptions,"enable"===e.type)})},toggle:function(e,l,t){if(l.toggleTS_isEnabled!==t){l.toggleTS_isEnabled=t;for(var i,s=e.$headers.length,o=0;o<s;o++)i=e.$headers.eq(o),g.setColumnSort(e,i,!t),g.setColumnAriaLabel(e,i,t);if(l.toggleTS_hideFilterRow)e.$table.find("."+g.css.filterRow).toggle(t);else if(g.hasWidget(e.$table,"filter"))for(s=e.$filters.length,o=0;o<s;o++)t&&!l.toggleTS_areDisabled.filters[o]?e.$filters.eq(o).find("input, select").removeClass(g.css.filterDisabled).prop("disabled",!1):t||((i=e.$filters.eq(o).find("input, select")).hasClass(g.css.filterDisabled)&&(l.toggleTS_areDisabled.filters[o]=!0),i.addClass(g.css.filterDisabled).prop("disabled",!0));l.filter_$externalFilters.toggleClass(g.css.filterDisabled,t).prop("disabled",!t)}"function"==typeof l.toggleTS_callback&&l.toggleTS_callback(e,t)}};g.addWidget({id:"toggle-ts",options:{toggleTS_hideFilterRow:!1,toggleTS_callback:null},init:function(e,l,t,i){s.init(t,i)},remove:function(e,l){l.$table.off("enable.toggleTS disable.toggleTS")}})}();return jQuery;}));
```
