## Supplementary Data SD1 for "Inferring binding specificities of human transcription factors with the wisdom of crowds": widget-uitheme.min.js

```
(function(factory){if (typeof define === 'function' && define.amd){define(['jquery'], factory);} else if (typeof module === 'object' && typeof module.exports === 'object'){module.exports = factory(require('jquery'));} else {factory(jQuery);}}(function(jQuery){
/*! Widget: uitheme - updated 2018-03-18 (v2.30.0) */
!function(D){"use strict";var A=D.tablesorter||{};A.themes={bootstrap:{table:"table table-bordered table-striped",caption:"caption",header:"bootstrap-header",sortNone:"",sortAsc:"",sortDesc:"",active:"",hover:"",icons:"",iconSortNone:"bootstrap-icon-unsorted",iconSortAsc:"glyphicon glyphicon-chevron-up",iconSortDesc:"glyphicon glyphicon-chevron-down",filterRow:"",footerRow:"",footerCells:"",even:"",odd:""},jui:{table:"ui-widget ui-widget-content ui-corner-all",caption:"ui-widget-content",header:"ui-widget-header ui-corner-all ui-state-default",sortNone:"",sortAsc:"",sortDesc:"",active:"ui-state-active",hover:"ui-state-hover",icons:"ui-icon",iconSortNone:"ui-icon-carat-2-n-s ui-icon-caret-2-n-s",iconSortAsc:"ui-icon-carat-1-n ui-icon-caret-1-n",iconSortDesc:"ui-icon-carat-1-s ui-icon-caret-1-s",filterRow:"",footerRow:"",footerCells:"",even:"ui-widget-content",odd:"ui-state-default"}},D.extend(A.css,{wrapper:"tablesorter-wrapper"}),A.addWidget({id:"uitheme",priority:10,format:function(e,o,t){var s,r,i,a,n,c,l,d,h,m,u,p,v=A.themes,f=o.$table.add(D(o.namespace+"_extra_table")),b=o.$headers.add(D(o.namespace+"_extra_headers")),C=o.theme||"jui",w=v[C]||{},S=D.trim([w.sortNone,w.sortDesc,w.sortAsc,w.active].join(" ")),g=D.trim([w.iconSortNone,w.iconSortDesc,w.iconSortAsc].join(" ")),j=A.debug(o,"uitheme");for(j&&(n=new Date),f.hasClass("tablesorter-"+C)&&o.theme===o.appliedTheme&&t.uitheme_applied||(t.uitheme_applied=!0,m=v[o.appliedTheme]||{},v=(p=!D.isEmptyObject(m))?[m.sortNone,m.sortDesc,m.sortAsc,m.active].join(" "):"",u=p?[m.iconSortNone,m.iconSortDesc,m.iconSortAsc].join(" "):"",p&&(t.zebra[0]=D.trim(" "+t.zebra[0].replace(" "+m.even,"")),t.zebra[1]=D.trim(" "+t.zebra[1].replace(" "+m.odd,"")),o.$tbodies.children().removeClass([m.even,m.odd].join(" "))),w.even&&(t.zebra[0]+=" "+w.even),w.odd&&(t.zebra[1]+=" "+w.odd),f.children("caption").removeClass(m.caption||"").addClass(w.caption),d=f.removeClass((o.appliedTheme?"tablesorter-"+(o.appliedTheme||""):"")+" "+(m.table||"")).addClass("tablesorter-"+C+" "+(w.table||"")).children("tfoot"),o.appliedTheme=o.theme,d.length&&d.children("tr").removeClass(m.footerRow||"").addClass(w.footerRow).children("th, td").removeClass(m.footerCells||"").addClass(w.footerCells),b.removeClass((p?[m.header,m.hover,v].join(" "):"")||"").addClass(w.header).not(".sorter-false").unbind("mouseenter.tsuitheme mouseleave.tsuitheme").bind("mouseenter.tsuitheme mouseleave.tsuitheme",function(e){D(this)["mouseenter"===e.type?"addClass":"removeClass"](w.hover||"")}),b.each(function(){var e=D(this);e.find("."+A.css.wrapper).length||e.wrapInner('<div class="'+A.css.wrapper+'" style="position:relative;height:100%;width:100%"></div>')}),o.cssIcon&&b.find("."+A.css.icon).removeClass(p?[m.icons,u].join(" "):"").addClass(w.icons||""),A.hasWidget(o.table,"filter")&&(r=function(){f.children("thead").children("."+A.css.filterRow).removeClass(p&&m.filterRow||"").addClass(w.filterRow||"")},t.filter_initialized?r():f.one("filterInit",function(){r()}))),s=0;s<o.columns;s++)c=o.$headers.add(D(o.namespace+"_extra_headers")).not(".sorter-false").filter('[data-column="'+s+'"]'),l=A.css.icon?c.find("."+A.css.icon):D(),(h=b.not(".sorter-false").filter('[data-column="'+s+'"]:last')).length&&(c.removeClass(S),l.removeClass(g),h[0].sortDisabled?l.removeClass(w.icons||""):(i=w.sortNone,a=w.iconSortNone,h.hasClass(A.css.sortAsc)?(i=[w.sortAsc,w.active].join(" "),a=w.iconSortAsc):h.hasClass(A.css.sortDesc)&&(i=[w.sortDesc,w.active].join(" "),a=w.iconSortDesc),c.addClass(i),l.addClass(a||"")));j&&console.log("uitheme >> Applied "+C+" theme"+A.benchmark(n))},remove:function(e,o,t,s){var r,i,a,n,c;t.uitheme_applied&&(r=o.$table,o=o.appliedTheme||"jui",i=A.themes[o]||A.themes.jui,a=r.children("thead").children(),n=i.sortNone+" "+i.sortDesc+" "+i.sortAsc,c=i.iconSortNone+" "+i.iconSortDesc+" "+i.iconSortAsc,r.removeClass("tablesorter-"+o+" "+i.table),t.uitheme_applied=!1,s||(r.find(A.css.header).removeClass(i.header),a.unbind("mouseenter.tsuitheme mouseleave.tsuitheme").removeClass(i.hover+" "+n+" "+i.active).filter("."+A.css.filterRow).removeClass(i.filterRow),a.find("."+A.css.icon).removeClass(i.icons+" "+c)))}})}(jQuery);return jQuery;}));
```
