## Supplementary Data SD1 for "Inferring binding specificities of human transcription factors with the wisdom of crowds": widget-vertical-group.min.js

```
(function(factory){if (typeof define === 'function' && define.amd){define(['jquery'], factory);} else if (typeof module === 'object' && typeof module.exports === 'object'){module.exports = factory(require('jquery'));} else {factory(jQuery);}}(function(jQuery){
/*! Widget: vertical-group (BETA) - updated 12/13/2017 (v2.29.1) */
!function(h){"use strict";var G=h.tablesorter,f=G.css;function g(r){r.removeClass(f.verticalGroupHide+" "+f.verticalGroupShow)}function C(r,e,a){e.parent().removeClass(r.zebra[(a+1)%2]).addClass(r.zebra[a%2])}function r(r,e,a){var o=-1,t=r.tBodies[0].rows,i=G.hasWidget(r,"zebra"),l=[],s=[];if(!a.vertical_group_lock){if(a.vertical_group_lock=!0,""===(l=h.map(e.$headerIndexed,function(r){return r.hasClass(f.verticalGroupHeader)?1:""})).join(""))g(h(t).find("."+f.verticalGroupHide+",."+f.verticalGroupShow));else for(var c=0;c<t.length;c++)for(var u,v,d,p=!1,n=0;n<e.columns;n++)l[n]&&t[c].cells[n]?(u=h(t[c].cells[n]),v=G.isValueInArray(n,e.sortList),d=u.html(),v<0?g(u):p||d!==s[n]?0===v&&(p=!0,u.hasClass(f.verticalGroupShow)||u.addClass(f.verticalGroupShow),u.removeClass(f.verticalGroupHide),i)&&C(a,u,v?o:++o):(u.hasClass(f.verticalGroupHide)||u.addClass(f.verticalGroupHide),i&&C(a,u,o),u.removeClass(f.verticalGroupShow)),s[n]=d):o++;a.vertical_group_lock=!1}}h.extend(G.css,{verticalGroupHeader:"tablesorter-vertical-group",verticalGroupHide:"tablesorter-vertical-group-hide",verticalGroupShow:"tablesorter-vertical-group-show"}),G.addWidget({id:"vertical-group",priority:99,init:r,format:r})}(jQuery);return jQuery;}));
```
