## Supplementary Data SD1 for "Inferring binding specificities of human transcription factors with the wisdom of crowds": widget-view.min.js

```
(function(factory){if (typeof define === 'function' && define.amd){define(['jquery'], factory);} else if (typeof module === 'object' && typeof module.exports === 'object'){module.exports = factory(require('jquery'));} else {factory(jQuery);}}(function(jQuery){
!function(l){"use strict";var i,a,o,n=l.tablesorter,c=!1,r=n.view={copyCaption:function(e,t){r.removeCaption(e,t),0<e.$table.find("caption").length&&l(t.view_caption).text(e.$table.find("caption").text())},removeCaption:function(e,t){l(t.view_caption).empty()},buildToolBar:function(t,a){r.removeToolBar(t,a),r.copyCaption(t,a);var o=l(a.view_toolbar);l.each(a.view_layouts,function(e,t){var i=a.view_switcher_class,i=(e===a.view_layout&&(i+=" active"),l("<a>",{href:"#",class:i,"data-view-type":e,title:t.title}));i.append(l("<i>",{class:t.icon})),o.append(i)}),o.find("."+a.view_switcher_class).on("click",function(e){if(e.preventDefault(),l(this).hasClass("active"))return!1;o.find("."+a.view_switcher_class).removeClass("active"),l(this).addClass("active"),a.view_layout=l(this).attr("data-view-type"),!0===a.view_layouts[a.view_layout].raw?(r.remove(t,a),r.buildToolBar(t,a)):(!1===c&&r.hideTable(t,a),r.buildView(t,a))})},removeToolBar:function(e,t){l(t.view_toolbar).empty(),r.removeCaption(e,t)},buildView:function(e,t){r.removeView(e,t);var a=t.view_layouts[t.view_layout],o=l(a.container,{class:t.view_layout});n.getColumnText(e.$table,0,function(e){var n=a.tmpl,i=(l.each(l(e.$row).find("td"),function(e,t){var i={},a="{col"+e+"}",o=(l.each(t.attributes,function(e,t){i[t.nodeName]=t.nodeValue}),l(t).html()),o=l("<span />").append(l("<span/>",i).append(o));n=(n=n.replace(new RegExp(a,"g"),o.html())).replace(new RegExp("{col"+e+":raw}","g"),l(t).text())}),l(n));l.each(e.$row[0].attributes,function(e,t){"class"===t.nodeName?i.attr(t.nodeName,i.attr(t.nodeName)+" "+t.nodeValue):i.attr(t.nodeName,t.nodeValue)}),o.append(i)}),l(t.view_container).append(o),e.$table.triggerHandler("viewComplete")},removeView:function(e,t){l(t.view_container).empty()},hideTable:function(e){i=e.$table.css("position"),a=e.$table.css("bottom"),o=e.$table.css("left"),e.$table.css({position:"absolute",top:"-10000px",left:"-10000px"}),c=!0},init:function(e,t){!1!==t.view_layout&&void 0!==t.view_layouts[t.view_layout]&&(!1===c&&r.hideTable(e,t),e.$table.on("tablesorter-ready",function(){r.buildToolBar(e,t),r.buildView(e,t)}))},remove:function(e,t){r.removeToolBar(e,t),r.removeView(e,t),e.$table.css({position:i,top:a,left:o}),c=!1}};n.addWidget({id:"view",options:{view_toolbar:"#ts-view-toolbar",view_container:"#ts-view",view_caption:"#ts-view-caption",view_switcher_class:"ts-view-switcher",view_layout:!1,view_layouts:{}},init:function(e,t,i,a){r.init(i,a)},remove:function(e,t,i){r.remove(t,i)}})}(jQuery);return jQuery;}));
```
