## Supplementary Data SD1 for "Inferring binding specificities of human transcription factors with the wisdom of crowds": README.md

tablesorter (FORK) is a jQuery plugin for turning a standard HTML table with THEAD and TBODY tags into a sortable table without page refreshes. tablesorter can successfully parse and sort many types of data including linked data in a cell. This forked version adds lots of new enhancements including: alphanumeric sorting, pager callback functons, multiple widgets providing column styling, ui theme application, sticky headers, column filters and resizer, as well as extended documentation with a lot more demos.

[![NPM Version][npm-image]][npm-url] [![devDependency Status][david-dev-image]][david-dev-url] [![zenhub-image]][zenhub-url]

### Notice!

* Because of the change to the internal cache, the tablesorter v2.16+ core, filter widget and pager (both plugin &amp; widget) will only work with the same version or newer files.

### [Documentation](https://mottie.github.io/tablesorter/docs/)

* See the [full documentation](https://mottie.github.io/tablesorter/docs/).
* All of the original documentation has been included.
* Information from my blog post on [undocumented options](https://wowmotty.blogspot.com/2011/06/jquery-tablesorter-missing-docs.html) and lots of new demos have also been included.
* Change log moved from included text file into the [wiki documentation](https://github.com/Mottie/tablesorter/wiki/Changes).

### Questions?

[![irc-image]][irc-url] [![slack-image]][slack-url] [![stackoverflow-image]][stackoverflow-url]

* Check the [FAQ](https://github.com/Mottie/tablesorter/wiki/FAQ) page.
* Search the [main documentation](https://mottie.github.io/tablesorter/docs/) (click the menu button in the upper left corner).
* Search the [issues](https://github.com/Mottie/tablesorter/issues) to see if the question or problem has been brought up before, and hopefully resolved.
* If someone is available, ask your question in the `#tablesorter` IRC channel at freenode.net.
* Ask your question at [Stackoverflow](https://stackoverflow.com/questions/tagged/tablesorter) using a tablesorter tag.
* Please don't open a [new issue](https://github.com/Mottie/tablesorter/issues) unless it really is an issue with the plugin, or a feature request. Thanks!

### Demos

* [Basic alpha-numeric sort Demo](https://mottie.github.io/tablesorter/).
* Links to demo pages can be found within the main [documentation](https://mottie.github.io/tablesorter/docs/).
* More demos & playgrounds - updated in the [wiki pages](https://github.com/Mottie/tablesorter/wiki).

### Features

* Multi-column alphanumeric sorting and filtering.
* Multi-tbody sorting - see the [options](https://mottie.github.io/tablesorter/docs/index.html#options) table on the main document page.
* Supports [Bootstrap v2-4](https://mottie.github.io/tablesorter/docs/example-option-theme-bootstrap-v3.html).
* Parsers for sorting text, alphanumeric text, URIs, integers, currency, floats, IP addresses, dates (ISO, long and short formats) &amp; time. [Add your own easily](https://mottie.github.io/tablesorter/docs/example-parsers.html).
* Inline editing - see [demo](https://mottie.github.io/tablesorter/docs/example-widget-editable.html).
* Support for ROWSPAN and COLSPAN on TH elements.
* Support secondary "hidden" sorting (e.g., maintain alphabetical sort when sorting on other criteria).
* Extensibility via [widget system](https://mottie.github.io/tablesorter/docs/example-widgets.html).
* Cross-browser: IE 6.0+, FF 2+, Safari 2.0+, Opera 9.0+, Chrome 5.0+.
* Small code size, starting at 25K minified.
* Works with jQuery 1.2.6+ (jQuery 1.4.1+ needed with some widgets).
* Works with jQuery 1.9+ (`$.browser.msie` was removed; needed in the original version).

### Licensing

* Copyright (c) 2007 Christian Bach.
* The original version can be found at [http://tablesorter.com](http://tablesorter.com), or on [GitHub](https://github.com/christianbach/tablesorter).
* Dual licensed under the [MIT](https://opensource.org/licenses/mit-license.php) or [GPLv2](https://www.gnu.org/licenses/gpl-2.0.html) licenses (pick one).

### Download

* Get all files: [zip](https://github.com/Mottie/tablesorter/archive/master.zip) or [tar.gz](https://github.com/Mottie/tablesorter/archive/master.tar.gz).
* Use [bower](https://bower.io/): `bower install jquery.tablesorter`.
* Use [node.js](https://nodejs.org/): `npm install tablesorter`.
* CDNJS: https://cdnjs.com/libraries/jquery.tablesorter
* jsDelivr: http://www.jsdelivr.com/?query=tablesorter

### Related Projects

* [Plugin for Rails](https://github.com/themilkman/jquery-tablesorter-rails). Maintained by [themilkman](https://github.com/themilkman).
* [UserFrosting](https://www.userfrosting.com) (A secure, modern user management system for PHP that uses tablesorter) by [@alexweissman](https://github.com/alexweissman).
* [Grav CMS](https://getgrav.org/): `bin/gpm install tablesorter` ([ref](https://github.com/Perlkonig/grav-plugin-tablesorter)).
* [tablesorter-pagercontrols](https://github.com/isg-software/tablesorter-pagercontrols) &ndash; programmatically adds pager controls below a table and applies the pager add-on for large HTML tables by [isg-software](https://github.com/isg-software).

### Contributing

If you would like to contribute, please...

1. Fork.
2. Make changes in a branch & add unit tests.
3. Run `grunt test` (if qunit fails, run it again - it's fickle).
4. Create a pull request.

### Special Thanks

* Big shout-out to [Nick Craver](https://github.com/NickCraver) for getting rid of the `eval()` function that was previously needed for multi-column sorting.
* Big thanks to [thezoggy](https://github.com/thezoggy) for helping with code, themes and providing valuable feedback.
* Big thanks to [ThsSin-](https://github.com/TheSin-) for taking over for a while and also providing valuable feedback.
* Thanks to [prijutme4ty](https://github.com/prijutme4ty) for numerous contributions!
* Also extra thanks to [christhomas](https://github.com/christhomas) and [Lynesth](https://github.com/Lynesth) for help with code.
* And, of course thanks to everyone else that has [contributed](https://github.com/Mottie/tablesorter/blob/master/AUTHORS), and continues to contribute through pull requests and open issues to this forked project!

[npm-url]: https://npmjs.org/package/tablesorter
[npm-image]: https://img.shields.io/npm/v/tablesorter.svg
[david-dev-url]: https://david-dm.org/Mottie/tablesorter?type=dev
[david-dev-image]: https://img.shields.io/david/dev/Mottie/tablesorter.svg
[zenhub-url]: https://zenhub.io
[zenhub-image]: https://cdn.rawgit.com/Mottie/tablesorter/master/docs/img/zenhub-badge.svg

[irc-url]: https://kiwiirc.com/client/irc.freenode.net#tablesorter
[irc-image]: https://img.shields.io/badge/irc-%23tablesorter-yellowgreen.svg
[slack-url]: https://join.slack.com/t/tablesorter/shared_invite/enQtNDc2NTczODg0NjkzLWRmYzE2NjIzNjM2ZWE1NzkyNzFkNjM4NjY0OTIyOTM3NDRmNWRhMjgxMTk5MjFhNzM3NWIzMWU1ZTc1NTZlOGE
[slack-image]: https://img.shields.io/badge/Slack-join%20chat-yellowgreen.svg
[stackoverflow-url]: http://stackoverflow.com/questions/tagged/tablesorter
[stackoverflow-image]: https://img.shields.io/badge/stackoverflow-tablesorter-blue.svg

### Recent Changes

View the [complete change log here](https://github.com/Mottie/tablesorter/wiki/Changes).

#### <a name="v2.32.0">Version 2.32.0</a> (2024-08-13)

* Core:
  * Remove ie8 support (Grunt limitation)
  * Initialize headers as `null`
* Grouping: Fix self-closing icon tags to support jQuery 3.5+
* Parser-input-select: Don't use updateCell if cell has `parser-false` class

#### <a name="v2.31.3">Version 2.31.3</a> (2020-03-03)

* Core:
  * Cache parsed value in colspan. Fixes [issue #1708](https://github.com/Mottie/tablesorter/issues/1708).
* AlignChar:
  * Tweak demo css. See [isssue #1713](https://github.com/Mottie/tablesorter/issues/1713).
* Pager:
  * Restore first row in cacheIndex. Fixes issues [#1714](https://github.com/Mottie/tablesorter/issues/1714) & [#1710](https://github.com/Mottie/tablesorter/issues/1710).
* Docs:
  * Update to jQuery 3.4.1 & migrate 3.1.0.
  * Update to Bootstrap 4.4.1.
  * CSS cleanup - code font size was off.

#### <a name="v2.31.2">Version 2.31.2</a> (2019-12-01)

* Column selector:
  * Fix scroller widget compatibility. See [issue #1682](https://github.com/Mottie/tablesorter/issues/1682).
* Filter:
  * Update select2 regexp. See [issue #1497](https://github.com/Mottie/tablesorter/issues/1497). Fixed in [PR #1677](https://github.com/Mottie/tablesorter/pull/1677); thanks [@cwisdo](https://github.com/cwisdo)!
  * Prevent search on enter if filters are unchanged. See [issue #1631](https://github.com/Mottie/tablesorter/issues/1631). Fixed in [PR #1642](https://github.com/Mottie/tablesorter/pull/1642); thanks [@larsbonczek](https://github.com/larsbonczek)!
* Math
  * Add math ignore to cells; it was documented, but not added previously. See [issue #1526](https://github.com/Mottie/tablesorter/issues/1526).
* Docs:
  * Various fixes.
  * Updated trigger sort page. See [issue #1641](https://github.com/Mottie/tablesorter/issues/1641).
  * Remove invalid widget option. See [issue #1581](https://github.com/Mottie/tablesorter/issues/1581).
