## Supplementary Data SD1 for "Inferring binding specificities of human transcription factors with the wisdom of crowds": SECURITY.md

# Security Policy

## Supported Versions
Security updates are applied only to the latest release.

## Reporting a Vulnerability
If you have discovered a security vulnerability in this project, please report it privately. **Do not disclose it as a public issue.** This gives us time to work with you to fix the issue before public exposure, reducing the chance that the exploit will be used before a patch is released.

Please disclose it at our [security advisory](https://github.com/Mottie/tablesorter/security/advisories/new).

This project is maintained by a team of volunteers on a reasonable-effort basis. As such, vulnerabilities will be disclosed in a best effort base.
