## Supplementary figures and images for "Inferring binding specificities of human transcription factors with the wisdom of crowds"

### black-asc.gif

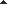

### black-desc.gif

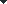

### black-unsorted.gif

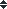

### bootstrap-black-unsorted.png

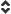

### bootstrap-white-unsorted.png

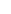

### dropbox-asc-hovered.png

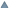

### dropbox-asc.png

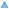

### dropbox-desc-hovered.png

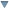

### dropbox-desc.png

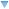

### first.png

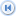

### green-asc.gif

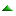

### green-desc.gif

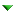

### green-unsorted.gif

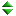

### ice-asc.gif

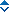

### ice-desc.gif

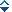

### ice-unsorted.gif

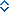

### last.png

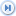

### loading.gif

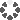

### metro-black-asc.png

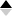

### metro-black-desc.png

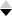

### metro-loading.gif

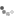

### metro-unsorted.png

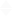

### metro-white-asc.png

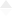

### metro-white-desc.png

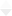

### next.png

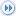

### prev.png

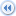

### white-asc.gif

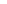

### white-desc.gif

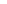

### white-unsorted.gif

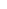
