## Supplementary Figures SF1-SF10 for "Inferring binding specificities of human transcription factors with the wisdom of crowds"

**Supplementary Figure SF1.** IBIS Challenge design and disciplines.
**A:** Overall data preparation workflow: data from five independent platforms is preprocessed with the Codebook MEX pipeline and then supplied to the BIBIS package to prepare the train/test slices. **B:** Setup of the disciplines: depending on the TF, either genomic or synthetic data are provided for training PWM and AAA models, which are then evaluated using the same test data and performance metrics.

**A**

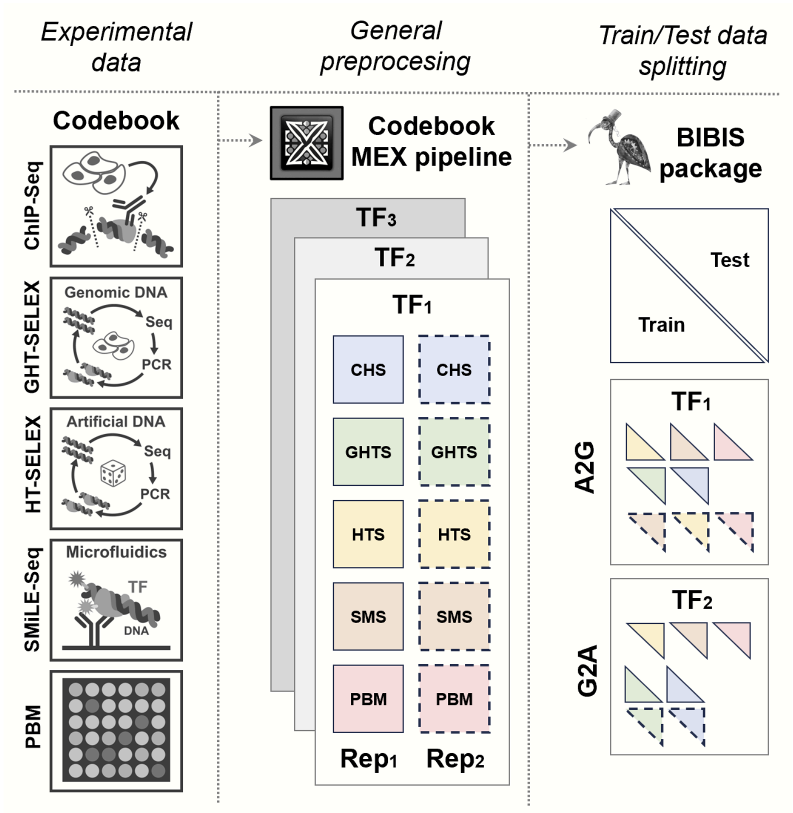


**B**

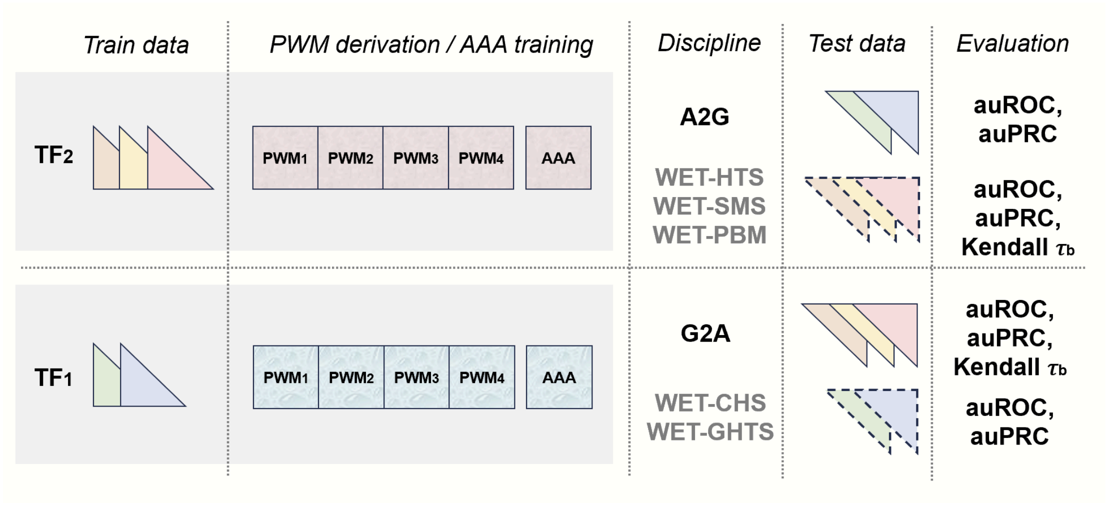

**Supplementary Figure SF2.** IBIS participants and winners in WET disciplines.
The bubble size is proportional to the percentage of cases where the team received a particular rank across 1000 random subsamples, each subsample had a 25% chance for each combination of a TF and a data type to be excluded from ranking. X-axis: ranks.

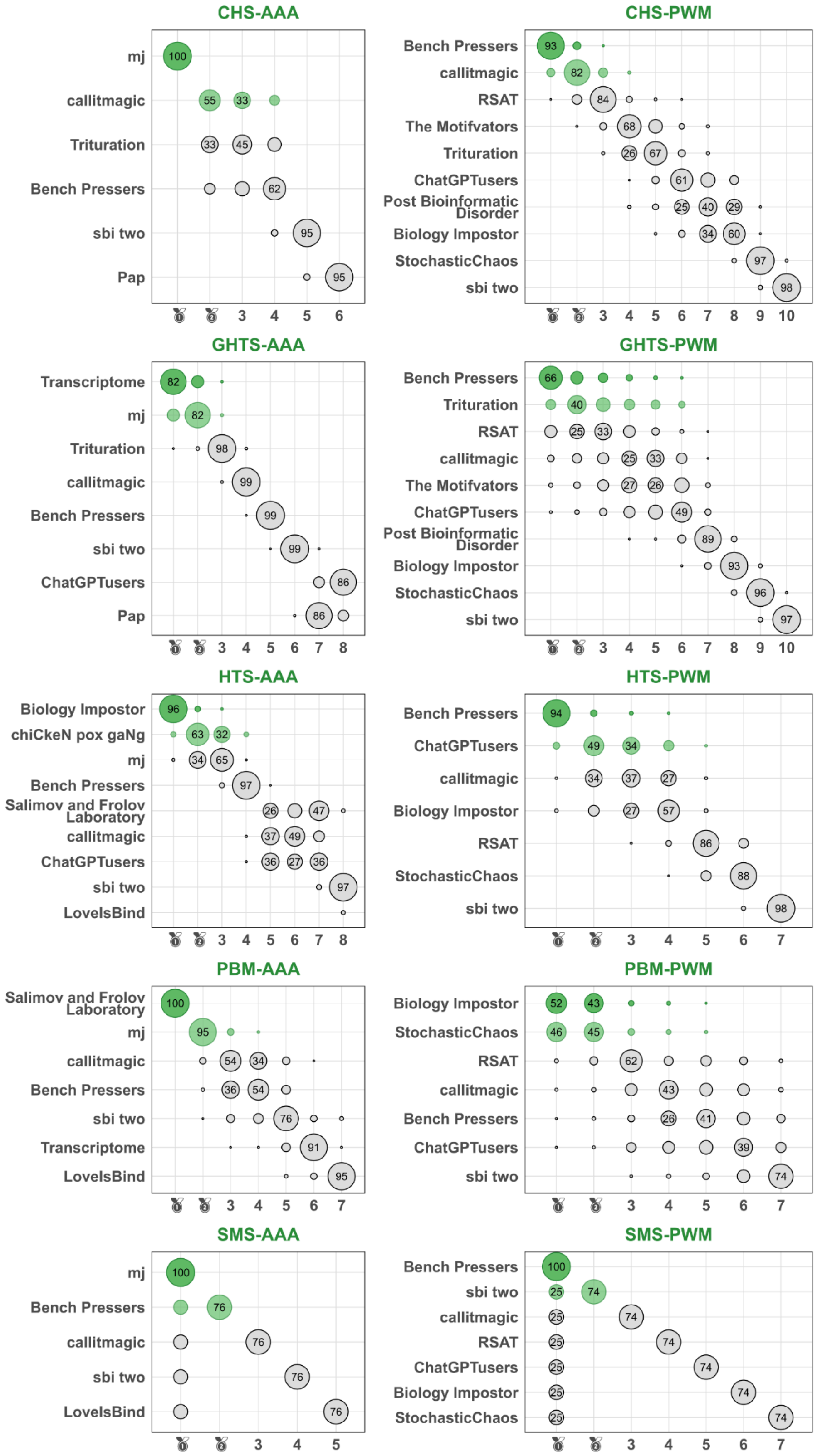

**Supplementary Figure SF3.** Normalized performance of different teams across TFs if trained and tested with the data from a single assay type. **A**: WET-HTS, -PBM, -SMS, **B**: WET-CHS, -GHTS. Color scale: mean value across test datasets and different performance metrics. The values are $\log_2$-ratios versus the values reached by the top-ranked PWM obtained from IBIS data by the challenge organizers; n.a.: solution for the combination of a TF and a data type not submitted for evaluation. Values exceeding zero (improvement over the strong baseline) are explicitly labeled.

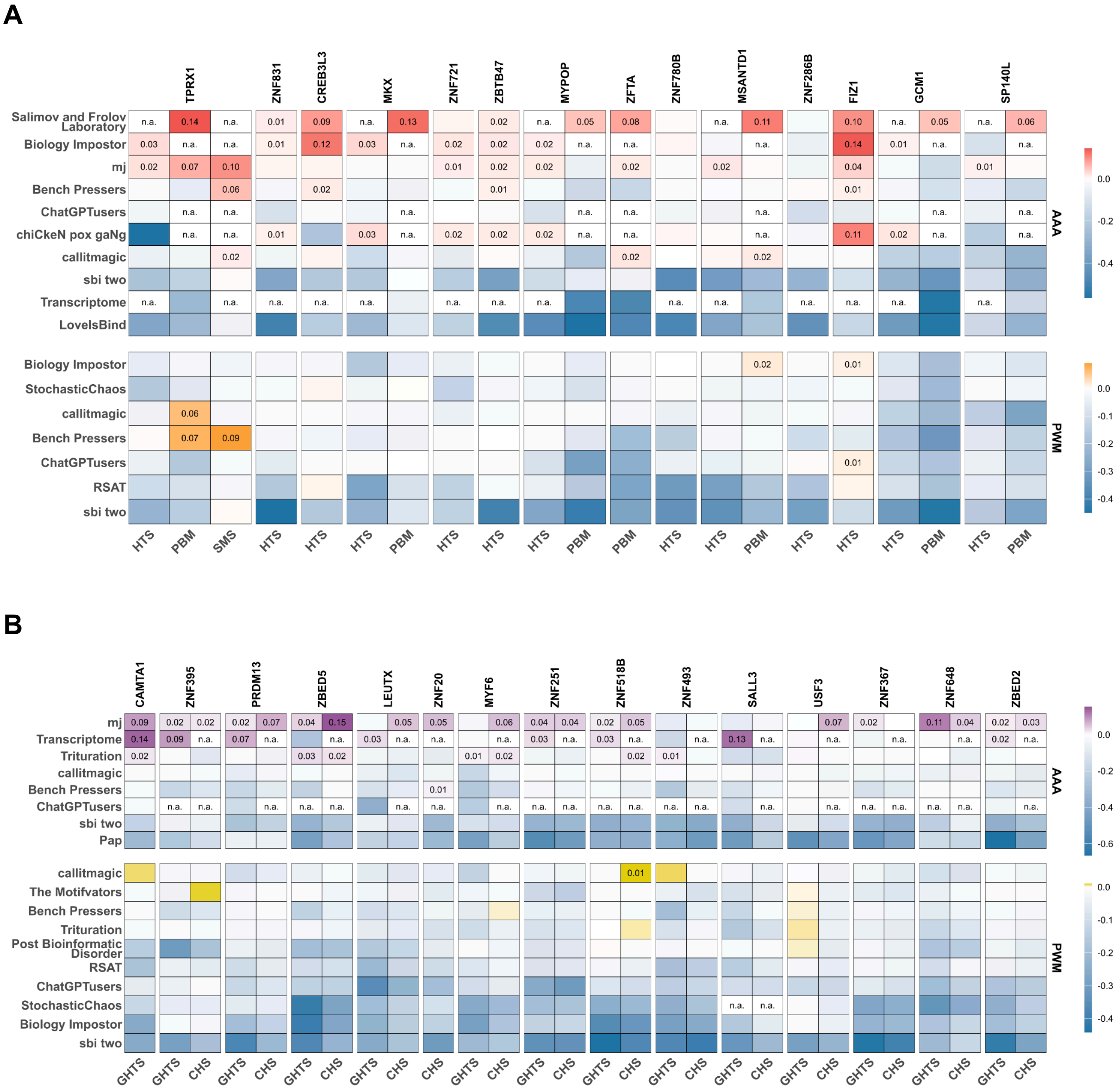

**Supplementary Figure SF4.** Raw values of performance metrics of IBIS solutions on the test data. For AUROC and AUPRC, the values are the mean across all types of genomic (A2G) or artificial (G2A) test data and all types of negative datasets. Teams that submitted incomplete solutions, skipping some TFs or some data types, are not shown. Maximal values for each TF are labeled on the plot. For a fair comparison, the MEX set of PWMs, constructed by the organizers, includes 4 PWMs per TF. **A**, **B**: A2G AUROC, AUPRC. **D**, **E**, **F**: G2A AUROC, AUPRC, and Kendall's $\tau_b$.

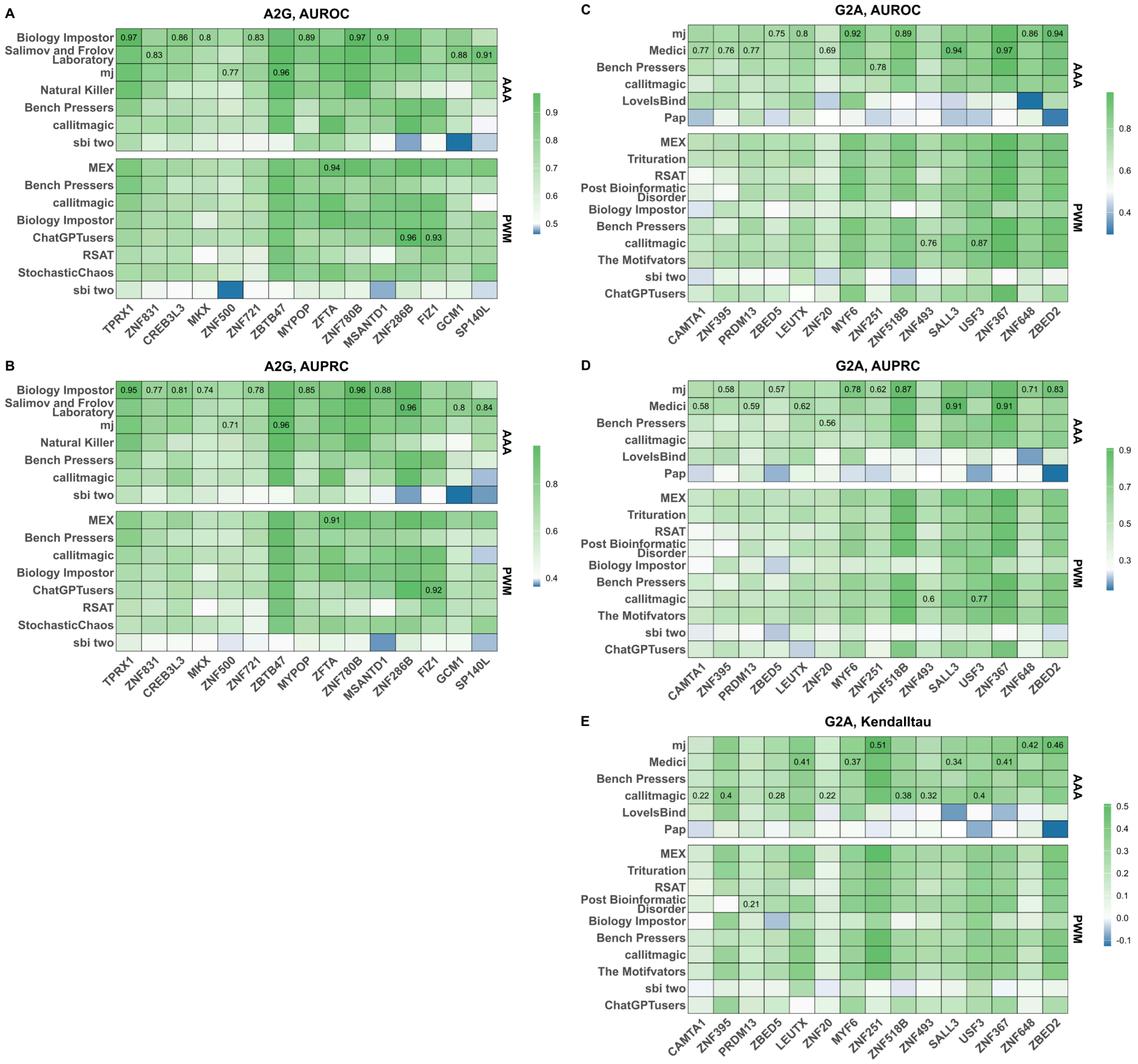

**Supplementary Figure SF5.** Examples of concordance curves obtained by applying selected IBIS models to recognize the allelic preferences of transcription factors at the allele-specific binding sites. **A**: A2G, GCM1 TF. **B**: G2A, MYF6 TF. Each series of plots includes the top 3 models (on average), MEX, and the lower-scoring model on average (sbi two), for comparison. An expectation across random predictions is the dashed line at 0.5 yielded by random guesses of allelic preferences at any number of ASBs (defined prediction threshold, which determines the number of ASBs with predicted binding sites, X axis).

**A**

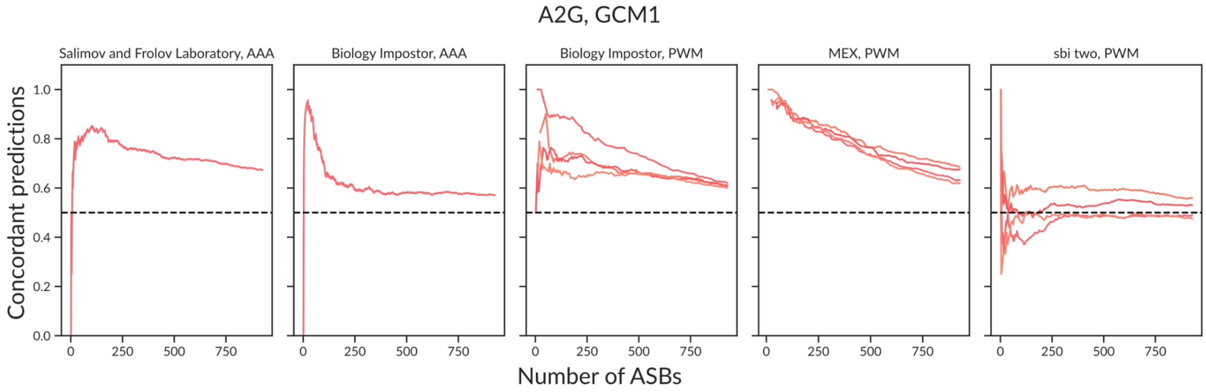


**B**

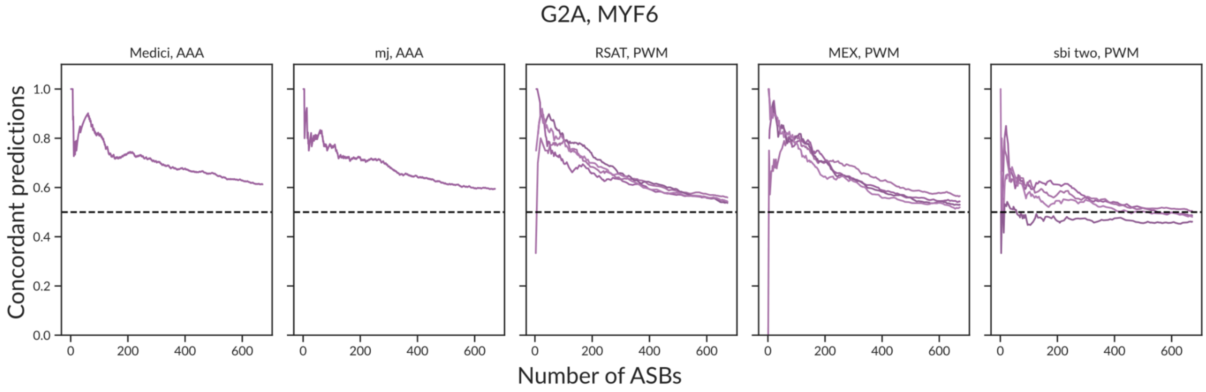

**Supplementary Figure SF6.** Regression and classification performance of TFBS models agree well on HT-SELEX data. **A**: Triple-A models, **B**: PWMs. Each point belongs to a pair of performance metrics for an evaluated model. AUROC and AUPRC (X-axes) and Kentall $\boldsymbol{\tau_b}$ (Y-axes) were averaged across all types of negative datasets; for PWMs, results for alternative scanning (sum-occupancy, best-hit) were also averaged. The Pearson correlation coefficient ρ and the number of points *n* are labeled on each plot.

**A**

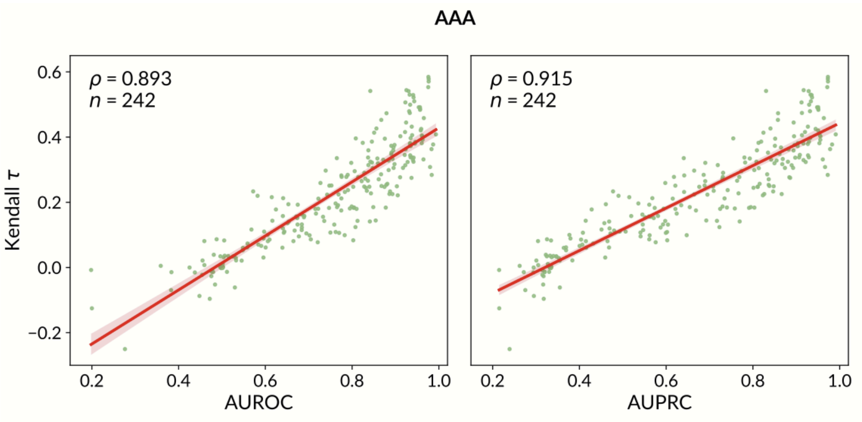


**B**

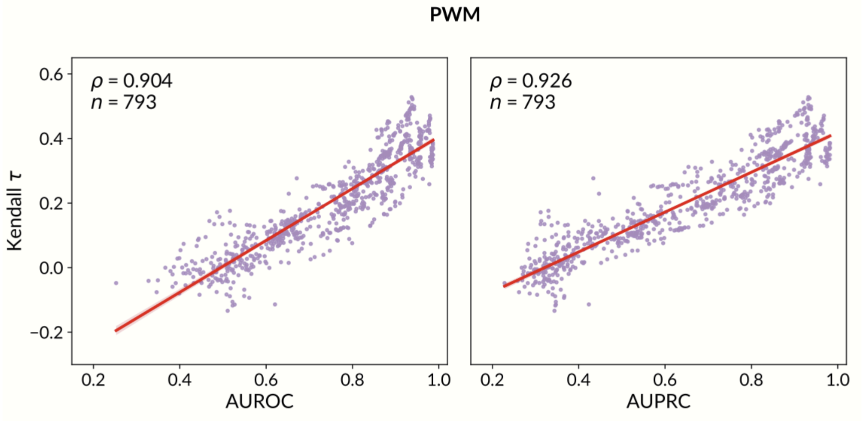

**Supplementary Figure SF7.** Comparison of PWM performance using two alternative scanning modes. **Top row**: AUPRC, **Bottom row**: AUROC. Each point belongs to a pair of performance metrics; metrics for alternative negative datasets are used independently. X-axis: *best-hit* scanning, Y-axis: *sum-occupancy* scanning. The number of points *n* (including the number above and below the diagonal) and p-value (*p*) of the Wilcoxon signed-rank test are labeled on each plot.

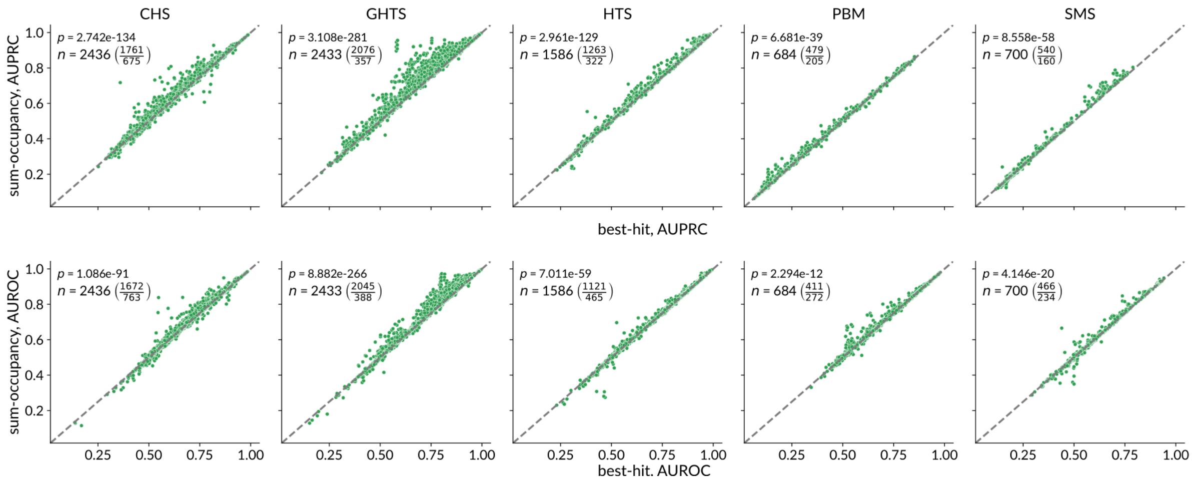


**Supplementary Figure SF8.** Distribution of achieved performance metrics when testing against the different types of negative data.

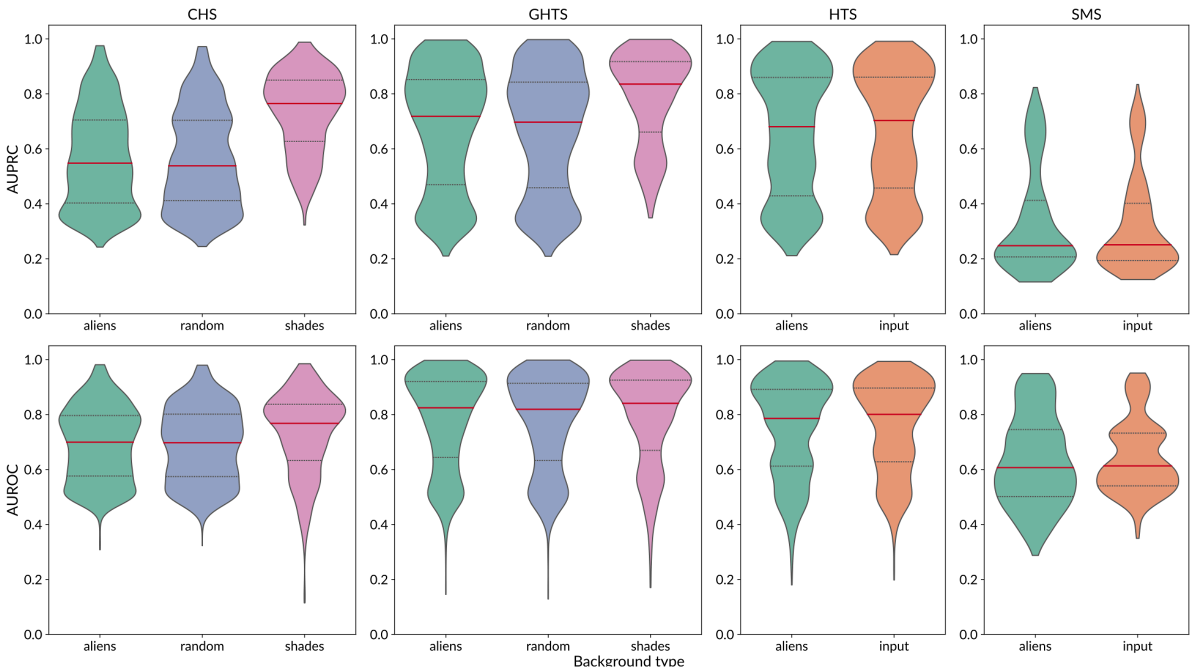

**Supplementary Figure SF9.** IBIS post-challenge benchmarking of LegNet variants.
**A, B**: Performance in the primary IBIS benchmarks. A - A2G, B - G2A. Color scale: mean value across test datasets and different performance metrics. The values are $log_2$-ratios versus the top-ranked PWMs obtained from IBIS data. Values exceeding zero (improvement over the strong PWM baseline) are explicitly labeled. **C, D:** Performance in recognizing the allelic preferences of individual transcription factors (C - A2G, D - G2A). Colors and plot structure are the same as in Figure 3.

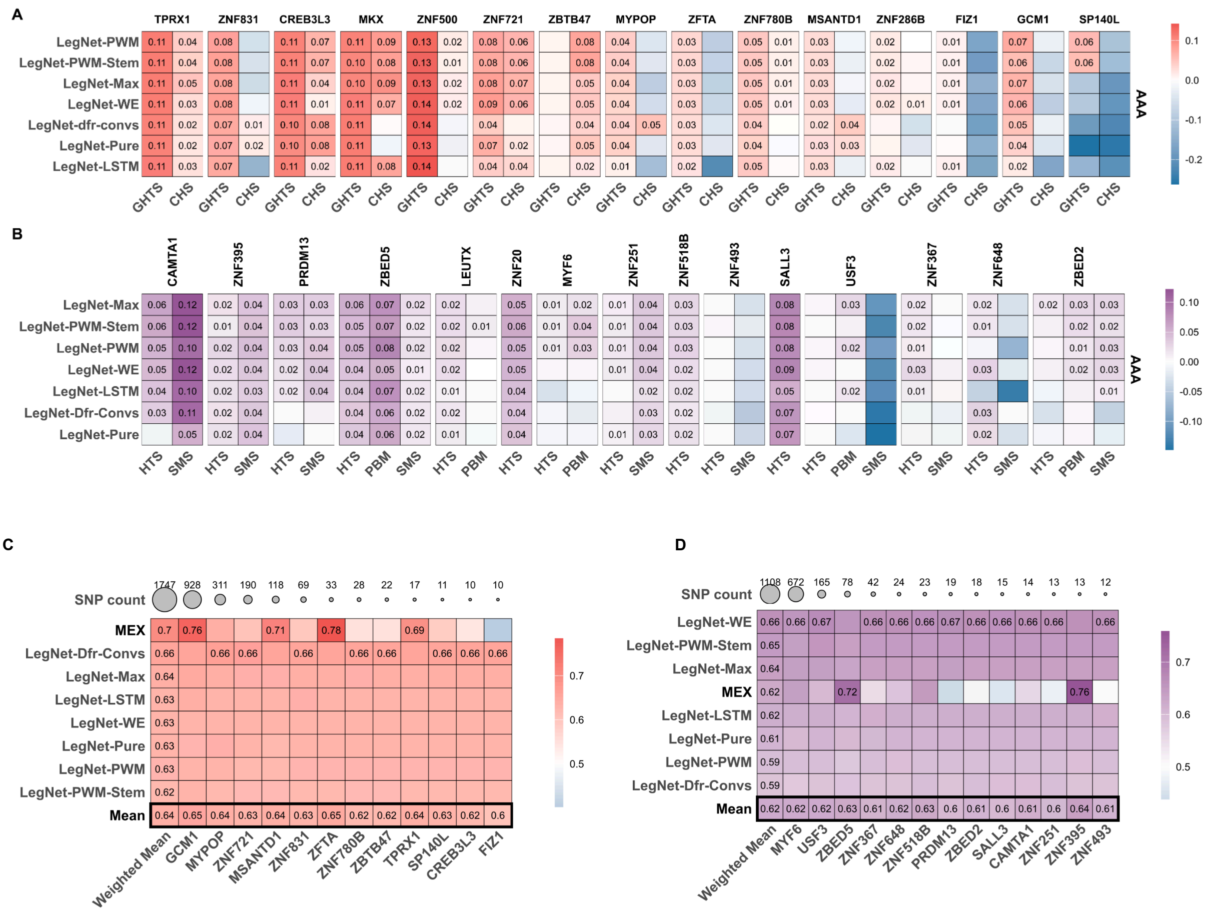

**Supplementary Figure SF10.** Rationale for replacing average pooling layers with max pooling counterparts. **Top:** The CNN model (or a hybrid architecture) is trained to predict TF binding using sequences of length *L*. The model learns that the sum of mean activations across feature maps should exceed 9 to indicate TF binding. **Bottom:** When tested on a sequence of length *2L*, where the TF binding site remains the same size, the mean activation decreases due to the extended non-signal region. This leads the model to incorrectly conclude that there is no TF binding site in the sequence (false negative).

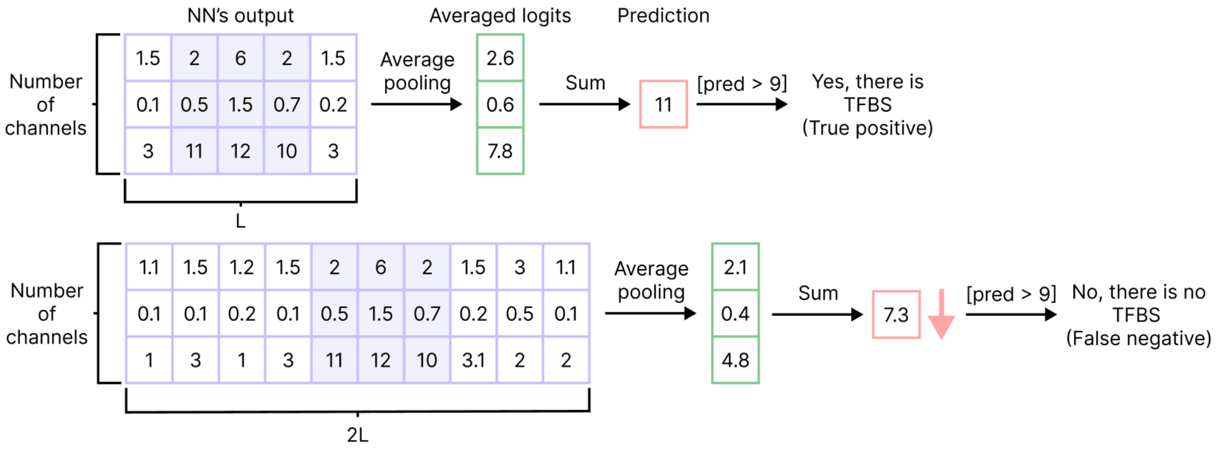
