## Supplementary Methods for "Inferring binding specificities of human transcription factors with the wisdom of crowds"

### Brief description of the IBIS challenge solutions designed by the participants

The detailed method descriptions as provided by the participants are included in the primary IBIS Zenodo Record[1]. The links to the GitHub code repositories, when available, are provided in **Supplementary Table ST2**.

#### PWM submissions

**Bench Pressers.** We sought to integrate classic and deep learning methods not only to maximize performance in the IBIS challenge, but also to learn the conditions under which one class of methods outperforms another. We applied a previously developed convolutional neural network (CNN)-based approach for motif discovery[2]. In the G2A-PWM task, we directly used our method to discover motifs in the CHS and GHTS datasets. We then used a classical algorithm, STREME[3], to refine the deep learned motifs. The pool of candidate motifs is then evaluated in a cross-validation scheme, and we included the 4 motifs most predictive of binding within and across assays in the final submission. For the A2G-PWM task, we focused exclusively on HTS data. We used our CNN method to discover motifs in all provided HT-SELEX cycles. As the first few cycles have relatively low enrichment of motif-containing sequences, we used a 2-component Gaussian mixture model to derive a threshold for input sequences to be included in the final motif. One motif for each cycle was included in the final A2G-PWM submission.

**Biology Impostor.** We generated PWMs using RAP[4] for PBM data, and HTS-IBIS[5] for HTS data. For the other data types, we developed and applied similar algorithms as follows. In each experiment, we ranked the k-mers of nucleotides according to their binding strength (where k was a hyperparameter of the process). For example, in HT-SELEX, we ranked the 8-mers based on their enrichment between cycle 1 and cycle 4. After ranking the k-mers, we aligned the top k-mers (where the number of top k-mers is another hyper-parameter of the process) to the top k-mer. Then, we used the alignment results to create the core PWM by weighing each nucleotide with the binding strength of the k-mer in which it appeared. Finally, if the flanking regions of the core positions were greater than a predefined information threshold (another hyperparameter), we extended the core PWM.

**callitmagic.** We used the MEME Suite (MEME for motif discovery, AME for motif enrichment analysis, and FIMO for motif finding)[6] to generate position weight matrices (PWMs) and identify motif occurrences. Our approach used predefined parameter grids for motif generation and cleaning the results based on E-value thresholds. A greedy search was conducted among numerous motifs generated by MEME on differently pre-processed and aggregated data. The decision rule for top-4 motifs selection was based on AME results using control sequences.

**ChatGPTusers.** To build PWMs from HTS data, we used the de novo motif discovery algorithm Autoseed[7]. Autoseed finds gapped and ungapped subsequences that represent local maxima, that is, are more enriched than closely related subsequences. For GHTS data, we employed HOMER[8].

**Post Bioinformatic Disorder.** For motif discovery, we employed ChIPMunk[9]. Different combinations and subsets of CHS and GHTS data were fed into the program and tested during the Leaderboard stage. In the end, three motifs were used in the submission: the first was based on merged CHS and GHTS datasets, the second was based on CHS data only, and the third was based on GHTS data only.

**RSAT.** We performed the analysis using the *Regulatory Sequence Analysis Tools* (RSAT)[10], Docker release 20240808. Training sets were processed with *peak-motifs*, which identifies overrepresented k-mers in the sequences, aligns them to generate seed position-count matrices (PCMs), derives secondary PCMs by collecting matching sites in the training sequences, and clusters the resulting matrices to remove redundancy. We further applied a novel genetic algorithm-based method to optimise matrix performance, assessed by their ability to discriminate positive from negative sequence sets using the area under the ROC curve.

**sbi two.** We used a structure-based learning approach to predict the binding preferences of each Transcription Factor (TF). For each TF, sequence information was downloaded from UniProt. PWMs were then generated using ModCRE[11] directly where possible. When no 3D structures were producible through ModCRE, they were generated with AlphaFold3[12] instead, before having PWMs predicted by ModCRE. In parallel, PWMs were predicted from experimental data using the MEME suite. These PWMs were then used to refine those predicted with ModCRE.  Upon completion of the refinement process, four PWMs were selected per TF for submission. To select these, we applied a scoring function that ranked refined PWMs according to their normalized width, number of experimental PWMs incorporated in the refinement process, and the similarity between the original ModCRE PWM and experimental PWMs used for refinement. After ranking, the top 4 PWMs were selected while maintaining uniqueness across the selection. PWMs predicted with the aid of AlphaFold3 modeling were not subjected to this Ranking, but their uniqueness was still ensured. Next, the prediction performance of the top 4 PWMs generated was compared to PWMs derived from AlphaFold3-derived binding sequences. If the AlphaFold3-derived PWM outperformed the ModCRE-created PWMs, it was included in the submission along with 3 remaining ModCRE motifs.

**StochasticChaos.** We used SeSiMCMC[13], a Gibbs sampler for de novo identification of similar nucleotide substrings in the input. The SeSIMCMC paper was published 20 years ago, and our main goal of participation in the challenge was to answer whether the algorithms of this class are still usable in the modern world of deep learning models. The default search parameters were used, with two adjustments. For SMiLE-seq, we set the probability for an input sequence to be empty (i.e, not to carry the motif) to 0.9 instead of the default 0.1. For all the runs, we set the number of chains (attempts) to 7 instead of the default 3. The time limit was set to 10000 seconds (the default is 1000). We added the --ibis command-line switch to the SeSiMCMC software to generate the IBIS challenge-formatted output.

**The Motifvators.** For motif discovery, we used HOMER[8] to identify enriched sequence motifs from repeat-masked hg38 genomic regions. Foreground and background sets were derived from GHT-SELEX peaks, with background sequences generated by shifting coordinates to avoid binding regions. The most significant motifs (motif1) were selected as position weight matrices (PWMs). These PWMs captured key transcription factor binding preferences and served as the foundation for downstream Triple-A modelling, see below.

**Trituration.** We developed a computational pipeline for de novo motif discovery in ChIP-seq data. The datasets were split into independent training and test sets by sorting peaks by significance and dividing them into even (test) and odd (training) indices. To ensure reliable background modeling, we generated negative samples by selecting random genomic regions with a dinucleotide composition closely matching that of the training set. For motif discovery, we employed STREME[3], scanning motif lengths from 6 to 24 bp (in steps of 2). The most enriched motif at each length was retained, and the performance was quantified using three complementary metrics: auROC, auPRC, and partial auROC (pAUC) at FPR=0.001; the pAUC score was normalized by 0.001 to fit into the range between 0 and 1. The optimal motif length was then chosen according to the highest combined score across these metrics.

### Triple-A submissions

**Bench Pressers.** The AAA submission was based on the PWM solution, see the description above.

**Biology Impostor.** We utilized HTS data only since SMS data was less informative, and PBM data was missing for several proteins. We assigned the sequences of each cycle an integer label, starting from 1 for cycle 1, while assigning alien and shuffled sequences the label 0. We removed duplicate sequences from each cycle. We used an ensemble of multiple convolutional neural networks. The convolutional layer had filters of variable widths applied directly to the original one-hot encoded sequences. This layer was followed by global max pooling and a cascade of fully connected layers. This architecture ensures that our models are largely insensitive to the length of the input sequences, which is crucial for the subsequent steps of our approach. To apply our models for prediction, we assumed that the binding site mostly occurs at the center of the sequence. This pattern can be represented as a probability distribution with the highest density of binding sites at the center. We developed a method that balances accuracy and speed, as predicting over shifting windows and selecting the maximum can become impractical for longer sequences. In the GHTS experiments, we averaged our model's predictions across the central 301, 101, and 51 nucleotides, while for the CHS experiment, we used the central 301, 151, and 51 nucleotides.

**callitmagic.** Top-performing PWMs (see above) were used to make predictions with FIMO for the IBIS AAA track.

**ChatGPTusers.** To obtain the predictions, PWMs (see above) were supplied to MOODS[14]. MOODS was executed with the P-value threshold of 1, which means it provided several scores for all reads in the test file. After that, the maximum score for each read was chosen. Min-max normalization of scores was applied for fitting data in the range from 0 to 1. This normalization was chosen as it showed the best results in internal benchmarking using the training data. The following GitHub repos host the code used in preparing the IBIS solution: https://github.com/jutaipal/kmercount_with_localmax, https://github.com/jutaipal/multinomial_motif_generator, https://github.com/jttoivon/moder2, https://github.com/jhkorhonen/MOODS.

**LoveIsBind.** We applied a Gradient Boosting approach using CatBoost[15] to predict PBM, HTS, and SMS binding peaks based on 3–7 mer frequency features, which were generated using the oligonucleotideFrequency function from the Biostrings package. We then used a hybrid modeling strategy: for TFs within well-defined families, a single model was trained by pooling the corresponding peaks and using the specific TF name as a categorical feature (to enable a TF-specific effect within the family-level model), while all other TFs used a dedicated, singular (i.e., non-pooled) model. Training data was sourced from the central 40 bp regions of ChIP-Seq and GHT-SELEX peaks. The 35 bp sequences for PBM models were derived by trimming the ends of the 40 bp sequences generated for the HTS and SMS models. The negative training set was constructed from two sources: GC-matched random genomic regions, generated using the *genNullSeqs* function from the gkmSVM package, and alien peaks from unrelated TF families. The multiplicity of random negatives was set according to the assay type: 10-fold for PBM, 2.5-fold for SMS, and 1-fold for HTS. For alien negatives, binding sites from a pool of unrelated TFs were combined. From this collection, a number of peaks were dynamically sampled to be approximately equal in number to the set of GC-matched random regions, ensuring the final negative set was approximately balanced between the two sources. Finally, to make the models invariant to strand orientation, all training data were augmented by including the reverse complement of each sequence, created using the reverseComplement function from the Biostrings package.

**Medici.** To identify transcription factor binding sites, we employed a convolutional neural network (CNN) with a fixed architecture and parallel convolutional kernels, depthwise separable convolutions, and squeeze-and-excitation blocks. Training incorporated *Reverse_Complement* and *Cyclic_Shift* augmentations, and the GC-content of negatives was matched to that of positives. Method-specific positive/negative ratios were used, with AdamW optimization and a linear learning rate scheduler. Separate models were trained for each protein and assay, with epochs adjusted to the respective dataset size. Simple central cropping of sequences proved as effective as motif localization methods, and in the end, the streamlined CNN consistently outperformed more complex variants.

**mj.** For each TF, a TCN[16] model was trained on all available sequences. The TCN model accumulates the context window of 31 nucleotides and then uses the global max pooling to predict a score for a given sequence. Alien sequences and randomly sampled regions from the hg38 genome assembly were used as negatives. During training, each sequence had a 50% chance to be replaced with its reverse complement. For predictions, each sequence and its reverse-complementary pair were processed by the model, and the resulting scores were used to calculate the weighted mean as the final prediction.

**Natural Killer.** Our method predicts transcription factor (A2G) binding using 1D ResNet[17] architectures trained on binary-encoded DNA sequences, where each nucleotide is represented by one of five input channels. ResNet18/34 models were trained on short fragments (40 bp) and applied to long genomic sequences using a sliding window (step 20) with maximum probability aggregation. Two rounds of pseudo-labeling with cross-validation were employed to enhance prediction quality. The models were implemented in PyTorch with BCEWithLogitsLoss.

**Pap.** The nucleotides were one-hot encoded, and the input sequence length was 100 bp. Zero-padding was employed in cases of shorter sequences. A validation set for hyper-parameter tuning comprised 10% of the training data. Convolutional neural networks were trained for each TF. Each network consisted of two convolutional layers followed by a global max-pooling layer. Then, two fully connected layers were used for classification. The output layer's two units used the softmax activation function and binary cross-entropy as a loss function. L1 was applied as an activity regularizer with a value of 5e-5, and L2 (5e-4) as a kernel regularization in the convolutional layers. A dropout probability of 0.5 was employed in the classifier layers, which helped mitigate overfitting. A batch size of 128 and an early-stopping criterion were set for fitting. Altogether, there were ~ 200k tunable parameters; the training was performed using the Adam optimizer.

**pwmsandme.** Both CHS and GHTS data were used for training. First, 41-bp-long DNA sequences that flanked the abs_summit coordinates were retrieved for all transcription factors (TFs). Then, the sequences were split into overlapping 8-mers with a 1-nt step. To achieve equivariance, reverse complement sequences were generated for each 8-mer, and then each pair of k-mers was sorted in alphabetical order. In each pair, only the first k-mer was retained to represent the whole sense-antisense DNA pair. The resulting list of tokens for each label was processed with a TfIdfVectorizer to obtain embeddings that reflect the frequency of k-mer (or a group of k-mers) occurrence in the DNA sequence flanking the abs_summit of the transcription factor (TF) of interest, as opposed to all TFs in the dataset. TF-IDF embeddings were created for a range of n-grams from 1 to 3 in order to account for longer TF binding sites. The data were fitted into a StackingClassifier that comprised LogisticRegression and KNN as initial estimators and CatBoost[15], a gradient boosting classifier, as the final one. All embeddings except those of the target class were used as negatives during model training. Model performance was evaluated using the F1-score. The objective was to maximize the weighted average while ensuring a macro average above 0.7 and minimizing its difference from the weighted average on the validation set, which was obtained after the initial train-test

split.

**Salimov and Frolov Laboratory.** We developed NovaBind, a hybrid convolutional–recurrent architecture. Two parallel convolutional layers (kernel sizes 7 and 15) extract sequence motifs from one-hot encoded sequences. Convolutions were implemented without bias terms and followed by ReLU activations. The outputs are merged and passed to a bidirectional

GRU[18] (default implementation) with batch normalization to mitigate positional bias. A global max pooling layer precedes a fully connected network with SiLU activations. Skip connections link the convolutional outputs to the dense layers, and the raw input directly to the GRU. Training used AdamW with cosine annealing; hyperparameters were selected by Bayesian search on PBM data. Reverse-complement augmentation was applied, with post-hoc max-conjunction during inference. For PBM, the task was formulated as a multi-task regression with MSE loss. Input sequences were extended with linker regions to provide additional context. The PBM signal was represented as a two-component Gaussian mixture, and the probability of one component was used to reweight the signal. For HTS, training was performed on the last available cycle using a multi-label classifier with categorical cross-entropy. Barcode-free sequences were used as positive examples, whereas alien sequences served as negatives. Inference on CHS and GHTS employed a sliding-window approach (~60 nt, stride 1–9), with predictions aggregated by maximum. Ensembling across multiple seeds and folds was used to improve predictions.

**sbi two.** Our ModCRE implementation for generating PWMs yields many per TF. By using these in generating input for a Neural Network to predict binding, we circumvent the need to select a single PWM from many options. Here, we used a subset of PWMs to scan binding sequences with FIMO for hits. These hit counts were then used to train a Neural Network to predict whether the corresponding TF binds to the sequence or not. We applied the models to predict binding in the supplied test sets. PWMs used for training models with higher accuracies were selected from amongst the original ModCRE-generated PWMs. This subset of ModCRE-predicted PWMs was then used to generate the input for our AAA model to make its binding predictions on the test set.

**The Motifvators.** We developed STIMULUS, a novel software enabling joint optimization of model hyperparameters and data processing methods to reach higher performance on unseen datasets. For IBIS, we applied STIMULUS to KAN (Kolmogorov-Arnold Networks), a promising deep learning paradigm that focuses on learning linear weights as a function (b-splines) coupled with Convolution Neural Networks (CNNs), an architecture that has long-lasting success in the field. KANs have shown high generalization performance on science-related tasks and are sensitive to design choices according to the original paper[19], which makes STIMULUS a promising framework for covering such weaknesses. To initialize the CNN+KAN model for each transcription factor, we used the top-ranked PWMs discovered by HOMER (see above). PyTorch was used for defining the model, and STIMULUS (https://github.com/mathysgrapotte/stimulus-py) for data augmentation, hyperparameter tuning, and model training. As for the data, we parsed sequences from GHT-SELEX with a balanced number of positive/negative (negative defined as aliens) sequences. The sequences are also cut to 40bps around the peak. STIMULUS automatically determined how to augment the data, which hyperparameters and optimizer to use, and selected the best-performing configuration for each TF, resulting in one optimized model per TF. The size of each of these models is in the order of tens of thousands of parameters.

**Transcriptome.** We used a hybrid CNN-LSTM architecture: we employed CNN layers to extract sequence features and an LSTM layer to capture long-range dependencies. Similar concepts were applied in previous studies[20,21], which used FCN+RNN architectures. Here, TensorFlow was used for implementation.

**Trituration.** We developed a motif discovery pipeline for ChIP-seq data that combines STREME and BaMMmotif2[22] to optimize de novo motif identification. The datasets were split into independent training and test sets by sorting peaks by significance and dividing them into even (test) and odd (training) indices. To ensure reliable background modeling, we generated negative samples by selecting random genomic regions with a dinucleotide composition closely matching that of the training set. Anchor motifs were first generated with STREME across motif lengths from 6 to 24 bp (in steps of 2), producing candidate PWMs to initialize BaMMmotif2 models. For each motif length, BaMMmotif2 was run across Markov chain orders from 1 to 5, and performance was evaluated with auROC, auPRC, and pAUC (at FPR=0.001, normalized to 0–1). The optimal motif model defined by the highest combined score across these three indices was then selected and submitted for evaluation.

**chiCkeN pox gaNg.** We used all available HTS data to train EfficientNet-like models, LegNet[23] for each transcription factor (TF), and a separate model was trained for each available replicate. We assembled datasets comprising positive sequences and negatives (GC-matched "aliens") with a 1:1 class balance. Sequences were one-hot encoded. During training, we treated the task as a binary classification problem; the models were trained using Binary Cross-Entropy loss and the OneCycleLR scheduler. Reverse complementary sequences were used for augmentation. During the inference, the predictions of all corresponding models were averaged to obtain the final predictions for each TF.
